# Cell-type-specific inhibitory circuitry from a connectomic census of mouse visual cortex

**DOI:** 10.1101/2023.01.23.525290

**Authors:** Casey M Schneider-Mizell, Agnes L. Bodor, Derrick Brittain, JoAnn Buchanan, Daniel J. Bumbarger, Leila Elabbady, Clare Gamlin, Daniel Kapner, Sam Kinn, Gayathri Mahalingam, Sharmishtaa Seshamani, Shelby Suckow, Marc Takeno, Russel Torres, Wenjing Yin, Sven Dorkenwald, J. Alexander Bae, Manuel A. Castro, Akhilesh Halageri, Zhen Jia, Chris Jordan, Nico Kemnitz, Kisuk Lee, Kai Li, Ran Lu, Thomas Macrina, Eric Mitchell, Shanka Subhra Mondal, Shang Mu, Barak Nehoran, Sergiy Popovych, William Silversmith, Nicholas L. Turner, William Wong, Jingpeng Wu, The MICrONS Consortium, Jacob Reimer, Andreas S. Tolias, H Sebastian Seung, R. Clay Reid, Forrest Collman, Nuno Maçarico da Costa

## Abstract

Mammalian cortex features a vast diversity of neuronal cell types, each with characteristic anatomical, molecular and functional properties. Synaptic connectivity powerfully shapes how each cell type participates in the cortical circuit, but mapping connectivity rules at the resolution of distinct cell types remains difficult. Here, we used millimeter-scale volumetric electron microscopy^1^ to investigate the connectivity of all inhibitory neurons across a densely-segmented neuronal population of 1352 cells spanning all layers of mouse visual cortex, producing a wiring diagram of inhibitory connections with more than 70,000 synapses. Taking a data-driven approach inspired by classical neuroanatomy, we classified inhibitory neurons based on the relative targeting of dendritic compartments and other inhibitory cells and developed a novel classification of excitatory neurons based on the morphological and synaptic input properties. The synaptic connectivity between inhibitory cells revealed a novel class of disinhibitory specialist targeting basket cells, in addition to familiar subclasses. Analysis of the inhibitory connectivity onto excitatory neurons found widespread specificity, with many interneurons exhibiting differential targeting of certain subpopulations spatially intermingled with other potential targets. Inhibitory targeting was organized into “motif groups,” diverse sets of cells that collectively target both perisomatic and dendritic compartments of the same excitatory targets. Collectively, our analysis identified new organizing principles for cortical inhibition and will serve as a foundation for linking modern multimodal neuronal atlases with the cortical wiring diagram.

## Introduction

In mammalian cortex, information processing involves a diverse population of neurons distributed across six layers in an arrangement described as a cortical column^2–5^. The function of this circuit depends on not only the properties of cells individually, but also the network of synaptic connectivity through which they interact. The concept of cell types has become central to understanding how this network is organized^6^. Originally classified on the basis of morphology^7^, cortical cell types have more recently been characterized also by molecular, electrophysiological, and functional properties believed to subserve specialized computational roles^8–18^. There are thus a variety of powerful methods to define cell types, but determining how these definitions might reflect differences in cortical connectivity remains difficult.

Inhibitory neurons, despite making up little more than 10% of cortical neurons^19^, have at least as much cell type diversity as excitatory neurons^12, 14, 20^, offering the potential for highly selective control of cortical activity. However, most of our understanding of inhibitory connectivity is based not on individual cell types, but on coarse molecular subclasses^21–23^ with certain shared developmental, functional, and synaptic properties. Parvalbumin (PV)-expressing neurons target the perisomatic region of excitatory cells, including basket cells that target the soma and proximal dendrites of excitatory neurons and chandelier cells that target the axon initial segment. Additionally, PV basket cells synapse with one another^24, 25^ and other basket cells^26^. Somatostatin (SST)-expressing neurons, such as Martinotti cells, target the distal and apical dendrites of excitatory cells and also inhibit non-SST inhibitory subclasses. The connectivity of vasoactive intestinal peptide (VIP)-expressing neurons is heterogeneous, including not only a subclass of disinhibitory specialists that preferentially target SST cells^25, 27^, but also excitatory-targeting small basket cells that coexpress cholecystokinin (CCK)^28–30^. A fourth subclass expresses Id2^23^, spanning *lamp5* and *sncg* transcriptomic groups^20, 22^, and includes neurogliaform cells with diffuse synaptic outputs as well as a variety of cell types in layer 1. Within these cardinal subclasses, individual cell types are highly diverse^12, 13, 31^ and functionally distinct^18^ but in most cases little is known about connectivity at that level.

Moreover, the structure of how inhibition is distributed across different excitatory cells is still a matter of debate, despite being a key determinant of how cortical activity is controlled. While some studies have observed largely unspecific connectivity onto nearby cells^32, 33^, other studies have found examples of selective targeting of certain subpopulations of excitatory cells based on their layer position^34^ or long-range axonal projection target^35–37^. It is not known whether such selectivity is common or rare relative to unspecific connectivity. Likewise, many other basic organizational properties remain unclear, for example which excitatory neurons receive inhibition from the same interneurons, and do somatic-targeting and dendrite-targeting neurons have similar or different connectivity patterns to one another.

The ideal data to address such questions would capture the synaptic connectivity of individual interneurons across a broad landscape of potential targets. To date, physiological^38, 39^ or viral^40^ approaches to measuring connectivity are still challenging to scale to the full diversity of potential cell type interactions. In smaller model organisms like *Caenorhabditis elegans*^41^ and *Drosophila melanogaster*^42, 43^, dense reconstruction using large-scale electron microscopy (EM) has been instrumental for discovering cell types and their connectivity. In mammalian cortex, technical limitations on EM volume sizes have meant that similar studies could not examine complete neuronal arbors, making the link between cellular morphology and connectivity difficult to address^44–48^. However, recent advances in data generation and machine learning^49–51^ have enabled the acquisition and dense segmentation of EM datasets at the scale of a cubic millimeter, making circuit-scale cortical EM volumes possible^1, 45^. In this study, we used a millimeter-scale EM volume of mouse primary visual cortex (VISp)^1^ to reconstruct anatomy and synaptic connectivity for a continuous population of 1352 neurons in a column spanning from layer 1 to white matter (WM).

The scale of this data, combined with the resolution provided by EM, led us to address the fundamental question of how the morphological cell types of the neocortex relate to the synaptic connectivity of inhibitory neurons. Taking a data-driven approach to classical neuroanatomical methods, we first classified inhibitory neurons into four subclasses largely overlapping conventional classes according to how each cell distributed its synaptic output across excitatory dendritic compartments and other inhibitory neurons. These subclasses, combined with synaptic connectivity between inhibitory neurons, revealed a novel category of disinhibitory specialist that targets basket cells, distinct from the expected category targeting SST-like neurons. Next, we developed a novel classification of excitatory neurons using morphological and synaptic input properties, capturing features that were not clear from morphology alone. By analyzing the synaptic output of inhibitory neurons at both the single cell and subclass levels, we found that inhibitory neuron output exhibited widespread specificity, targeting few postsynaptic subclasses, and we identified groups of interneurons with similar subclass-specific targeting, but with different compartmental targeting. This specificity was achieved with combination of both spatially-precise axonal arbors and cell type selectivity, synapsing more or less onto particular cell types than expected. Our data suggest an organizing principle for inhibitory connectivity that is complementary to, but distinct from, cell types: diverse groups of inhibitory neurons that are positioned to collectively control activity of the same target populations with remarkable precision.

## Results

### A millimeter-scale reconstruction of visual cortex with synaptic resolution

In order to measure synaptic connectivity and neuronal anatomy within a large neuronal population, we analyzed a serial section transmission EM volume of mouse visual cortex acquired as part of the broader MICrONs project^1^. The volume was imaged from pia to white matter (WM), spanning 870 *×* 1300 *×* 820 *µm* (anteroposterior *×* mediolateral *×* radial depth), split into two subvolumes along the anteroposterior direction^1^. Here, we analyzed the larger subvolume, which covers 523 *µm* of the anteropostior extent and includes parts of VISp and higher order visual areas AL and RL^52^(Fig. 1a–c). Importantly, these dimensions were sufficient to capture the entire dendritic arbor of typical cortical neurons (Fig. 1a) at a resolution capable of resolving ultrastructural features such as synaptic vesicles (Fig. 1b). Convolutional networks generated an initial autosegmentation of all cells, segmented nuclei, detected synapses and assigned synaptic partners^1, 50, 51^. Due to reduced alignment quality near the edge of tissue, segmentation began approximately 10 *µm* from the pial surface and continued into WM. To simultaneously perform analysis and correct segmentation errors, we used a scalable, centralized proofreading platform integrated with a spatial database to dynamically query annotations such as synapses across edits^53–55^.

**Figure 1.**
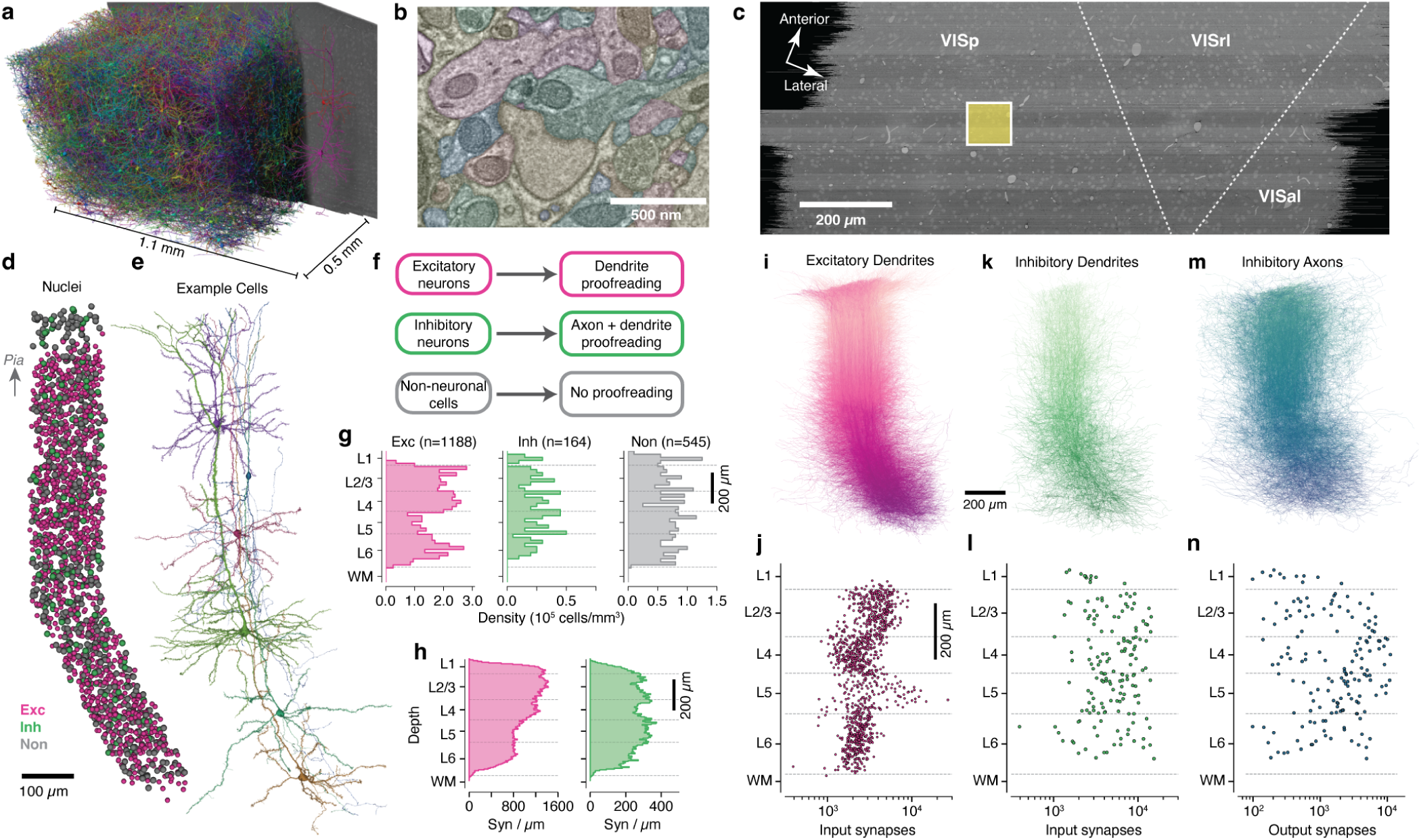
A columnar reconstruction of mouse visual cortex. **a)** The millimeter-scale EM volume is large enough to capture complete dendrites of cells across all layers. Neurons shown are a random subset of the volume, with a single example at right for clarity. **b)** The autosegmented EM data reveals ultrastructural features such as membranes, synapses, and mitochondria. **c)** Top view of EM data with approximate regional boundaries indicated. The yellow box indicates the 100 *µm ×* 100 *µm* column of interest. **d)** All nuclei within the column colored by cell class. **e)** Example neurons from along the column. Note that anatomical continuity required adding a slant in deeper layers. **f)** Proofreading workflow by cell class. **g)** Cell density for column cells along cortical depth by cell class. **h)** Input synapse count per *µm* of depth across all excitatory (purple) and inhibitory (green) column cells along cortical depth by target neuronal cell class. **i)** All excitatory dendrites, with arbors of cells with deeper somata colored darker. Same orientation as in **d**. **j)** Number of input synapses for each excitatory neuron as a function of soma depth. **k)** As in **j**, but for inhibitory neurons. **l)** As in **k**, but for inhibitory neurons. **m)** As in **j**, but for the proofread axons of inhibitory neurons. **n)** As in **k**, but for number of synaptic outputs on inhibitory neuron axons.

To generate an unbiased sample of cells across all layers, we focused our proofreading and analysis on cells whose soma was within a 100*×*100*µm* wide column spanning pia to WM and centered on the VISp portion of the volume (Fig. 1c). Note that reconstructions extended well past the column bounds, and this location was chosen to be far from dataset edges to avoid truncated arbors. A total of 1886 cells had their nucleus centroid inside the column. In order to follow a continuous population of neurons, the column slanted starting in layer 5, allowing the apical dendrites of deep-layer cells to be intermingled with the cell bodies of superficial cells (Figure 1d,e; see Methods). This slanted trajectory was also followed by primary axons of superficial cells and the translaminar axons of inhibitory neurons, suggesting that the pia–WM direction is not simply orthogonal to the pial surface in deep layers, but is shared across cell classes.

### A dense neuronal population sample across all layers

We classified all cells in the column as excitatory neurons, inhibitory neurons, or non-neuronal cells on the basis of morphology (Fig. 1d). For neurons, we performed extensive manual proofreading – more than 46,000 edits in all (Fig. 1f), guided by computational tools to focus attention on potential error locations (see Methods). Our proofreading strategy was designed to measure the connectivity of inhibitory neurons across all possible target cell types. Proofreading of excitatory neurons aimed to reconstruct complete dendritic arbors, combining both manual edits and computational filtering of false axonal merges onto dendrites (see Methods), while for inhibitory neurons we reconstructed both complete dendritic arbors and extensive but incomplete axonal arbors. Non-neuronal cells were not proofread, but we manually labeled non-neuronal subclasses as a data reference.

Consistent with previous reports^56^, excitatory cell densities varied between layers, while inhibitory neurons and non-neuronal cells were more uniform (Fig. 1g). The reconstructions included the locations of a total of 4,490,649 synaptic inputs across all cells. Synaptic inputs onto excitatory cell dendrites were more numerous in layers 1–4 compared to layers 5–6 (Spearman correlation of synapse count with depth: *r* = *−*0.92, *p* = 1.3 *×* 10*^−^*^1^^1^), while inputs onto inhibitory cells were relatively uniform across depths (Fig. 1h) (Spearman correlation of synapse count with depth: *r* = *−*0.06, *p* = 0.76).

Individual reconstructions captured rich anatomical properties across all layers. Excitatory cell dendrites (Fig. 1i) typically had thousands of synaptic inputs, with laminar differences in total synaptic input per cell (ANOVA for laminar effect: *F* = 82.9, *p* = 1.6 *×* 10*^−^*^48^, Fig. 1j). Typical inhibitory neurons had 10^3^–10^4^ synaptic inputs (Fig. 1k,l) and 10^2^–10^4^ outputs (Fig. 1m,n), but did not show strong laminar patterns (ANOVA for laminar effect: *F* = 0.72, *p* = 0.53). Collectively, inhibitory axons had 427,294 synaptic outputs. Attempts were made to follow every major inhibitory axon branch, but for large inhibitory arbors not every tip was reconstructed to completion and axonal properties should be treated as a lower bound. Based on comparisons to similar neurons where reconstruction aimed for completeness^57^, we estimate that typical axonal reconstructions captured 50-75% of their total synaptic output compared to exhaustive proofreading. There was also some variability; in particular, axons in layer 1 and deep layer 6 were generally less complete due to alignment quality, while a small number cells were more exhaustively reconstructed.

### Connectivity-based inhibitory subclasses

Molecular expression has emerged as a powerful organizing principle for inhibitory neurons, with four cardinal subclasses having distinct connectivity rules, synaptic dynamics, and developmental origins^21^. However, EM data has no direct molecular information, and no simple rules map morphology to molecular identity. Classical neuroanatomical studies often used the postsynaptic compartments targeted by an inhibitory neuron as a key feature of its subclass^30, 58, 59^, for example distinguishing soma-targeting basket cells from dendrite-targeting Martinotti cells.

Inspired by this approach, we used a data-driven approach based on the targeting properties of inhibitory neurons to assign cells to anatomical subclasses (Fig. 2a). For all excitatory neurons, we divided the dendritic arbor into four compartments: soma, proximal dendrite (<50 *µm* from the soma), apical dendrite, and distal basal dendrite (Fig. 2b) (Extended Data Fig. 1) (see Methods). Inhibitory cells were treated as a fifth target compartment. For each inhibitory neuron we measured the distribution of synaptic outputs across compartments (Fig. 2c). To further capture targeting properties, we devised two measures of how a cell distributes multiple synapses onto an individual target: 1) the fraction of all synapses that were part of a multisynaptic connection and 2) the fraction of synapses in a multisynaptic connection that were near another synapse in the same connection along the axon (“clumped”) (Fig. 2d). We used a distance threshold of 15 *µm*, approximately a quarter of the circumference of a typical cell body, although measurements are robust to this value (Extended Data Fig. 2). We use the term “connection” to indicate a pre- and postsynaptic pair of cells connected by one or more distinct synapses, and “multisynaptic connection” for a connection with at least two synapses. We trained a linear classifier based on expert annotations of the four cardinal subclasses for a subset of inhibitory neurons and applied it all cells (Fig. 2d, Extended Data Fig. 3).

**Figure 2.**
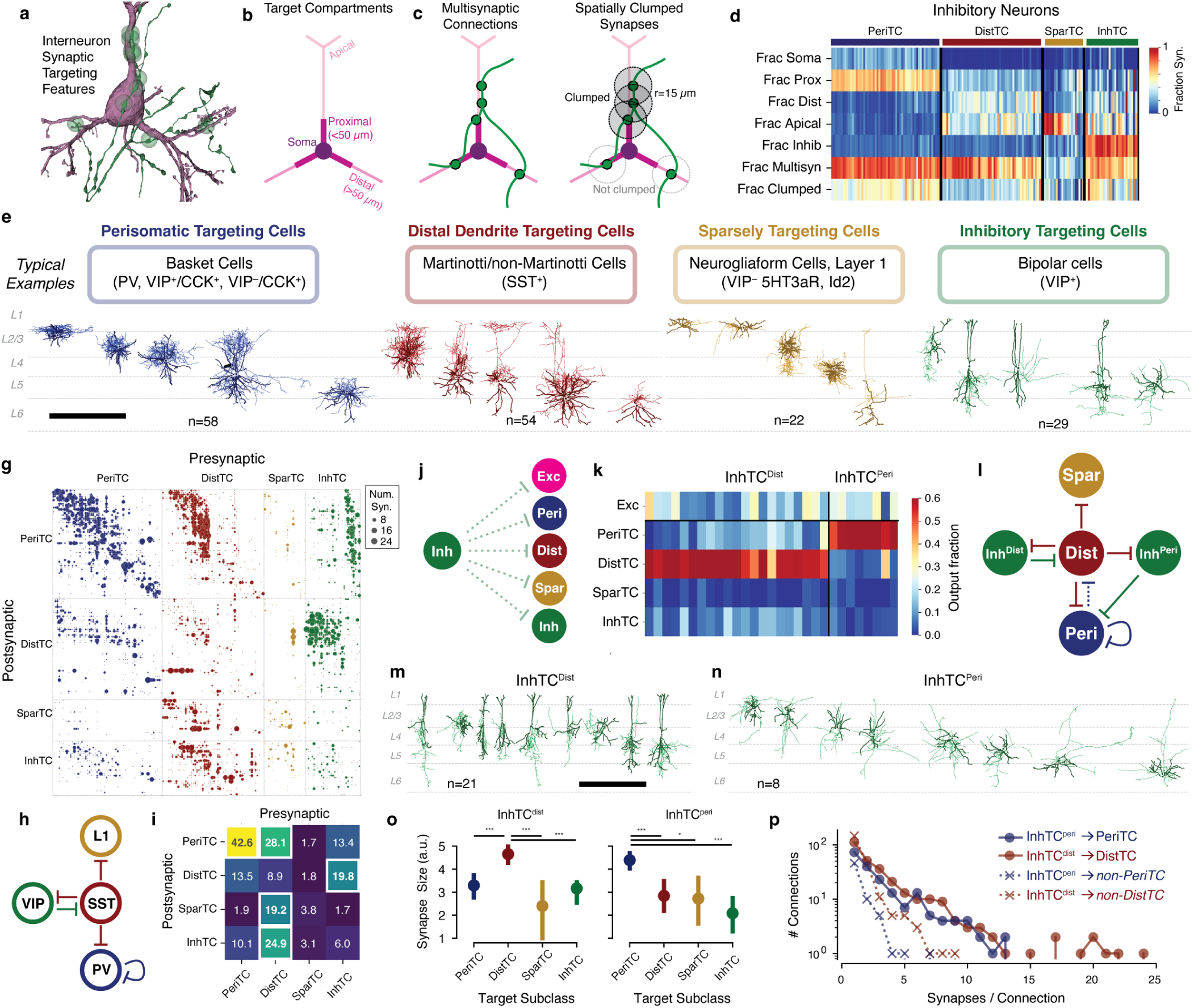
Data-driven characterization of inhibitory cell subclasses and their connectivity with one another. **a)** For determining inhibitory cell subclasses, connectivity properties were used such as an axon (green) making synapses (green dots) the perisomatic region of a target pyramidal cell (purple). **b)** Dendritic compartment definitions for excitatory neurons. **c)** Cartoon of a multisynaptic connection (left) and the synapses within the multisynaptic connection considered “clumped” along the presynaptic axon (right). **d)** Targeting features for all inhibitory neurons, measured as fraction of synapses onto column cells (for “Fraction clumped” only, all synapses in multisynaptic connections), sorted by target subclass. **e)** Relationship between anatomical connectivity categories (top), typical associated classical cell categories (middle), and anatomical examples (bottom) of the four inhibitory subclasses. Dendrite is darker, axon is lighter. Scale bar is 500 *µm*. **g)** Inhibition of inhibition connectivity. Each dot represents a connection from a presynaptic to a postsynaptic cell, with dot size proportional to synapse count. Dots are colored by presynaptic subclass and ordered by subclass, connectivity group (see Figure 5), and soma depth. **h)** Standard connectivity model of inhibition of inhibition based on molecular subclasses. **i)** Heatmap showing the mean number of synaptic inputs a postsynaptic cell received from all cells of a given presynaptic subclass. Note that the top five connections (highlighted in bold and white outlines) align with those arrows in **(h)**. **j)** Diagram of potential InhTC connectivity. **k)** Heatmap showing the fraction of synaptic outputs each InhTC places onto cells of other subclasses. InhTCs are clustered into two subtypes, one that targets DistTCs (InhTC^dist^) and one that targets PeriTCs (InhTC^peri^s). **l)** Connectivity diagram for InhTC^peri^s suggested by data. **m)** Morphology of example InhTC^dist^s. Scale bar is 500 *µm*. **n)** Morphology of all InhTC^peri^s. Scale bar same as **(m)**. **o)** Median synapse size from InhTC^dist^ (left) and InhTC^peri^ (right) onto inhibitory subclasses. Error bars indicate 95% confidence interval. T-test p-values indicated; *: p<0.05, ***: p<0.005 after Holm-Sidak correction. **p)** Distribution of synapses per connection for InhTC^peri^ and InhTC^dist^ onto their preferred and non-preferred targets.

We named each subclass based on its dominant anatomical property: perisomatic targeting cells (PeriTC) that primarily target soma or proximal dendrites, distal dendrite targeting cells (DistTC) that primarily target distal basal or apical dendrites, sparsely targeting cells (SparTC) that make few multisynaptic connections, and inhibitory targeting cells (InhTC) that primarily target other inhibitory neurons instead of much more numerous excitatory neurons (Fig. 2e). Typical examples of each subclass correspond approximately to classical or molecular subclasses (Fig. 2e), but there is not a one-to-one match. For example, PeriTCs would include soma-targeting cells from multiple molecular subclasses (e.g. both PV and CCK+ basket cells)^21, 28, 60^ and DistTCs could include any molecular class that targets apical dendrites, not only SST neurons. The SparTC subclass included both neurogliaform cells and all layer 1 interneurons, suggesting it largely contained cells from the *lamp5* and *sncg* transcriptomic classes^13^ or the collective *Id2* class^23^.

Some cell types, such as chandelier cells^61^, had no examples in the column (although were found elsewhere in the EM volume^62^), and other cells in the column either did not fall cleanly into classical categories. For example, among PeriTCs we found that the degree to which cells targeted soma versus proximal dendrites fell along a continuum (Extended Data Fig. 4). A small number of PeriTCs made very few synapses onto somata but many onto proximal dendrites, a feature inconsistent with the standard definition of a basket cell^28^. Among DistTCs, we found a similar range of distal basal versus apical dendrite targeting (Extended Data Fig. 4). Even those DistTCs with substantial output in layer 1, a defining feature of Martinotti cells across layers^11, 63^, showed substantial targeting of basal dendrites in layers 2–6 (Extended Data Fig. 5), suggesting an underappreciated additional aspect of Martinotti cell connectivity (See also Gamlin et al.^57^ and Bodor et al.^64^). Interestingly, the DistTC with the highest fraction of apical-targeting outputs (88%) did not significantly arborize in layer 1, but instead projected a descending horse-tail axon across layers 2–5 (see Fig. 6j).

### Inhibition of inhibitory neurons

Inhibitory neurons, even those that primarily target excitatory cells, also target other inhibitory neurons. The EM data offered a detailed view into the inhibition of inhibition, with 9,235 synapses between pairs of inhibitory neurons across 3,569 distinct connections (Fig. 2g, Extended Data Fig. 6). Numerous studies have identified a standard architecture for the inhibition of inhibition at the subclass level^25, 65, 66^: PV neurons inhibit other PV neurons, SST neurons inhibit all other subclasses (but not themselves), and VIP neurons inhibit SST neurons (Fig. 2h). Many variations on this broad pattern have been found, however^67^. For example, VIP+ neurons have been shown to target both SST and PV cells^27, 68^, but little is known about the relationship between these connections and the diversity of cells expressing VIP.

To measure inhibitory connectivity at the level of cardinal subclasses, we computed the average number of synapses between all neurons of a presynaptic subclass onto each single neuron of the postsynaptic subclass. The five expected subclass-level connections align with the five strongest connections measured from EM (Fig. 2i), based on the approximate correspondence (Fig. 2e). Both cardinal subclass identification and neuronal reconstructions were thus consistent with established connectivity.

At the level of individual cells, however, the data revealed novel connectivity patterns. We focused on InhTCs, “inhibitory specialists” that almost exclusively target other inhibitory neurons rather than excitatory cells (mean: 82% of synaptic outputs). For each InhTC we computed its distribution of synaptic outputs across inhibitory subclasses (Fig. 2j, k). Extensive literature has found VIP-positive inhibitory specialists that preferentially target SST cells^25, 39, 69^, and thus we expected InhTCs would largely target DistTCs.

Indeed, we found 21/29 InhTCs whose synaptic output was principally onto DistTCs (mean of 74% of those synapses onto inhibitory neurons), a group we denote InhTC^Dist^ (Fig. 2k). Single-neuron consideration of InhTC^Dist^ connectivity showed striking laminar organization (Extended Data Fig. 7). We found that InhTC^Dist^ in layers 2–4 targeted those DistTCs in layers 4-5, but not those in layer 2/3. Similarly, those DistTCs in layer 2/3 made few synapses onto InhTC^Dist^ in return. Interestingly, layer 2/3 DistTCs typically targeted excitatory neurons in upper (“layer 2”) but not lower (“layer 3”) layer 2/3, suggesting InhTC-mediated disinhibition differs across layer 2/3 pyramidal cells.

Unexpectedly, we also found a second population of inhibitory specialists. This smaller group of InhTCs (8/29) specifically targeted PeriTCs (mean of 82% of those synapses onto inhibitory neurons), hence we called them InhTC^Peri^) (Fig. 2k). While InhTC^Dist^ had bipolar or multipolar dendrites and were concentrated in layers 2–4 (Extended Data Fig. 7), consistent with typical VIP neurons (Fig. 2m), InhTCs^Peri^ all had multipolar dendrites and were distributed across layers (Fig. 2n). The eight InhTC^Peri^ in the column targeted 56/58 PeriTCs with a mean of 10.5 net synapses per target cell, suggesting that this connectivity is likely to include both PV and other molecular subclasses (Extended Data Fig. 6). Because the reciprocal inhibition between VIP and SST neurons is thought to play an important part in their functional role^70^, we next asked if InhTC^Peri^ receive reciprocal inhibition from PeriTCs. However, we found few reciprocal synapses from PeriTCs back onto InhTC^Peri^s, but numerous inhibitory inputs from DistTCs (Extended Data Fig. 8), suggesting an addition to the standard inhibitory diagram (Fig. 2k).

The targeting preference of InhTCs was seen across multiple aspects of their connectivity. We first looked at a measure of synapse size based on the automatic synapse detection (see Methods). InhTC^Dist^*→* DistTC synapses had a median size 44% larger than those onto other inhibitory subclasses (Fig. 2o). Similarly, the median InhTC^Peri^ *→* PeriTC synapse being 69% larger than synapses onto other inhibitory subclasses (Fig. 2o). In addition, the mean number of synapses per unique connection was significantly higher between InhTC^Dist^ *→* DistTC compared to other targets (Fig. 2p) (3.6 vs 1.6 syn./connection, *p* = 1.5 *∗* 10*^−^*^10^) and between InhTC^Peri^ *→* PeriTC compared to other targets (3.1 vs 1.5 syn./connection; *p* = 1.1 *∗* 10*^−^*^5^, Student’s t-test). The location of synapses onto of their preferred targets was similar for the two InhTC subgroups, with a median distance from soma of 83.5 *µm* (InhTC^Dist^) and 86.2 *µm* (InhTC^Peri^) and no significant difference in distribution (Kolmogorov-Smirnov test, p=0.25) (Extended Data Fig. 8). Taken together, both InhTC^Dist^ and InhTC^Peri^ express their distinct targeting preferences in a common manner: through increased synapse count, larger synapses and more synapses per connection.

### Dendritic synaptic input properties define excitatory subclasses

While inhibitory neurons have frequently been described as having dense, nonspecific connectivity onto nearby neurons^32, 71^, many studies have revealed examples not only of layer-specific connectivity^72^, but also selectivity within spatially intermingled excitatory subpopulations^36, 37, 73^. It is unclear the degree to which inhibition is specific or not, and, in general, the principles underlying which excitatory neurons are inhibited by which populations inhibitory neurons is not well understood.

To address these questions, we first anatomically characterized excitatory neurons subclasses in the EM data. Previous approaches to data-driven clustering of excitatory neuron morphology could only use information about the skeleton^14, 15, 17, 74^, but the EM data also has the location and size of all synaptic inputs (Fig. 3a). We reasoned that synaptic features in addition to skeleton features would better characterize the landscape of excitatory neurons, since synapses directly reflect how neurons interact with one another. We assembled a suite of 29 features to describe each cell, including synapse properties such median synapse size, skeleton qualities such as total branch length, and spatial properties characterizing the distribution of synapses with depth (Fig. 3b, Extended Data Fig. 9–11, see Methods). The synapse detection algorithm did not distinguish between excitatory and inhibitory synapses, and thus all synapse-based measures potentially include both types of synapses. We performed unsupervised clustering of these features (Fig. 3b–d), identifying 17 “morphological types” or “M-types”. Briefly, we repeatedly applied a graph-based clustering algorithm^75^ on subsets of the data to compute a matrix of co-clustering frequency between cells, followed by agglomerative clustering to obtain a consensus result (see Methods).

**Figure 3.**
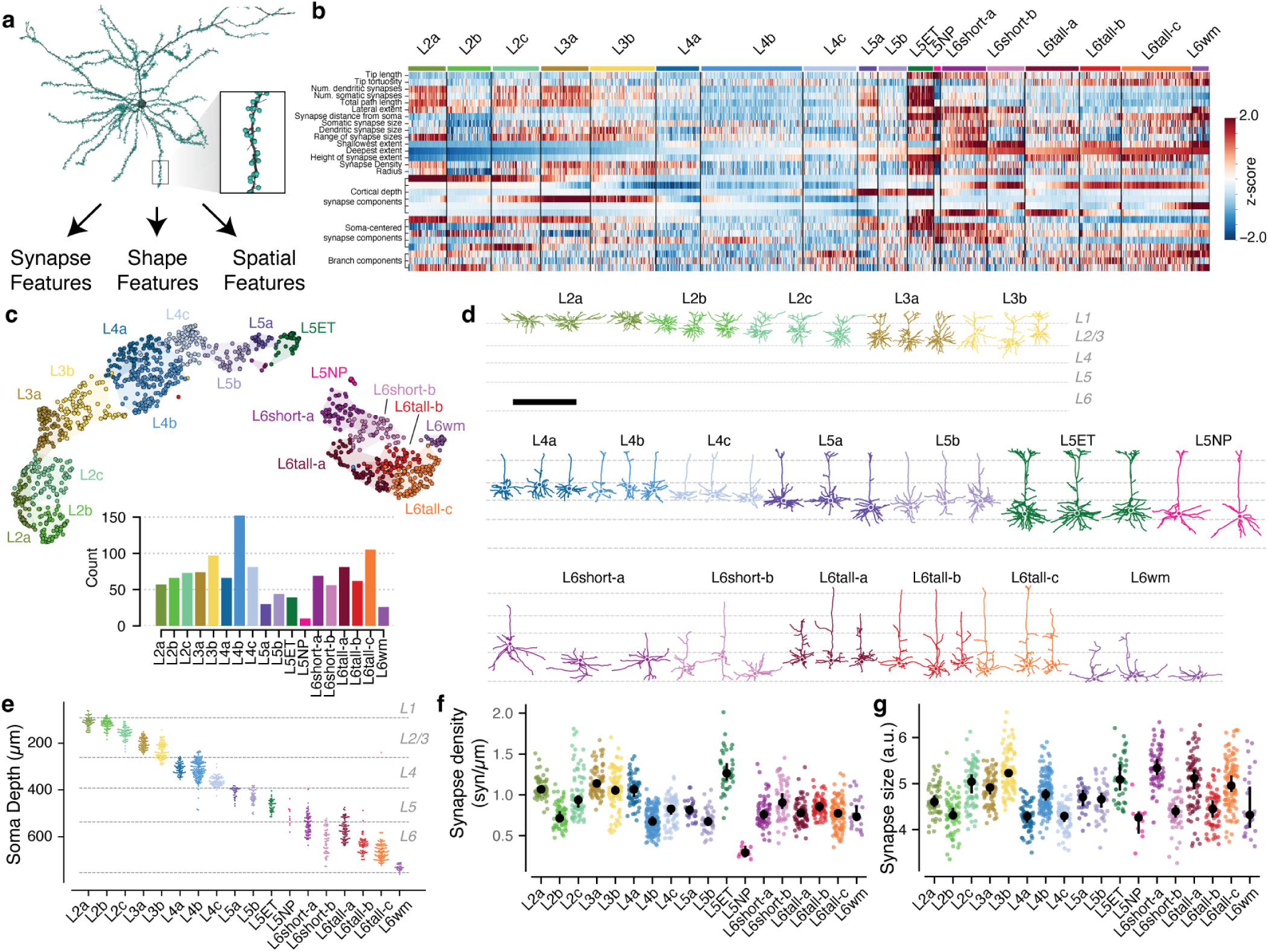
Data-driven characterization of excitatory neuron morphological types (M-types). **a)** Morphology (black) and synapse (cyan dots) properties were used to extract features for each excitatory neuron, such as this layer 2/3 pyramidal cell shown. **b)** Heatmap of z-scored feature values for all excitatory neurons, ordered by anatomical cluster (see text) and soma depth. See Methods for detailed feature descriptions. **c)** UMAP projection of neuron features colored by anatomical cluster. Inset shows number of cells per cluster. **d)** Example morphologies for each cluster. See (Extended Data Fig. 12) for all excitatory neurons. Scale bar is 500 *µm*. **e)** Soma depth of cells in each anatomical cluster. **f)** Median linear density of input synapses across dendrites by M-type. **g)** Median synapse size (arbitrary units, see Methods). In **f** and **g**, colored dots indicate single cells, black dots and error bars indicate a bootstrapped (n=1000) estimate of the median and 95% confidence interval.

To relate this landscape to known cell types, expert neuroanatomists labeled cells by the layer of the cell body and long-range projection type (IT: intratelencephalic or intracortical; ET: extratelencephalic or subcortical projecting, NP: near-projecting, CT: cortico-thalamic)^76^. Each layer contained multiple M-types, some spatially intermingled and others separating into subdomains within layer (Fig. 3e). M-types were named by the dominant expert label (Extended Data Fig. 13), with M-types within the same layer being ordered by projection class and average soma depth. For clarity, we use the letter “L” in the name of M-types (which may include some cells outside the given layer) and the word “layer” to refer to a spatial region. Upper and lower layer 2/3 emerged as having distinct clusters, which we denoted “L2” and “L3” respectively. Layer 6 had the most distinct M-types. These broadly split into two categories, those with shorter ascending or inverted apical dendrites (L6short), consistent with IT subclasses, and those with taller ascending apicals and narrow basal dendrites (L6tall), consistent with CT subclasses^14^. It was difficult to unambiguously label some layer 6 neurons as either IT or CT on the basis of anatomy alone, but 99% (n=142/143) manually assigned CT cells fell into one of the L6tall M-types. While we expect that L6short cells are IT and layer 6 CT cells fall into the L6tall M-types, we thus do not have the confidence to assign these names.

Most M-types had visually distinguishable characteristics, consistent with previous studies^14, 15, 17^ (Fig. 3d, Extended Data Fig. 12), but in some cases subtle differences in skeleton features were further differentiated by synaptic properties. For example, the two layer 2 M-types are visually similar, although L2a had a 29% higher overall dendritic length (L2a, 4532 *µm*; L2b, 3510 *µm*) (Extended Data Fig. 9). However, L2a cells had 80% more synaptic inputs than L2b cells (L2a, 4758; L2b, 2649), a 40% higher median synapse density (L2a, 1.04 syn/*µm*; L2b, 0.72 syn/*µm*) (Fig. 3f,g), and a wider distribution of synapse sizes (Extended Data Fig. 9). Median synapse size turned out to differ across M-types, often matching layer transitions (Fig. 3g, (Extended Data Fig. 11)). Strikingly, L5NP cells were outliers across synaptic properties, with the fewest total dendritic inputs, lowest synaptic input density, and among the smallest synapses (Fig. 3g,h). Excitatory M-types thus not only differ in morphology, but also in cell-level synaptic properties like total synaptic input and local properties like synapse size.

### Coordination of inhibition across excitatory M-types

Excitatory M-types are defined by their different structural properties, but this may or may not be meaningful to cortical circuitry. One piece of evidence that M-types do matter would be if different M-types received input from different inhibitory populations. Having classified inhibitory subclasses and excitatory M-types, we thus analyzed how inhibition is distributed across the landscape of excitatory neurons. The column reconstructions included 70,884 synapses from inhibitory neurons onto excitatory neurons (Fig. 4a). PeriTCs and DistTCs were by far the dominant source of inhibition, with individual cells having as many as 2,118 synapses onto excitatory cells in the column (mean PeriTC: 581 synapses per presynaptic cell; mean DistTC: 596 synapses per presynaptic cell), while SparTCs and InhTCs made far fewer synapses per presynaptic cell (mean SparTC: 74 synapses, mean InhTC: 16 synapses) (Fig. 4b). Inhibition was distributed unequally across M-types (Fig. 4c). Much of this difference in inhibition was related to differences in overall synaptic input. Across M-types, synaptic input at the soma, which is almost completely inhibitory, was strongly correlated (r=0.96, p=5 *×* 10*^−^*^10^) with net synaptic input onto dendrites, which is primarily excitatory (Extended Data Fig. 14). Notably, this structural balance of dendritic and somatic input also remained significant across individual cells within 16/18 M-types (all except L5NP and L5wm).

**Figure 4.**
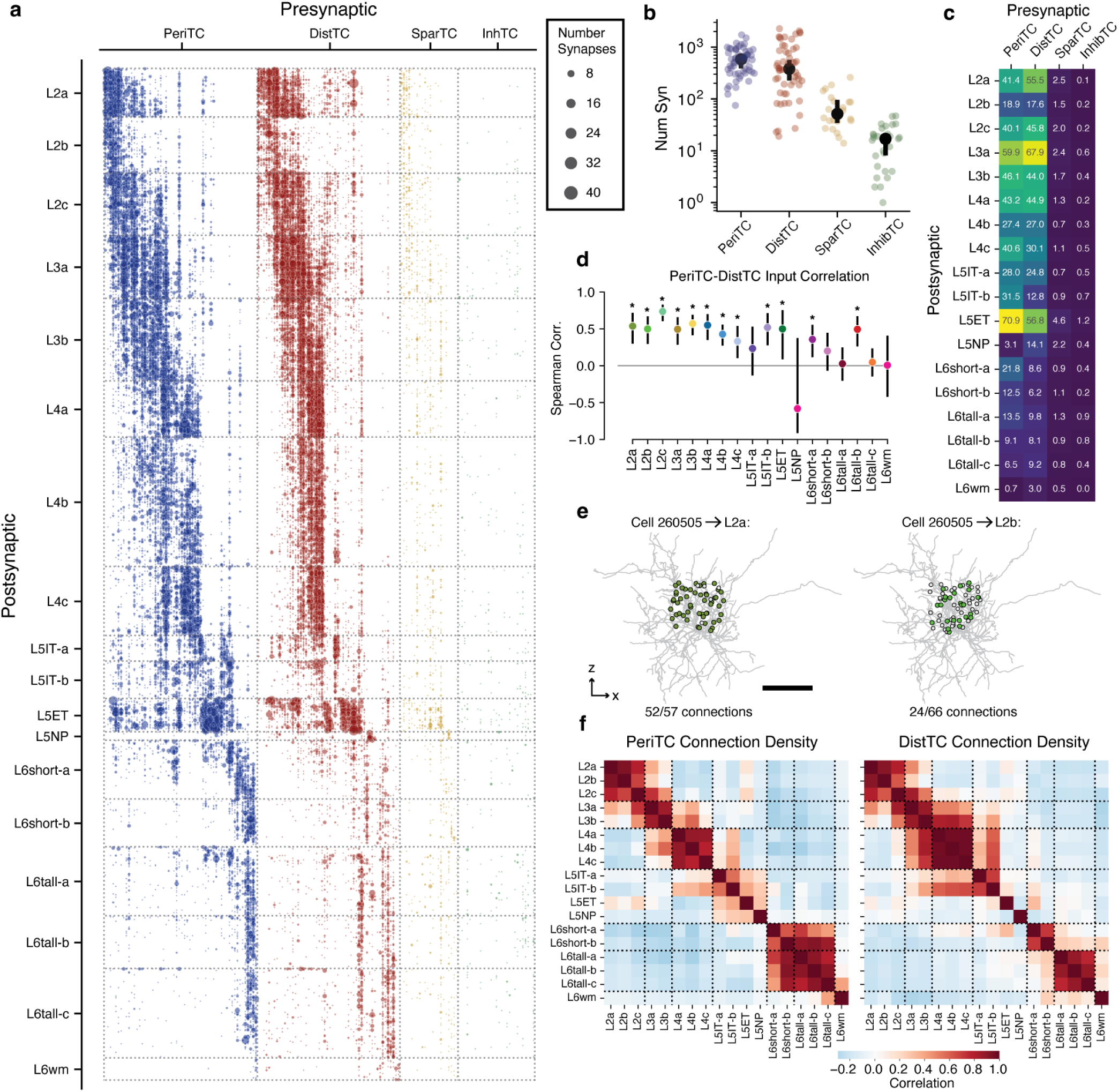
Inhibition of excitatory neurons. **a)** Connectivity from all inhibitory neurons (columns) onto all excitatory neurons, sorted by M-type and soma depth. Dot size indicates net number of synapses observed. **b)** Net synapses onto column cells for each inhibitory subclass. Black dots indicate median, bars show 5% confidence interval. **c)** Mean net synapses per target cell from each inhibitory subclass onto each excitatory M-type. **d)** Spearman correlation of PeriTC and DistTC net input onto individual cells, measured within each M-type. Bars indicate 95% confidence interval based on bootstrapping (n=2000). Stars indicate M-types significantly different from zero with a p-value < 0.05 after Holm-Sidak multiple test correction. **e)** Example of connectivity density calculation. Connectivity density from a single interneuron (gray) onto all cells within two example M-types (left: L2a right, L2b). Potential target cell body positions shown as dots, filled if synaptically connected and gray otherwise. Scale bar is 100*µm*. **f)** Pearson correlation of connectivity density between excitatory M-types, based on PeriTCs (left) and DistTCs (right). Dotted lines indicate groups of cells roughly within a layer.

Similarly, synaptic input from PeriTC and DistTC was also typically balanced onto individual cells for each M-type. We examined the number of PeriTC and DistTC inputs onto individual excitatory neurons for each M-type and found that PeriTC and DistTC input was significantly positively correlated for 12/18 M-types (Fig. 4d), suggesting coordinated amounts of inhibitory synaptic inputs across the entire arbor of target cells. M-types in upper layers had particularly heterogeneous amounts of inhibitory input, with L2b cells receiving 60% fewer synapses from intracolumnar interneurons as spatially intermingled L2a cells (L2b: 37.7 *±* 0.27 syn; L2a: 94.8 *±* 0.58 syn), while L3b cells had nearly as many intracolumnar inhibitory inputs as much larger L5ET cells. All layer 6 M-types had relatively few intracolumnar inhibitory inputs compared to upper layers (Fig. 4c). However, note that due to the columnar sampling, this reflects only local sources of inhibition and does not eliminate the possibility that deep layer neurons receive inhibitory input from more distant cells compared to upper layers.

Individual inhibitory neurons often targeted multiple M-types, suggesting that certain cell type combinations are inhibited together. For each inhibitory neuron we computed the connection density onto each M-type, that is the fraction of cells within the column that received synaptic input (Fig. 4e). To measure the structure of co-inhibition, we computed the correlation of inhibitory connection density between M-types across PeriTCs and DistTCs separately (Fig. 4f). A high correlation would indicate that the same inhibitory neurons that connected more (or less) to one M-type also connect more (or less) to another, while zero correlation would suggest independent sources of inhibition between M-types.

These correlations revealed several notable features of the structure of inhibition across layers. In superficial cortex, the layer 2 and layer 3 M-types were strongly correlated within layer, but had relatively weak correlation with between layers, suggesting different sources of inhibition. Layer 4 M-types, in contrast, were highly correlated with one another. Layer 5 M-types were more complex, with weak correlation with one another, suggesting largely non-overlapping sources of inhibition, particularly among neurons with different long-range projection targets. Interestingly, layer 5 IT cells shared co-inhibition with layer 4 M-types. Layer 6 inhibition was virtually independent from other layers, with DistTC connectivity also distinct between IT-like L6short cells and CT-like L6tall cells. Importantly, most co-targeting relationships were consistent for both PeriTC and DistTC output, suggesting that cardinal inhibitory subclasses distribute their output across excitatory neurons with similar patterns of connectivity.

### Cellular contributions of inhibition

How do individual neurons distribute their output to produce the patterns of inhibition described above? To compare patterns of output across inhibitory neurons, for every inhibitory neuron, we measured the fraction of synaptic outputs made onto each M-type (Fig. 5a). This normalized synaptic output budget reflected factors such as the number of synapses per connection and the number of potential targets, but is not strongly affected by partial arbors. We performed a consensus clustering (see Methods), identifying 18 “motif groups”, sets of cells with similar patterns of output connectivity (Fig. 5a, Extended Data Fig. 15). While this measurement only included synapses with cells within the column, interneurons made more than 4 times more synapses onto cells outside the column than within (Extended Data Fig. 15s,t). To check if these results would hold with data outside the column, we used a prediction of neuronal M-types based on perisomatic features and trained on column M-type labels^62^. We found that within-column and predicted dataset-wide synaptic output budgets were highly correlated (pearson r=0.90), confirming that the columnar sampling provided a good estimate of overall neuronal connectivity (Extended Data Fig. 15u,v).

**Figure 5.**
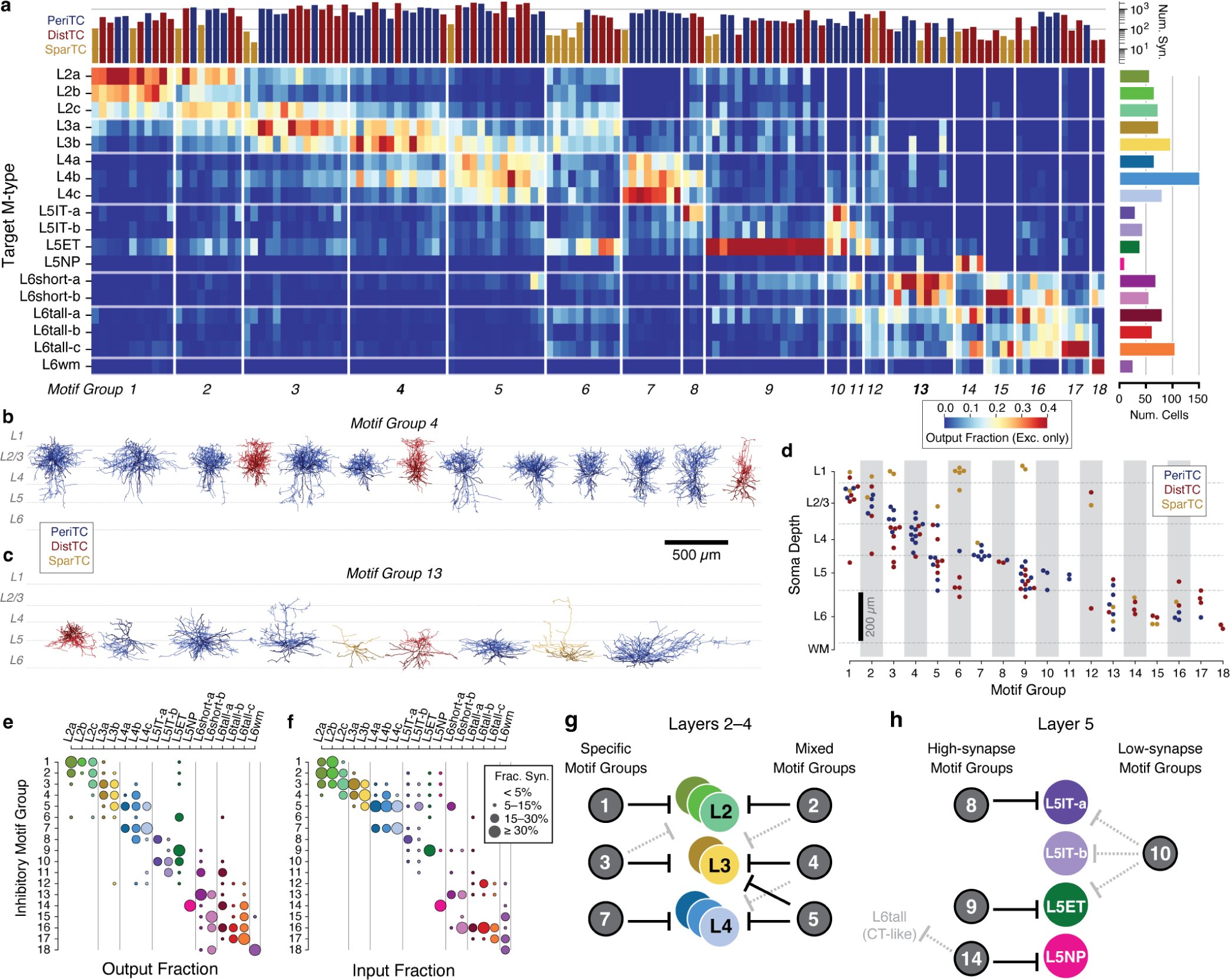
Inhibitory motif groups organize inhibitory connectivity. **a)** Distribution of synaptic output for all interneurons, clustered into motif groups with common target distributions. Each row is an excitatory target M-type, each column is an interneuron, and color indicates fraction of observed synapses from the interneuron onto the target M-type. Only synapses onto excitatory neurons are used to compute the fraction. Neurons are ordered by motif group and soma depth. Bar plots along top indicate number of synapses onto column cells, with color showing subclass (as in **d**). Bar plots along right indicate number of cells in target M-type. **b)** All cells in Group 4. Colors as in **d**. **c)** All cells in Group 13. Colors as in **d**. **d)** Soma depth and subclass for cells in each motif group. **e)** Net synaptic output distribution across M-types for each motif group. **f)** Synaptic input for each M-type from each motif group as a fraction of all within-column inhibition. **g)** Schematic of motif group connectivity in upper layers. **h)** Schematic of motif group connectivity in Layer 5.

Each motif group represented a collection of cells that targeted the same pattern of excitatory cell types. While some motif groups focused their output onto single excitatory M-types (e.g. Group 9) or layers (Group 7), others spanned broadly (e.g. Group 6). However, motif groups were not simply individual cell types. Individual motif groups (Fig. 5b,c) showed diversity in both individual cell morphology and connectivity subclasses (Extended Data Fig. 15, Extended Data Fig. 1). Indeed, 15/18 groups (comprising 156/163 cells) included neurons from at least two subclasses, often aligned in cortical depth (Fig. 5d). This aligns with the observation of similar co-targeting between PeriTCs and DistTCs.

Presynaptic specificity alone does not tell us about the importance of a connection to its postsynaptic partners. To get a view of the relationship between motif groups and M-types, we computed both the average output fraction from each motif group onto each M- type, (Fig. 5e) and the input fraction, the fraction of all within-column inhibitory synaptic inputs onto a given M-type that come from each motif group (Fig. 5f). These offered different perspectives, as an individual neuron could be both very selective but also contribute much less input than another less selective cell. Input fraction often followed output fraction for particularly strong connections, but not always. For example, while Group 3 more strongly targeted layer M-types in layer 3 than layer 2, it still contributed a substantial fraction of all inhibitory layer 2 input. In addition, we found that dominant connections for motif groups had both high connectivity density and multiple synapses per connection (Extended Data Fig. 16), properties that suggest a strong functional role in the circuit.

Inhibitory circuits were organized differently in upper layers compared to layer 5. In layers 2–4, each excitatory M-type received strong inhibition from 2-3 motif groups with overlapping combinations of targets, some specific within layers and others that cross layer boundaries (Fig. 5g). In contrast, most motif groups targeted only single M-types in layer 5, although in some cases also targeted cells in other layers (Fig. 5h). Connectivity patterns in layer 6 included clear examples of IT-specific and putatively CT-specific cells, similar to the layer 5 projection classes, but also had cells, particularly PeriTCs, that targeted widely in layer 6. Taken together, our data suggest that cortical inhibition is comprised of groups of neurons which target specific collections of M-types across their perisomatic and dendritic compartments with sub-laminar precision.

### Synaptic selectivity

Our data suggest that cell-type-specific targeting is a widespread property of cortical interneurons, in that individual neurons target only a small number of excitatory subclasses. Many factors could go into achieving such specific connectivity. Neurons of different types have varying dendritic and axonal morphologies that physically constrain potential interactions^77, 78^. Neurons also vary in the density of synaptic outputs or synaptic inputs and exhibit different compartmental targeting preferences. In addition, they can exhibit cell type selectivity, which we define as making synapses with particular cell types more or less than might be expected based on other factors such as axon/dendrite overlap, Note that we explicitly distinguish this from specificity, which we use to describe how concentrated the output connectivity is onto particular cell types. To differentiate the effects of these contributing factors, we assembled a rich collection of information about each interneuron: morphology (Fig. 6a), synaptic connectivity, and how output is distributed across compartments (Fig. 6b) or excitatory M-types (Fig. 6c) within the column. To quantify the effect of selectivity in particular, we developed a Selectivity Index (SI) comparing observed synaptic connectivity to a null model capturing key properties outside of M-type (Fig. 6d). The null model needed to capture not only space, but also compartment preference and postsynaptic factors such as number of synapses a cell typically receives and the spatial heterogeneity of potential targets^78–80^. Because many of these factors are correlated with target cell type, such a null model aims to address confounding between the cell type label itself and those structural properties that affect connectivity more generally, for example if cells with higher input synapse density receive more inhibitory inputs irrespective of their cell type.

**Figure 6.**
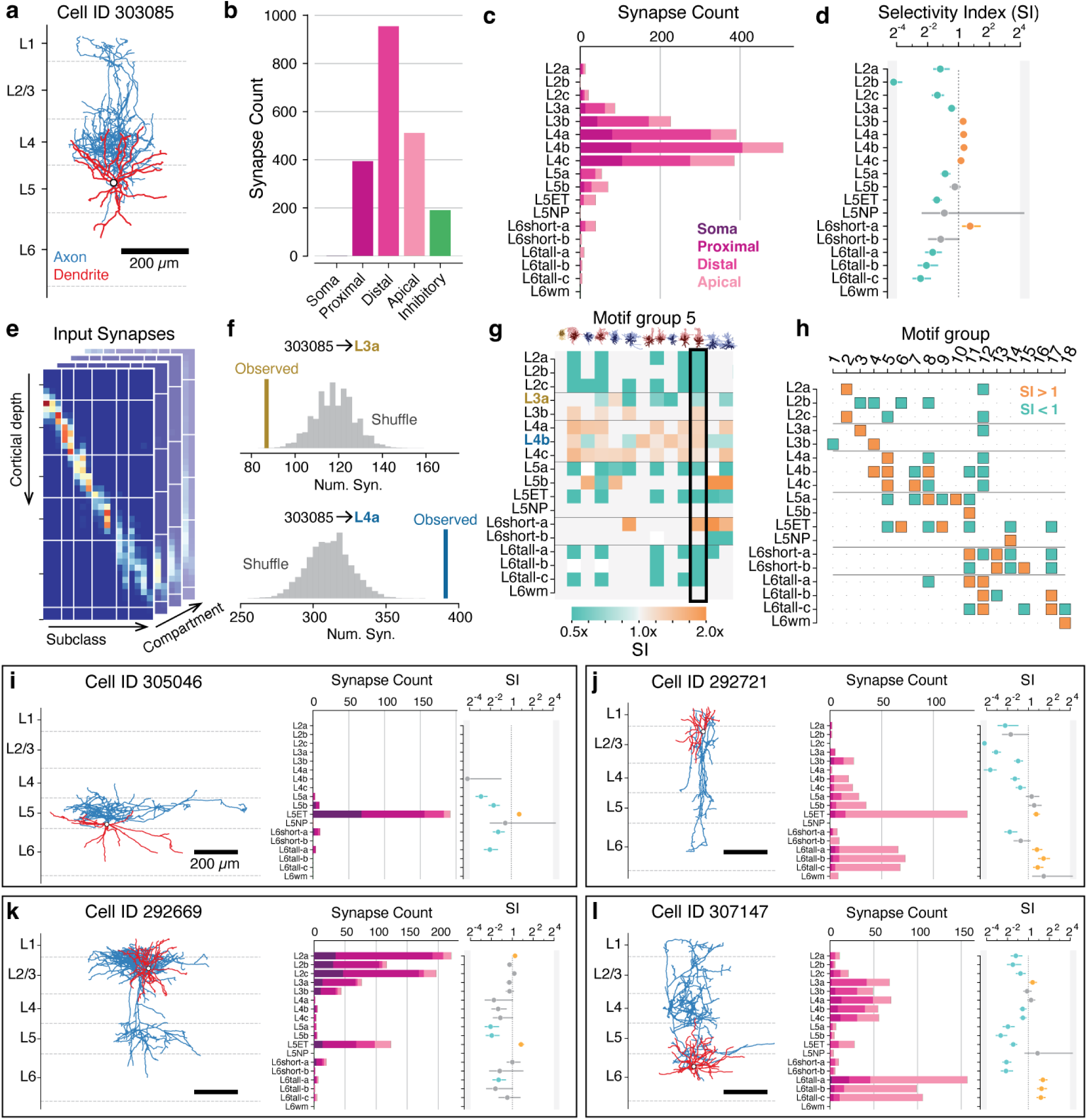
Synaptic selectivity and cell connectivity cards. **a)** Example inhibitory neuron (Cell ID 303085). Axon in blue, dendrite in red. **b)** Distribution of synaptic outputs across target compartments for the cell in **a**. **c)** Distribution of synaptic outputs across M-types (bar length) and compartments (bar colors) for the cell in **a**. **d)** Selectivity index (SI) values for the cell in **a**, measured as the ratio of observed synapse count to median shuffled synapse count for a null model as described below. Error bars indicate 95 percentile interval. Colored dots (blue: low, orange: high) indicate significant differences (two-sided p<0.05) relative to the shuffle distribution after Holm-Sidak multiple test correction. **e)** As a baseline synapse distribution for null models, all synaptic inputs onto all cells in the column were binned by compartment, depth, and M-type. See (Extended Data Fig. 17) for more details. **f)** Shuffled connectivity for the cell in **a** was computed by sampling from the baseline synapse distribution with the observed depth and compartment bins and counting (N=1,000) and counting synapses onto each M-type. Example shuffle values for L3a (top) and L4a (bottom) M-types vs. observed synapses are shown. **g)** SI for all cells in Motif group 5. Non-significant values are assigned a value of 1. The cell in **a** is highlighted by a black box. **h)** Direction of the median cell’s SI from each motif group onto each M-type. Orange indicates more connected, blue less connected. Connections where the median SI was non-significant are indicated with a dot. **i––l)** Compact cell connectivity cards encapsulating anatomy (left), M-type target distribution (middle, bar length), compartment targeting (middle, bar colors as in **d**), and SI (right, as in g) for four example neurons. Full connectivity cards for all cells can be found in Extended Data Cell Atlas.

We computed a baseline distribution by considering all synaptic inputs onto all column neuron dendrites and binning them by cortical depth (20 *µm* depth bins), M-type, and target compartment (Fig. 6e) (Extended Data Fig. 17). For each interneuron, we computed a shuffled output distribution across M-types by repeatedly sampling connectivity from the baseline distribution based on that cell’s synapse depths and compartment targeting distribution. For each connection from an interneuron onto a potential target M-type, we defined the SI as the ratio of observed connectivity to the median of the distribution of shuffled connectivity (N=1000 repeats) (Fig. 6f), reflecting the amount of cell-type-dependent selectivity beyond the factors included in the null model. While this sampling included both excitatory and inhibitory synapses, previous studies^81, 82^ and our data suggest that excitatory and inhibitory inputs are proportional to one another, even at the level of individual cells (Extended Data Fig. 14).

Because motif groups distributed their output with similar specificity, we asked if cells in motif groups had common patterns of selectivity as well (Fig. 6g). To ask where required common selectivity to produce their observed connectivity, we computed the median SI for each M-type/motif group pair, setting non-significant SI values to 1 (Fig. 6h). We found that while 17/18 motif groups showed consistent positive or negative selectivity for some targets, in many cases highly specific connectivity was not associated with increased selectivity. For example, Group 1 is highly specific to layer 2 targets but surprisingly did not show consistent positive selectivity for them (Extended Data Fig. 17). Examination of each constraint of the null model — synapse abundance of different targets, presynaptic compartment specificity, and presynaptic depth distribution — suggested that this was due to Group 1 axons having a narrow spatial distribution of axons that strongly overlapped layer 2 targets, which for many cells was sufficient to explain their connectivity (Extended Data Fig. 17). This effect was particularly pronounced for PeriTCs, which tended to target a more compact spatial domain with less overlap between M-types. In contrast, for DistTCs the increased spatial overlap of distal and apical dendrites of different M-types required additional selectivity to explain their connectivity (Extended Data Fig. 17). Collectively, this suggests that to achieve specific targeting patterns, interneurons both project their axons to precise spatial domains and selectively favor or disfavor making synapses with specific targets, with the relative contribution of these factors differing across cell types.

### Cell connectivity cards

While motif groups describe the broad organization of groups of cells, individual interneurons showed fascinating but idiosyncratic structural properties. To quickly assess individual cell properties, we devised a compact collection of measures that summarize morphology and connectivity at the level of M-types and compartments that were assembled into “connectivity cards” (Fig. 6i–l). Individual cards can reveal unique features that were not clear from groups alone, such as extreme specificity (Fig. 6i) and different patterns of translaminar connectivity (Fig. 6j–l). A full atlas of cards for all interneurons can be found in Extended Data Cell Atlas.

## Discussion

While excitatory neurons display morphological and transcriptomic diversity beyond layer boundaries or even long range projection classes^14, 83^, it has been unclear in general how or if these differences have functional consequences in the local inhibitory circuit. Here, we generated an EM reconstruction of neuronal anatomy and inhibitory connectivity across a column of visual cortex. This data offered new insights into not only the connectivity of individual neurons, but also the broader organization of inhibition across neuronal subclasses. Using synaptic properties in addition to traditional morphological features, we found a collection of excitatory M-types with distinct patterns of inhibitory input, demonstrating that anatomical distinctions, including both projection classes and more subtle sublaminar differences, are reflected in the cortical circuit. Inhibitory neurons were organized into motif groups, heterogeneous collections of cells that convergently targeted both perisomatic and dendritic compartments of particular combinations of M-types. We also identified new aspects to the inhibition of inhibition, including a novel type of disinhibitory specialist that mainly targets basket cells.

### A. Sublaminar specificity of inhibitory targeting

This question of interneuronal specificity has been perhaps best studied in layer 5, with its highly distinct ET and IT excitatory projection subclasses^84^. For dendrite targeting cells, precise genetic targeting of layer 5 SST subtypes identified distinct cell types that targeted ET versus IT cells^37^. In addition, developmental perturbation altering ET or IT neurons has suggested that they have different perisomatic input as well, with PV cells preferentially targeting ET and CCK neurons targeting IT^85–87^. The EM reconstruction presented here identified additional structure agreed to this picture. We found that ET cells received inputs from a larger number and diversity of inhibitory cells than IT, suggesting that despite being less numerous, ET cells have a larger and more complex inhibitory network than IT cells. ET cells were also frequently involved in translaminar circuits, with several examples of both ascending and descending translaminar PeriTCs and ascending DistTCs that targeted both L2/3 neurons and ETs but not ITs, suggesting bidirectional pathways for coordinated inhibition. Morphology and inhibitory connectivity also identified diversity within the L5IT population: shallow L5ITa and deeper L5ITb. L5ITa had a small collection of highly specific PeriTC and DistTC inputs, including a Martinotti cell distinct from previously established IT-specific SST cell types^37^ (see also Gamlin et al.^57^). L5ITb, in contrast, received highly selective inhibition from only one inhibitory neuron, and otherwise largely shared inhibition with layer 4 pyramidal cells. In addition, layer 5 NP cells had yet another distinct collection of inhibitory neurons. Collectively, the data suggests several non-overlapping inhibitory networks within layer 5 alone, with few sources of inhibition that collectively inhibited all layer 5 neurons. This affords the network the potential to selectively inhibit each projection class both at the soma and across apical and basal dendrites, potentially with different cell types active under different network conditions and behavioral states^18^, or with synapses changing via different plasticity rules^88^.

Subtype-specific inhibitory circuits were found across all other layers as well. Layer 6 also has distinct IT and CT excitatory projection subclasses and, similar to layer 5, we found distinct inhibitory subpopulations targeting each. In contrast to layer 5, however, there was a combination of both projection-specific and broad layer 6 inhibition. We speculate that this indicates that projection subnetworks in layer 6 might interact with one another more than in layer 5, in order for collective inhibition to be useful.

Even in layers 2–4, with only IT cells, there was significant sublaminar specificity. The differential inhibition of layer 2 versus layer 3 cells suggests that are functionally distinct subnetworks with independent modulation. This could mirror depth-dependent differences in in intracortical projection patterns^89^, similar to prefrontal cortex, where amygdala-projecting layer 2 cells receive inhibition that selectively avoids neighboring cortical-projecting cells^36, 73^. Another possibility is that they are well-posed to differentially modulate top-down versus sensory-driven activity^90, 91^, as layer 3 receives more sensory thalamic input than layer 2^92, 93^. More generally, the distinct inhibitory environments of upper and lower layer 2/3 have been observed across cortex, from primary sensory areas^44, 94^ to higher-order association areas^36, 73, 95^, suggesting that it may reflect a general functional specialization.

Interestingly, most inhibitory neurons that targeted main thalamorecipient layer 4 also targeted layer 3, possibly reflecting the need to collectively control populations that share common thalamic input. The most specific layer 4 neurons were a collection of PeriTCs that also all showed particularly selective innervation of a morphologically distinct excitatory subpopulation in deep layer 4 (see also Weis *et al.*^96^). Such sublaminar organization within layer 4 is apparent in primates and cat visual cortex on the basis of thalamic axon projection and cytoarchitecture^97, 98^, and was seen in EM reconstruction of barrel cortex^99^. The data here points to a graded transition in mouse visual cortex rather than sharp sublamina, but with a similar principle of specialized thalamic inputs and local circuits.

Collectively, pervasive sublaminar specificity of inhibition hints that their excitatory targets have different circuit roles as well, and suggests that it will be fruitful to pay close attention to sublaminar patterns of activity. However, the functional consequences of these inhibitory networks will depend on how different groups of inhibitory neurons are recruited and how local excitation feeds back into the inhibitory network. Interestingly, a companion study found that L5-ET neurons target those inhibitory neurons that synapse back onto them^64^, raising the question if M-types generally target those motif groups that exert inhibitory influence on them. It will also be important to understand the extent to which inhibitory cells with overlapping targets are active simultaneously, collectively setting an inhibitory tone, or instead different populations that target the same cells are active under different contexts, as in hippocampal basket cells^26^.

### B. Basket cell disinhibition

Extensive functional experiments have demonstrated that a class of VIP interneurons are specialized to target other inhibitory neurons, with little output onto excitatory neurons. Such disinhibitory VIP neurons have been shown to strongly target SST neurons across cortical areas^25, 27, 39, 69, 100^ and, to a lesser extent, fast spiking or PV neurons^27, 100^. Here, we found two classes of disinhibitory specialists: one that targeted putative SST cells and a second that targeted basket cells. Bipolar cells, a typical VIP interneuron morphology, were all putatively SST-targeting, while multipolar cells were found in both. Future experiments will be required to determine which molecular subclass the basket-targeting disinhibitory specialists are, and they do not fit any well-established connectivity profile.

This novel class of cells specialized to control basket cells adds yet another pathway for the cortical network to control perisomatic inhibition. Fast spiking or PV basket cells have been shown to be inhibited by other PV cells^25, 39^, SST cells^101^ and even by neurogliaform cells via volume transmission^102^. The basket-targeting disinhibitory specialists differ from these other pathways in their specificity — not only do they distribute the majority of synaptic output onto basket cells instead of any other inhibitory or excitatory targets, they do so with larger synapses and more synapses per connection. This highly specific targeting offers an intriguing pathway to potentially enhance basket-cell-mediated excitatory gain or synchrony without significantly affecting other neuronal populations. Determining what conditions cause these cells to be active will be important for understanding their functional effect.

### C. Limitations

One principal concern is the generalizability of data, since it comes from a single animal, is located toward the edge of VISp, and has at most a few examples per cell type. Companion work from the same dataset focusing on multiple examples of a few morphologically-defined cell types shows consistent target preferences^57, 64^, and our data also agrees with recent functional measurements of type-specific connectivity of SST cells^37^. We thus believe that the connectivity results will apply generally, although it will be important to measure the variability across individual cells, distinct animals, and locations in cortex. However, it is an interesting question if the connection specificity we observed is entirely determined by molecularly-defined cell types or if other factors such as developmental timing, exact cortical depth, or activity-dependent plasticity^103^ play a significant role in shaping these patterns. Such additional factors could interact with cell types to produce a combination discrete and continuous rules underlying synaptic connectivity, while this study focused only on discrete cell type and compartmental rules. Integrating the functional imaging from this same dataset^1^ could provide a valuable foundation to investigate these potential interactions.

This study focused on the architecture of classical synaptic connectivity. However, gap junctions, non-synaptic neurotransmitter release and neuromodulator release are all used by inhibitory interneurons, and enriching this data with such information will yield a more complete picture of functional interactions^104^. Even among synapses, this study only considered cells and connectivity within a narrow range of distances and limited volume. If cells change their connectivity with distance, as has been seen in excitatory neurons^105^, or if some regions of the axon were within the column but others were not, this would bias the observed connectivity distributions. Further, while the columnar approach was provided large samples of excitatory subgroups, it resulted in few examples for any given inhibitory cell type; some cell types such as chandelier cells were not observed at all. Extending a similar analysis across a much wider extent will be important for building a complete map of inhibitory cell types and firmly establishing the nature of inhibitory motif groups. The lack of segmentation in the top 10 *µm* of layer 1 truncates some apical tufts as well as limiting reconstruction quality of layer 1 interneurons. For those excitatory neurons with extensive apical tufts, particularly layer 2 and L5ET cells, the reconstructions here might miss both distinguishing characteristics and sources of inhibitory input in that region.

### D. Comparison to other cell type approaches

Recent surveys aimed at discovering excitatory cell types have used morphology alone^96, 106^, morphology and electrophysiology^14^, transcriptomics^20, 107^, or all three together using Patch-seq^17^. The M-types found here from morphology and synaptic properties generally agree with these other approaches, in particular distinguishing cells in upper layer and lower layer 2/3 and differentiating between projection subclasses. Transcriptomic studies have found multiple excitatory clusters in upper layer 2/3 in VISp^107, 108^, although it is not clear if they correspond to exactly two morphological clusters observed here. In other cases, the M-types described here can likely be divided further. For example, transcriptomic clustering subdivides layer 5 ET cells into three groups^22^, while we only have one. Improved sampling of rare cell types, incorporation of axonal morphology, and multimodal analysis will be important to help refine the landscape of cell types.

For inhibitory neurons, we focused on generating rich descriptions of individual neurons rather than clustering cell types, since we did not have enough examples of most specific cell types. However, in many cases there are striking morphological similarities between interneurons reconstructed here in EM and examples from the literature, such as particular layer 6 ascending translaminar basket cells^109, 110^ and Cell IDs 302999 and 305182. In addition, a companion study quantitatively matched EM reconstructions to Patch-seq reconstructions^31^ for a subset of SST neurons^57^, demonstrating that these matches can be algorithmically discovered with sufficient data.

### E. Peters’ rule and White’s exceptions

Since the seminal work of Braitenberg and Schüz^80^ and the experiments of Alan Peters^111^ and Edward White^112^, two ideas have been pitched as competing principles: One is that cell type connectivity simply reflects the distribution of pre- and post-synaptic elements of each cell type (“Peters’ rule”) and the other is that cells preferentially target certain cells over others (“White’s exceptions”). These concepts have been of great value in estimating the cortical wiring diagram^79, 113^ and offering null hypotheses for targeted studies^114^. Our data suggests both mechanisms are likely to be used by cortical neurons. While some inhibitory cell types present remarkably selective connectivity, other cell types appear to broadly target available partners. Importantly, some interneurons that connect with extremely specific connectivity, for example onto layer 2 M-types, have no evidence of selectivity beyond tight control of their axonal arbors. However, such tight spatial organization can follow synaptic specificity, for example if earlier in development or maturation, axonal exploration were followed by pruning branches with synapses onto non-preferred targets^115^. Our data suggest that neurons use a combination of spatial overlap, morphology, and synaptic selectivity together, with cell-type specific differences in the degree to which each aspect contributes to observed connectivity.

### F. A foundation for the multimodal study of cell types

This work provides a key step in dissecting the connectivity of cortical cell types^6^, but to make the most use of this type of data will require following in the footsteps of *Drosophila* and linking EM to genetic tools^116^. Patch-seq, which generates both transciptomic and morphological data, is a promising route to generating these correspondences. In addition to striking visual matches between cells reconstructed in EM and individual samples from a VISp Patch-seq data collection^31^, a companion study used established morphometric features to to quantitatively link EM reconstructions of layer 5 Martinotti cells to specific transcriptomic subtypes^57^. Currently, however, many cell subclasses determined based on transcriptomic or multimodal clusters have diverse morphologies and likely diverse connectivity^17, 31^. This suggests that the process of linking structural and molecular datasets should aim to become bi-directional, not only decorating EM reconstructions with transcriptomic information, but also using EM to identify cell types with distinct connectivity and analyzing Patch-seq data to identify distinguishing transcriptomic markers or collecting additional examples.

The anatomical data presented here are exceptionally rich, and this study offered just one approach to its analysis. To facilitate subsequent analysis of anatomy, connectivity, and ultrastructure, all EM data, segmentations, skeletons and tables of synapses and cell types are available are available via MICrONs-Explorer^1^. In addition to being a rich resource for detailed neuroanatomy, this highly-curated population within a larger volume will serve an important role for future analyses. The data presented here can serve as training data for anatomical classifiers^62, 117^ or improvements in automated proofreading^118^. Moreover, it can serve as a rich seed from which to consider other aspects of the cortical circuit in the same dataset, such as thalamic input, excitatory connectivity, and functional properties.

## Supporting information

Extended Data Cell Atlas

## Author Contributions

We use the CRediT system for author roles. Conceptualization: NMdC, CMSM, AB. Methodology: CMSM, NMdC, FC, AB. Software: CMSM, DB, SD, CJ, WS, FC. Validation: AB, NMdC, FC, CMSM, MT, JB, CG. Formal analysis: CMSM, FC, NMdC. Investigation: NMdC, FC, CMSM, AB, JB. Resources: JAB, MAC, AH, ZJ, KiL, KaL, RL, TM, EM, SSM, SM, BN, SP, WS, NT, WW, JW, FC, CMSM, JB, DB, DJB, DK, SK, GM, MT, RT, WY, SD, NMdC, RCR. Data curation: NMdC, FC, CMSM, AB, JB, CG, LE, SSe, MT. Writing - Original draft: NMdC, CMSM, FC. Writing - Review & editing: CMSM, FC, JB, RCR, NMdC, MT. Visualization: CMSM, FC. Supervision: NMdC, RCR, FC, HSS. Project administration: SSu, NMdC. Funding acquisition: NMdC, RCR, HSS, JR, AST.

## Acknowledgements

The work was supported by the Intelligence Advanced Research Projects Activity (IARPA) via Department of Interior/ Interior Business Center (DoI/IBC) contract numbers D16PC00003, D16PC00004, and D16PC0005. NMdC, FC and RCR also acknowledge support from NIH RF1MH125932 and from NSF NeuroNex 2 award 2014862. AST also acknowledges support from NSF NeuroNex grant 1707400 and from NIH (U19MH114830, R01 EY026927, T32-EY-002520-37).

The authors thank David Markowitz, the IARPA MICrONS Program Manager, who coordinated this work during all three phases of the MICrONS program, IARPA program managers Jacob Vogelstein and David Markowitz for conceiving of and advocating for the MICrONS program, and Wade Leonard and Jennifer Wang, IARPA SETAs, for their assistance. We thank Brock Wester, William Gray-Roncal, Sandy Hider, Tim Gion, Daniel Xenes, Jordan Matelsky, Caitlyn Bishop, Derek Pryor, Dean Kleissas, Luis Rodriguez and Miller Wilt from John Hopkins University Applied Physics lab for providing data assessments on the neural circuit reconstruction and infrastructure through BOSSdb. We also thank Frances Chance, Brad Aimone, Kristofor Carlson, Warren Davis, Kim Pfeiffer, David Robinson, Timothy Shead, Craig Vineyard and everyone at Sandia National Laboratories for their support and assistance.

The Allen Institute for Brain Science (AIBS) team thanks many people and teams who supported this project: Rob Young for managing the stitching and alignment pipeline at the AIBs; John Philips, Sill Coulter and the Program Management team at the AIBS for their guidance for project strategy and operations; Hongkui Zeng, Ed Lein, Christof Koch and Allan Jones for their support and leadership; Manufacturing and Processing Engineering team at the AIBS for their help in implementing the EM imaging and sectioning pipeline; Brian Youngstrom, Stuart Kendrick and the Allen Institute IT team for support with infrastructure, data management and data transfer; Facilities, Finance, and Legal teams at the AIBS for their support on the MICrONS contract; Stephan Saalfeld, Khaled Khairy and Eric Trautman for help with the parameters for 2D stitching and rough alignment of the dataset. Finally, we thank the Allen Institute for Brain Science founder, Paul G. Allen, for his vision, encouragement and support.

We would like to thank the “Connectomics at Google” team for developing Neuroglancer and computational resource donations, in particular J. Maitin-Shepard for authoring Neuroglancer and help creating the reformatted sharded multi-resolution meshes and imagery files used to display the data. We would also like to thank Amazon and the AWS Open Science platform for providing computational resources and Intel for their assistance.

The U.S. Government is authorized to reproduce and distribute reprints for Governmental purposes notwithstanding any copyright annotation thereon. Disclaimer: The views and conclusions contained herein are those of the authors and should not be interpreted as necessarily representing the official policies or endorsements, either expressed or implied, of IARPA, DoI/IBC, or the U.S. Government.

## Methods

This dataset was acquired, aligned, and segmented as part of the larger MICRONS project. Methods underlying dataset acquisition are described in detail elsewhere^1, 49–51^ and the primary data resource is described in a separate publication^1^. We repeat some of the methodological details for the dataset here for convenience.

### Animal preparation for EM

All animal procedures were approved by the Institutional Animal Care and Use Committee at the Allen Institute for Brain Science or Baylor College of Medicine. Neurophysiology data acquisition was conducted at Baylor College of Medicine prior to EM imaging, afterwards the mice were transferred to the Allen Institute in Seattle and kept in a quarantine facility for 1–3 days, after which they were euthanized and perfused. All results described here are from a single male mouse, age 64 days at onset of experiments, expressing GCaMP6s in excitatory neurons via SLC17a7-Cre and Ai162 heterozygous transgenic lines (recommended and generously shared by Hongkui Zeng at Allen Institute for Brain Science; JAX stock 023527 and 031562, respectively). Two-photon functional imaging took place between P75 and P80 followed by two-photon structural imaging of cell bodies and blood vessels at P80. The mouse was perfused at P87. Details of animal preparation are described in detail elsewhere^1^ and summarized below.

### Tissue preparation

After optical imaging at Baylor College of Medicine, candidate mice were shipped via overnight air freight to the Allen Institute. Mice were transcardially perfused with a fixative mixture of 2.5% paraformaldehyde, 1.25% glutaraldehyde, and 2 mM calcium chloride, in 0.08 M sodium cacodylate buffer, pH 7.4. A thick (1200 µm) slice was cut with a vibratome and post-fixed in perfusate solution for 12–48 h. Slices were extensively washed and prepared for reduced osmium treatment (rOTO) based on the protocol of Hua and colleagues^119^. All steps were performed at room temperature, unless indicated otherwise. 2% osmium tetroxide (78 mM) with 8% v/v formamide (1.77 M) in 0.1 M sodium cacodylate buffer, pH 7.4, for 180 minutes, was the first osmication step. Potassium ferricyanide 2.5% (76 mM) in 0.1 M sodium cacodylate, 90 minutes, was then used to reduce the osmium. The second osmium step was at a concentration of 2% in 0.1 M sodium cacodylate, for 150 minutes. Samples were washed with water, then immersed in thiocarbohydrazide (TCH) for further intensification of the staining (1% TCH (94 mM) in water, 40 °C, for 50 minutes). After washing with water, samples were immersed in a third osmium immersion of 2% in water for 90 minutes. After extensive washing in water, lead aspartate (Walton’s (20 mM lead nitrate in 30 mM aspartate buffer, pH 5.5), 50°, 120 minutes) was used to enhance contrast. After two rounds of water wash steps, samples proceeded through a graded ethanol dehydration series (50%, 70%, 90% w/v in water, 30 minutes each at 4 °C, then 3 x 100%, 30 minutes each at room temperature). Two rounds of 100% acetonitrile (30 minutes each) served as a transitional solvent step before proceeding to epoxy resin (EMS Hard Plus). A progressive resin infiltration series (1:2 resin:acetonitrile (e.g. 33% v/v), 1:1 resin:acetonitrile (50% v/v), 2:1 resin acetonitrile (66% v/v), then 2 x 100% resin, each step for 24 hours or more, on a gyrotary shaker) was done before final embedding in 100% resin in small coffin molds. Epoxy was cured at 60° for 96 hours before unmolding and mounting on microtome sample stubs. The sections were then collected at a nominal thickness of 40 nm using a modified ATUMtome (RMC/Boeckeler^49^) onto 6 reels of grid tape^49, 120^.

### Transmission electron microscopy imaging

The parallel imaging pipeline used in this study^49^ used a fleet of transmission electron microscopes that had been converted to continuous automated operation. It is built upon a standard JEOL 1200EXII 120kV TEM that had been modified with customized hardware and software, including an extended column and a custom electron-sensitive scintillator. A single large-format CMOS camera outfitted with a low distortion lens was used to grab image frames at an average speed of 100 ms. The autoTEM was also equipped with a nano-positioning sample stage that offered fast, high-fidelity montaging of large tissue sections and a reel-to-reel tape translation system that locates each section using index barcodes. During imaging, the reel-to-reel GridStage moved the tape and located the targeting aperture through its barcode and acquired a 2D montage. We performed image QC on all data and reimaged sections that failed the screening.

### Image processing: Volume assembly

The volume assembly pipeline is described in detail elsewhere^50, 51^. Briefly, the images collected by the autoTEMs are first corrected for lens distortion effects using non-linear transformations computed from a set of 10×10 highly overlapping images collected at regular intervals. Overlapping image pairs are identified within each section and point correspondences are extracted using SIFT features. Montage transformation parameters are estimated per image to minimize the sum of squared distances between the point correspondences between these tile images, with regularization. A downsampled version of these stitched sections are produced for estimating a per-section transformation that roughly aligns these sections in 3D. he rough aligned volume is rendered to disk for further fine alignment. The software tools used to stitch and align the dataset is available in our github repository (https://github.com/AllenInstitute/render-modules). To fine align the volume it was required to make the image processing pipeline robust to image and sample artifacts. Cracks larger than 30 um (in 34 sections) were corrected by manually defining transforms. The smaller and more numerous cracks and folds in the dataset were automatically identified using convolutional networks trained on manually labeled samples using 64 × 64 × 40 nm^3^ resolution image. The same was done to identify voxels which were considered tissue. The rough alignment was iteratively refined in a coarse-to-fine hierarchy^121^, using an approach based on a convolutional network to estimate displacements between a pair of images^122^. Displacement fields were estimated between pairs of neighboring sections, then combined to produce a final displacement field for each image to further transform the image stack. Alignment was first refined using 1024 × 1024 × 40 nm^3^ images, then 64 × 64 × 40 nm^3^ images. The composite image of the partial sections was created using the tissue mask previously computed.

### Image processing: Segmentation

The image segmentation pipeline is fully described in Macrina et al^50^. Remaining misalignments were detected by cross-correlating patches of image in the same location between two sections, after transforming into the frequency domain and applying a high-pass filter. Combining with the tissue map previously computed, a “segmentation output mask” was generated that sets the output of later processing steps to zero in locations with poor alignment. Using previously described methods^123^, a convolutional network was trained to estimate inter-voxel affinities that represent the potential for neuronal boundaries between adjacent image voxels. A convolutional network was also trained to perform a semantic segmentation of the image for neurite classifications, including (1) soma+nucleus, (2) axon, (3) dendrite, (4) glia, and (5) blood vessel. Following the methods described in Wu et al^124^, both networks were applied to the entire dataset at 8 × 8 × 40 nm^3^ in overlapping chunks to produce a consistent prediction of the affinity and neurite classification maps and the segmentation output mask was applied to predictions. The affinity map was processed with a distributed watershed and clustering algorithm to produce an over-segmented image, where the watershed domains are agglomerated using single-linkage clustering with size thresholds^125, 126^. The over-segmentation was then processed by a distributed mean affinity clustering algorithm^125, 126^ to create the final segmentation.

For synapse detection and assignment, a convolutional network was trained to predict whether a given voxel participated in a synaptic cleft. Inference on the entire dataset was processed using the methods described in Wu et al^124^ using 8 × 8 × 40 nm^3^ images. These synaptic cleft predictions were segmented using connected components, and components smaller than 40 voxels were removed. A separate network was trained to perform synaptic partner assignment by predicting the voxels of the synaptic partners given the synaptic cleft as an attentional signal^127^. This assignment network was run for each detected cleft, and coordinates of both the presynaptic and postsynaptic partner predictions were logged along with each cleft prediction.

For nucleus detection^1^ a convolutional network was trained to predict whether a voxel participated in a cell nucleus. Following the methods described in Wu et al^124^, a nucleus prediction map was produced on the entire dataset at 64 × 64 × 40 nm^3^.

### Column description and cell classes

The column borders were found by manually identifying a region in primary visual cortex that was far from both dataset boundaries and the boundaries with higher order visual areas. A 100 *µm ×* 100 *µm* box was placed based on layer 2/3 and was extended along the negative y axis of the dataset.

While analyzing data, we observed that deep layer neurons had apical dendrites that were not oriented along the most direct pia-to-white-matter direction, and we adapted the definition of the column to accommodate these curved neuronal streamlines. Using a collection of layer 5 ET cells, we placed points along the apical dendrite to the cell body and then along the primary descending axon towards white matter. We computed the slant angle as two piecewise linear segments, one along the negative y axis to lower layer 5 where little slant was observed, and one along the direction defined by the vector averaged direction of the labeled axons. We believe the slant to be a biological feature of the tissue and not a technical artifact for several reasons:

1. The curvature is not aligned to a sectioning plane or associated with shearing or other distortion in the imagery, making it unlikely to be a result of the alignment process.
2. Blood vessel segmentation does not show a large correlated distortion in deep layers, making it unlikely to be a result of mechanical stress on the tissue (see https://ngl.microns-explorer.org/#!gs://microns-static-links/mm3/blood_vessels.json). Moreover, it is unclear why such stress would affect only layer 5b and below.
3. Individual examples of neurons with slanted morphologies can be found among single cell reconstructions in the literature, for example several descending bipolar VIP interneurons and layer 6 pyramidal cells in [31]. It is not possible to determine if these individual cases correspond to a larger population of correlated arbors, but it suggests these morphologies are not atypical.
4. Similar curvature has been observed in other large EM datasets from visual cortex.

Using these boundaries and previously computed nucleus centroids^1^, we identified all cells inside the columnar volume. Coarse cell classes (excitatory, inhibitory, and non-neuronal) were assigned based on brief manual examination and rechecked by subsequent proofreading and automated cell typing^62^. To facilitate concurrent analysis and proofreading, we split all false merges connecting any column neurons to other cells (as defined by detected nuclei) before continuing with other work.

### Proofreading

Proofreading was performed primarily by five expert neuroanatomists using the PyChunkedGraph^53, 54^ infrastructure and a modified version of Neuroglancer^128^. Proofreading was aided by on-demand highlighting of branch points and tips on user-defined regions of a neuron based on rapid skeletonization (https://github.com/AllenInstitute/Guidebook). This approach quickly directed proofreader attention to potential false merges and locations for extension, as well as allowed a clear record of regions of an arbor that had been evaluated.

For dendrites, we checked all branch points for correctness and all tips to see if they could be extended. False merges of simple axon fragments onto dendrites were often not corrected in the raw data, since they could be computationally filtered for analysis after skeletonization (see below). Detached spine heads were not comprehensively proofread, and previous estimates place the rate of detachment at approximately 10–15%. Using this method, dendrites could be proofread in approximately ten minutes per cell.

For inhibitory axons, we began by “cleaning” axons of false merges by looking at all branch points. We then performed extension of axonal tips until either their biological completion or data ambiguities, particularly emphasizing all thick branches or tips that were well-suited to project to new laminar regions. For axons with many thousand synaptic outputs, we followed some but not all tips to completion once major branches were cleaned and established. For smaller neurons, particularly those with bipolar or multipolar morphology, most tips were extended to the point of completion or ambiguity. Axon proofreading time differed significantly by cell type not only because of differential total axon length, but axon thickness differences that resulted in differential quality of autosegmentations, with thicker axons being of higher initial quality. Typically, inhibitory axon cleaning and extension took 3–10 hours per neuron.

### Manual cell subclass and layer labels

Expert neuroanatomists further labeled excitatory and inhibitory neurons into subclasses. Layer definitions were based on considerations of both cell body density (in analogy with nuclear staining) supplemented by identifying kinks in the depth distribution of nucleus size near expected layer boundaries^62^.

For excitatory neurons, the categories used were: Layer 2/3-IT, Layer 4-IT, Layer 5-IT, Layer 5-ET, Layer 5-NP, Layer 6-IT, Layer 6-CT, and Layer 6b (“L6-WM”) cells. Excitatory expert labels did not affect analysis, but were used as the basis for naming morphological clusters. Layer 2/3 and upper Layer 4 cells were defined on the basis of dendritic morphology and cell body depth. Layer 5 cells were similarly defined by cell body depth, with projection subclasses distinguished by dendritic morphology following Gouwens, Sorenson, and Berg, 2019^14^ and classical descriptions of thick (ET) and thin-tufted (IT) cells. Layer 5 ET cells had thick apical dendrites, large cell bodies, numerous spines, a pronounced apical tuft, and deeper ET cells had many oblique dendrites. Layer 5 IT cells had more slender apical dendrites and smaller tufts, fewer spines, and fewer dendritic branches overall. Layer 5 NP cells corresponded to the “Spiny 10” subclass described in Gouwens, Sorenson, and Berg; these cells had few basal dendritic branches, each very long and with few spines or intermediate branch points. Layer 6 neurons were defined by cell body depth, but only some cells were able to be labeled as IT or CT by human experts. Layer 6 pyramidal cell with stellate dendritic morphology, inverted apical dendrites, or wide dendritic arbors were classified as IT cells. Layer 6 pyramidal cells with small and narrow basal dendrites, an apical dendrite ascending to Layer 4 or Layer 1, and a myelinated primary axon projecting into white matter were labeled as CT cells.

For inhibitory neurons, manual cell typing considered axonal and dendritic morphology as well as connectivity. Cells that primarily contacted soma or perisomatic regions were labeled as basket cells (BC). Cells that made arbors that extended up to layer 1 or formed a dense plexus and primarily targeted distal dendrites were labeled as putative SST cells. Cells that remained mostly in layer 1 or had extensive arborization and many non-synaptic boutons were labeled as putative Id2 or neurogliaform cells. Finally, cells with a bipolar dendritic morphology or a multipolar dendritic morphology and output onto other inhibitory neurons were labeled as putative VIP cells. Several cells, particularly in layer 6, had an ambiguous subclass assignment, typically when their connectivity was not basket-like but their morphology was also not similar to upper layer Martinotti or non-Martinotti cells.

### Skeletonization

To rapidly skeletonize dynamic data, we took advantage of the PyChunkedGraph data structure that collects all supervoxels belonging to the same neuronal segmentation into 2 *µm ×* 2 *µm ×* 20 *µm* “chunks” with a unique id and precisely defined topological adjacency with neighboring chunks of the same object. Each chunk is called a “level 2 chunk” and the complete set of chunks for a neuron and their adjacency we call the “level 2 graph,” based on its location in the hierarchy of the PyChunkedGraph data structure^54^. We precompute and cache a representative central point in space, the volume, and the surface area for each level 2 chunk and update this data when new chunks are created due to proofreading edits. Using the level 2 graph and assigning edge lengths corresponding to the distance between the representative points for each vertex (i.e. each level 2 chunk), we run the TEASAR^129^ algorithm (10 *µm* invalidation radius) to extract a loop-free skeleton. Each of the level 2 vertices removed by the TEASAR algorithm is associated with its closest remaining skeleton, making it possible to map surface area and volume data to the skeleton. Typical edges between skeleton vertices are about 1.7 *µm*, and new skeletons can be computed *de novo* in approximately 10 seconds, making them useful for analysis over length scales of tens of *µm* or larger.

To represent the cell body, an additional vertex was placed at the location of the nucleus centroid and all vertices within an initial radius and topologically connected to centroid were collapsed into this vertex with associated data mapping. The radius was determined for each neuron separately by consideration of the volume of each cell body. A companion work^62^ computed the volume of each cell body, and we generated an effective radius based on the sphere with the same volume. To ensure that our values captured potentially lopsided cell bodies, we padded this effective radius by an additional factor of 1.25. Skeletons were rooted at the cell body, with “downstream” meaning away from soma and “upstream” meaning towards soma. Each synapse was assigned to skeleton vertices based on the level 2 chunk of its associated supervoxel. For each unbranched segment of the skeleton (i.e. between two branch points or between a branch point and end point), we computed an approximate radius *r* based on a cylinder with the same path length *L* and total volume *V* associated with that segment 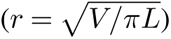.

### Axon/dendrite classification

To detect axons, we took advantage of the skeleton morphology, the location of presynaptic and postsynaptic synapses, and the clear segregation between inputs and outputs of cortical neurons. For inhibitory cells, we used synapse flow centrality^130^ to identify the start of the axon as the location of maximum paths along the skeleton between sites of synaptic input and output. Two inhibitory neurons had two distinct, biologically correct axons after proofreading (Cell IDs 258362 and 307059). For these cells we ran this method twice, masking off the axon found after the first run, in order to identify both. For excitatory neurons that did not have extended axons, there were often insufficient synaptic outputs on their axon for this approach to be reliable. Excitatory neurons with a segregation index^130^ of 0.7 (on a scale with 0 indicating random distribution of input and output synapses and 1 indicating perfect input/output segregation) or above were considered well-separated and the synapse flow centrality solution was used. For cells with a segregation index less than 0.7, we instead looked for branches near the soma with few synaptic inputs. Specifically, we took identified all skeleton vertices within 30 *µm* from the cell body and looked at the distinct branches downstream of this region. For each branch, we computed the total path length and the total number of synaptic inputs in order to get a linear input density. Branches with a path length both more than 20 *µm* and with an input density less than 0.1 synaptic inputs per *µm* were labeled as being axonal and filtered out of subsequent analysis.

We further filtered out any remaining axon fragments merged onto pyramidal cell dendrites using a similar approach. We identified all unbranched segments (regions between two branch points or between a branch point and end point) on the non-axonal region of the skeleton and computed their input synapse density. Starting from terminal segments (i.e. those with no downstream segments), we labeled a segment as a “false merge” if it had an input density less than 0.1 synaptic inputs per *µm*. This process iterated across terminal segments until all remaining had an input density of at least 0.1 inputs per *µm*. Falsely merged segments were masked out of the skeleton for all analysis.

### Excitatory dendrite compartments

We assigned all synaptic inputs onto excitatory neurons to one of four compartments: soma, proximal dendrite, distal basal dendrite, and distal apical dendrite. The most complex part was distinguishing the basal dendrite from the apical dendrite. While easy in most cases for neurons in layer 3–5 due to the consistent nature of apical dendrites being single branches reaching toward layer 1, this is not true everywhere. In upper layer 2/3, cells often have multiple branches in layer 1 equally consistent with apical dendrites and in layer 6 there are often cells with apical dendrites that stop in layer 4, that point toward white matter, or even that lack a clear apical branch entirely. To objectively and scalably define apical dendrites, we built a classifier that could detect between 0–3 distinct apical branches per cell. Following the intuition from neuroanatomical experts, we used features based on the branch orientation, location in space, relative location compared to the cell body, and branch-level complexity. Specifically, we trained a random forest classifier to predict whether a skeleton vertex belonged to an apical dendrite based on several features: Depth of vertex, depth of soma, difference in depth between soma and vertex, vertex distance to soma along the skeleton, vertex distance to farthest tip, normalized vertex distance to tip (between 0 and 1), tortuosity of path to root, number of branch points along the path to root, radial distance from soma, absolute distance from soma, and angle relative to vertical between the vector from soma to vertex. We aggregated predictions within each branch by summing the log odds ratio from the model prediction, with the net log-odds ratio saturating at *±*200.

Finally, for each branch *i* with aggregated odds ratio *R_i_*, we compare branches to one another via a soft-max operation: 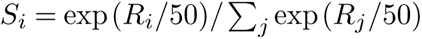. Branches with a maximum tip length of less than 50 *µm* were considered too short to be a potential apical dendrite and excluded from consideration and not included in the denominator. Branches with both *R_i_ >* 0 (evidence is positive towards being apical) and *S_i_ >* 0.25 (allowing no more than 4 apical branches with equal weight to be possible, and more likely 2-3 at most). Training data was selected from an initial 50 random cells, followed by an additional 33 cells chosen representing cases where the classifier did not perform correctly. Performance on both random and difficult cells had an F1 score of 0.9297 (86 true positives, 599 true negatives, 2 false positives, and 11 false negatives) based on leave-one-out cross validation, with at least one apical dendrite correctly classified for all cells.

Compartment labels were propagated to synapses based on the associated skeleton vertices. Soma synapses were all those associated with level 2 chunks within the soma collapse region (see Skeletonization section). Proximal dendrites were those outside of the soma, but within 50 *µm* after the start of the branch. Distal basal synapses were all those associated with vertices more distant than the proximal threshold, but not on an apical branch. Apical synapses were all those associated with vertices more distant than the proximal threshold and on an apical branch.

### Inhibitory feature extraction and clustering

Many classical methods of distinguishing interneuron classes are based on how cells distribute their synapses across target compartments. Following proofreading, expert neuroanatomists attempted to classify all inhibitory neurons broadly into “basket cells,” “SST-like cells”, “VIP-like cells”, and “neurogliaform/layer 1” cells based on connectivity properties and morphology. While 150 cells were labeled on this basis, an additional 13 neurons were considered uncertain (primarily in layer 6) and in some cases manual labels were low confidence. To classify inhibitory neurons in a data driven manner, we thus measured four properties of how cells distribute their synaptic outputs:

1. The fraction of synapses onto inhibitory neurons.
2. The fraction of synapses onto excitatory neurons that are onto soma.
3. The fraction of synapses onto excitatory neurons that are onto proximal dendrites.
4. The fraction of synapses onto excitatory neurons that are onto distal apical dendrites.

Because the fraction of synapses targeting all compartments sums to one, the last remaining property, synapses onto distal basal dendrites, was not independent and thus was measured but not included as a feature. Inspection of the data suggested two additional properties that characterized synaptic output across inhibitory neurons:

5. The fraction of synapses that are part of multisynaptic connections, those with at least two synapses between the same presynaptic neuron and target neuron.
6. The fraction of multisynaptic connection synapses that were also within 15 *µm* of another synapse with the same target, as measured between skeleton nodes. Note that we robustness of this parameter and found that intersynapse distances between from 5 to more than 100 *µm* have qualitatively similar results.

Using these six features, we trained a linear discriminant classifier on cells with manual annotations and applied it to all inhibitory cells. Differences from manual annotations were treated not as inaccurate classifications, but rather a different view of the data.

### Excitatory feature extraction and clustering

To characterize excitatory neuron morphology, we computed features based only on excitatory neuron dendrites and soma. The features were:

1. Median distance from branch tips to soma per cell.
2. Median tortuosity of the path from branch tips to soma per cell. Tortuosity is measured as the ratio of path length to the Euclidean distance from tip to soma centroid.
3. Number of synaptic inputs on the dendrite.
4. Number of synaptic inputs on the soma.
5. Net path length across all dendritic branches.
6. Radial extent of dendritic arbor. We define “radial distance” to be the distance within the same plane as the pial surface. For every neuron, we computed a pia-to-white-matter line, including slanted region in deep layers, passing through its cell body. For each skeleton vertex, we computed the radial distance to the pia-to-white-matter line at the same depth. To avoid any outliers, the radial extent of the neuron was defined to be the 97th percentile distance across all vertices.
7. Median distance to soma across all synaptic inputs.
8. Median synapse size of synaptic inputs onto the soma.
9. Median synapse size of synaptic inputs onto the dendrites.
10. Dynamic range of synapse size of dendrite synaptic inputs. This was measured as the difference between 95th and 5th percentile synapse sizes.
11. Shallowest extent of synapses, based on the 5th percentile of synapse depths.
12. Deepest extent of synapses, based on the 95th percentile of synapse depths.
13. Vertical extent of synapses, based on the difference between 95th and 5th percentile of synapse depths.
14. Median linear density of synapses. This was measured by computing the net path length and number of synapses along 50 depth bins from layer 1 to white matter and computing the median. A linear density was found by dividing synapse count by path length per bin, and the median was found across all bins with nonzero path length.
15. Median radius across dendritic skeleton vertices. To avoid the region immediately around the soma from having a potential outlier effect, we only considered skeleton vertices at least 30 *µm* from the soma.

Three additional sets of features used component decompositions. To more fully characterize the absolute depth distribution of synaptic inputs, for each excitatory neuron, we computed the number of synapses in each of 50 depth bins from the top of layer 1 to surface of white matter (bin width *≈* 20 *µm*). We z-scored synapse counts for each cell and computed the top six components using SparsePCA. The loadings for each of these components based on the net synapse distribution were used as features.

To characterize the distribution of synaptic inputs relative to the cell body instead of cortical space, we computed the number of synapses in 13 soma-adjusted depth bins starting 100 *µm* above and below the soma. As before, synapse counts were z-scored and we computed the top five components using SparsePCA. The loadings for each of these components were used as additional features.

To characterize the relationship with branching to distance, we measured the number of distinct branches as a function of distance from the soma at ten distances, every 30 *µm* starting at 30 *µm* from the soma and continuing to 300 *µm*. For robustness relative to precise branch point locations, the number of branches were computed by finding the number of distinct connected components of the skeleton found in the subgraph formed by the collection of vertices between each distance value and 10 *µm* toward the soma. We computed the top three singular value components of the matrix based on branch count vs distance for all excitatory neurons, and the loadings were used as features.

All features were computed after a rigid rotation of 5 degrees to flatten the pial surface and translation to zero the pial surface on the y axis. Features based on apical classification were not explicitly used to avoid ambiguities based on both biology and classification.

Using this collection of features, we clustered excitatory neurons by running phenograph^75^ 500 times with 95% of cells included each time. Phenograph finds a nearest neighborhood graph based on proximity in the feature space and clusters by running the Leiden algorithm for community detection on the graph. Here, we used a graph based on 10 nearest neighbors and clustered with a resolution parameter of 1.3. These values were chosen to consistently separate layer 5 ET, IT, and NP cells from one another, a well established biological distinction. A co-clustering matrix was assembled with each element corresponding to the number of times two cells were placed in the same cluster. To compute the final consensus clusters, we performed agglomerative clustering with complete linkage based on the co-clustering matrix, with the target number of clusters set by a minimum Davies-Bouldin score and a maximum Silhouette score. Clusters were then named based on the most frequent manually defined cell type within the cluster and reordered based on median soma depth. The labeling of cells as layer 2 and layer 3 was formed on the basis of soma depth and a morphology with a relatively flat morphology often with no distinct apical trunk, although often apical-tuft-like branches emitting directly from the cell body. The L2c subclass was ambiguously defined between the two categories, with cells that had a distinct apical trunk, but with connectivity and other properties seemed more similar to layer 2 subclasses.

To compute the importance of each feature for each M-type, for each M-type we trained a random forest classifier to predict whether or not a cell belonged to it using scikit-learn^131^. Because the classes were strongly imbalanced, we used SMOTE resampling to over-sample datapoints from the smaller class. We used the Mean Decrease in Impurity metric, which quantifies how often a given feature was used in the decision tree ensemble.

### Inhibitory connectivity and selectivity

To measure intracolumnar inhibitory connectivity, we first restricted synaptic outputs to the axon of each inhibitory neuron as we have not observed any correctly classified synaptic outputs on dendritic arbors in this dataset. One cell with fewer than 30 synaptic outputs was omitted due to size. All remaining synaptic outputs across all interneurons were then filtered to include only those that target cells within the column, unless otherwise specified. Each output synapse was also labeled with the target skeleton vertex, dendritic compartment, and M-type of the target neuron based on the compartment definitions above.

For measuring the synaptic output budget across cell types across the dataset (i.e. inside and outside the column), we used a hierarchical classifier based on a collection of perisomatic features that was trained on the data-driven clustering from the column sample^62^. Only synapses onto object segmentation associated with a single nucleus and a cell type classification were used. Although most of these additional targets are not proofread, estimates based on proofread neurons suggest that 99% of non-proofread input synapses are accurate^62^.

To measure inhibitory selectivity within the column, we compared the M-type distribution of its synaptic outputs to the M-type distribution of synaptic inputs according to a null model accounting for cell abundance, synapse abundance, and depth. We first generated a baseline distribution of all 4,504,935 somatic or dendritic synaptic inputs to all column cells, where each synapse was associated with a precise depth, target compartment, and an M-type. We discretized synaptic inputs into 50 depth bins spanning pia to white matter, each covering *≈* 20*µm*, and each of the five compartments: soma, proximal dendrite, basal dendrite, apical dendrite, or inhibitory neuron. For each interneuron, we similarly discretized its synaptic output into the same bins, compartments, and M-types. To generate a randomized output distribution preserving both observed depth and compartment distributions, we randomly picked synapses from the baseline distribution with the observed depth bins and compartment targets but without regard to M-type. We computed 10,000 randomized distributions per interneuron. To get a preference index, we compared the observed number of synapses onto a given M-type to the median of the number of synapses from the shuffle distribution. To get a significance for the preference index for a given M-type, we directly computed the two-sided p-value of the observed number of synapses relative to the shuffle distribution for that M-type. P-values were corrected for multiple comparisons using the Holm-Sidak method within each interneuron for those M-types with non-zero potential connectivity. Selectivity was only measured within the column because we did not generate compartment labels for unproofread dendrites outside of the column.

On connectivity cards, we also show a similar preference index based on compartment rather than M-type. In that case, the shuffled distribution preserves observed depth and M-type output distributions, but not compartments.

### Software and data availability

Data for this paper was analyzed at materialization version 795. Synapse tables for column cells, neuronal skeletons, and tables for manual and automatic cell types and connectivity groups are available at https://doi.org/10.5281/zenodo.7641780. Analysis code will be made publicly available on an Allen Institute Github repository (forthcoming). EM imagery and an older version of segmentation is publicly available via https://www.microns-explorer.org/cortical-mm3 and will be updated closer to publication. All analysis was performed in Python 3.9 using custom code, making extensive use of CAVEclient (https://github.com/seung-lab/CAVEclient) and CloudVolume^132^ to interact with data infrastructure, MeshParty^133^ to analyze skeletons, and libraries Matplotlib^134^, Numpy^135^, Pandas^136^, Scikit-learn^131^, Scipy^137^, stats-models^138^ and VTK^139^ for general computation, machine learning and data visualization.

## Supplemental Information

**Extended Data Figure 1.**
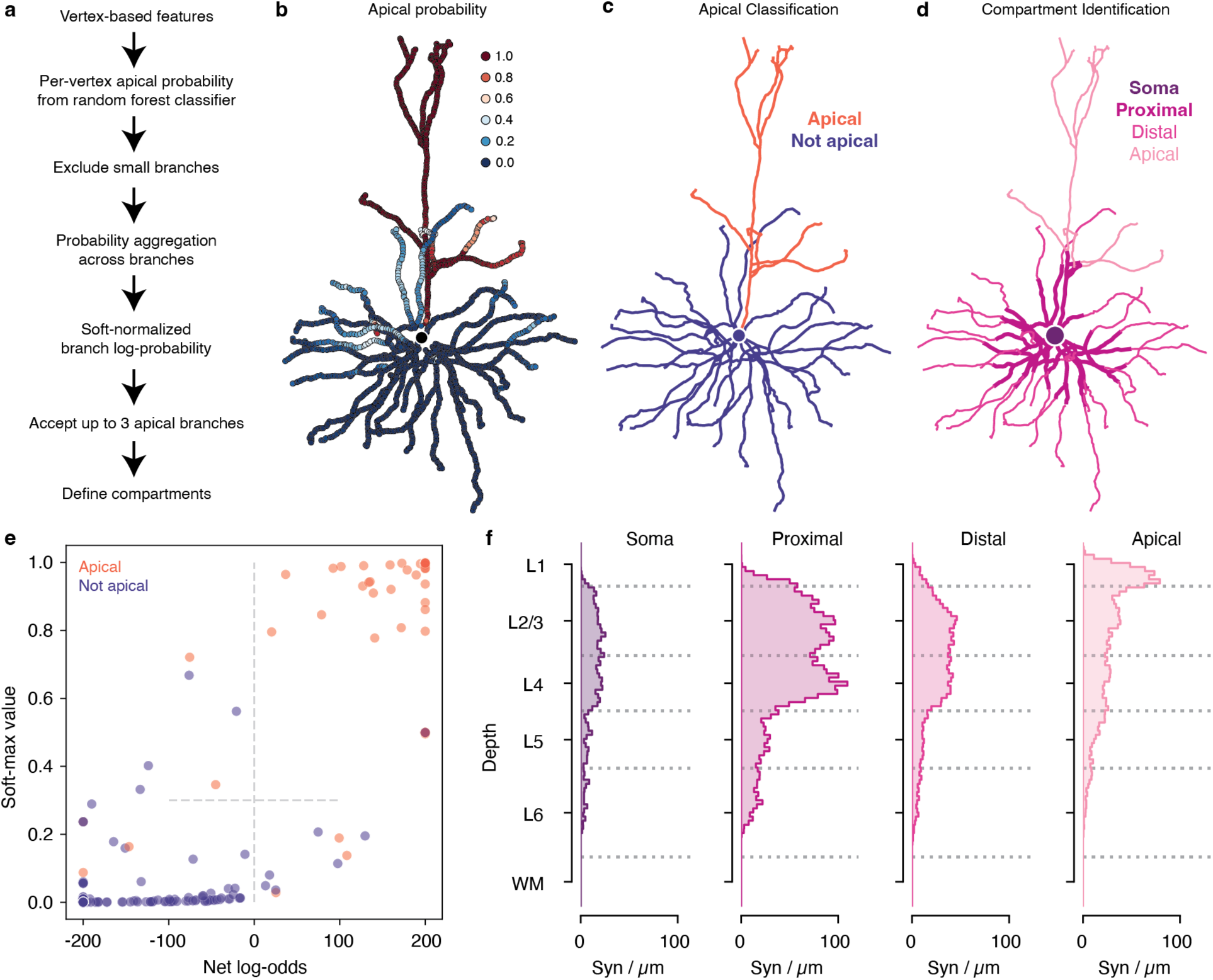
Compartment classification pipeline. **a)** Description of the compartment classification pipeline. **b–d)** Pipeline applied to an example layer 3 pyramidal cell. **b)** Apical probability per vertex. **c)** Branch-level apical classification. **d)** Final organization into four dendritic compartments based on apical classification and distance rules. **e)** Quantification of quality of apical branch classification based on leave-one-out classification with a training set based on 50 randomly selected cells and 23 cells chosen to improve difficult classifications. Each dot is a branch of a test pyramidal cell, colored red if apical and blue if not apical. X-axis is the net log-odds of the branch being apical (capped at *±*200) and the y-axis is the relative apical quality based on a soft-max operation (see Methods for details). Branches in the upper right quadrant were classified as apical. The method was able to correctly classify at least one apical branch for all cells, and “false positives” were often associated with borderline cases. **f)** Distribution of synaptic inputs onto excitatory neurons with depth by dendritic compartment. Values are based on counting synapses in bins at a given depth, but at any location laterally.

**Extended Data Figure 2.**
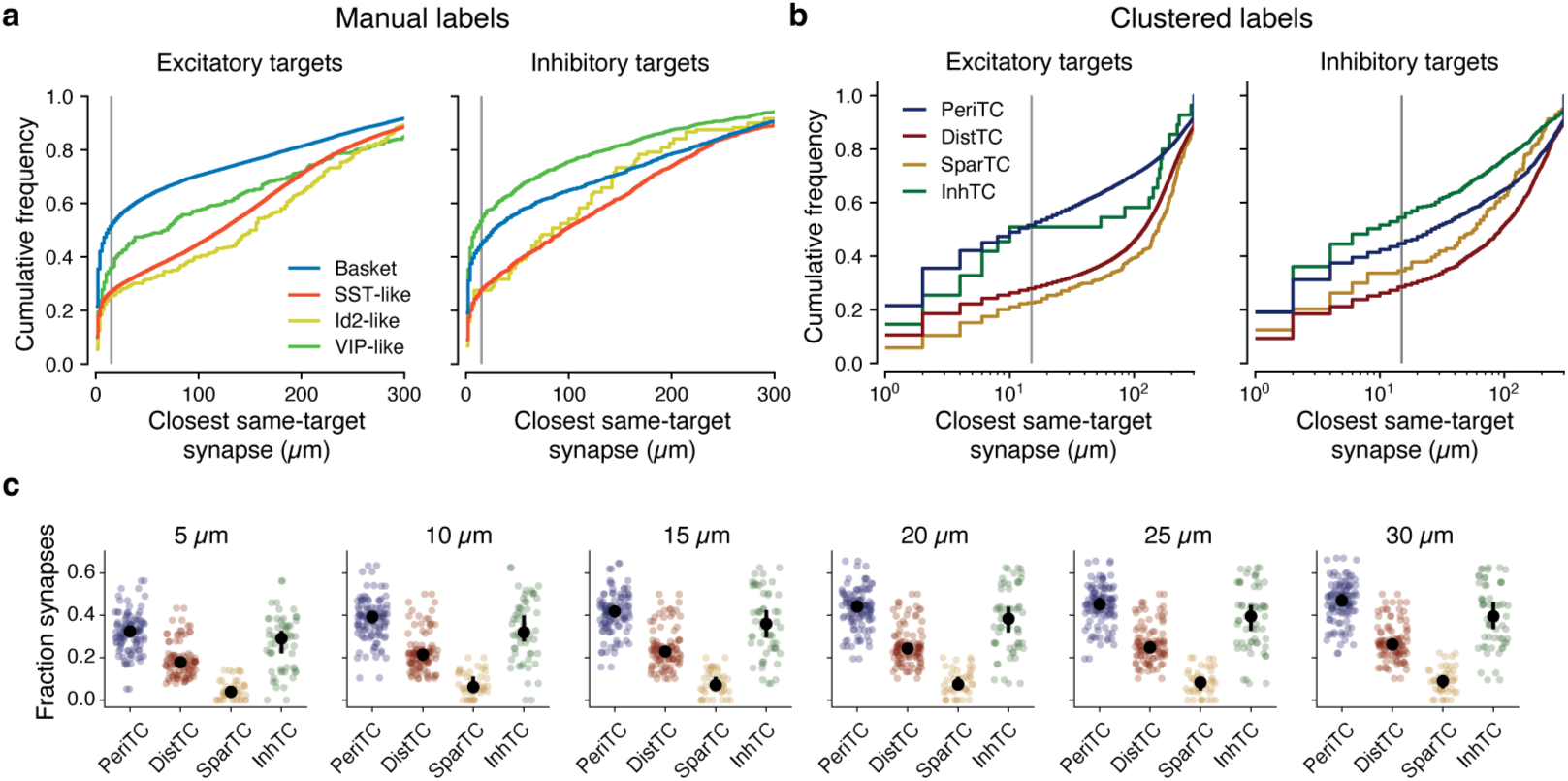
Closest distances between synapses in multisynaptic connectincs. **a)** Cumulative distributions of the closest synapse onto the same target along the axonal arbor per manually labeled inhibitory neuron subclass. Excitatory (left) and inhibitory (right) targets shown separately. Vertical gray line indicates the value used for “clumpiness” in the main text. **b)** Same as **a**, but for the cluster-based labels and with log scale to highlight shorter distances. **c)** The “clumpiness” metric using different distance thresholds. The qualitative relationships are extremely robust to distance thresholds.

**Extended Data Figure 3.**
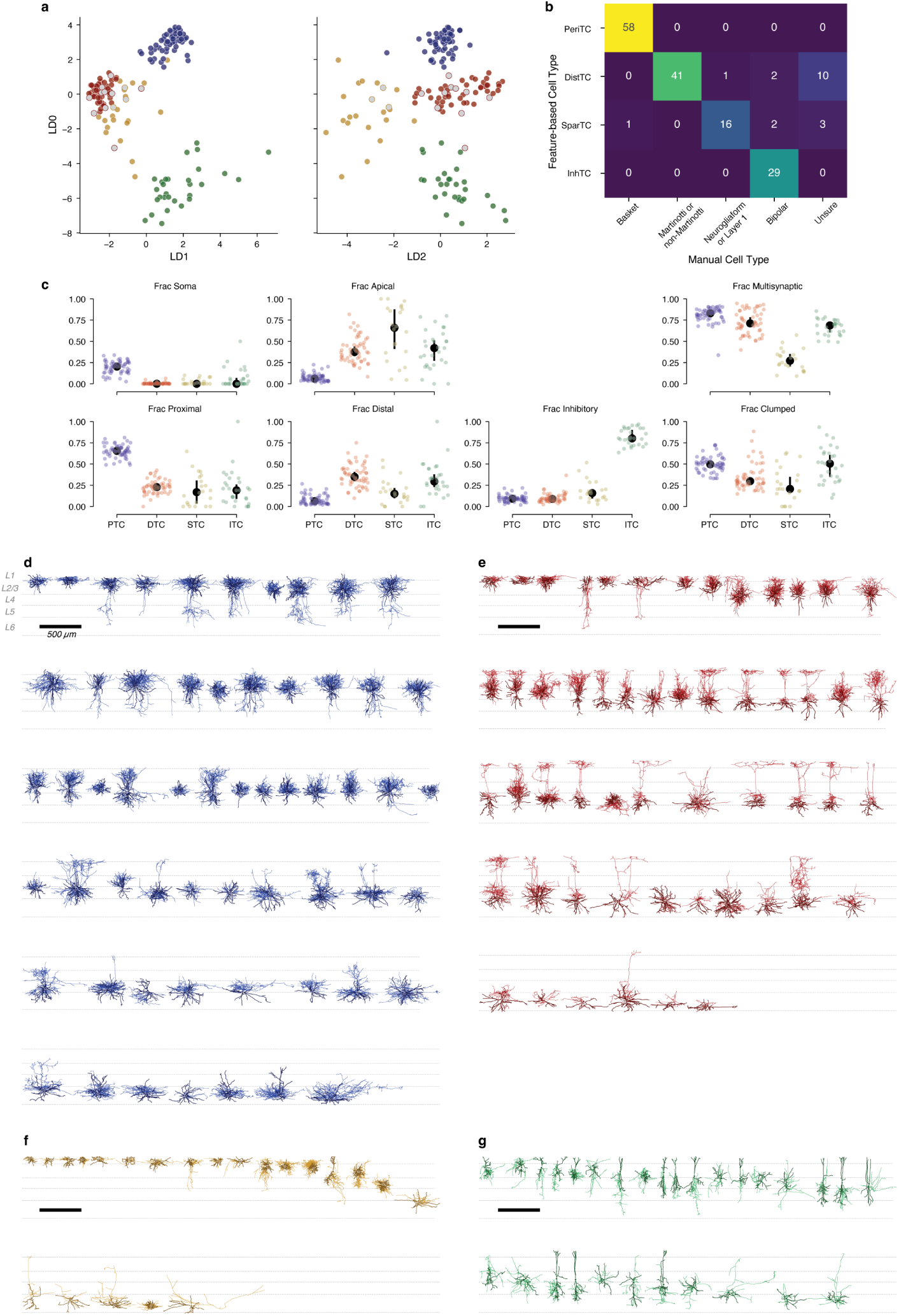
Inhibitory neuron properties. **a)** Projections of all analyzed interneurons (n=163) projected on a 3-d space based on linear discriminant analysis (LDA) using connectivity features (shown in **c**). Fully colored dots indicate manually classified cells used as training data for LDA, while dots with grey centers were labeled based on this classification. **b)** Matrix showing relationship between anatomical subclasses and manual classifications. **c)** Individual connectivity features, organized by subclass. Colored dots are individual cells, black dots indicate median with error bars showing a bootstrapped 95% confidence interval. **d–g)** Morphology of all PeriTCs (**d**), DistTCs (**e**), SparTCs (**f**), and InhTCs (**g**). Scale bars are 500 *µm*. Dark and thick lines are dendrite, thinner and lighter are axon. Cells are ordered by soma depth.

**Extended Data Figure 4.**
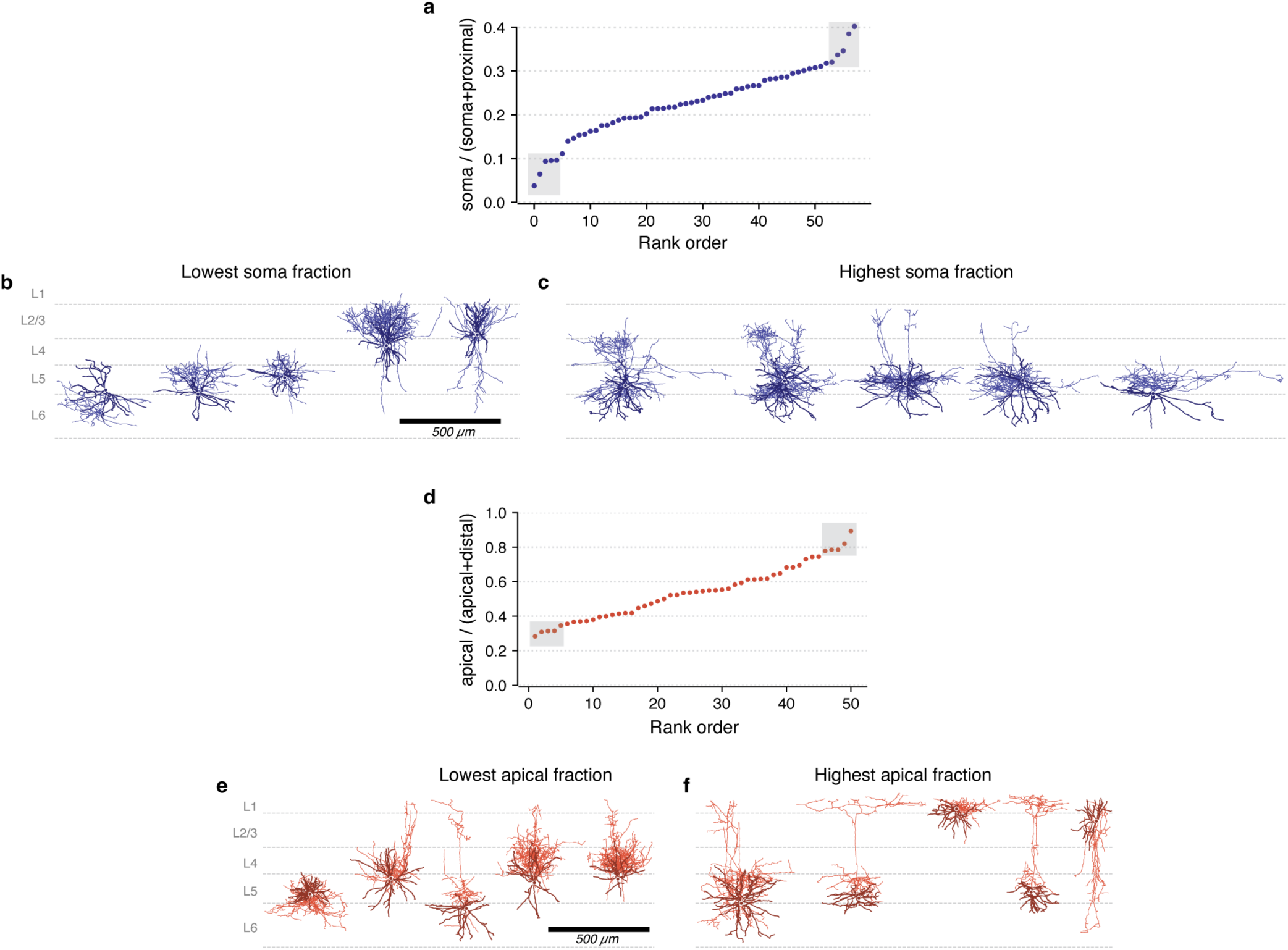
Diversity of compartment targeting within perisomatic targeting cells and distal dendrite targeting cells. **a)** Among PeriTCs, the rank-ordered fraction of output synapses targeting somata as a fraction of synapses targeting either somata or proximal dendrites (but not distal or apical dendrites). Boxes indicate the top and bottom five cells, shown below. **b)** The five PeriTC cells with the lowest soma fraction, ordered as in **a**. **c)** The five PeriTC cells with the highest soma fraction, ordered as in **a**. **d)** Among DistTCs, the rank-ordered fraction of output synapses targeting apical synapses among only those synapses targeting apical or distal dendrites. **e)** The five DistTC cells with the lowest apical fraction, ordered as in **a**. **f)** The five DistTC cells with the highest apical fraction, ordered as in **a**.

**Extended Data Figure 5.**
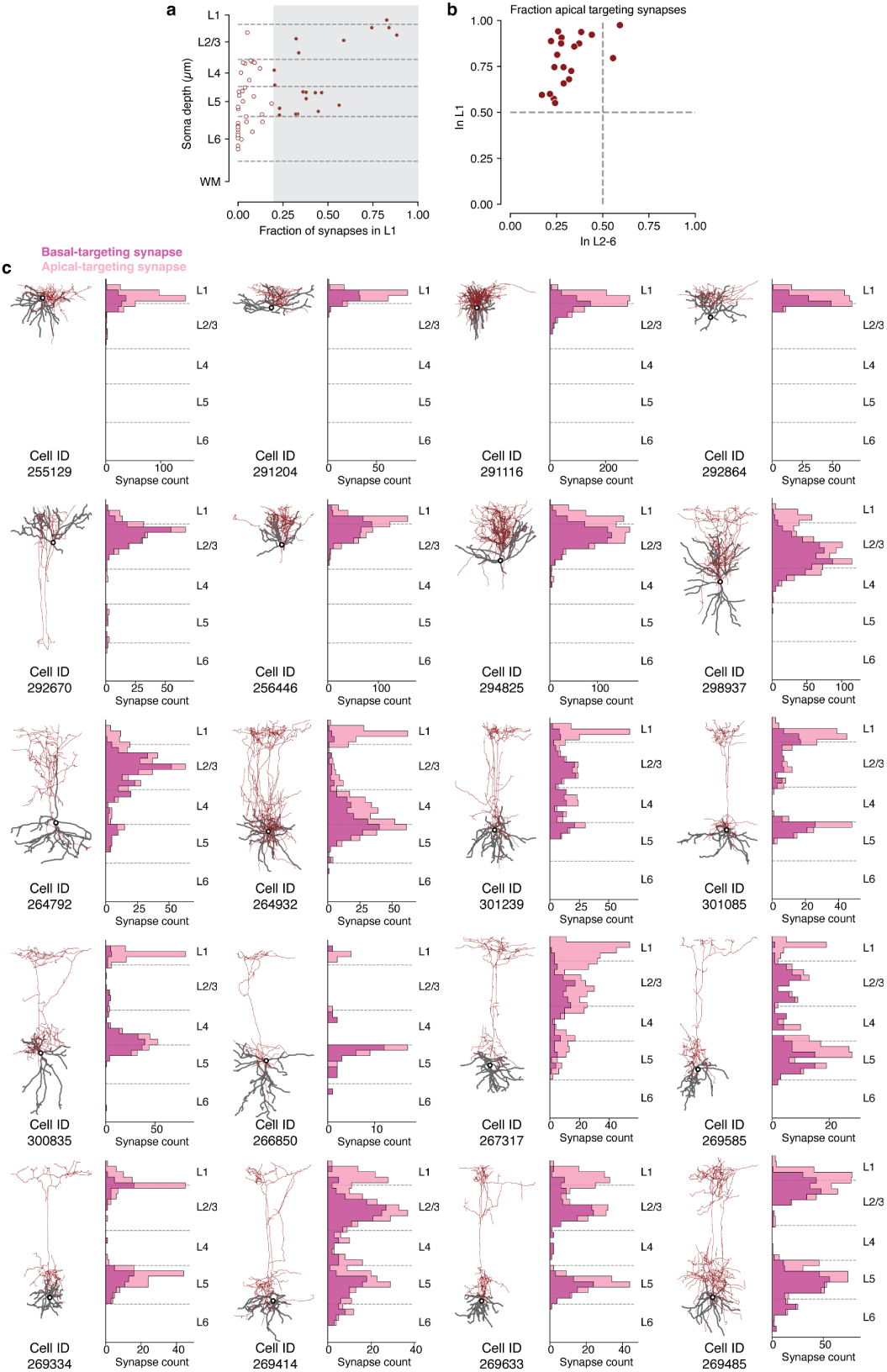
Compartment targeting of Martinotti cells across layers. **a)** DistTC cells have differing fractions of synaptic outputs in layer 1. For the purposes of this figure, we focus on cells with more than 20% of their outputs in layer 1 (gray region, filled circles). **b)** For each DistTC with 20%+ layer 1 output, we measured the fraction of outputs onto apical dendrites among target excitatory cells in the column as a function of synapse depth. The y-axis reflects the apical fraction among synapses in layer 1, while the x-axis reflects the apical fraction among synapses in all deeper layers. For all such cells, the majority of layer 1 targets were onto apical dendrites, while in 18/20 cells the output in other layers was majority of targets were onto basal dendrites. **c)**. Target distributions of individual cells. For each DistTC with 20%+ layer 1 output (morphology at left), we computed the histogram of synapses targeting apical vs basal compartments on excitatory neurons. Note that for most cells with cell bodies in layer 4 and below, there is both a layer 1 arborization that primarily targets apical dendrites and a deeper arborization that largely targets basal dendrites.

**Extended Data Figure 6.**
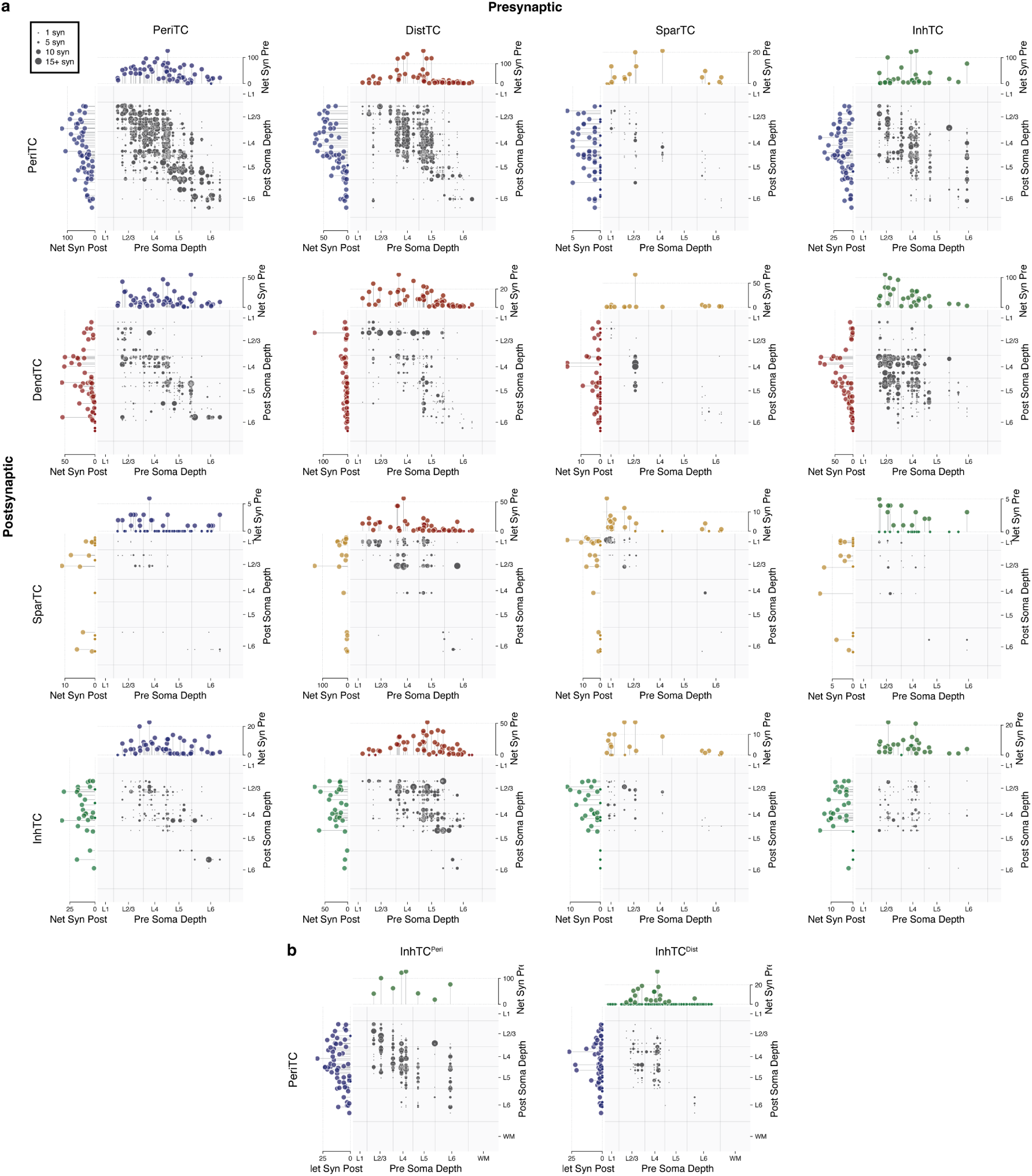
Inhibition of inhibition. **a)** Connectivity dotplot between inhibitory neurons, organized by inhibitory subclasses, organized by soma depth. For each panel, the scatterplot reflects the connectivity from cells in the presynaptic subclass (x-axis) to cells in the postsynaptic subclass (y-axis). Each dot is a single connection, with larger dots having more synapses. The location of each dot corresponds to the depth of the pre- and post-synaptic cell bodies. Stem plots on top and side indicate the net synaptic inputs and net synaptic outputs of each cell in each subclass within the column sample. **b)** Same as **a**, but for InhTC^Peri^ and InhTC^Dist^ onto PeriTCs separately.

**Extended Data Figure 7.**
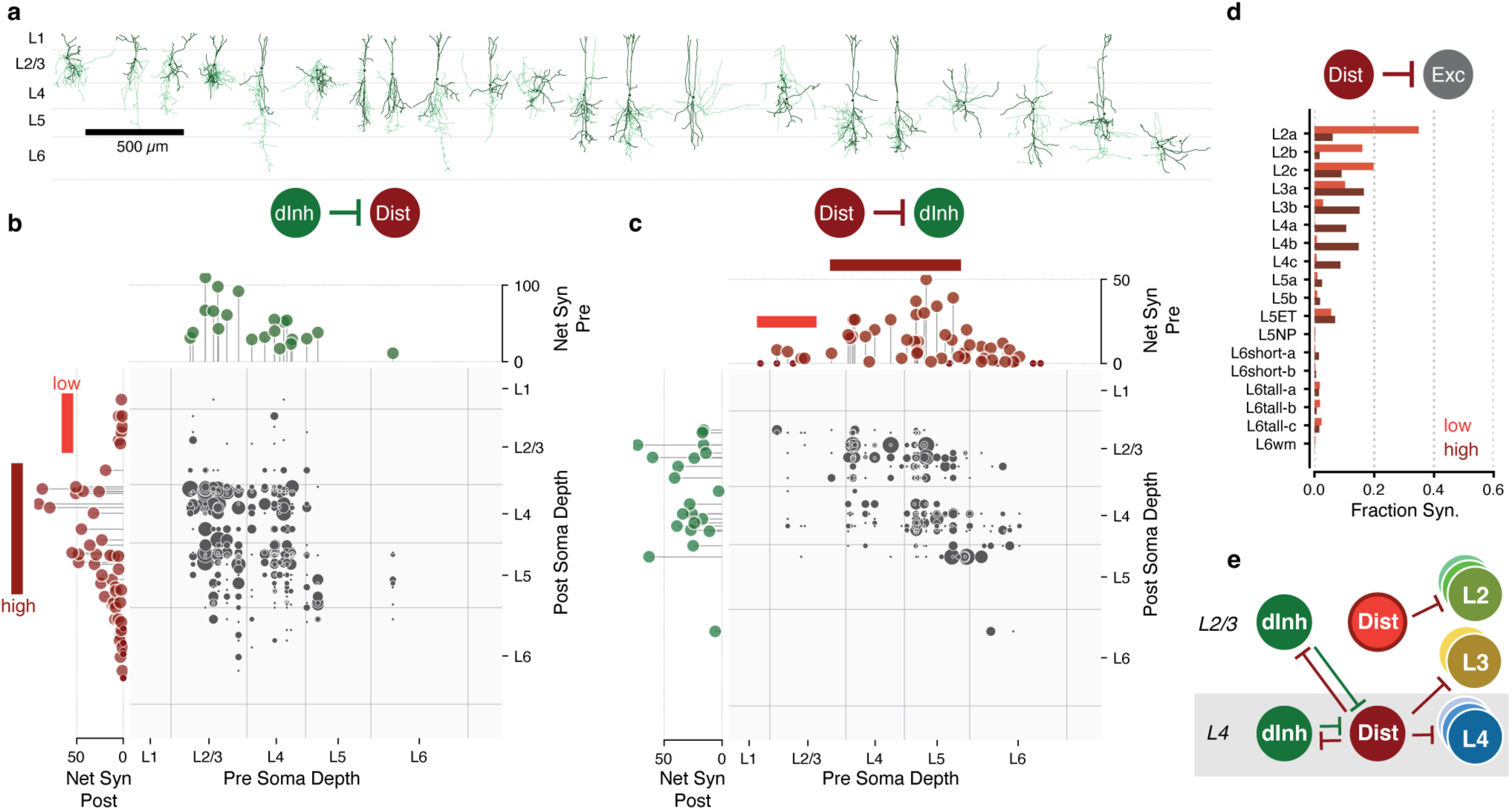
A laminar-specific circuit for InhTC^dist^ cells. **a)** Morphology of all InhTC that preferentially target DistTCs. Cells are sorted by soma depth. **b)** Connectivity dotplot for synapses from InhTC^dist^ onto DistTCs. In the grid, each dot represents a connection from one InhTC onto one DistTC, with the number of synapses indicated by dot size. The location of the dot corresponds to the soma depth of the pre- and post-synaptic cells. Stem plots on top and side indicate the net synaptic inputs and net synaptic outputs of each InhTC^dist^ and DistTC. Note that DistTCs in layer 2/3 receive little input from InhTCs, compared to those in layer 4 and upper layer 5. **c)** Connectivity scatterplot for synapses from DistTCs onto InhTC^dist^, as in **b**. Note that the DistTCs in layer 2/3 also form few synaptic outputs onto InhTC^dist^. **d)** Distribution across M-types of synaptic outputs across low-connection DistTCs and high-connection DistTCs. **e)** Connectivity cartoon suggested by this data.

**Extended Data Figure 8.**
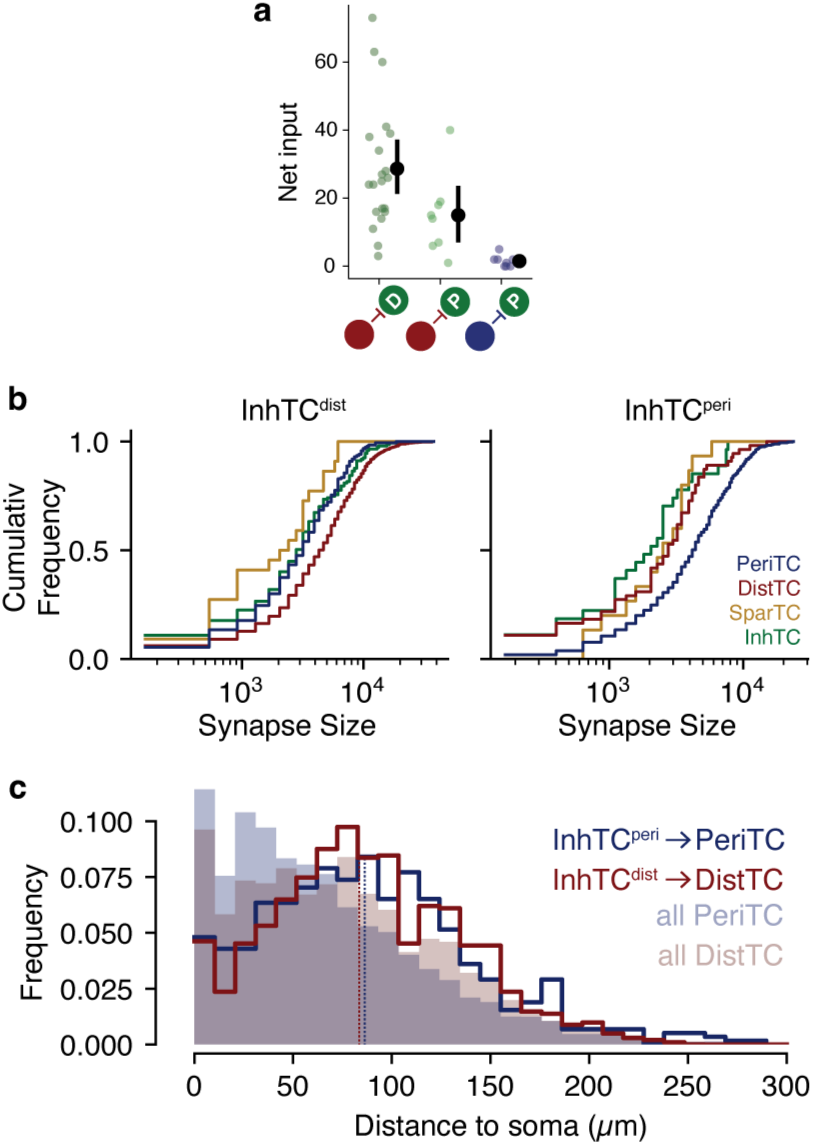
Additional InhTC targeting properties. **a)** Net synapse count onto subtypes of InhTCs. Red indicates DistTCs, blue indicates PeriTCs, and the D and P indicate InhTC^dist^ and InhTC^peri^ respectively. Colored dots are individual cells, black dots are mean, and error bars are bootstrapped 95% confidence interval for the mean. **b)** Cumulative distribution of synapse sizes of InhTC cells onto different target inhibitory subclasses. Left: InhTC^dist^, Right: InhTC^peri^. **c)** Distribution of synapse location from InhTC subtypes onto preferred targets. Lines indicate observed distribution of synapse distances from soma for the preferred targets of InhTC^dist^ and ^peri^. Vertical dashed lines indicate median distances. Filled distributions indicate the synapse distance distribution of all synaptic inputs for the two target Inhibitory subclasses. Synapse locations onto target cells are nearly identical in both average and distribution between the two InhTC subclasses.

**Extended Data Figure 9.**
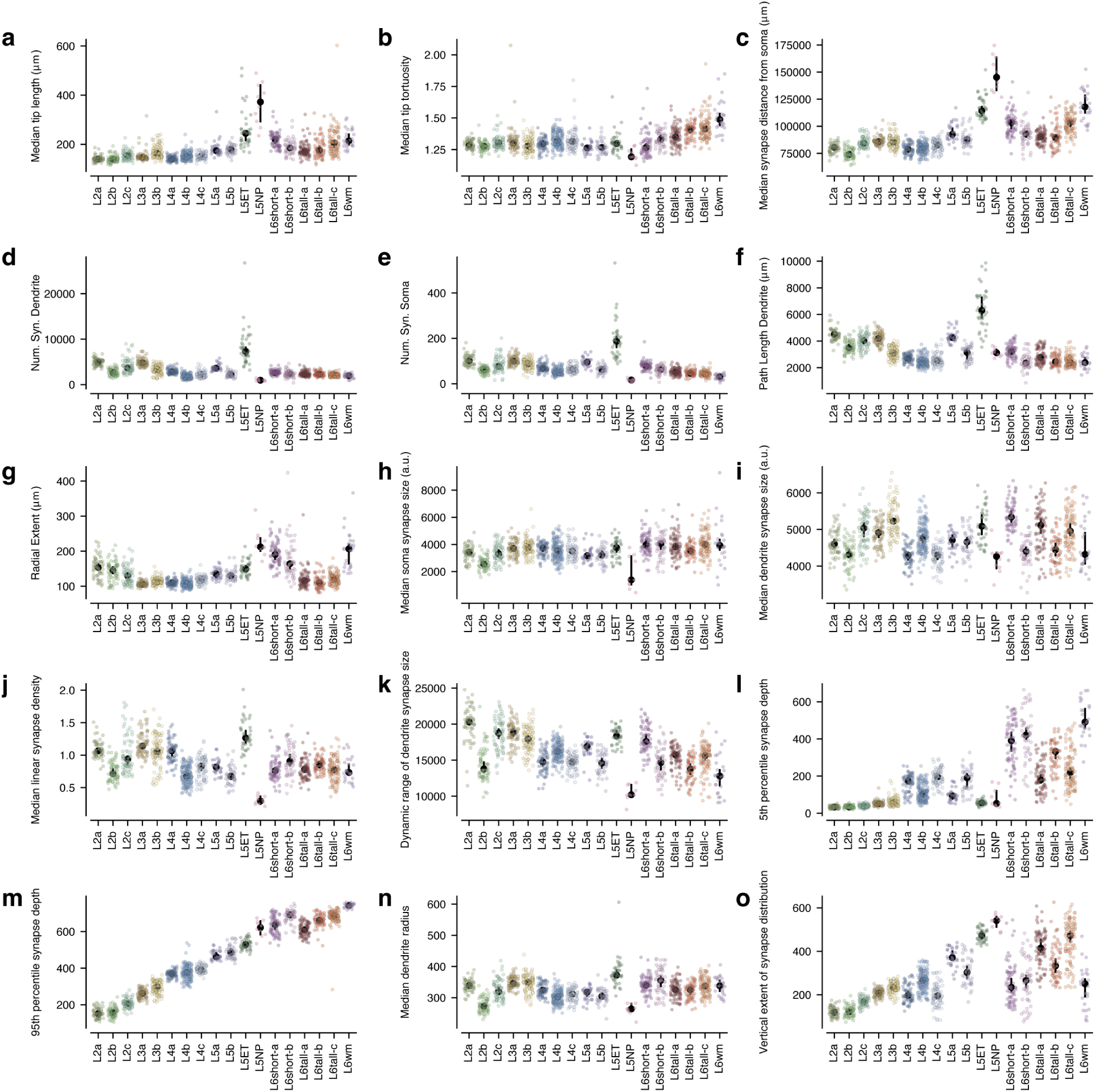
Dendritic features — properties. Features used for classifying M-types, in addition to data-driven components shown in Extended Data Figure 10. See Methods for full definitions of each feature. Colored dots are individual cells, black dots indicate median, and error bars are a bootstrapped 95% confidence interval of the median. **a)** Median dendritic branch tip length. **b)** Median tip tortuosity. **c)** Median input synapse distance from soma. **d)** Number of dendritic synaptic inputs (not including soma). **e)** Number of synaptic inputs on soma. **f)** Net path length of the dendritic arbor. **g)** Radial extent of dendritic arbor, from slanted streamline. **h)** Median size of input synapse onto soma. **i)** Median size of input synapse onto dendrites. **j)** Median linear input synapse density. **k)** Range of synapse sizes, measured as difference between 95th to 5th percentile. **l)** Shallowest extent of synapse distribution, measured as the 5th percentile of input synapse depth. **m)** Deepest extent of synapse distribution, measured as the 95th percentile of synapse depth. **n)** Median dendrite radius. **o)** Height of synapse distribution, measured as the difference between 95th and 5th percentile of synapse depth.

**Extended Data Figure 10.**
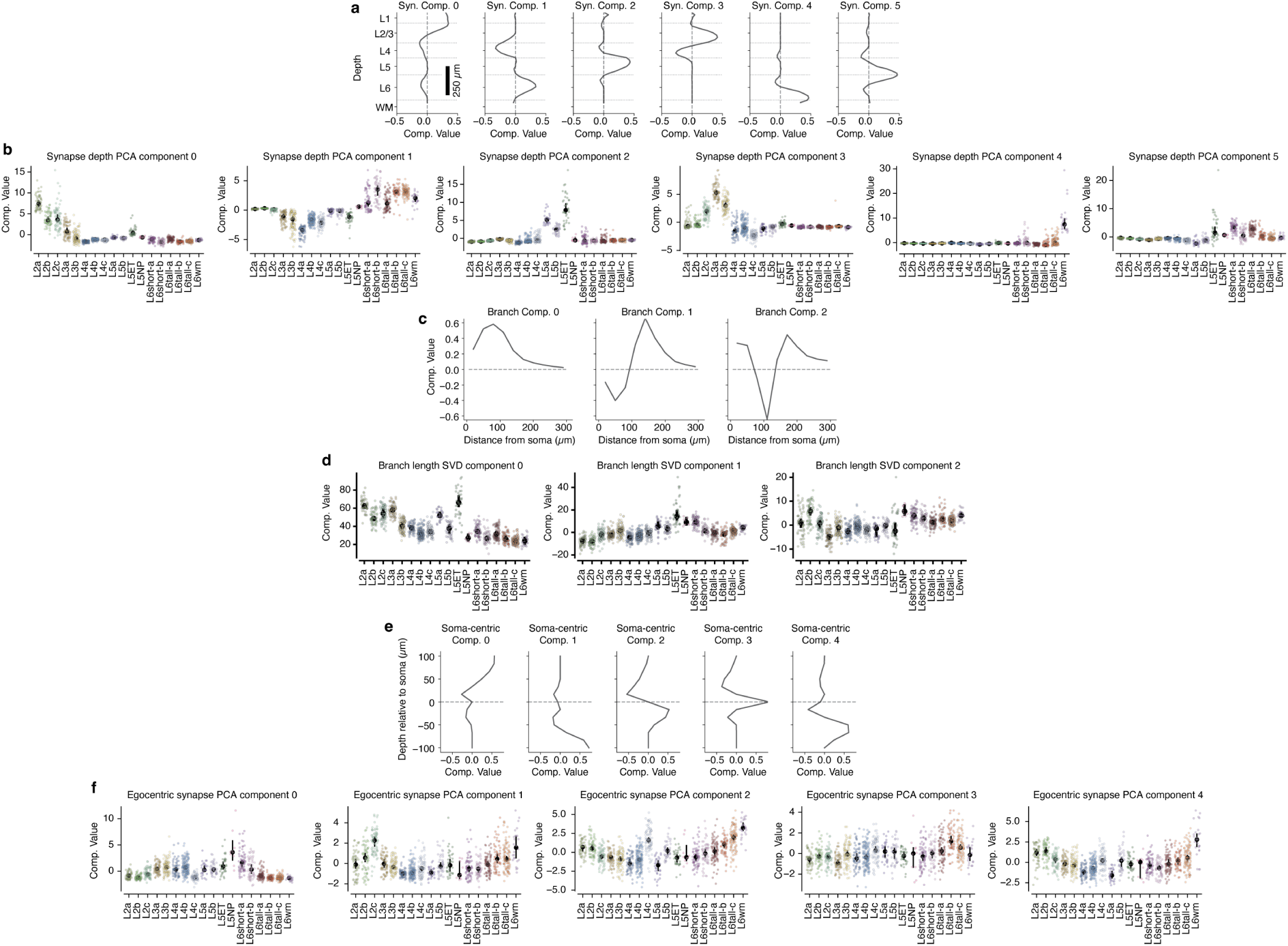
Dendritic features — compartment loadings. Several of the M-type features were based on data-driven components in addition to the basic features shown in Extended Data Figure 9. **a)** SparsePCA components of synapse count across 50 depth bins (approximately 20*µm* per bin) across all excitatory neurons. **b)** Loadings of each resulting M-type cluster onto the components in **a**. Each colored dot is a cell, black dots are median with error bars indicating bootstrapped confidence interval of the median. **c)** Singular value decomposition (SVD) components of the number of distinct branches as a function of distance from the soma in 30 *µm* increments. **d)** Loadings of each resulting M-type cluster onto the branching SVD components in **c**. **e)** SparsePCA components of synapse count as a function of depth relative to the soma location, measured in 13 uniform bins from 100 *µm* below the soma to 100 *µm* above the soma. **f)** Loadings of each resulting M-type cluster onto the soma-centric synapse components in **e**.

**Extended Data Figure 11.**
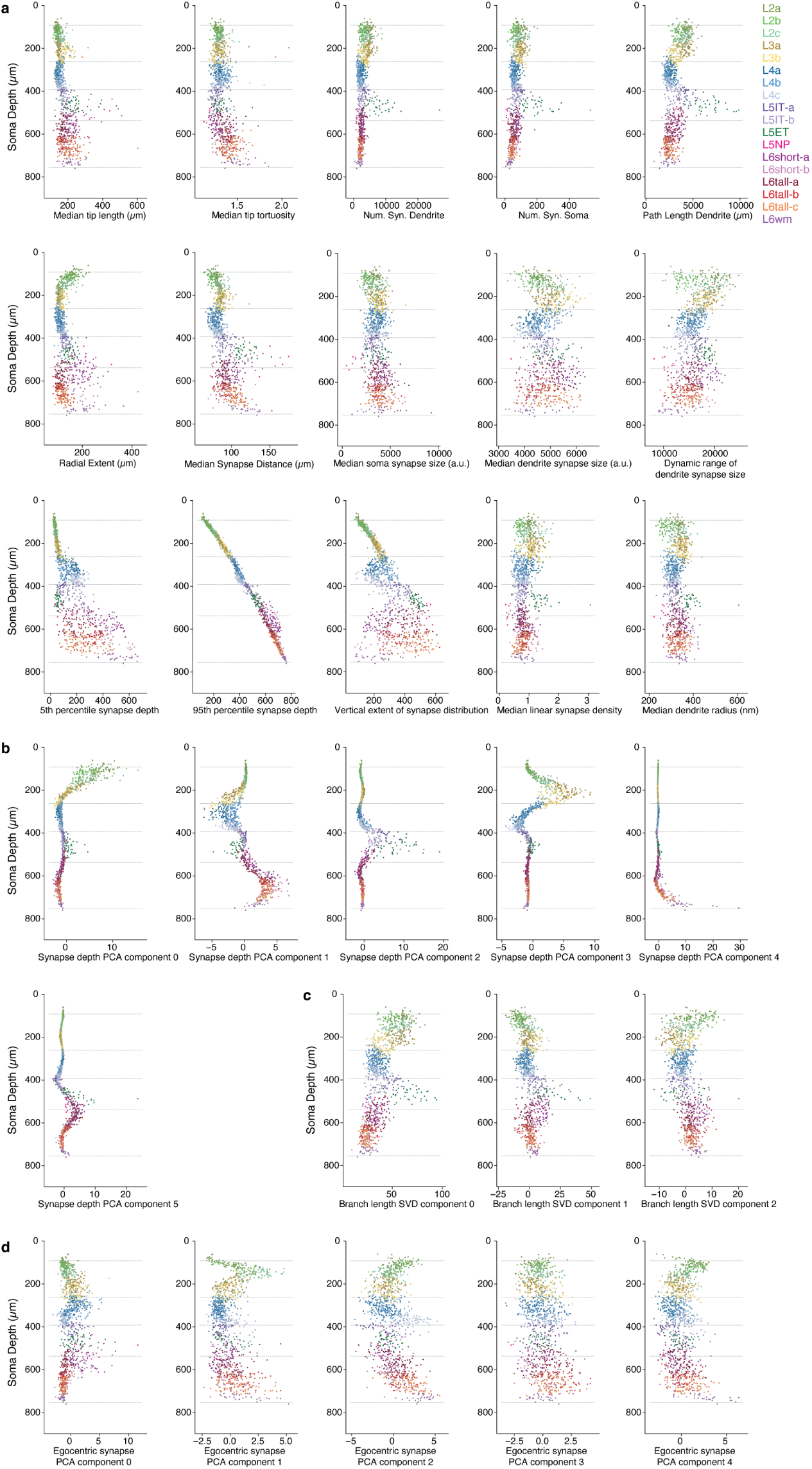
Dendritic properties as a function of depth. For each plot, each dot is an excitatory neuron colored by M-type. The depth of each cell is along the y-axis, while the value of the property is along the x-axis. Approximate layer boundaries are shown with dashed lines. **a)** Individual cell properties. **b)** Synapse depth components. **c)** Branch distribution components. **d)** Soma-centered synapse depth components.

**Extended Data Figure 12.**
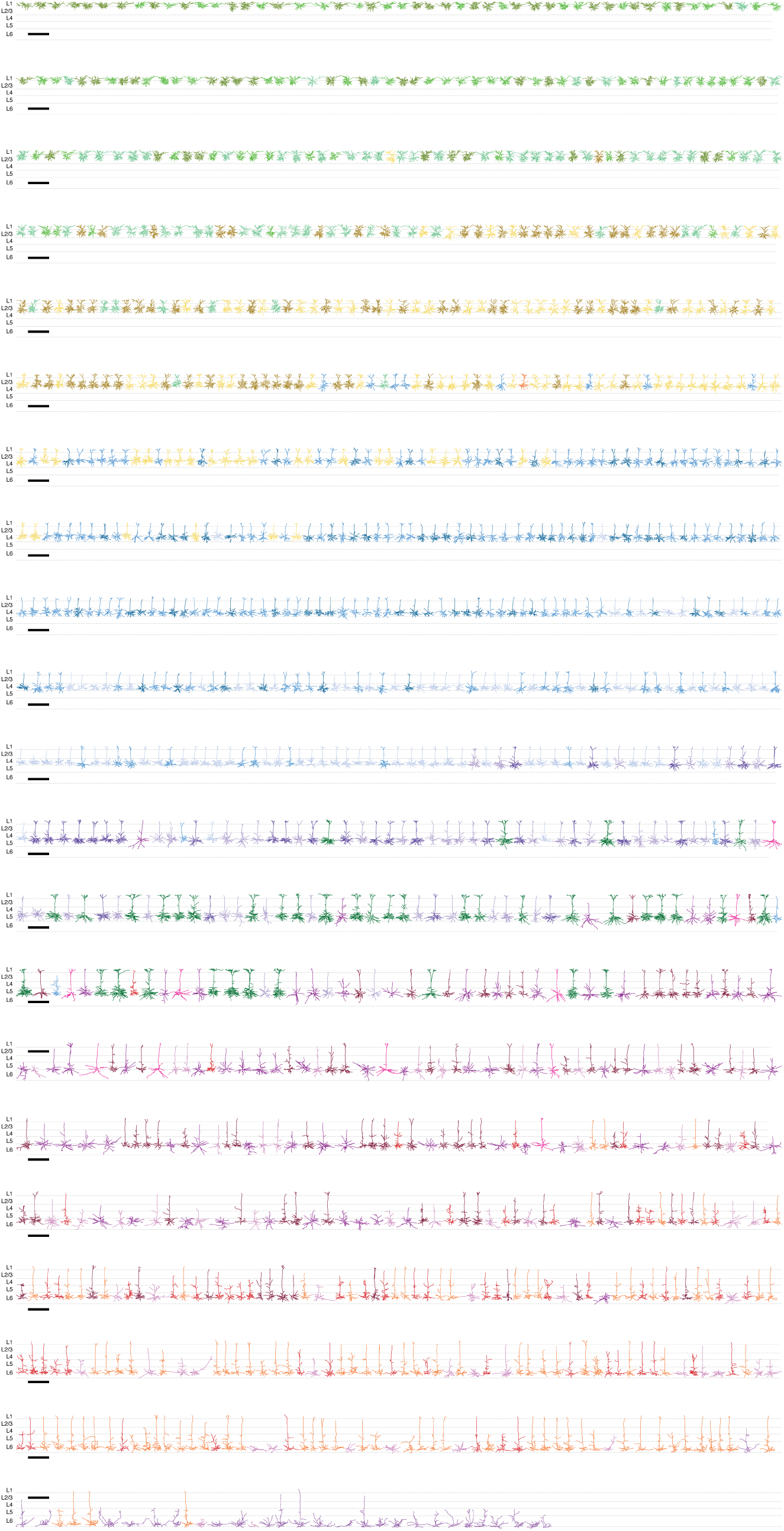
Morphology of all excitatory neurons, sorted by soma depth and colored by anatomical class. Scale bar is 500 *µm*.

**Extended Data Figure 13.**
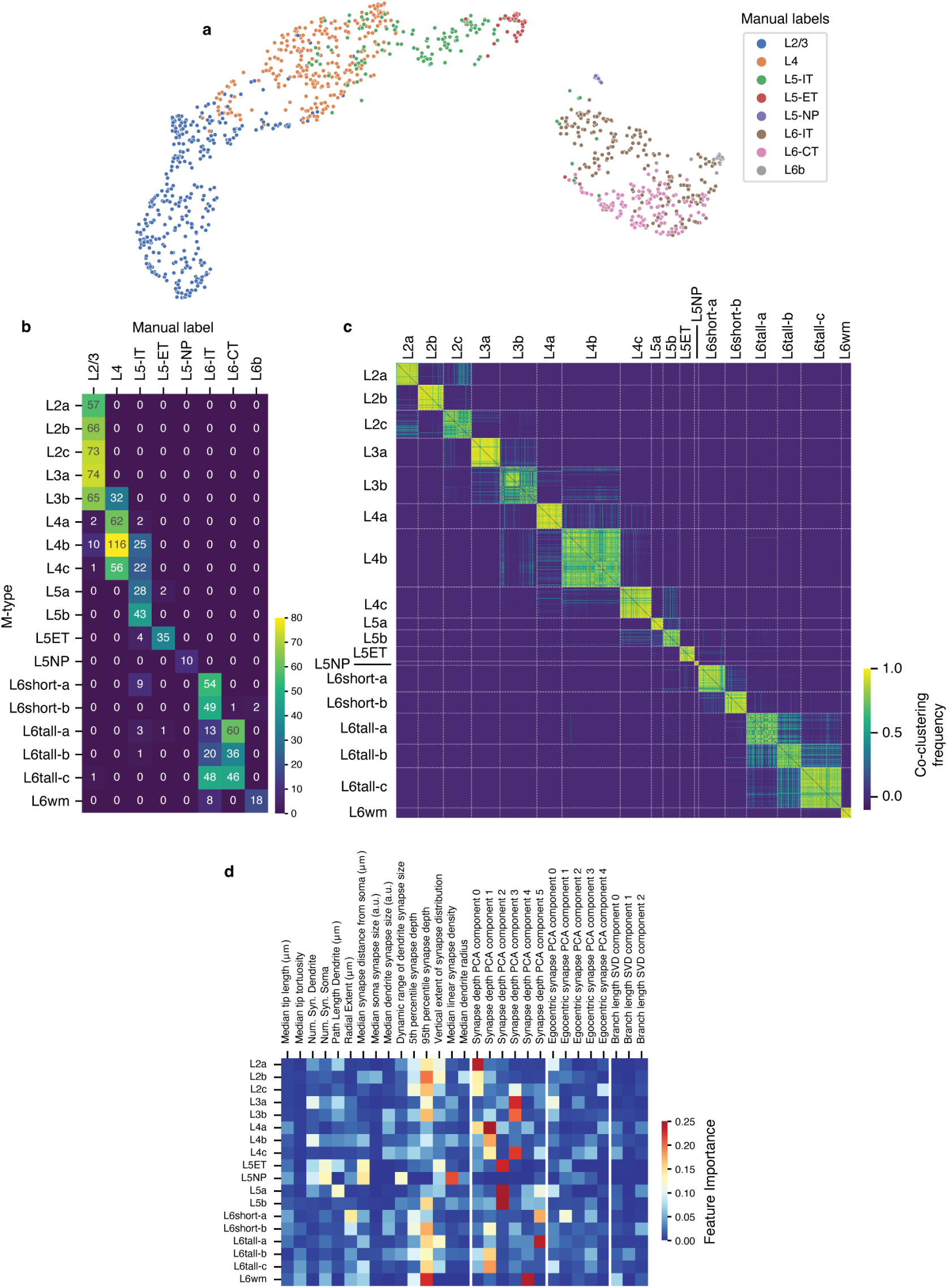
M-type clustering and manual labels. **a)** Matrix of manual labels (x-axis) vs M-types. **b)** UMAP representation of features, colored by manually labeled cell types. **c)** Co-clustering matrix of excitatory cells, indicating ther number of times a pair of cells was clustered together by iterations of phenograph. Cells are ordered by subsequent agglomerative clustering on this matrix. **d)** Feature importance for each M-type, based on training binary random forest classifiers to predict each M-type separately and computing the mean decrease in impurity for each feature.

**Extended Data Figure 14.**
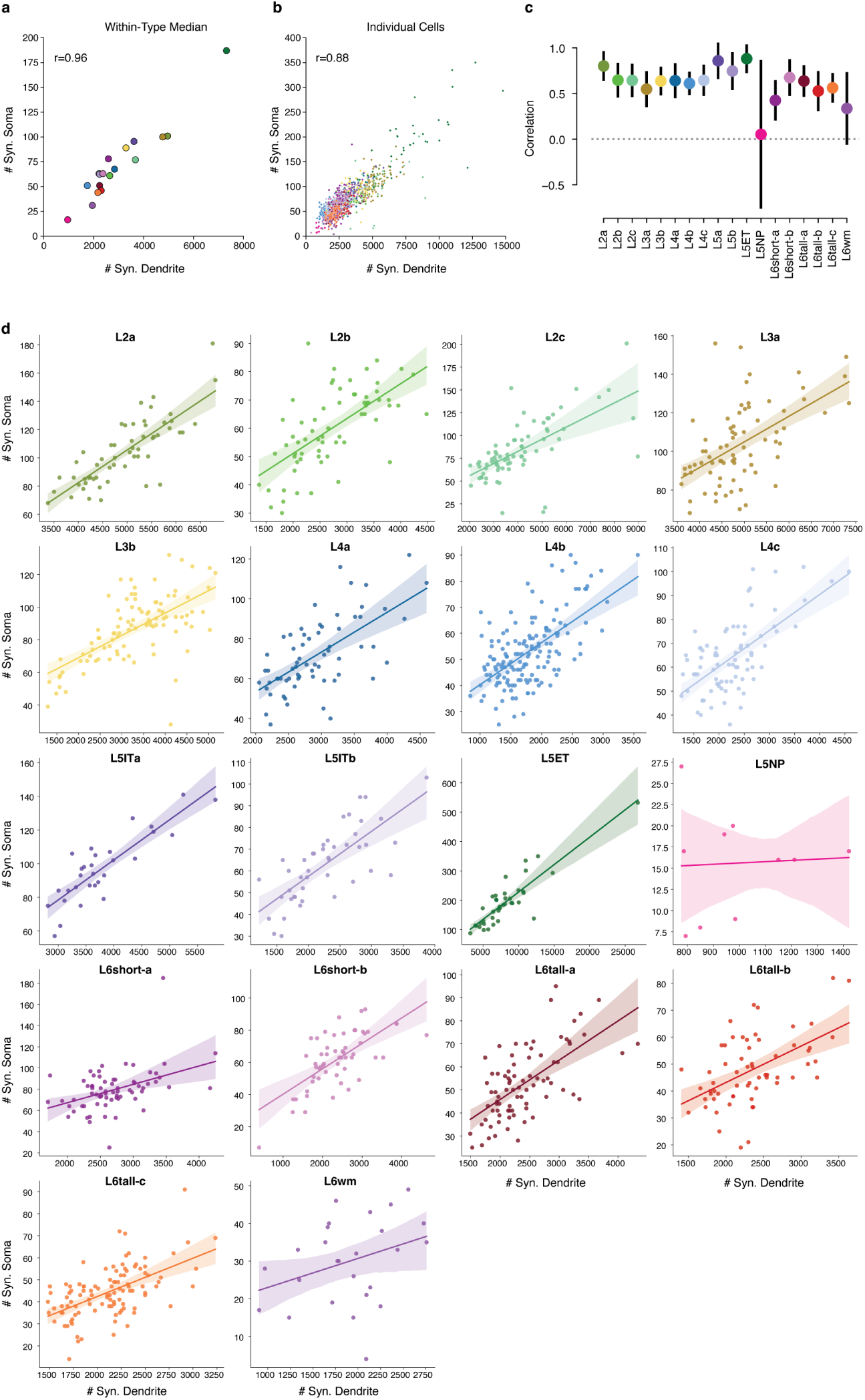
Somatic versus dendritic synapses across all excitatory M-types. a) Median number of dendritic and somatic synapses for excitatory neurons of all M-types. Pearson r=0.96, p=5 *×* 10*^−^*^10^. b) Number of dendritic and somatic input synapses across all excitatory neurons, colored by M-type. Pearson r=0.86, p<1 *×* 10*^−^*^10^. Black line indicates linear fit with 95% confidence intervals from bootstrapping. c) Individual ordinary least square fits (with 95% confidence interval) for each M-type of z-scored dendritic synapses vs z-scored somatic synapses. With the exception of L5NP cells and deep layer 6b L6wm cells, all M-types have a positive relationship between predominantly-inhibitory somatic synapses and mostly excitatory dendritic synapses. d) Number of dendritic vs somatic input synapses for each M-type separately, linear fit line and 95% confidence interval.

**Extended Data Figure 15.**
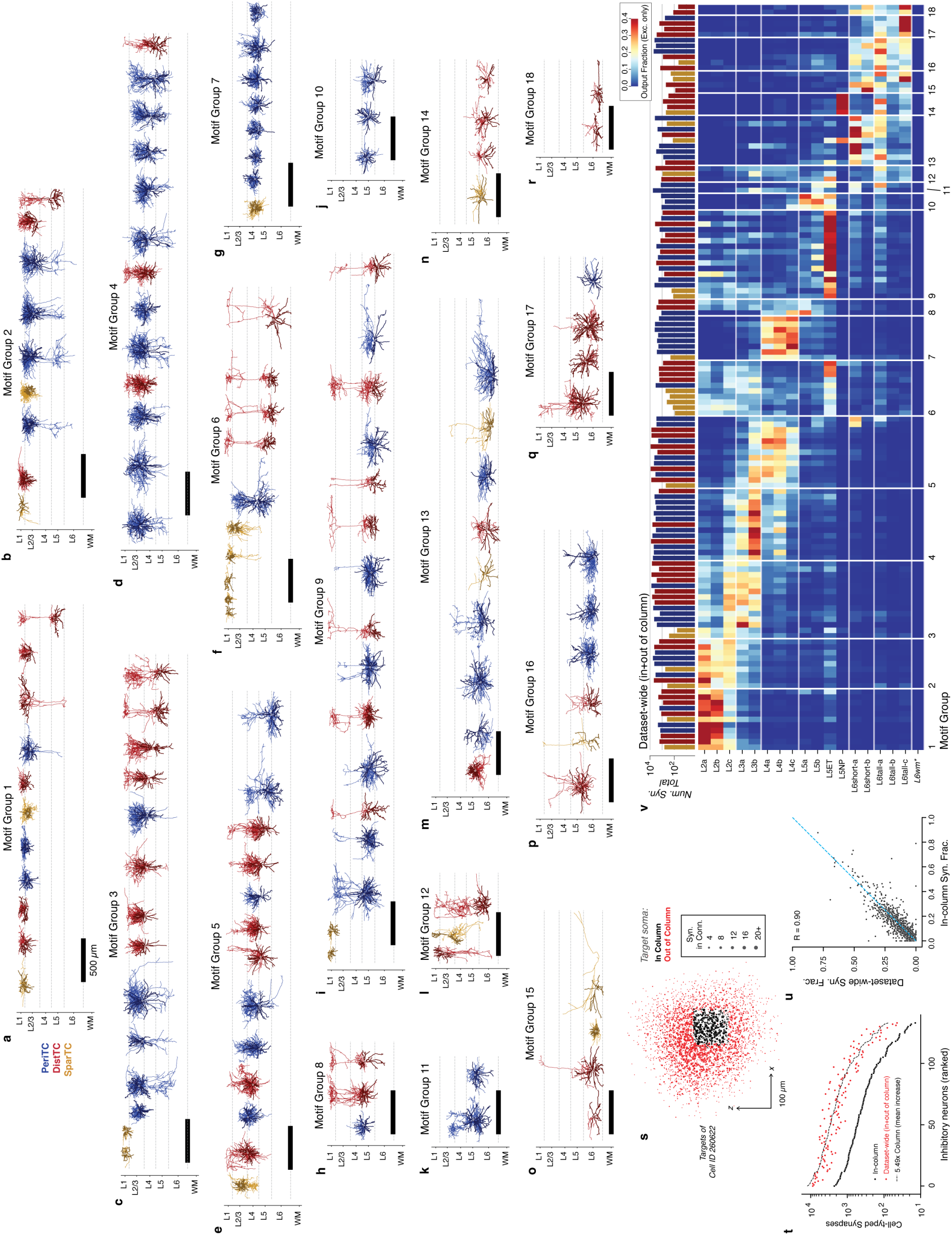
Additional characterization of motif groups. **a–r)** Morphology of all cells, organized by motif group. Within each group, cells are ordered by soma depth. Colors indicate M-type, darker lines indicate dendrites. **s)** The arbors of cells extend well beyond the columnar data. The scatterplot depicts a top-down view of soma locations of all synaptic targets of Cell ID 260622. Black dots are cells within the column, red dots are cells outside the column sample; dot size is proportional to number of synapses. **t)** The number of synapses from each interneuron onto target neurons within the column (black) and anywhere the dataset (red). Interneurons were ordered by within-column synapse count. The mean cell had 5.49 times more synapses across the dataset than onto column targets alone (black dashed line). Only targets passing basic quality control criteria were included. Note that while cells outside the sampled column are not necessarily proofread, synapses onto unproofread dendrites are nearly always correct (see Methods). **u)** Scatterplot of output synapse budget values within-column and dataset-wide (see **v**). The blue line indicates equality. The Pearson correlation between within-column measurements with the dataset-wide measurements was R=0.9, not including trivial zeros (see Methods). **v)** Output synapse budget for for each interneurons onto dataset-wide target M-types, using predictions from perisomatic features from Elabbady et al.^62^. Note that the L6wm M-type was not included in predictions, and is thus trivially zero for all interneurons.

**Extended Data Figure 16.**
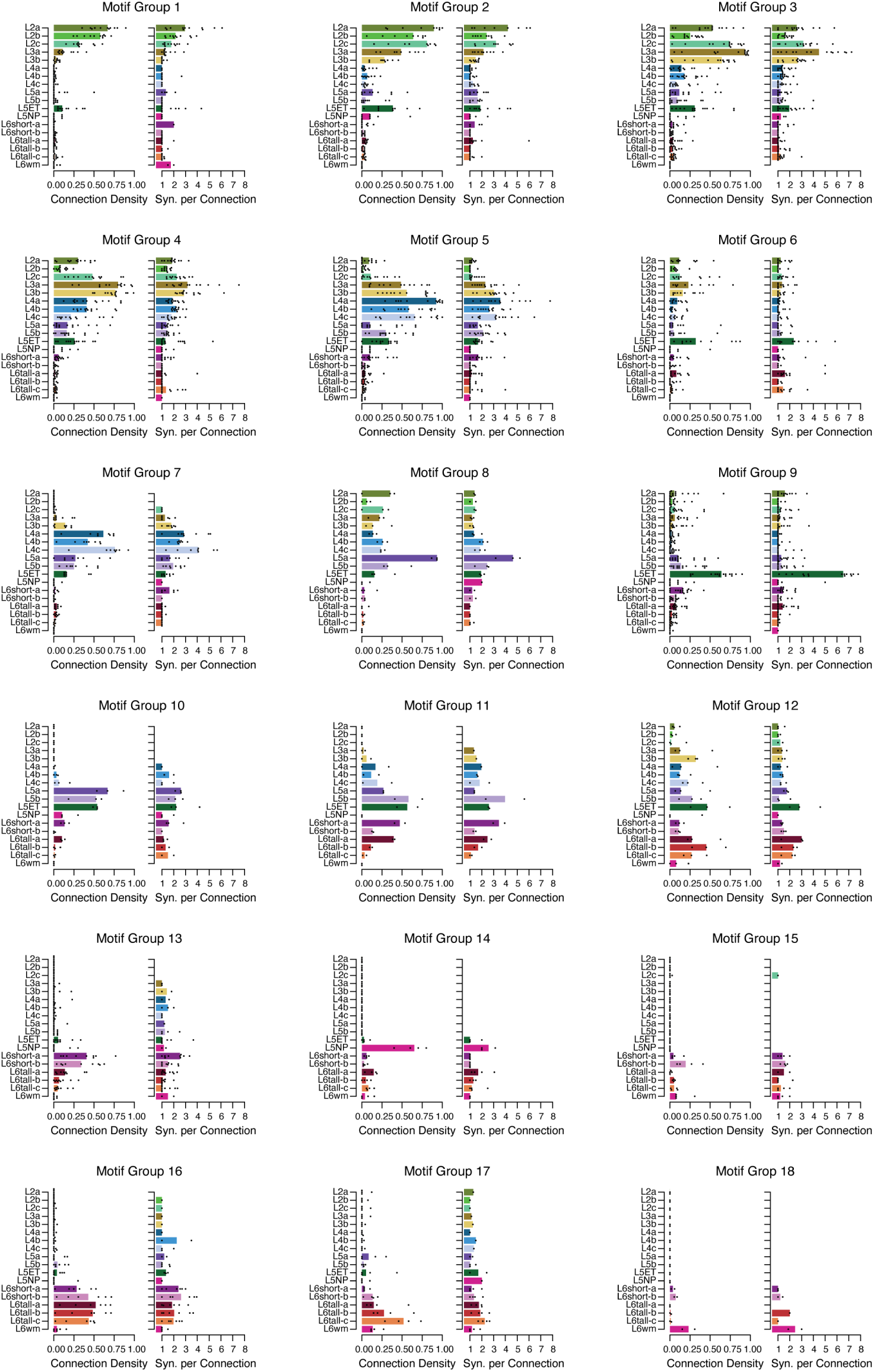
Additional connectivity statistics within motif groups. Connection density (left) measures the fraction of cells for a given M-type within the column targeted with at least 1 synapse. Synapses per connection (right) measures the average number of synapses in each observed connection. Single cell values are represented by dots, median values are shown with bars.

**Extended Data Figure 17.**
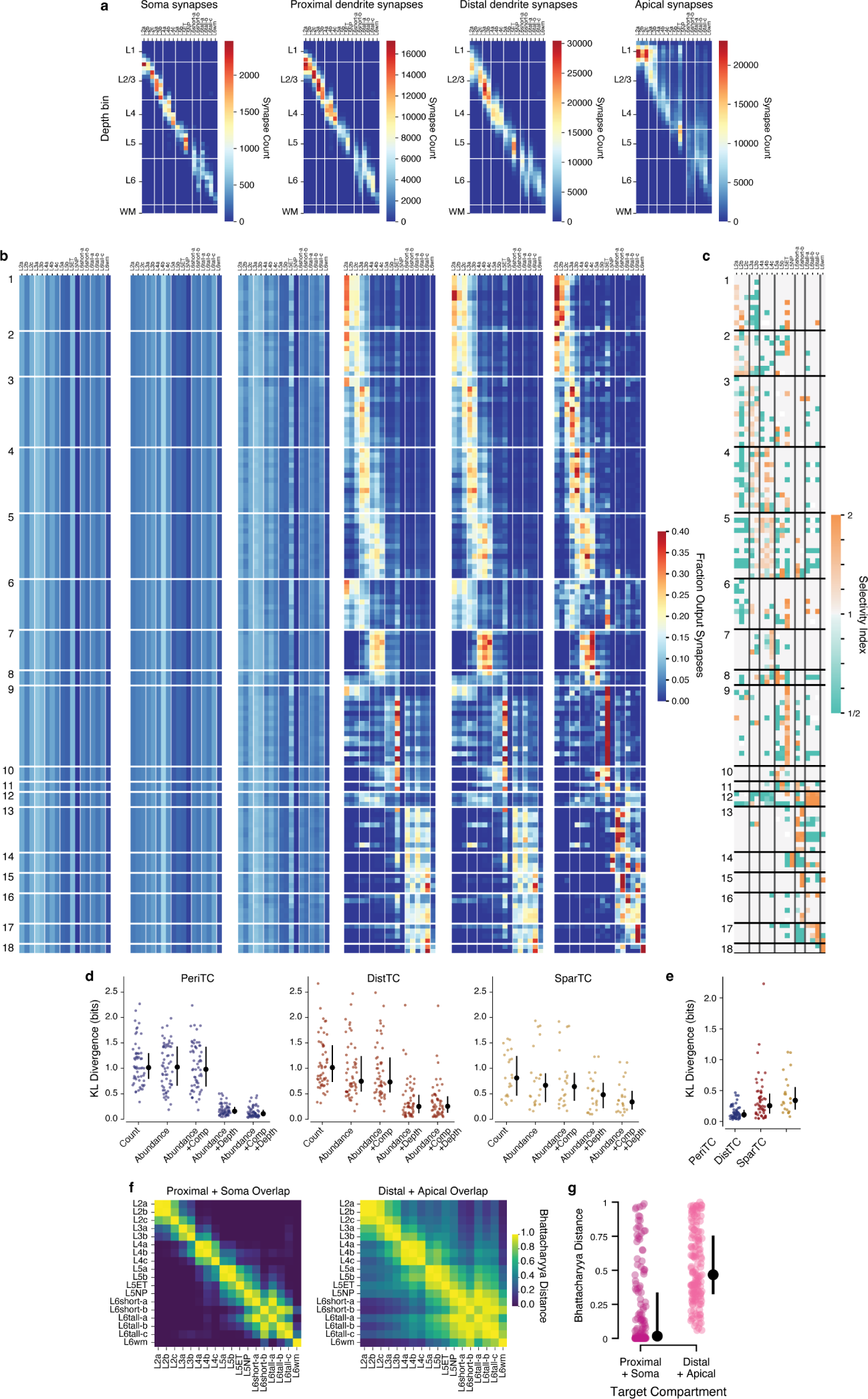
Selectivity and null models for inhibitory connectivity. **a)** Number of synapses per M-type, compartment, and depth bin. These values were used as a the baseline against which to compare synaptic output distributions for each inhibitory neuron. **b)** Expected value of each presynaptic inhibitory neuron according to an increasingly complex set of null models. Each row represents the fraction of synaptic outputs from a given inhibitory neuron (ordered as in Figure 5e), distributed across excitatory M-types. From the left: 1) Synaptic outputs were proportional to the number of cells in a given M-type, regardless of location in space. This approach accounts for the differing cell frequency for each M-type. 2) Synaptic outputs were proportional to the net number of input synapses for a given M-type, regardless of location in space. This approach accounts for the diversity in synaptic inputs for each M-type. 3) Synaptic outputs were distributed across compartments for each inhibitory cell as observed, and distributed across M-types for each compartment separately. This approach accounts for the observed differences in compartment targeting for different interneurons. 4) Synaptic outputs were distributed across M-types within each of 50 depth bins, matching the observed depth distribution of synaptic outputs for each inhibitory neuron. This approach accounts for the spatial distribution of synapses, but not compartment targeting. 5) Synaptic outputs were distributed across M-types within both depth bins and compartments, matching the observed distribution of both. This approach accounts for both the spatial distribution of synapses and compartment targeting, and is the most complete model considered here. At the far right, the observed distribution on the same scale, repeating the data in Figure 5. **c)** Selectivity index (SI) for all cells, as described in the main text. Purple values have the observed number of output synapses significantly higher than a null model with matched compartment and depth targeting, while green are significantly less. Non-significant SI values are treated as 1. **d)** Difference between the observed distribution and the null model distribution for each cell as measured by the Kullback-Leibler divergence (from observed distribution to null distribution), by inhibitory subclass. Each colored dot is a cell, black dots are median with error bars indicating a 95% confidence interval based on a bootstrap. **e)** Comparison between the most complete null model across inhibitory subclasses. The PeriTCs have the lowest KL divergence of all types, indicating that the null model best predicts their connectivity. Note also that the individual cells exhibit a range of specificity relative to null models. **f)** Similarity of M-type synapse distributions in space, using the Bhattacharyya distance between the depth distribution of synaptic inputs onto soma and proximal dendrites (left) and distal and apical dendrite (right). Values closer to 1 indicate more similar distributions, values closer to 0 indicate more distinct distributions. **g)** All Bhattacharyya distance comparisons in **e**, with colored dots indicating pairs of distinct M-types, black dots indicating the median, and error bars showing a bootstrapped 95% confidence interval. Across all pairs, synaptic inputs onto the perisomatic and somatic compartments are more spatially segregated across different M-types than synaptic inputs onto distal and apical dendrites (*p* = 3.0 *×* 10*^−^*^19^, Mann-Whitney U test).

## References

1. Consortium, M. et al. Functional connectomics spanning multiple areas of mouse visual cortex. bioRxiv, 2021.07.28.454025 (July 29, 2021).

2. Lorente de Nò, R. in *Physiology of the Nervous System* 288–312 (Oxford University Press, 1949).

3. Mountcastle, V. Modality and topographic properties of single neurons of cat’s somatic sensory cortex. J Neurophysiol 20, 408–434 (July 1957).

4. Hubel, D. & Wiesel, T. Receptive fields, binocular interaction and functional architecture in the cat’s visual cortex. J Physiol 160, 106–154 (1962).

5. Da Costa, N. M. & Martin, K. A. C. Whose Cortical Column Would that Be? Frontiers in neuroanatomy 4, 16 (2010).

6. Zeng, H. What is a cell type and how to define it? Cell 185, 2739–2755 (July 2022).

7. Ramon y Cajal, S. *Histology of the Nervous System of Man and Vertebrates* (A. Maloine, Paris, 1909).

8. Jones, E. G. Varieties and distribution of non-pyramidal cells in the somatic sensory cortex of the squirrel monkey. Journal of Comparative Neurology 160, 205–267 (1975).

9. Gilbert, C. D. & Wiesel, T. N. Morphology and intracortical projections of functionally characterised neurones in the cat visual cortex. Nature 280, 120–125 (July 1979).

10. Martin, K. A. & Whitteridge, D. Form, function and intracortical projections of spiny neurones in the striate visual cortex of the cat. The Journal of Physiology 353, 463–504 (1984).

11. Wang, Y. et al. Anatomical, physiological and molecular properties of Martinotti cells in the somatosensory cortex of the juvenile rat. The Journal of Physiology 561, 65–90 (Nov. 1, 2004).

12. Ascoli, G. A. et al. Petilla terminology: nomenclature of features of GABAergic interneurons of the cerebral cortex. Nature Reviews Neuroscience 9, 557–568 (July 2008).

13. Tasic, B. et al. Shared and distinct transcriptomic cell types across neocortical areas. Nature 563, 72–78 (Nov. 2018).

14. Gouwens, N. W. et al. Classification of electrophysiological and morphological neuron types in the mouse visual cortex. Nature Neuroscience 22, 1182–1195 (July 2019).

15. Kanari, L. et al. Objective Morphological Classification of Neocortical Pyramidal Cells. Cerebral Cortex 29, 1719–1735 (Apr. 1, 2019).

16. De Vries, S. E. J. et al. A large-scale standardized physiological survey reveals functional organization of the mouse visual cortex. Nature Neuroscience 23, 138–151 (Jan. 2020).

17. Scala, F. et al. Phenotypic variation of transcriptomic cell types in mouse motor cortex. Nature 598, 144–150 (Oct. 7, 2021).

18. Bugeon, S. et al. A transcriptomic axis predicts state modulation of cortical interneurons. Nature 607, 330–338 (July 2022).

19. Meyer, H. S. et al. Inhibitory interneurons in a cortical column form hot zones of inhibition in layers 2 and 5A. Proceedings of the National Academy of Sciences of the United States of America 108, 16807–16812 (Oct. 4, 2011).

20. Tasic, B. et al. Adult mouse cortical cell taxonomy revealed by single cell transcriptomics. Nature Neuroscience 19, 335–346 (Feb. 2016).

21. Tremblay, R., Lee, S. & Rudy, B. GABAergic Interneurons in the Neocortex: From Cellular Properties to Circuits. Neuron 91, 260–292 (July 2016).

22. Yao, Z. et al. A taxonomy of transcriptomic cell types across the isocortex and hippocampal formation. Cell 184, 3222– 3241.e26 (June 2021).

23. Machold, R. et al. Id2 GABAergic interneurons comprise a neglected fourth major group of cortical inhibitory cells. eLife 12, e85893 (Sept. 4, 2023).

24. Somogyi, P., Tamás, G., Lujan, R. & Buhl, E. H. Salient features of synaptic organisation in the cerebral cortex. Brain Research Reviews 26, 113–135 (May 1998).

25. Pfeffer, C. K., Xue, M., He, M., Huang, Z. J. & Scanziani, M. Inhibition of inhibition in visual cortex: the logic of connections between molecularly distinct interneurons. Nature Neuroscience 16, 1068–1076 (Aug. 2013).

26. Dudok, B. et al. Alternating sources of perisomatic inhibition during behavior. Neuron 109, 997–1012.e9 (Mar. 2021).

27. Pi, H.-J. et al. Cortical interneurons that specialize in disinhibitory control. Nature 503, 521–524 (Nov. 2013).

28. Kubota, Y. Untangling GABAergic wiring in the cortical microcircuit. Current Opinion in Neurobiology 26, 7–14 (June 1, 2014).

29. Peters, A., Meinecke, D. L. & Karamanlidis, A. N. Vasoactive intestinal polypeptide immunoreactive neurons in the primary visual cortex of the cat. en. J. Neurocytol. 16, 23–38 (Feb. 1987).

30. Peters, A. The axon terminals of vasoactive intestinal polypeptide (VIP)-containing bipolar cells in rat visual cortex. en. J. Neurocytol. 19, 672–685 (Oct. 1990).

31. Gouwens, N. W. et al. Integrated Morphoelectric and Transcriptomic Classification of Cortical GABAergic Cells. Cell 183, 935–953.e19 (Nov. 2020).

32. Fino, E. & Yuste, R. Dense Inhibitory Connectivity in Neocortex. Neuron 69, 1188–1203 (Mar. 2011).

33. Karnani, M. M., Agetsuma, M. & Yuste, R. A blanket of inhibition: functional inferences from dense inhibitory connectivity. Current Opinion in Neurobiology 26, 96–102 (June 2014).

34. Muñoz, W., Tremblay, R., Levenstein, D. & Rudy, B. Layer-specific modulation of neocortical dendritic inhibition during active wakefulness. Science 355, 954–959 (Mar. 3, 2017).

35. Hilscher, M. M., Leão, R. N., Edwards, S. J., Leão, K. E. & Kullander, K. Chrna2-Martinotti Cells Synchronize Layer 5 Type A Pyramidal Cells via Rebound Excitation. PLOS Biology 15, e2001392 (Feb. 9, 2017).

36. Lu, J. et al. Selective inhibitory control of pyramidal neuron ensembles and cortical subnetworks by chandelier cells. Nature Neuroscience 20, 1377–1383 (Oct. 2017).

37. Wu, S. J. et al. Cortical somatostatin interneuron subtypes form cell-type-specific circuits. Neuron 111, 2675–2692.e9 (Sept. 6, 2023).

38. Jiang, X. et al. Principles of connectivity among morphologically defined cell types in adult neocortex. Science 350, aac9462–aac9462 (Nov. 27, 2015).

39. Campagnola, L. et al. Local connectivity and synaptic dynamics in mouse and human neocortex. Science 375, eabj5861 (Mar. 11, 2022).

40. Yetman, M. J. et al. Intersectional monosynaptic tracing for dissecting subtype-specific organization of GABAergic interneuron inputs. Nature Neuroscience, 1 (Jan. 28, 2019).

41. White, J. G., Southgate, E., Thomson, J. N. & Brenner, S. The structure of the nervous system of the nematode Caenorhabditis elegans. Philosophical Transactions of the Royal Society of London. Series B, Biological Sciences 314, 1–340 (Nov. 12, 1986).

42. Zheng, Z. et al. A Complete Electron Microscopy Volume of the Brain of Adult Drosophila melanogaster. Cell 174, 730–743.e22 (July 2018).

43. Scheffer, L. K. et al. A connectome and analysis of the adult Drosophila central brain. eLife 9 (eds Marder, E., Eisen, M. B., Pipkin, J. & Doe, C. Q.) e57443 (Sept. 3, 2020).

44. Schneider-Mizell, C. M. et al. Structure and function of axo-axonic inhibition. eLife 10 (eds Calabrese, R. L., Callaway, E., Huang, Z. J. & Oberlaender, M.) e73783 (Dec. 1, 2021).

45. Shapson-Coe, A. et al. A connectomic study of a petascale fragment of human cerebral cortex. bioRxiv, 2021.05.29.446289 (May 30, 2021).

46. Kasthuri, N. et al. Saturated Reconstruction of a Volume of Neocortex. Cell 162, 648–661 (July 30, 2015).

47. Lee, W.-C. A. et al. Anatomy and function of an excitatory network in the visual cortex. Nature 532, 370–374 (Apr. 21, 2016).

48. Motta, A. et al. Dense connectomic reconstruction in layer 4 of the somatosensory cortex. Science 366, eaay3134 (Nov. 29, 2019).

49. Yin, W. et al. A petascale automated imaging pipeline for mapping neuronal circuits with high-throughput transmission electron microscopy. Nature Communications 11, 4949 (Dec. 2020).

50. Macrina, T., et al. Petascale neural circuit reconstruction: automated methods. bioRxiv, 2021.08.04.455162 (2021).

51. Mahalingam, G. et al. A scalable and modular automated pipeline for stitching of large electron microscopy datasets. eLife 11 (eds Cardona, A., Calabrese, R. L., Cardona, A., Arganda-Carreras, I. & Tischer, C.) e76534 (July 26, 2022).

52. Glickfeld, L. L. & Olsen, S. R. Higher-Order Areas of the Mouse Visual Cortex. Annual Review of Vision Science 3, 251–273 (2017).

53. Dorkenwald, S. et al. Binary and analog variation of synapses between cortical pyramidal neurons. eLife 11, e76120 (Nov. 16, 2022).

54. Dorkenwald, S. et al. FlyWire: online community for whole-brain connectomics. Nature Methods 19, 119–128 (Jan. 2022).

55. Dorkenwald, S., et al. CAVE: Connectome Annotation Versioning Engine. bioRxiv, 2023.07.26.550598 (July 28, 2023).

56. Keller, D., Erö, C. & Markram, H. Cell Densities in the Mouse Brain: A Systematic Review. Frontiers in Neuroanatomy 12, 83 (2018).

57. Gamlin, C. Integrating EM and Patch-seq data: Synaptic connectivity and target specificity of predicted Sst transcriptomic types. *In prep*. (2023).

58. Somogyi, P., Kisvárday, Z., Martin, K. & Whitteridge, D. Synaptic connections of morphologically identified and physiologically characterized large basket cells in the striate cortex of cat. Neuroscience 10, 261–294 (1983).

59. Kubota, Y., Karube, F., Nomura, M. & Kawaguchi, Y. The Diversity of Cortical Inhibitory Synapses. Frontiers in Neural Circuits 10 (2016).

60. Freund, T. F. & Katona, I. Perisomatic Inhibition. Neuron 56, 33–42 (Oct. 4, 2007).

61. Somogyi, P., Freund, T. & Cowey, A. The axo-axonic interneuron in the cerebral cortex of the rat, cat and monkey. Neuroscience 7, 2577–2607 (Nov. 1982).

62. Elabbady, L. et al. Quantitative Census of Local Somatic Features in Mouse Visual Cortex. bioRiv, 2022.07.20.499976 (July 22, 2022).

63. Nigro, M. J., Hashikawa-Yamasaki, Y. & Rudy, B. Diversity and Connectivity of Layer 5 Somatostatin-Expressing Interneurons in the Mouse Barrel Cortex. The Journal of Neuroscience 38, 1622–1633 (Feb. 14, 2018).

64. Bodor, A. L., et al. The Synaptic Architecture of Layer 5 Thick Tufted Excitatory Neurons in the Visual Cortex of Mice preprint (Neuroscience, Oct. 20, 2023).

65. Lee, S., Kruglikov, I., Huang, Z. J., Fishell, G. & Rudy, B. A disinhibitory circuit mediates motor integration in the somatosensory cortex. Nature Neuroscience 16, 1662–1670 (Nov. 2013).

66. Kepecs, A. & Fishell, G. Interneuron cell types are fit to function. Nature 505, 318–326 (Jan. 2014).

67. Kullander, K. & Topolnik, L. Cortical disinhibitory circuits: cell types, connectivity and function. Trends in Neurosciences 44, 643–657 (Aug. 2021).

68. Dávid, C., Schleicher, A., Zuschratter, W. & Staiger, J. F. The innervation of parvalbumin-containing interneurons by VIP-immunopositive interneurons in the primary somatosensory cortex of the adult rat. European Journal of Neuroscience 25, 2329–2340 (2007).

69. Fu, Y. et al. A Cortical Circuit for Gain Control by Behavioral State. Cell 156, 1139–1152 (Mar. 13, 2014).

70. Millman, D. J. et al. VIP interneurons in mouse primary visual cortex selectively enhance responses to weak but specific stimuli. eLife 9, e55130 (Oct. 27, 2020).

71. Packer, A. M. & Yuste, R. Dense, Unspecific Connectivity of Neocortical Parvalbumin-Positive Interneurons: A Canonical Microcircuit for Inhibition? Journal of Neuroscience 31, 13260–13271 (Sept. 14, 2011).

72. Naka, A. et al. Complementary networks of cortical somatostatin interneurons enforce layer specific control. eLife 8 (eds Svoboda, K., Colgin, L., Yu, J. & Petersen, C. C.) e43696 (Mar. 18, 2019).

73. McGarry, L. M. & Carter, A. G. Inhibitory Gating of Basolateral Amygdala Inputs to the Prefrontal Cortex. The Journal of Neuroscience 36, 9391–9406 (Sept. 7, 2016).

74. Narayanan, R. T. et al. Beyond Columnar Organization: Cell Type- and Target Layer-Specific Principles of Horizontal Axon Projection Patterns in Rat Vibrissal Cortex. Cerebral Cortex 25, 4450–4468 (Nov. 2015).

75. Levine, J. H. et al. Data-Driven Phenotypic Dissection of AML Reveals Progenitor-like Cells that Correlate with Prognosis. Cell 162, 184–197 (July 2, 2015).

76. Kim, E. J., Juavinett, A. L., Kyubwa, E. M., Jacobs, M. W. & Callaway, E. M. Three Types of Cortical Layer 5 Neurons That Differ in Brain-wide Connectivity and Function. Neuron 88, 1253–1267 (Dec. 16, 2015).

77. Stepanyants, A., Hof, P. R. & Chklovskii, D. B. Geometry and Structural Plasticity of Synaptic Connectivity. Neuron 34, 275–288 (Apr. 11, 2002).

78. Udvary, D. et al. The impact of neuron morphology on cortical network architecture. Cell Reports 39, 110677 (Apr. 12, 2022).

79. Binzegger, T., Douglas, R. J. & Martin, K. A. C. A quantitative map of the circuit of cat primary visual cortex. The Journal of Neuroscience: The Official Journal of the Society for Neuroscience 24, 8441–8453 (Sept. 29, 2004).

80. Braitenberg, V. & Schüz, A. *Cortex: Statistics and Geometry of Neuronal Connectivity* 2nd (Springer-Verlag, Berlin, 1998).

81. Xue, M., Atallah, B. V. & Scanziani, M. Equalizing excitation–inhibition ratios across visual cortical neurons. Nature 511, 596–600 (July 2014).

82. Iascone, D. M. et al. Whole-Neuron Synaptic Mapping Reveals Spatially Precise Excitatory/Inhibitory Balance Limiting Dendritic and Somatic Spiking. Neuron 106, 566–578.e8 (May 2020).

83. Hodge, R. D. et al. Conserved cell types with divergent features in human versus mouse cortex. Nature 573, 61–68 (Sept. 5, 2019).

84. Naka, A. & Adesnik, H. Inhibitory Circuits in Cortical Layer 5. Frontiers in Neural Circuits 10 (2016).

85. Ye, Z. et al. Instructing Perisomatic Inhibition by Direct Lineage Reprogramming of Neocortical Projection Neurons. Neuron 88, 475–483 (Nov. 4, 2015).

86. Lodato, S. et al. Excitatory Projection Neuron Subtypes Control the Distribution of Local Inhibitory Interneurons in the Cerebral Cortex. Neuron 69, 763–779 (Feb. 2011).

87. Wester, J. C. et al. Neocortical Projection Neurons Instruct Inhibitory Interneuron Circuit Development in a Lineage-Dependent Manner. Neuron 102, 960–975.e6 (June 2019).

88. Hennequin, G., Agnes, E. J. & Vogels, T. P. Inhibitory Plasticity: Balance, Control, and Codependence. Annual Review of Neuroscience 40, 557–579 (July 25, 2017).

89. Kim, E. J. et al. Extraction of Distinct Neuronal Cell Types from within a Genetically Continuous Population. Neuron (May 2020).

90. Condylis, C. et al. Dense functional and molecular readout of a circuit hub in sensory cortex. Science 375, eabl5981 (Jan. 7, 2022).

91. O’Toole, S. M., Oyibo, H. K. & Keller, G. B. Molecularly targetable cell types in mouse visual cortex have distinguishable prediction error responses. Neuron 111, 2918–2928.e8 (Sept. 2023).

92. Zhuang, J. et al. Laminar distribution and arbor density of two functional classes of thalamic inputs to primary visual cortex. Cell Reports 37, 109826 (Oct. 12, 2021).

93. Harris, K. D. & Shepherd, G. M. G. The neocortical circuit: themes and variations. Nature Neuroscience 18, 170–181 (Feb. 2015).

94. Karimi, A., Odenthal, J., Drawitsch, F., Boergens, K. M. & Helmstaedter, M. Cell-type specific innervation of cortical pyramidal cells at their apical dendrites. eLife 9 (ed Mason, C. A.) e46876 (Feb. 28, 2020).

95. Anastasiades, P. G. & Carter, A. G. Circuit organization of the rodent medial prefrontal cortex. Trends in Neurosciences 44, 550–563 (July 1, 2021).

96. Weis, M. A. et al. Large-scale unsupervised discovery of excitatory morphological cell types in mouse visual cortex. bioRxiv, 2022.12.22.521541 (Dec. 22, 2022).

97. Lund, J. S. Anatomical Organization of Macaque Monkey Striate Visual Cortex. Annual Review of Neuroscience 11, 253–288 (Mar. 1988).

98. Leventhal, A. Evidence that the different classes of relay cells of the cat’s lateral geniculate nucleus terminate in different layers of the striate cortex. Experimental Brain Research 37 (Oct. 1979).

99. Hua, Y. et al. Connectomic analysis of thalamus-driven disinhibition in cortical layer 4. Cell Reports 41, 111476 (Oct. 11, 2022).

100. Yu, J., Hu, H., Agmon, A. & Svoboda, K. Recruitment of GABAergic Interneurons in the Barrel Cortex during Active Tactile Behavior. Neuron 104, 412–427.e4 (Oct. 23, 2019).

101. Xu, H., Jeong, H.-Y., Tremblay, R. & Rudy, B. Neocortical Somatostatin-Expressing GABAergic Interneurons Disinhibit the Thalamorecipient Layer 4. Neuron 77, 155–167 (Jan. 2013).

102. Chittajallu, R., Pelkey, K. A. & McBain, C. J. Neurogliaform cells dynamically regulate somatosensory integration via synapse-specific modulation. Nature Neuroscience 16, 13–15 (Jan. 2013).

103. Huang, Z. J., Di Cristo, G. & Ango, F. Development of GABA innervation in the cerebral and cerebellar cortices. Nature Reviews Neuroscience 8, 673–686 (Sept. 2007).

104. Bargmann, C. I. & Marder, E. From the connectome to brain function. Nature Methods 10, 483–490 (June 2013).

105. Schmidt, H. et al. Axonal synapse sorting in medial entorhinal cortex. Nature 549, 469–475 (Sept. 2017).

106. Oberlaender, M. et al. Cell Type–Specific Three-Dimensional Structure of Thalamocortical Circuits in a Column of Rat Vibrissal Cortex. Cerebral Cortex 22, 2375–2391 (Oct. 2012).

107. Zhang, M. et al. Spatially resolved cell atlas of the mouse primary motor cortex by MERFISH. Nature 598, 137–143 (Oct. 7, 2021).

108. Cheng, S. et al. Vision-dependent specification of cell types and function in the developing cortex. Cell 185, 311–327.e24 (Jan. 2022).

109. Bortone, D. S., Olsen, S. R. & Scanziani, M. Translaminar Inhibitory Cells Recruited by Layer 6 Corticothalamic Neurons Suppress Visual Cortex. Neuron 82, 474–485 (Apr. 2014).

110. Frandolig, J. E. et al. The Synaptic Organization of Layer 6 Circuits Reveals Inhibition as a Major Output of a Neocortical Sublamina. Cell Reports 28, 3131–3143.e5 (Sept. 2019).

111. Peters, A. & Feldman, M. L. The projection of the lateral geniculate nucleus to area 17 of the rat cerebral cortex. I. General description. J. Neurocytol. 5, 63–84 (Feb. 1976).

112. White, E. L. & Keller, A. Intrinsic circuitry involving the local axon collaterals of corticothalamic projection cells in mouse SmI cortex. J. Comp. Neurol. 262, 13–26 (Aug. 1987).

113. Egger, R., Dercksen, V. J., Udvary, D., Hege, H.-C. & Oberlaender, M. Generation of dense statistical connectomes from sparse morphological data. Frontiers in Neuroanatomy 8, 129 (2014).

114. Da Costa, N. M. & Martin, K. A. C. Selective targeting of the dendrites of corticothalamic cells by thalamic afferents in area 17 of the cat. J. Neurosci. 29, 13919–13928 (Nov. 2009).

115. Gour, A. et al. Postnatal connectomic development of inhibition in mouse barrel cortex. Science 371, eabb4534 (Jan. 29, 2021).

116. Bates, A. S., Janssens, J., Jefferis, G. S. & Aerts, S. Neuronal cell types in the fly: single-cell anatomy meets single-cell genomics. Current Opinion in Neurobiology 56, 125–134 (June 2019).

117. Dorkenwald, S. et al. Multi-Layered Maps of Neuropil with Segmentation-Guided Contrastive Learning. bioRxiv, 2022.03.29.486320 (Mar. 30, 2022).

118. Celii, B. et al. NEURD: automated proofreading and feature extraction for connectomics. bioRxiv, 2023.03.14.532674 (Mar. 15, 2023).

## References

119. Hua, Y., Laserstein, P. & Helmstaedter, M. Large-volume en-bloc staining for electron microscopy-based connectomics. Nature Communications 6, 7923 (Aug. 3, 2015).

120. Phelps, J. S. et al. Reconstruction of motor control circuits in adult Drosophila using automated transmission electron microscopy. en. Cell 184, 759–774.e18 (Feb. 2021).

121. Wetzel, A. W., et al. Registering large volume serial-section electron microscopy image sets for neural circuit recon-struction using FFT signal whitening in 2016 IEEE Applied Imagery Pattern Recognition Workshop (AIPR) (Oct. 2016), 1–10.

122. Mitchell, E., Keselj, S., Popovych, S., Buniatyan, D. & Sebastian Seung, H. Siamese Encoding and Alignment by Multiscale Learning with Self-Supervision (Apr. 2019).

123. Lee, K., Zung, J., Li, P., Jain, V. & Sebastian Seung, H. Superhuman Accuracy on the SNEMI3D Connectomics Challenge (May 2017).

124. Wu, J., Silversmith, W. M., Lee, K. & Sebastian Seung, H. Chunkflow: hybrid cloud processing of large 3D images by convolutional nets 2021.

125. Zlateski, A. & Seung, H. S. Image Segmentation by Size-Dependent Single Linkage Clustering of a Watershed Basin Graph 2015.

126. Lu, R., Zlateski, A. & Sebastian Seung, H. Large-scale image segmentation based on distributed clustering algorithms (June 2021).

127. Turner, N. et al. Synaptic Partner Assignment Using Attentional Voxel Association Networks. arXiv:1904.09947 [cs] (Nov. 21, 2019).

128. Maitin-Shepard, J., et al. *google/neuroglancer:* version v2.23. Oct. 16, 2021.

129. Sato, M., Bitter, I., Bender, M., Kaufman, A. & Nakajima, M. *TEASAR: tree-structure extraction algorithm for accurate and robust skeletons* in *Proceedings the Eighth Pacific Conference on Computer Graphics and Applications* Proceedings the Eighth Pacific Conference on Computer Graphics and Applications (Oct. 2000), 281–449.

130. Schneider-Mizell, C. M. et al. Quantitative neuroanatomy for connectomics in Drosophila. eLife 5 (ed Calabrese, R. L.) e12059 (Mar. 18, 2016).

131. Pedregosa, F. et al. Scikit-learn: Machine Learning in Python. Journal of Machine Learning Research 12, 2825–2830 (2011).

132. Silversmith, W. et al. seung-lab/cloud-volume: Zenodo Release *v1* version 5.3.2. Nov. 10, 2021.

133. Dorkenwald, S., Schneider-Mizell, C. & Collman, F. sdorkenw/MeshParty: v1.9.0 version v1.9.0. Mar. 13, 2020.

134. Hunter, J. D. Matplotlib: A 2D Graphics Environment. Computing in Science & Engineering 9, 90–95 (2007).

135. Harris, C. R. et al. Array programming with NumPy. Nature 585, 357–362 (Sept. 17, 2020).

136. Team, T. P. D. pandas-dev/pandas: Pandas version v1.5.2. Nov. 22, 2022.

137. Virtanen, P. et al. SciPy 1.0: fundamental algorithms for scientific computing in Python. Nature Methods 17, 261–272 (Mar. 2, 2020).

138. Seabold, S. & Perktold, J. statsmodels: Econometric and statistical modeling with python in 9th Python in Science Conference (2010).

139. Schroeder, W., Martin, K. & Lorensen, B. *The visualization toolkit* 4. ed. In collab. with Kitware, I. 512 pp. (Kitware, Inc, Clifton Park, NY, 2006).

