## Extended Data Cell Atlas for "Cell-type-specific inhibitory circuitry from a connectomic census of mouse visual cortex"

### Motif Group 1

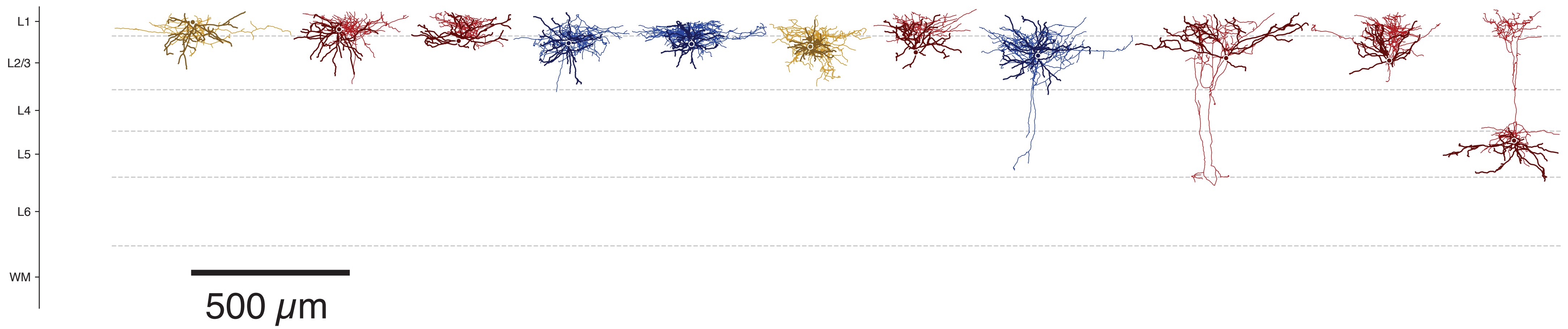

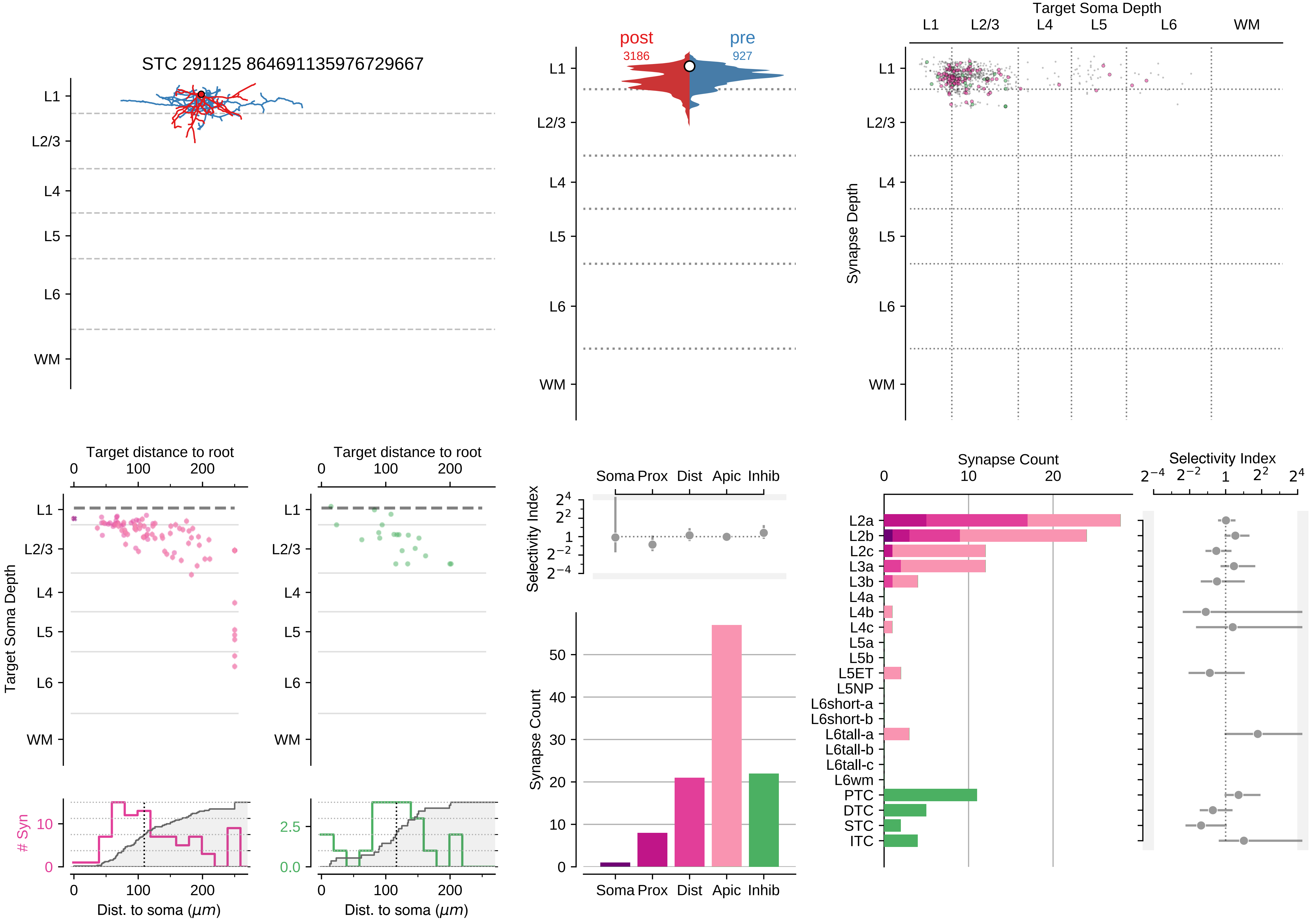

STC 291125 864691135976729667
L1
L2/3
L4
L5
L6
WM
post
3186
pre
927
Target Soma Depth
Synapse Depth
Target distance to root
Target Soma Depth
### Syn
Dist. to soma (µm)
Selectivity Index
Soma Prox Dist Apic Inhib
Synapse Count
Synapse Count
Selectivity Index
L2a
L2b
L2c
L3a
L3b
L4a
L4b
L4c
L5a
L5b
L5ET
L5NP
L6short-a
L6short-b
L6tall-a
L6tall-b
L6tall-c
L6wm
PTC
DTC
STC
ITC

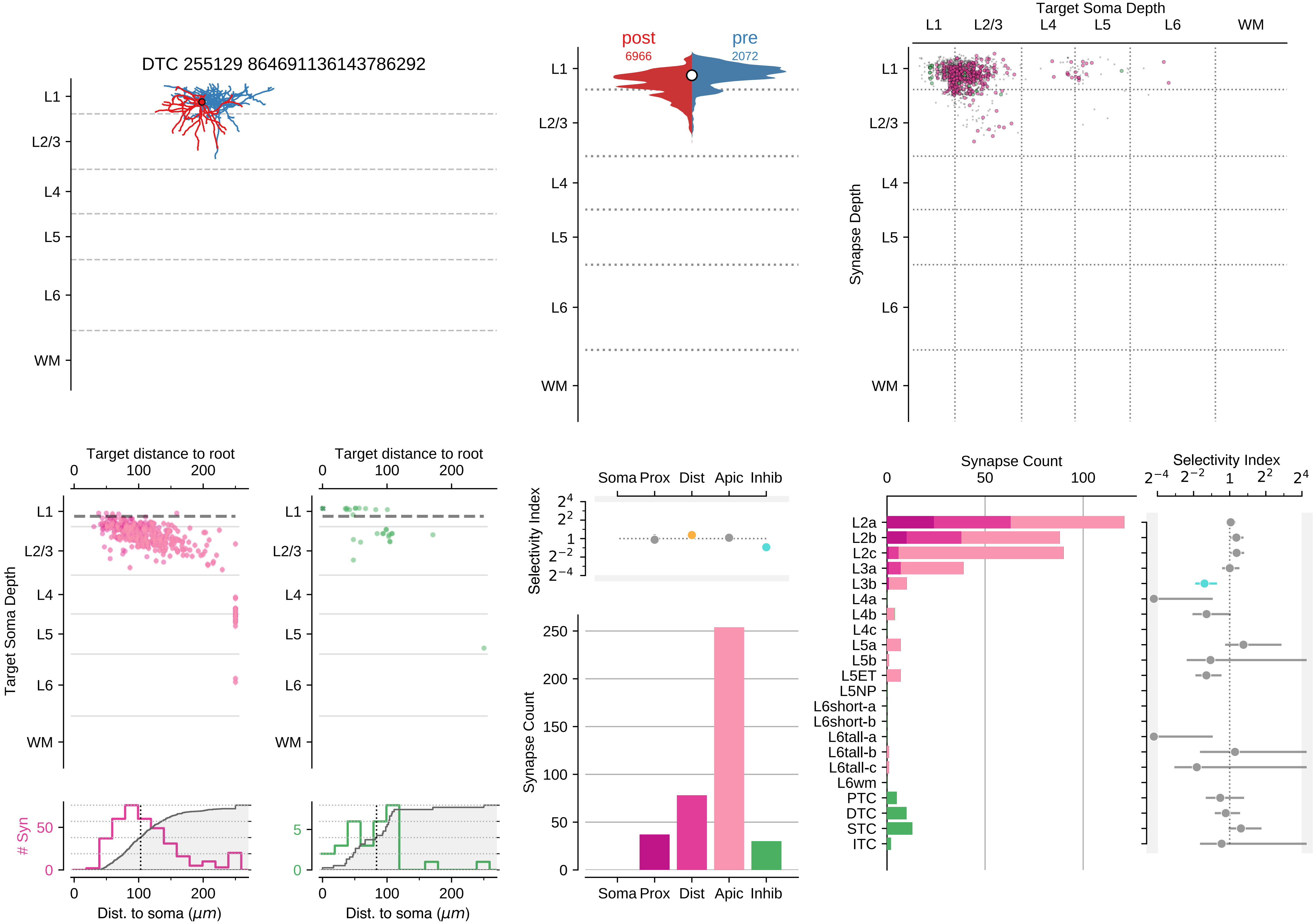

DTC 255129 864691136143786292
L1
L2/3
L4
L5
L6
WM
post
6966
pre
2072
Target Soma Depth
Synapse Depth
Target distance to root
Target Soma Depth
### Syn
Dist. to soma ($\mu m$)
Selectivity Index
Soma Prox
Dist
Apic
Inhib
Synapse Count
L2a
L2b
L2c
L3a
L3b
L4a
L4b
L4c
L5a
L5b
L5ET
L5NP
L6short-a
L6short-b
L6tall-a
L6tall-b
L6tall-c
L6wm
PTC
DTC
STC
ITC

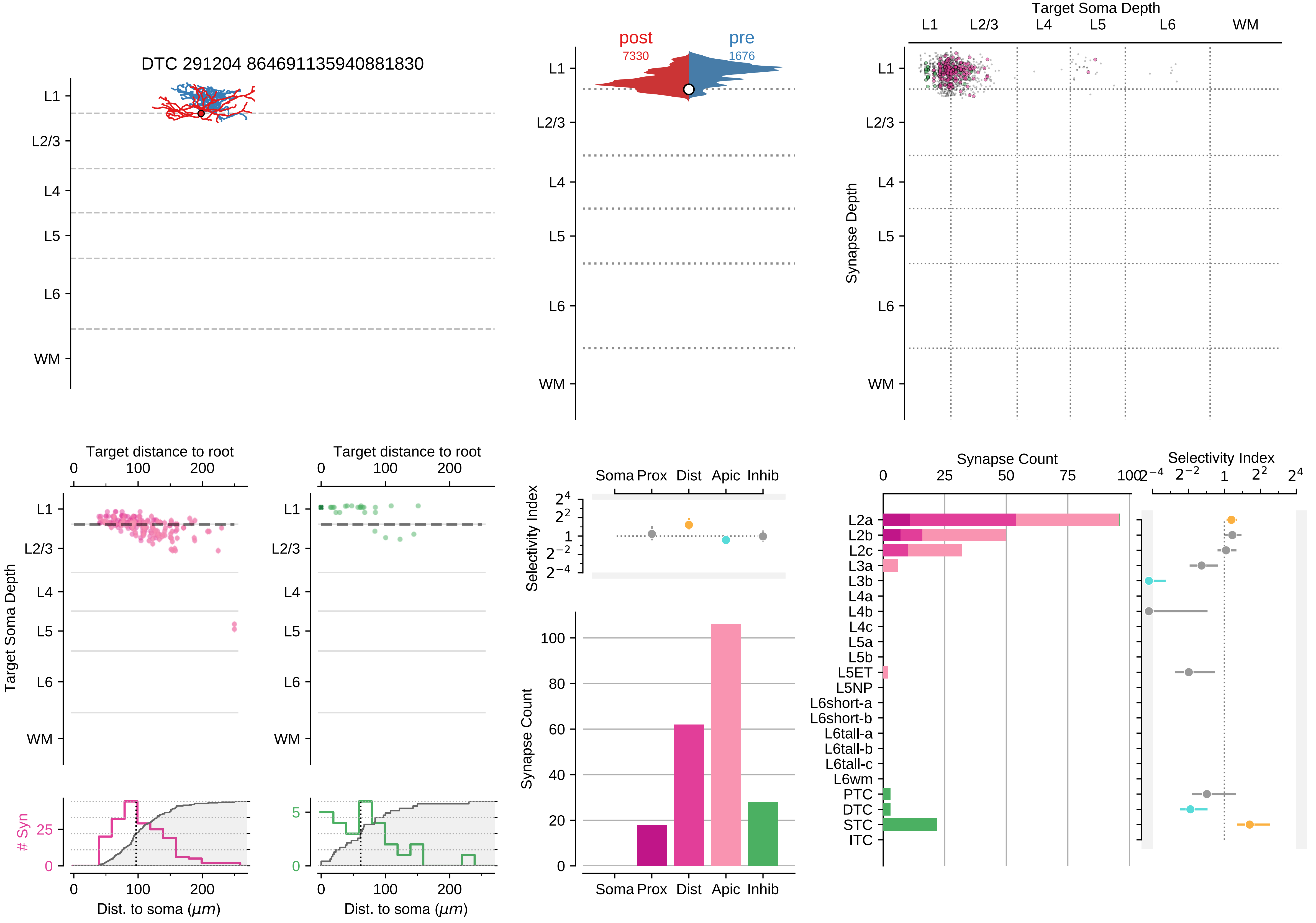

DTC 291204 864691135940881830
L1
L2/3
L4
L5
L6
WM
post
7330
pre
1676
Target Soma Depth
Synapse Depth
Target distance to root
Target Soma Depth
### Syn
Dist. to soma ($\mu m$)
Selectivity Index
Soma Prox
Dist
Apic
Inhib
Synapse Count
L2a
L2b
L2c
L3a
L3b
L4a
L4b
L4c
L5a
L5b
L5ET
L5NP
L6short-a
L6short-b
L6tall-a
L6tall-b
L6tall-c
L6wm
PTC
DTC
STC
ITC

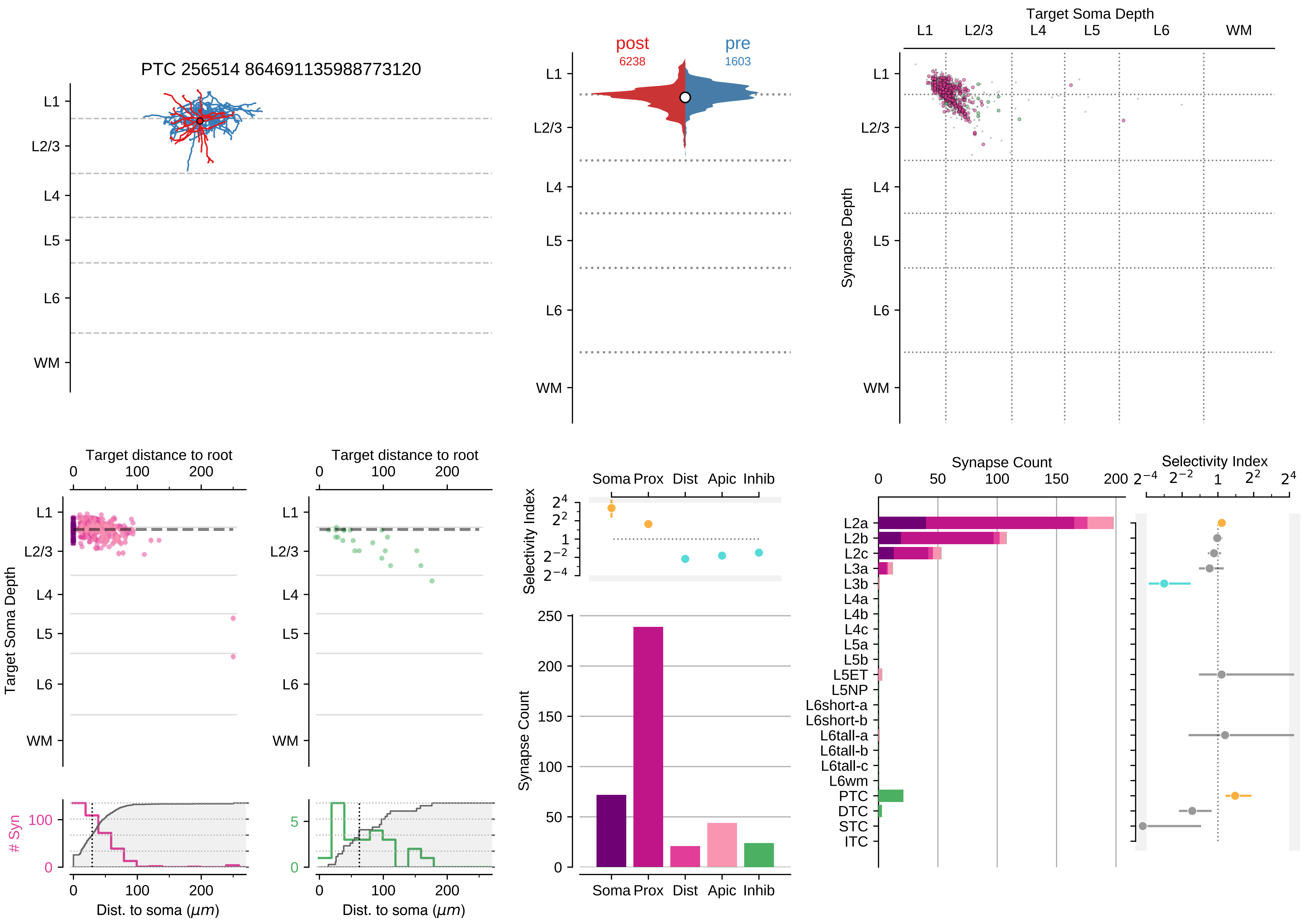

PTC 256514 864691135988773120
L1
L2/3
L4
L5
L6
WM
post
6238
pre
1603
Target Soma Depth
Synapse Depth
Target distance to root
0
100
200
Target Soma Depth
### Syn
Dist. to soma ($\mu m$)
Selectivity Index
Soma
Prox
Dist
Apic
Inhib
Synapse Count
Synapse Count
L2a
L2b
L2c
L3a
L3b
L4a
L4b
L4c
L5a
L5b
L5ET
L5NP
L6short-a
L6short-b
L6tall-a
L6tall-b
L6tall-c
L6wm
PTC
DTC
STC
ITC
Selectivity Index

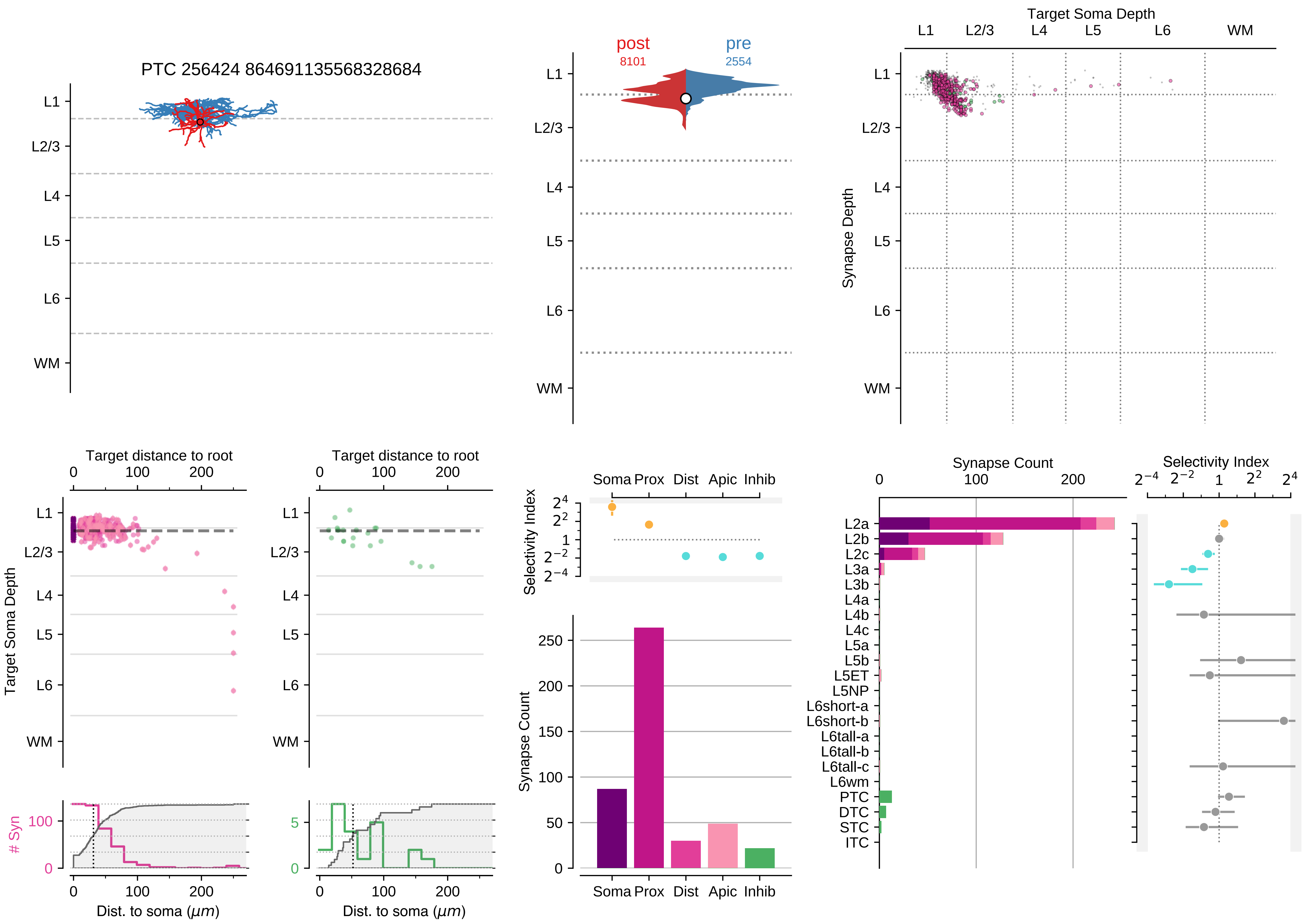
PTC 256424 864691135568328684
L1
L2/3
L4
L5
L6
WM
post
8101
pre
2554
Target Soma Depth
Synapse Depth
Target distance to root
0
100
200
Target Soma Depth
### Syn
Dist. to soma (μm)
Selectivity Index
Soma Prox Dist Apic Inhib
Synapse Count
L2a
L2b
L2c
L3a
L3b
L4a
L4b
L4c
L5a
L5b
L5ET
L5NP
L6short-a
L6short-b
L6tall-a
L6tall-b
L6tall-c
L6wm
PTC
DTC
STC
ITC

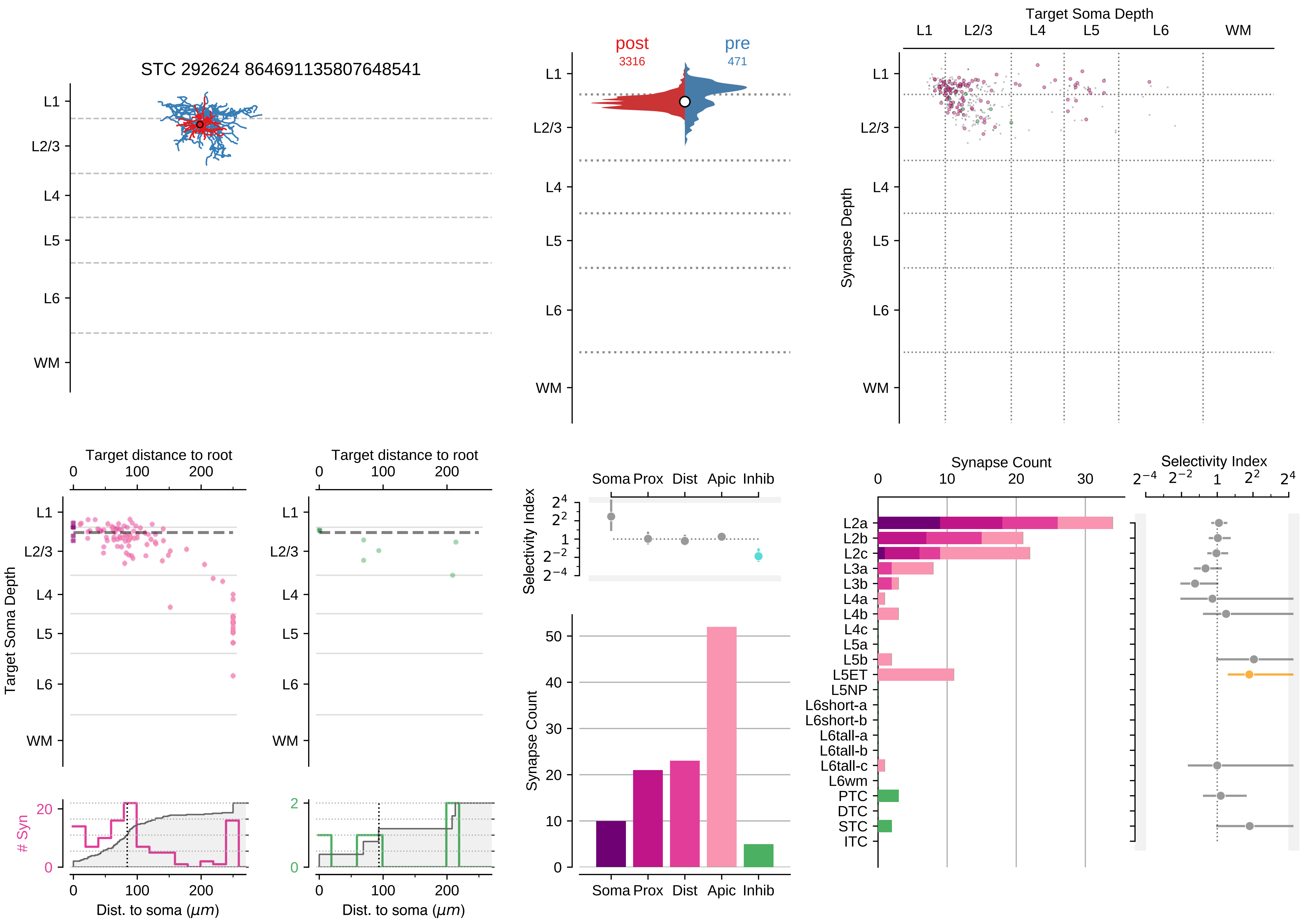

STC 292624 864691135807648541
L1
L2/3
L4
L5
L6
WM
post
3316
pre
471
Target Soma Depth
Synapse Depth
Target distance to root
Target Soma Depth
### Syn
Dist. to soma (μm)
Selectivity Index
Soma Prox Dist Apic Inhib
Synapse Count
Synapse Count
Selectivity Index
L2a
L2b
L2c
L3a
L3b
L4a
L4b
L4c
L5a
L5b
L5ET
L5NP
L6short-a
L6short-b
L6tall-a
L6tall-b
L6tall-c
L6wm
PTC
DTC
STC
ITC

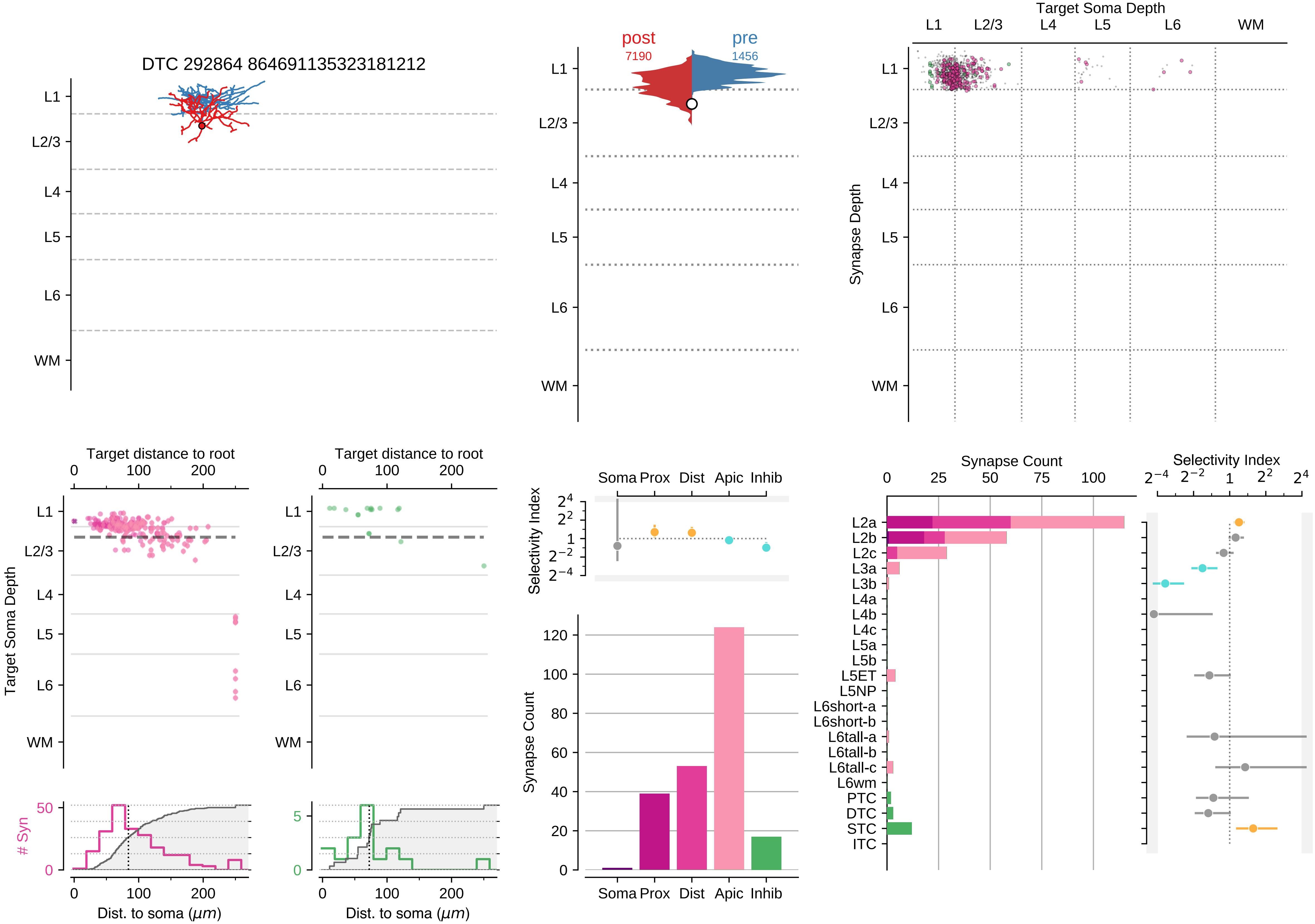

DTC 292864 864691135323181212
post
7190
pre
1456
Target Soma Depth
Synapse Depth
L1
L2/3
L4
L5
L6
WM
Target distance to root
Target Soma Depth
Dist. to soma (μm)
### Syn
Selectivity Index
Soma
Prox
Dist
Apic
Inhib
Synapse Count
L2a
L2b
L2c
L3a
L3b
L4a
L4b
L4c
L5a
L5b
L5ET
L5NP
L6short-a
L6short-b
L6tall-a
L6tall-b
L6tall-c
L6wm
PTC
DTC
STC
ITC

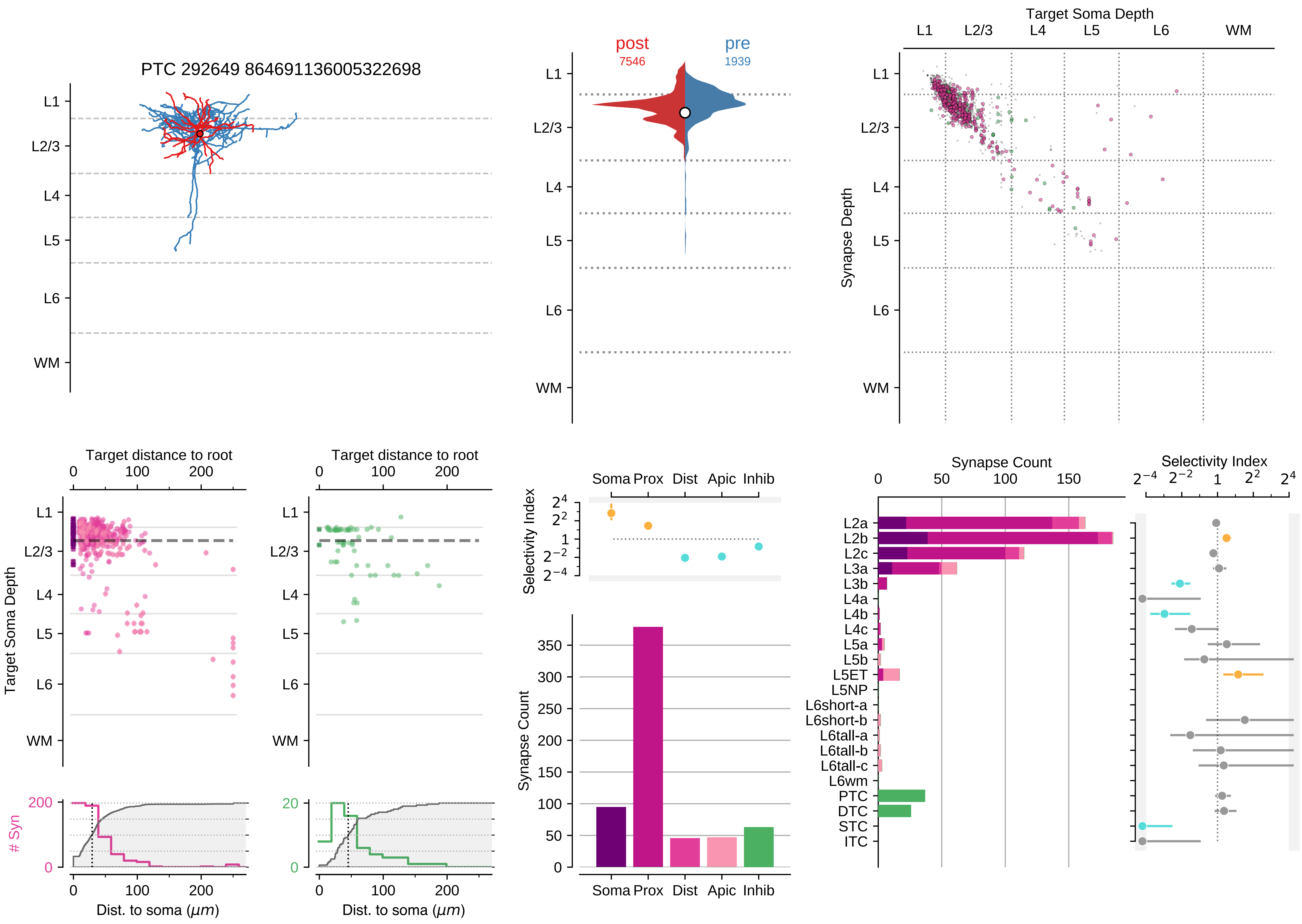
PTC 292649 864691136005322698
L1
L2/3
L4
L5
L6
WM
post
7546
pre
1939
Target Soma Depth
Synapse Depth
Target distance to root
Dist. to soma (μm)
### Syn
Selectivity Index
Soma Prox Dist Apic Inhib
Synapse Count
L2a
L2b
L2c
L3a
L3b
L4a
L4b
L4c
L5a
L5b
L5ET
L5NP
L6short-a
L6short-b
L6tall-a
L6tall-b
L6tall-c
L6wm
PTC
DTC
STC
ITC

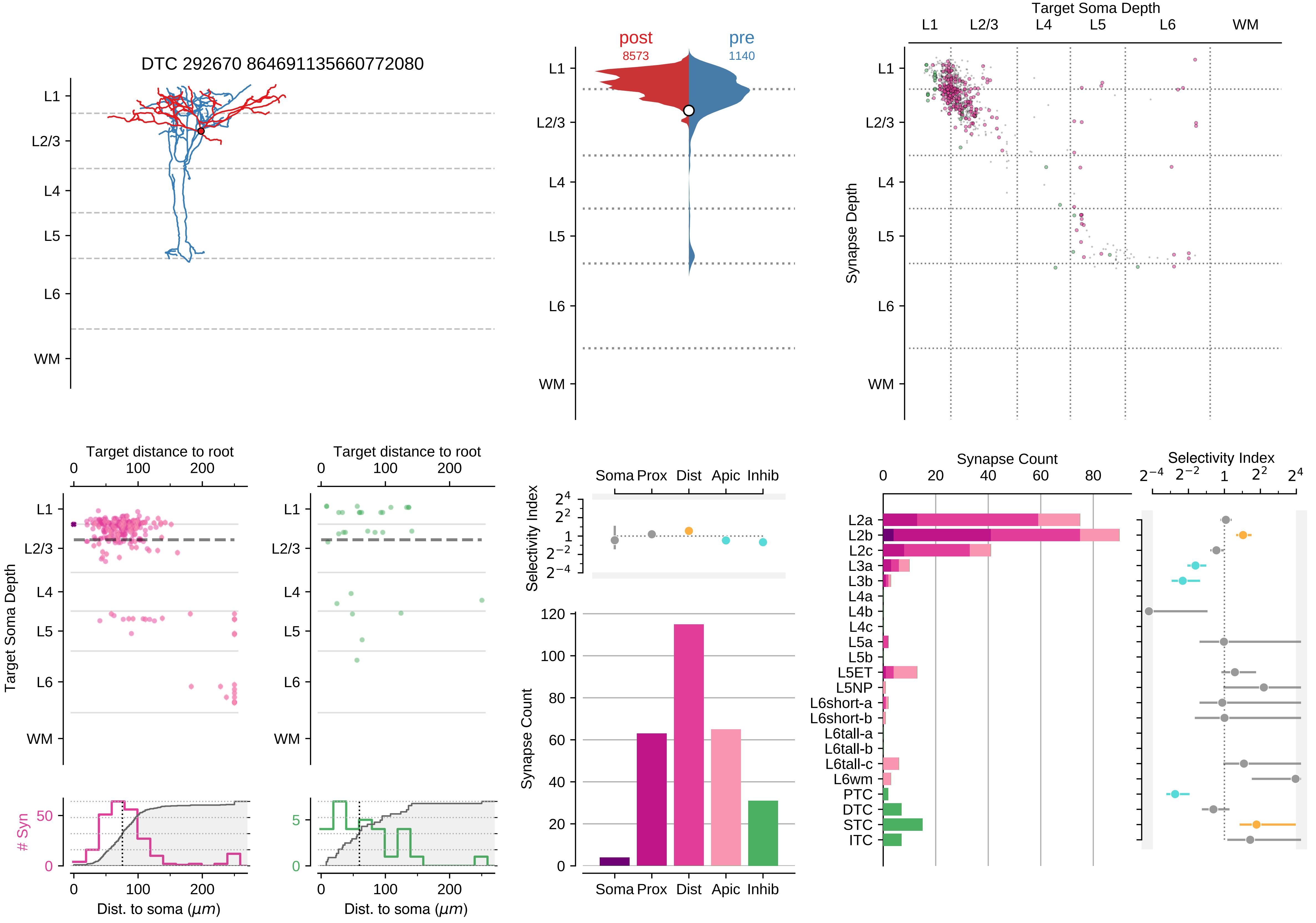

DTC 292670 864691135660772080
post
8573
pre
1140
Target Soma Depth
Synapse Depth
L1
L2/3
L4
L5
L6
WM
Target distance to root
Target Soma Depth
### Syn
Dist. to soma ($\mu m$)
Selectivity Index
Soma
Prox
Dist
Apic
Inhib
Synapse Count
L2a
L2b
L2c
L3a
L3b
L4a
L4b
L4c
L5a
L5b
L5ET
L5NP
L6short-a
L6short-b
L6tall-a
L6tall-b
L6tall-c
L6wm
PTC
DTC
STC
ITC

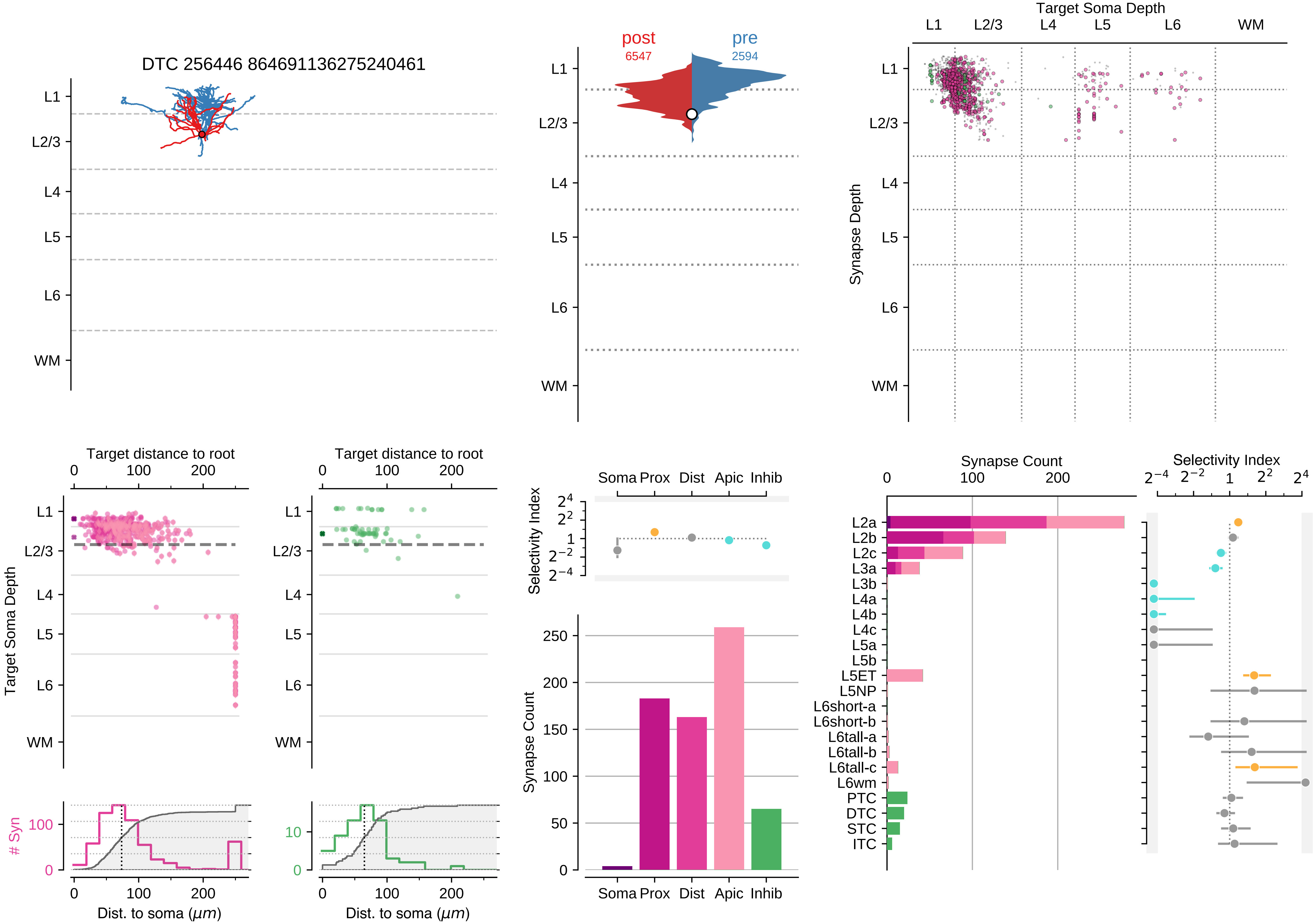
DTC 256446 864691136275240461
L1
L2/3
L4
L5
L6
WM
post
6547
pre
2594
Target Soma Depth
Synapse Depth
Target distance to root
Target Soma Depth
### Syn
Dist. to soma (μm)
Selectivity Index
Soma
Prox
Dist
Apic
Inhib
Synapse Count
L2a
L2b
L2c
L3a
L3b
L4a
L4b
L4c
L5a
L5b
L5ET
L5NP
L6short-a
L6short-b
L6tall-a
L6tall-b
L6tall-c
L6wm
PTC
DTC
STC
ITC

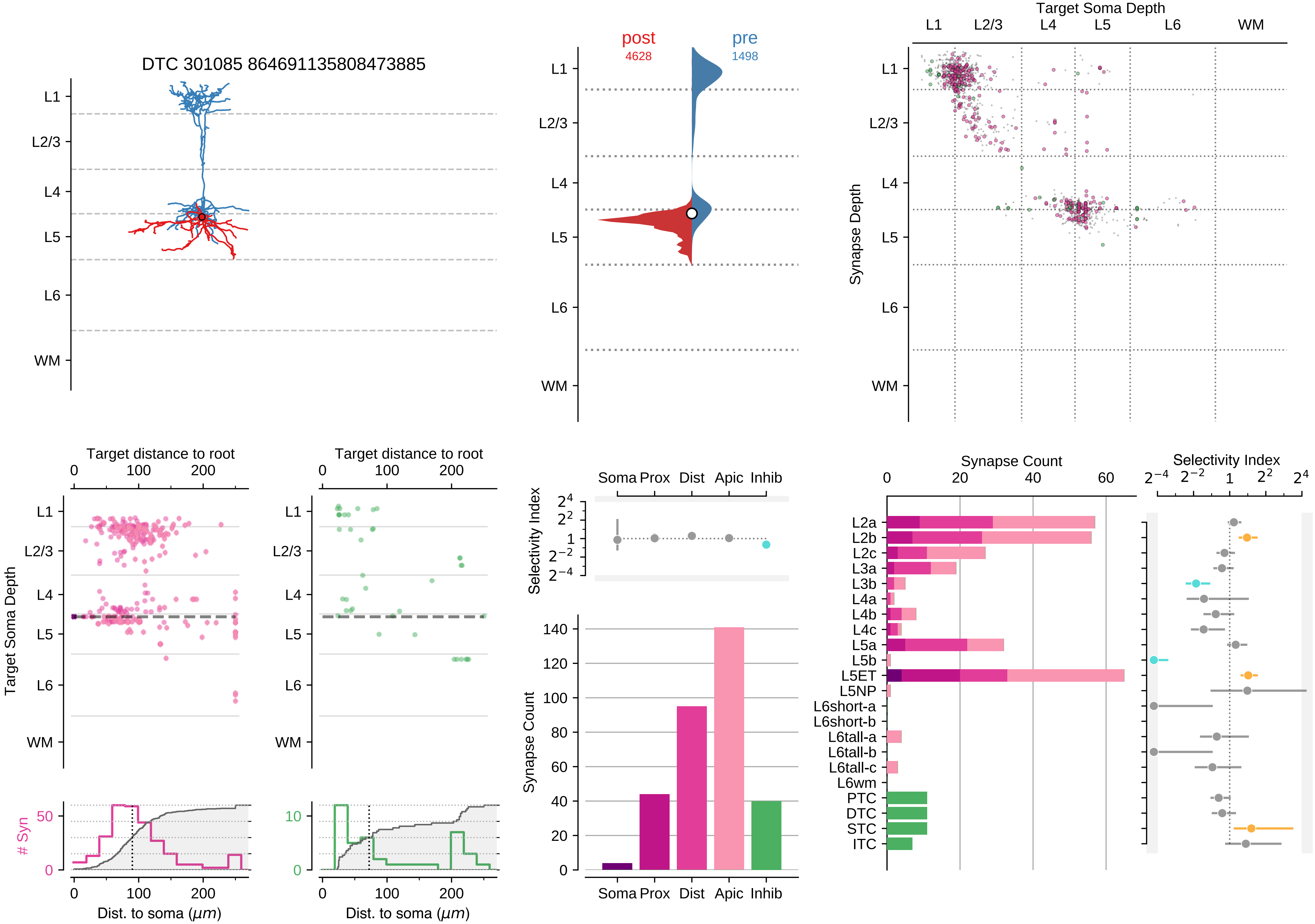

DTC 301085 864691135808473885
L1
L2/3
L4
L5
L6
WM
post
4628
pre
1498
Target Soma Depth
Synapse Depth
Target distance to root
Target Soma Depth
### Syn
Dist. to soma ($\mu m$)
Selectivity Index
Soma
Prox
Dist
Apic
Inhib
Synapse Count
L2a
L2b
L2c
L3a
L3b
L4a
L4b
L4c
L5a
L5b
L5ET
L5NP
L6short-a
L6short-b
L6tall-a
L6tall-b
L6tall-c
L6wm
PTC
DTC
STC
ITC

### Motif Group 2

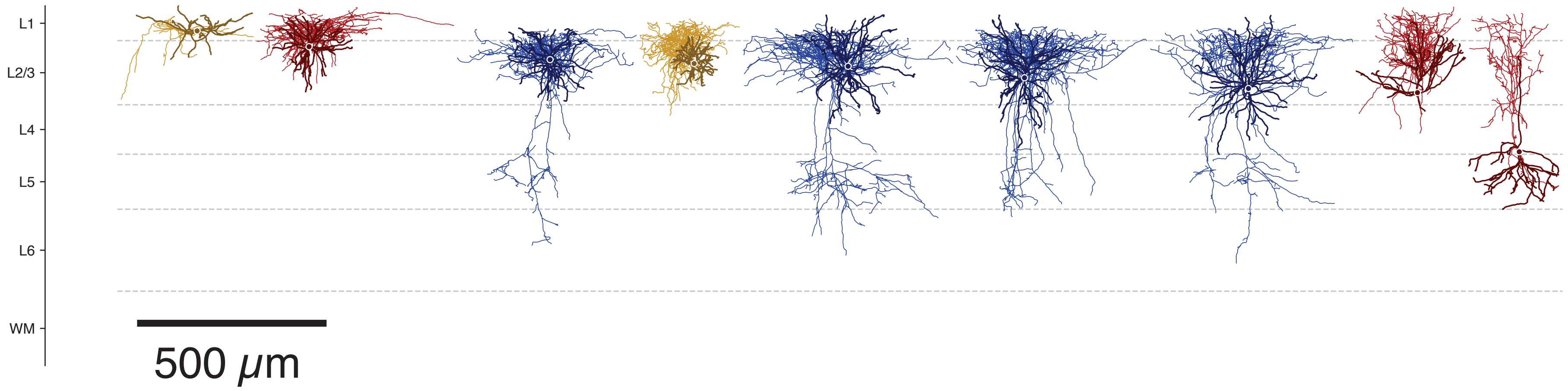

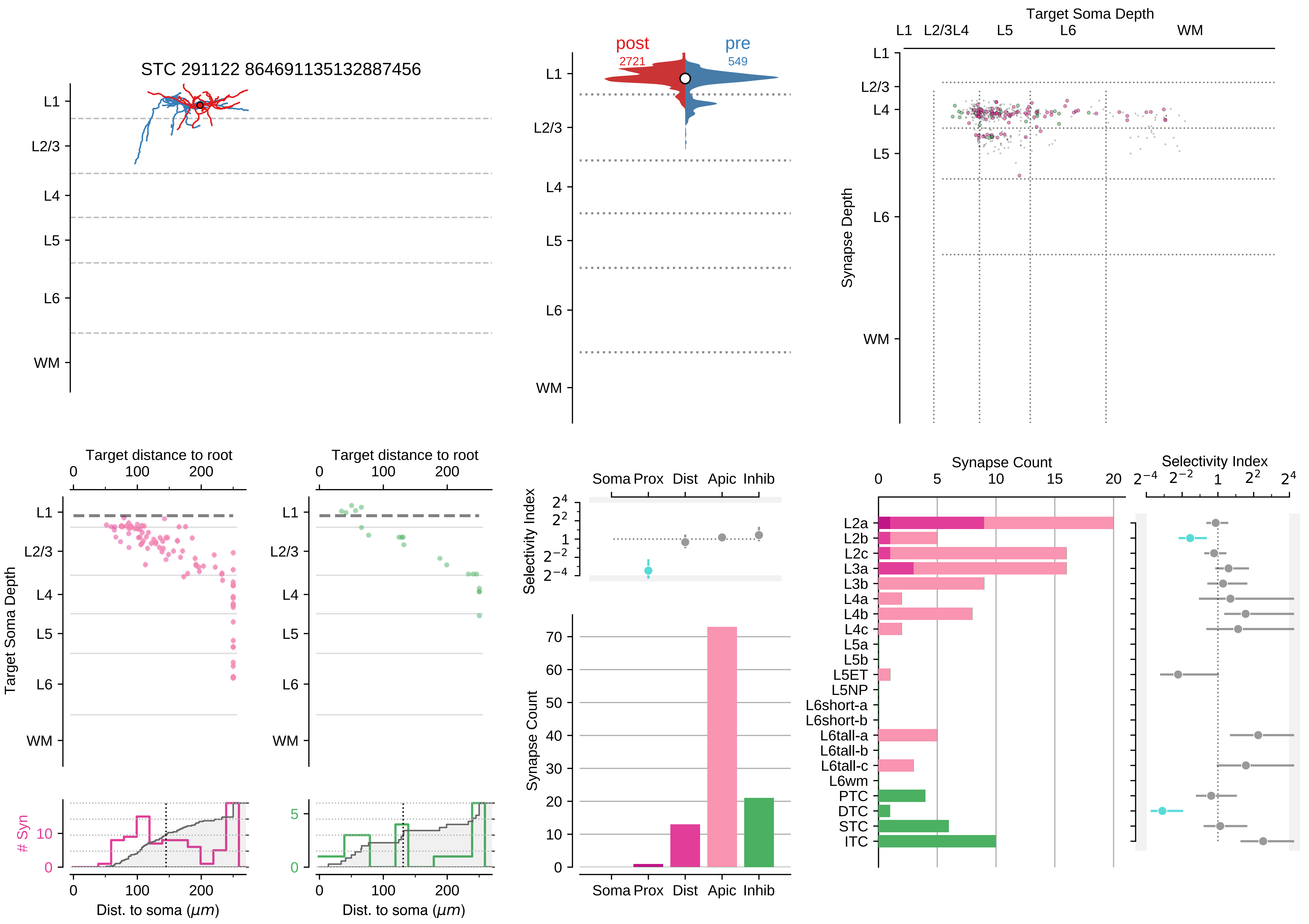
STC 291122 864691135132887456
post
2721
pre
549
Target Soma Depth
Synapse Depth
L1
L2/3
L4
L5
L6
WM
Target distance to root
Target Soma Depth
### Syn
Dist. to soma (μm)
Selectivity Index
Synapse Count
Soma
Prox
Dist
Apic
Inhib
L2a
L2b
L2c
L3a
L3b
L4a
L4b
L4c
L5a
L5b
L5ET
L5NP
L6short-a
L6short-b
L6tall-a
L6tall-b
L6tall-c
L6wm
PTC
DTC
STC
ITC

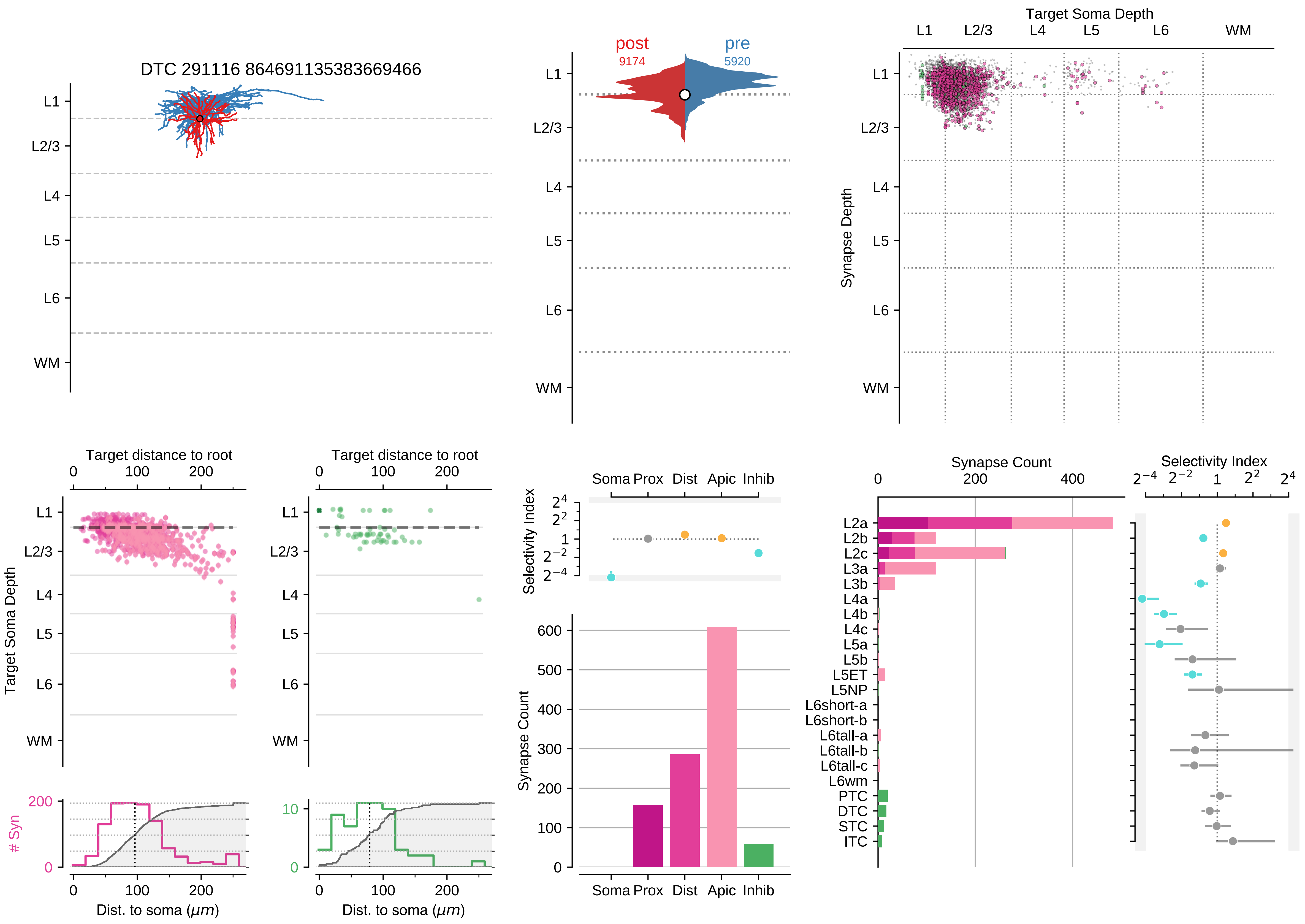

DTC 291116 864691135383669466
L1
L2/3
L4
L5
L6
WM
post
9174
pre
5920
Target Soma Depth
Synapse Depth
Target distance to root
Target Soma Depth
### Syn
Dist. to soma (μm)
Selectivity Index
Soma Prox Dist Apic Inhib
Synapse Count
L2a
L2b
L2c
L3a
L3b
L4a
L4b
L4c
L5a
L5b
L5ET
L5NP
L6short-a
L6short-b
L6tall-a
L6tall-b
L6tall-c
L6wm
PTC
DTC
STC
ITC

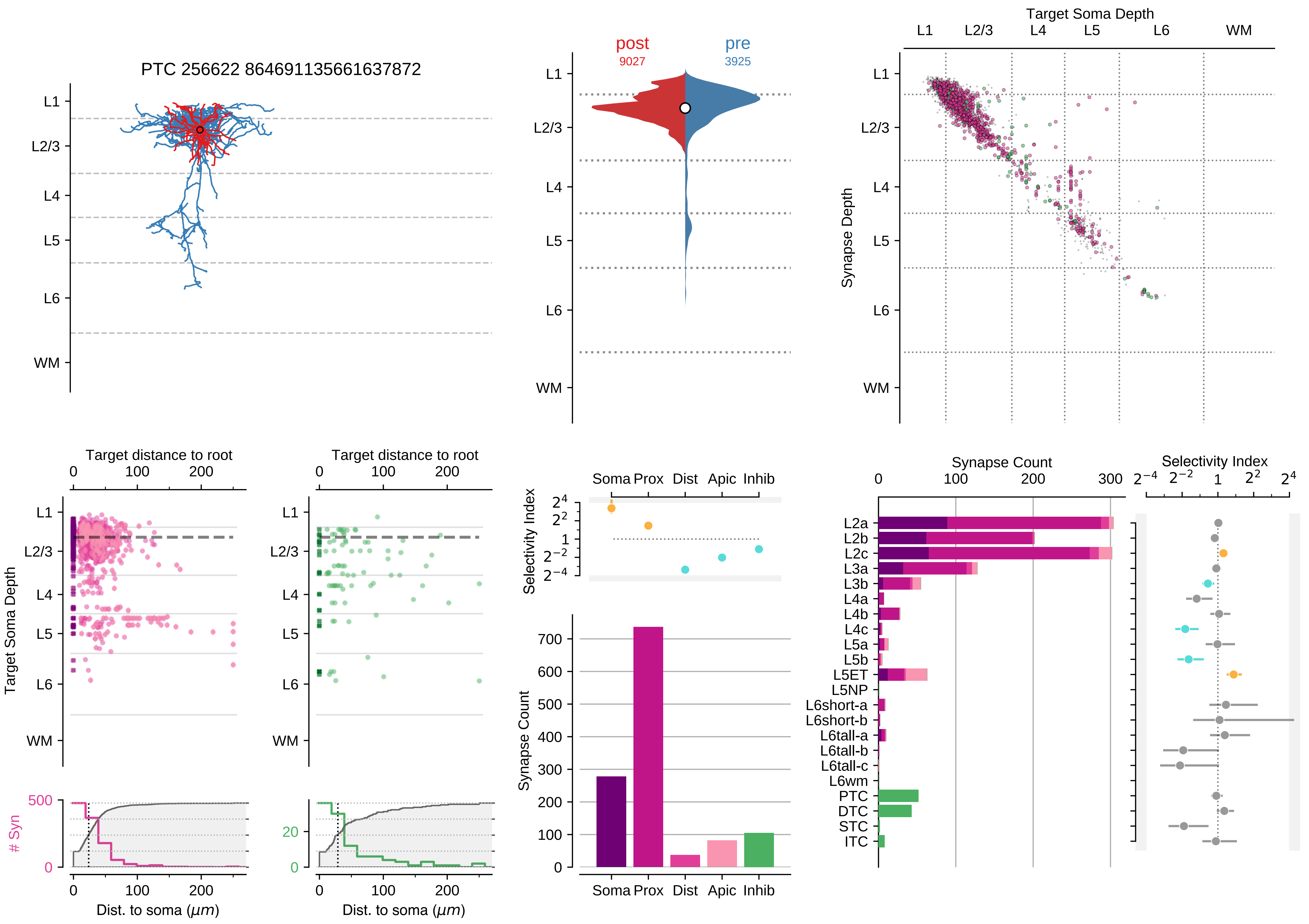
PTC 256622 864691135661637872
L1
L2/3
L4
L5
L6
WM
post
9027
pre
3925
Target Soma Depth
Synapse Depth
Target distance to root
Target Soma Depth
### Syn
Dist. to soma (μm)
Selectivity Index
Soma
Prox
Dist
Apic
Inhib
Synapse Count
L2a
L2b
L2c
L3a
L3b
L4a
L4b
L4c
L5a
L5b
L5ET
L5NP
L6short-a
L6short-b
L6tall-a
L6tall-b
L6tall-c
L6wm
PTC
DTC
STC
ITC

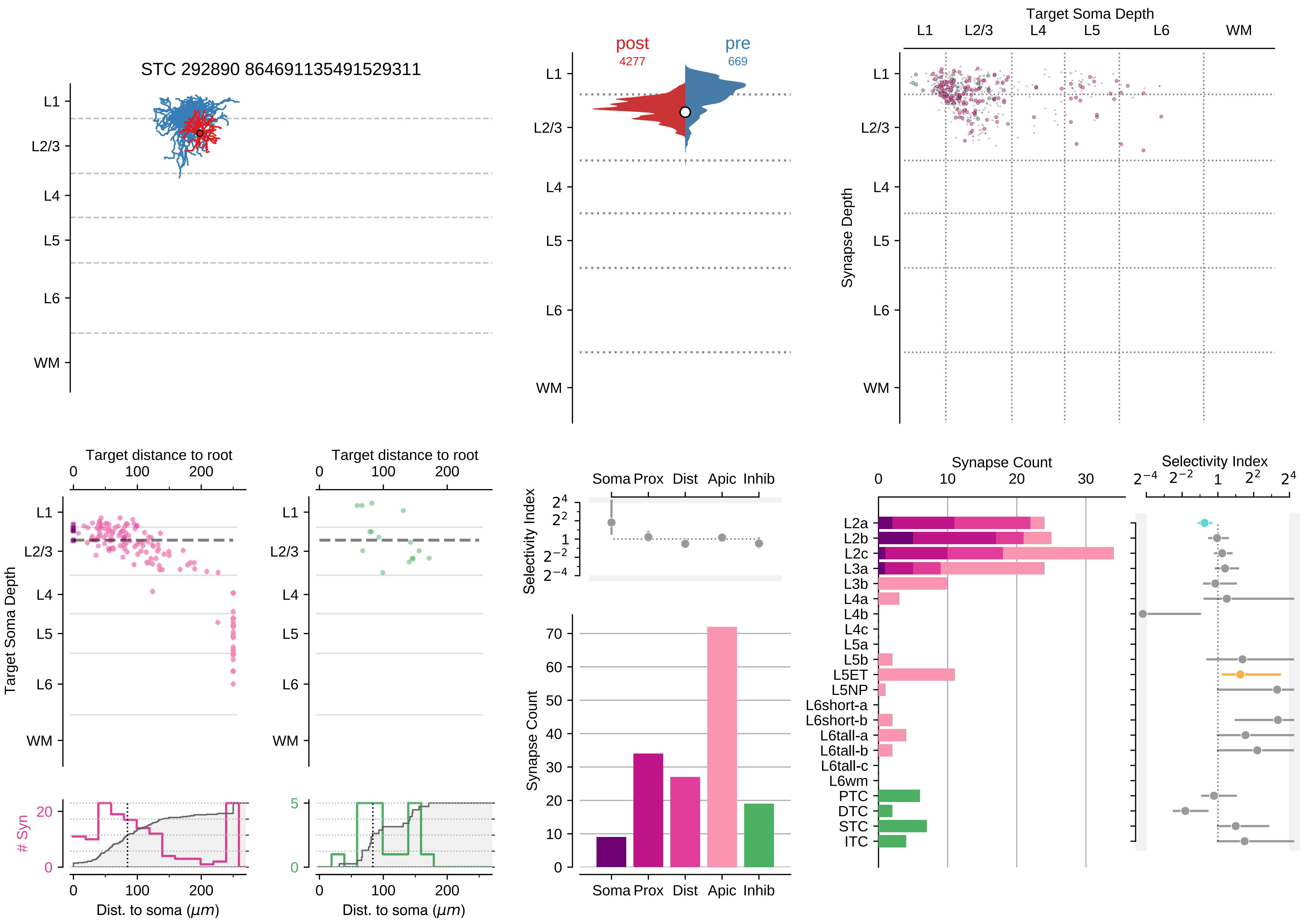

STC 292890 864691135491529311
L1
L2/3
L4
L5
L6
WM
post
4277
pre
669
Target Soma Depth
Synapse Depth
Target distance to root
Target Soma Depth
### Syn
Dist. to soma (μm)
Selectivity Index
Soma Prox Dist Apic Inhib
Synapse Count
L2a
L2b
L2c
L3a
L3b
L4a
L4b
L4c
L5a
L5b
L5ET
L5NP
L6short-a
L6short-b
L6tall-a
L6tall-b
L6tall-c
L6wm
PTC
DTC
STC
ITC

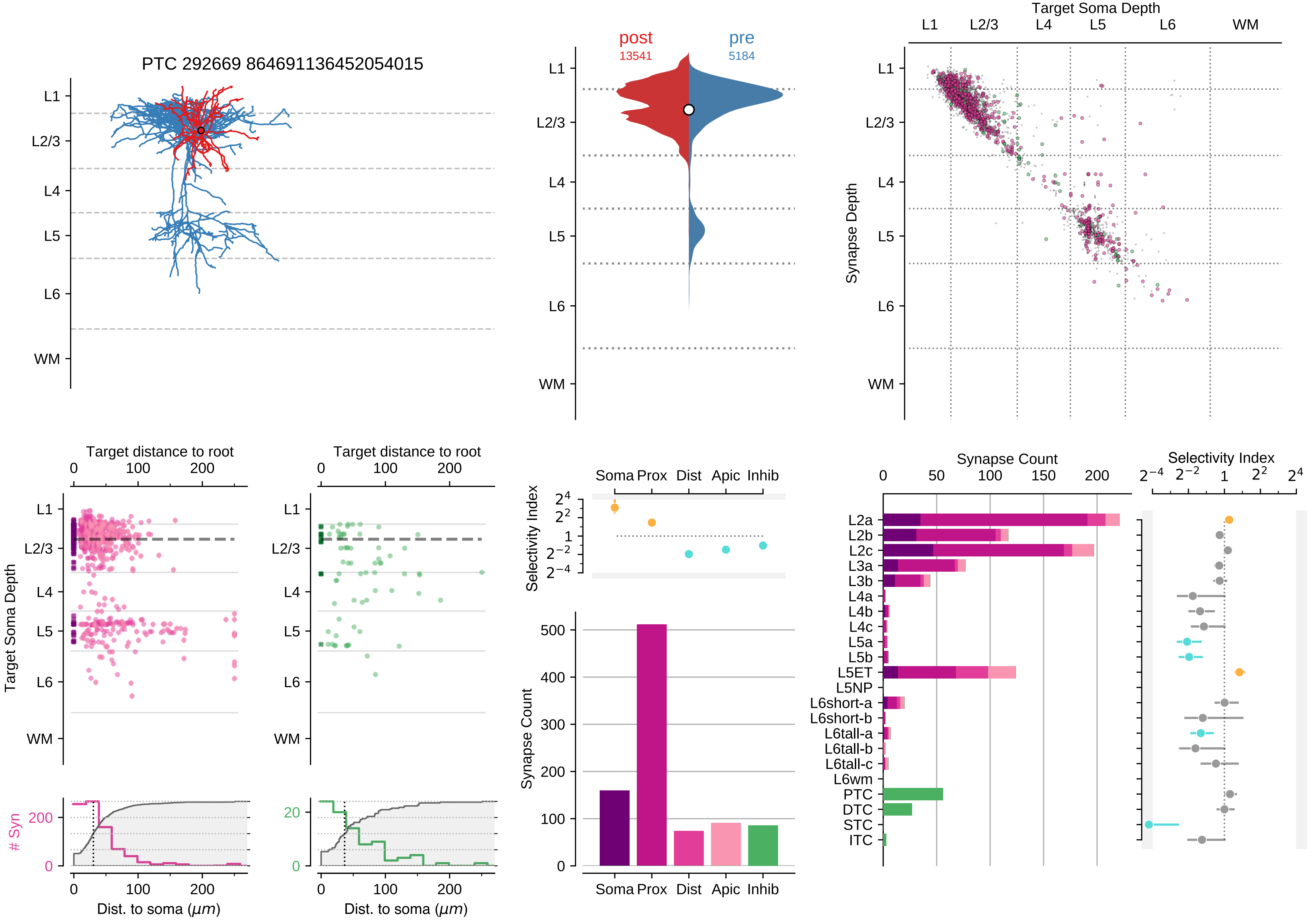

PTC 292669 864691136452054015
L1
L2/3
L4
L5
L6
WM
post
13541
pre
5184
Target Soma Depth
Synapse Depth
Target distance to root
Target Soma Depth
### Syn
Dist. to soma (μm)
Selectivity Index
Soma Prox Dist Apic Inhib
Synapse Count
Synapse Count
Selectivity Index
L2a
L2b
L2c
L3a
L3b
L4a
L4b
L4c
L5a
L5b
L5ET
L5NP
L6short-a
L6short-b
L6tall-a
L6tall-b
L6tall-c
L6wm
PTC
DTC
STC
ITC

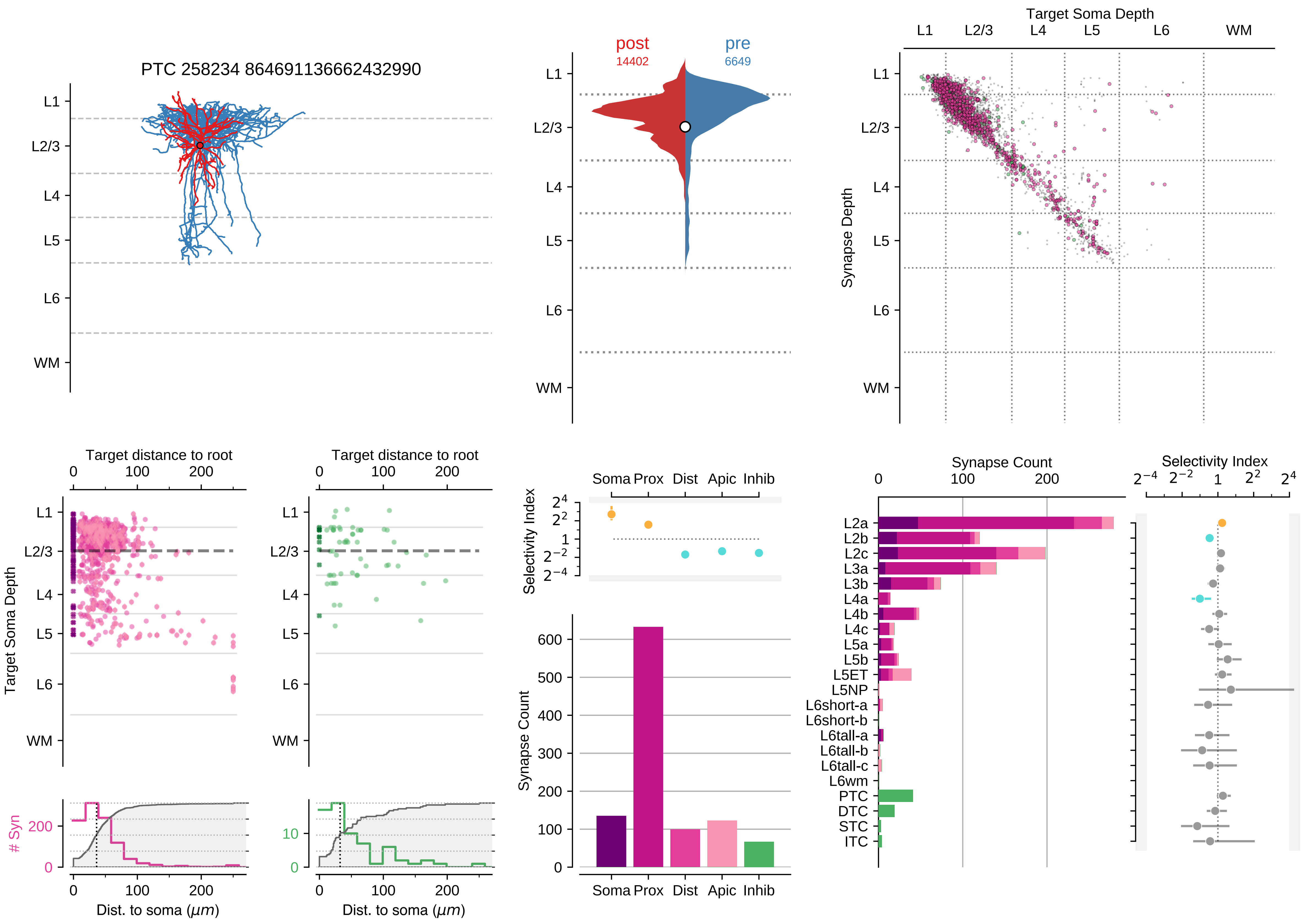

PTC 258234 864691136662432990
L1
L2/3
L4
L5
L6
WM
post
14402
pre
6649
Target Soma Depth
Synapse Depth
Target distance to root
Target Soma Depth
Dist. to soma ($\mu m$)
### Syn
Selectivity Index
Soma Prox Dist Apic Inhib
Synapse Count
L2a
L2b
L2c
L3a
L3b
L4a
L4b
L4c
L5a
L5b
L5ET
L5NP
L6short-a
L6short-b
L6tall-a
L6tall-b
L6tall-c
L6wm
PTC
DTC
STC
ITC

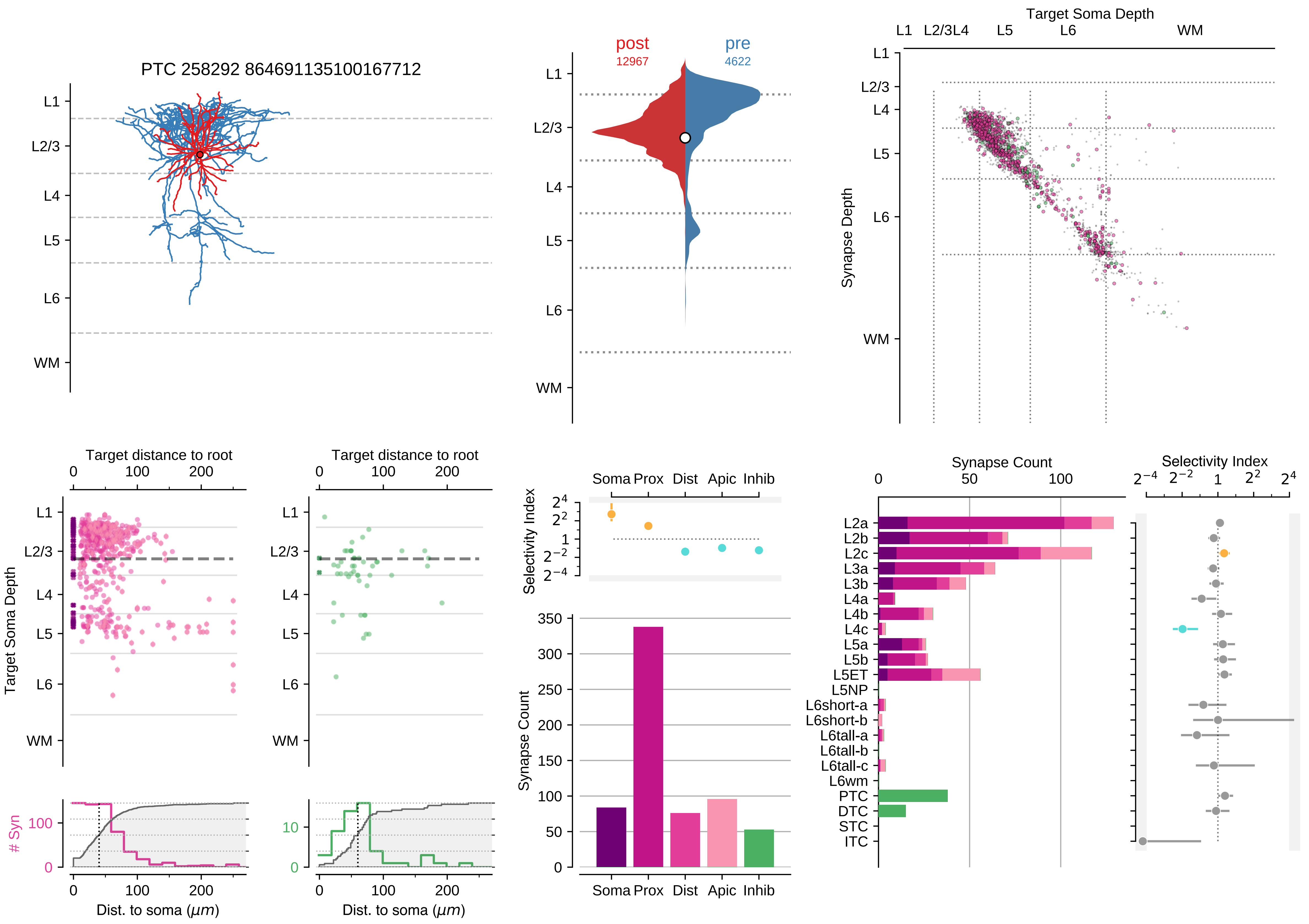
PTC 258292 864691135100167712
L1
L2/3
L4
L5
L6
WM
post
12967
pre
4622
Target Soma Depth
Synapse Depth
Target distance to root
Target Soma Depth
### Syn
Dist. to soma (μm)
Selectivity Index
Soma Prox Dist Apic Inhib
Synapse Count
L2a
L2b
L2c
L3a
L3b
L4a
L4b
L4c
L5a
L5b
L5ET
L5NP
L6short-a
L6short-b
L6tall-a
L6tall-b
L6tall-c
L6wm
PTC
DTC
STC
ITC

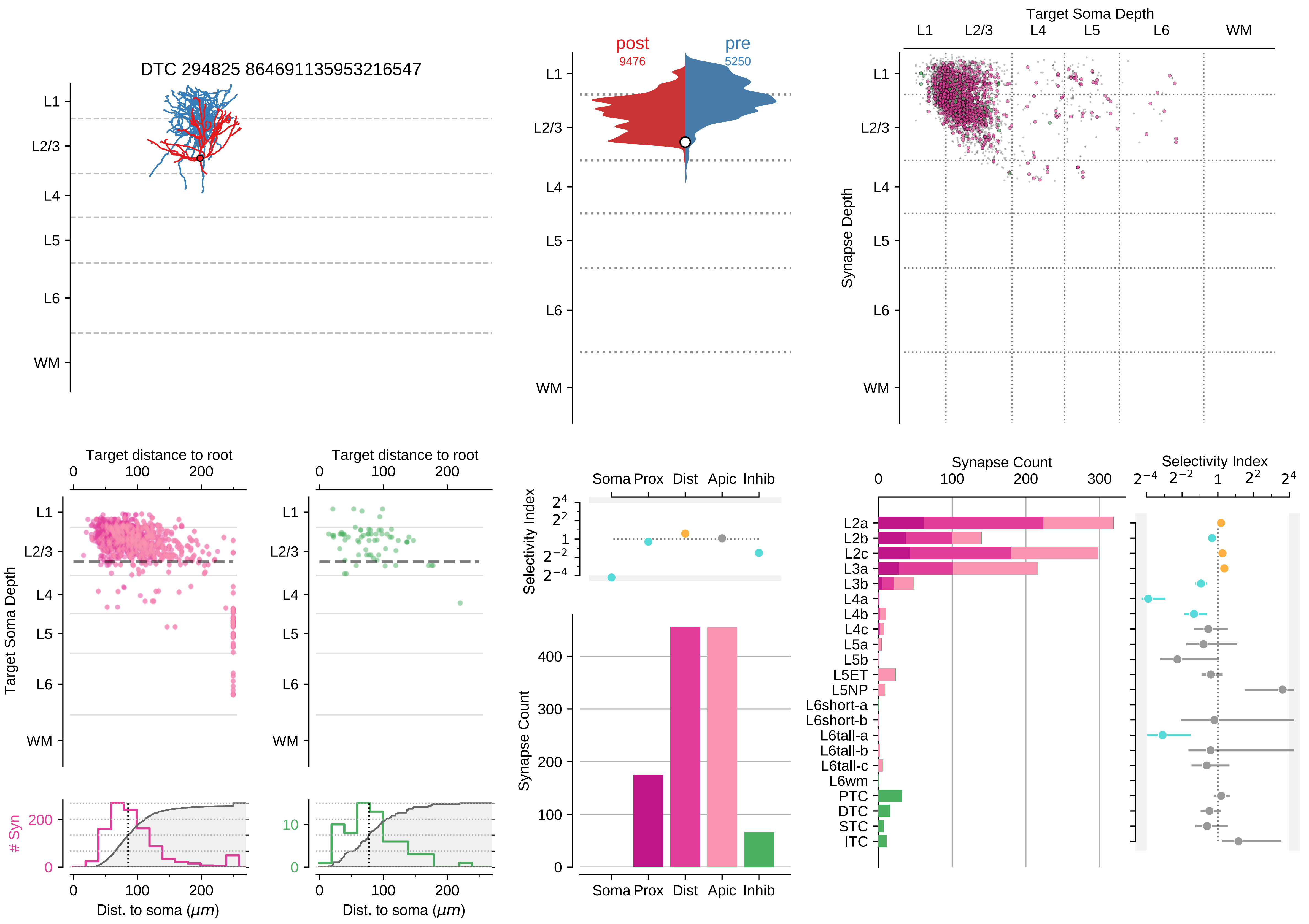

DTC 294825 864691135953216547
post
9476
pre
5250
Target Soma Depth
Synapse Depth
L1
L2/3
L4
L5
L6
WM
Target distance to root
Target Soma Depth
### Syn
Dist. to soma ($\mu m$)
Selectivity Index
Synapse Count
Soma
Prox
Dist
Apic
Inhib
L2a
L2b
L2c
L3a
L3b
L4a
L4b
L4c
L5a
L5b
L5ET
L5NP
L6short-a
L6short-b
L6tall-a
L6tall-b
L6tall-c
L6wm
PTC
DTC
STC
ITC

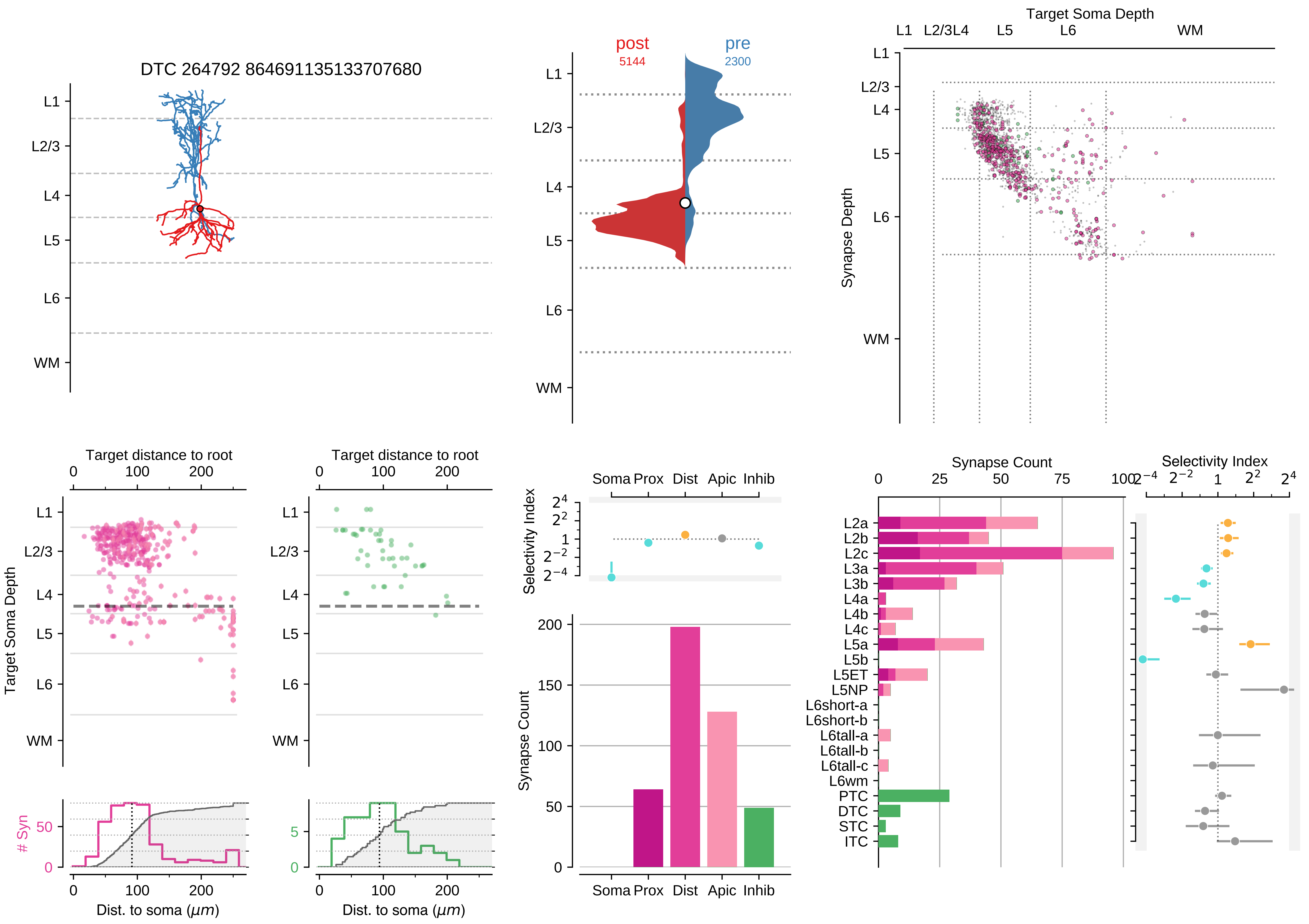

DTC 264792 864691135133707680
L1
L2/3
L4
L5
L6
WM
post
5144
pre
2300
Target Soma Depth
Synapse Depth
Target distance to root
Target Soma Depth
Dist. to soma ($\mu m$)
### Syn
Selectivity Index
Soma Prox Dist Apic Inhib
Synapse Count
L2a
L2b
L2c
L3a
L3b
L4a
L4b
L4c
L5a
L5b
L5ET
L5NP
L6short-a
L6short-b
L6tall-a
L6tall-b
L6tall-c
L6wm
PTC
DTC
STC
ITC

### Motif Group 3

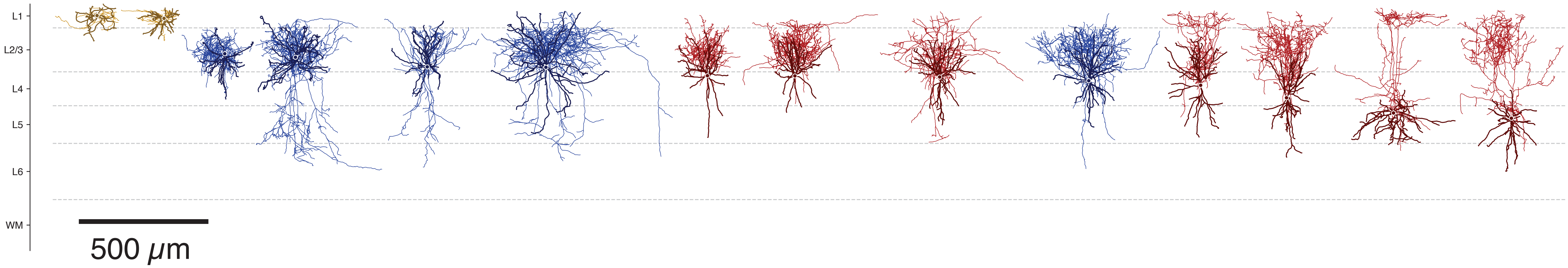

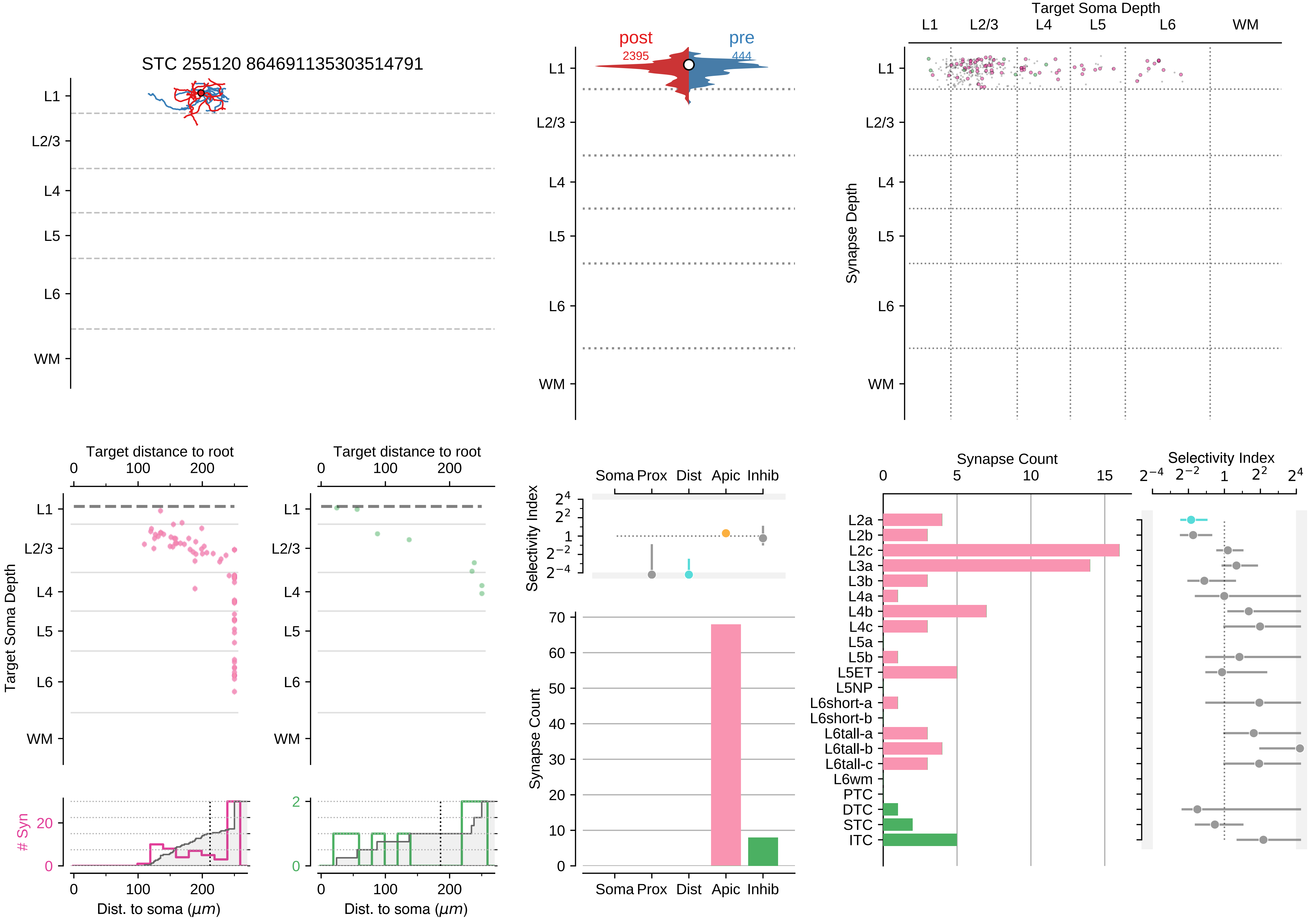

STC 255120 864691135303514791
post
2395
pre
444
Target Soma Depth
Synapse Depth
L1
L2/3
L4
L5
L6
WM
Target distance to root
Target Soma Depth
### Syn
Dist. to soma (μm)
Selectivity Index
Synapse Count
Soma Prox Dist Apic Inhib
L2a
L2b
L2c
L3a
L3b
L4a
L4b
L4c
L5a
L5b
L5ET
L5NP
L6short-a
L6short-b
L6tall-a
L6tall-b
L6tall-c
L6wm
PTC
DTC
STC
ITC

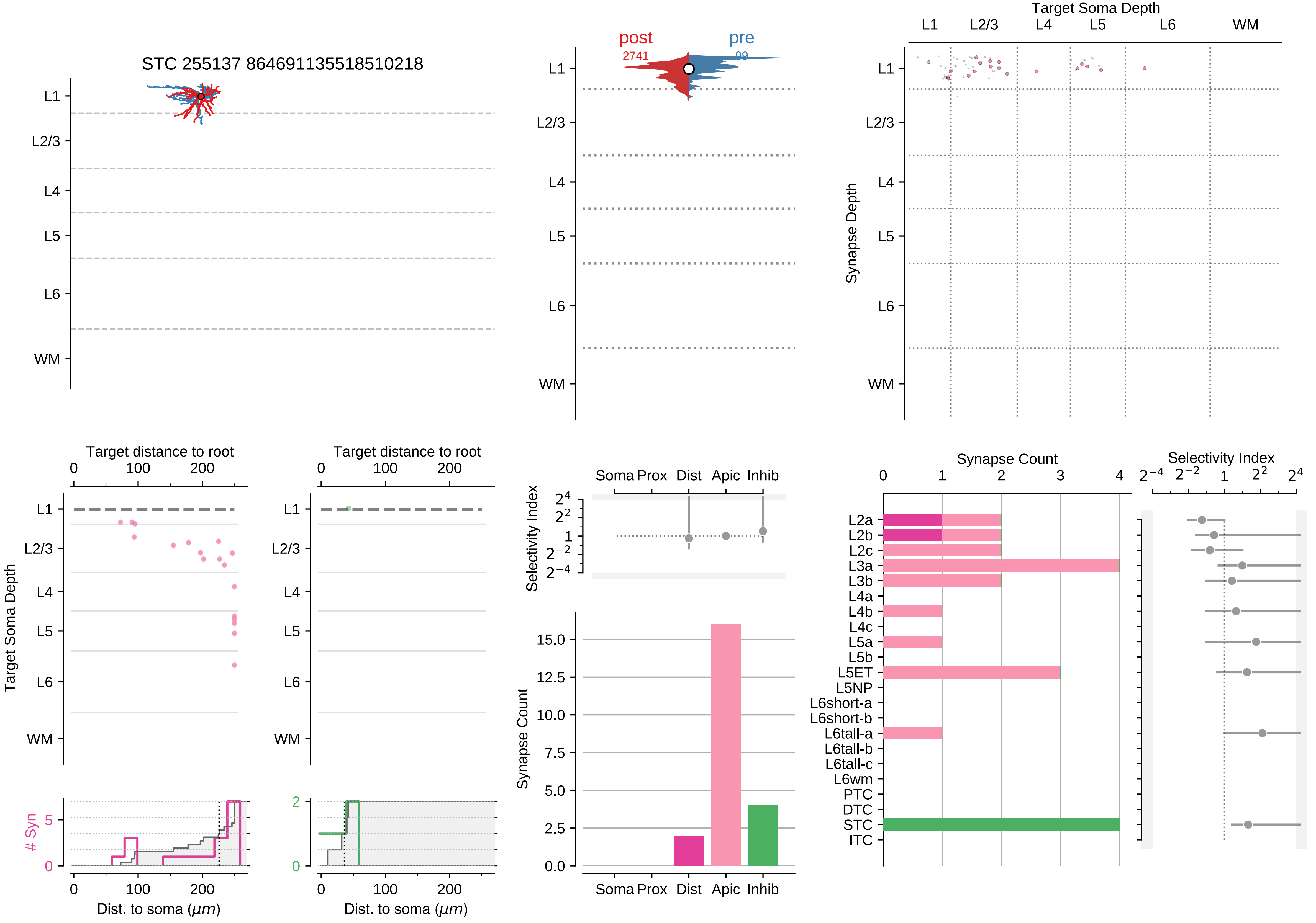

STC 255137 864691135518510218
post
2741
pre
99
Target Soma Depth
Synapse Depth
L1
L2/3
L4
L5
L6
WM
Target distance to root
Dist. to soma (μm)
### Syn
Soma Prox
Dist
Apic
Inhib
Selectivity Index
Synapse Count
L2a
L2b
L2c
L3a
L3b
L4a
L4b
L4c
L5a
L5b
L5ET
L5NP
L6short-a
L6short-b
L6tall-a
L6tall-b
L6tall-c
L6wm
PTC
DTC
STC
ITC

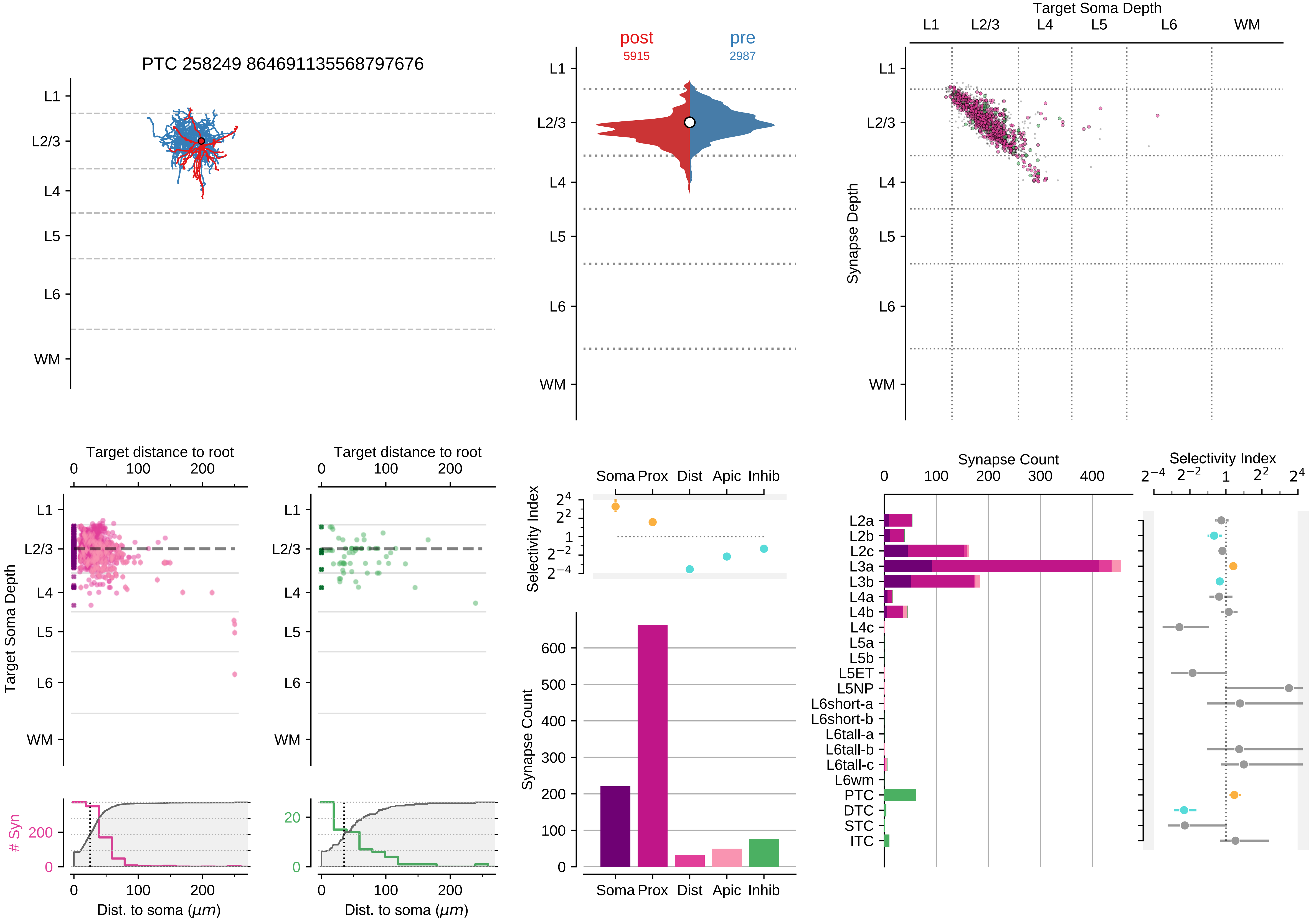
PTC 258249 864691135568797676
L1
L2/3
L4
L5
L6
WM
post
5915
pre
2987
Target Soma Depth
Synapse Depth
Target distance to root
Target Soma Depth
### Syn
Dist. to soma (μm)
Selectivity Index
Soma
Prox
Dist
Apic
Inhib
Synapse Count
L2a
L2b
L2c
L3a
L3b
L4a
L4b
L4c
L5a
L5b
L5ET
L5NP
L6short-a
L6short-b
L6tall-a
L6tall-b
L6tall-c
L6wm
PTC
DTC
STC
ITC

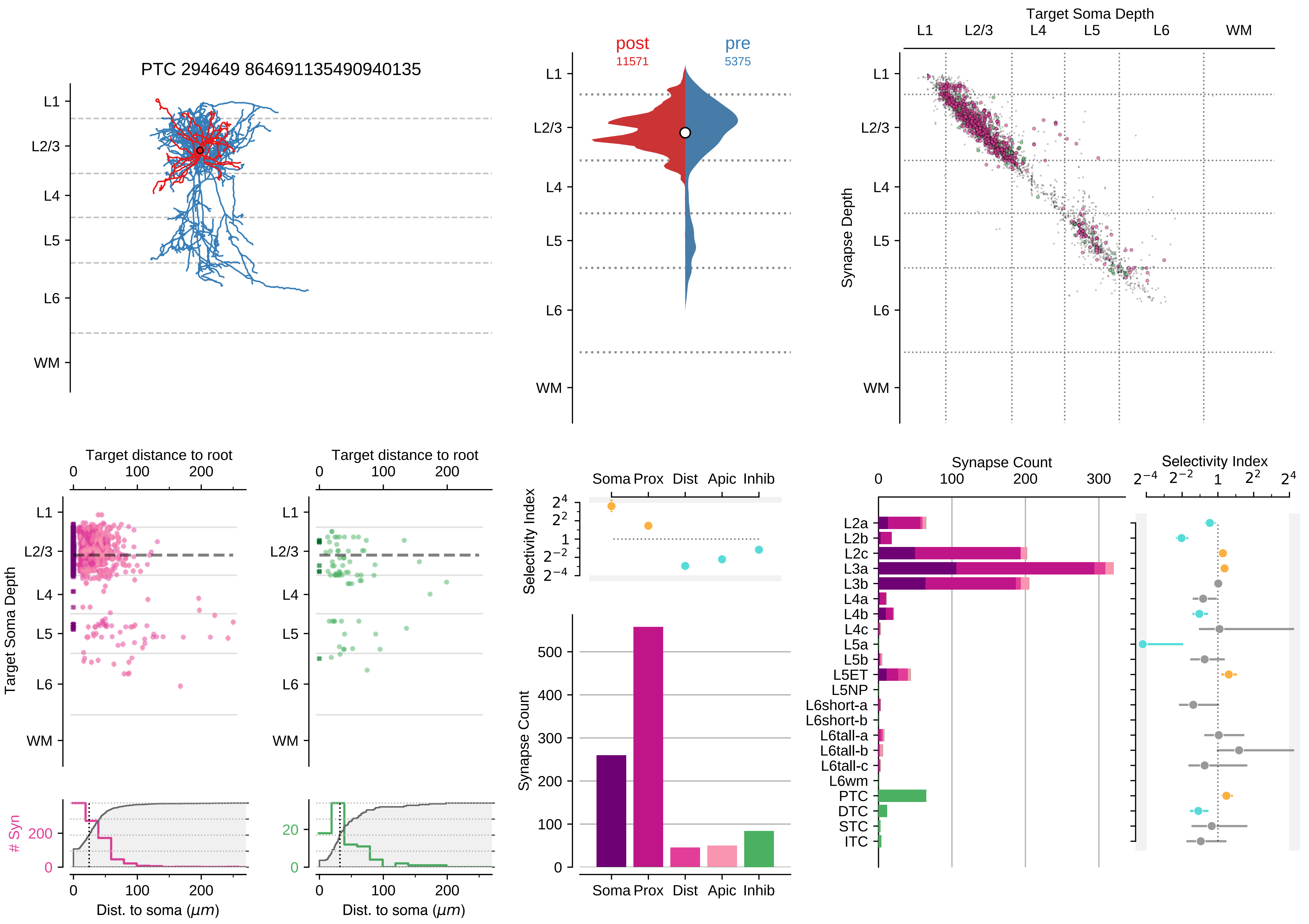
PTC 294649 864691135490940135
L1
L2/3
L4
L5
L6
WM
post
11571
pre
5375
Target Soma Depth
Synapse Depth
Target distance to root
Target Soma Depth
### Syn
Dist. to soma (μm)
Selectivity Index
Soma Prox Dist Apic Inhib
Synapse Count
L2a
L2b
L2c
L3a
L3b
L4a
L4b
L4c
L5a
L5b
L5ET
L5NP
L6short-a
L6short-b
L6tall-a
L6tall-b
L6tall-c
L6wm
PTC
DTC
STC
ITC

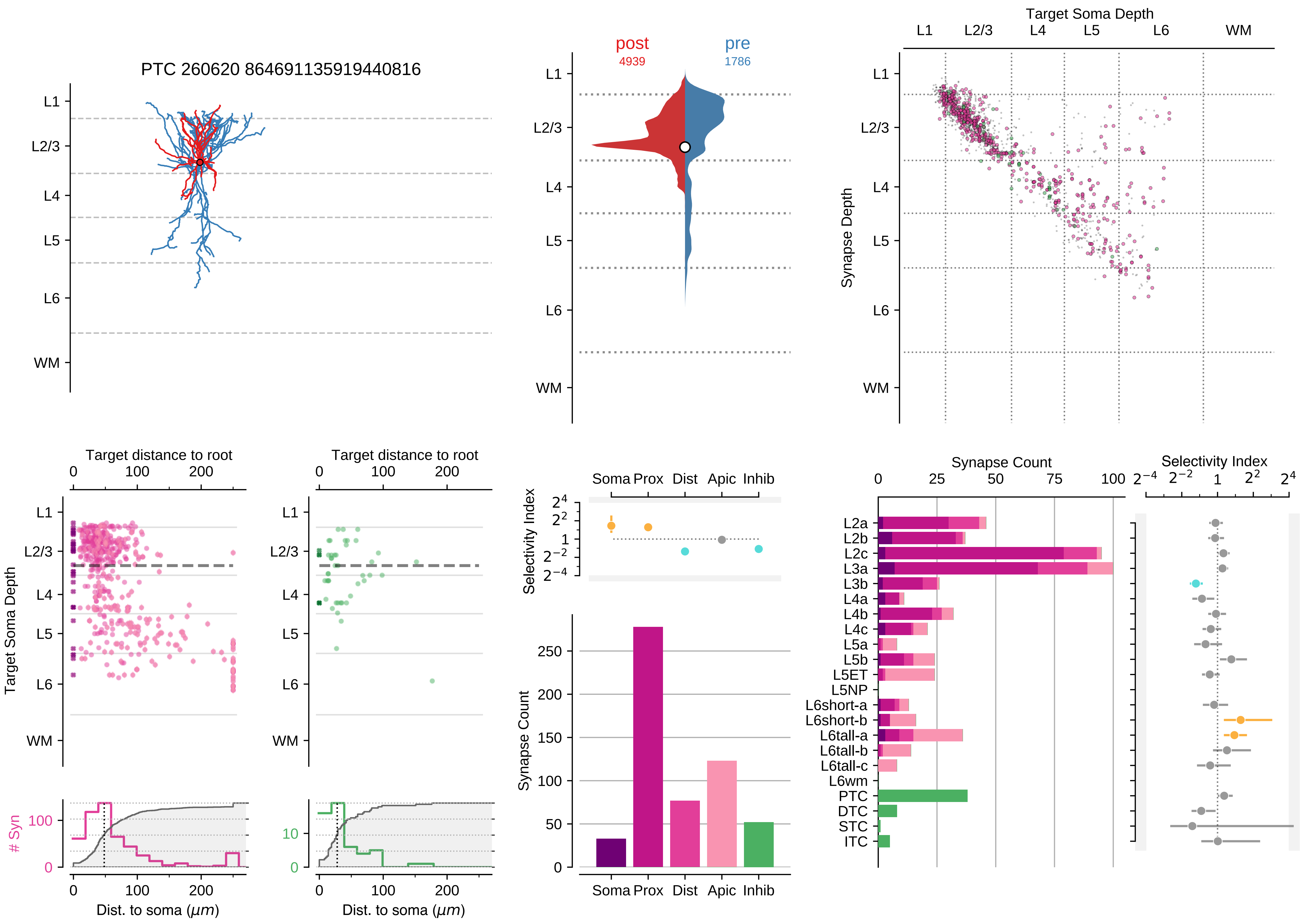

PTC 260620 864691135919440816
post
4939
pre
1786
Target Soma Depth
L1
L2/3
L4
L5
L6
WM
Synapse Depth
Target distance to root
Target Soma Depth
### Syn
Dist. to soma ($\mu m$)
Selectivity Index
Soma
Prox
Dist
Apic
Inhib
Synapse Count
L2a
L2b
L2c
L3a
L3b
L4a
L4b
L4c
L5a
L5b
L5ET
L5NP
L6short-a
L6short-b
L6tall-a
L6tall-b
L6tall-c
L6wm
PTC
DTC
STC
ITC

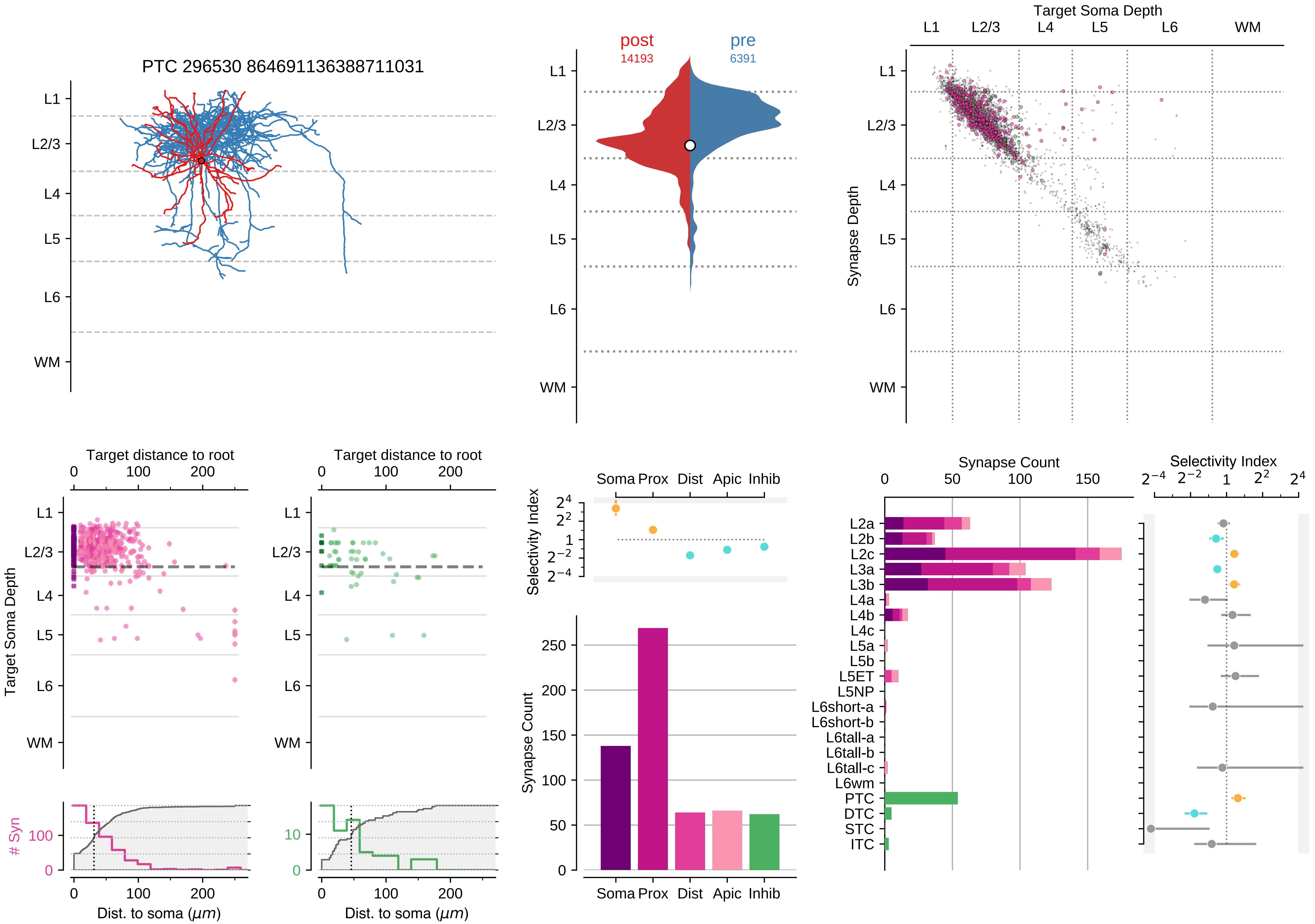
PTC 296530 864691136388711031
L1
L2/3
L4
L5
L6
WM
post
14193
pre
6391
Target Soma Depth
Synapse Depth
Target distance to root
0
100
200
Target Soma Depth
### Syn
Dist. to soma (μm)
Selectivity Index
Soma
Prox
Dist
Apic
Inhib
Synapse Count
0
50
100
150
200
250
Synapse Count
Selectivity Index
L2a
L2b
L2c
L3a
L3b
L4a
L4b
L4c
L5a
L5b
L5ET
L5NP
L6short-a
L6short-b
L6tall-a
L6tall-b
L6tall-c
L6wm
PTC
DTC
STC
ITC

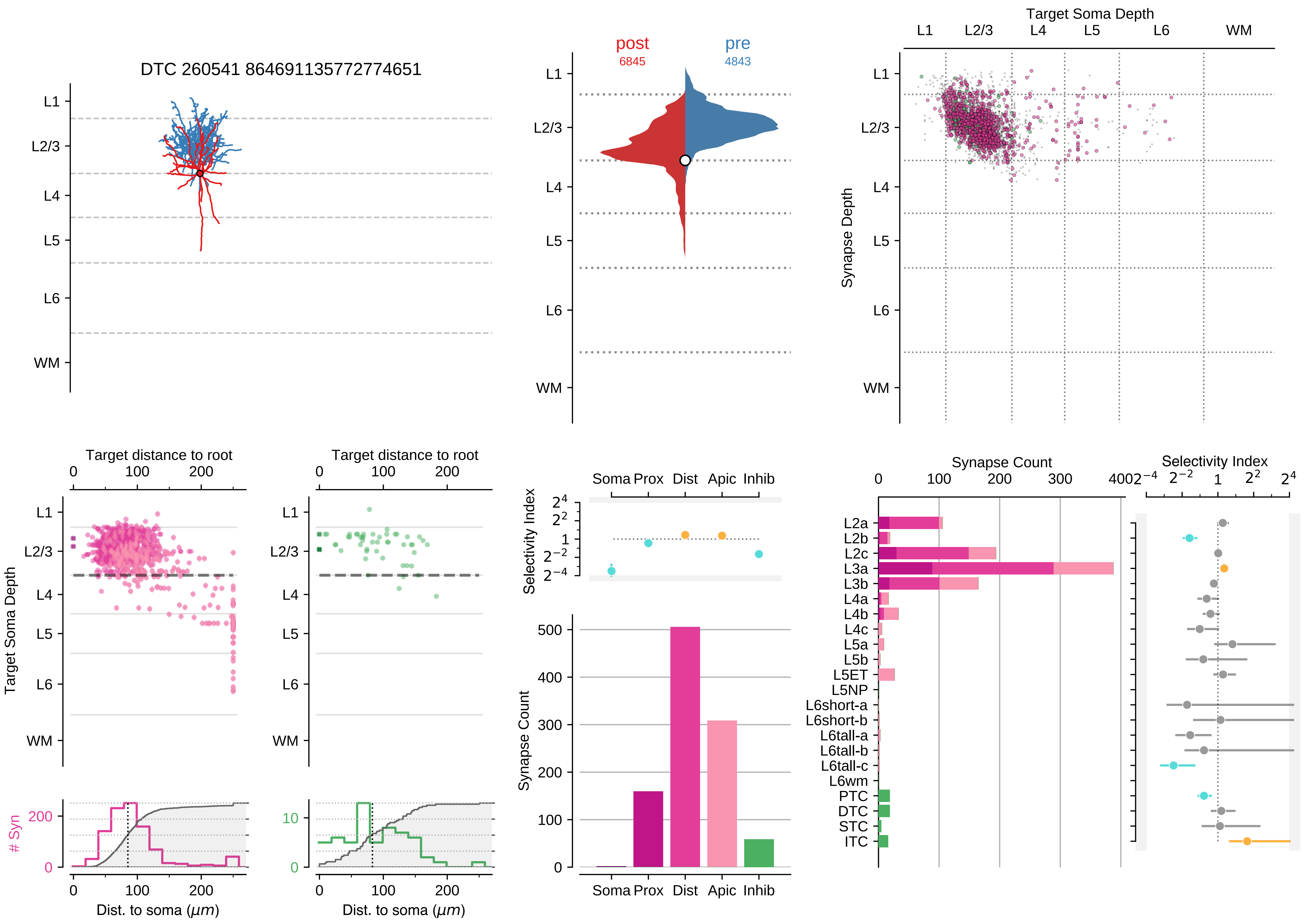

DTC 260541 864691135772774651
post
6845
pre
4843
Target Soma Depth
Synapse Depth
L1
L2/3
L4
L5
L6
WM
Target distance to root
Target Soma Depth
### Syn
Dist. to soma ($\mu m$)
Selectivity Index
Synapse Count
Soma Prox Dist Apic Inhib
L2a
L2b
L2c
L3a
L3b
L4a
L4b
L4c
L5a
L5b
L5ET
L5NP
L6short-a
L6short-b
L6tall-a
L6tall-b
L6tall-c
L6wm
PTC
DTC
STC
ITC

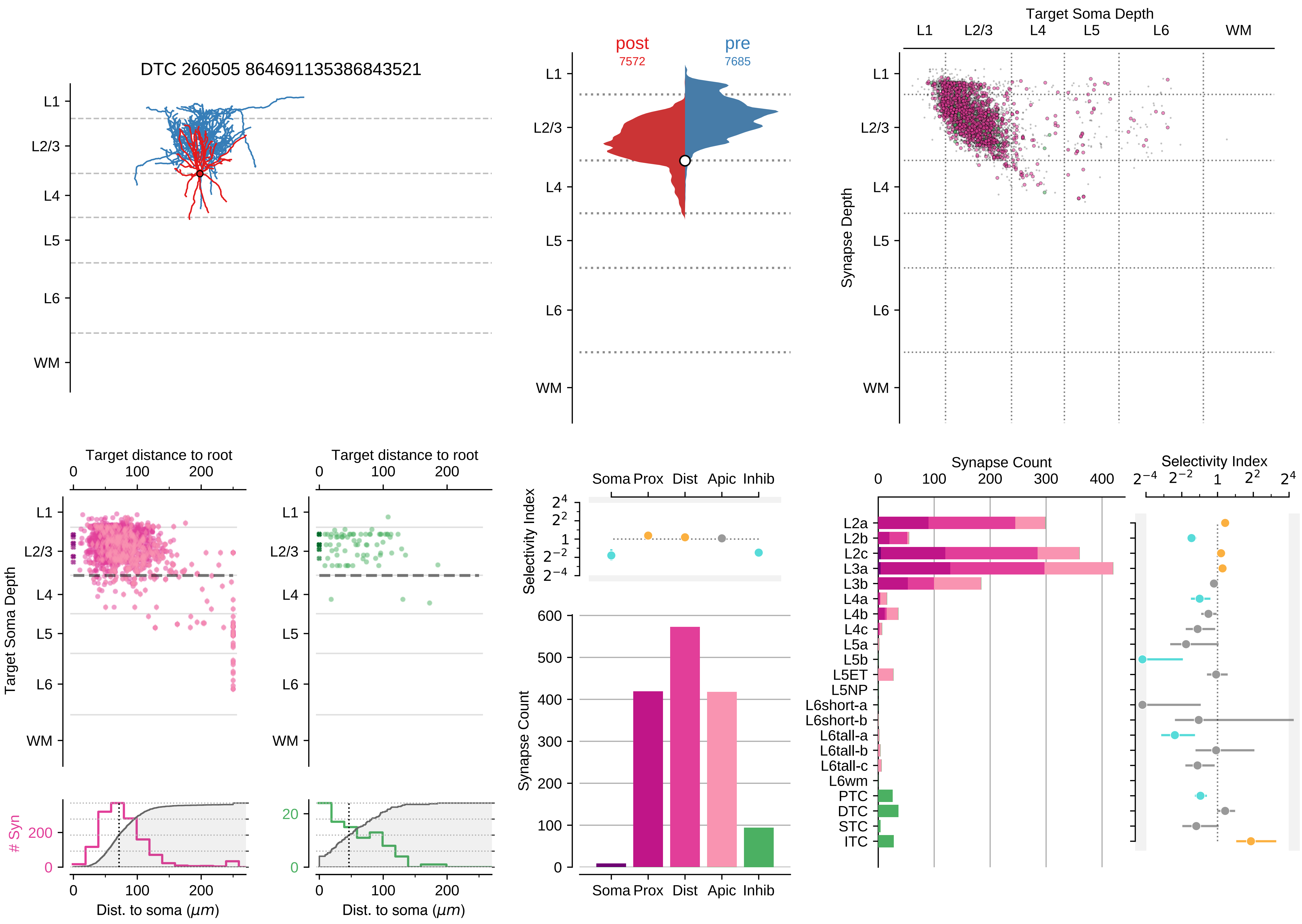

DTC 260505 864691135386843521
post 7572
pre 7685
Target Soma Depth
Synapse Depth
L1
L2/3
L4
L5
L6
WM
Target distance to root
Target Soma Depth
### Syn
Dist. to soma (μm)
Selectivity Index
Synapse Count
Soma Prox Dist Apic Inhib
L2a
L2b
L2c
L3a
L3b
L4a
L4b
L4c
L5a
L5b
L5ET
L5NP
L6short-a
L6short-b
L6tall-a
L6tall-b
L6tall-c
L6wm
PTC
DTC
STC
ITC

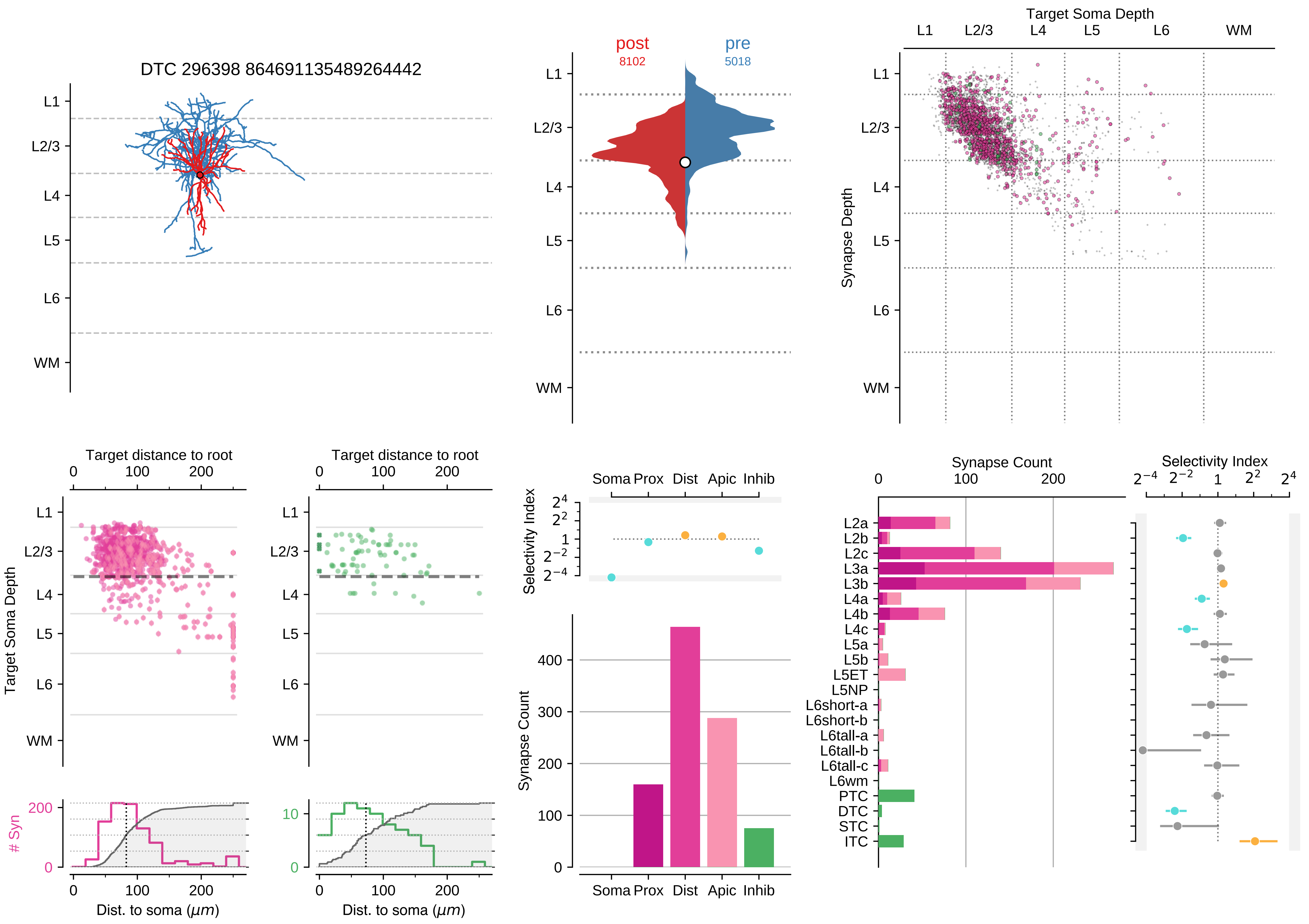
DTC 296398 864691135489264442
L1
L2/3
L4
L5
L6
WM
post
8102
pre
5018
Target Soma Depth
Synapse Depth
Target distance to root
Target Soma Depth
### Syn
Dist. to soma ($\mu m$)
Selectivity Index
Soma Prox Dist Apic Inhib
Synapse Count
L2a
L2b
L2c
L3a
L3b
L4a
L4b
L4c
L5a
L5b
L5ET
L5NP
L6short-a
L6short-b
L6tall-a
L6tall-b
L6tall-c
L6wm
PTC
DTC
STC
ITC

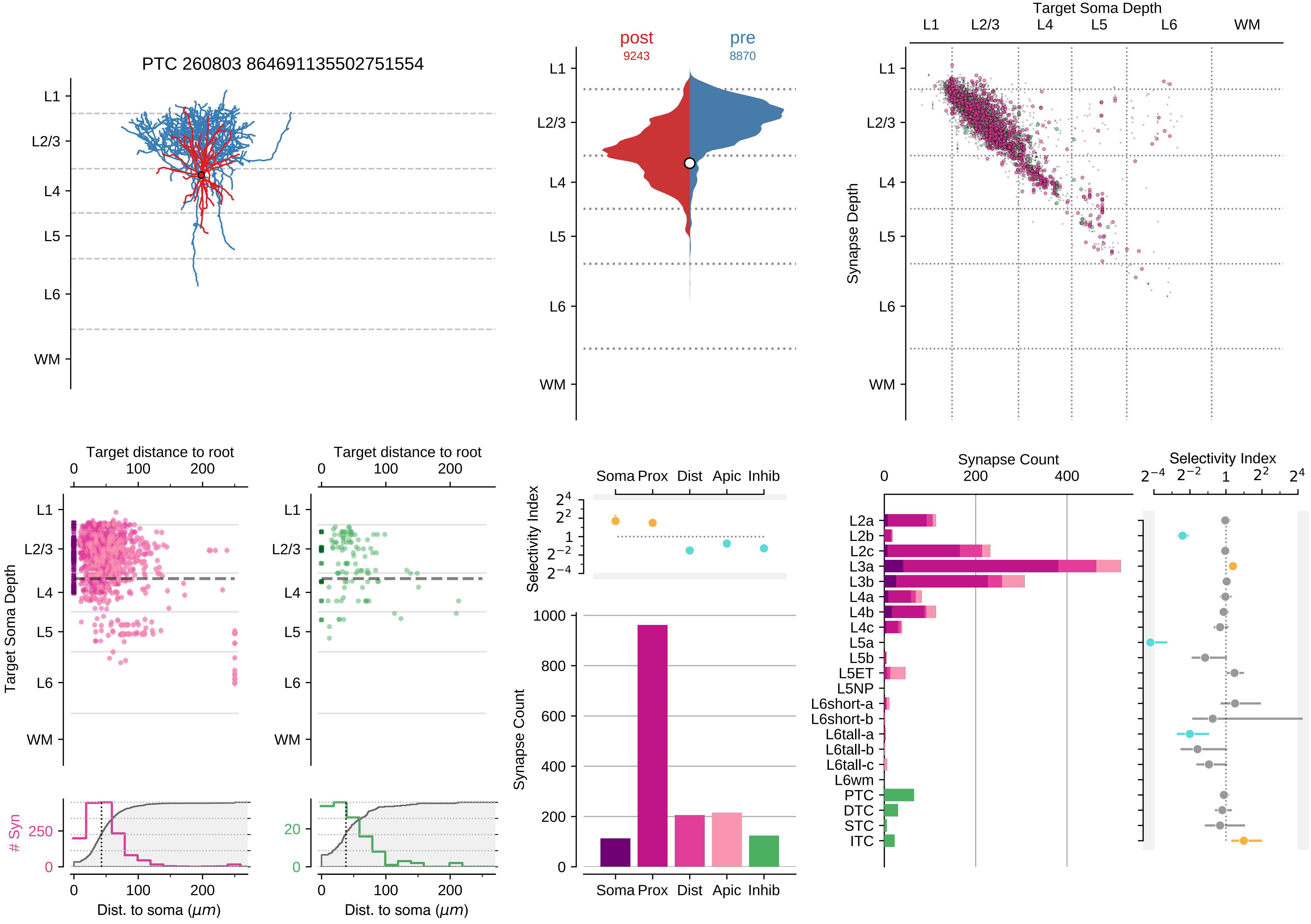

PTC 260803 864691135502751554
L1
L2/3
L4
L5
L6
WM
post
9243
pre
8870
Target Soma Depth
Synapse Depth
Target distance to root
0
100
200
Target Soma Depth
Dist. to soma (μm)
### Syn
250
0
20
Selectivity Index
Soma
Prox
Dist
Apic
Inhib
Synapse Count
1000
800
600
400
200
L2a
L2b
L2c
L3a
L3b
L4a
L4b
L4c
L5a
L5b
L5ET
L5NP
L6short-a
L6short-b
L6tall-a
L6tall-b
L6tall-c
L6wm
PTC
DTC
STC
ITC

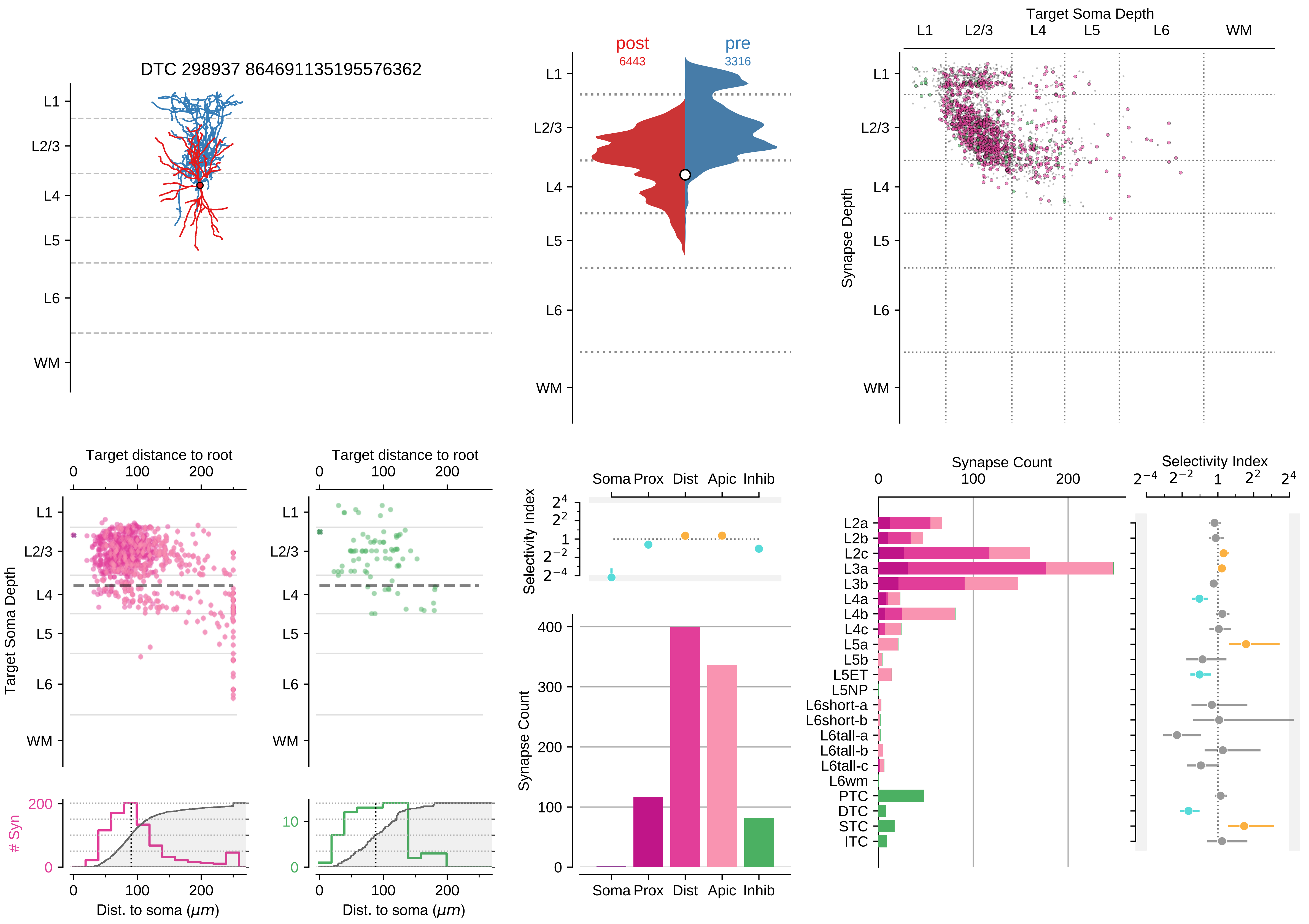

DTC 298937 864691135195576362
L1
L2/3
L4
L5
L6
WM
post
6443
pre
3316
Target Soma Depth
Synapse Depth
Target distance to root
Dist. to soma ($\mu m$)
### Syn
Selectivity Index
Soma Prox Dist Apic Inhib
Synapse Count
L2a
L2b
L2c
L3a
L3b
L4a
L4b
L4c
L5a
L5b
L5ET
L5NP
L6short-a
L6short-b
L6tall-a
L6tall-b
L6tall-c
L6wm
PTC
DTC
STC
ITC

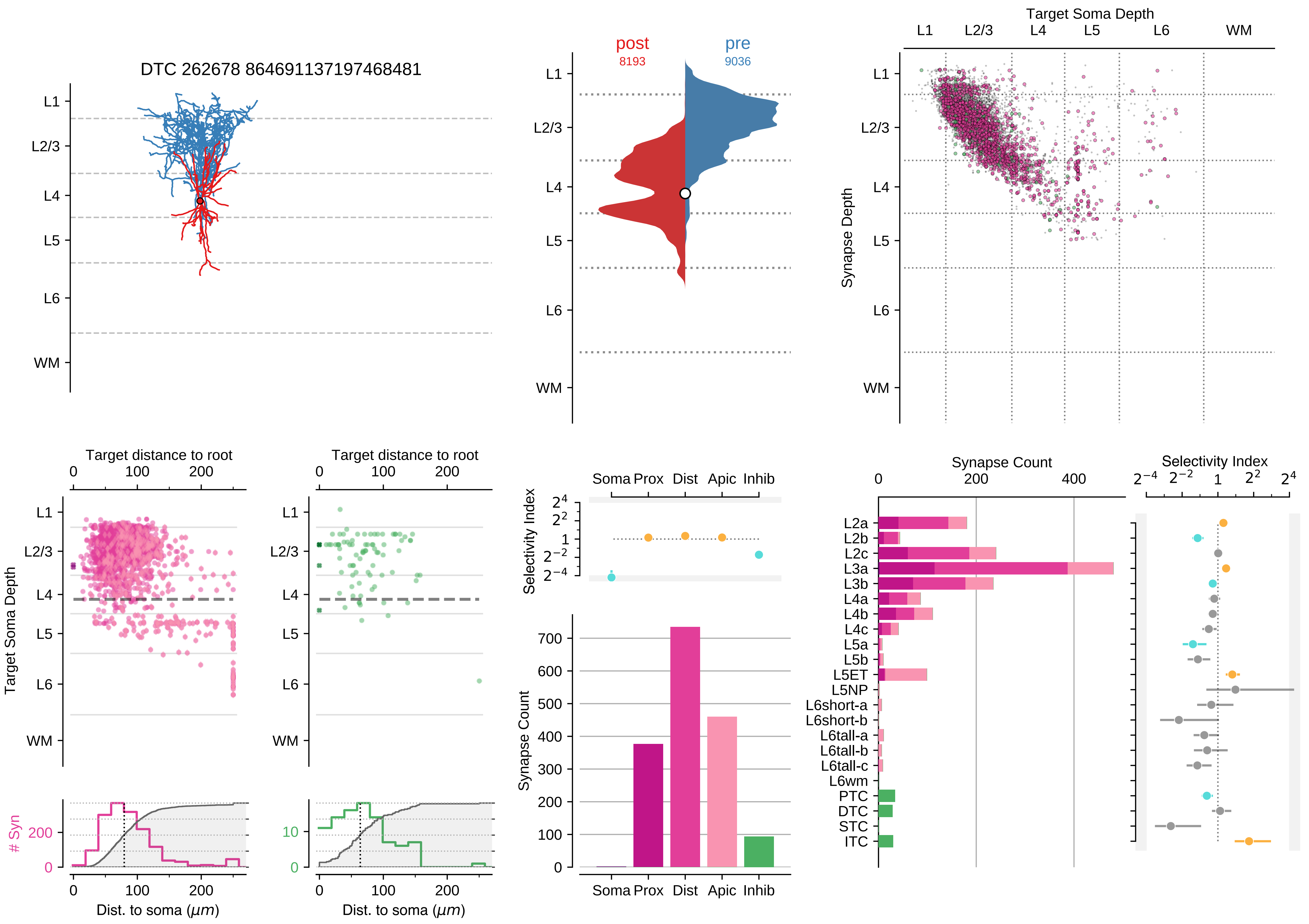
DTC 262678 864691137197468481
L1
L2/3
L4
L5
L6
WM
post
8193
pre
9036
Target Soma Depth
Synapse Depth
Target distance to root
Target Soma Depth
### Syn
Dist. to soma (μm)
Selectivity Index
Soma Prox Dist Apic Inhib
Synapse Count
L2a
L2b
L2c
L3a
L3b
L4a
L4b
L4c
L5a
L5b
L5ET
L5NP
L6short-a
L6short-b
L6tall-a
L6tall-b
L6tall-c
L6wm
PTC
DTC
STC
ITC

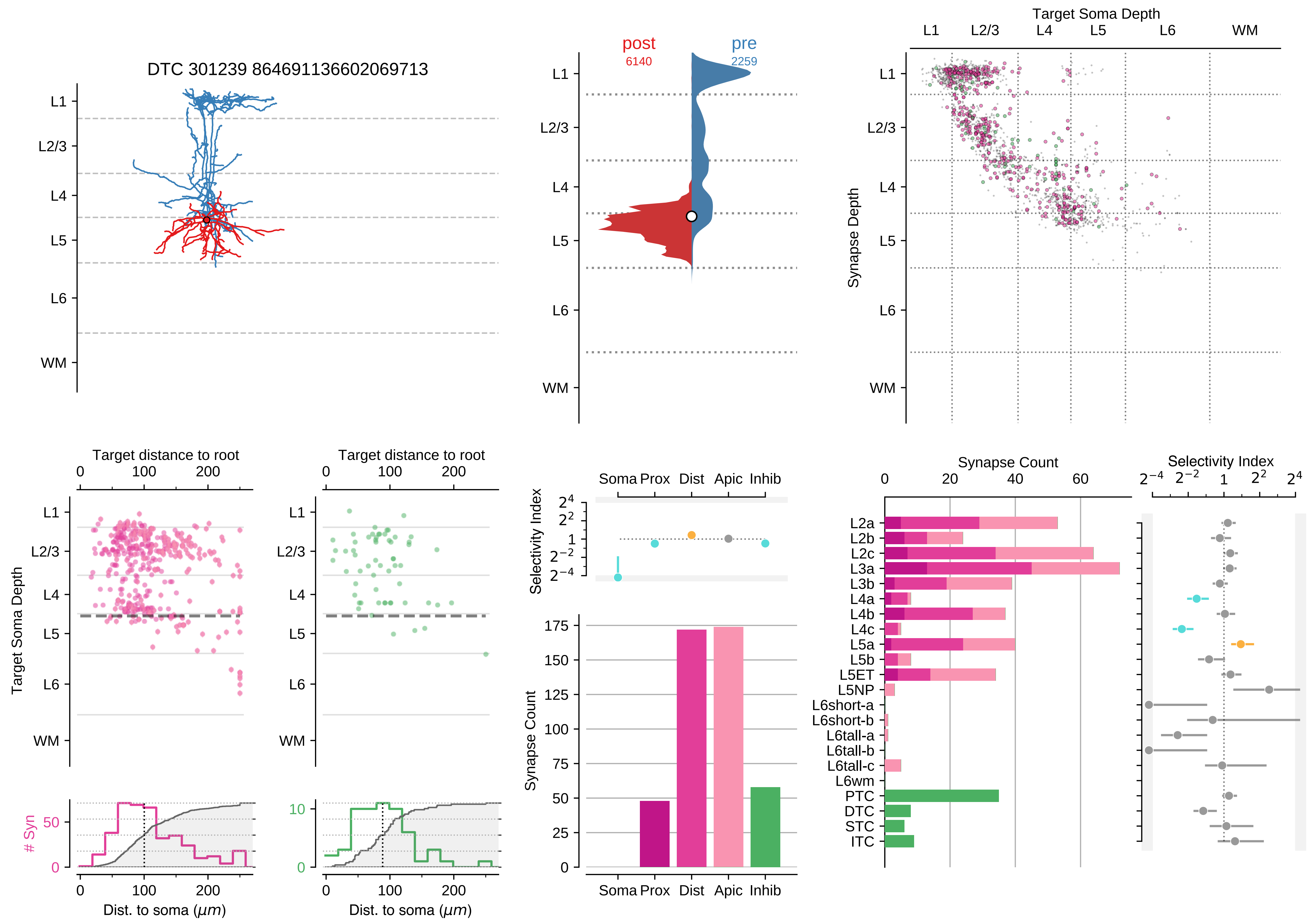

DTC 301239 864691136602069713
L1
L2/3
L4
L5
L6
WM
post
6140
pre
2259
Target Soma Depth
Synapse Depth
Target distance to root
Target Soma Depth
### Syn
Dist. to soma (μm)
Selectivity Index
Soma Prox Dist Apic Inhib
Synapse Count
Selectivity Index
L2a
L2b
L2c
L3a
L3b
L4a
L4b
L4c
L5a
L5b
L5ET
L5NP
L6short-a
L6short-b
L6tall-a
L6tall-b
L6tall-c
L6wm
PTC
DTC
STC
ITC

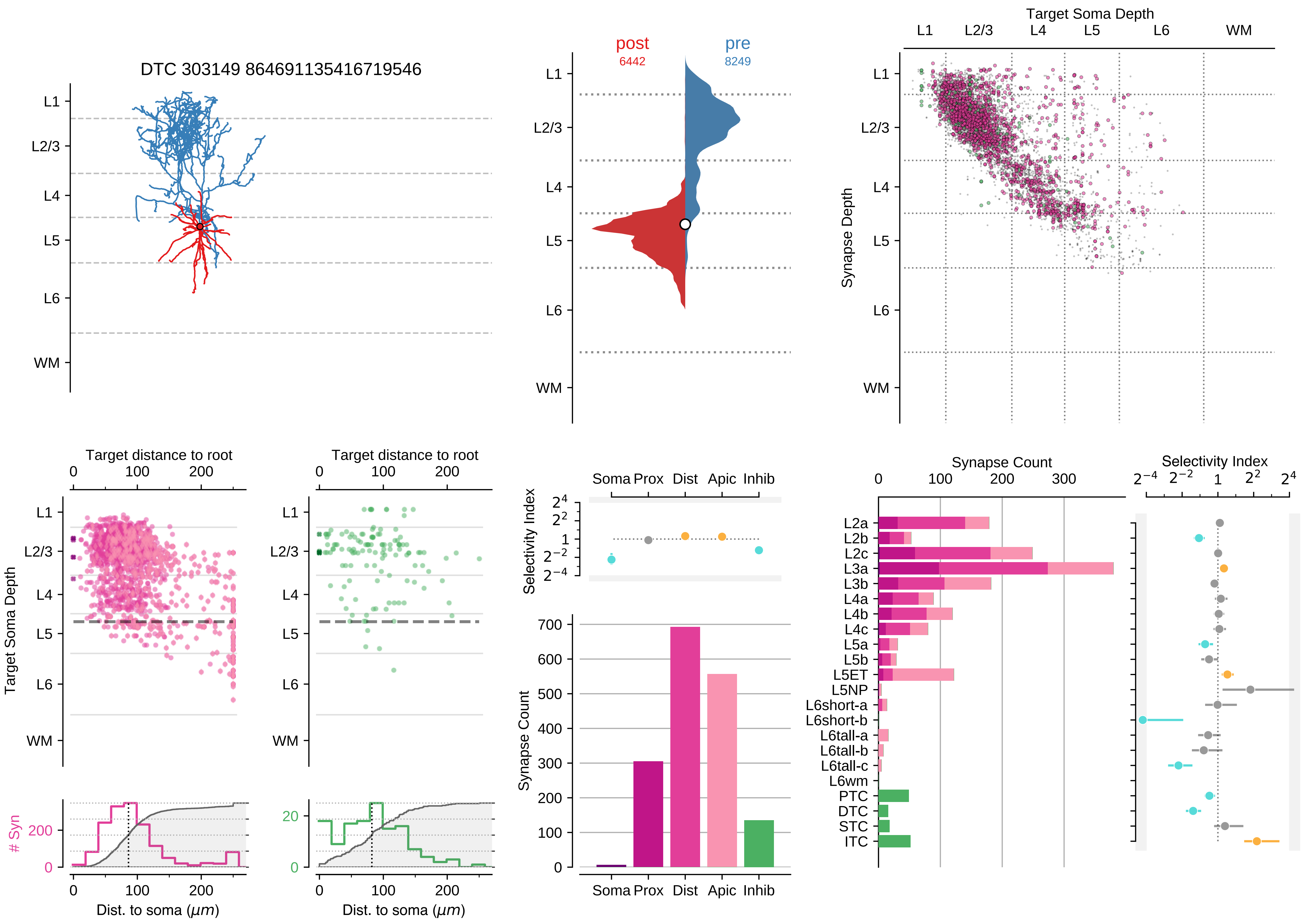

DTC 303149 864691135416719546
L1
L2/3
L4
L5
L6
WM
post
6442
pre
8249
Target Soma Depth
Synapse Depth
Target distance to root
Target Soma Depth
### Syn
Dist. to soma (μm)
Selectivity Index
Synapse Count
Soma Prox Dist Apic Inhib
L2a
L2b
L2c
L3a
L3b
L4a
L4b
L4c
L5a
L5b
L5ET
L5NP
L6short-a
L6short-b
L6tall-a
L6tall-b
L6tall-c
L6wm
PTC
DTC
STC
ITC

### Motif Group 4

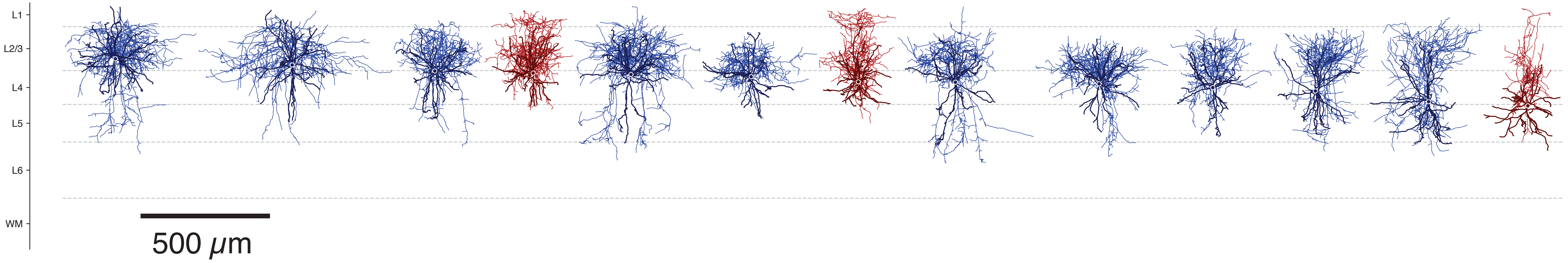

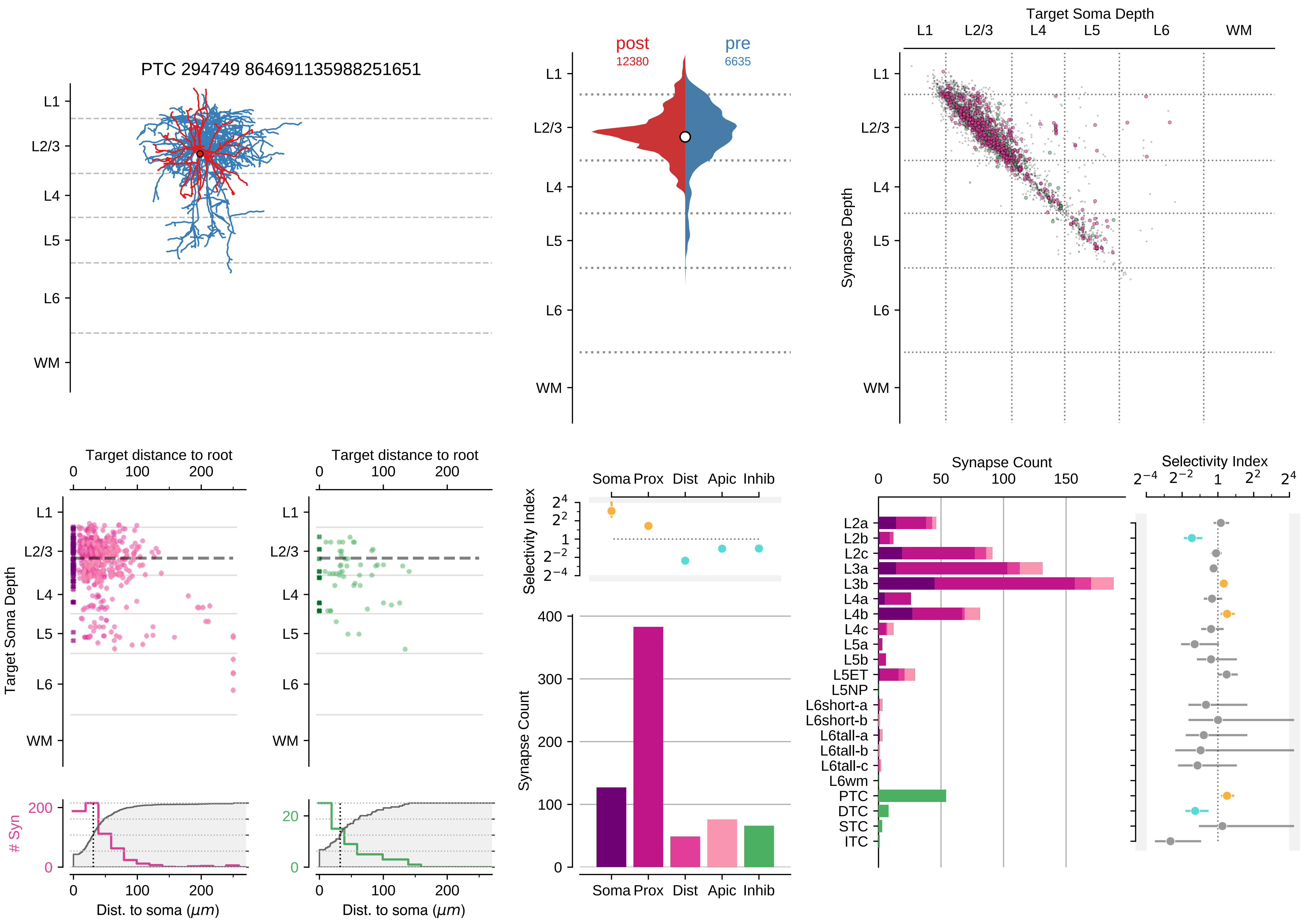

PTC 294749 864691135988251651
L1
L2/3
L4
L5
L6
WM
post
12380
pre
6635
Target Soma Depth
Synapse Depth
Target distance to root
0
100
200
Target Soma Depth
### Syn
Dist. to soma (μm)
Selectivity Index
Soma
Prox
Dist
Apic
Inhib
Synapse Count
L2a
L2b
L2c
L3a
L3b
L4a
L4b
L4c
L5a
L5b
L5ET
L5NP
L6short-a
L6short-b
L6tall-a
L6tall-b
L6tall-c
L6wm
PTC
DTC
STC
ITC

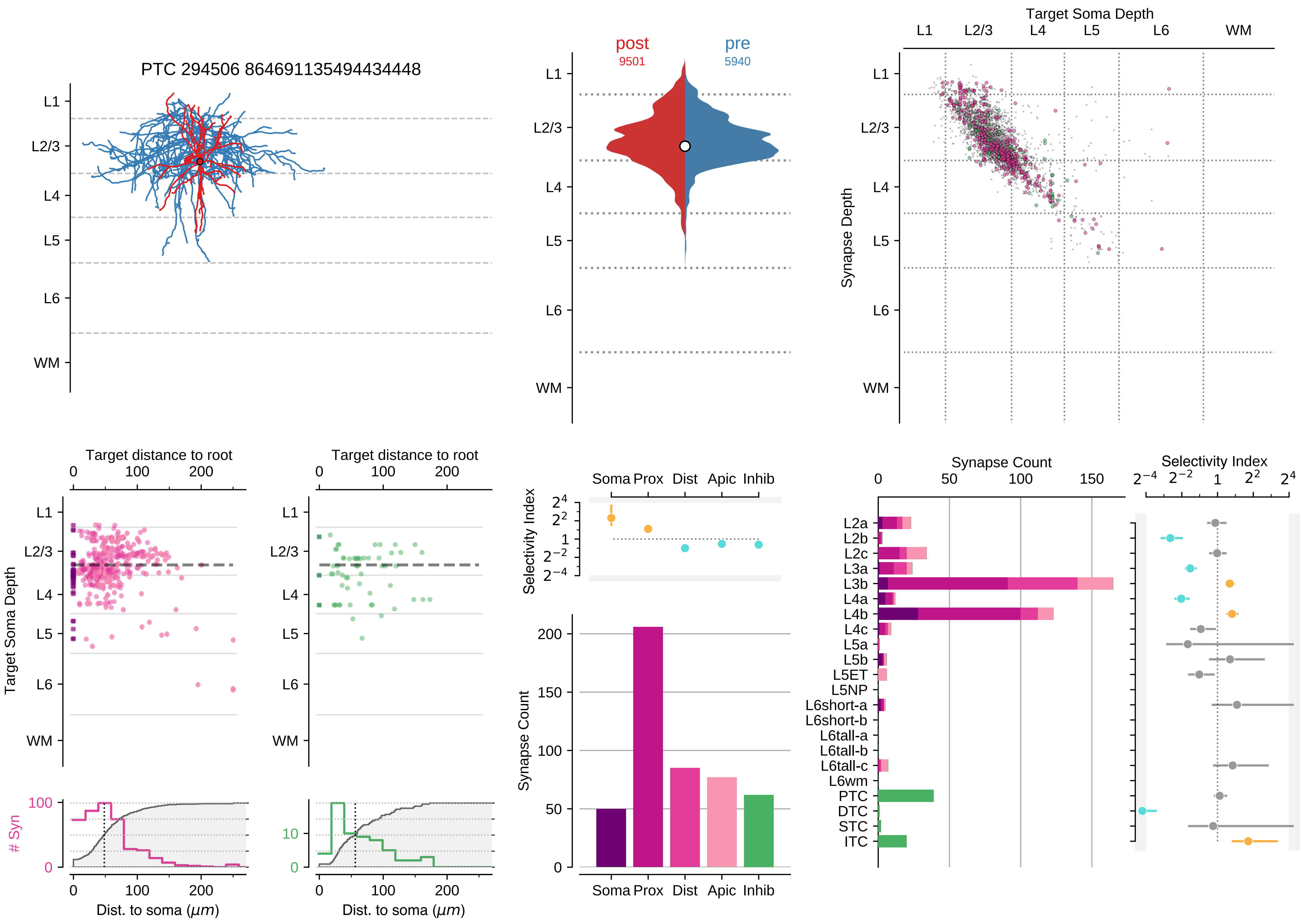

PTC 294506 864691135494434448
post
9501
pre
5940
Target Soma Depth
Synapse Depth
L1
L2/3
L4
L5
L6
WM
Target distance to root
Target Soma Depth
### Syn
Dist. to soma ($\mu m$)
Selectivity Index
Soma
Prox
Dist
Apic
Inhib
Synapse Count
L2a
L2b
L2c
L3a
L3b
L4a
L4b
L4c
L5a
L5b
L5ET
L5NP
L6short-a
L6short-b
L6tall-a
L6tall-b
L6tall-c
L6wm
PTC
DTC
STC
ITC

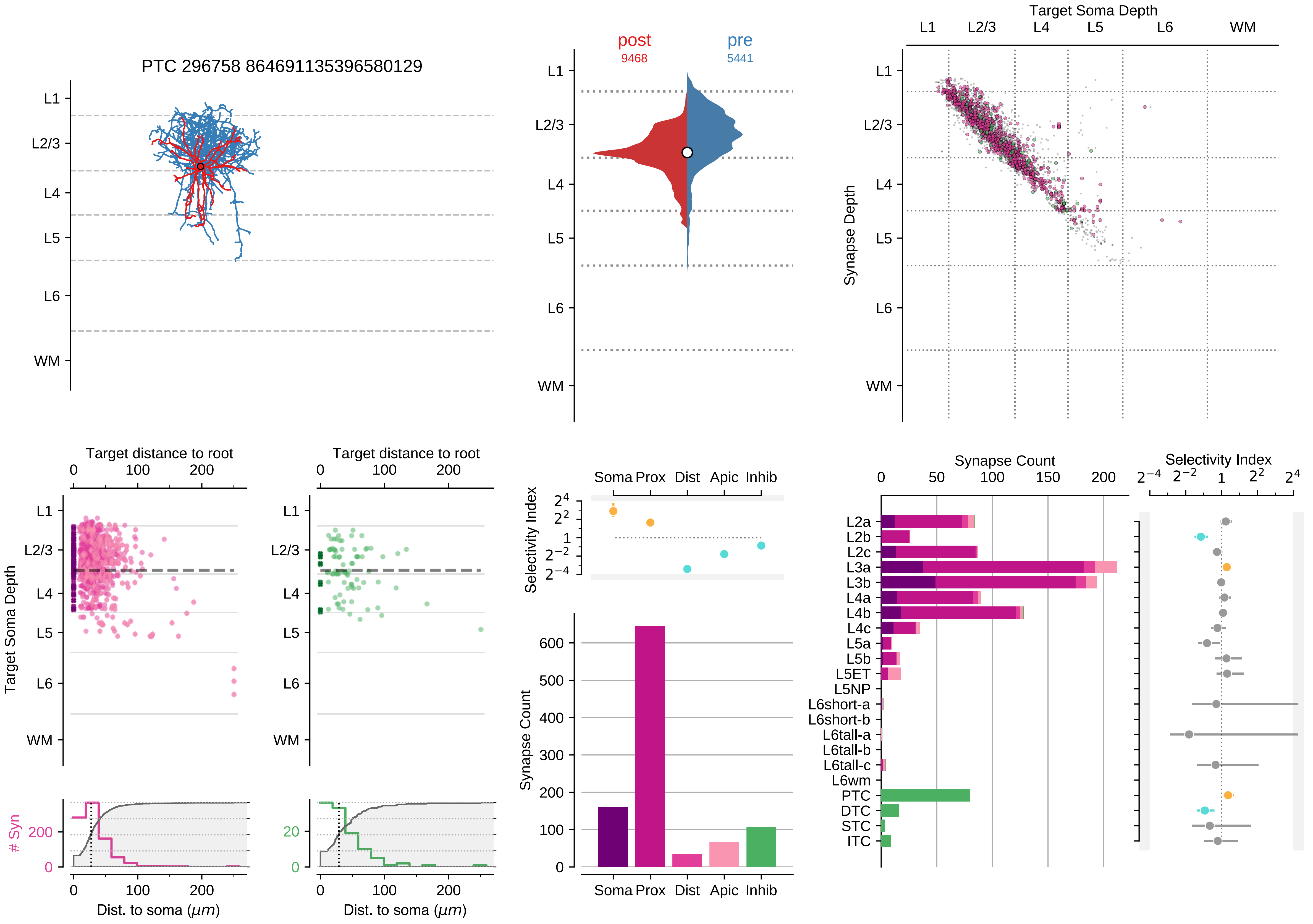

PTC 296758 864691135396580129
L1
L2/3
L4
L5
L6
WM
post
9468
pre
5441
Target Soma Depth
Synapse Depth
Target distance to root
Target Soma Depth
### Syn
Dist. to soma ($\mu m$)
Selectivity Index
Soma Prox Dist Apic Inhib
Synapse Count
L2a
L2b
L2c
L3a
L3b
L4a
L4b
L4c
L5a
L5b
L5ET
L5NP
L6short-a
L6short-b
L6tall-a
L6tall-b
L6tall-c
L6wm
PTC
DTC
STC
ITC

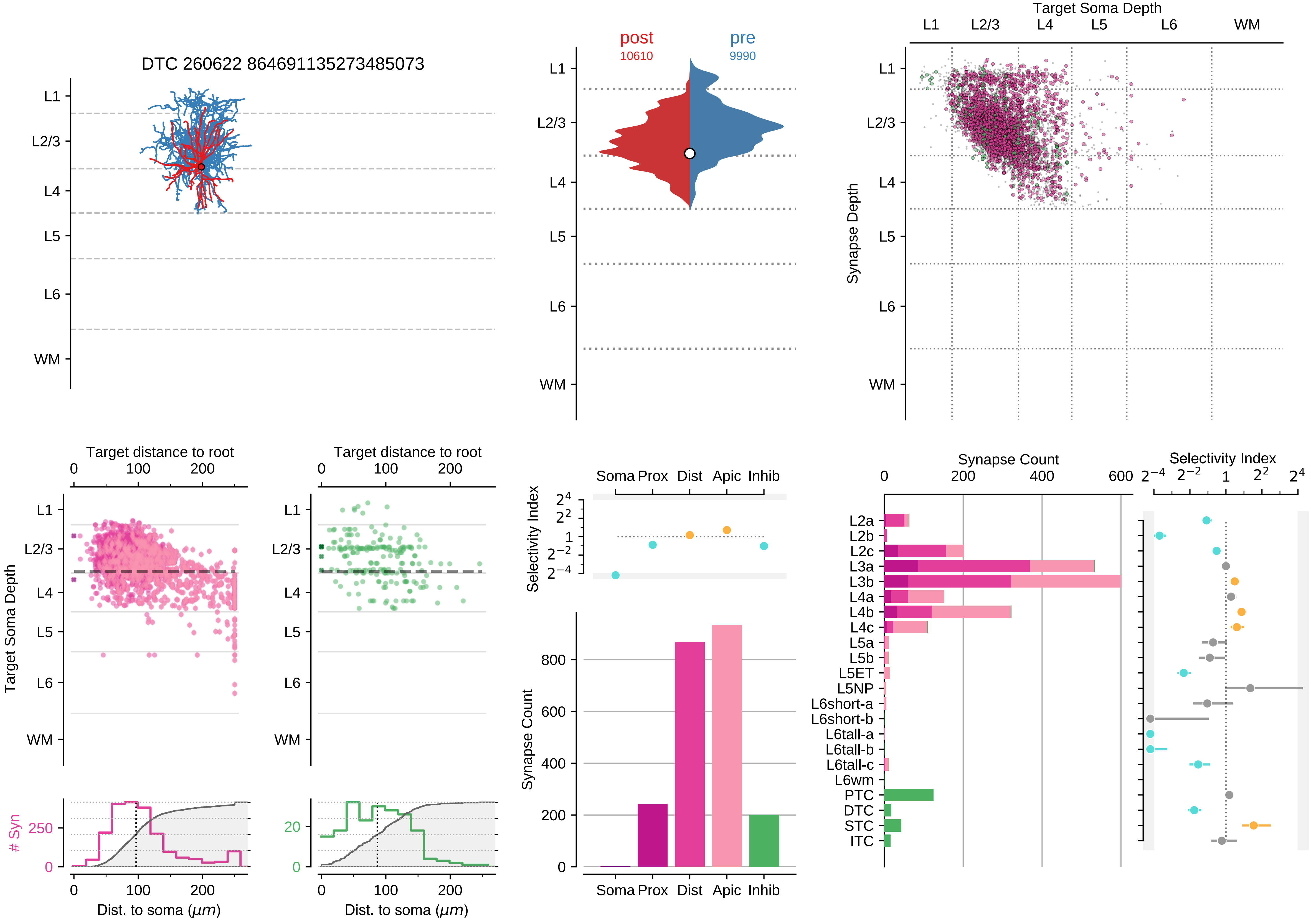
DTC 260622 864691135273485073
post
10610
pre
9990
Target Soma Depth
Synapse Depth
L1
L2/3
L4
L5
L6
WM
Target distance to root
Target Soma Depth
### Syn
Dist. to soma (μm)
Selectivity Index
Synapse Count
Soma Prox Dist Apic Inhib
L2a
L2b
L2c
L3a
L3b
L4a
L4b
L4c
L5a
L5b
L5ET
L5NP
L6short-a
L6short-b
L6tall-a
L6tall-b
L6tall-c
L6wm
PTC
DTC
STC
ITC

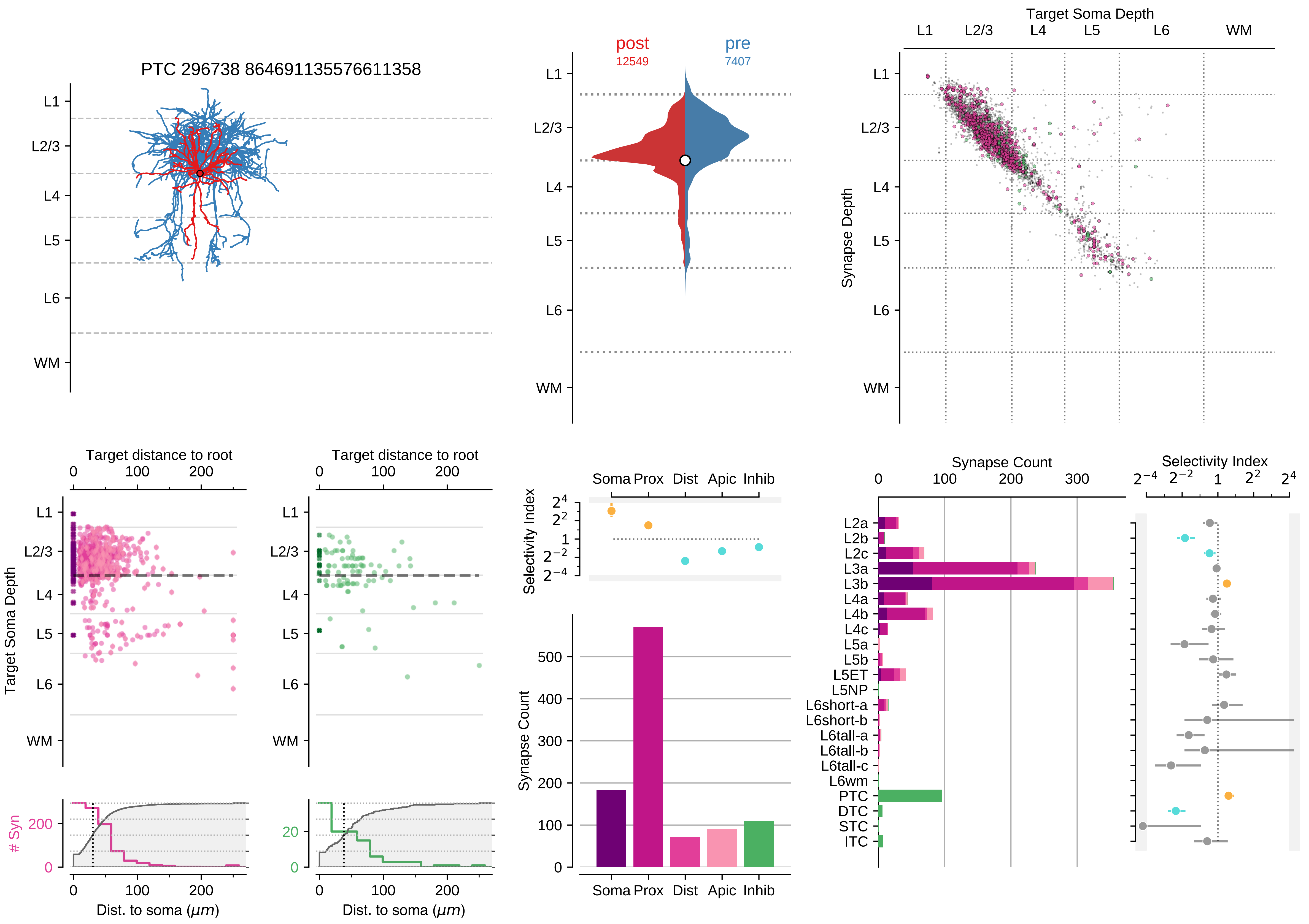

PTC 296738 864691135576611358
L1
L2/3
L4
L5
L6
WM
post
12549
pre
7407
Target Soma Depth
Synapse Depth
Target distance to root
Target Soma Depth
### Syn
Dist. to soma (μm)
Selectivity Index
Soma
Prox
Dist
Apic
Inhib
Synapse Count
L2a
L2b
L2c
L3a
L3b
L4a
L4b
L4c
L5a
L5b
L5ET
L5NP
L6short-a
L6short-b
L6tall-a
L6tall-b
L6tall-c
L6wm
PTC
DTC
STC
ITC

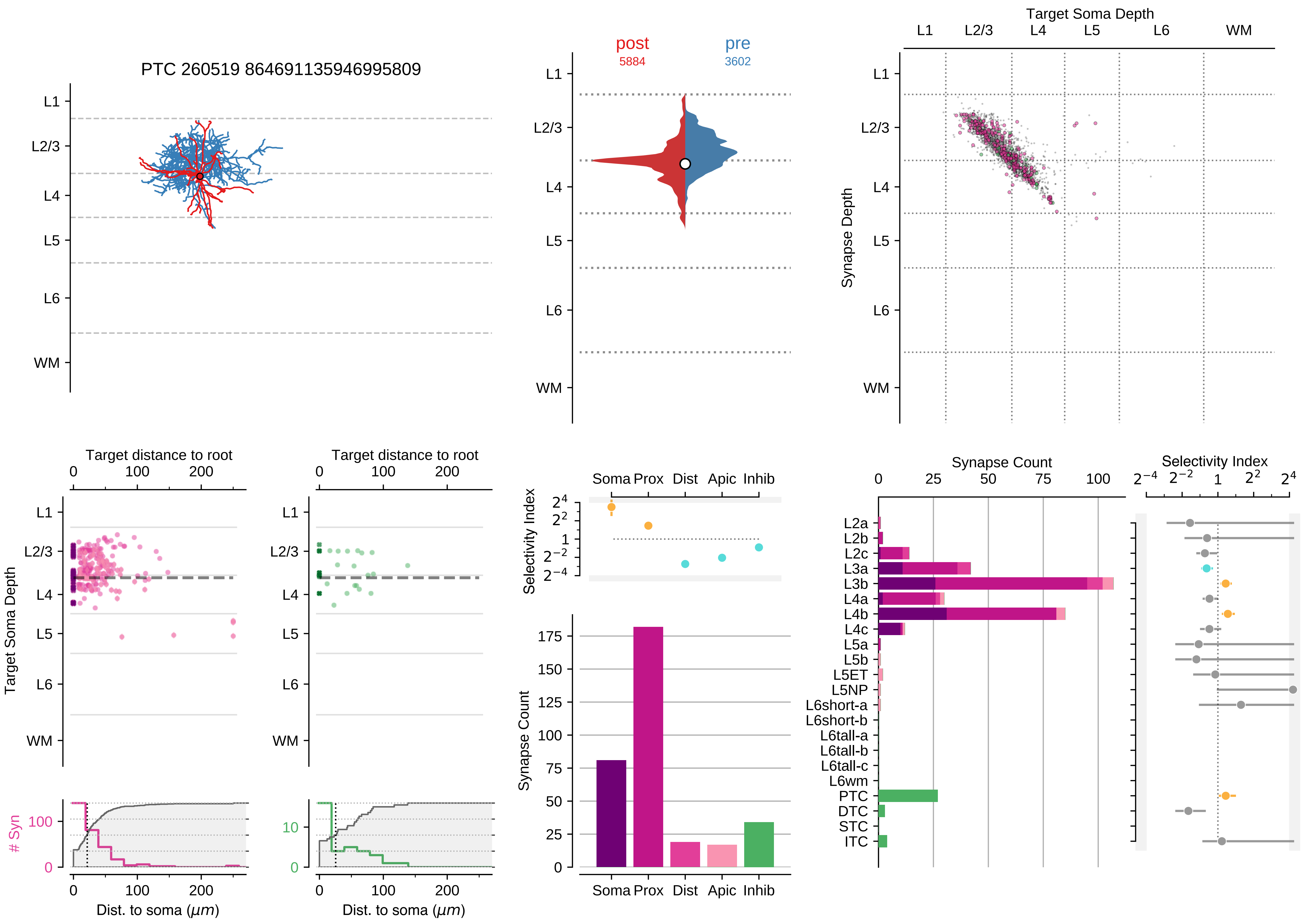

PTC 260519 864691135946995809
L1
L2/3
L4
L5
L6
WM
post
5884
pre
3602
Target Soma Depth
Synapse Depth
Target distance to root
Target Soma Depth
Dist. to soma (μm)
### Syn
Selectivity Index
Soma Prox Dist Apic Inhib
Synapse Count
L2a
L2b
L2c
L3a
L3b
L4a
L4b
L4c
L5a
L5b
L5ET
L5NP
L6short-a
L6short-b
L6tall-a
L6tall-b
L6tall-c
L6wm
PTC
DTC
STC
ITC

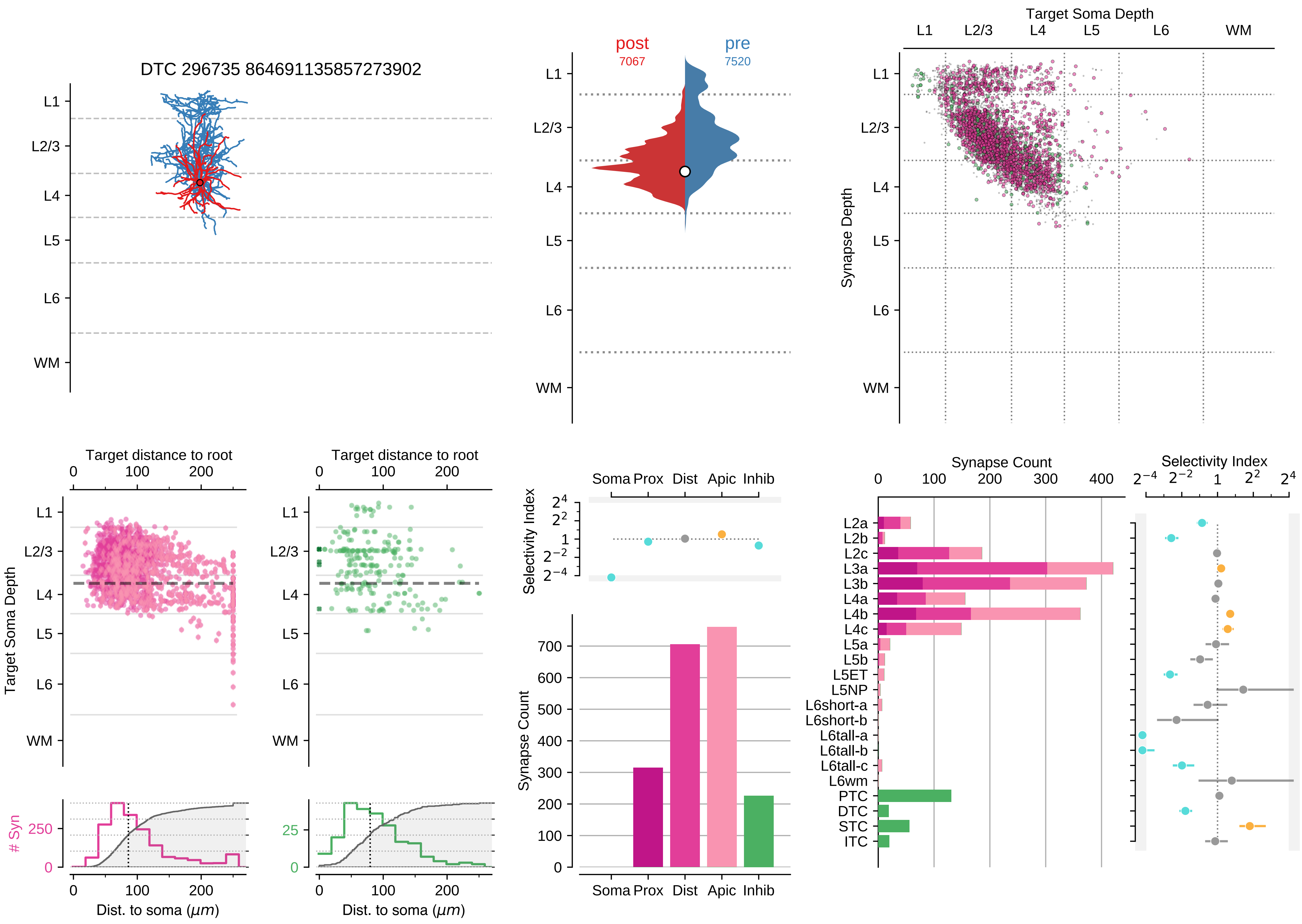

DTC 296735 864691135857273902
L1
L2/3
L4
L5
L6
WM
post
7067
pre
7520
Target Soma Depth
Synapse Depth
Target distance to root
Target Soma Depth
### Syn
Dist. to soma (μm)
Selectivity Index
Soma
Prox
Dist
Apic
Inhib
Synapse Count
L2a
L2b
L2c
L3a
L3b
L4a
L4b
L4c
L5a
L5b
L5ET
L5NP
L6short-a
L6short-b
L6tall-a
L6tall-b
L6tall-c
L6wm
PTC
DTC
STC
ITC

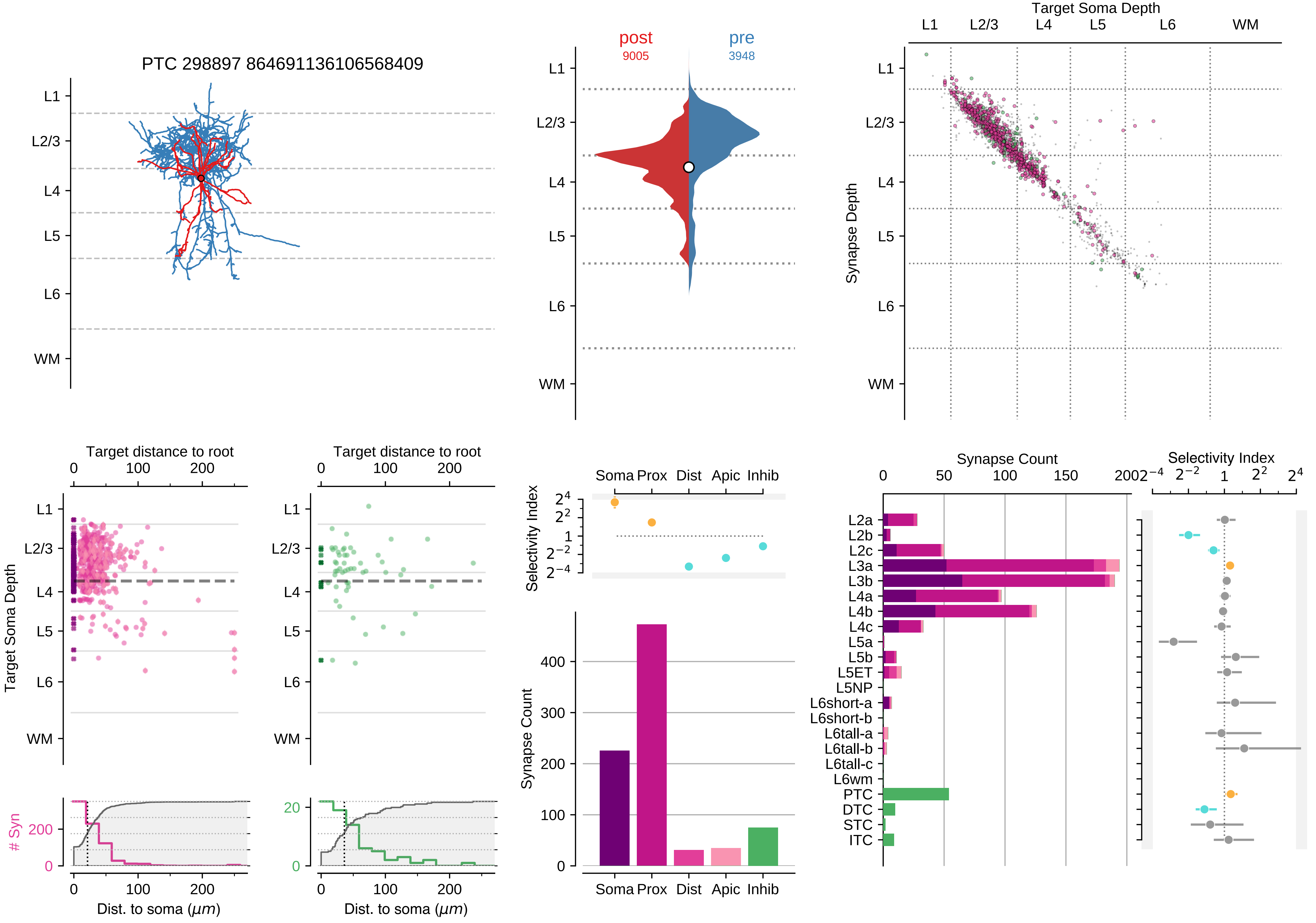

PTC 298897 864691136106568409
L1
L2/3
L4
L5
L6
WM
post
9005
pre
3948
Target Soma Depth
Synapse Depth
Target distance to root
Target Soma Depth
### Syn
Dist. to soma (μm)
Selectivity Index
Soma Prox Dist Apic Inhib
Synapse Count
L2a
L2b
L2c
L3a
L3b
L4a
L4b
L4c
L5a
L5b
L5ET
L5NP
L6short-a
L6short-b
L6tall-a
L6tall-b
L6tall-c
L6wm
PTC
DTC
STC
ITC

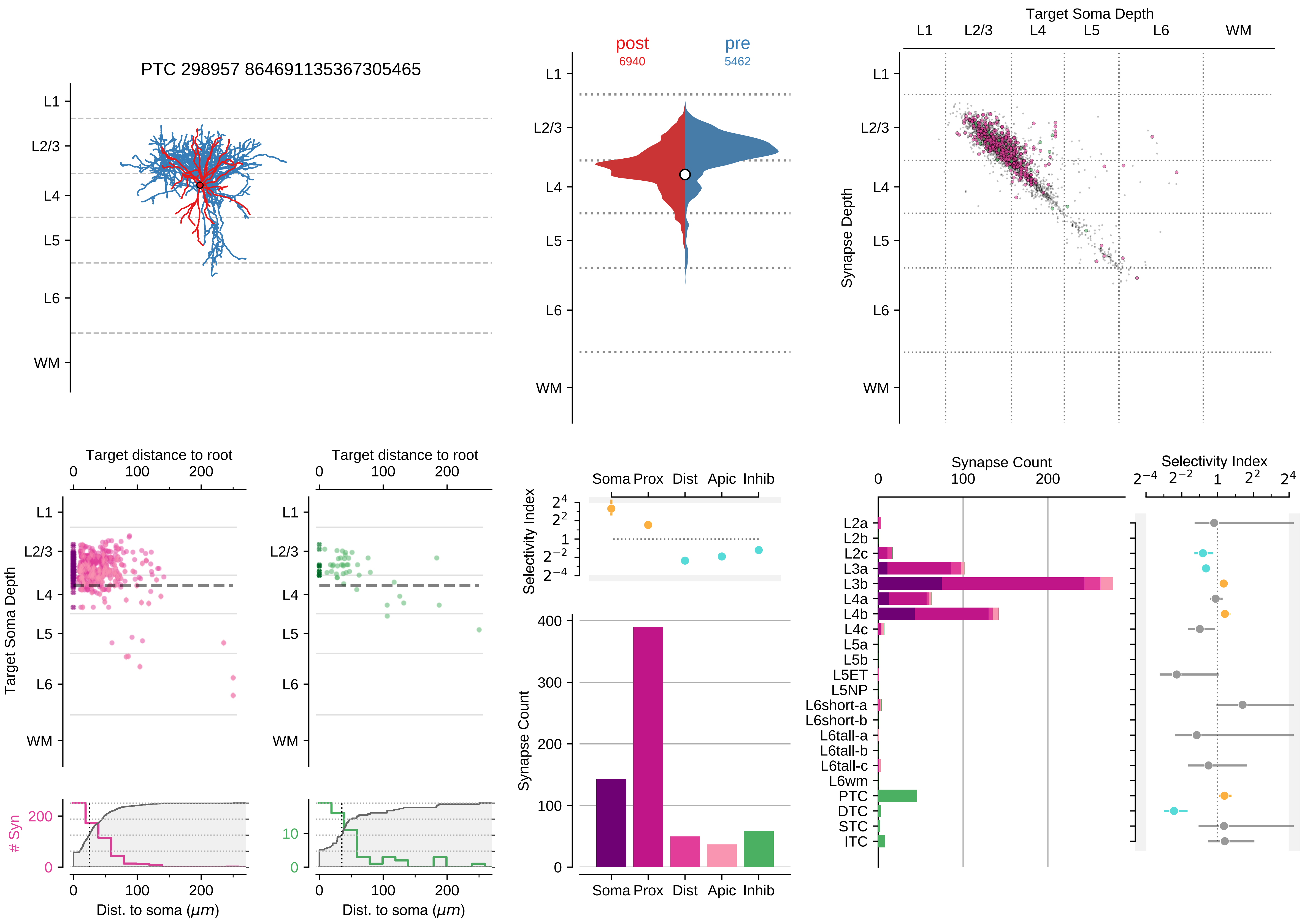

PTC 298957 864691135367305465
post
6940
pre
5462
Target Soma Depth
Synapse Depth
L1
L2/3
L4
L5
L6
WM
Target distance to root
Target Soma Depth
### Syn
Dist. to soma ($\mu m$)
Selectivity Index
Soma Prox Dist Apic Inhib
Synapse Count
L2a
L2b
L2c
L3a
L3b
L4a
L4b
L4c
L5a
L5b
L5ET
L5NP
L6short-a
L6short-b
L6tall-a
L6tall-b
L6tall-c
L6wm
PTC
DTC
STC
ITC

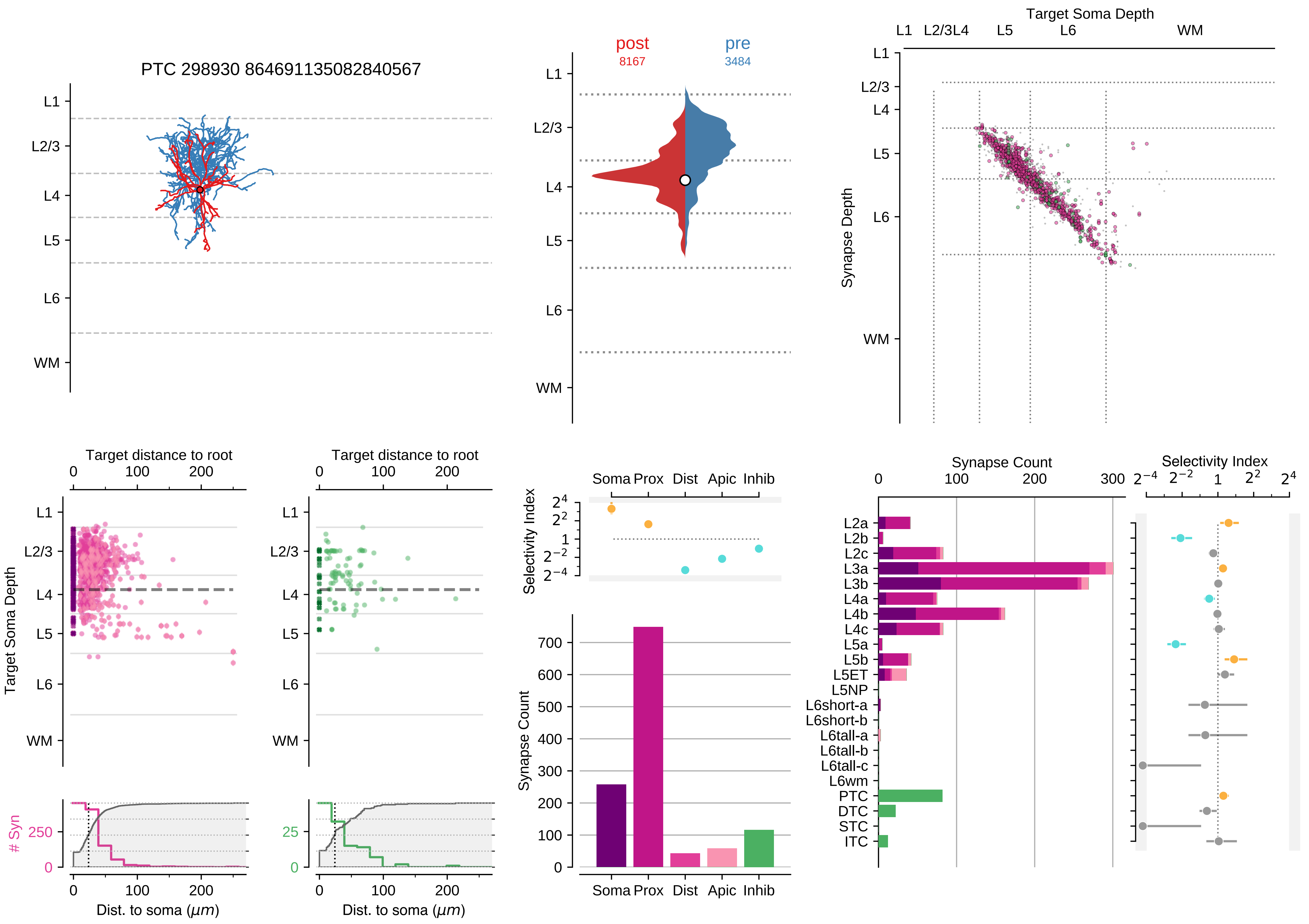

PTC 298930 864691135082840567
L1
L2/3
L4
L5
L6
WM
post
8167
pre
3484
Target Soma Depth
Synapse Depth
Target distance to root
Target Soma Depth
### Syn
Dist. to soma (μm)
Selectivity Index
Soma Prox Dist Apic Inhib
Synapse Count
Synapse Count
Selectivity Index
L2a
L2b
L2c
L3a
L3b
L4a
L4b
L4c
L5a
L5b
L5ET
L5NP
L6short-a
L6short-b
L6tall-a
L6tall-b
L6tall-c
L6wm
PTC
DTC
STC
ITC

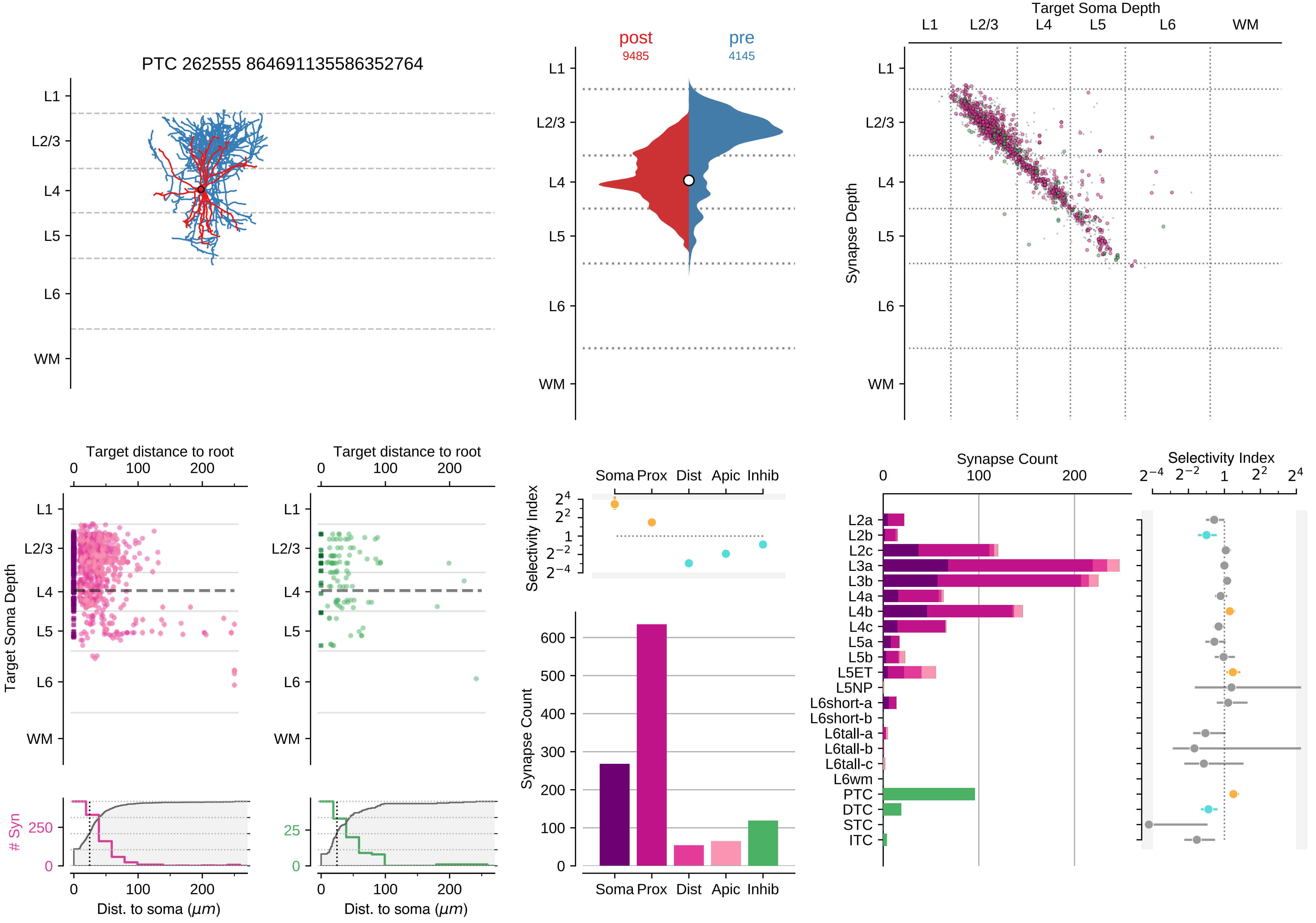

PTC 262555 864691135586352764
L1
L2/3
L4
L5
L6
WM
post
9485
pre
4145
Target Soma Depth
Synapse Depth
Target distance to root
Target Soma Depth
### Syn
Dist. to soma (μm)
Selectivity Index
Soma Prox Dist Apic Inhib
Synapse Count
L2a
L2b
L2c
L3a
L3b
L4a
L4b
L4c
L5a
L5b
L5ET
L5NP
L6short-a
L6short-b
L6tall-a
L6tall-b
L6tall-c
L6wm
PTC
DTC
STC
ITC

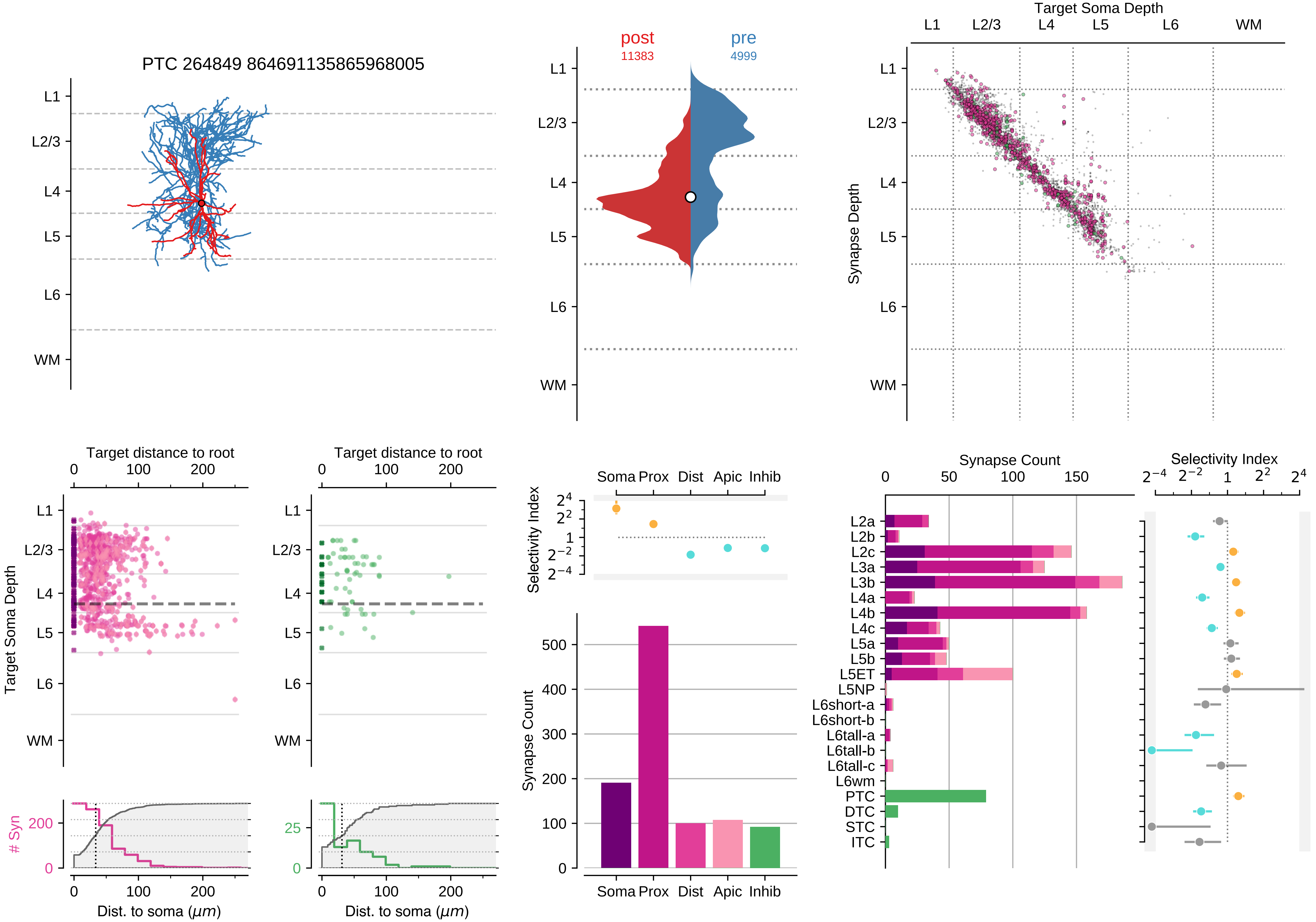

PTC 264849 864691135865968005
L1
L2/3
L4
L5
L6
WM
post
11383
pre
4999
Target Soma Depth
Synapse Depth
Target distance to root
Target Soma Depth
### Syn
Dist. to soma ($\mu m$)
Selectivity Index
Soma Prox Dist Apic Inhib
Synapse Count
L2a
L2b
L2c
L3a
L3b
L4a
L4b
L4c
L5a
L5b
L5ET
L5NP
L6short-a
L6short-b
L6tall-a
L6tall-b
L6tall-c
L6wm
PTC
DTC
STC
ITC

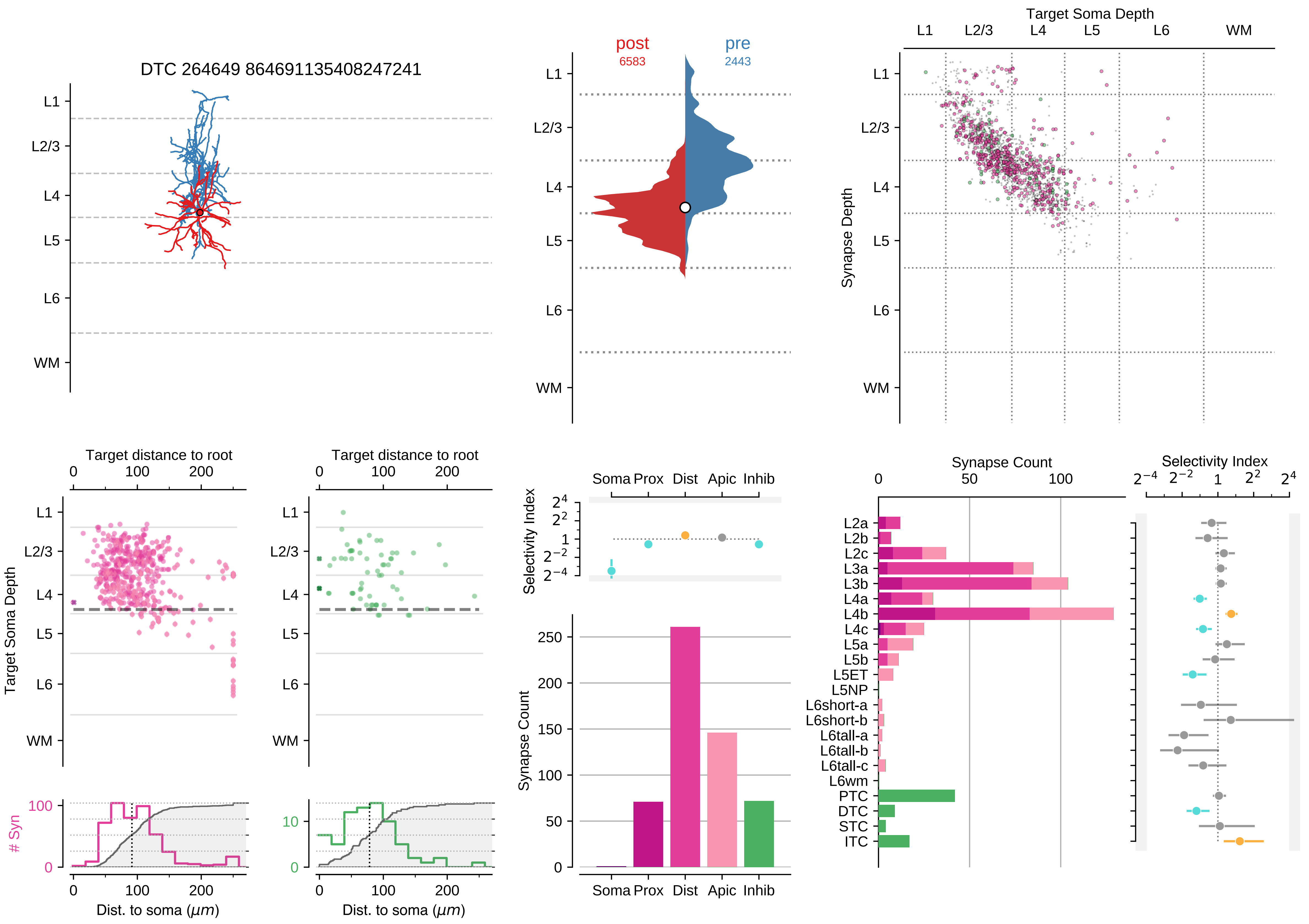
DTC 264649 864691135408247241
L1
L2/3
L4
L5
L6
WM
post
6583
pre
2443
Target Soma Depth
Synapse Depth
Target distance to root
Target Soma Depth
### Syn
Dist. to soma ($\mu m$)
Selectivity Index
Soma Prox Dist Apic Inhib
Synapse Count
Synapse Count
Selectivity Index
L2a
L2b
L2c
L3a
L3b
L4a
L4b
L4c
L5a
L5b
L5ET
L5NP
L6short-a
L6short-b
L6tall-a
L6tall-b
L6tall-c
L6wm
PTC
DTC
STC
ITC

### Motif Group 5

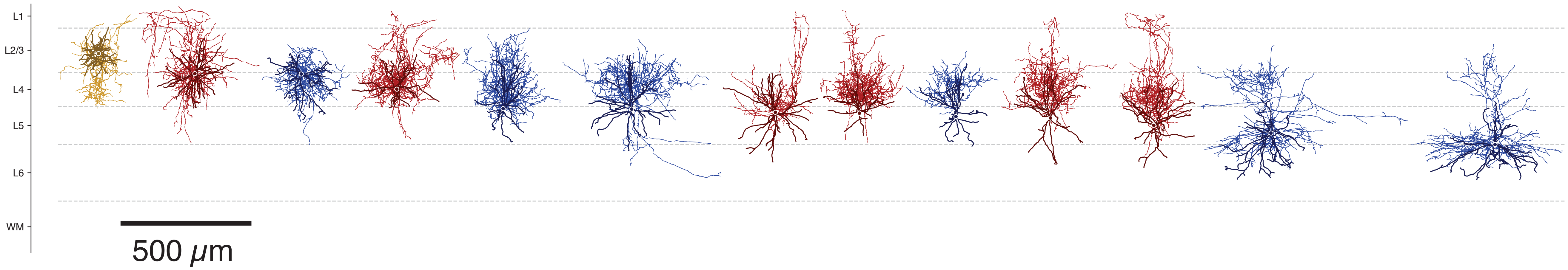

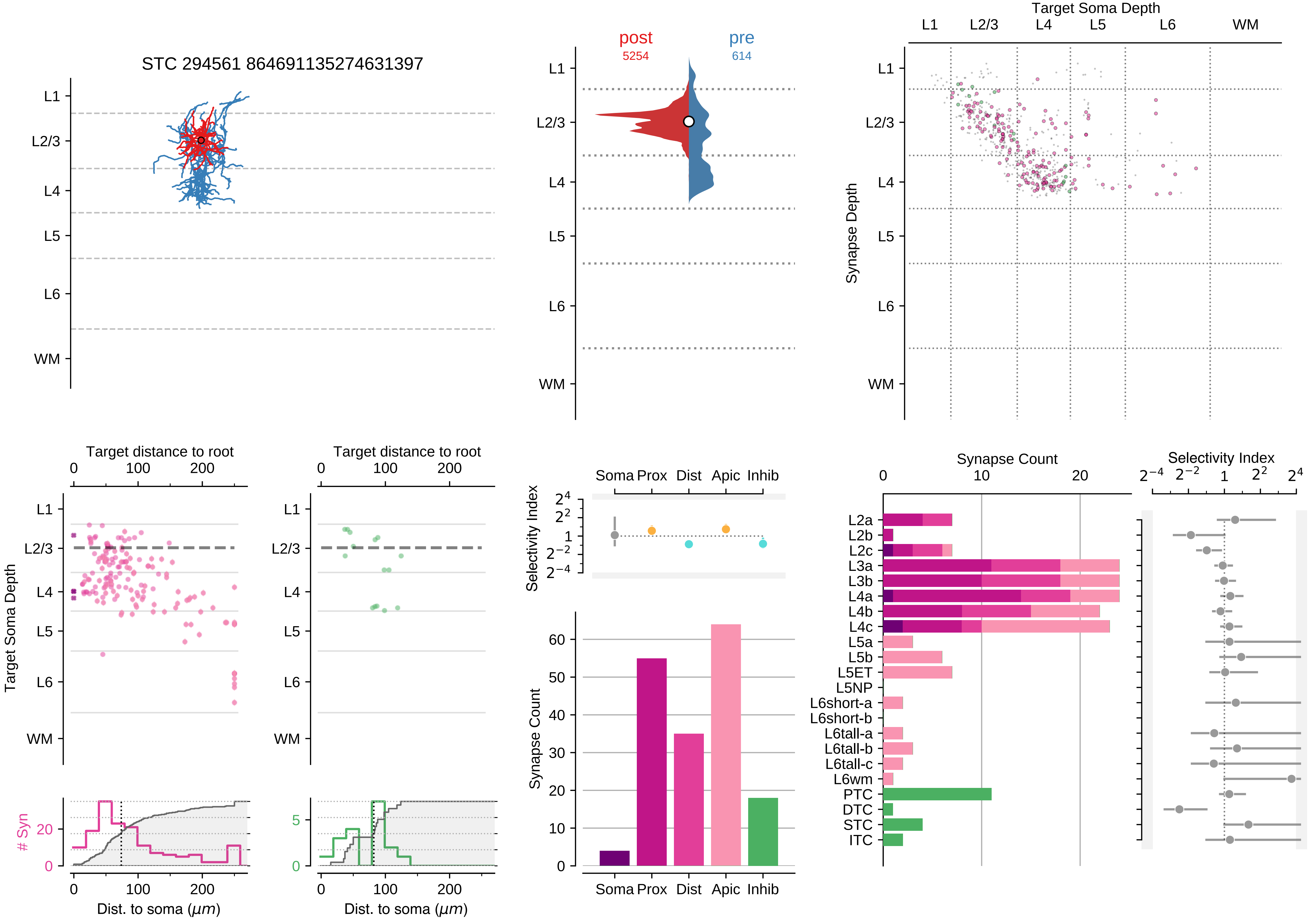

STC 294561 864691135274631397
L1
L2/3
L4
L5
L6
WM
post
5254
pre
614
Target Soma Depth
Synapse Depth
Target distance to root
Target Soma Depth
### Syn
Dist. to soma (μm)
Selectivity Index
Soma Prox Dist Apic Inhib
Synapse Count
L2a
L2b
L2c
L3a
L3b
L4a
L4b
L4c
L5a
L5b
L5ET
L5NP
L6short-a
L6short-b
L6tall-a
L6tall-b
L6tall-c
L6wm
PTC
DTC
STC
ITC

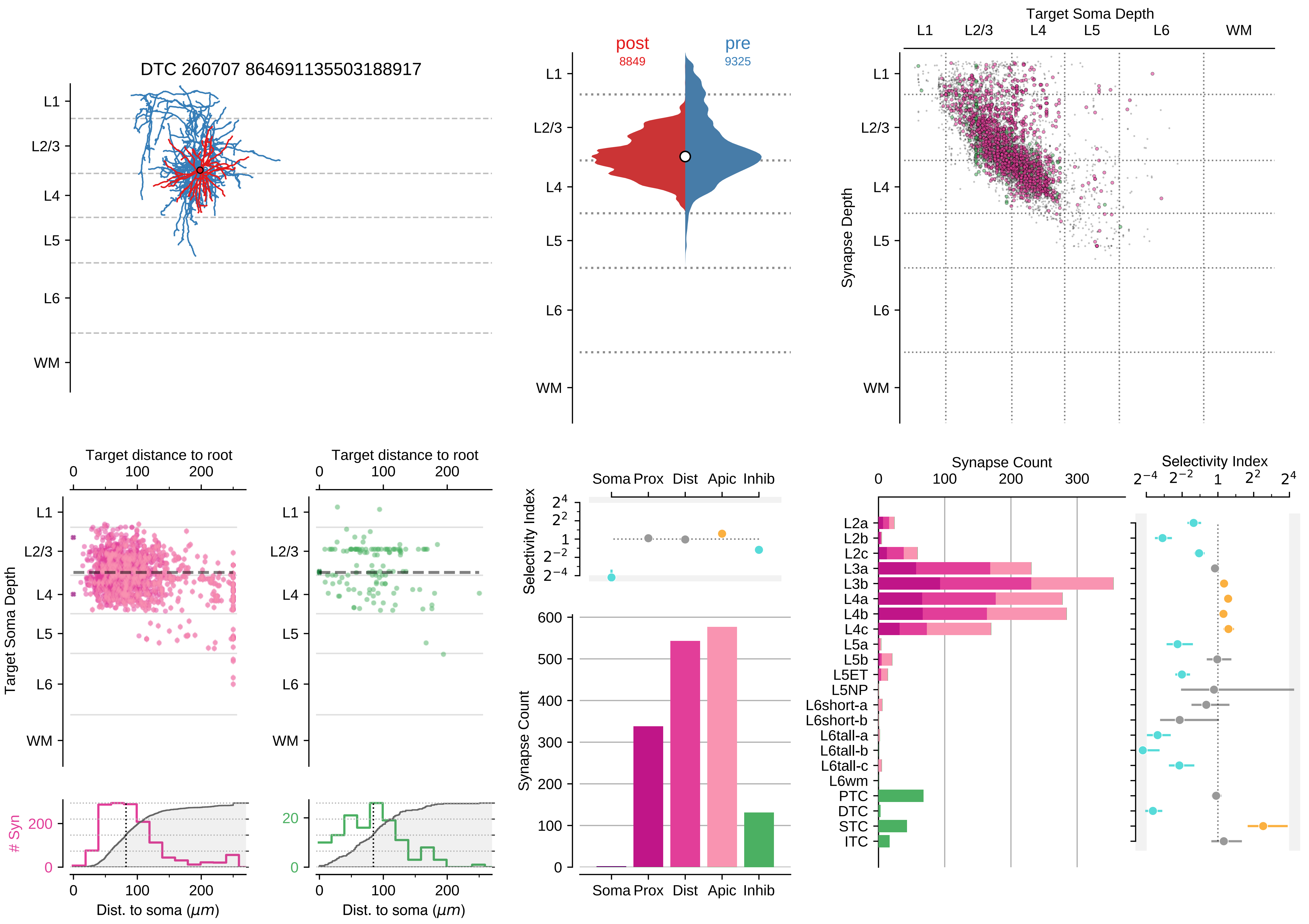

DTC 260707 864691135503188917
L1
L2/3
L4
L5
L6
WM
post
8849
pre
9325
Target Soma Depth
Synapse Depth
Target distance to root
Target Soma Depth
### Syn
Dist. to soma (μm)
Selectivity Index
Soma Prox Dist Apic Inhib
Synapse Count
L2a
L2b
L2c
L3a
L3b
L4a
L4b
L4c
L5a
L5b
L5ET
L5NP
L6short-a
L6short-b
L6tall-a
L6tall-b
L6tall-c
L6wm
PTC
DTC
STC
ITC

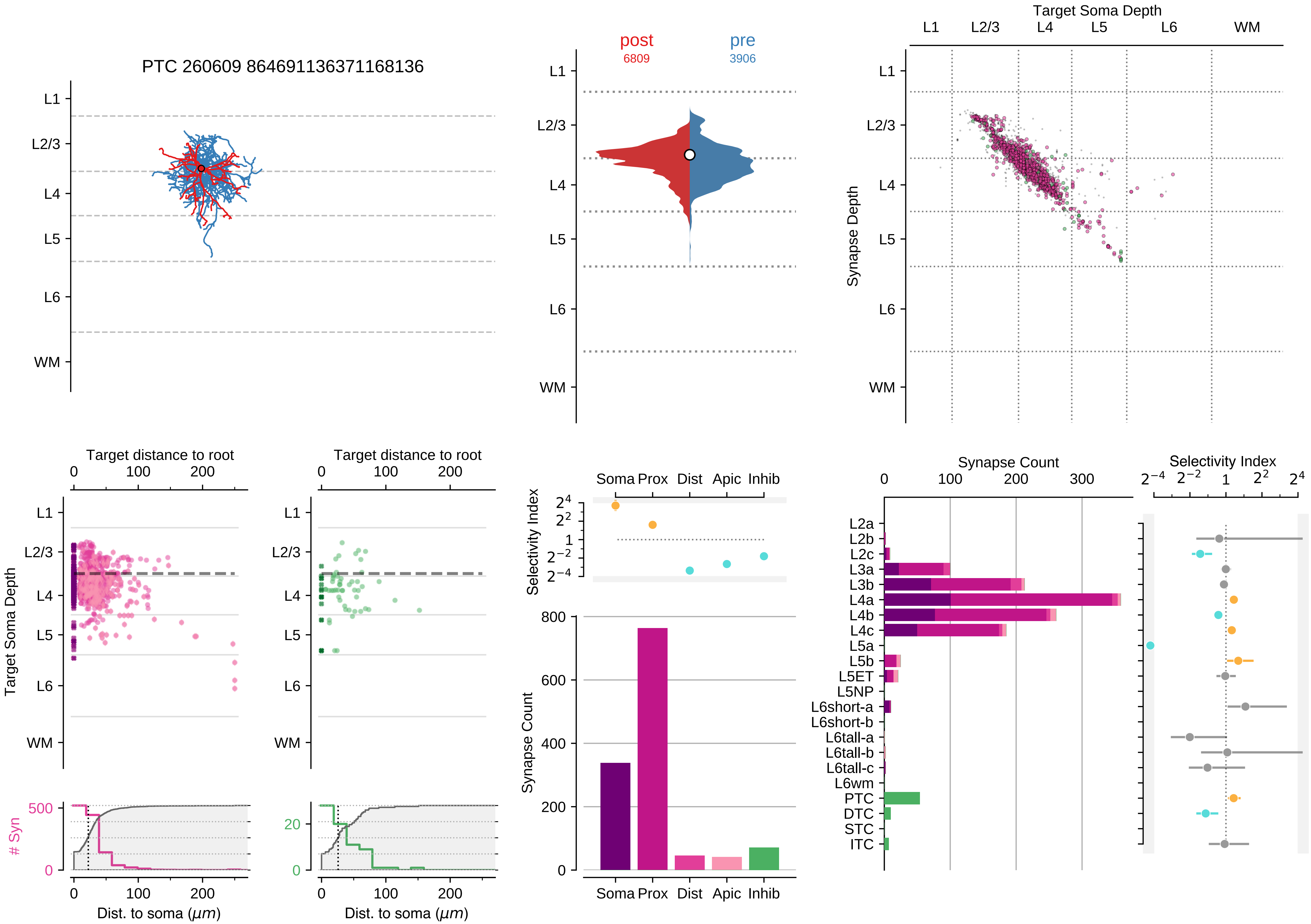

PTC 260609 864691136371168136
L1
L2/3
L4
L5
L6
WM
post
6809
pre
3906
Target Soma Depth
Synapse Depth
Target distance to root
Target Soma Depth
### Syn
Dist. to soma (μm)
Selectivity Index
Soma Prox Dist Apic Inhib
Synapse Count
Selectivity Index
L2a
L2b
L2c
L3a
L3b
L4a
L4b
L4c
L5a
L5b
L5ET
L5NP
L6short-a
L6short-b
L6tall-a
L6tall-b
L6tall-c
L6wm
PTC
DTC
STC
ITC

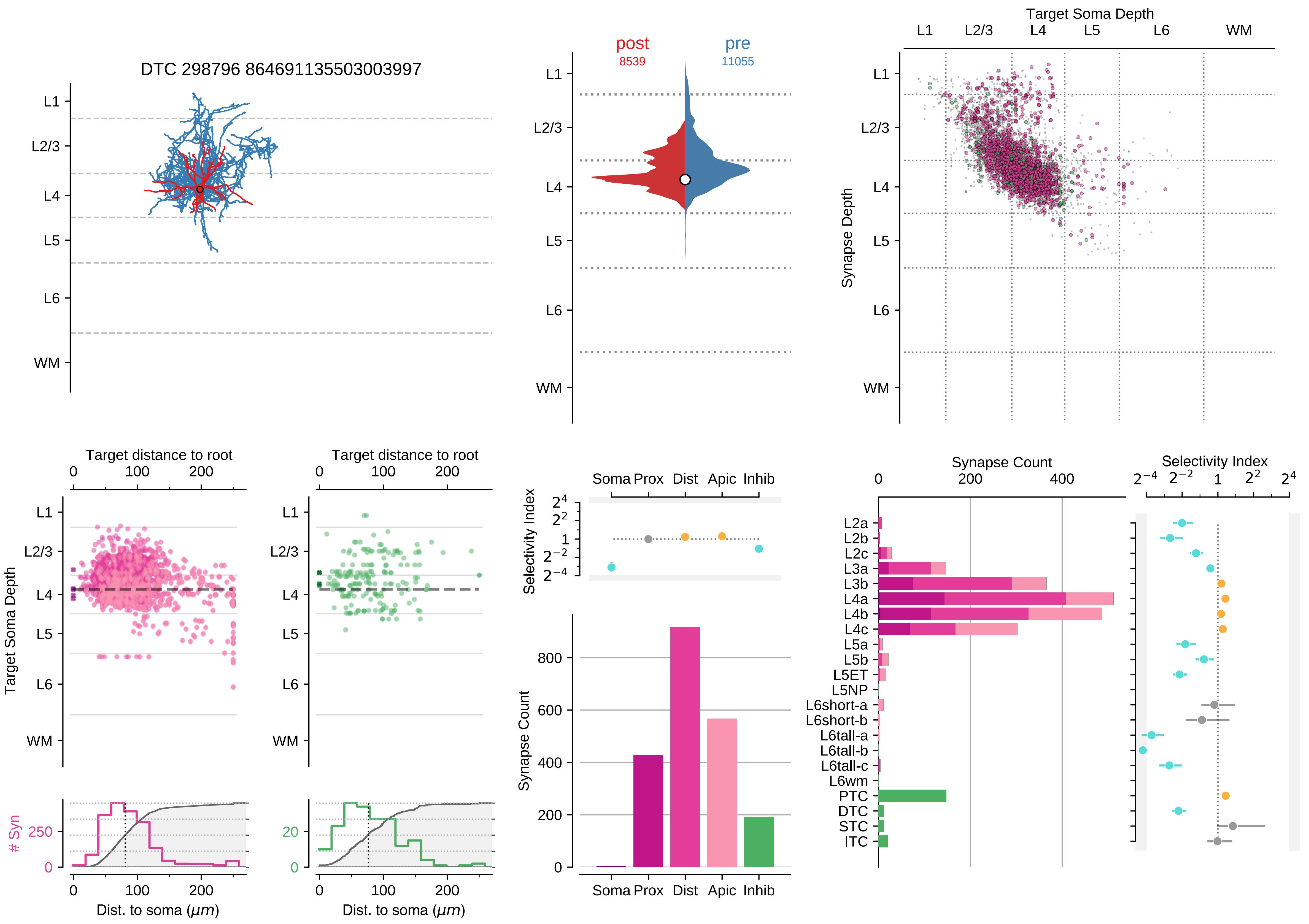

DTC 298796 864691135503003997
L1
L2/3
L4
L5
L6
WM
post
8539
pre
11055
Target Soma Depth
Synapse Depth
Target distance to root
Dist. to soma ($\mu m$)
### Syn
Selectivity Index
Soma Prox
Dist
Apic
Inhib
Synapse Count
L2a
L2b
L2c
L3a
L3b
L4a
L4b
L4c
L5a
L5b
L5ET
L5NP
L6short-a
L6short-b
L6tall-a
L6tall-b
L6tall-c
L6wm
PTC
DTC
STC
ITC

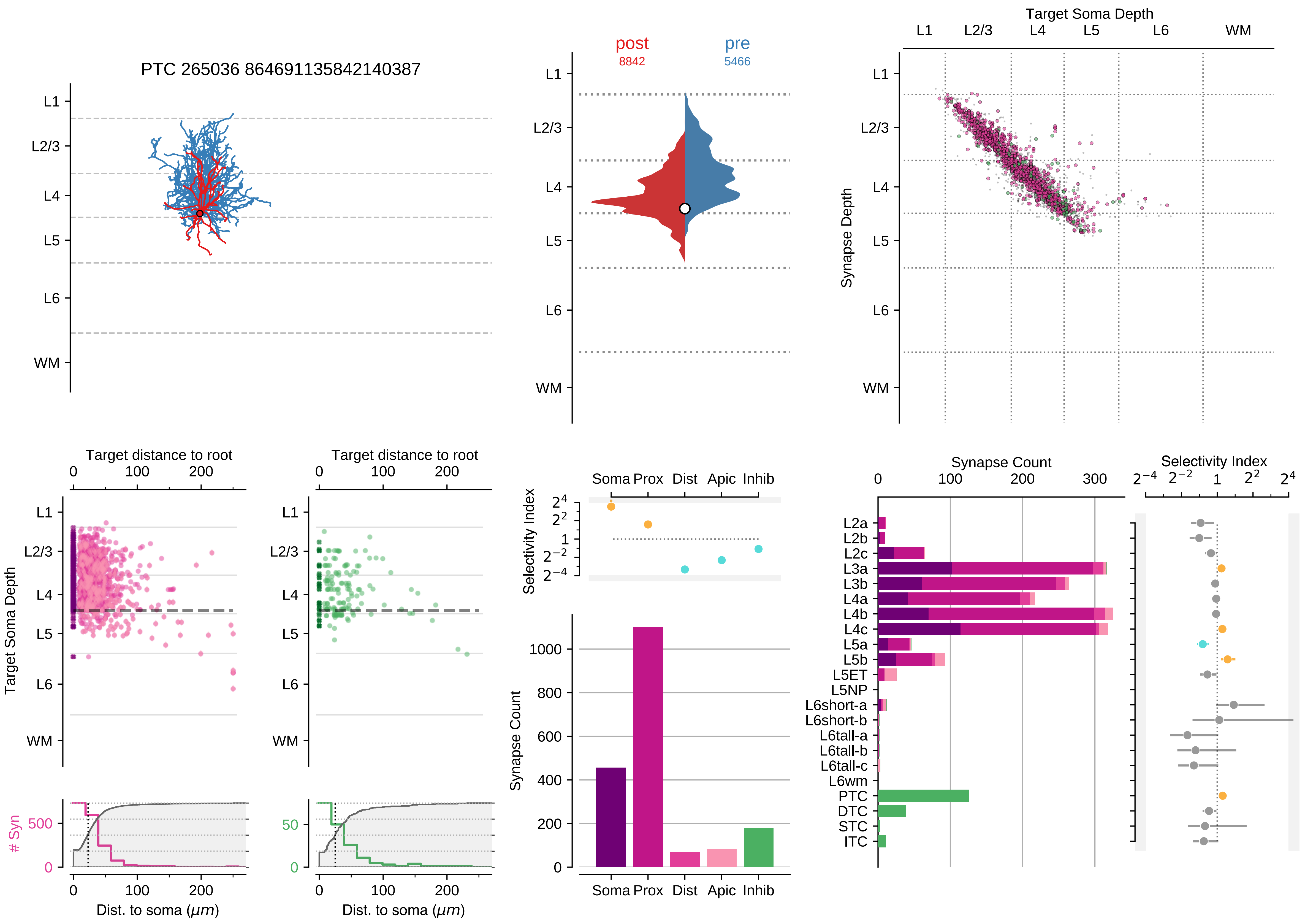

PTC 265036 864691135842140387
L1
L2/3
L4
L5
L6
WM
post
8842
pre
5466
Target Soma Depth
Synapse Depth
Target distance to root
Target Soma Depth
### Syn
Dist. to soma (μm)
Selectivity Index
Soma Prox Dist Apic Inhib
Synapse Count
L2a
L2b
L2c
L3a
L3b
L4a
L4b
L4c
L5a
L5b
L5ET
L5NP
L6short-a
L6short-b
L6tall-a
L6tall-b
L6tall-c
L6wm
PTC
DTC
STC
ITC

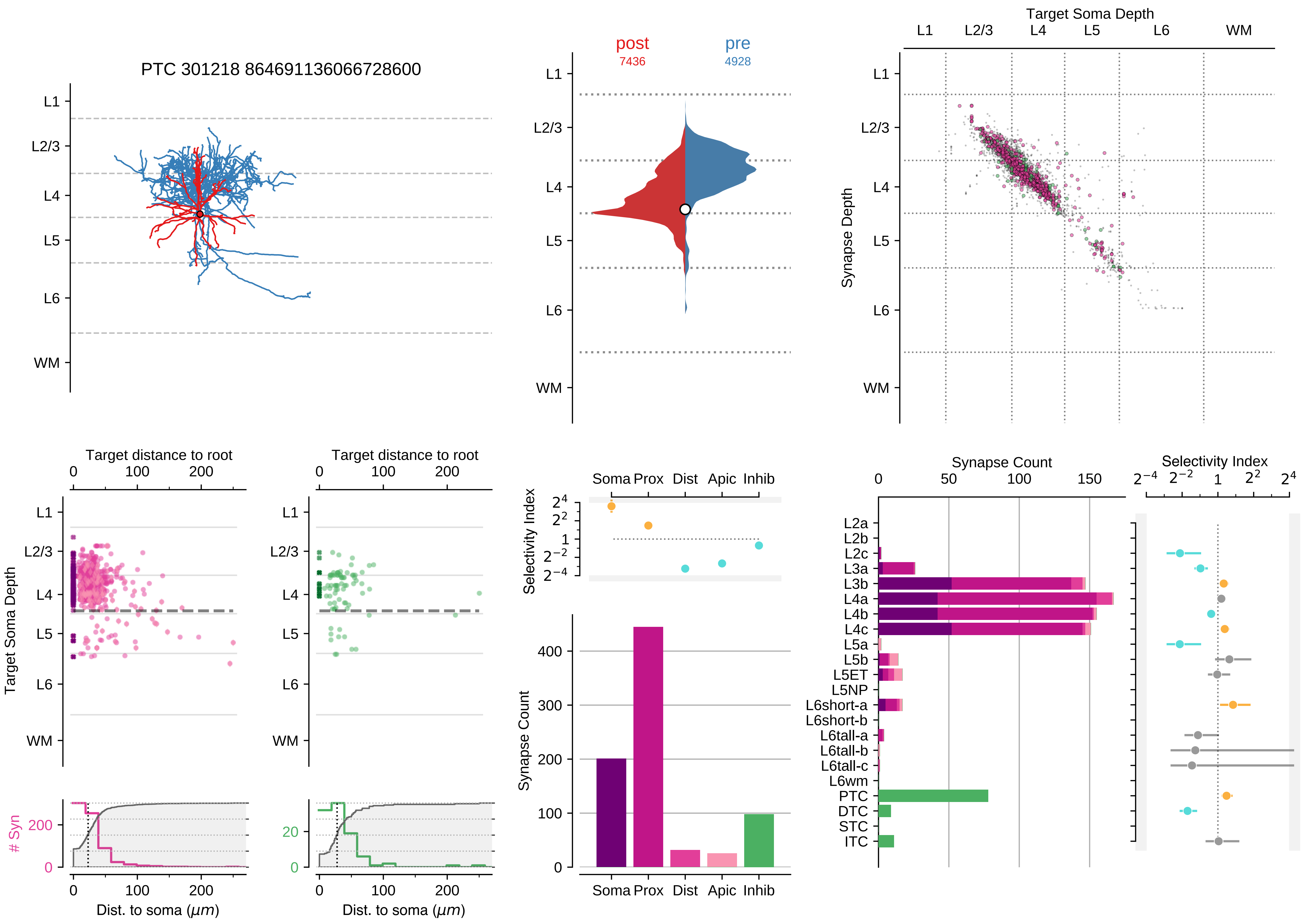

PTC 301218 864691136066728600
L1
L2/3
L4
L5
L6
WM
post
7436
pre
4928
Target Soma Depth
Synapse Depth
Target distance to root
0
100
200
Target Soma Depth
### Syn
Dist. to soma (μm)
Selectivity Index
Soma Prox Dist Apic Inhib
Synapse Count
Synapse Count
0
50
100
150
Selectivity Index
L2a
L2b
L2c
L3a
L3b
L4a
L4b
L4c
L5a
L5b
L5ET
L5NP
L6short-a
L6short-b
L6tall-a
L6tall-b
L6tall-c
L6wm
PTC
DTC
STC
ITC

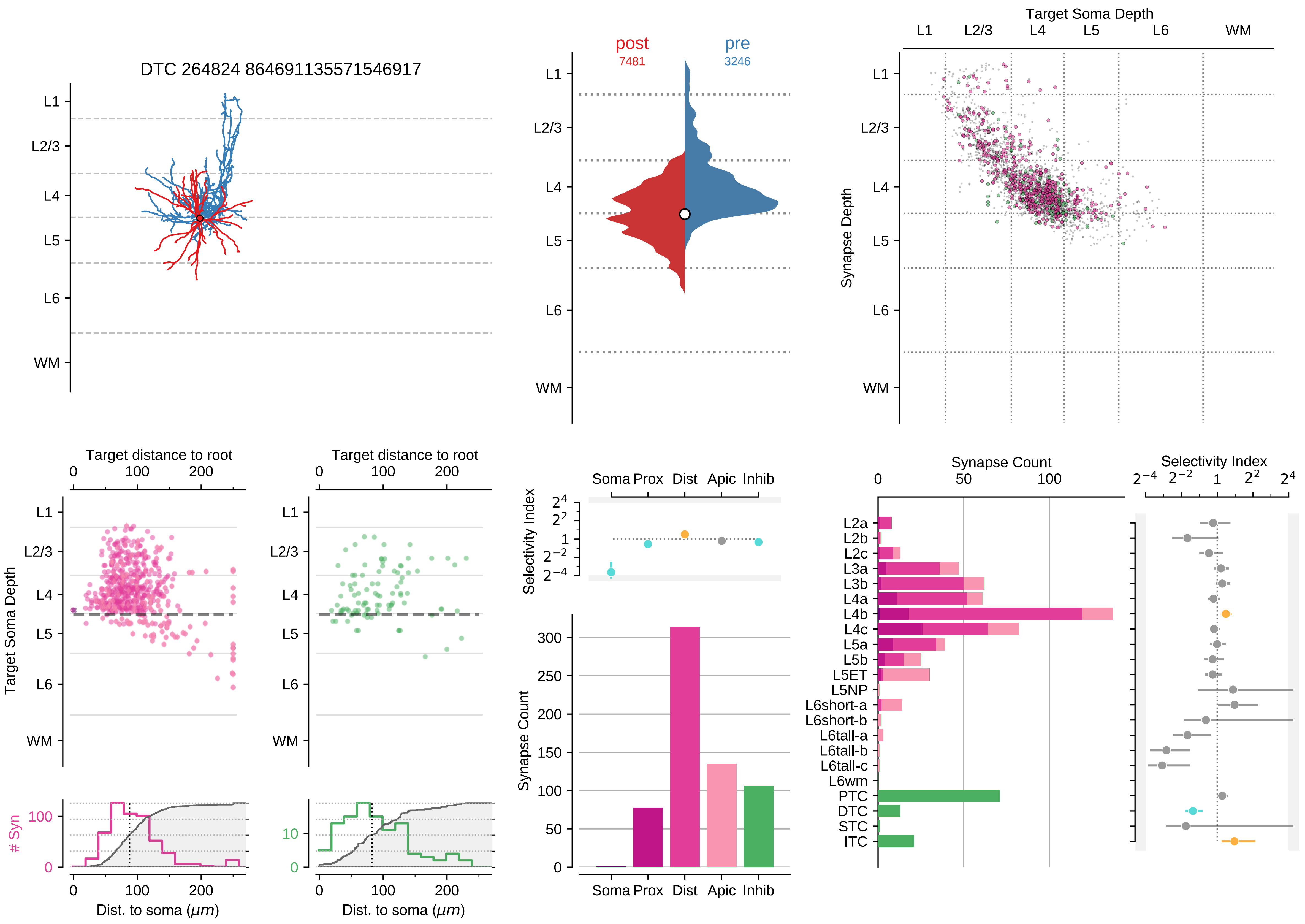
DTC 264824 864691135571546917
L1
L2/3
L4
L5
L6
WM
post
7481
pre
3246
Target Soma Depth
Synapse Depth
Target distance to root
Target Soma Depth
### Syn
Dist. to soma ($\mu m$)
Selectivity Index
Soma Prox Dist Apic Inhib
Synapse Count
L2a
L2b
L2c
L3a
L3b
L4a
L4b
L4c
L5a
L5b
L5ET
L5NP
L6short-a
L6short-b
L6tall-a
L6tall-b
L6tall-c
L6wm
PTC
DTC
STC
ITC

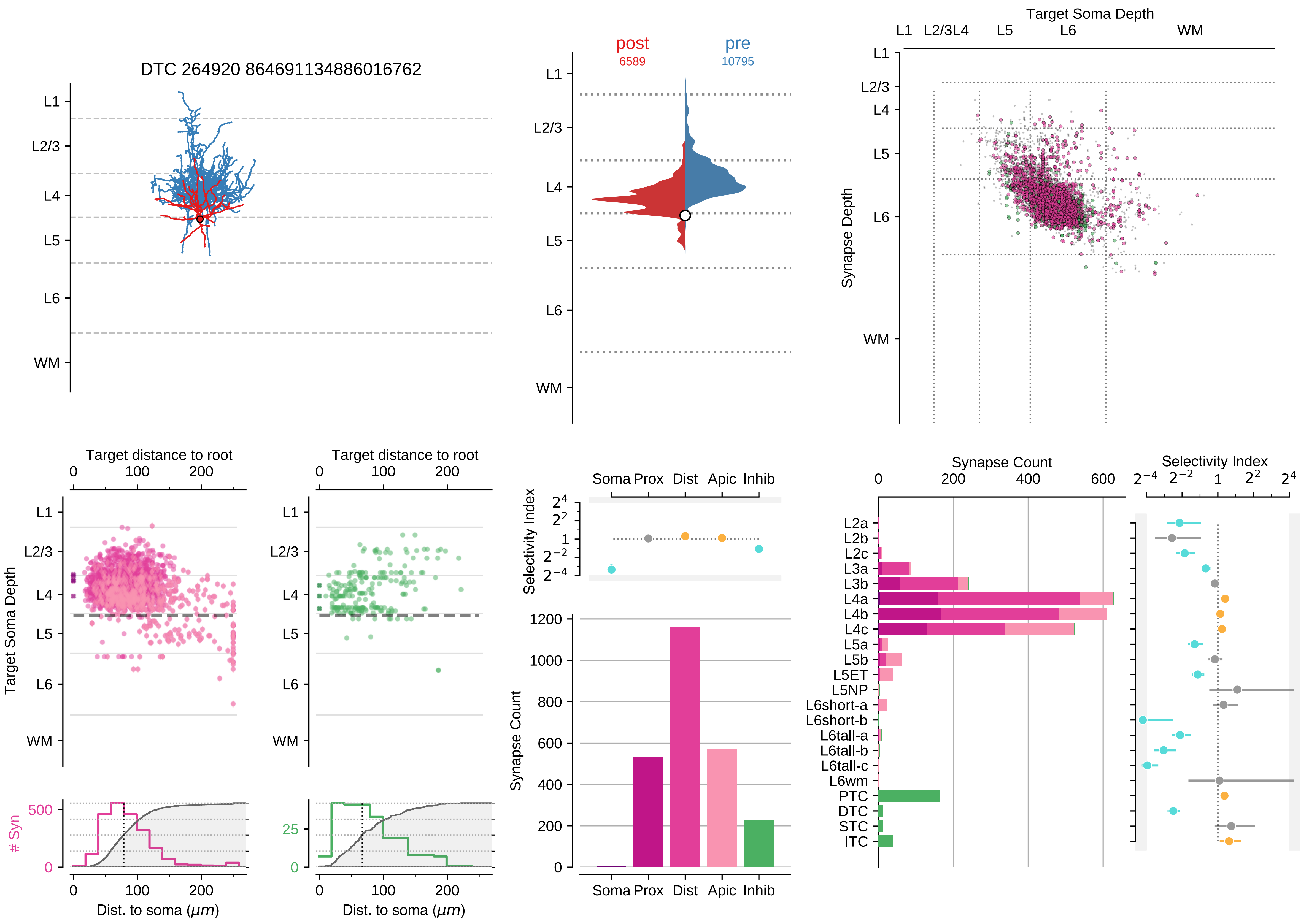

DTC 264920 864691134886016762
post
6589
pre
10795
Target Soma Depth
L1
L2/3
L4
L5
L6
WM
Synapse Depth
Target distance to root
Target Soma Depth
### Syn
Dist. to soma (μm)
Selectivity Index
Soma
Prox
Dist
Apic
Inhib
Synapse Count
L2a
L2b
L2c
L3a
L3b
L4a
L4b
L4c
L5a
L5b
L5ET
L5NP
L6short-a
L6short-b
L6tall-a
L6tall-b
L6tall-c
L6wm
PTC
DTC
STC
ITC

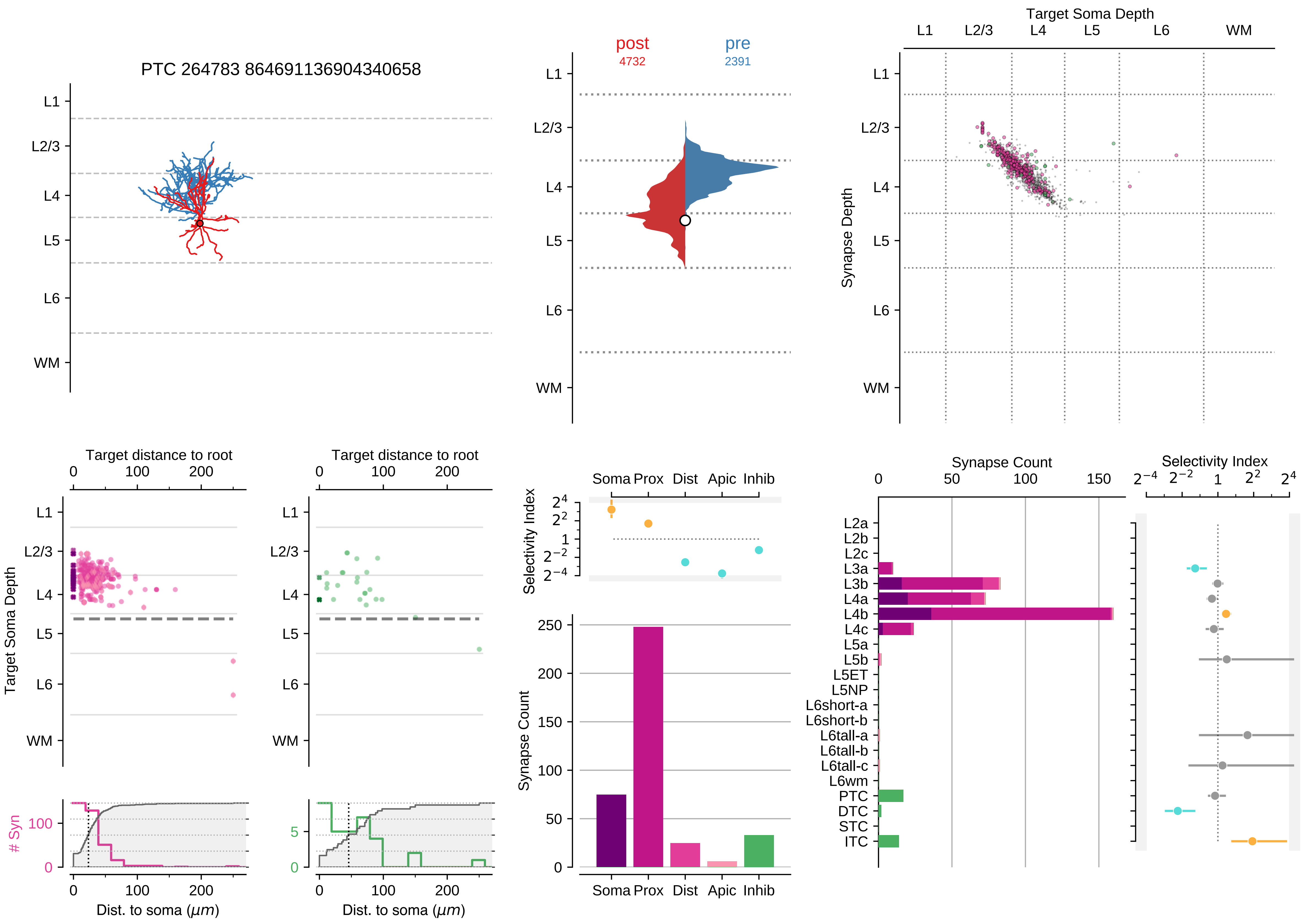

PTC 264783 864691136904340658
L1
L2/3
L4
L5
L6
WM
post
4732
pre
2391
Target Soma Depth
Synapse Depth
Target distance to root
0
100
200
Target Soma Depth
### Syn
Dist. to soma (μm)
Selectivity Index
Soma Prox Dist Apic Inhib
Synapse Count
Synapse Count
0
50
100
150
Selectivity Index
L2a
L2b
L2c
L3a
L3b
L4a
L4b
L4c
L5a
L5b
L5ET
L5NP
L6short-a
L6short-b
L6tall-a
L6tall-b
L6tall-c
L6wm
PTC
DTC
STC
ITC

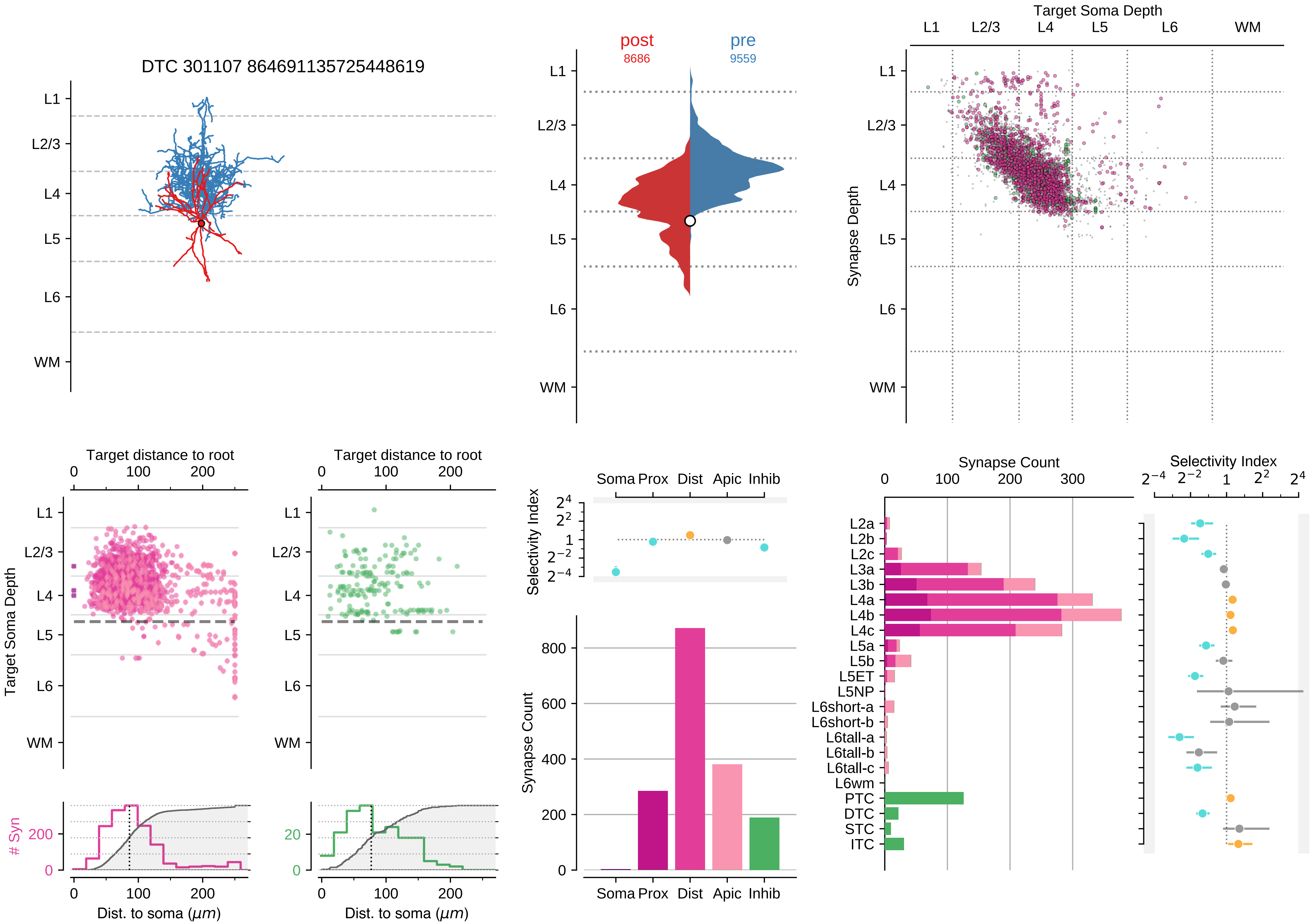

DTC 301107 864691135725448619
L1
L2/3
L4
L5
L6
WM
post
8686
pre
9559
Target Soma Depth
Synapse Depth
Target distance to root
Target Soma Depth
### Syn
Dist. to soma ($\mu m$)
Selectivity Index
Soma Prox Dist Apic Inhib
Synapse Count
L2a
L2b
L2c
L3a
L3b
L4a
L4b
L4c
L5a
L5b
L5ET
L5NP
L6short-a
L6short-b
L6tall-a
L6tall-b
L6tall-c
L6wm
PTC
DTC
STC
ITC

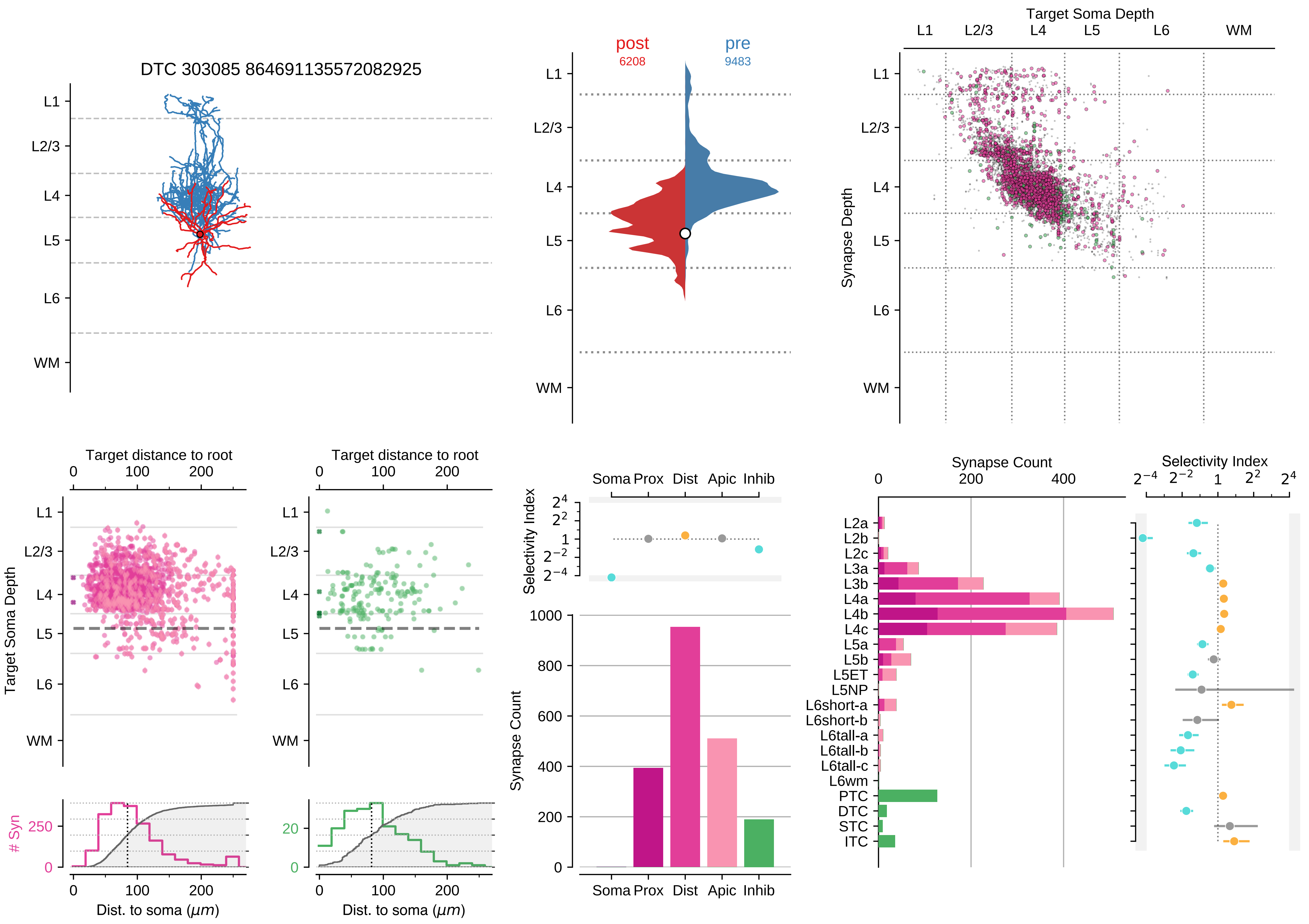

DTC 303085 864691135572082925
L1
L2/3
L4
L5
L6
WM
post
6208
pre
9483
Target Soma Depth
Synapse Depth
Target distance to root
Target Soma Depth
### Syn
Dist. to soma ($\mu m$)
Soma Prox Dist Apic Inhib
Selectivity Index
Synapse Count
L2a
L2b
L2c
L3a
L3b
L4a
L4b
L4c
L5a
L5b
L5ET
L5NP
L6short-a
L6short-b
L6tall-a
L6tall-b
L6tall-c
L6wm
PTC
DTC
STC
ITC

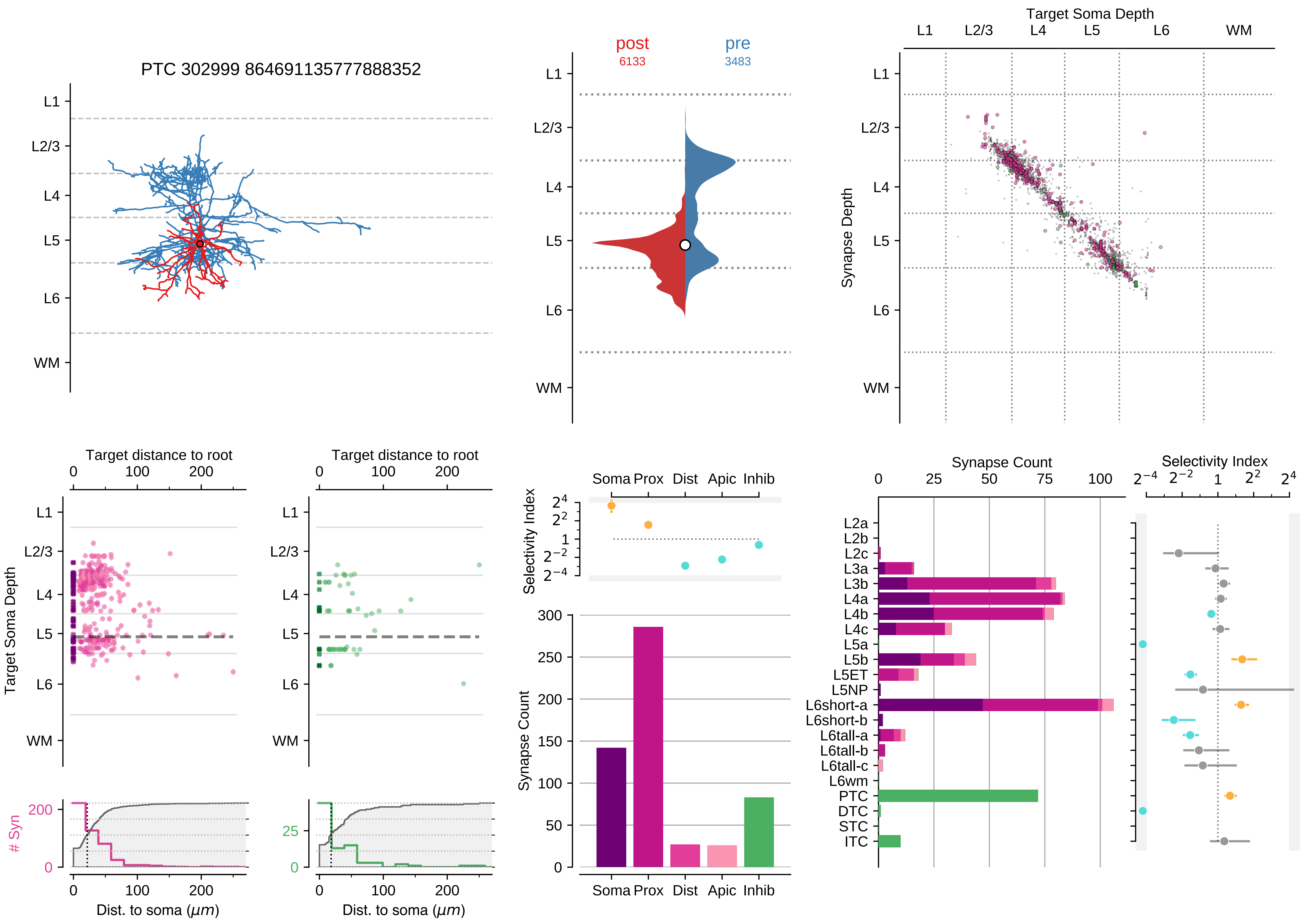

PTC 302999 864691135777888352
post
6133
pre
3483
Target Soma Depth
Synapse Depth
L1
L2/3
L4
L5
L6
WM
Target distance to root
Target Soma Depth
### Syn
Dist. to soma ($\mu m$)
Selectivity Index
Soma Prox Dist Apic Inhib
Synapse Count
L2a
L2b
L2c
L3a
L3b
L4a
L4b
L4c
L5a
L5b
L5ET
L5NP
L6short-a
L6short-b
L6tall-a
L6tall-b
L6tall-c
L6wm
PTC
DTC
STC
ITC

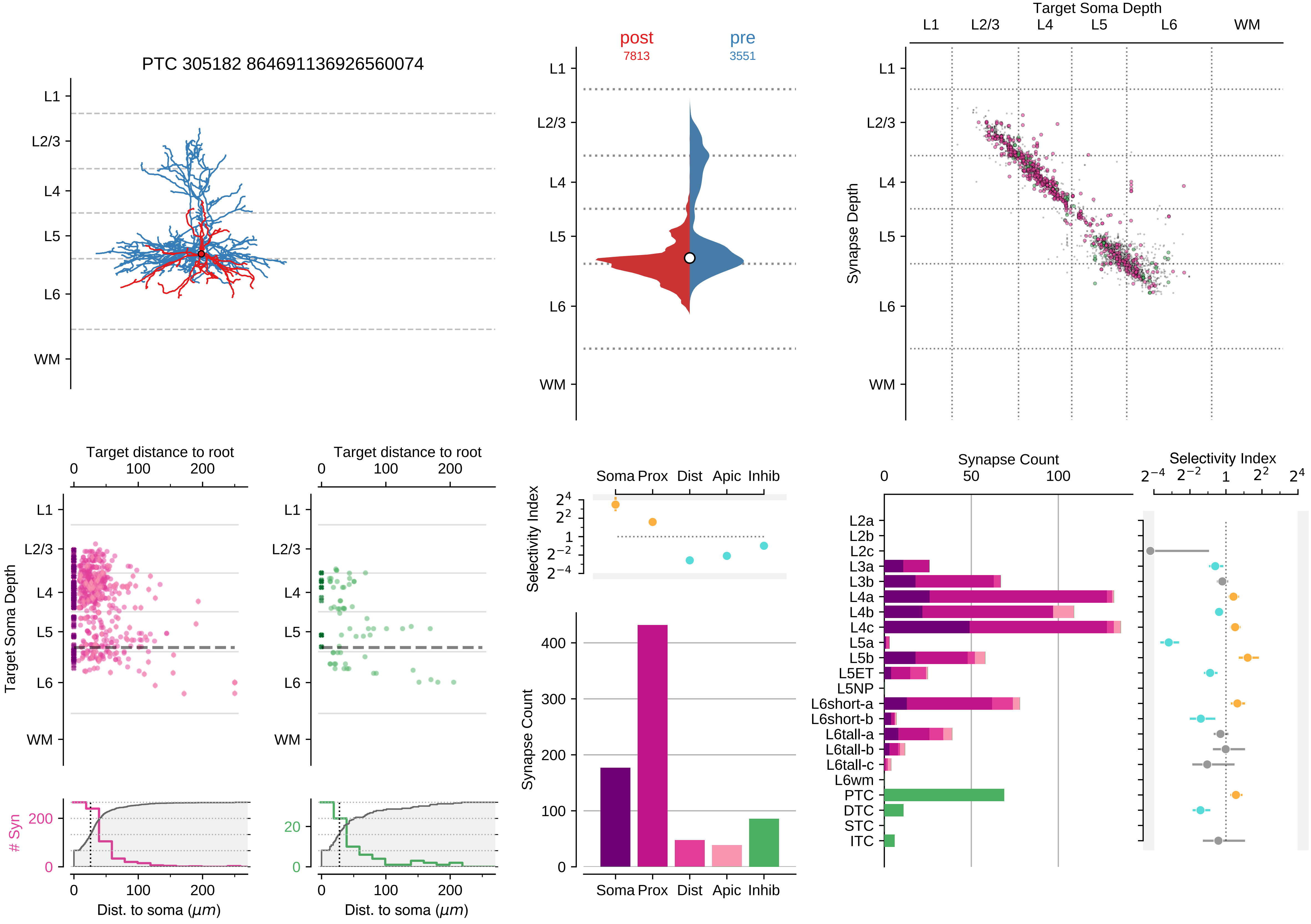
PTC 305182 864691136926560074
L1
L2/3
L4
L5
L6
WM
post
7813
pre
3551
Target Soma Depth
Synapse Depth
Target distance to root
0
100
200
Target Soma Depth
### Syn
Dist. to soma (μm)
Selectivity Index
Soma
Prox
Dist
Apic
Inhib
Synapse Count
400
300
200
100
Synapse Count
50
100
Selectivity Index
L2a
L2b
L2c
L3a
L3b
L4a
L4b
L4c
L5a
L5b
L5ET
L5NP
L6short-a
L6short-b
L6tall-a
L6tall-b
L6tall-c
L6wm
PTC
DTC
STC
ITC

### Motif Group 6

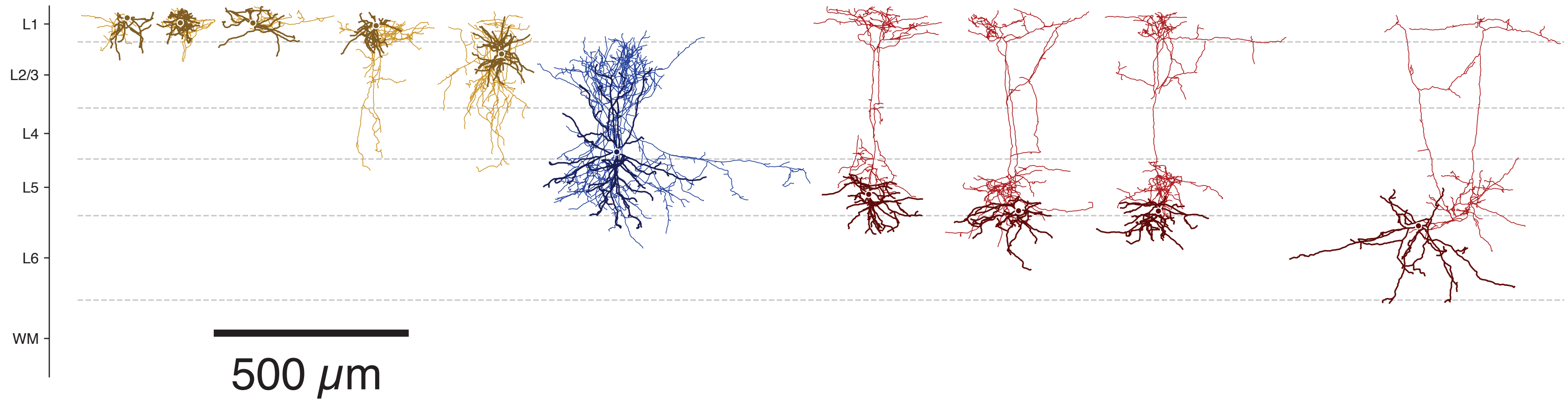

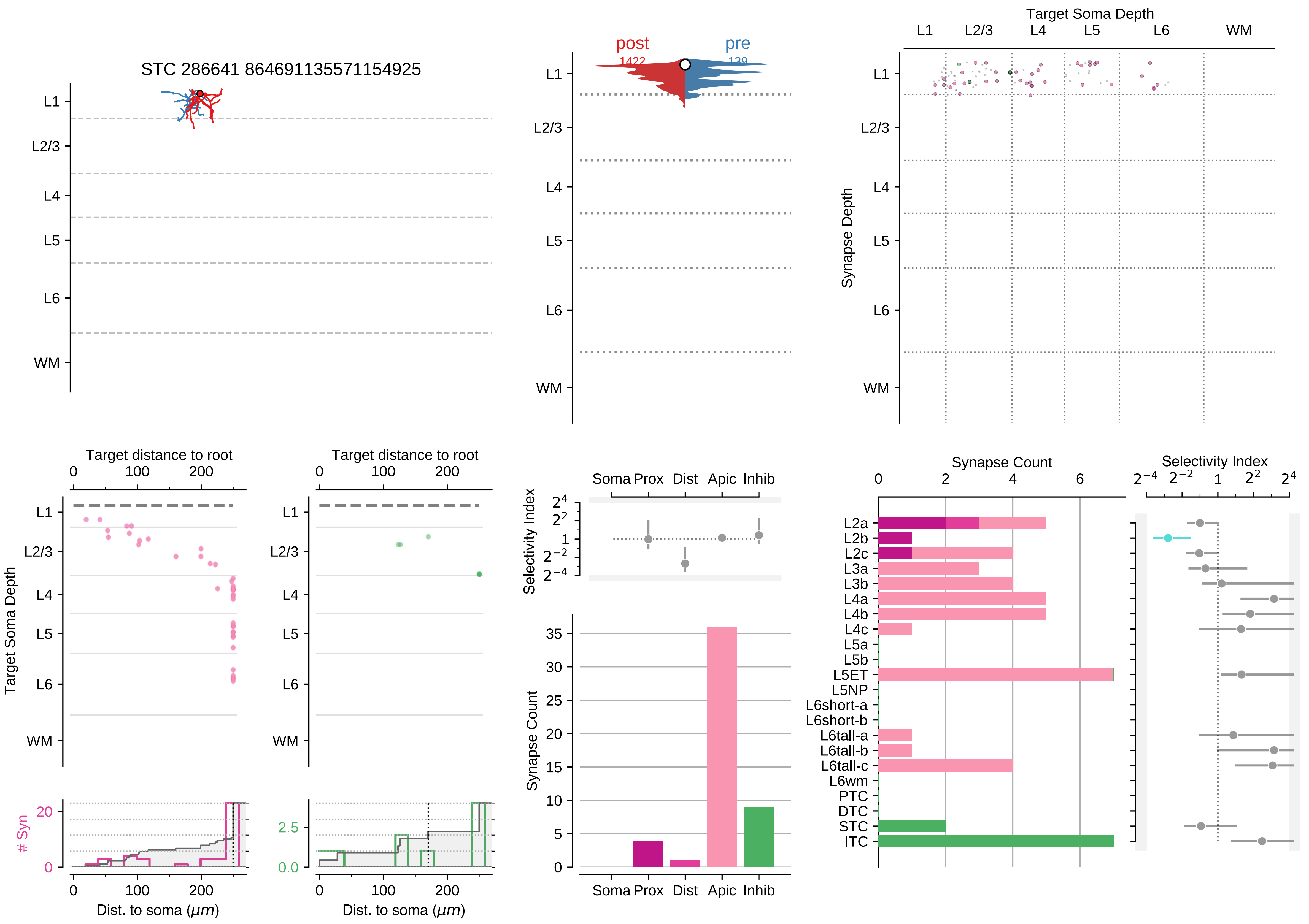

STC 286641 864691135571154925
L1
L2/3
L4
L5
L6
WM
post
1422
pre
139
Target Soma Depth
Synapse Depth
Target distance to root
Target Soma Depth
Dist. to soma (μm)
### Syn
Selectivity Index
Soma Prox
Dist
Apic
Inhib
Synapse Count
L2a
L2b
L2c
L3a
L3b
L4a
L4b
L4c
L5a
L5b
L5ET
L5NP
L6short-a
L6short-b
L6tall-a
L6tall-b
L6tall-c
L6wm
PTC
DTC
STC
ITC

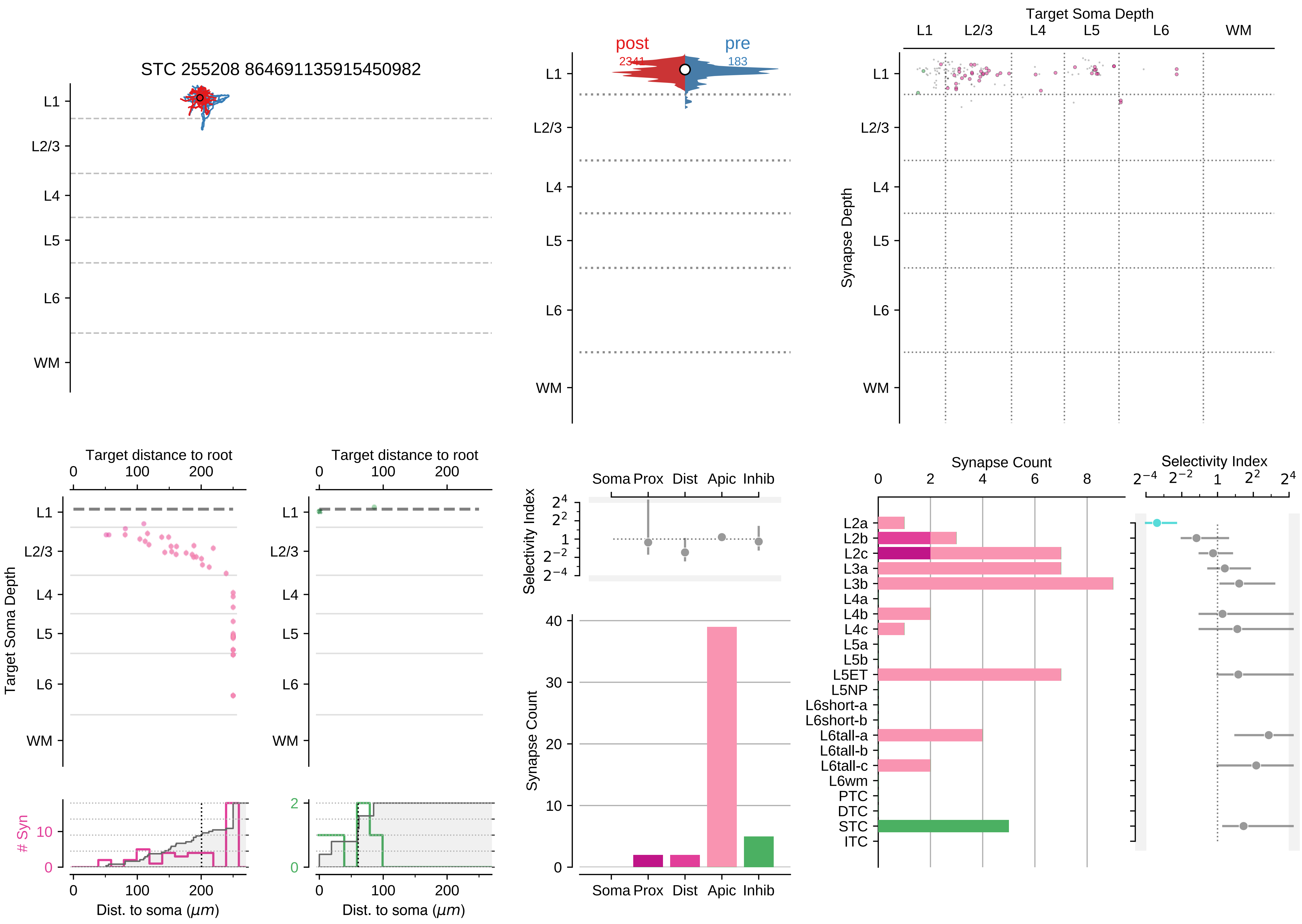
STC 255208 864691135915450982
post
2341
pre
183
Target Soma Depth
Synapse Depth
L1
L2/3
L4
L5
L6
WM
Target distance to root
Target Soma Depth
### Syn
Dist. to soma ($\mu m$)
Selectivity Index
Soma Prox Dist Apic Inhib
Synapse Count
Synapse Count
Selectivity Index
L2a
L2b
L2c
L3a
L3b
L4a
L4b
L4c
L5a
L5b
L5ET
L5NP
L6short-a
L6short-b
L6tall-a
L6tall-b
L6tall-c
L6wm
PTC
DTC
STC
ITC

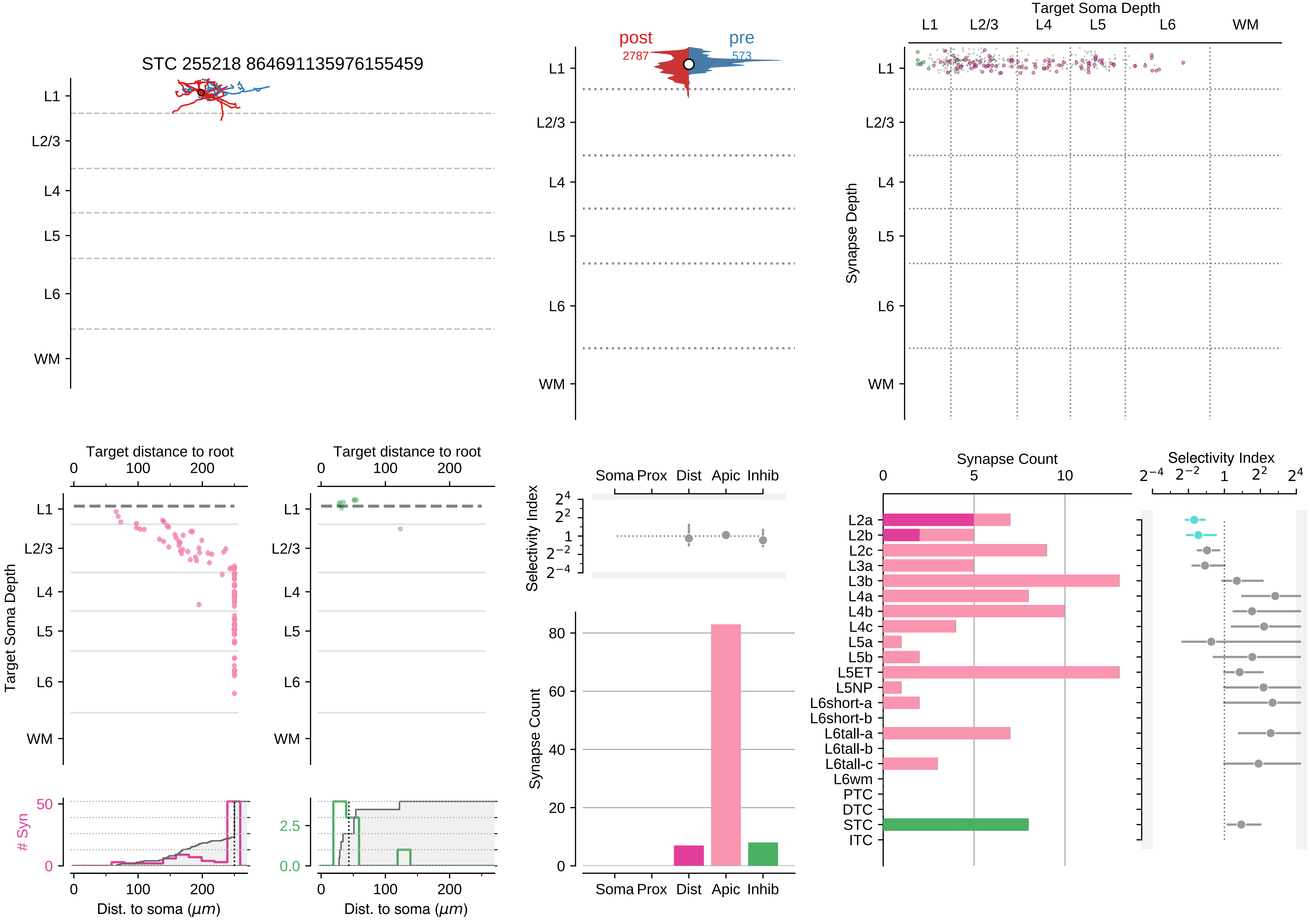

STC 255218 864691135976155459
L1
L2/3
L4
L5
L6
WM
post
2787
pre
573
Target Soma Depth
Synapse Depth
Target distance to root
Dist. to soma ($\mu m$)
### Syn
Selectivity Index
Soma Prox
Dist
Apic
Inhib
Synapse Count
L2a
L2b
L2c
L3a
L3b
L4a
L4b
L4c
L5a
L5b
L5ET
L5NP
L6short-a
L6short-b
L6tall-a
L6tall-b
L6tall-c
L6wm
PTC
DTC
STC
ITC

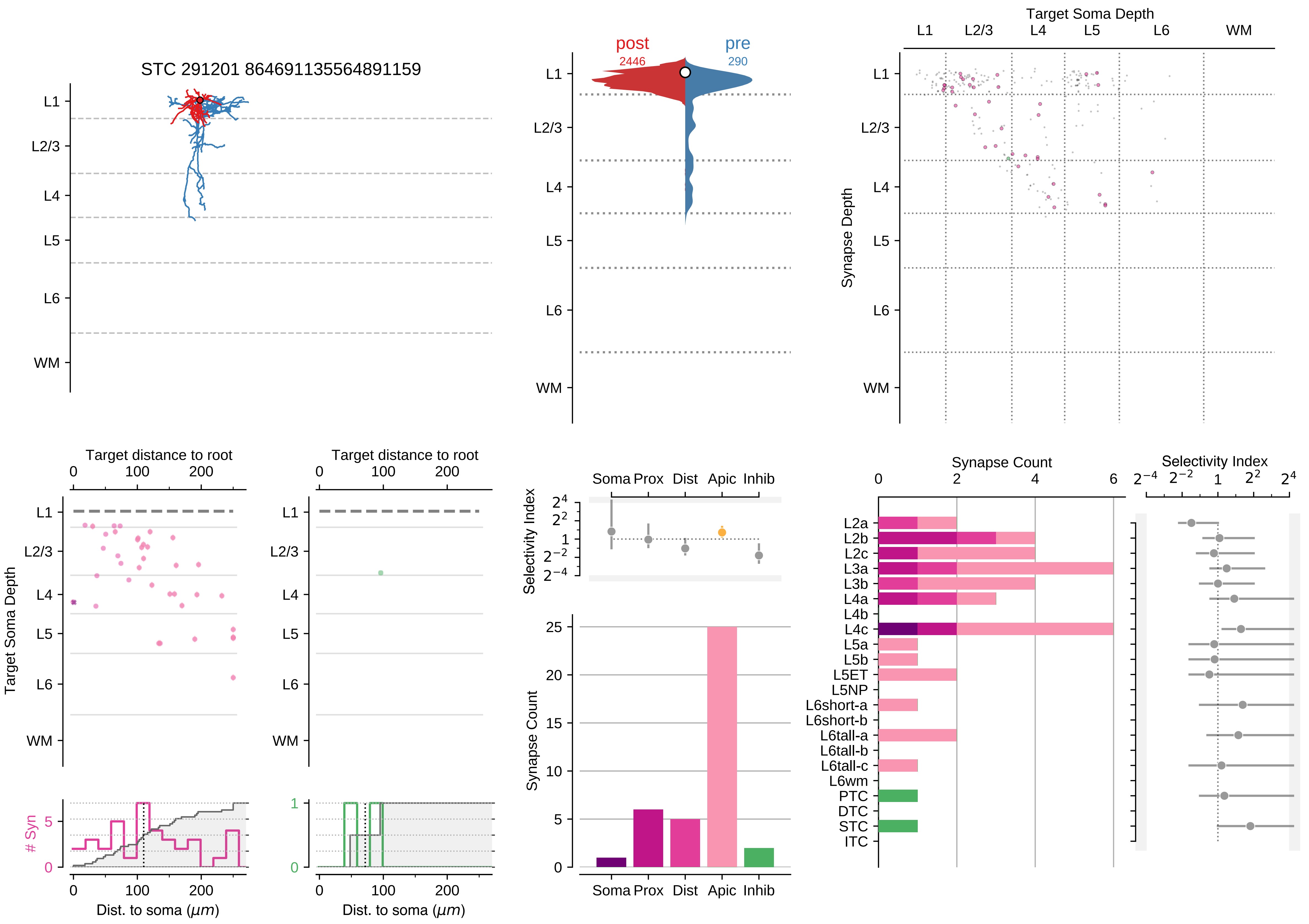

STC 291201 864691135564891159
L1
L2/3
L4
L5
L6
WM
post
2446
pre
290
Target Soma Depth
Synapse Depth
Target distance to root
Dist. to soma (μm)
### Syn
Selectivity Index
Synapse Count
Soma
Prox
Dist
Apic
Inhib
L2a
L2b
L2c
L3a
L3b
L4a
L4b
L4c
L5a
L5b
L5ET
L5NP
L6short-a
L6short-b
L6tall-a
L6tall-b
L6tall-c
L6wm
PTC
DTC
STC
ITC

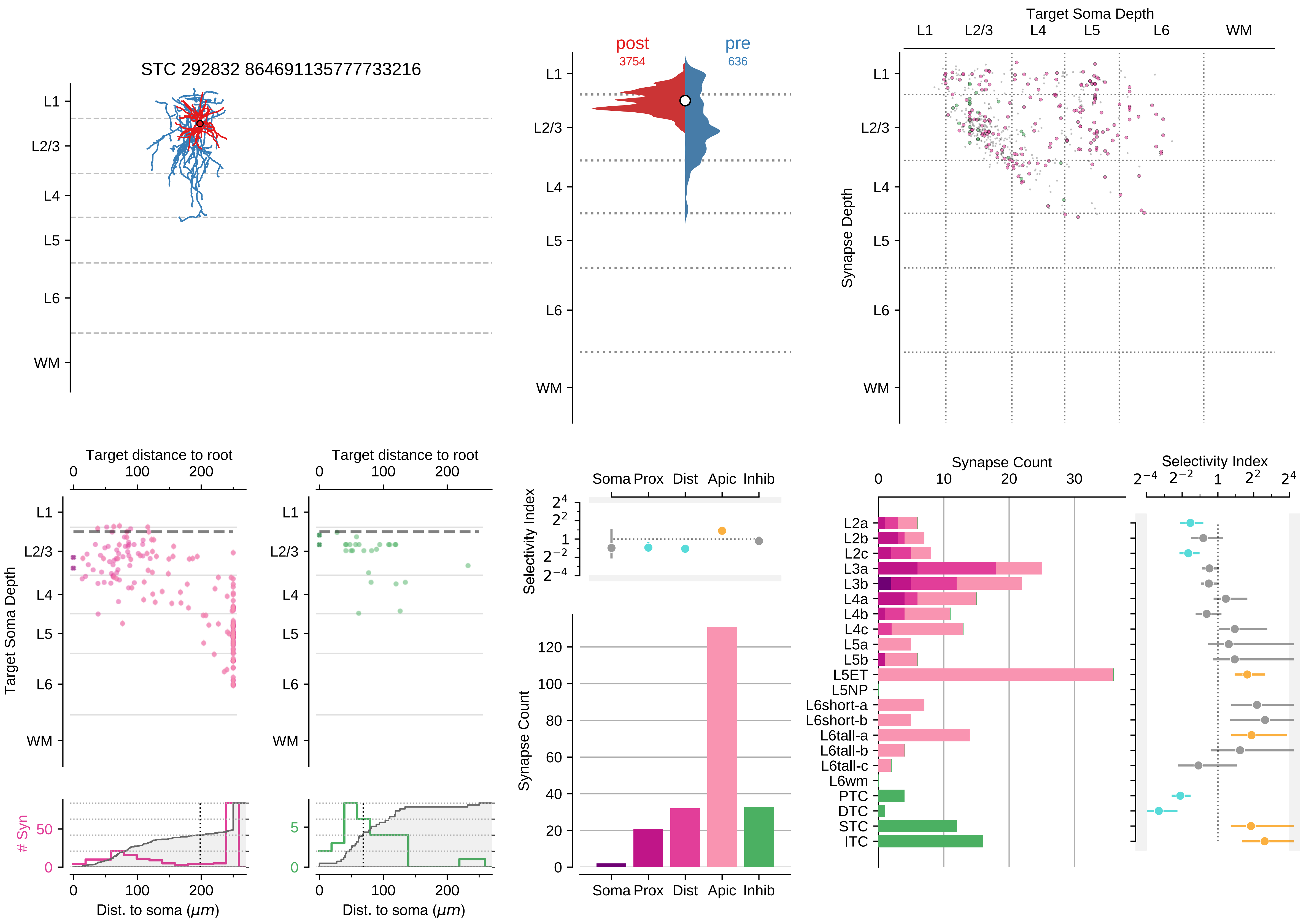

STC 292832 864691135777733216
L1
L2/3
L4
L5
L6
WM
post
3754
pre
636
Target Soma Depth
Synapse Depth
Target distance to root
Target Soma Depth
### Syn
Dist. to soma (μm)
Selectivity Index
Soma Prox Dist Apic Inhib
Synapse Count
L2a
L2b
L2c
L3a
L3b
L4a
L4b
L4c
L5a
L5b
L5ET
L5NP
L6short-a
L6short-b
L6tall-a
L6tall-b
L6tall-c
L6wm
PTC
DTC
STC
ITC

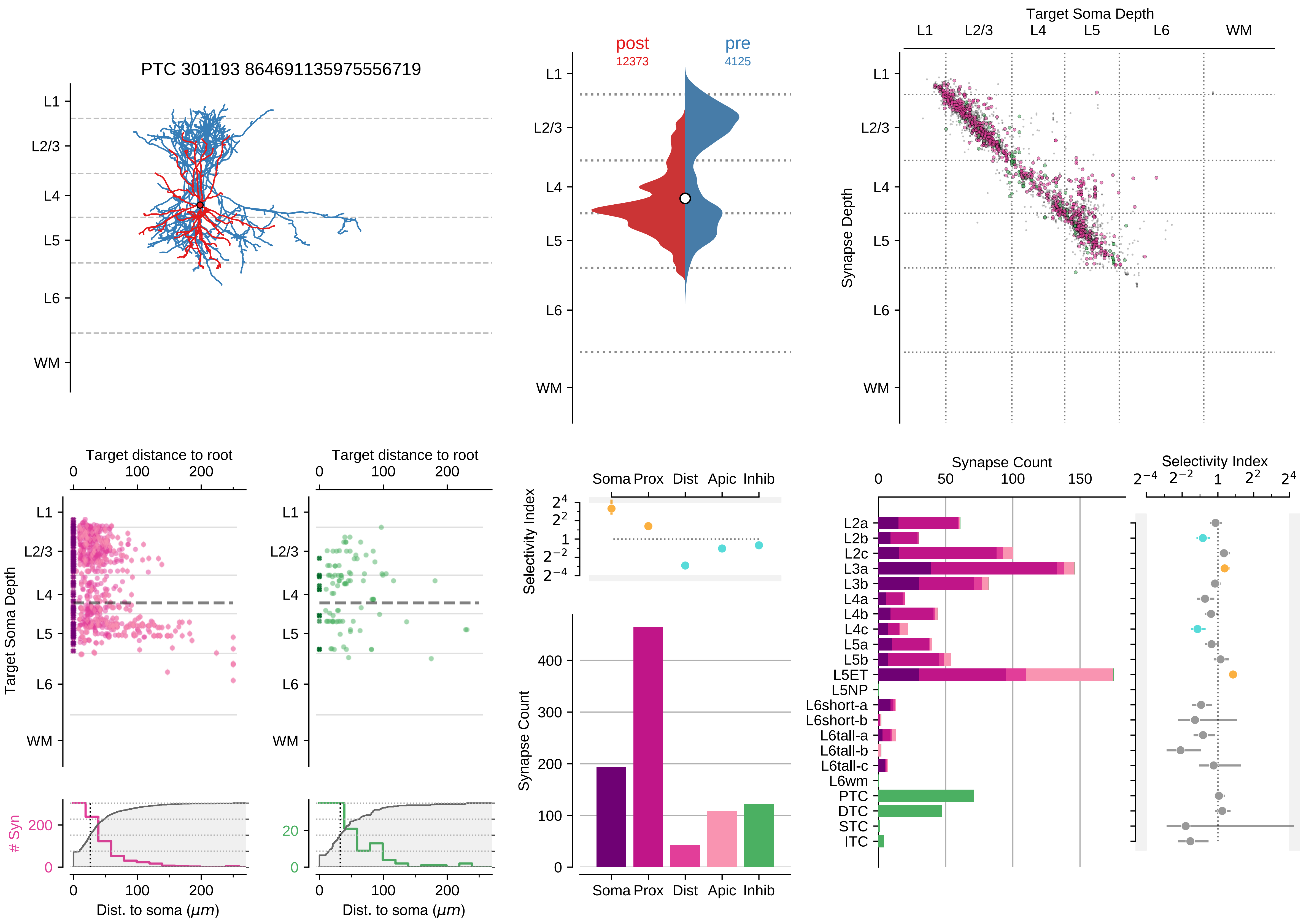

PTC 301193 864691135975556719
L1
L2/3
L4
L5
L6
WM
post
12373
pre
4125
Target Soma Depth
Synapse Depth
Target distance to root
Target Soma Depth
### Syn
Dist. to soma ($\mu m$)
Selectivity Index
Soma Prox Dist Apic Inhib
Synapse Count
L2a
L2b
L2c
L3a
L3b
L4a
L4b
L4c
L5a
L5b
L5ET
L5NP
L6short-a
L6short-b
L6tall-a
L6tall-b
L6tall-c
L6wm
PTC
DTC
STC
ITC

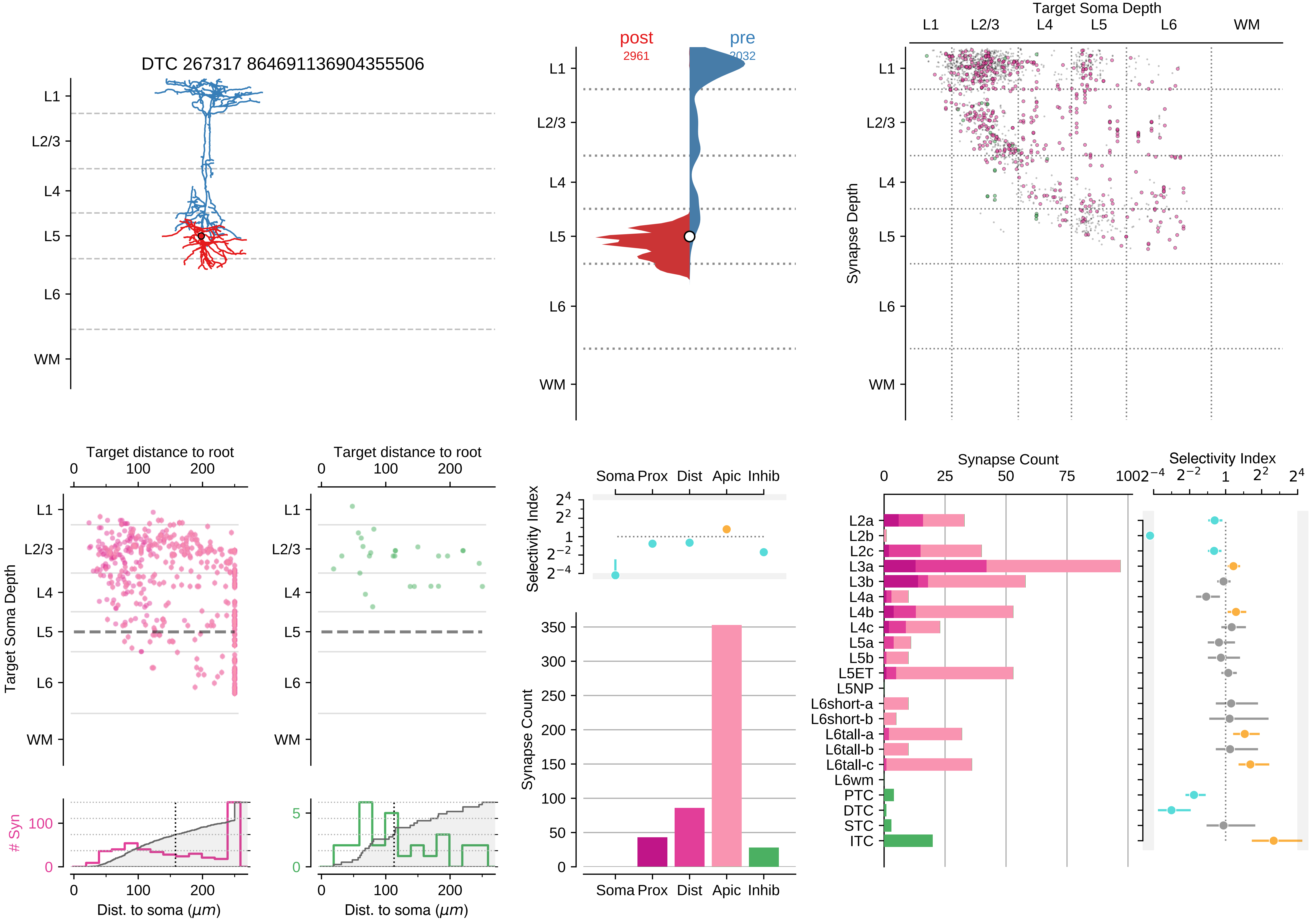
DTC 267317 864691136904355506
L1
L2/3
L4
L5
L6
WM
post
2961
pre
2032
Target Soma Depth
Synapse Depth
Target distance to root
Target Soma Depth
### Syn
Dist. to soma ($\mu m$)
Selectivity Index
Soma Prox Dist Apic Inhib
Synapse Count
L2a
L2b
L2c
L3a
L3b
L4a
L4b
L4c
L5a
L5b
L5ET
L5NP
L6short-a
L6short-b
L6tall-a
L6tall-b
L6tall-c
L6wm
PTC
DTC
STC
ITC

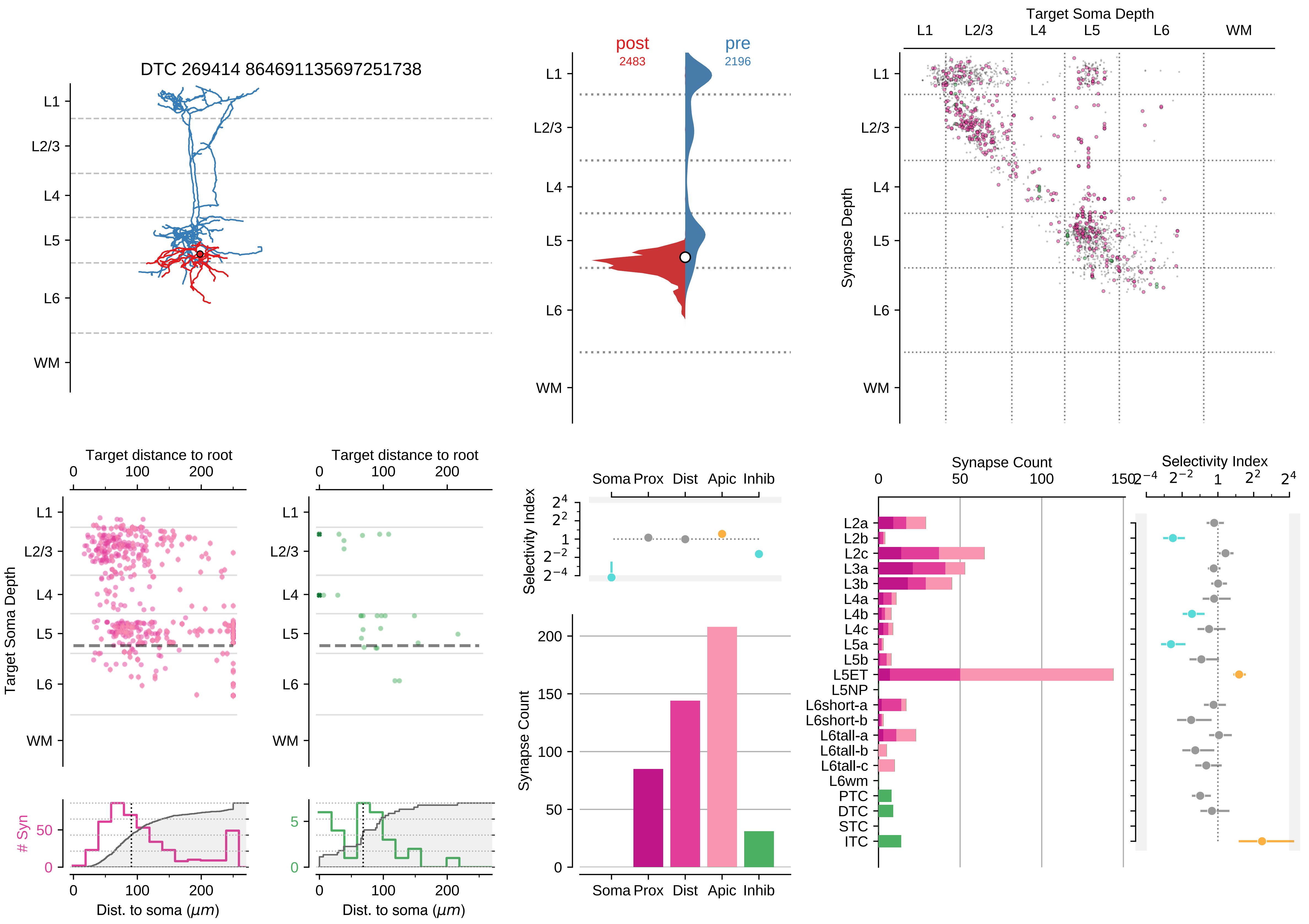
DTC 269414 864691135697251738
L1
L2/3
L4
L5
L6
WM
post
2483
pre
2196
Target Soma Depth
Synapse Depth
Target distance to root
Target Soma Depth
Dist. to soma (μm)
### Syn
Selectivity Index
Soma Prox Dist Apic Inhib
Synapse Count
L2a
L2b
L2c
L3a
L3b
L4a
L4b
L4c
L5a
L5b
L5ET
L5NP
L6short-a
L6short-b
L6tall-a
L6tall-b
L6tall-c
L6wm
PTC
DTC
STC
ITC

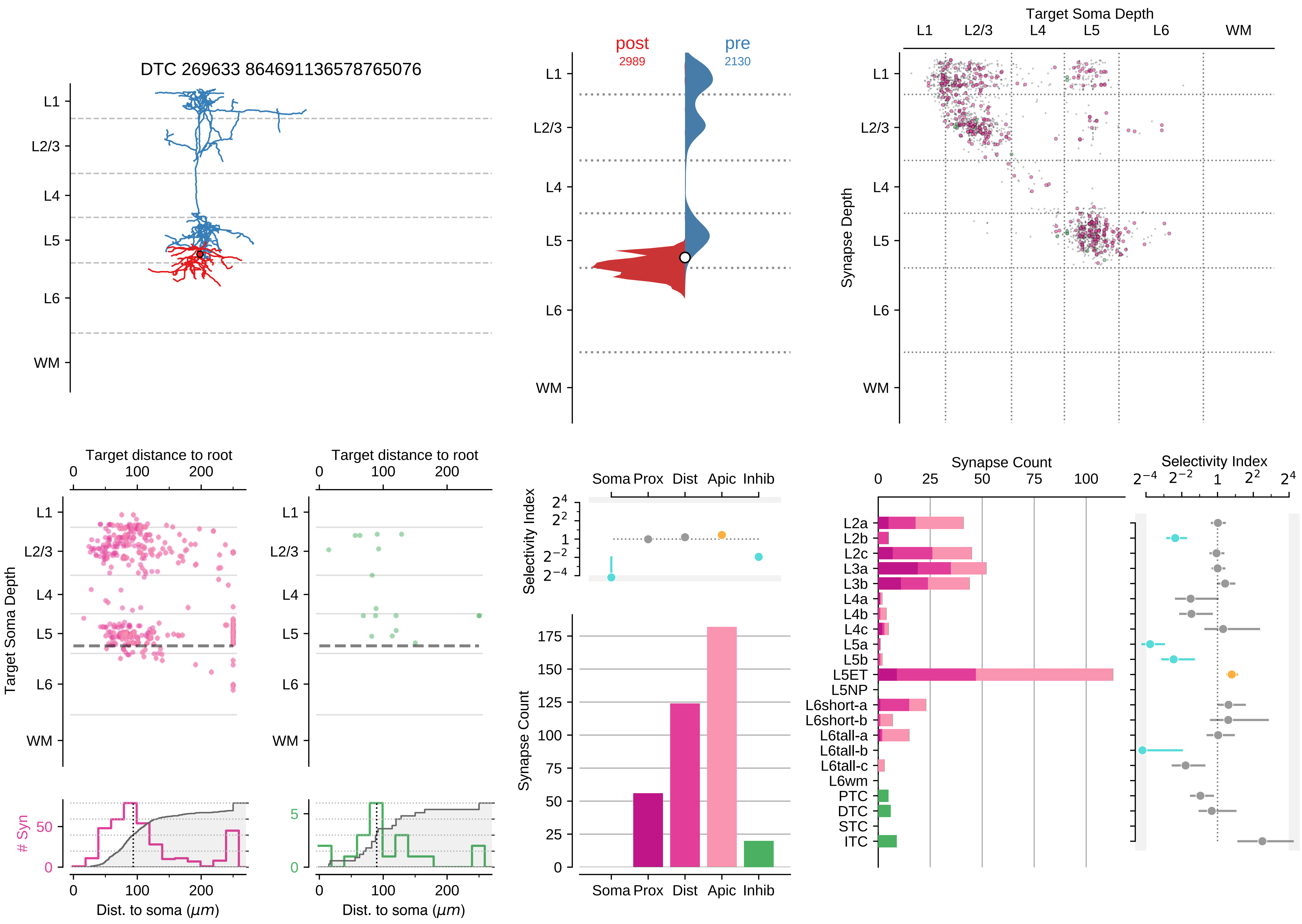
DTC 269633 864691136578765076
L1
L2/3
L4
L5
L6
WM
post
2989
pre
2130
Target Soma Depth
Synapse Depth
Target distance to root
Target Soma Depth
### Syn
Dist. to soma (μm)
Selectivity Index
Soma Prox
Dist
Apic
Inhib
Synapse Count
L2a
L2b
L2c
L3a
L3b
L4a
L4b
L4c
L5a
L5b
L5ET
L5NP
L6short-a
L6short-b
L6tall-a
L6tall-b
L6tall-c
L6wm
PTC
DTC
STC
ITC

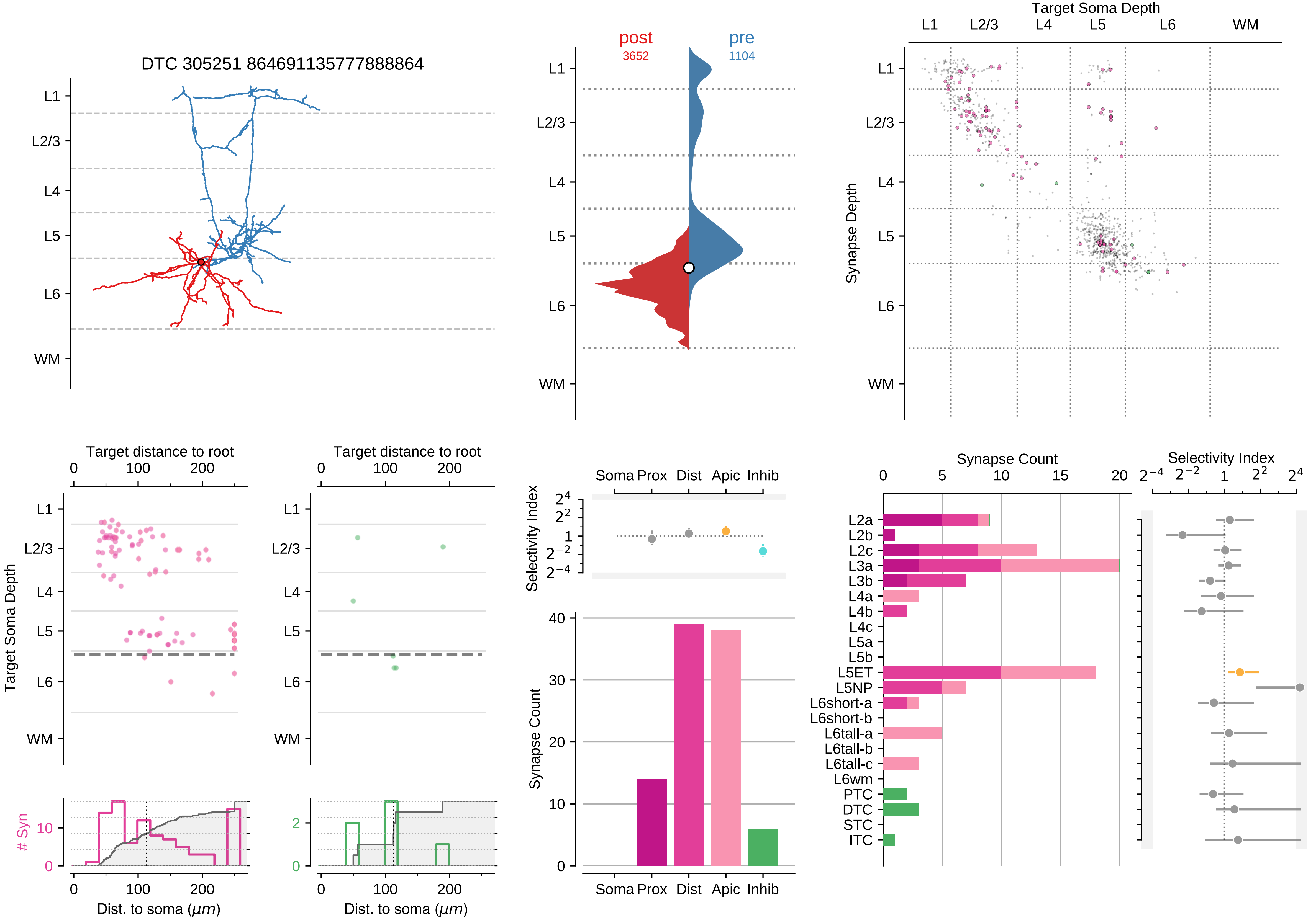
DTC 305251 864691135777888864
L1
L2/3
L4
L5
L6
WM
post
3652
pre
1104
Target Soma Depth
Synapse Depth
Target distance to root
Dist. to soma (μm)
### Syn
Selectivity Index
Synapse Count
Soma Prox
Dist
Apic
Inhib
L2a
L2b
L2c
L3a
L3b
L4a
L4b
L4c
L5a
L5b
L5ET
L5NP
L6short-a
L6short-b
L6tall-a
L6tall-b
L6tall-c
L6wm
PTC
DTC
STC
ITC

### Motif Group 7

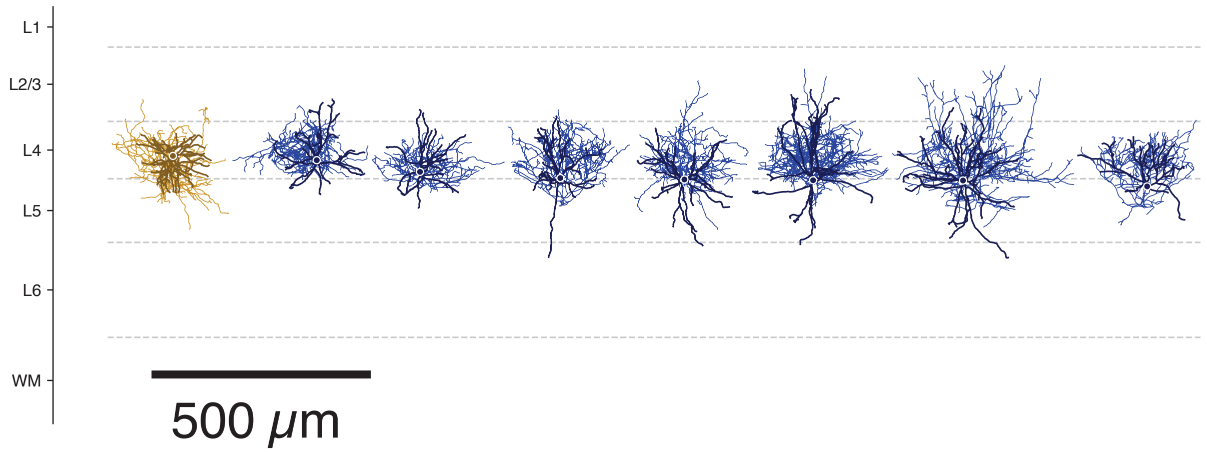

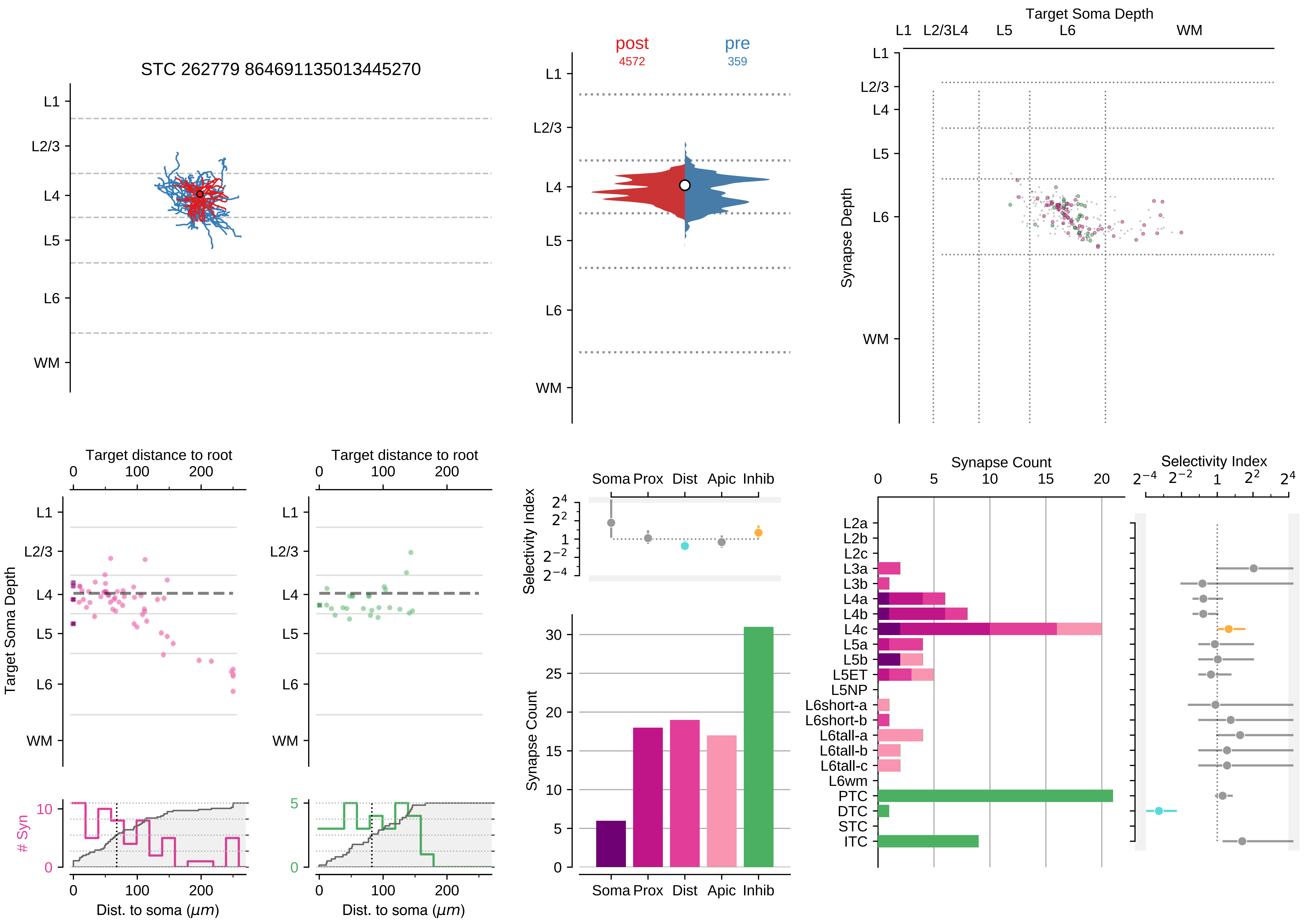
STC 262779 864691135013445270
L1
L2/3
L4
L5
L6
WM
post
4572
pre
359
Target Soma Depth
Synapse Depth
Target distance to root
Target Soma Depth
### Syn
Dist. to soma ($\mu m$)
Selectivity Index
Soma Prox Dist Apic Inhib
Synapse Count
L2a
L2b
L2c
L3a
L3b
L4a
L4b
L4c
L5a
L5b
L5ET
L5NP
L6short-a
L6short-b
L6tall-a
L6tall-b
L6tall-c
L6wm
PTC
DTC
STC
ITC

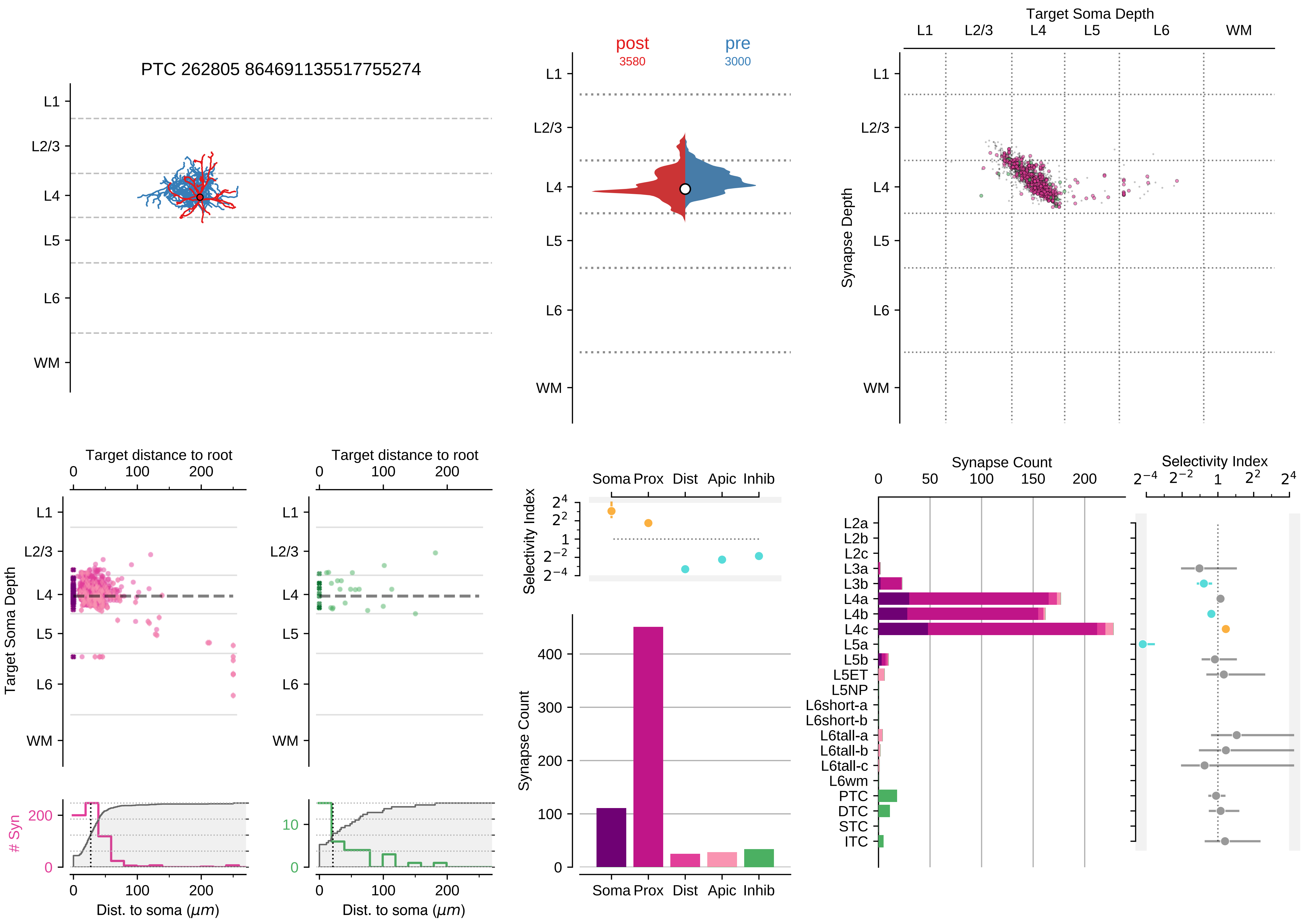
PTC 262805 864691135517755274
L1
L2/3
L4
L5
L6
WM
post
3580
pre
3000
Target Soma Depth
Synapse Depth
Target distance to root
Target Soma Depth
### Syn
Dist. to soma (μm)
Selectivity Index
Soma Prox Dist Apic Inhib
Synapse Count
L2a
L2b
L2c
L3a
L3b
L4a
L4b
L4c
L5a
L5b
L5ET
L5NP
L6short-a
L6short-b
L6tall-a
L6tall-b
L6tall-c
L6wm
PTC
DTC
STC
ITC

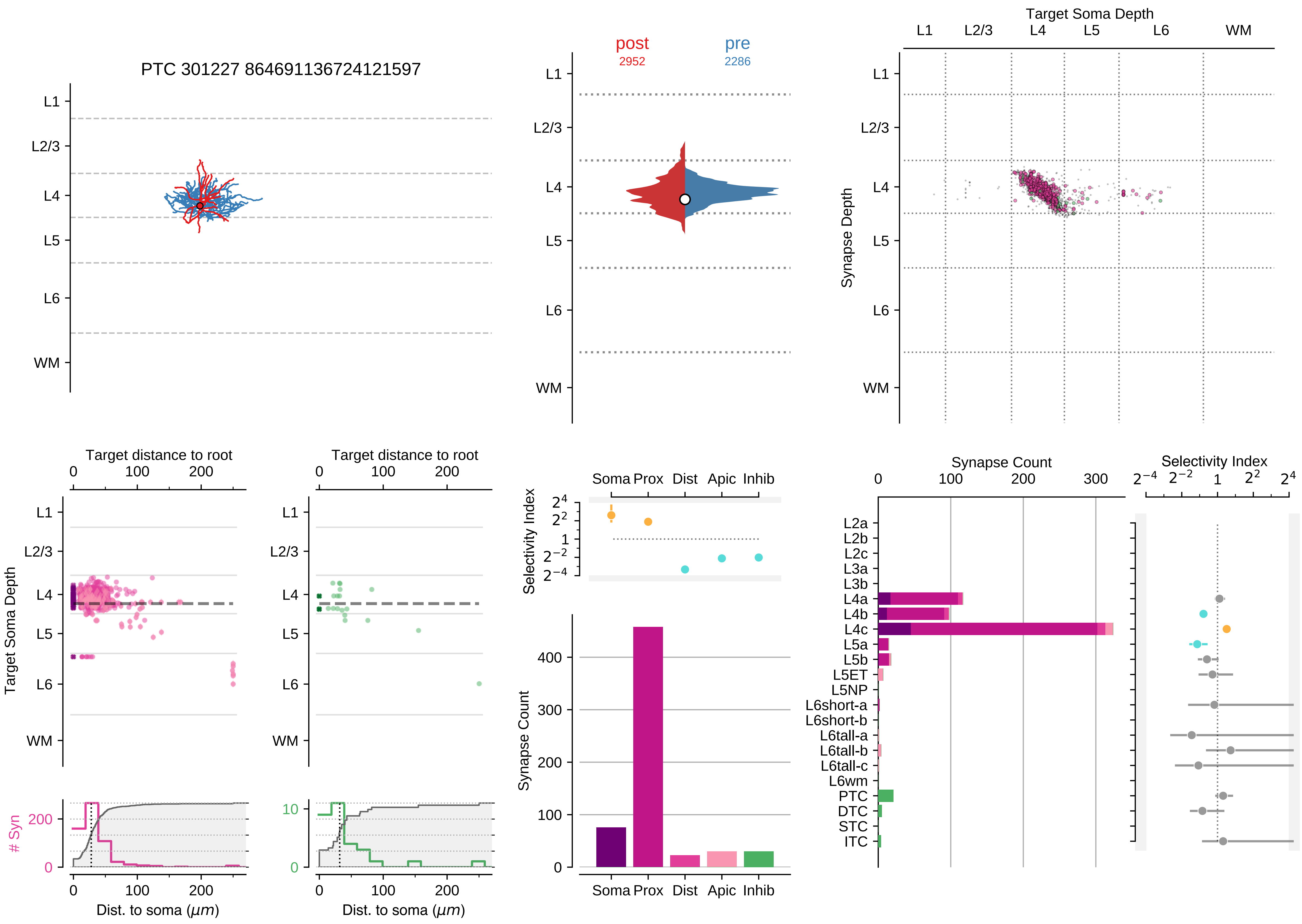

PTC 301227 864691136724121597
L1
L2/3
L4
L5
L6
WM
post
2952
pre
2286
Target Soma Depth
Synapse Depth
Target distance to root
0
100
200
Target Soma Depth
### Syn
Dist. to soma ($\mu m$)
Selectivity Index
Soma Prox Dist Apic Inhib
Synapse Count
400
300
Synapse Count
Selectivity Index
L2a
L2b
L2c
L3a
L3b
L4a
L4b
L4c
L5a
L5b
L5ET
L5NP
L6short-a
L6short-b
L6tall-a
L6tall-b
L6tall-c
L6wm
PTC
DTC
STC
ITC

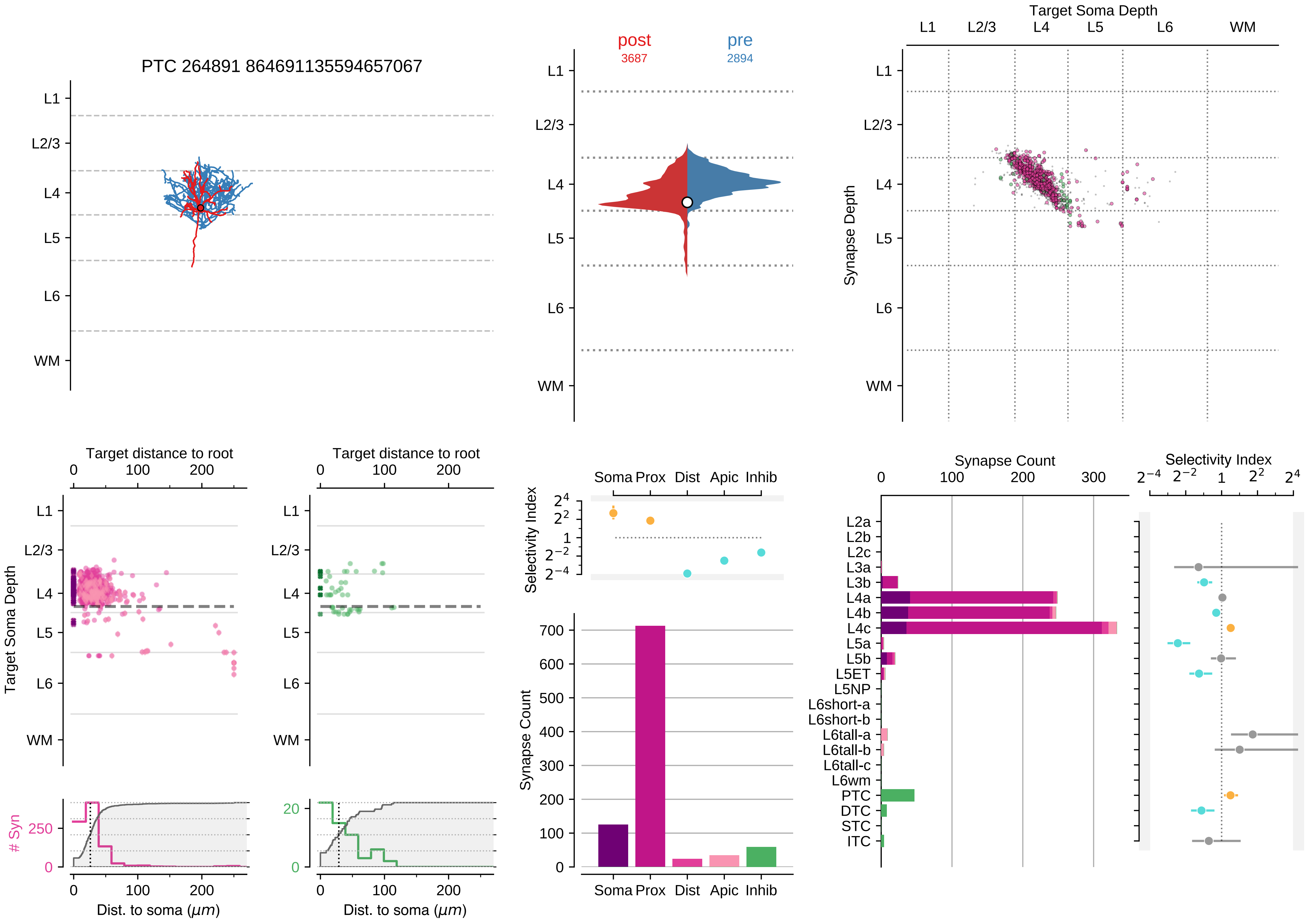

PTC 264891 864691135594657067
L1
L2/3
L4
L5
L6
WM
post
3687
pre
2894
Target Soma Depth
Synapse Depth
Target distance to root
Target Soma Depth
### Syn
Dist. to soma ($\mu m$)
Selectivity Index
Soma Prox Dist Apic Inhib
Synapse Count
Synapse Count
Selectivity Index
L2a
L2b
L2c
L3a
L3b
L4a
L4b
L4c
L5a
L5b
L5ET
L5NP
L6short-a
L6short-b
L6tall-a
L6tall-b
L6tall-c
L6wm
PTC
DTC
STC
ITC

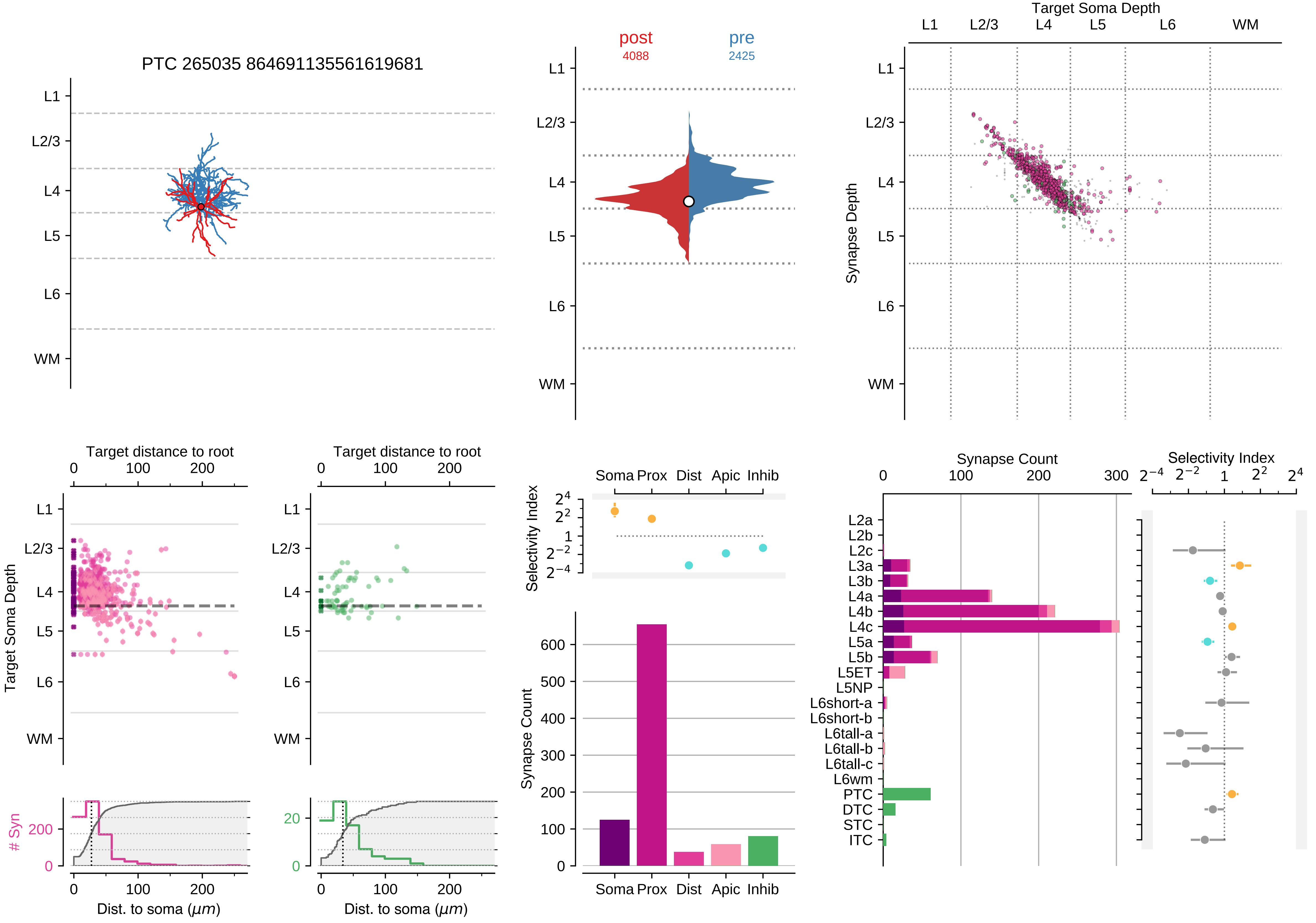

PTC 265035 864691135561619681
L1
L2/3
L4
L5
L6
WM
post
4088
pre
2425
Target Soma Depth
Synapse Depth
Target distance to root
0
100
200
Target Soma Depth
### Syn
Dist. to soma ($\mu m$)
Selectivity Index
Soma
Prox
Dist
Apic
Inhib
Synapse Count
Synapse Count
0
100
200
300
Selectivity Index
$2^{-4}$
$2^{-2}$
1
$2^{2}$
$2^{4}$
L2a
L2b
L2c
L3a
L3b
L4a
L4b
L4c
L5a
L5b
L5ET
L5NP
L6short-a
L6short-b
L6tall-a
L6tall-b
L6tall-c
L6wm
PTC
DTC
STC
ITC

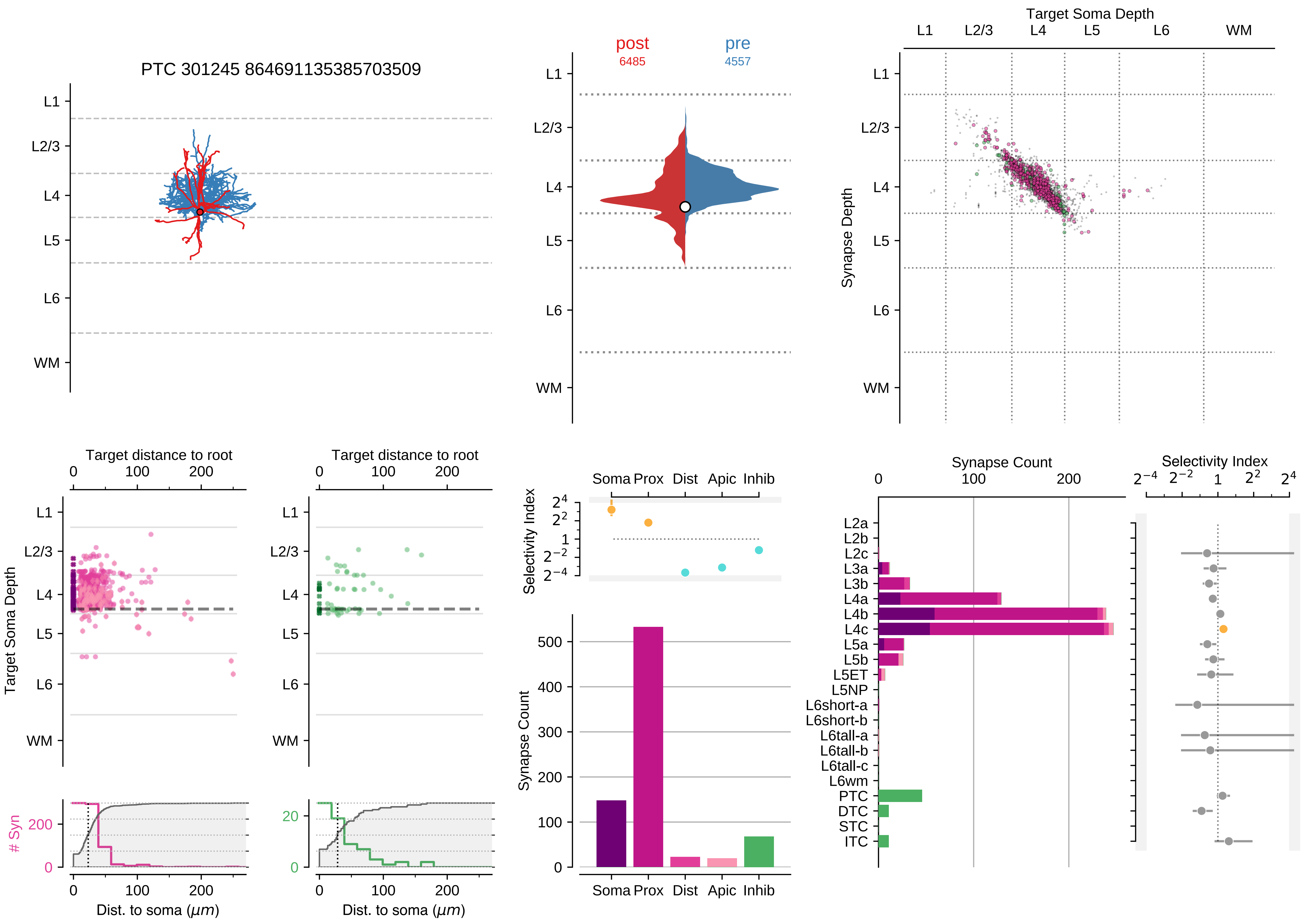

PTC 301245 864691135385703509
L1
L2/3
L4
L5
L6
WM
post
6485
pre
4557
Target Soma Depth
Synapse Depth
Target distance to root
Target Soma Depth
Dist. to soma (μm)
### Syn
Selectivity Index
Soma
Prox
Dist
Apic
Inhib
Synapse Count
L2a
L2b
L2c
L3a
L3b
L4a
L4b
L4c
L5a
L5b
L5ET
L5NP
L6short-a
L6short-b
L6tall-a
L6tall-b
L6tall-c
L6wm
PTC
DTC
STC
ITC

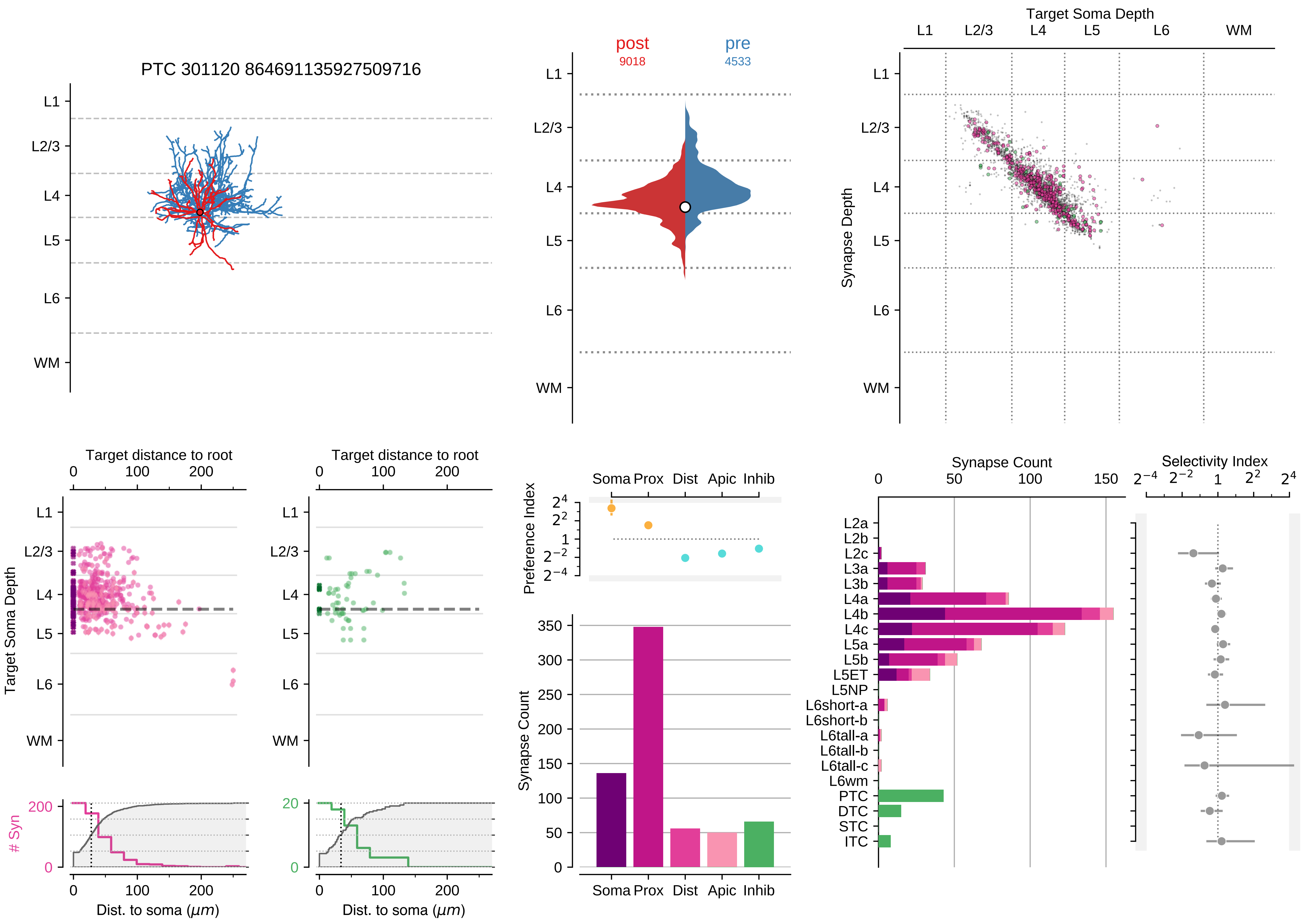

PTC 301120 864691135927509716
post
9018
pre
4533
Target Soma Depth
Synapse Depth
L1
L2/3
L4
L5
L6
WM
Target distance to root
Target Soma Depth
### Syn
Dist. to soma ($\mu m$)
Preference Index
Soma
Prox
Dist
Apic
Inhib
Synapse Count
Selectivity Index
L2a
L2b
L2c
L3a
L3b
L4a
L4b
L4c
L5a
L5b
L5ET
L5NP
L6short-a
L6short-b
L6tall-a
L6tall-b
L6tall-c
L6wm
PTC
DTC
STC
ITC

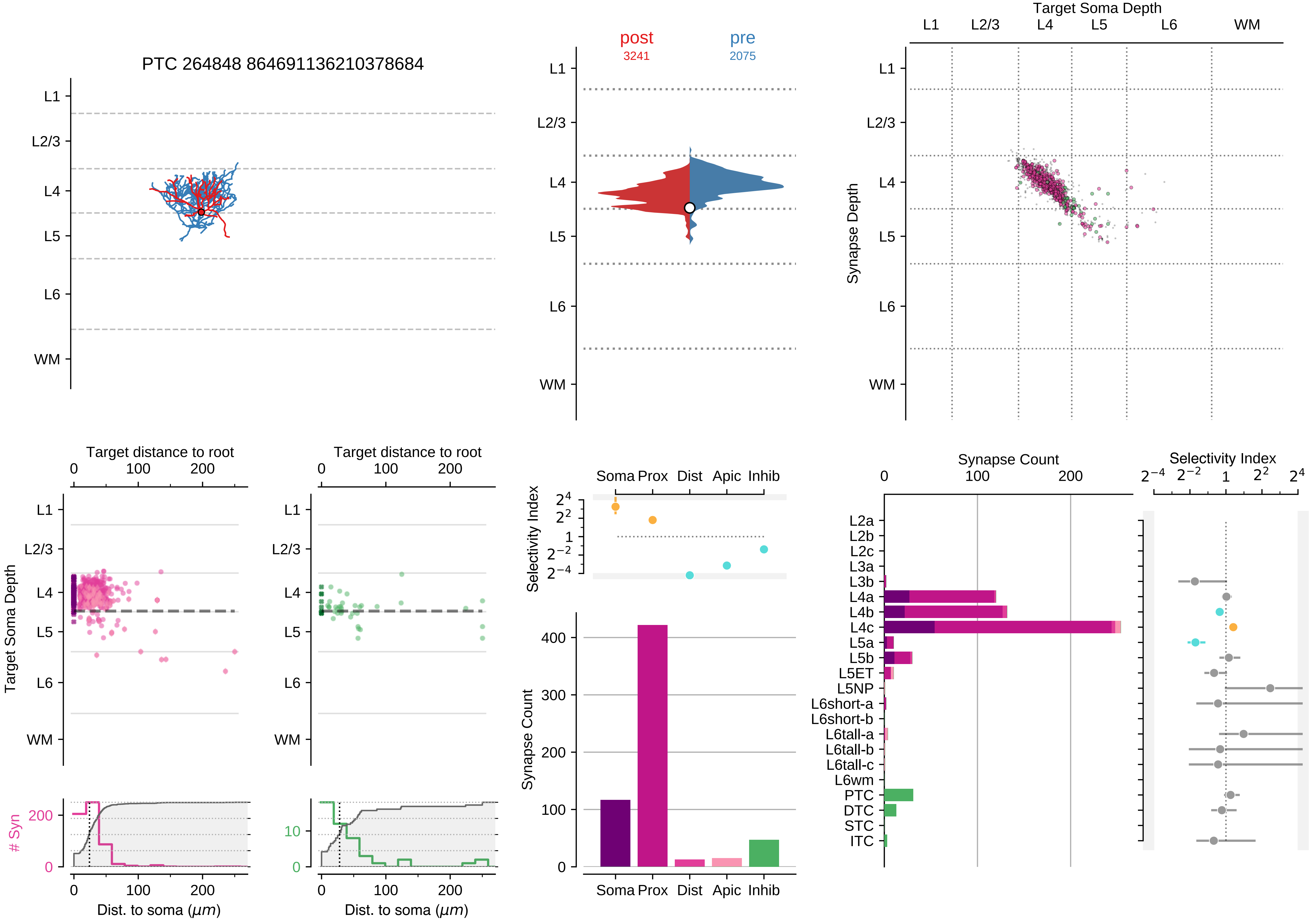
PTC 264848 864691136210378684
L1
L2/3
L4
L5
L6
WM
post
3241
pre
2075
Target Soma Depth
Synapse Depth
Target distance to root
Dist. to soma ($\mu m$)
### Syn
Selectivity Index
Soma
Prox
Dist
Apic
Inhib
Synapse Count
L2a
L2b
L2c
L3a
L3b
L4a
L4b
L4c
L5a
L5b
L5ET
L5NP
L6short-a
L6short-b
L6tall-a
L6tall-b
L6tall-c
L6wm
PTC
DTC
STC
ITC

### Motif Group 8

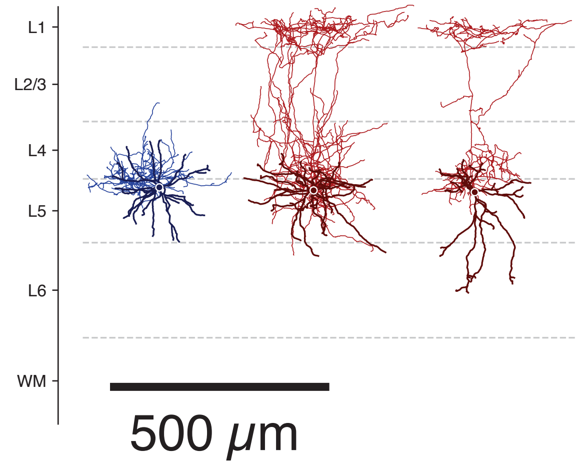

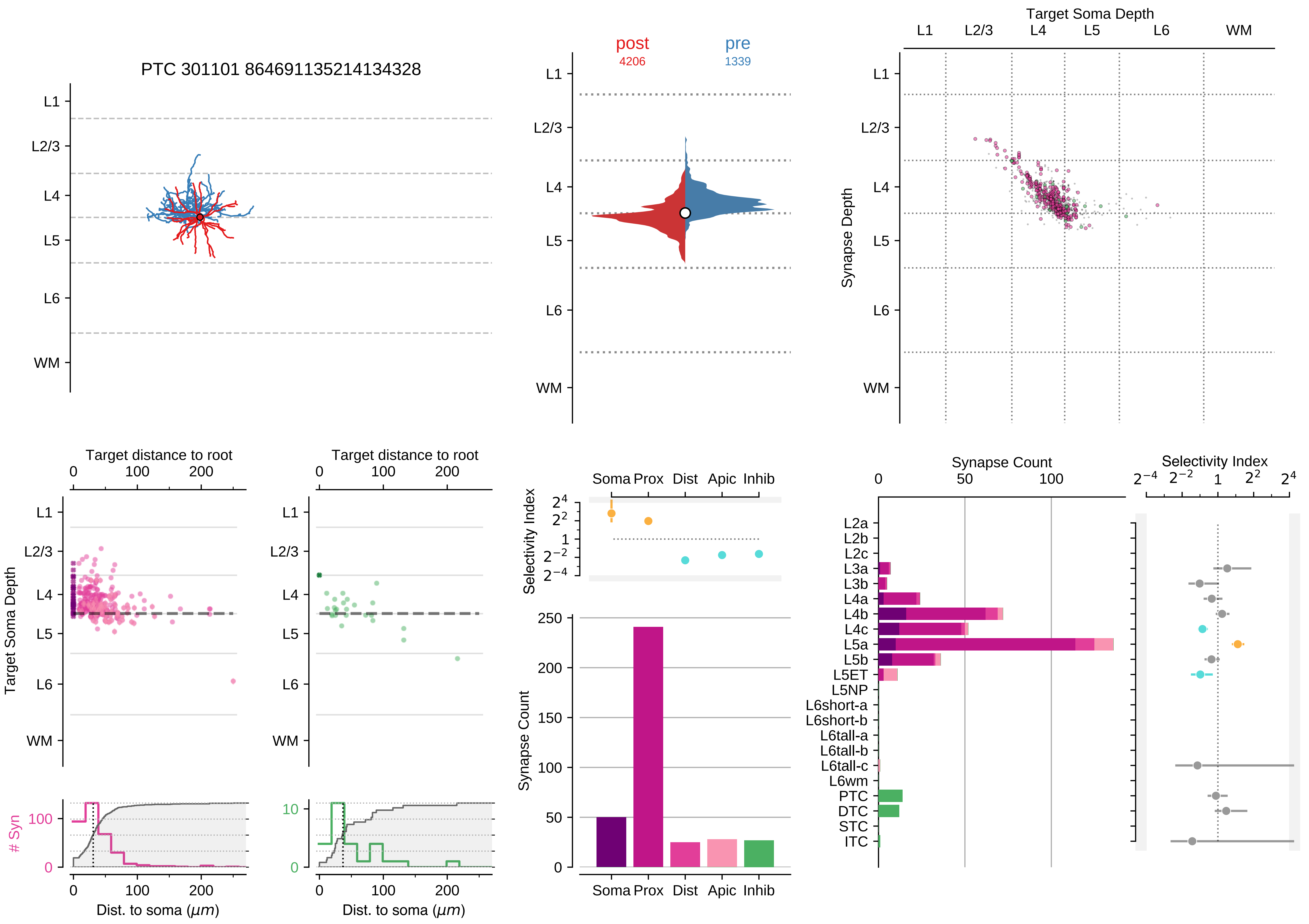

PTC 301101 864691135214134328
L1
L2/3
L4
L5
L6
WM
post
4206
pre
1339
Target Soma Depth
Synapse Depth
Target distance to root
Dist. to soma ($\mu m$)
### Syn
Selectivity Index
Synapse Count
Soma
Prox
Dist
Apic
Inhib
L2a
L2b
L2c
L3a
L3b
L4a
L4b
L4c
L5a
L5b
L5ET
L5NP
L6short-a
L6short-b
L6tall-a
L6tall-b
L6tall-c
L6wm
PTC
DTC
STC
ITC

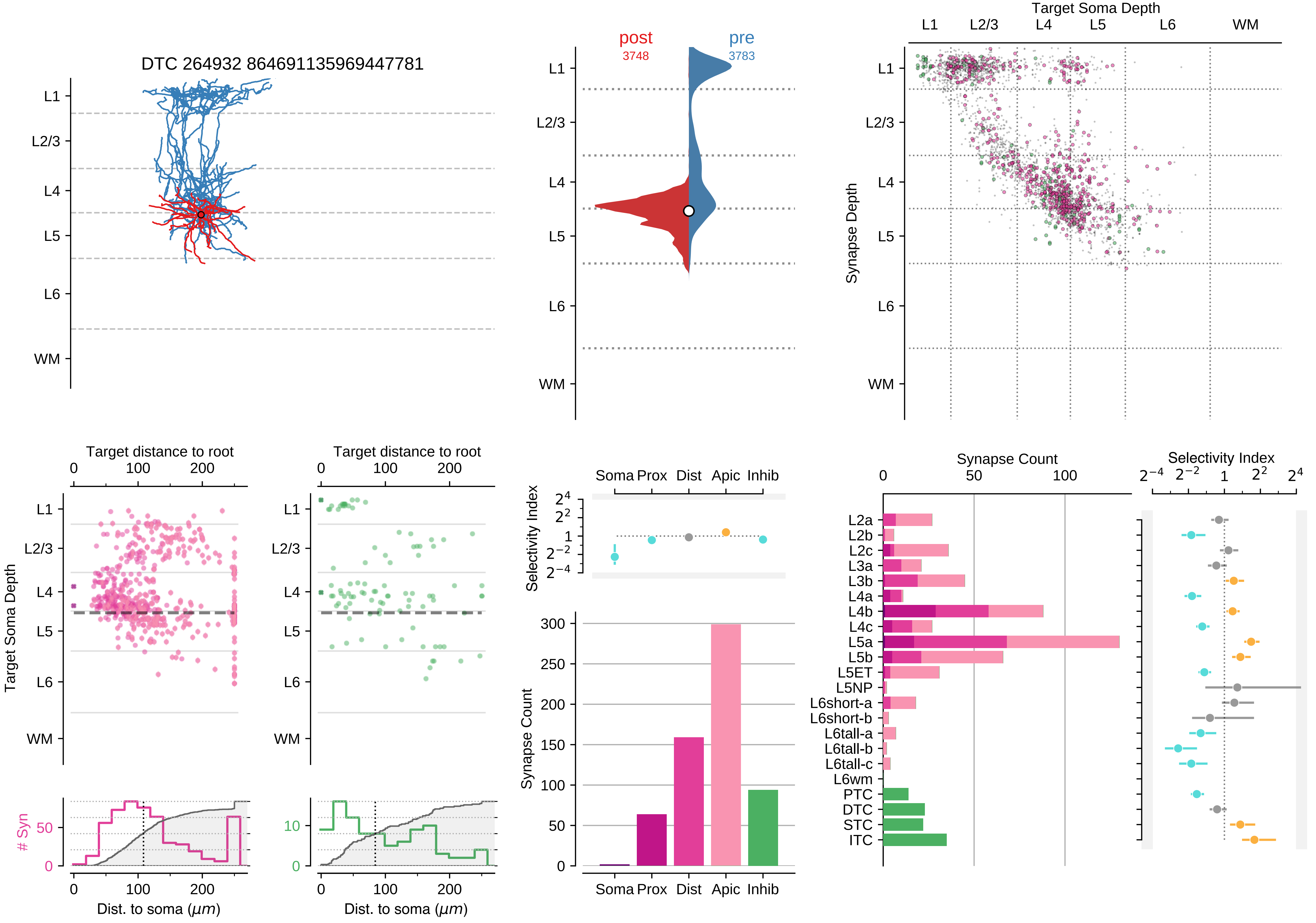

DTC 264932 864691135969447781
post
3748
pre
3783
Target Soma Depth
Synapse Depth
L1
L2/3
L4
L5
L6
WM
Target distance to root
Target Soma Depth
### Syn
Dist. to soma ($\mu m$)
Selectivity Index
Soma Prox Dist Apic Inhib
Synapse Count
L2a
L2b
L2c
L3a
L3b
L4a
L4b
L4c
L5a
L5b
L5ET
L5NP
L6short-a
L6short-b
L6tall-a
L6tall-b
L6tall-c
L6wm
PTC
DTC
STC
ITC

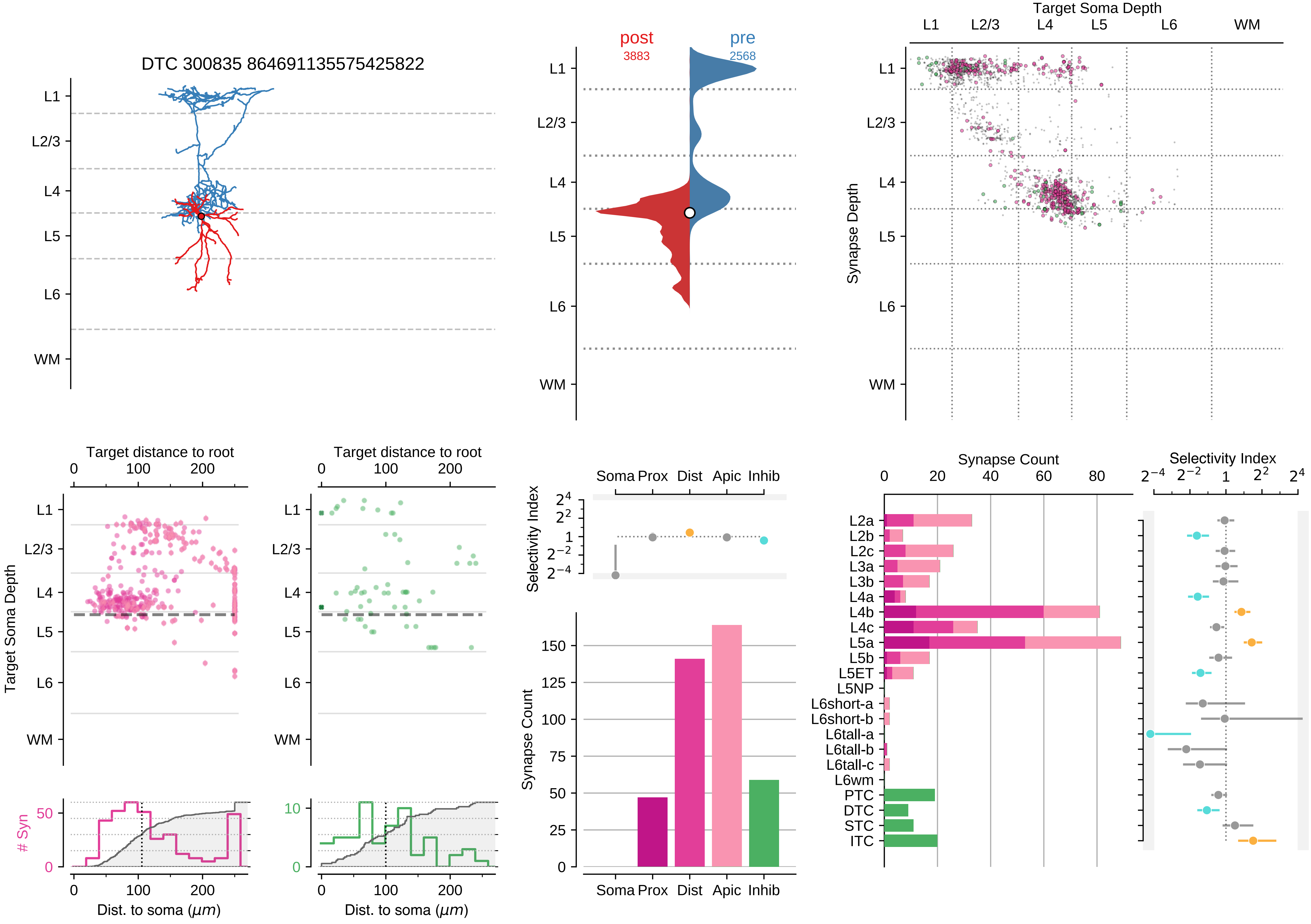

DTC 300835 864691135575425822
post
3883
pre
2568
Target Soma Depth
L1
L2/3
L4
L5
L6
WM
Synapse Depth
Target distance to root
Target Soma Depth
### Syn
Dist. to soma (μm)
Soma Prox Dist Apic Inhib
Selectivity Index
Synapse Count
L2a
L2b
L2c
L3a
L3b
L4a
L4b
L4c
L5a
L5b
L5ET
L5NP
L6short-a
L6short-b
L6tall-a
L6tall-b
L6tall-c
L6wm
PTC
DTC
STC
ITC

### Motif Group 9

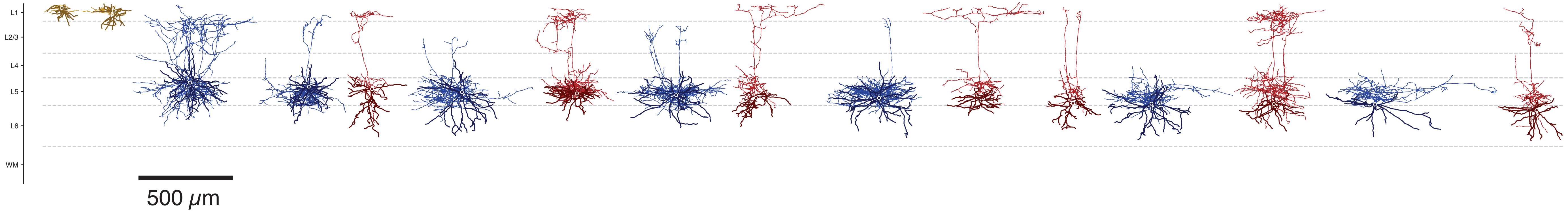

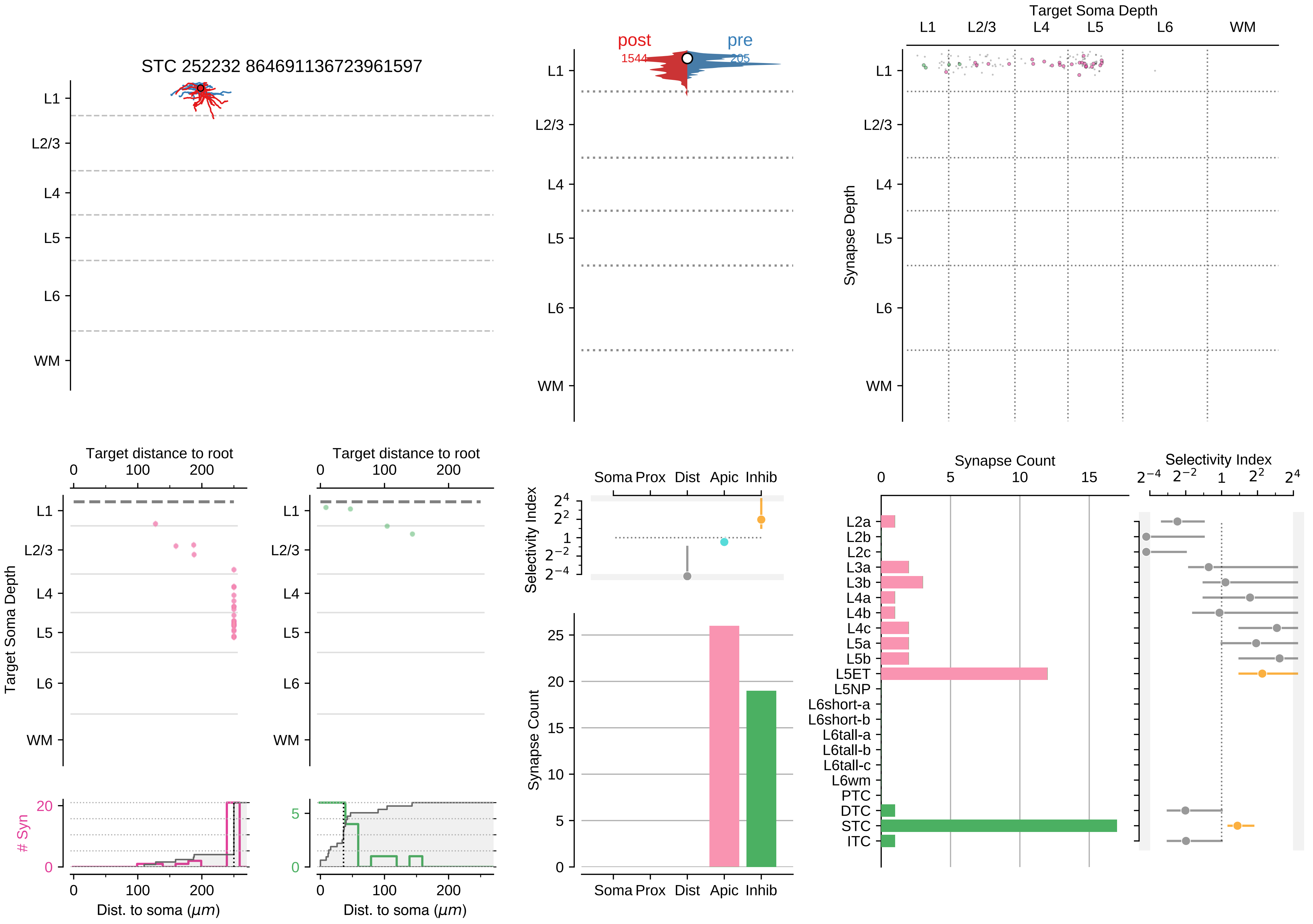
STC 252232 864691136723961597
L1
L2/3
L4
L5
L6
WM
post
1544
pre
205
Target Soma Depth
Synapse Depth
Target distance to root
Target Soma Depth
Dist. to soma (μm)
### Syn
Selectivity Index
Soma Prox Dist Apic Inhib
Synapse Count
L2a
L2b
L2c
L3a
L3b
L4a
L4b
L4c
L5a
L5b
L5ET
L5NP
L6short-a
L6short-b
L6tall-a
L6tall-b
L6tall-c
L6wm
PTC
DTC
STC
ITC

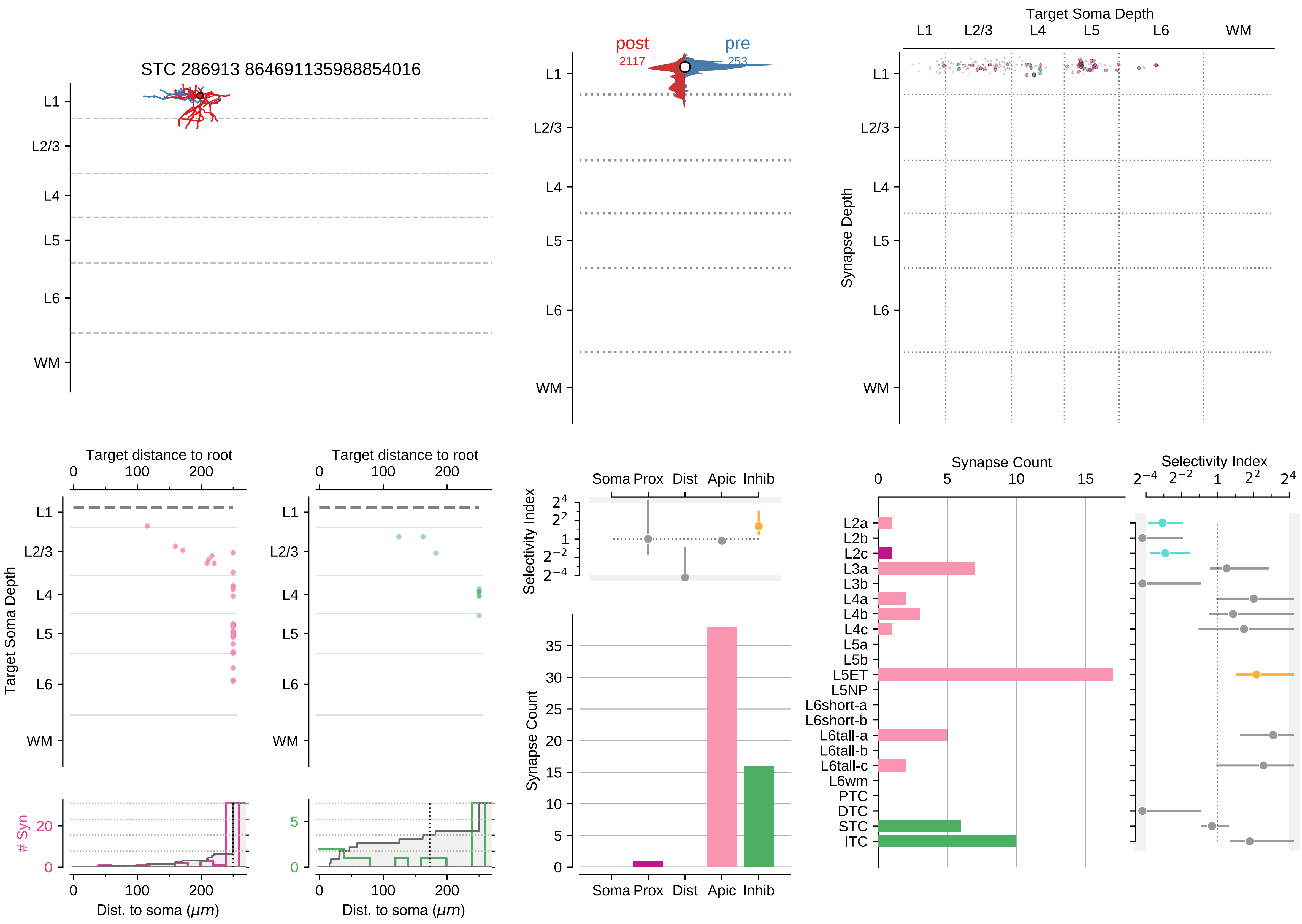
STC 286913 864691135988854016
L1
L2/3
L4
L5
L6
WM
post
2117
pre
253
Target Soma Depth
Synapse Depth
Target distance to root
Target Soma Depth
### Syn
Dist. to soma (μm)
Selectivity Index
Soma Prox Dist Apic Inhib
Synapse Count
Synapse Count
Selectivity Index
L2a
L2b
L2c
L3a
L3b
L4a
L4b
L4c
L5a
L5b
L5ET
L5NP
L6short-a
L6short-b
L6tall-a
L6tall-b
L6tall-c
L6wm
PTC
DTC
STC
ITC

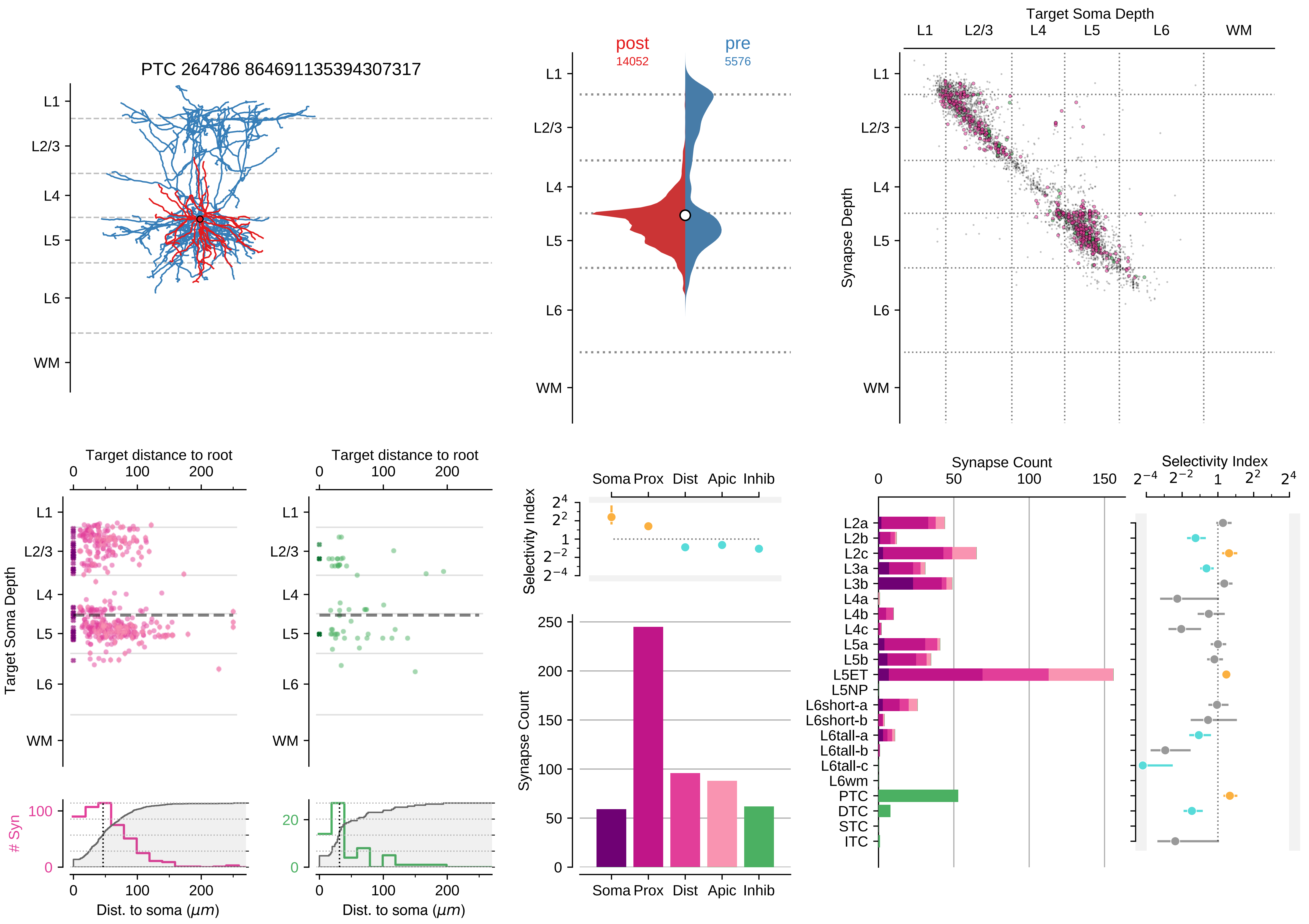

PTC 264786 864691135394307317
L1
L2/3
L4
L5
L6
WM
post
14052
pre
5576
Target Soma Depth
Synapse Depth
Target distance to root
Target Soma Depth
### Syn
Dist. to soma ($\mu m$)
Soma
Prox
Dist
Apic
Inhib
Selectivity Index
Synapse Count
L2a
L2b
L2c
L3a
L3b
L4a
L4b
L4c
L5a
L5b
L5ET
L5NP
L6short-a
L6short-b
L6tall-a
L6tall-b
L6tall-c
L6wm
PTC
DTC
STC
ITC

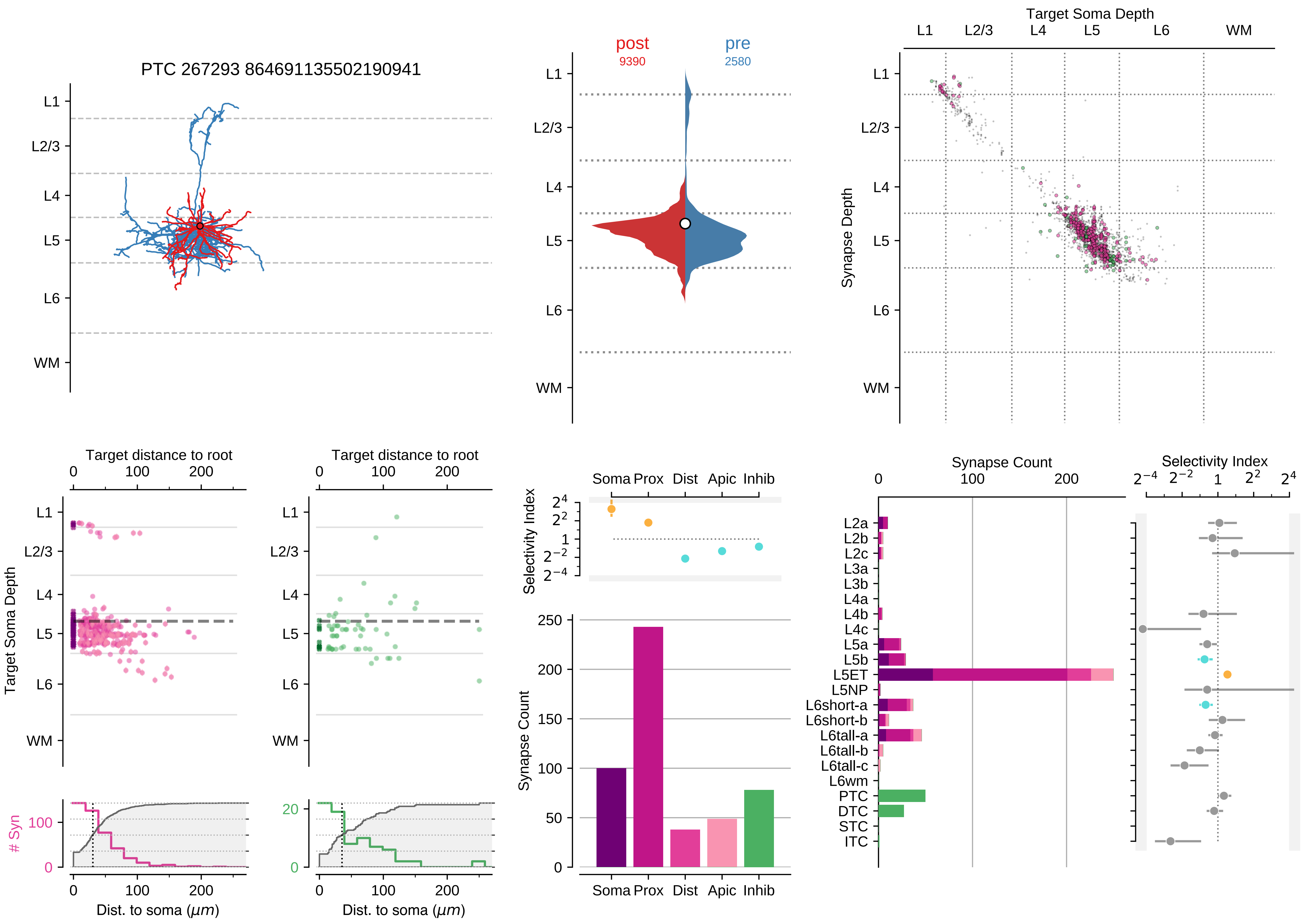

PTC 267293 864691135502190941
L1
L2/3
L4
L5
L6
WM
post
9390
pre
2580
Target Soma Depth
Synapse Depth
Target distance to root
Target Soma Depth
### Syn
Dist. to soma (μm)
Selectivity Index
Synapse Count
Soma
Prox
Dist
Apic
Inhib
L2a
L2b
L2c
L3a
L3b
L4a
L4b
L4c
L5a
L5b
L5ET
L5NP
L6short-a
L6short-b
L6tall-a
L6tall-b
L6tall-c
L6wm
PTC
DTC
STC
ITC

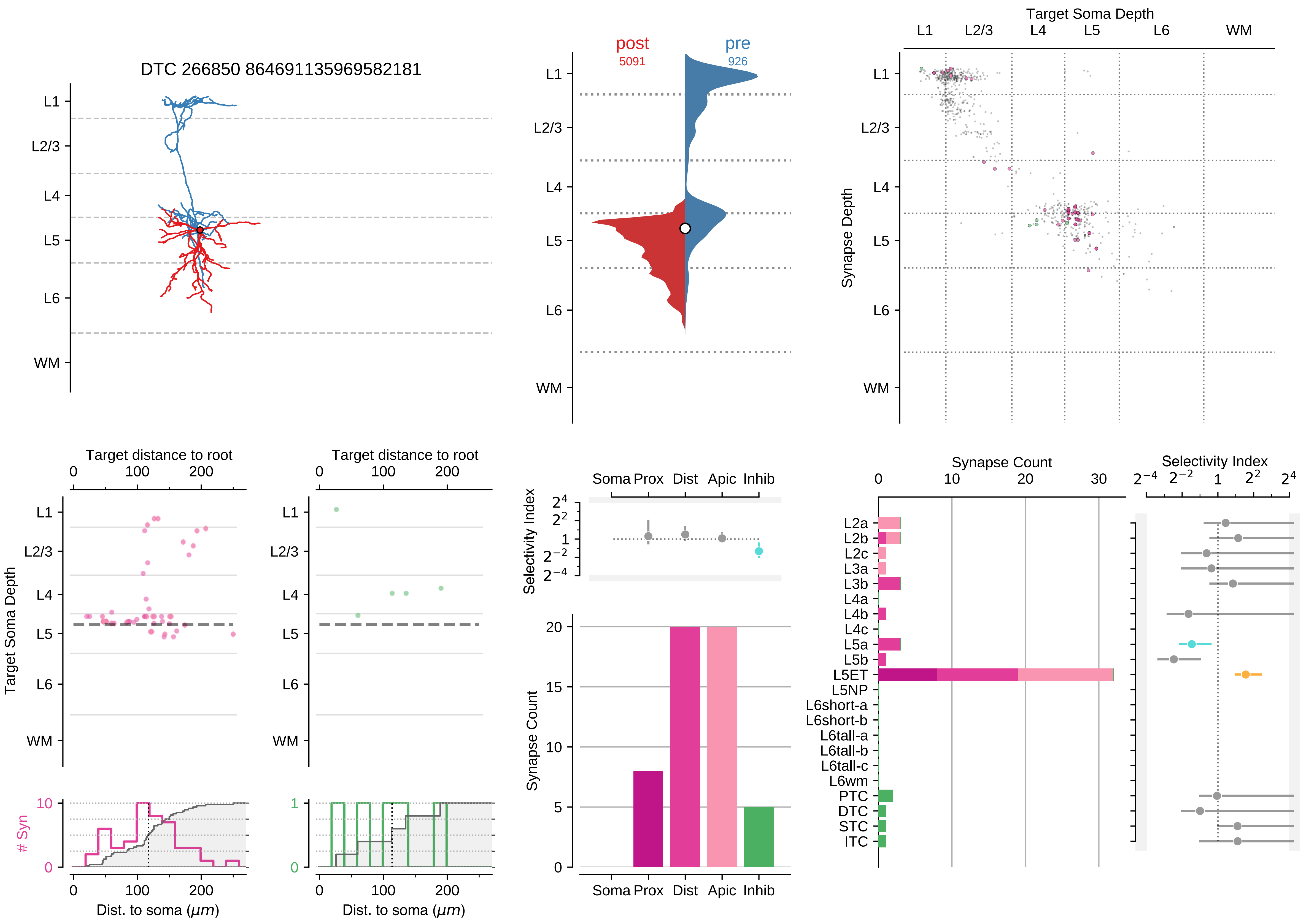

DTC 266850 864691135969582181
L1
L2/3
L4
L5
L6
WM
post
5091
pre
926
Target Soma Depth
Synapse Depth
Target distance to root
0
100
200
Target Soma Depth
### Syn
10
0
1
Dist. to soma ($\mu m$)
Selectivity Index
Soma Prox
Dist
Apic
Inhib
$2^4$
$2^2$
1
$2^{-2}$
$2^{-4}$
Synapse Count
20
15
10
5
0
30
L2a
L2b
L2c
L3a
L3b
L4a
L4b
L4c
L5a
L5b
L5ET
L5NP
L6short-a
L6short-b
L6tall-a
L6tall-b
L6tall-c
L6wm
PTC
DTC
STC
ITC

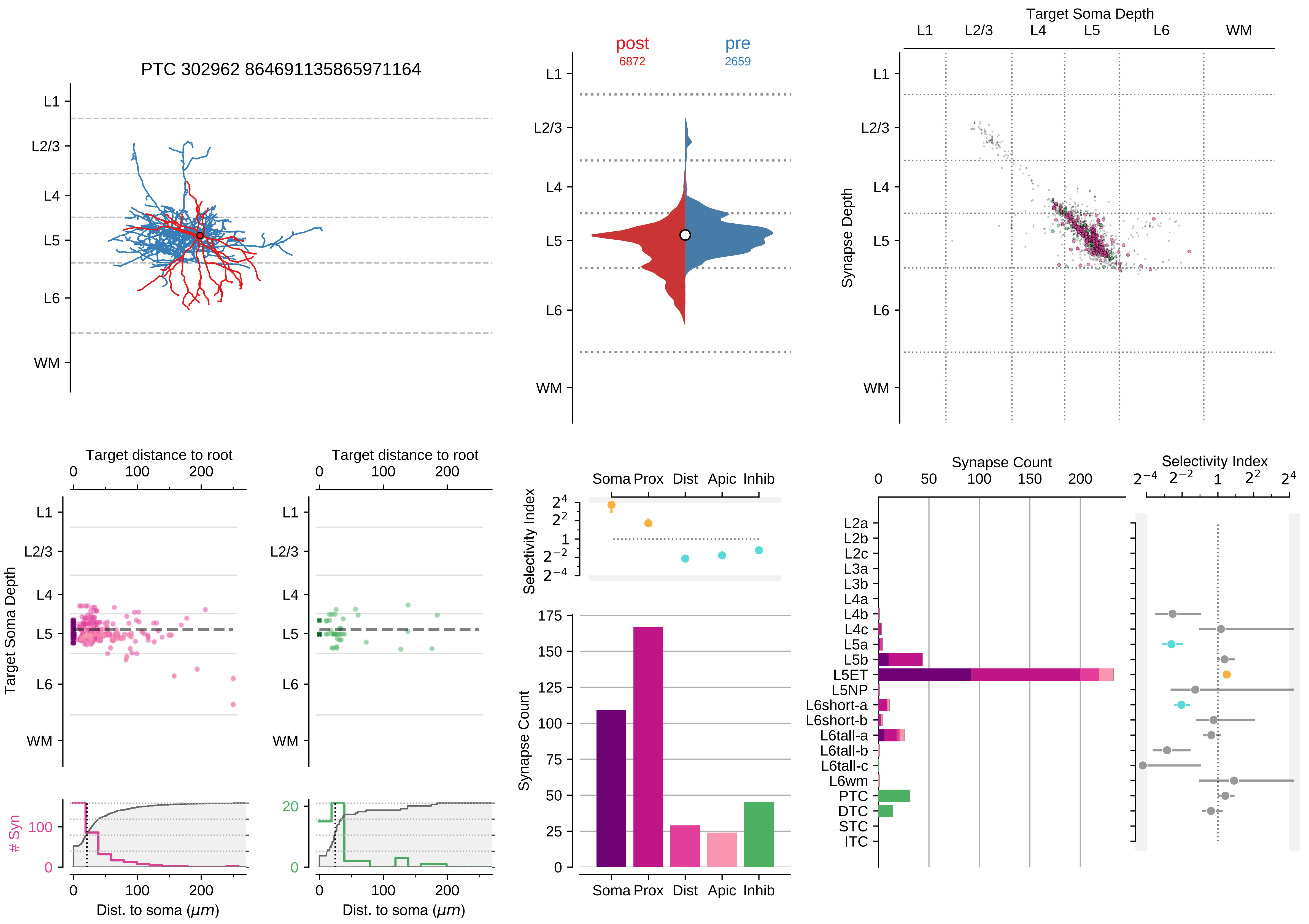
PTC 302962 864691135865971164
L1
L2/3
L4
L5
L6
WM
post
6872
pre
2659
Target Soma Depth
Synapse Depth
Target distance to root
Target Soma Depth
### Syn
Dist. to soma ($\mu m$)
Selectivity Index
Soma Prox Dist Apic Inhib
Synapse Count
L2a
L2b
L2c
L3a
L3b
L4a
L4b
L4c
L5a
L5b
L5ET
L5NP
L6short-a
L6short-b
L6tall-a
L6tall-b
L6tall-c
L6wm
PTC
DTC
STC
ITC

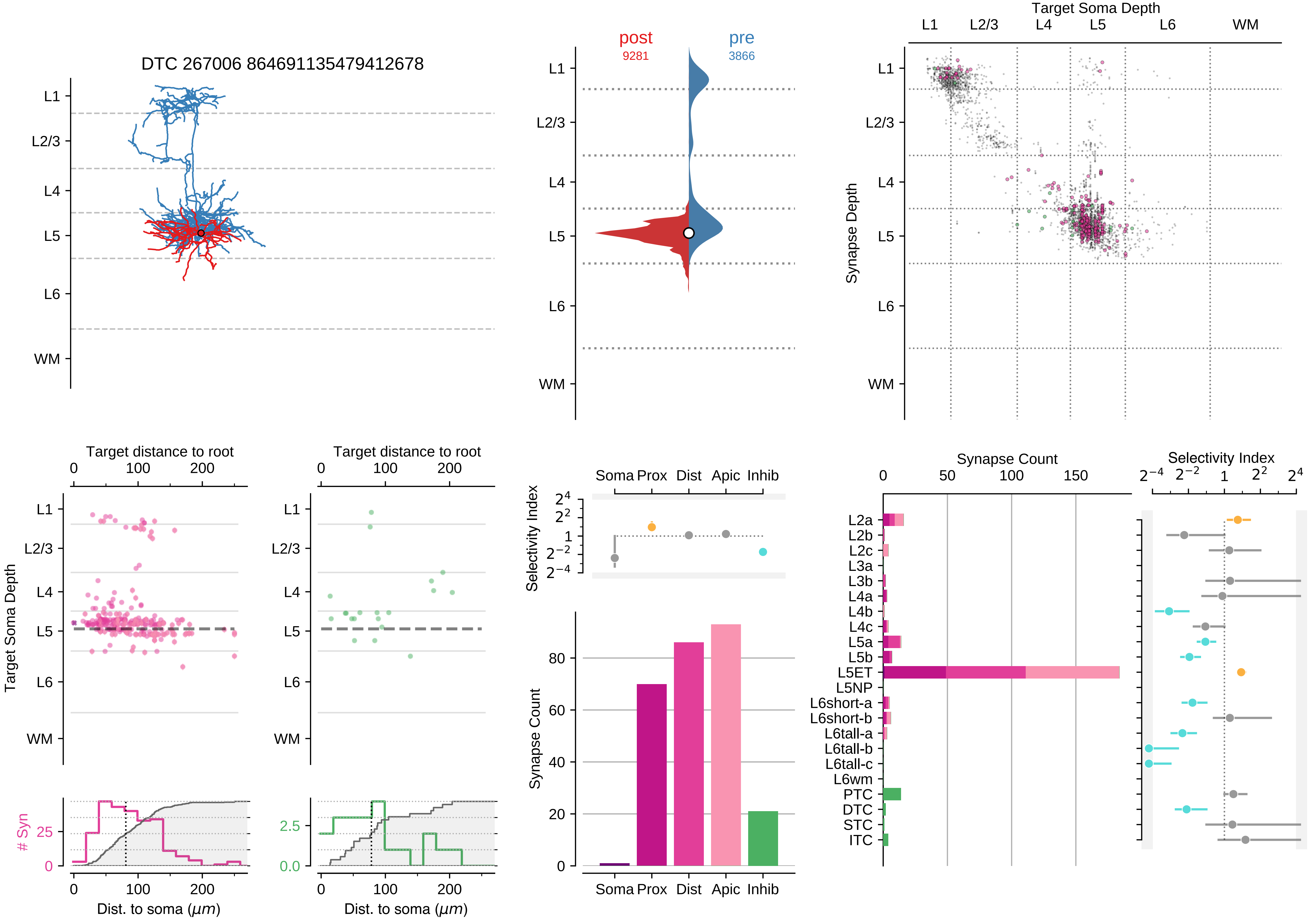

DTC 267006 864691135479412678
post
9281
pre
3866
Target Soma Depth
Synapse Depth
L1
L2/3
L4
L5
L6
WM
Target distance to root
Target Soma Depth
### Syn
Dist. to soma (μm)
Selectivity Index
Soma Prox Dist Apic Inhib
Synapse Count
L2a
L2b
L2c
L3a
L3b
L4a
L4b
L4c
L5a
L5b
L5ET
L5NP
L6short-a
L6short-b
L6tall-a
L6tall-b
L6tall-c
L6wm
PTC
DTC
STC
ITC

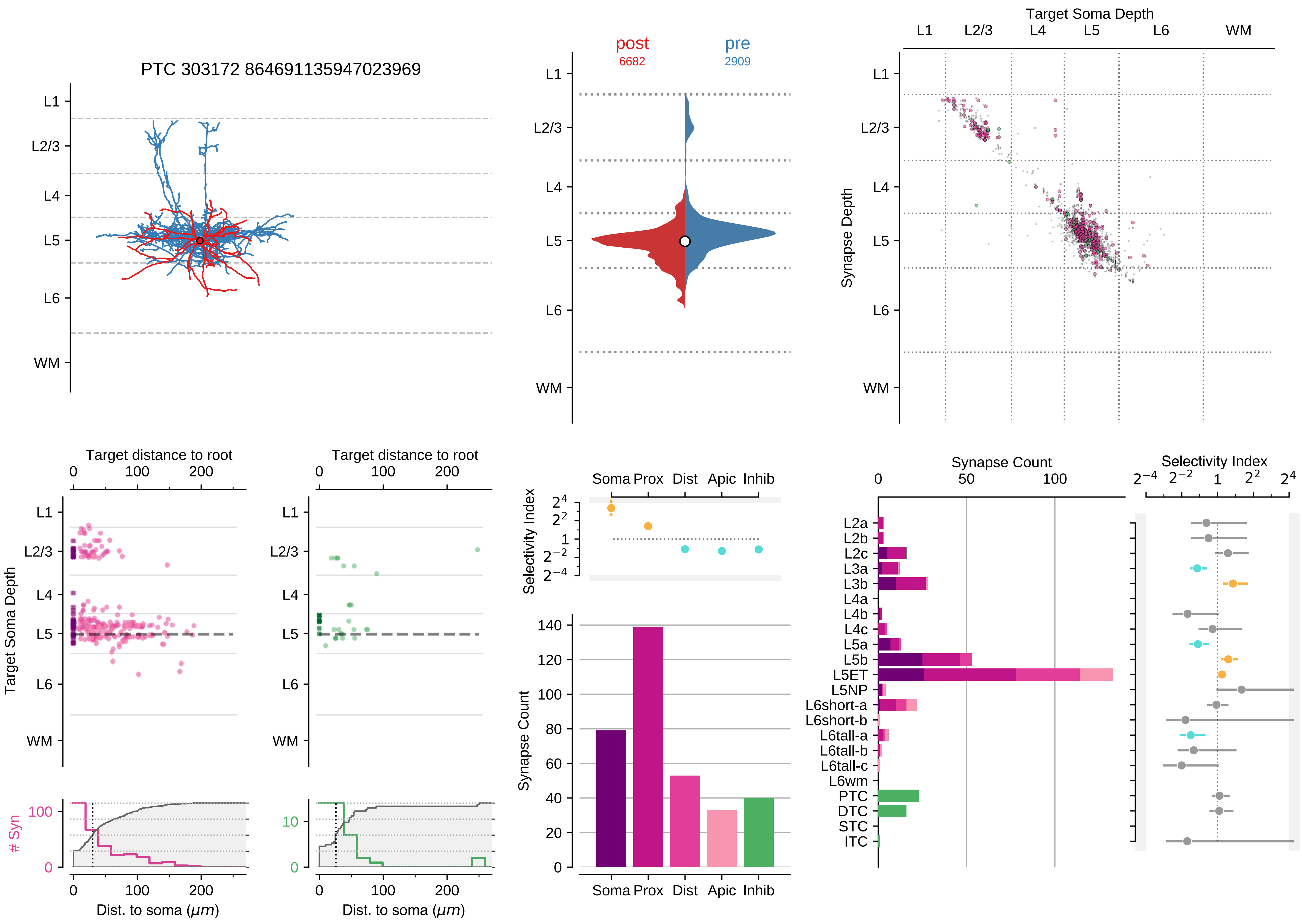

PTC 303172 864691135947023969
L1
L2/3
L4
L5
L6
WM
post
6682
pre
2909
Target Soma Depth
Synapse Depth
Target distance to root
Target Soma Depth
### Syn
Dist. to soma ($\mu m$)
Soma
Prox
Dist
Apic
Inhib
Selectivity Index
Synapse Count
L2a
L2b
L2c
L3a
L3b
L4a
L4b
L4c
L5a
L5b
L5ET
L5NP
L6short-a
L6short-b
L6tall-a
L6tall-b
L6tall-c
L6wm
PTC
DTC
STC
ITC

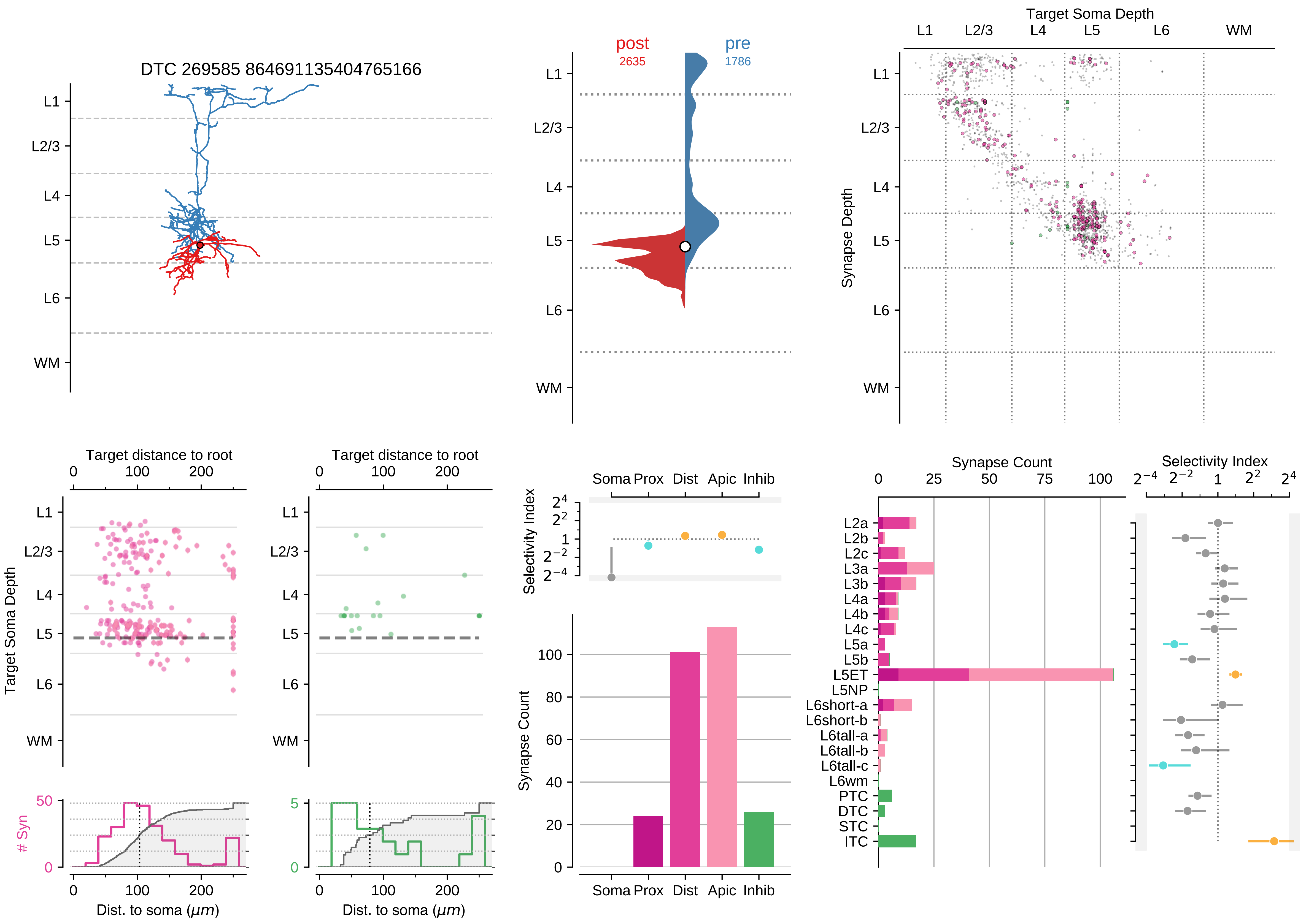
DTC 269585 864691135404765166
L1
L2/3
L4
L5
L6
WM
post
2635
pre
1786
Target Soma Depth
Synapse Depth
Target distance to root
Target Soma Depth
### Syn
Dist. to soma (μm)
Selectivity Index
Soma Prox Dist Apic Inhib
Synapse Count
L2a
L2b
L2c
L3a
L3b
L4a
L4b
L4c
L5a
L5b
L5ET
L5NP
L6short-a
L6short-b
L6tall-a
L6tall-b
L6tall-c
L6wm
PTC
DTC
STC
ITC

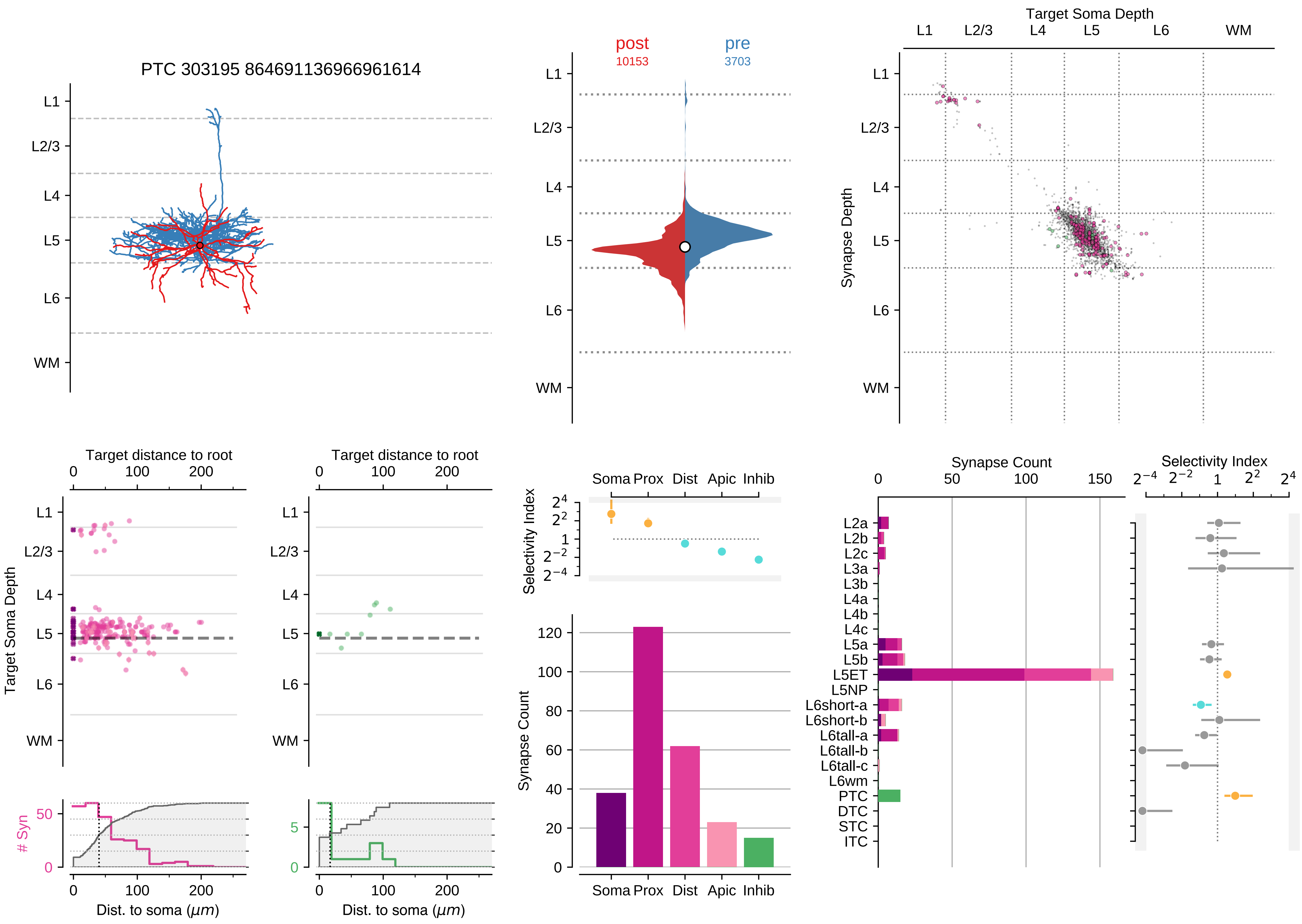

PTC 303195 864691136966961614
post
10153
pre
3703
Target Soma Depth
Synapse Depth
L1
L2/3
L4
L5
L6
WM
Target distance to root
Target Soma Depth
### Syn
Dist. to soma (μm)
Selectivity Index
Soma Prox Dist Apic Inhib
Synapse Count
L2a
L2b
L2c
L3a
L3b
L4a
L4b
L4c
L5a
L5b
L5ET
L5NP
L6short-a
L6short-b
L6tall-a
L6tall-b
L6tall-c
L6wm
PTC
DTC
STC
ITC

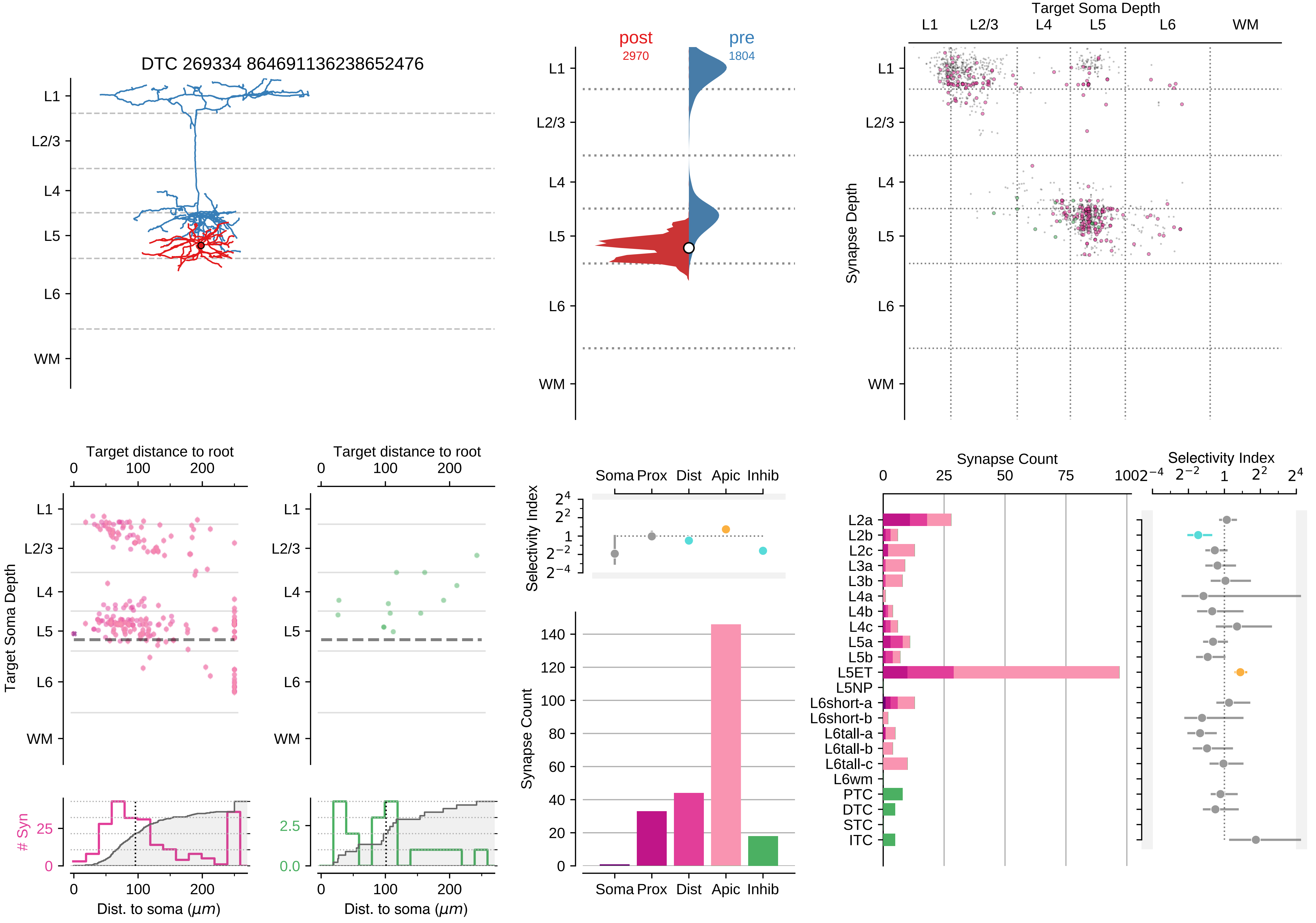

DTC 269334 864691136238652476
L1
L2/3
L4
L5
L6
WM
post
2970
pre
1804
Target Soma Depth
Synapse Depth
Target distance to root
Target Soma Depth
### Syn
Dist. to soma (μm)
Selectivity Index
Soma Prox Dist Apic Inhib
Synapse Count
L2a
L2b
L2c
L3a
L3b
L4a
L4b
L4c
L5a
L5b
L5ET
L5NP
L6short-a
L6short-b
L6tall-a
L6tall-b
L6tall-c
L6wm
PTC
DTC
STC
ITC

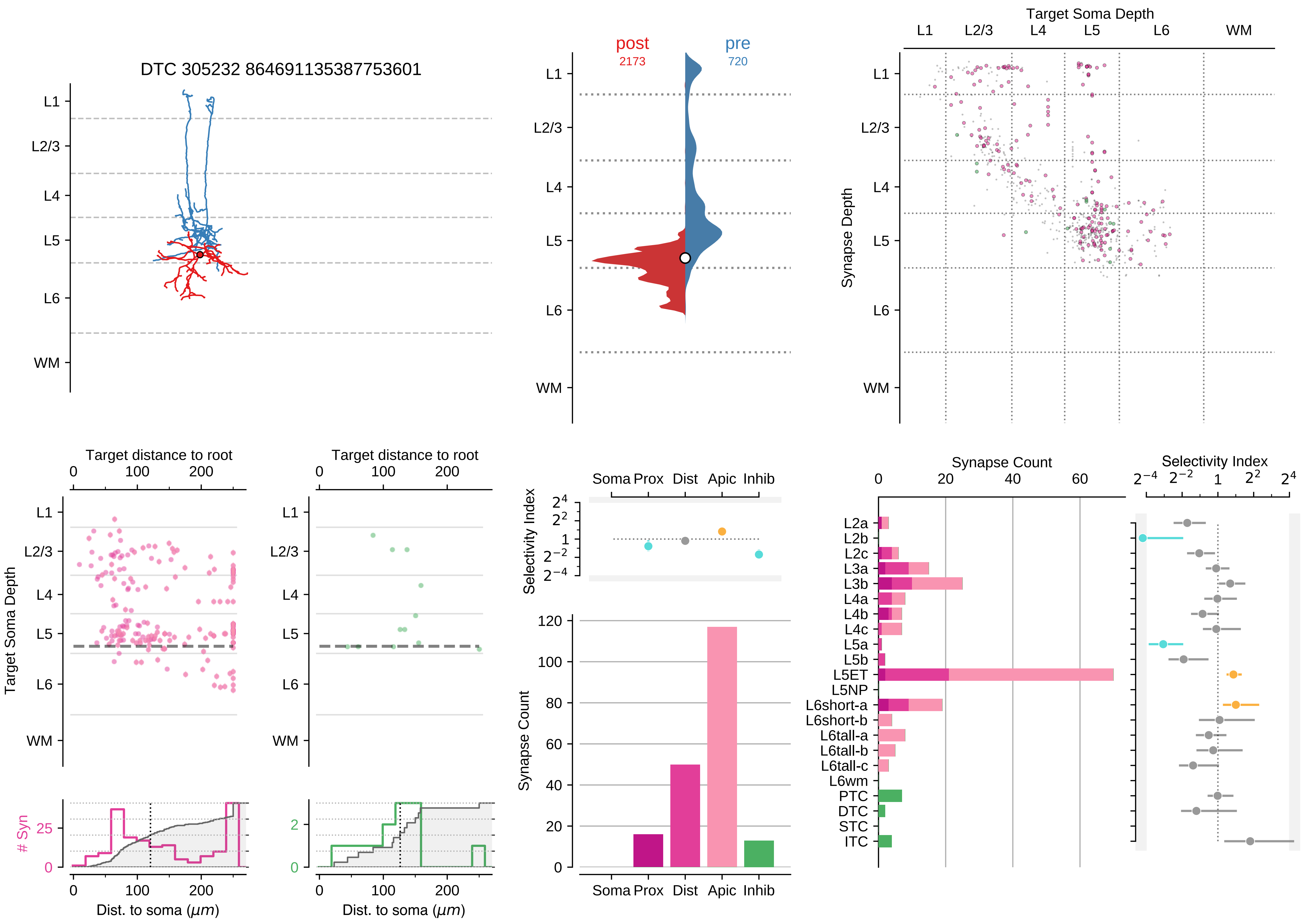

DTC 305232 864691135387753601
L1
L2/3
L4
L5
L6
WM
post
2173
pre
720
Target Soma Depth
Synapse Depth
Target distance to root
Target Soma Depth
### Syn
Dist. to soma (μm)
Selectivity Index
Soma Prox Dist Apic Inhib
Synapse Count
L2a
L2b
L2c
L3a
L3b
L4a
L4b
L4c
L5a
L5b
L5ET
L5NP
L6short-a
L6short-b
L6tall-a
L6tall-b
L6tall-c
L6wm
PTC
DTC
STC
ITC

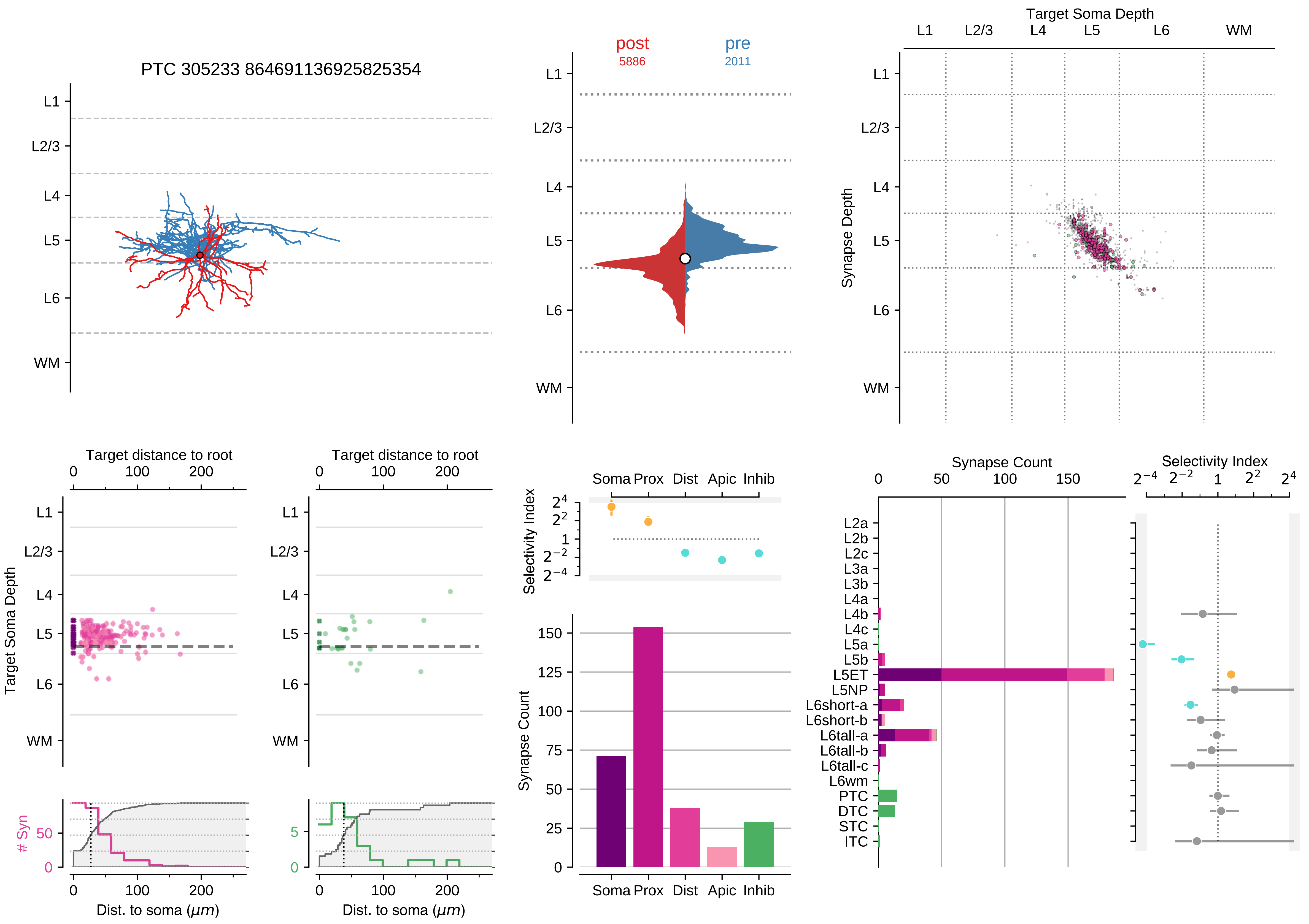
PTC 305233 864691136925825354
L1
L2/3
L4
L5
L6
WM
post
5886
pre
2011
Target Soma Depth
Synapse Depth
Target distance to root
Target Soma Depth
### Syn
Dist. to soma ($\mu m$)
Selectivity Index
Soma Prox Dist Apic Inhib
Synapse Count
Synapse Count
L2a
L2b
L2c
L3a
L3b
L4a
L4b
L4c
L5a
L5b
L5ET
L5NP
L6short-a
L6short-b
L6tall-a
L6tall-b
L6tall-c
L6wm
PTC
DTC
STC
ITC

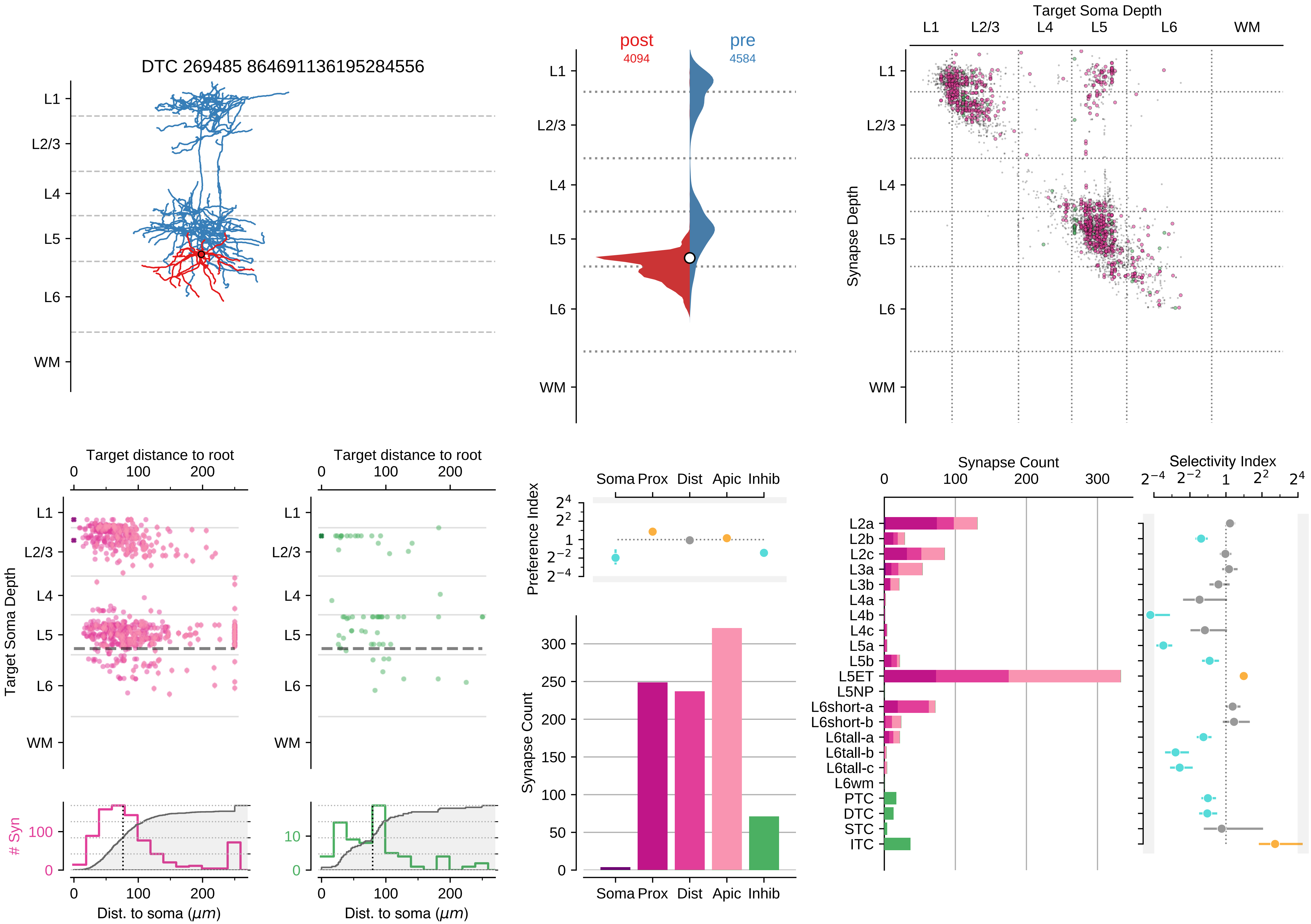

DTC 269485 864691136195284556
L1
L2/3
L4
L5
L6
WM
post
4094
pre
4584
Target Soma Depth
Synapse Depth
Target distance to root
Target Soma Depth
### Syn
Dist. to soma ($\mu m$)
Preference Index
Soma Prox Dist Apic Inhib
Synapse Count
Synapse Count
Selectivity Index
L2a
L2b
L2c
L3a
L3b
L4a
L4b
L4c
L5a
L5b
L5ET
L5NP
L6short-a
L6short-b
L6tall-a
L6tall-b
L6tall-c
L6wm
PTC
DTC
STC
ITC

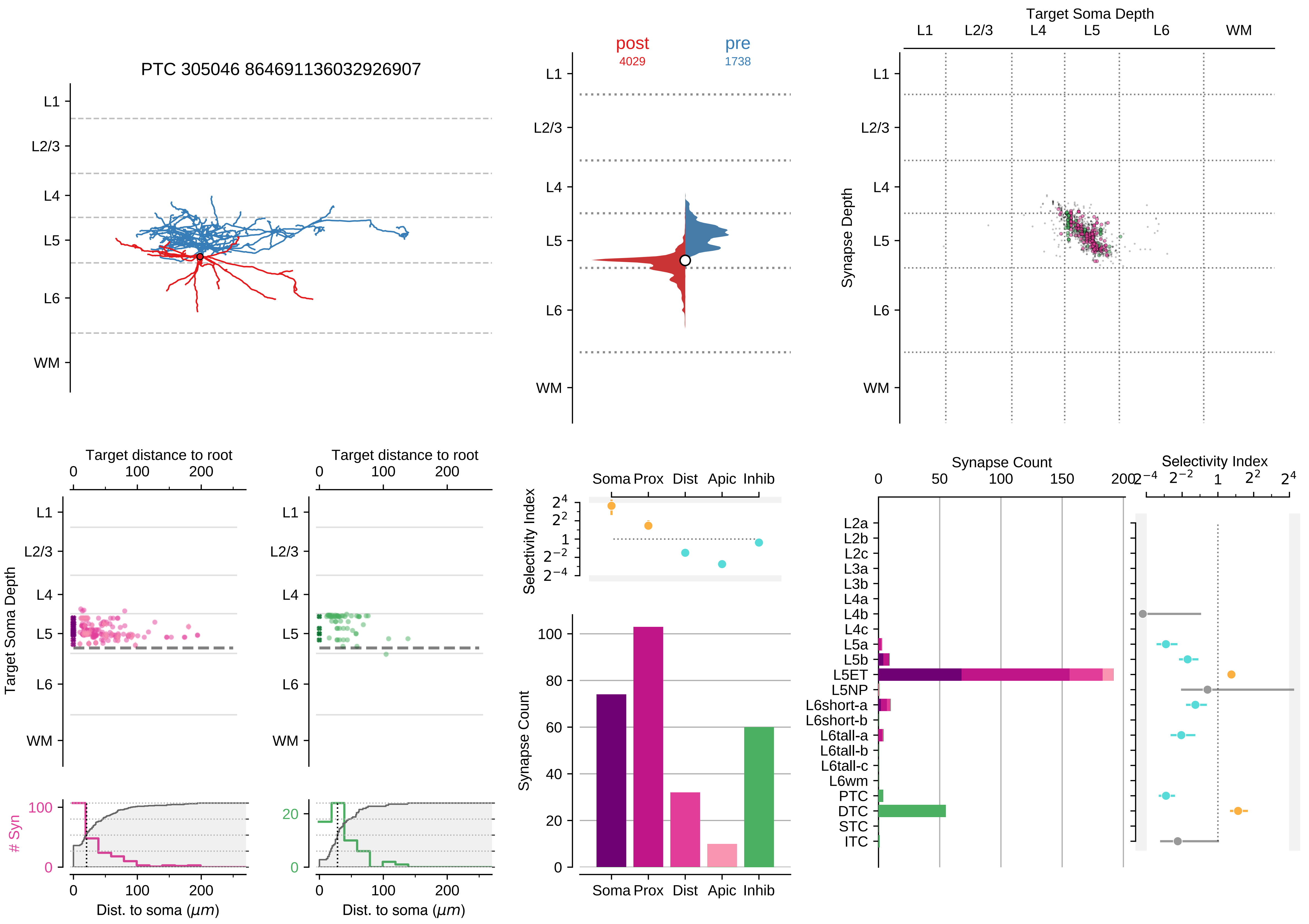

PTC 305046 864691136032926907
L1
L2/3
L4
L5
L6
WM
post
4029
pre
1738
Target Soma Depth
Synapse Depth
Target distance to root
Target Soma Depth
### Syn
Dist. to soma ($\mu m$)
Selectivity Index
Soma Prox Dist Apic Inhib
Synapse Count
Synapse Count
Selectivity Index
L2a
L2b
L2c
L3a
L3b
L4a
L4b
L4c
L5a
L5b
L5ET
L5NP
L6short-a
L6short-b
L6tall-a
L6tall-b
L6tall-c
L6wm
PTC
DTC
STC
ITC

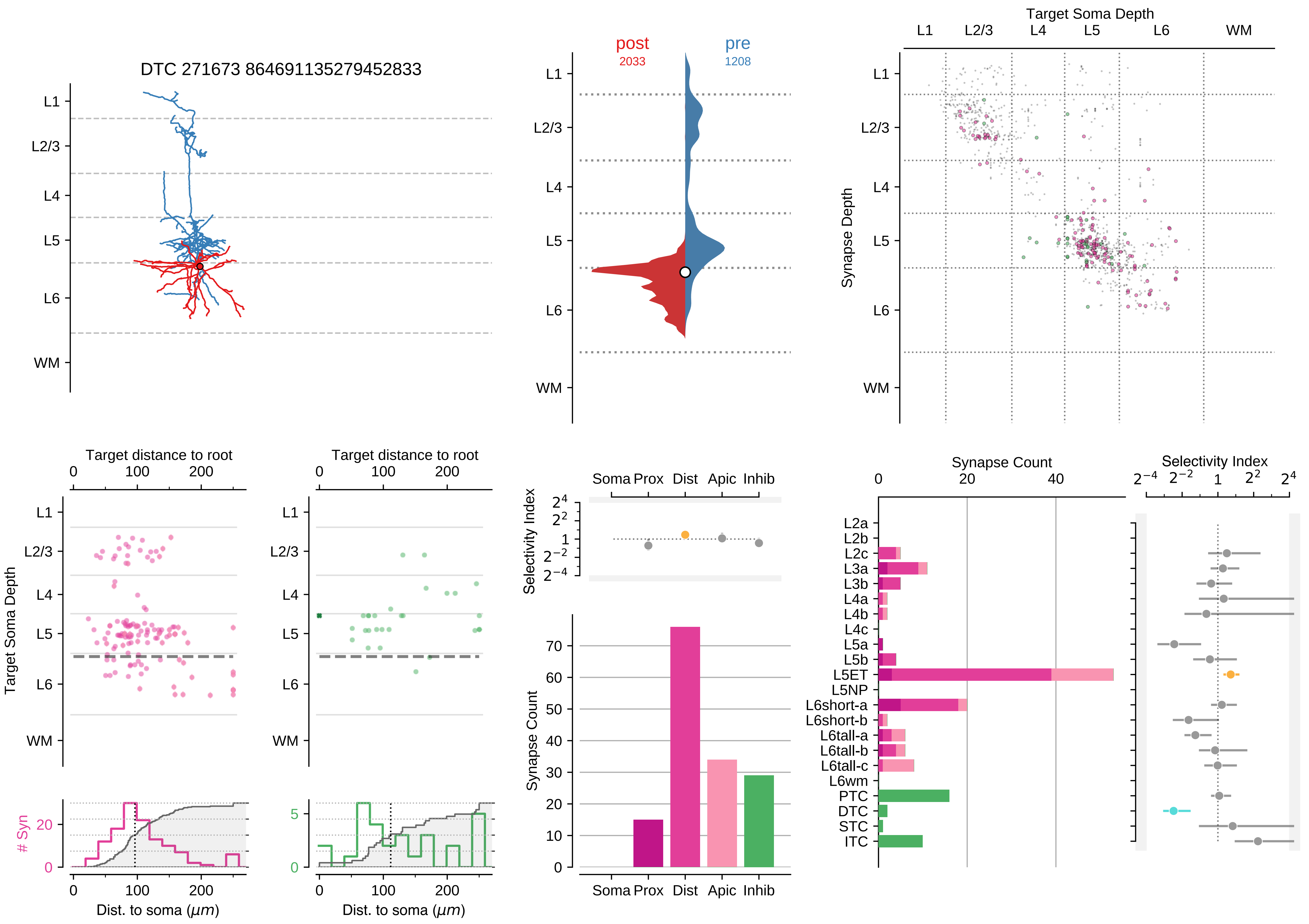
DTC 271673 864691135279452833
L1
L2/3
L4
L5
L6
WM
post
2033
pre
1208
Target Soma Depth
Synapse Depth
Target distance to root
Dist. to soma ($\mu m$)
### Syn
Selectivity Index
Synapse Count
Soma Prox
Dist
Apic
Inhib
L2a
L2b
L2c
L3a
L3b
L4a
L4b
L4c
L5a
L5b
L5ET
L5NP
L6short-a
L6short-b
L6tall-a
L6tall-b
L6tall-c
L6wm
PTC
DTC
STC
ITC

### Motif Group 10

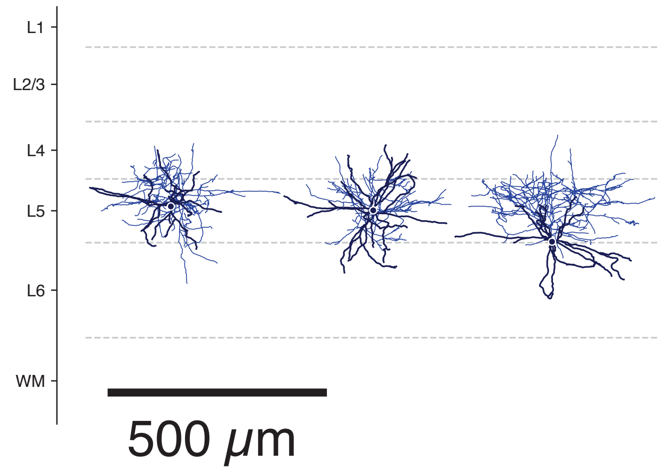

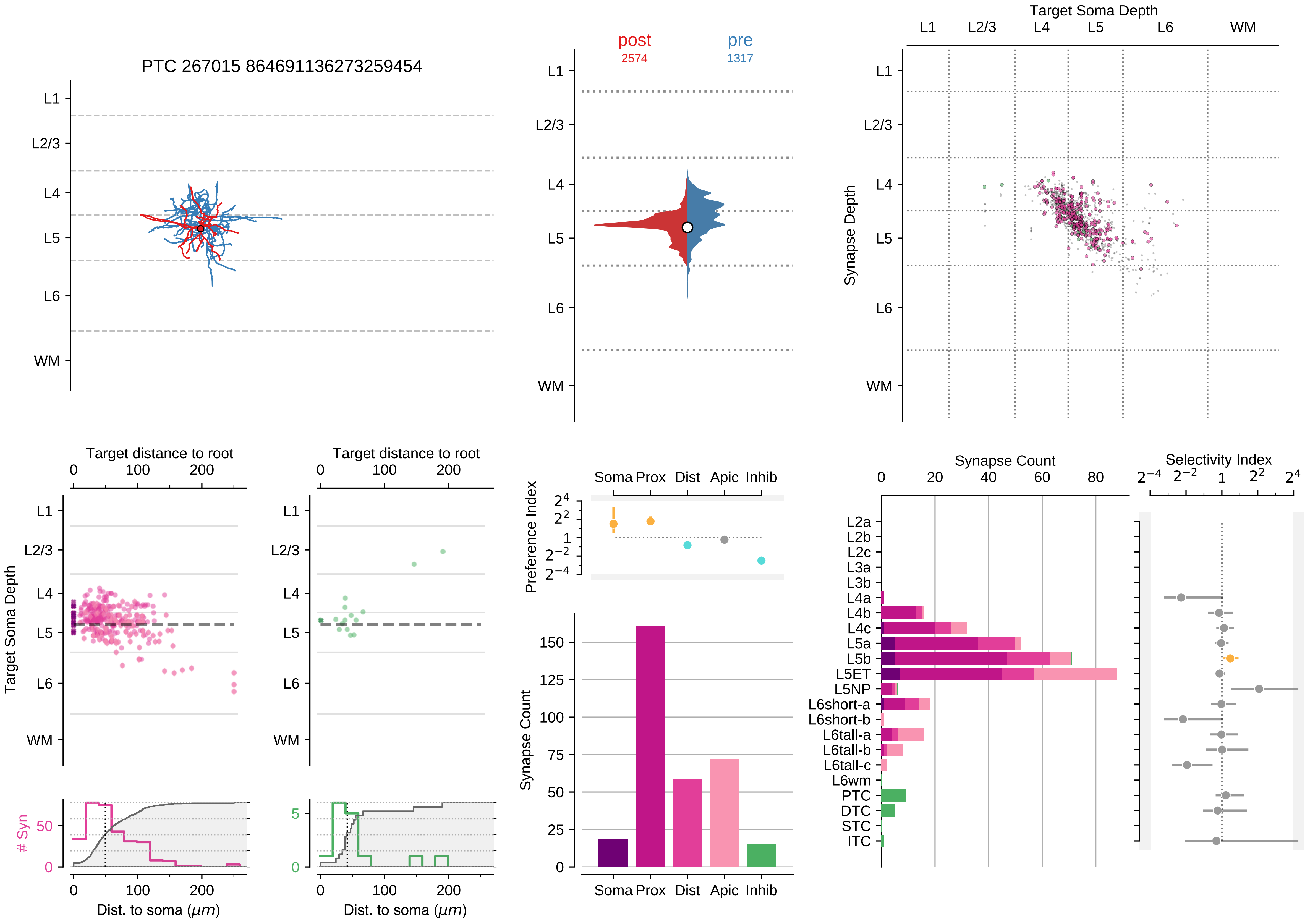
PTC 267015 864691136273259454
L1
L2/3
L4
L5
L6
WM
post
2574
pre
1317
Target Soma Depth
Synapse Depth
Target distance to root
0
100
200
Target Soma Depth
### Syn
50
0
5
Dist. to soma ($\mu m$)
Soma
Prox
Dist
Apic
Inhib
Preference Index
$2^4$
$2^2$
1
$2^{-2}$
$2^{-4}$
Synapse Count
150
125
100
75
50
25
0
20
40
60
80
Selectivity Index
L2a
L2b
L2c
L3a
L3b
L4a
L4b
L4c
L5a
L5b
L5ET
L5NP
L6short-a
L6short-b
L6tall-a
L6tall-b
L6tall-c
L6wm
PTC
DTC
STC
ITC

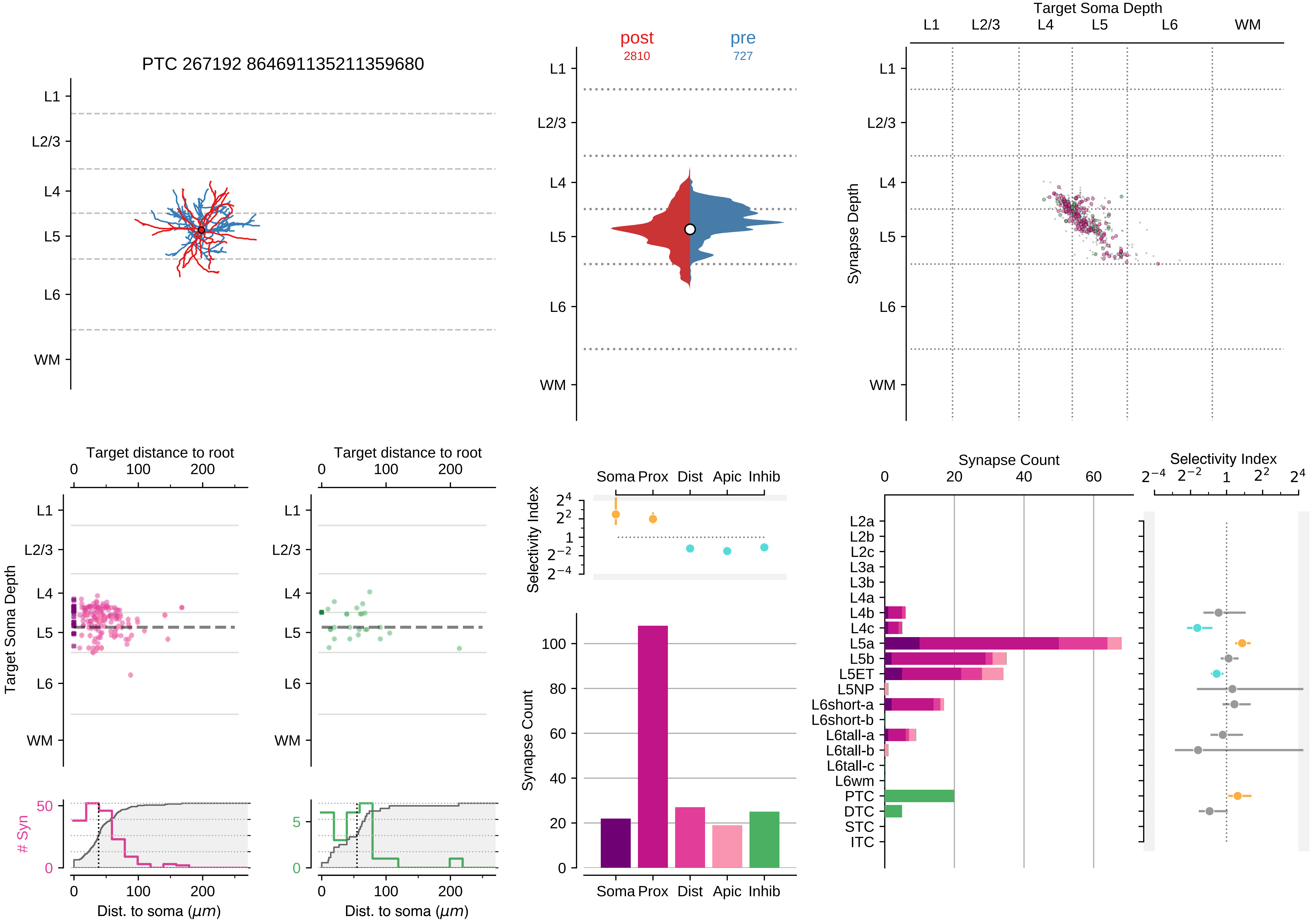

PTC 267192 864691135211359680
L1
L2/3
L4
L5
L6
WM
post
2810
pre
727
Target Soma Depth
Synapse Depth
Target distance to root
Target Soma Depth
### Syn
Dist. to soma (μm)
Selectivity Index
Soma
Prox
Dist
Apic
Inhib
Synapse Count
L2a
L2b
L2c
L3a
L3b
L4a
L4b
L4c
L5a
L5b
L5ET
L5NP
L6short-a
L6short-b
L6tall-a
L6tall-b
L6tall-c
L6wm
PTC
DTC
STC
ITC

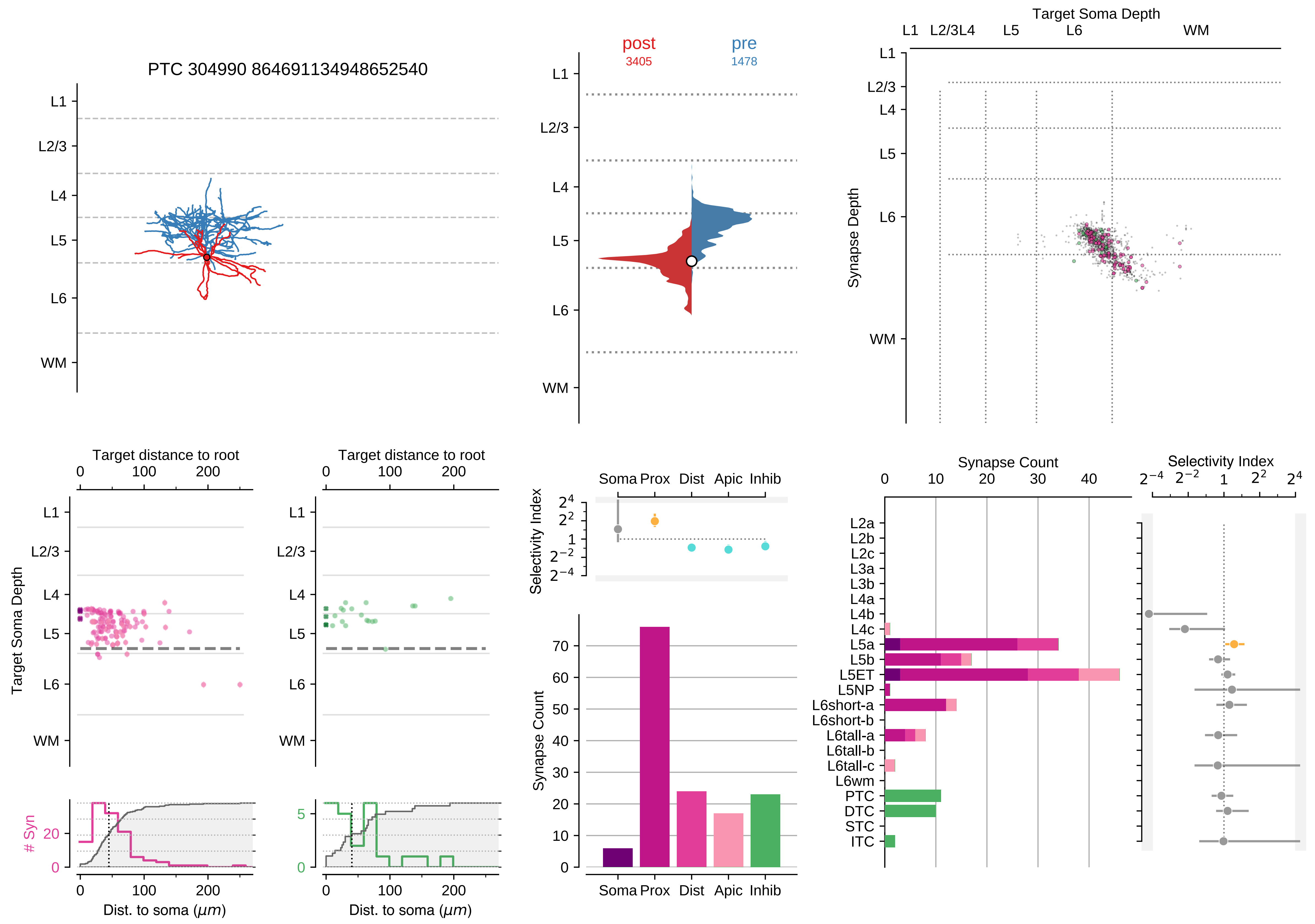

PTC 304990 864691134948652540
L1
L2/3
L4
L5
L6
WM
post
3405
pre
1478
Target Soma Depth
Synapse Depth
Target distance to root
Target Soma Depth
### Syn
Dist. to soma (μm)
Selectivity Index
Soma
Prox
Dist
Apic
Inhib
Synapse Count
L2a
L2b
L2c
L3a
L3b
L4a
L4b
L4c
L5a
L5b
L5ET
L5NP
L6short-a
L6short-b
L6tall-a
L6tall-b
L6tall-c
L6wm
PTC
DTC
STC
ITC

### Motif Group 11

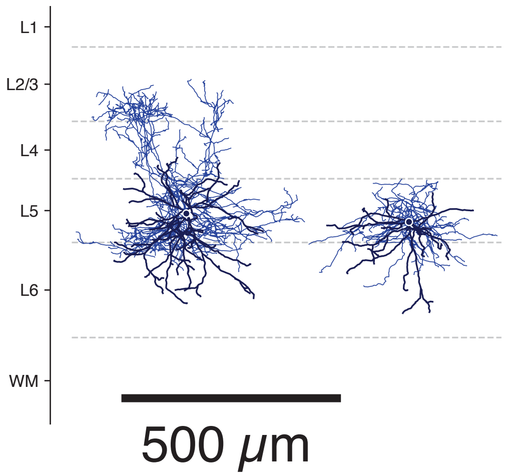

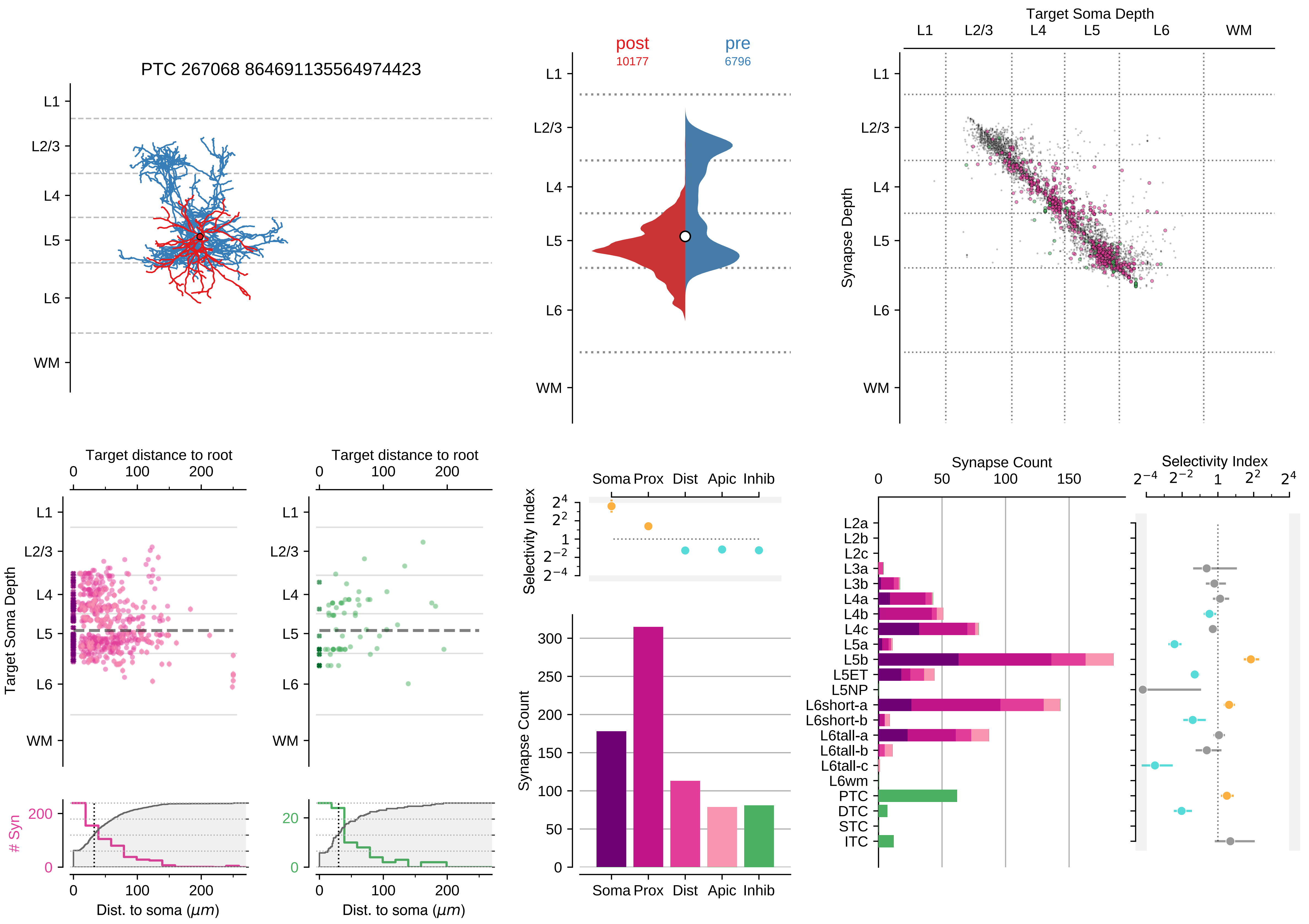

PTC 267068 864691135564974423
L1
L2/3
L4
L5
L6
WM
post
10177
pre
6796
Target Soma Depth
Synapse Depth
Target distance to root
Target Soma Depth
### Syn
Dist. to soma ($\mu m$)
Selectivity Index
Soma Prox Dist Apic Inhib
Synapse Count
L2a
L2b
L2c
L3a
L3b
L4a
L4b
L4c
L5a
L5b
L5ET
L5NP
L6short-a
L6short-b
L6tall-a
L6tall-b
L6tall-c
L6wm
PTC
DTC
STC
ITC

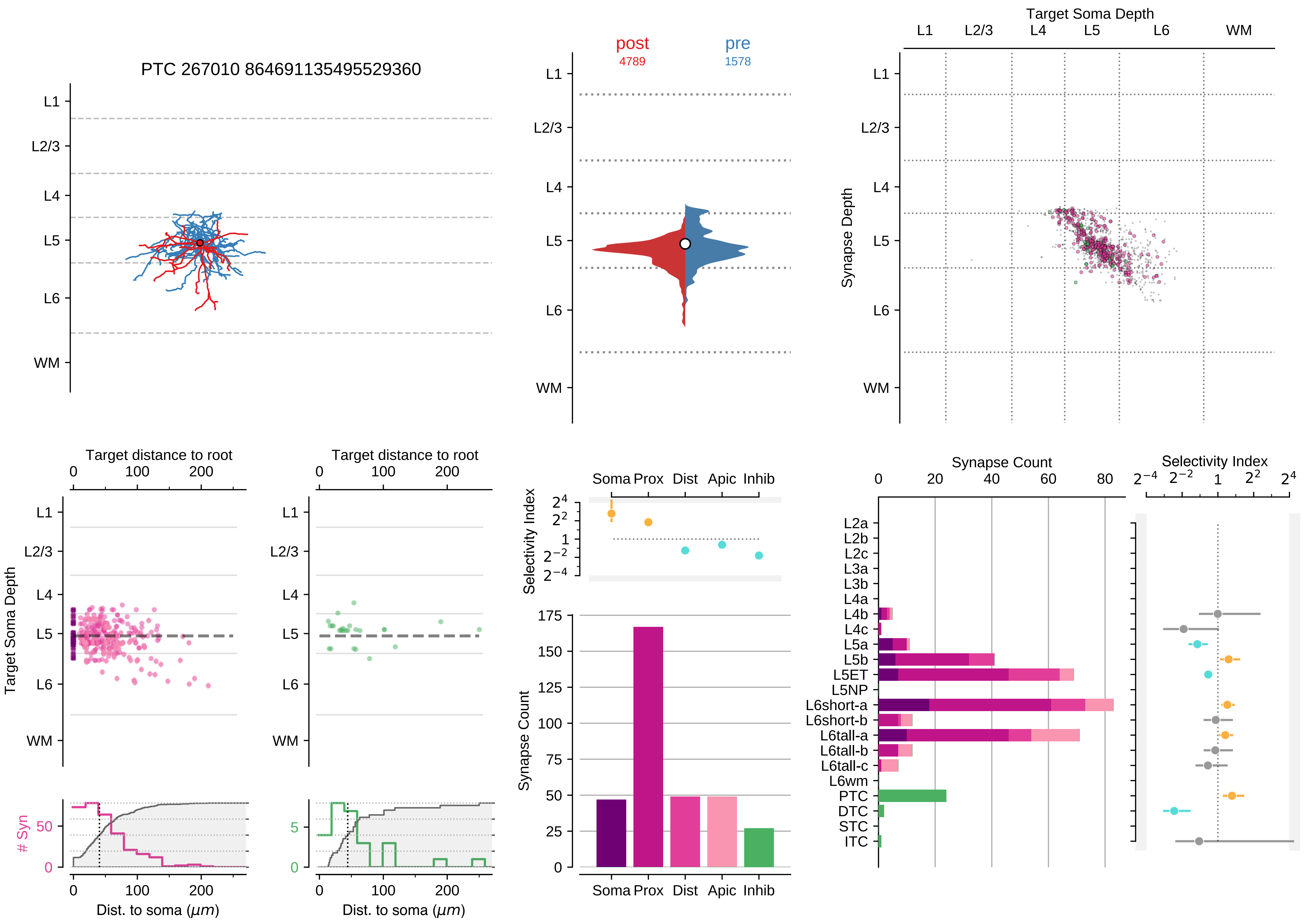

PTC 267010 864691135495529360
post
4789
pre
1578
Target Soma Depth
L1
L2/3
L4
L5
L6
WM
Synapse Depth
Target distance to root
Target Soma Depth
### Syn
Dist. to soma ($\mu m$)
Selectivity Index
Soma
Prox
Dist
Apic
Inhib
Synapse Count
L2a
L2b
L2c
L3a
L3b
L4a
L4b
L4c
L5a
L5b
L5ET
L5NP
L6short-a
L6short-b
L6tall-a
L6tall-b
L6tall-c
L6wm
PTC
DTC
STC
ITC

### Motif Group 12

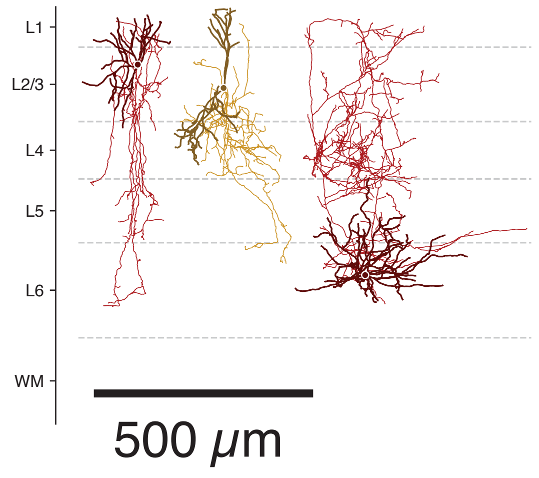

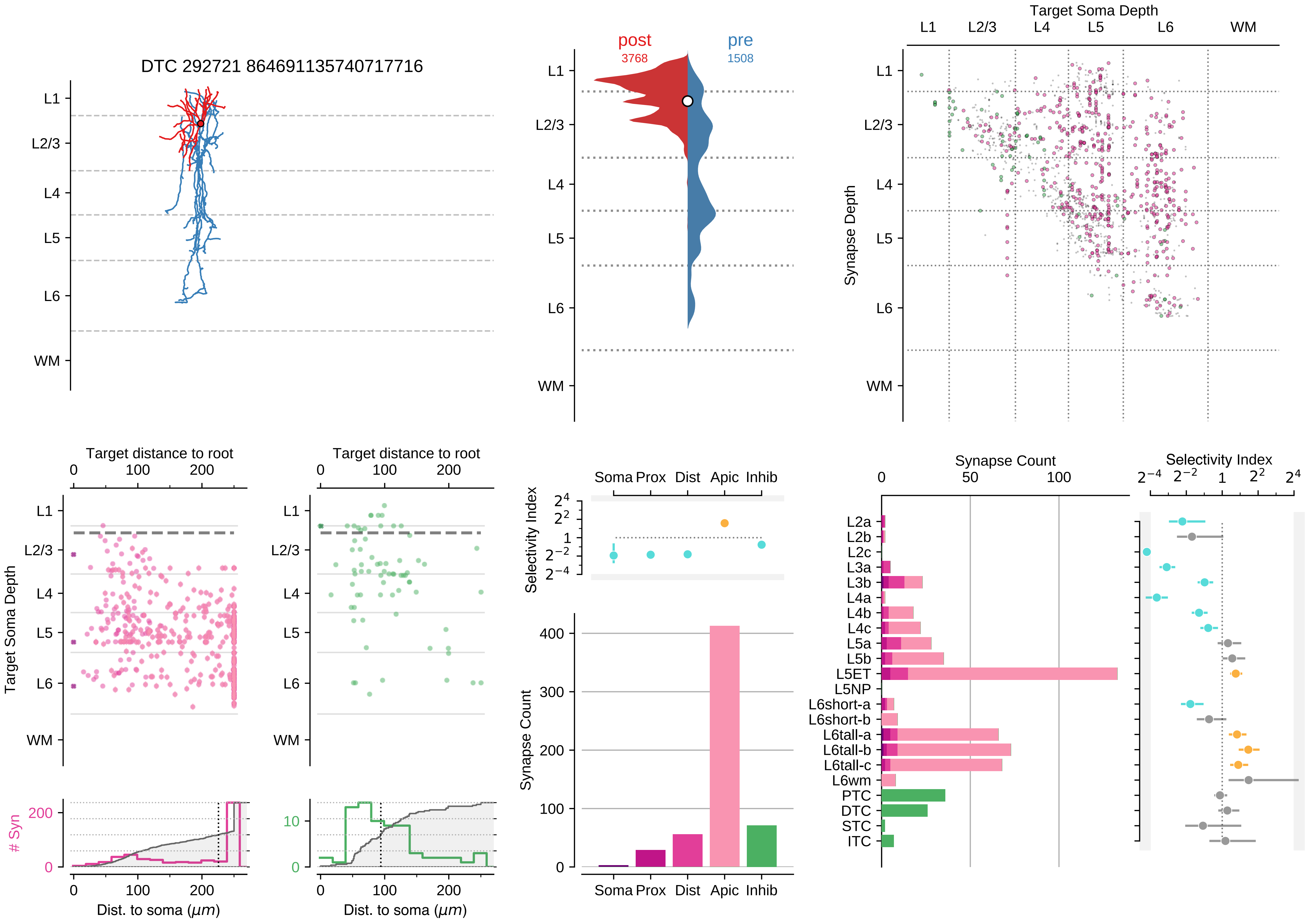

DTC 292721 864691135740717716
L1
L2/3
L4
L5
L6
WM
post
3768
pre
1508
Target Soma Depth
Synapse Depth
Target distance to root
Target Soma Depth
### Syn
Dist. to soma ($\mu m$)
Soma Prox Dist Apic Inhib
Selectivity Index
Synapse Count
L2a
L2b
L2c
L3a
L3b
L4a
L4b
L4c
L5a
L5b
L5ET
L5NP
L6short-a
L6short-b
L6tall-a
L6tall-b
L6tall-c
L6wm
PTC
DTC
STC
ITC

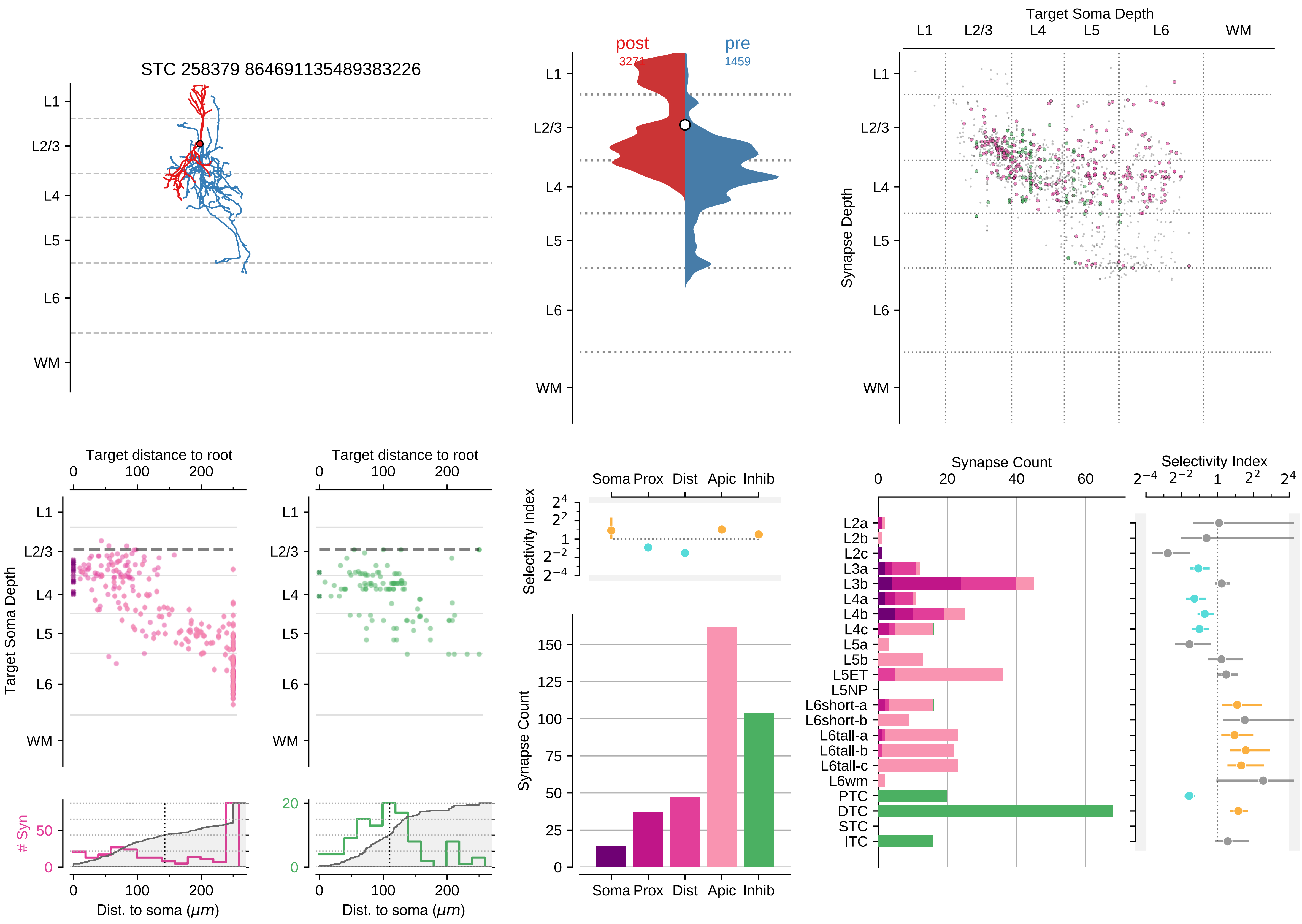

STC 258379 864691135489383226
post
pre
1459
Target Soma Depth
Synapse Depth
Target distance to root
Target Soma Depth
Dist. to soma (μm)
### Syn
Selectivity Index
Soma Prox Dist Apic Inhib
Synapse Count
L1
L2/3
L4
L5
L6
WM
L2a
L2b
L2c
L3a
L3b
L4a
L4b
L4c
L5a
L5b
L5ET
L5NP
L6short-a
L6short-b
L6tall-a
L6tall-b
L6tall-c
L6wm
PTC
DTC
STC
ITC

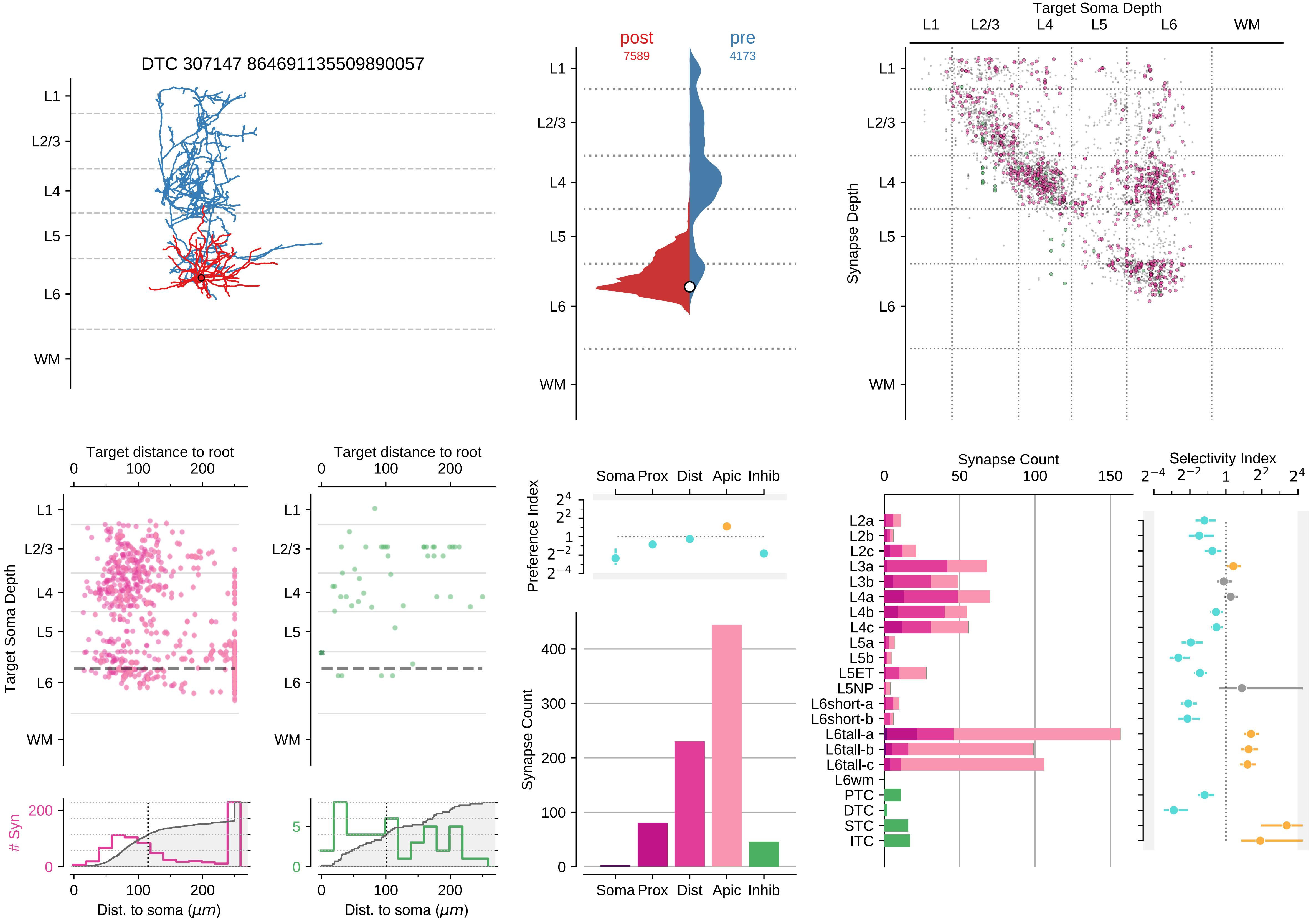

DTC 307147 864691135509890057
L1
L2/3
L4
L5
L6
WM
post
7589
pre
4173
Target Soma Depth
Synapse Depth
Target distance to root
Target Soma Depth
### Syn
Dist. to soma (μm)
Preference Index
Soma Prox Dist Apic Inhib
Synapse Count
Selectivity Index
L2a
L2b
L2c
L3a
L3b
L4a
L4b
L4c
L5a
L5b
L5ET
L5NP
L6short-a
L6short-b
L6tall-a
L6tall-b
L6tall-c
L6wm
PTC
DTC
STC
ITC

### Motif Group 13

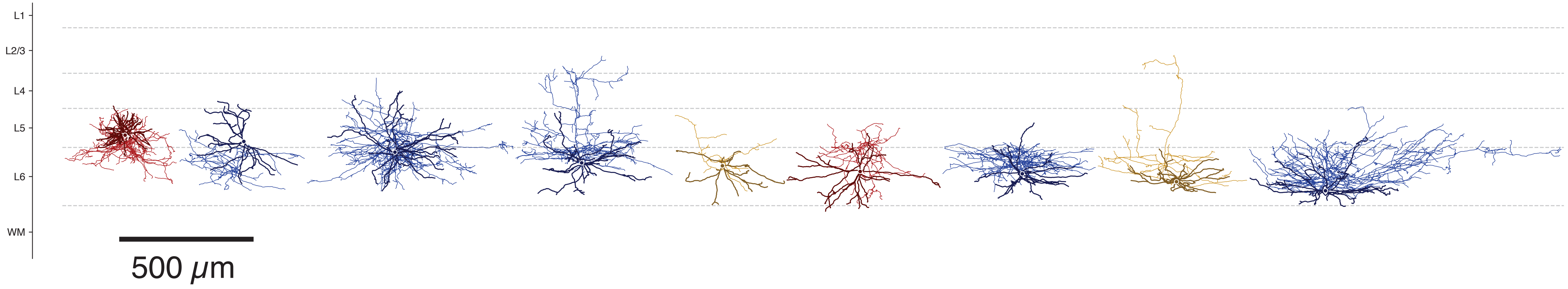

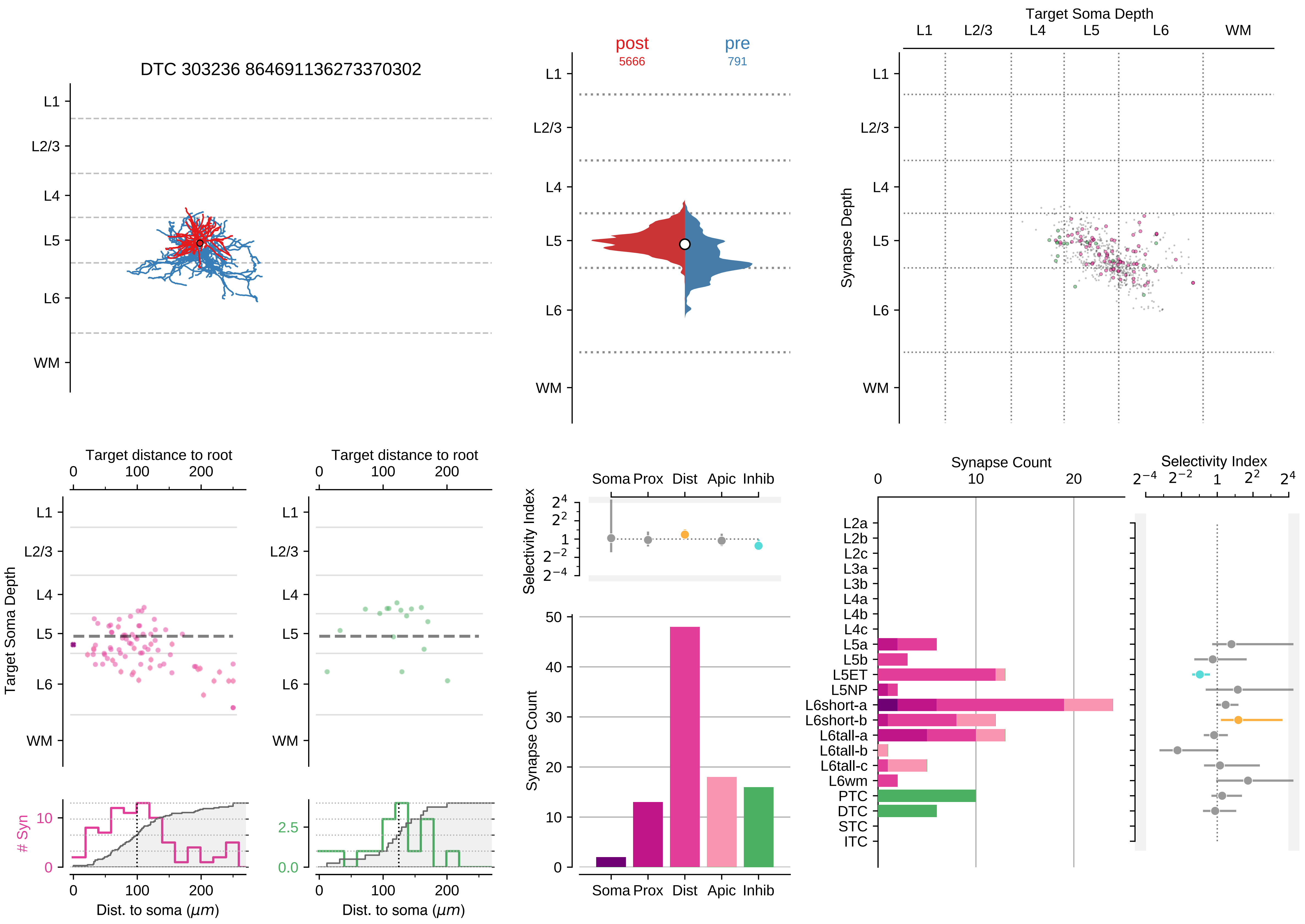

DTC 303236 864691136273370302
L1
L2/3
L4
L5
L6
WM
post
5666
pre
791
Target Soma Depth
Synapse Depth
Target distance to root
0
100
200
### Syn
Dist. to soma ($\mu m$)
Selectivity Index
Soma
Prox
Dist
Apic
Inhib
Synapse Count
$2^{-4}$
$2^{-2}$
1
$2^{2}$
$2^{4}$
L2a
L2b
L2c
L3a
L3b
L4a
L4b
L4c
L5a
L5b
L5ET
L5NP
L6short-a
L6short-b
L6tall-a
L6tall-b
L6tall-c
L6wm
PTC
DTC
STC
ITC

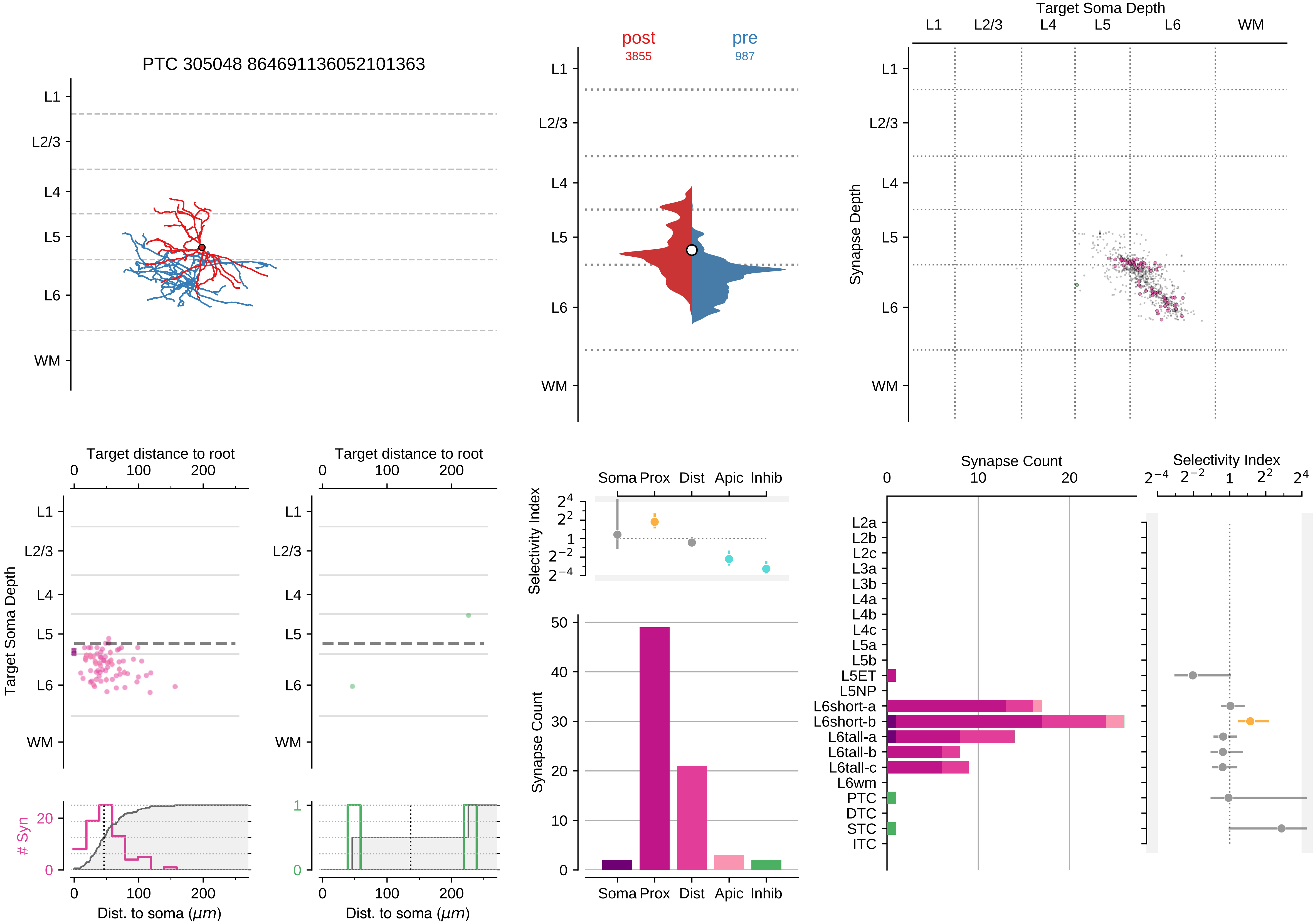

PTC 305048 864691136052101363
L1
L2/3
L4
L5
L6
WM
post
3855
pre
987
Target Soma Depth
Synapse Depth
Target distance to root
Target Soma Depth
### Syn
Dist. to soma ($\mu m$)
Selectivity Index
Soma
Prox
Dist
Apic
Inhib
Synapse Count
Selectivity Index
L2a
L2b
L2c
L3a
L3b
L4a
L4b
L4c
L5a
L5b
L5ET
L5NP
L6short-a
L6short-b
L6tall-a
L6tall-b
L6tall-c
L6wm
PTC
DTC
STC
ITC

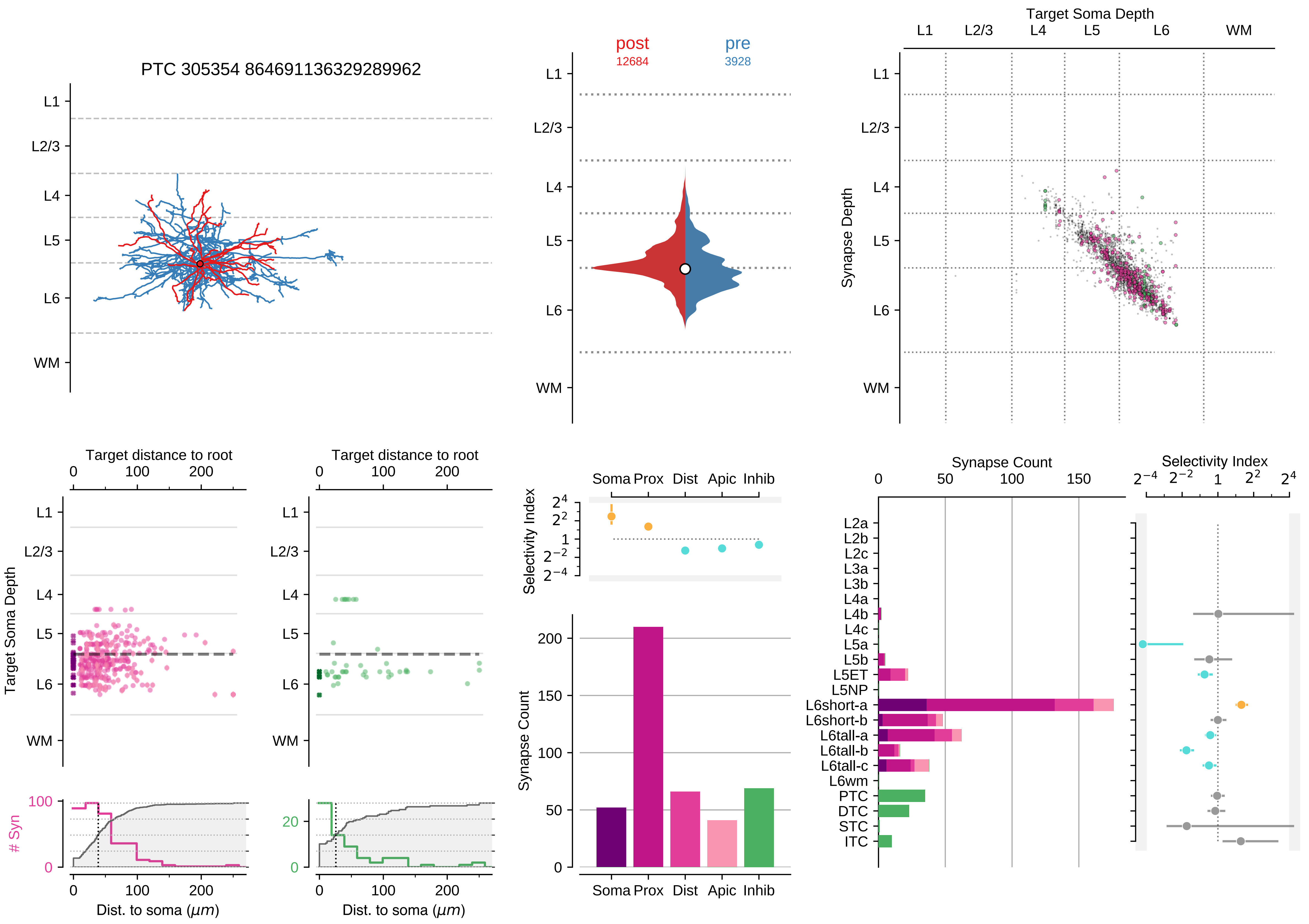

PTC 305354 864691136329289962
L1
L2/3
L4
L5
L6
WM
post
12684
pre
3928
Target Soma Depth
Synapse Depth
Target distance to root
0
100
200
Target Soma Depth
### Syn
Dist. to soma ($\mu m$)
Selectivity Index
Soma Prox Dist Apic Inhib
Synapse Count
Synapse Count
Selectivity Index
L2a
L2b
L2c
L3a
L3b
L4a
L4b
L4c
L5a
L5b
L5ET
L5NP
L6short-a
L6short-b
L6tall-a
L6tall-b
L6tall-c
L6wm
PTC
DTC
STC
ITC

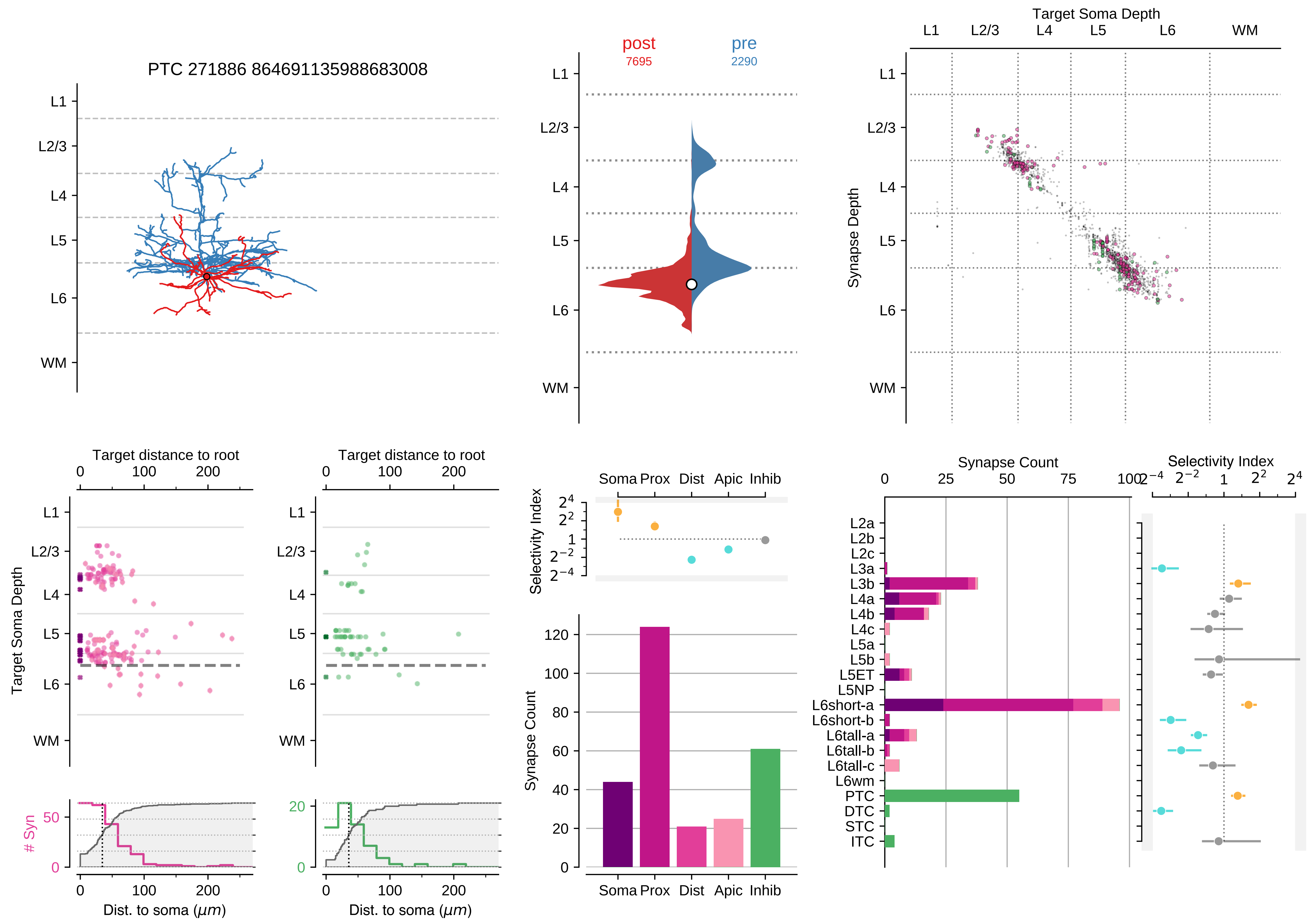

PTC 271886 864691135988683008
L1
L2/3
L4
L5
L6
WM
post
7695
pre
2290
Target Soma Depth
Synapse Depth
Target distance to root
Target Soma Depth
Dist. to soma ($\mu m$)
### Syn
Soma Prox Dist Apic Inhib
Selectivity Index
Synapse Count
L2a
L2b
L2c
L3a
L3b
L4a
L4b
L4c
L5a
L5b
L5ET
L5NP
L6short-a
L6short-b
L6tall-a
L6tall-b
L6tall-c
L6wm
PTC
DTC
STC
ITC

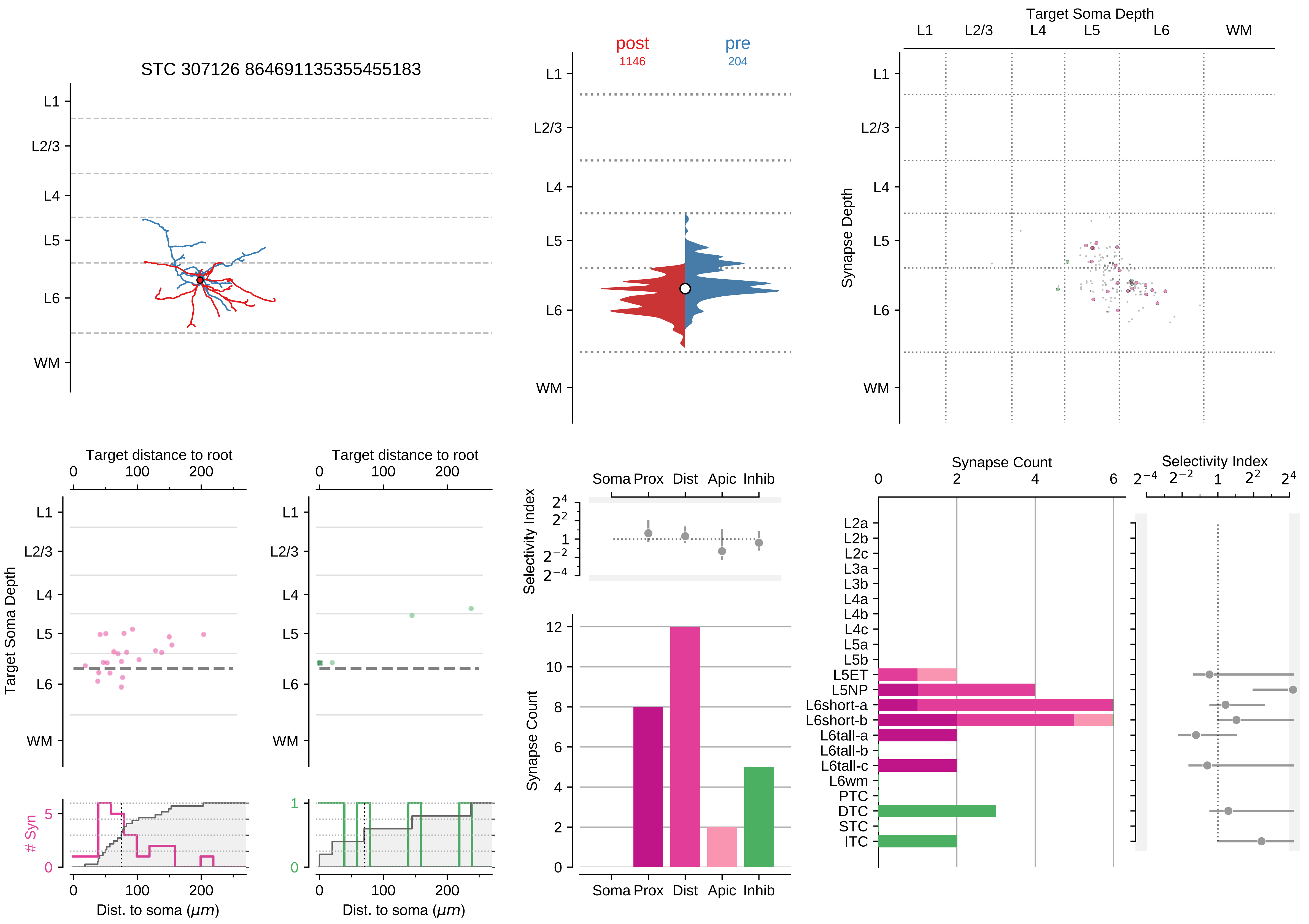

STC 307126 864691135355455183
L1
L2/3
L4
L5
L6
WM
post
1146
pre
204
Target Soma Depth
Synapse Depth
Target distance to root
Dist. to soma (μm)
### Syn
Selectivity Index
Soma Prox Dist Apic Inhib
Synapse Count
L2a
L2b
L2c
L3a
L3b
L4a
L4b
L4c
L5a
L5b
L5ET
L5NP
L6short-a
L6short-b
L6tall-a
L6tall-b
L6tall-c
L6wm
PTC
DTC
STC
ITC

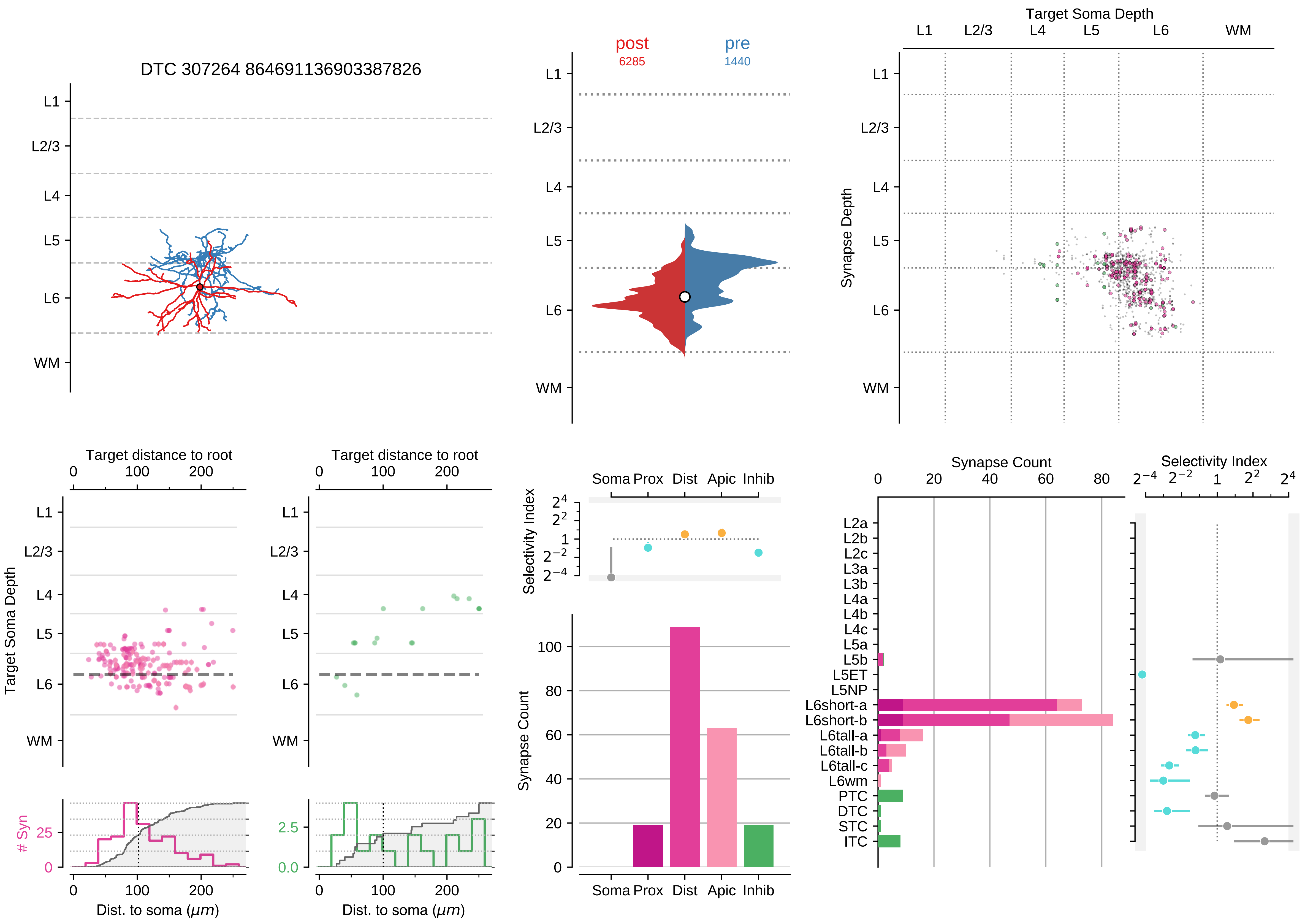
DTC 307264 864691136903387826
post
6285
pre
1440
Target Soma Depth
Synapse Depth
L1
L2/3
L4
L5
L6
WM
Target distance to root
Target Soma Depth
### Syn
Dist. to soma (μm)
Selectivity Index
Synapse Count
Soma
Prox
Dist
Apic
Inhib
L2a
L2b
L2c
L3a
L3b
L4a
L4b
L4c
L5a
L5b
L5ET
L5NP
L6short-a
L6short-b
L6tall-a
L6tall-b
L6tall-c
L6wm
PTC
DTC
STC
ITC

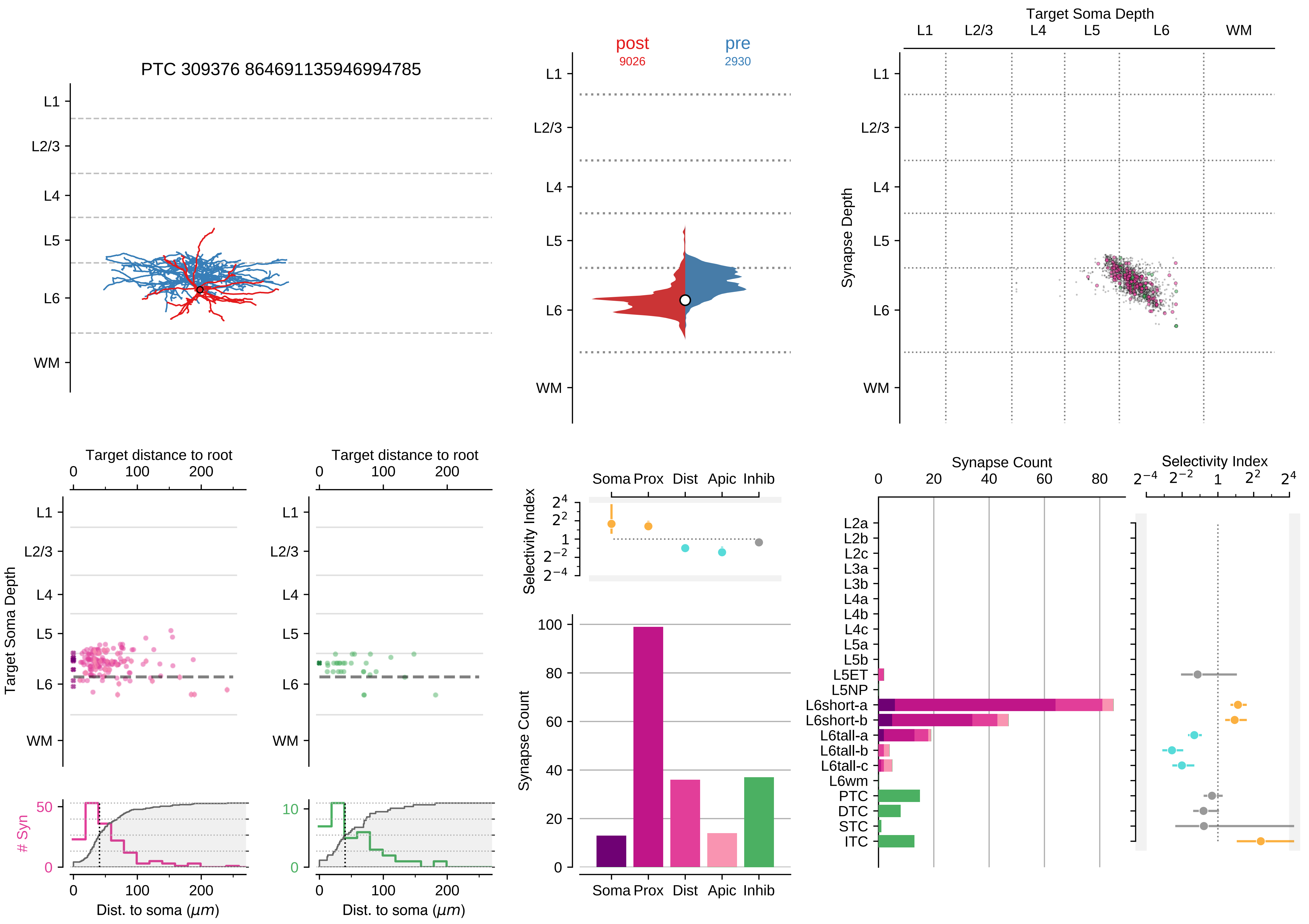

PTC 309376 864691135946994785
post
9026
pre
2930
Target Soma Depth
L1
L2/3
L4
L5
L6
WM
Synapse Depth
Target distance to root
Target Soma Depth
### Syn
Dist. to soma (μm)
Selectivity Index
Soma
Prox
Dist
Apic
Inhib
Synapse Count
L2a
L2b
L2c
L3a
L3b
L4a
L4b
L4c
L5a
L5b
L5ET
L5NP
L6short-a
L6short-b
L6tall-a
L6tall-b
L6tall-c
L6wm
PTC
DTC
STC
ITC

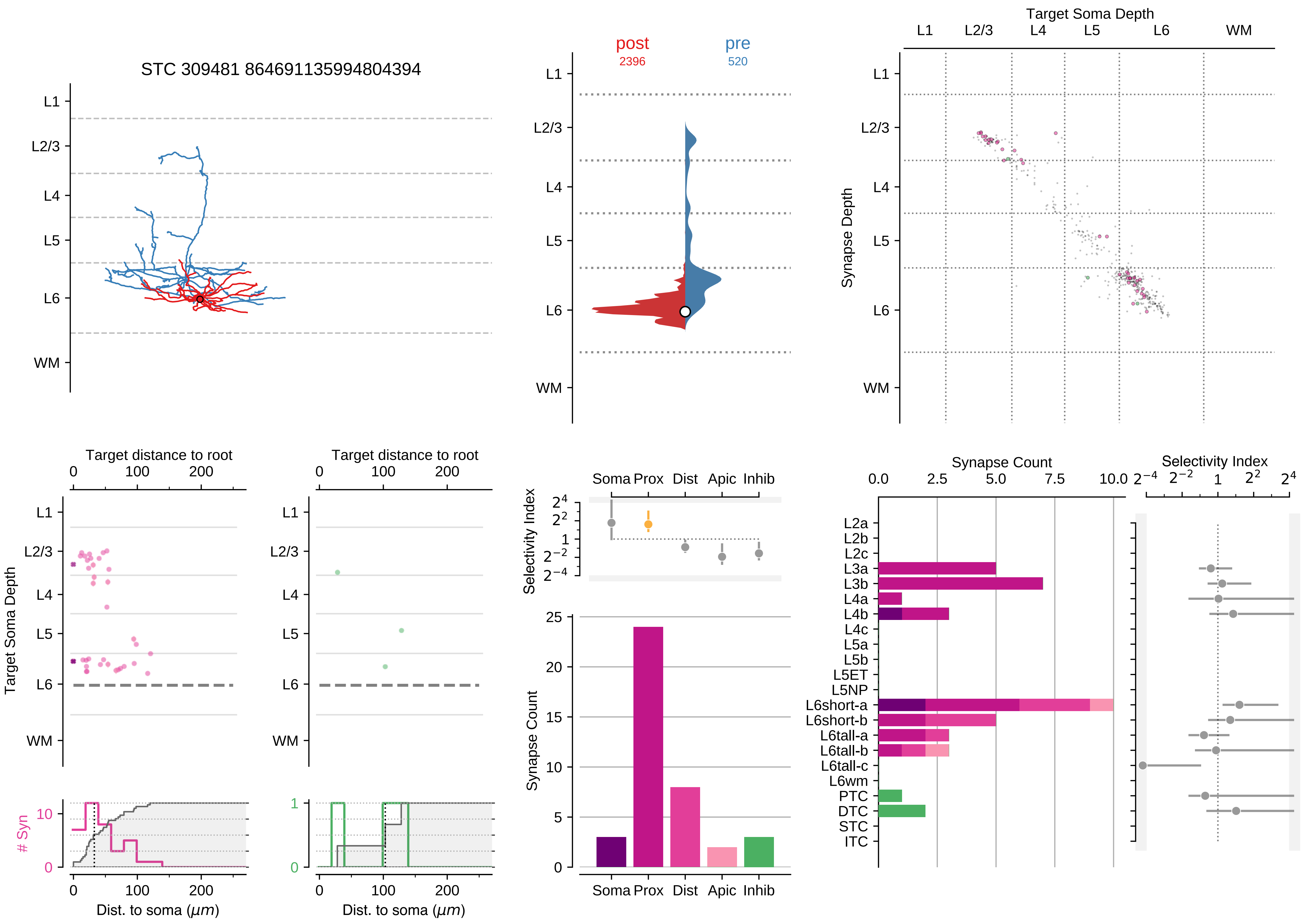

STC 309481 864691135994804394
L1
L2/3
L4
L5
L6
WM
post
2396
pre
520
Target Soma Depth
Synapse Depth
Target distance to root
Target Soma Depth
### Syn
Dist. to soma (μm)
Selectivity Index
Soma Prox Dist Apic Inhib
Synapse Count
L2a
L2b
L2c
L3a
L3b
L4a
L4b
L4c
L5a
L5b
L5ET
L5NP
L6short-a
L6short-b
L6tall-a
L6tall-b
L6tall-c
L6wm
PTC
DTC
STC
ITC

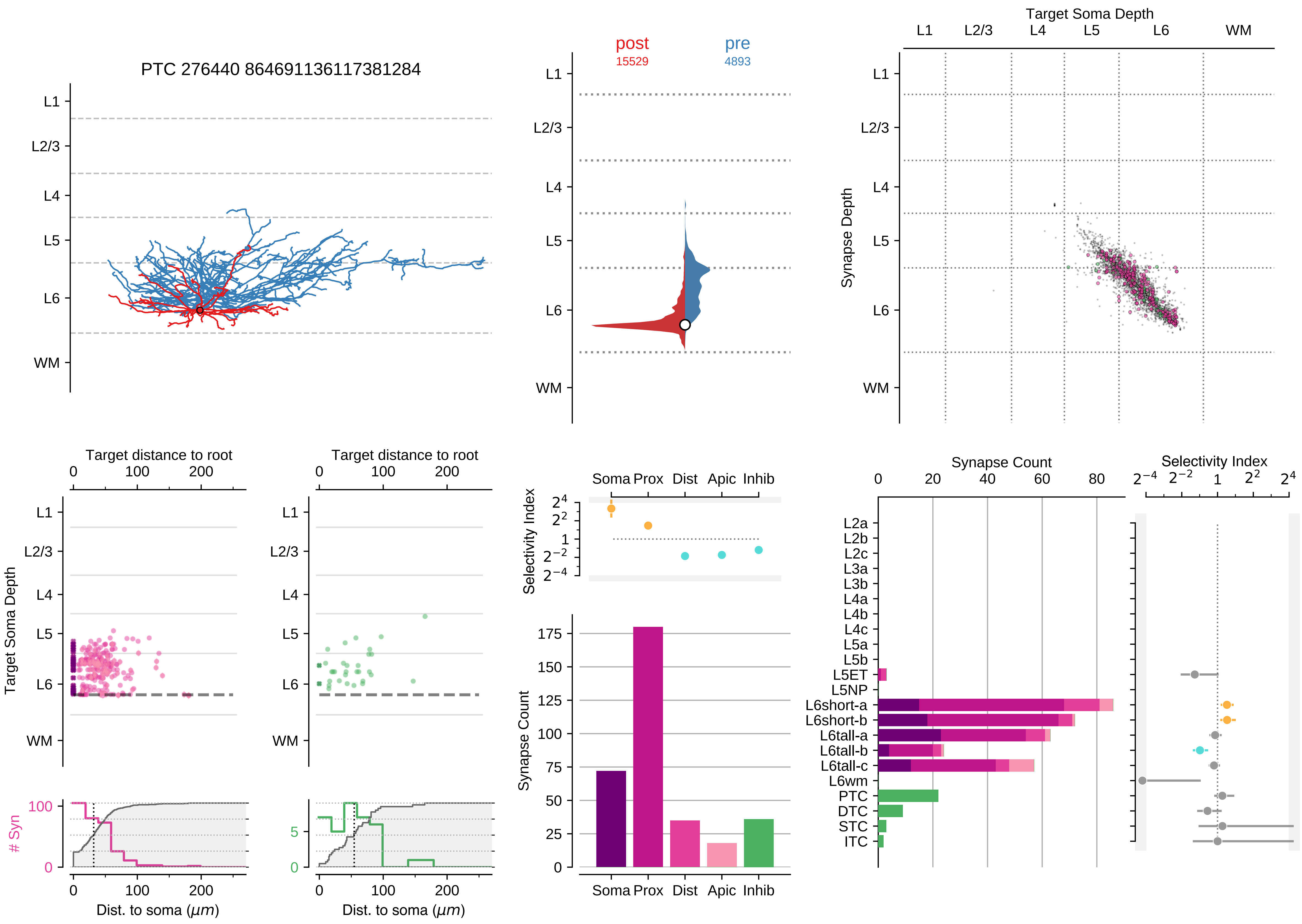

PTC 276440 864691136117381284
L1
L2/3
L4
L5
L6
WM
post
15529
pre
4893
Target Soma Depth
Synapse Depth
Target distance to root
Dist. to soma ($\mu m$)
### Syn
Selectivity Index
Soma Prox Dist Apic Inhib
Synapse Count
L2a
L2b
L2c
L3a
L3b
L4a
L4b
L4c
L5a
L5b
L5ET
L5NP
L6short-a
L6short-b
L6tall-a
L6tall-b
L6tall-c
L6wm
PTC
DTC
STC
ITC

### Motif Group 14

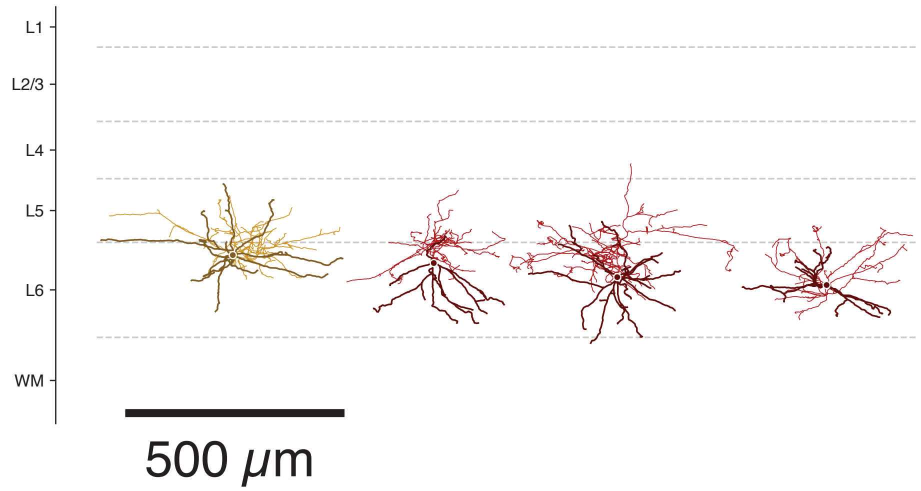

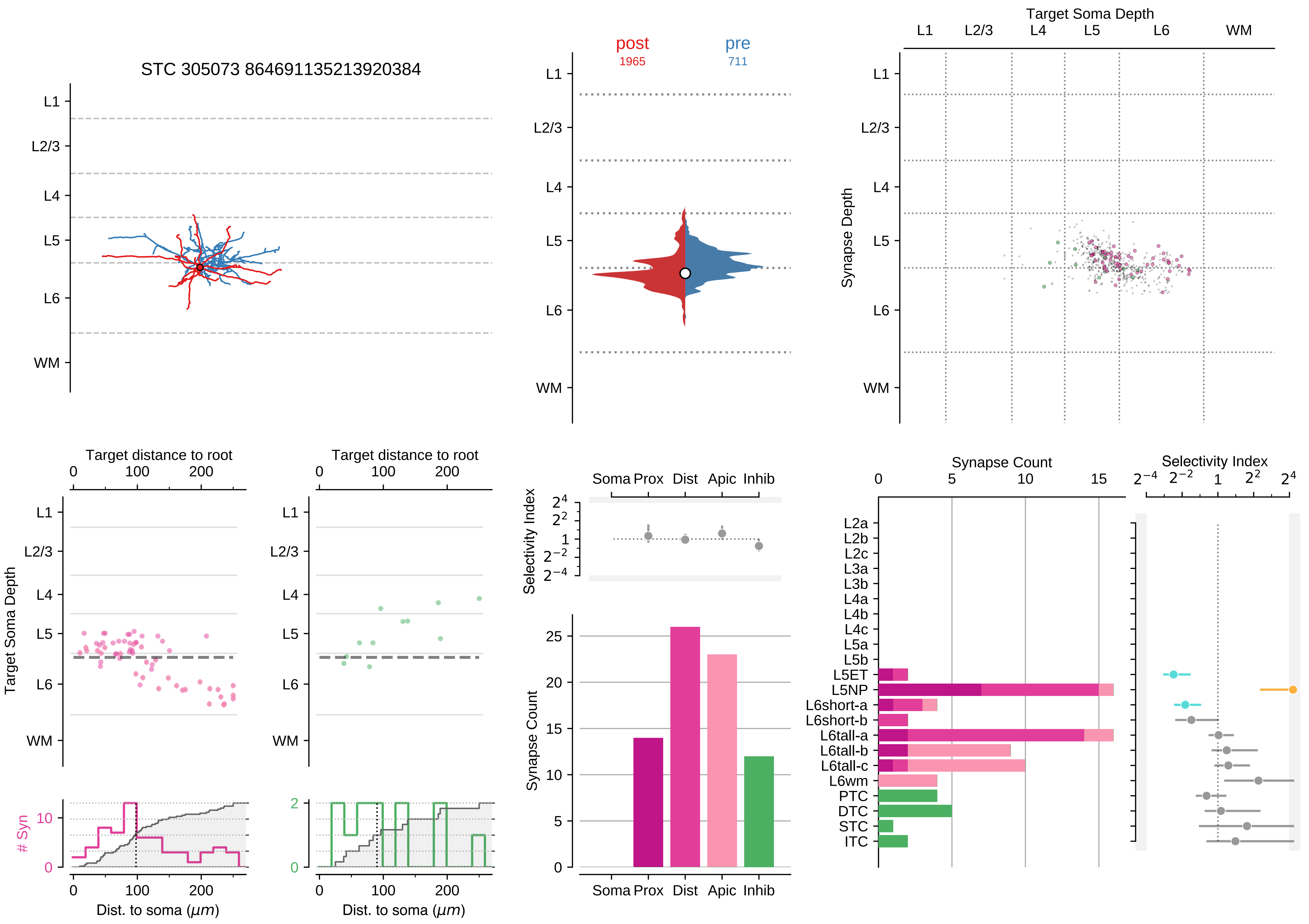

STC 305073 864691135213920384
post
1965
pre
711
Target Soma Depth
Synapse Depth
L1
L2/3
L4
L5
L6
WM
Target distance to root
Target Soma Depth
### Syn
Dist. to soma ($\mu m$)
Selectivity Index
Soma Prox Dist Apic Inhib
Synapse Count
L2a
L2b
L2c
L3a
L3b
L4a
L4b
L4c
L5a
L5b
L5ET
L5NP
L6short-a
L6short-b
L6tall-a
L6tall-b
L6tall-c
L6wm
PTC
DTC
STC
ITC

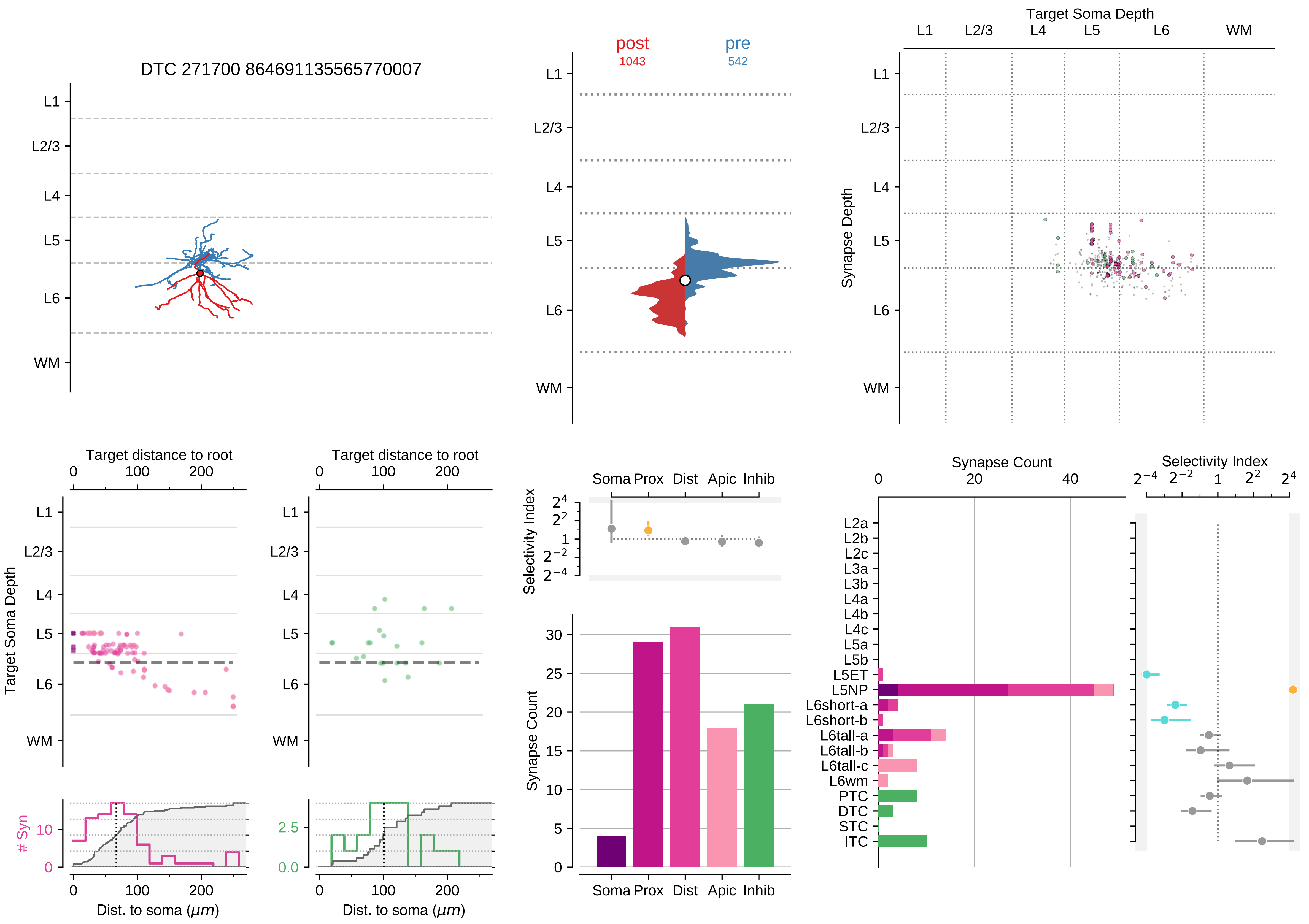

DTC 271700 864691135565770007
L1
L2/3
L4
L5
L6
WM
post
1043
pre
542
Target Soma Depth
Synapse Depth
Target distance to root
Dist. to soma ($\mu m$)
### Syn
Selectivity Index
Soma Prox Dist Apic Inhib
Synapse Count
L2a
L2b
L2c
L3a
L3b
L4a
L4b
L4c
L5a
L5b
L5ET
L5NP
L6short-a
L6short-b
L6tall-a
L6tall-b
L6tall-c
L6wm
PTC
DTC
STC
ITC

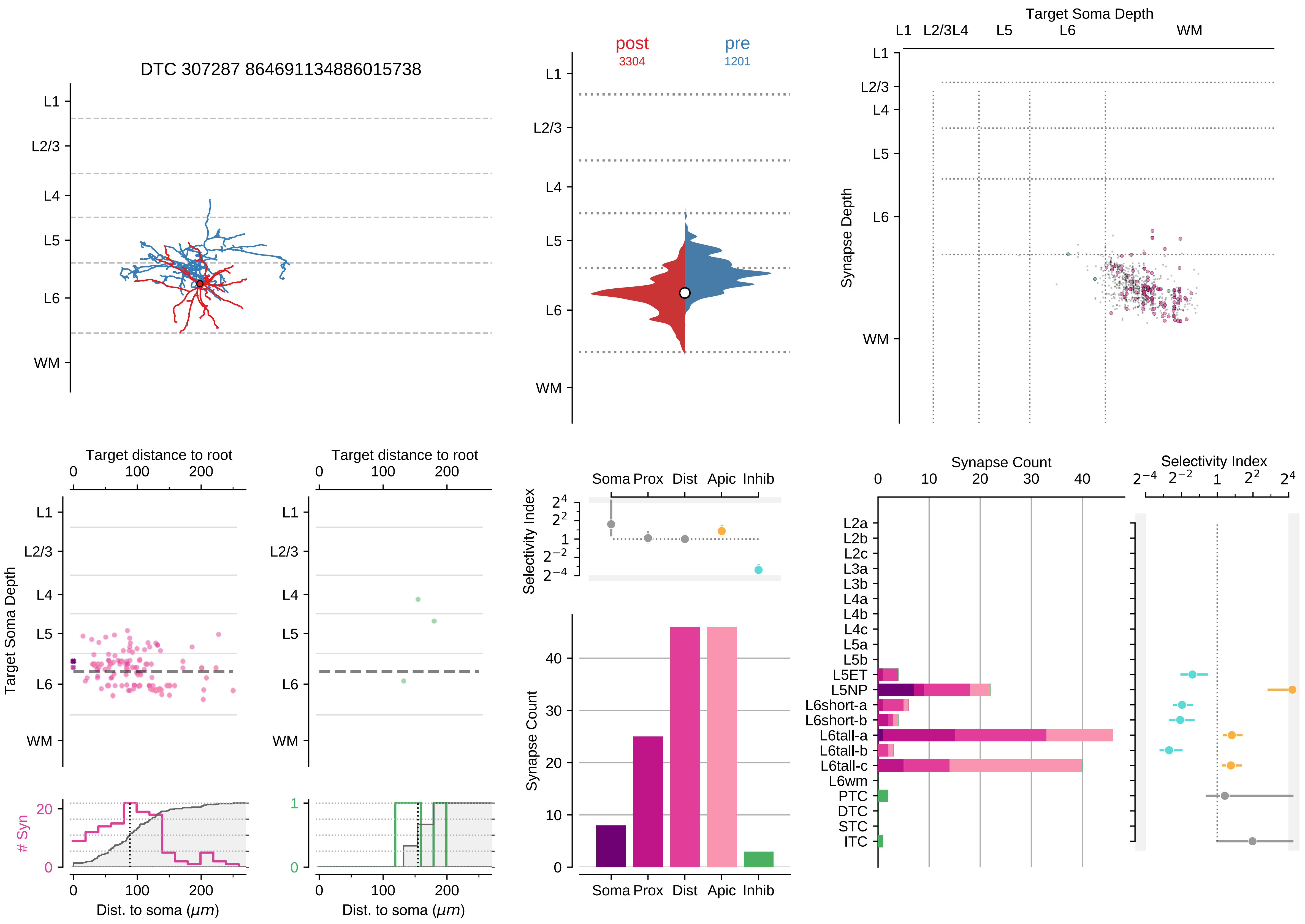

DTC 307287 864691134886015738
post
3304
pre
1201
Target Soma Depth
Synapse Depth
L1
L2/3
L4
L5
L6
WM
Target distance to root
Target Soma Depth
### Syn
Dist. to soma (μm)
Selectivity Index
Synapse Count
Soma
Prox
Dist
Apic
Inhib
L2a
L2b
L2c
L3a
L3b
L4a
L4b
L4c
L5a
L5b
L5ET
L5NP
L6short-a
L6short-b
L6tall-a
L6tall-b
L6tall-c
L6wm
PTC
DTC
STC
ITC

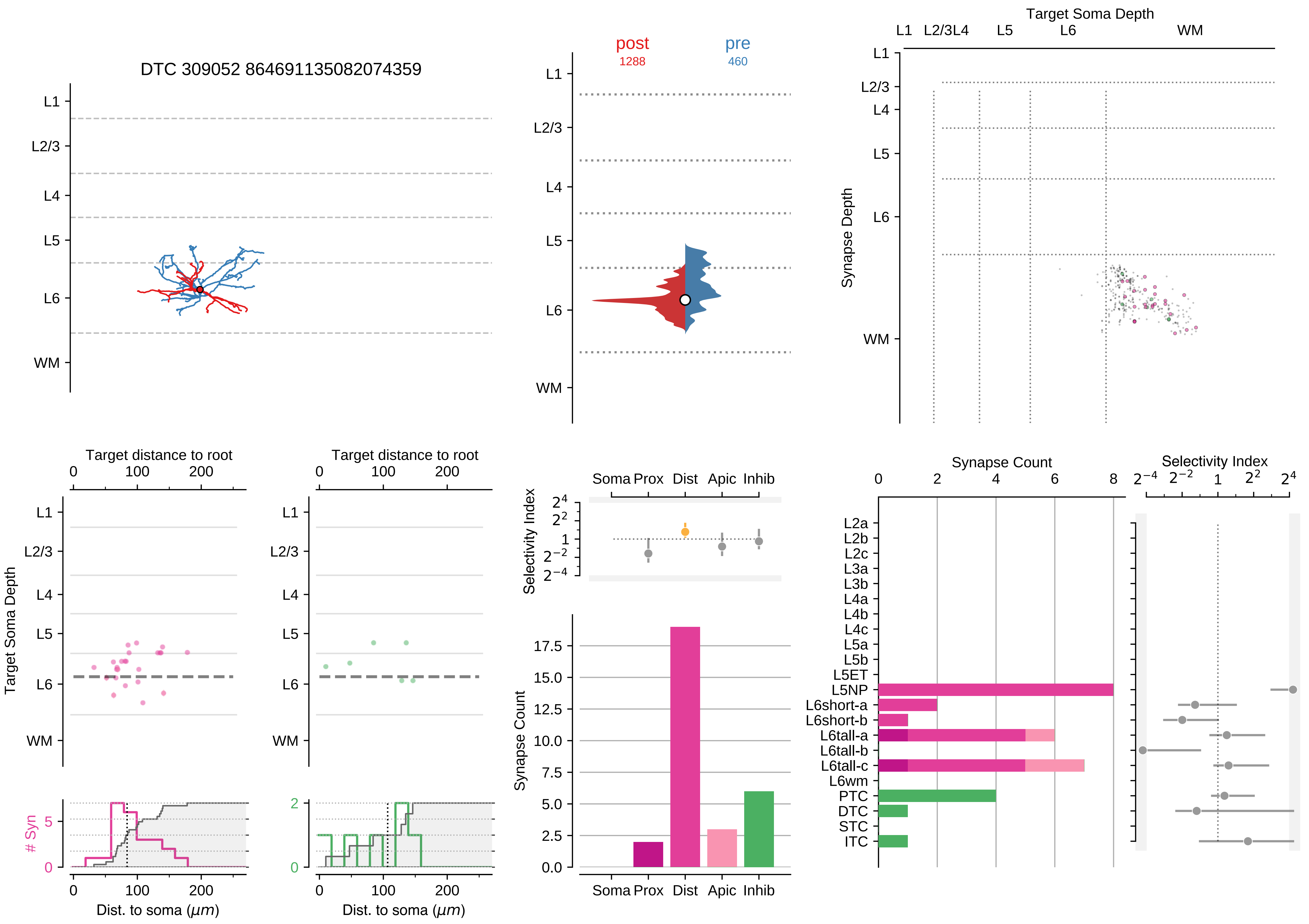

DTC 309052 864691135082074359
post
1288
pre
460
Target Soma Depth
Synapse Depth
L1
L2/3
L4
L5
L6
WM
Target distance to root
Target Soma Depth
### Syn
Dist. to soma ($\mu m$)
Selectivity Index
Soma Prox Dist Apic Inhib
Synapse Count
Synapse Count
Selectivity Index
L2a
L2b
L2c
L3a
L3b
L4a
L4b
L4c
L5a
L5b
L5ET
L5NP
L6short-a
L6short-b
L6tall-a
L6tall-b
L6tall-c
L6wm
PTC
DTC
STC
ITC

### Motif Group 15

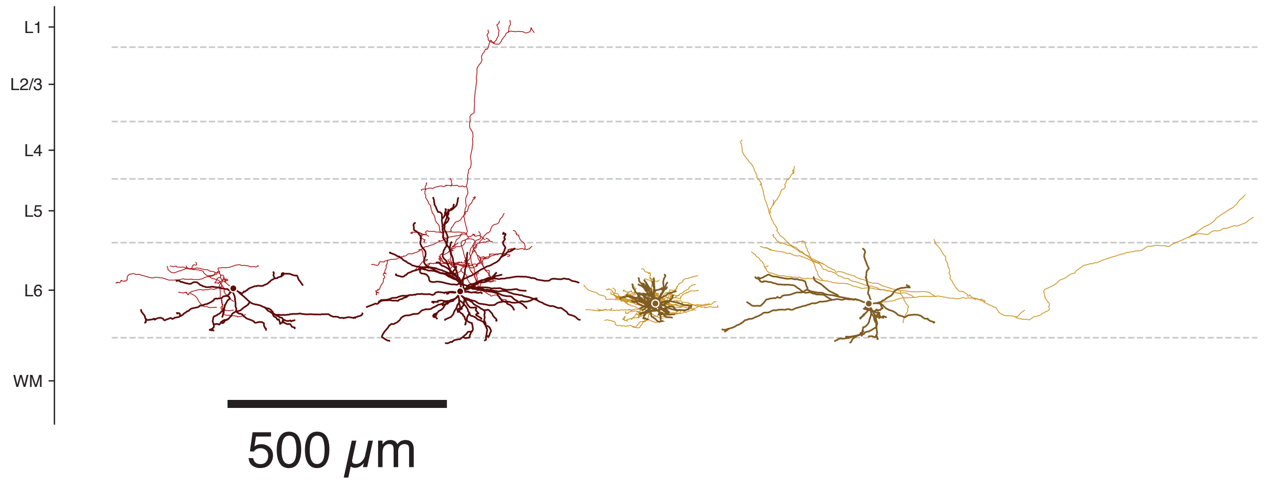

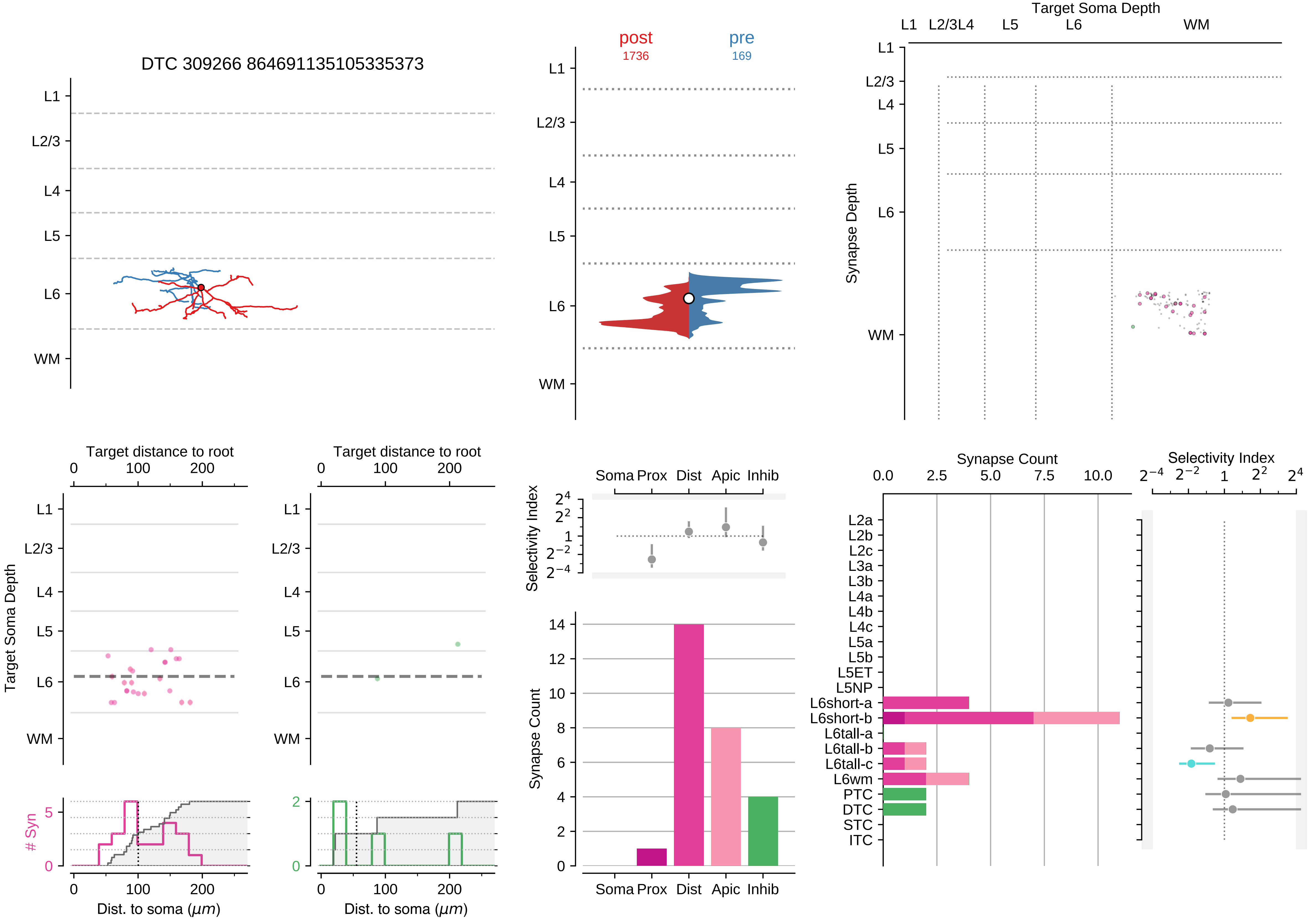

DTC 309266 864691135105335373
L1
L2/3
L4
L5
L6
WM
post
1736
pre
169
Target Soma Depth
Synapse Depth
Target distance to root
Dist. to soma (μm)
### Syn
Selectivity Index
Synapse Count
Soma
Prox
Dist
Apic
Inhib
L2a
L2b
L2c
L3a
L3b
L4a
L4b
L4c
L5a
L5b
L5ET
L5NP
L6short-a
L6short-b
L6tall-a
L6tall-b
L6tall-c
L6wm
PTC
DTC
STC
ITC

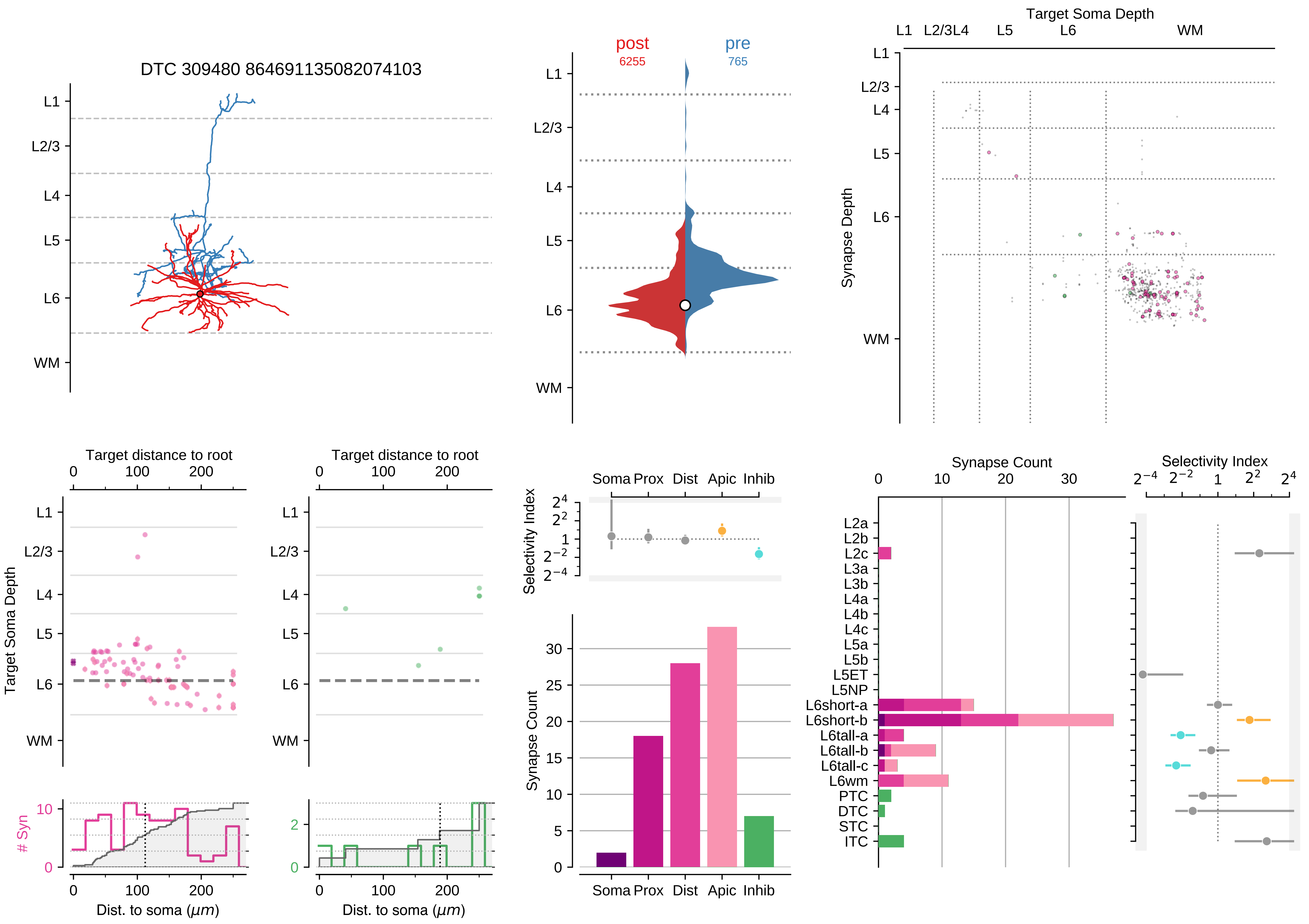

DTC 309480 864691135082074103
post
6255
pre
765
Target Soma Depth
Synapse Depth
L1
L2/3
L4
L5
L6
WM
Target distance to root
Target Soma Depth
### Syn
Dist. to soma (μm)
Selectivity Index
Soma Prox Dist Apic Inhib
Synapse Count
L2a
L2b
L2c
L3a
L3b
L4a
L4b
L4c
L5a
L5b
L5ET
L5NP
L6short-a
L6short-b
L6tall-a
L6tall-b
L6tall-c
L6wm
PTC
DTC
STC
ITC

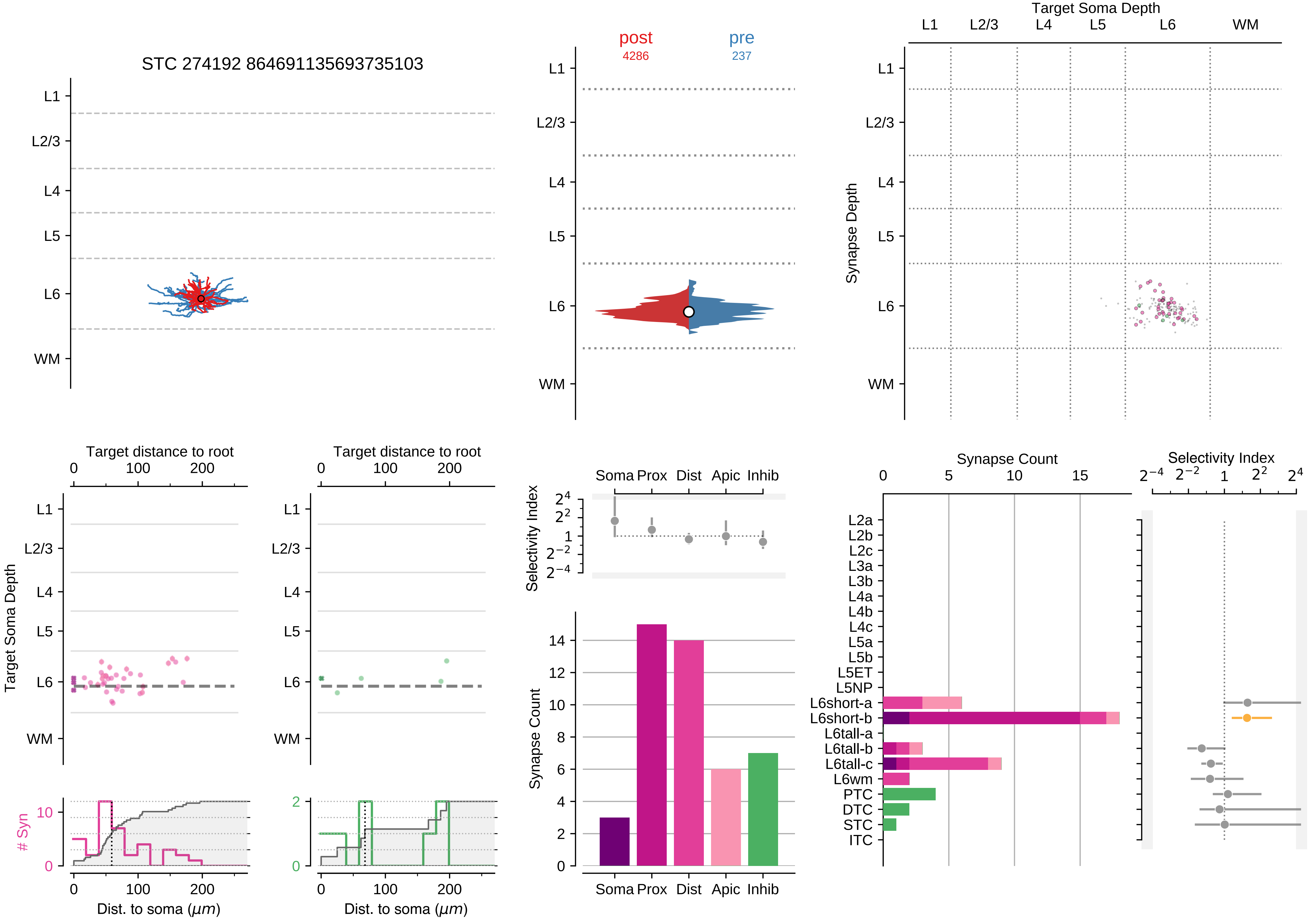

STC 274192 864691135693735103
post
4286
pre
237
Target Soma Depth
Synapse Depth
L1
L2/3
L4
L5
L6
WM
Target distance to root
Dist. to soma ($\mu m$)
### Syn
Selectivity Index
Synapse Count
Soma
Prox
Dist
Apic
Inhib
L2a
L2b
L2c
L3a
L3b
L4a
L4b
L4c
L5a
L5b
L5ET
L5NP
L6short-a
L6short-b
L6tall-a
L6tall-b
L6tall-c
L6wm
PTC
DTC
STC
ITC

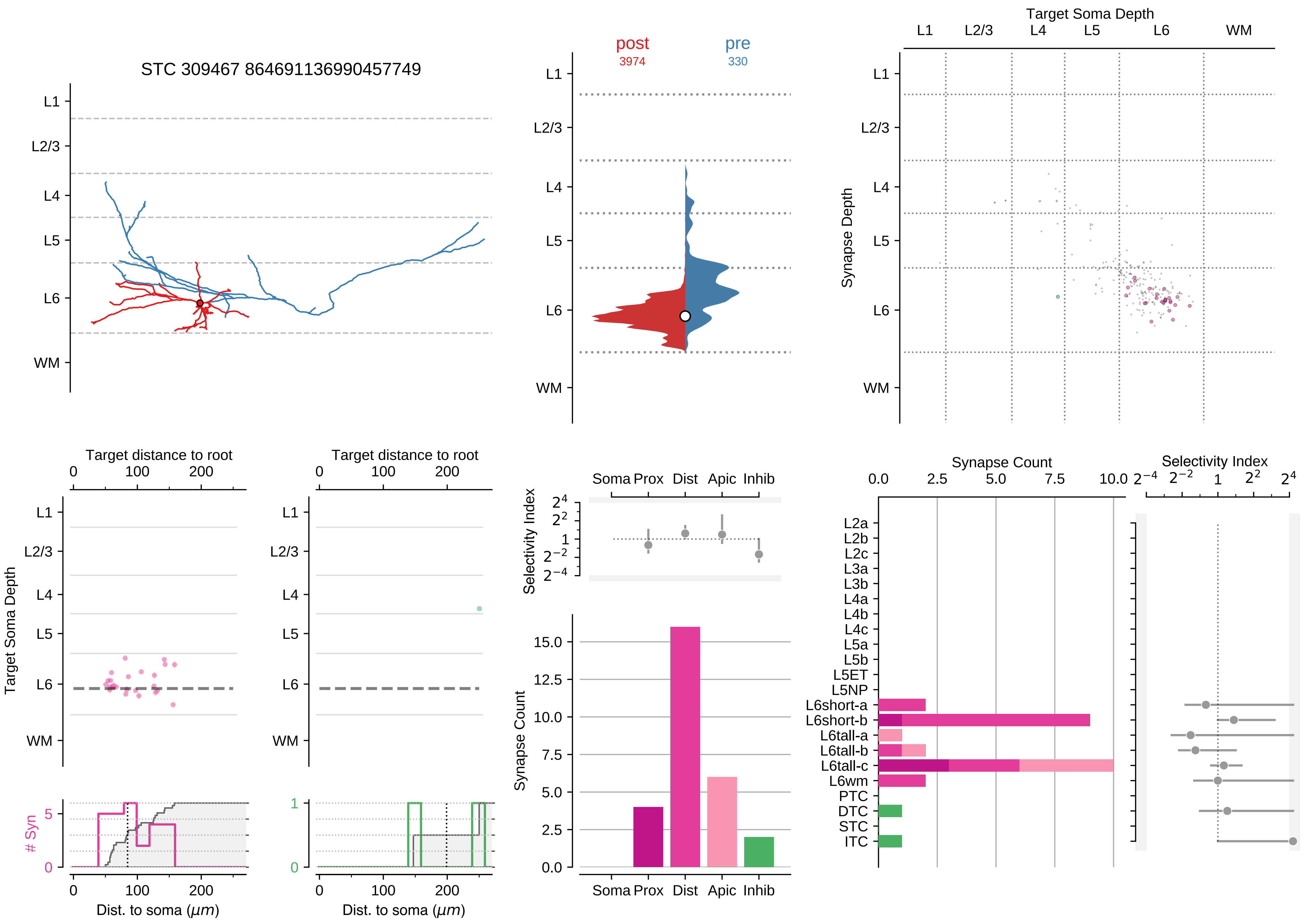
STC 309467 864691136990457749
L1
L2/3
L4
L5
L6
WM
post
3974
pre
330
Target Soma Depth
Synapse Depth
Target distance to root
0
100
200
Target Soma Depth
### Syn
Dist. to soma (μm)
Selectivity Index
Soma Prox Dist Apic Inhib
Synapse Count
Synapse Count
0.0
2.5
5.0
7.5
10.0
Selectivity Index
L2a
L2b
L2c
L3a
L3b
L4a
L4b
L4c
L5a
L5b
L5ET
L5NP
L6short-a
L6short-b
L6tall-a
L6tall-b
L6tall-c
L6wm
PTC
DTC
STC
ITC

### Motif Group 16

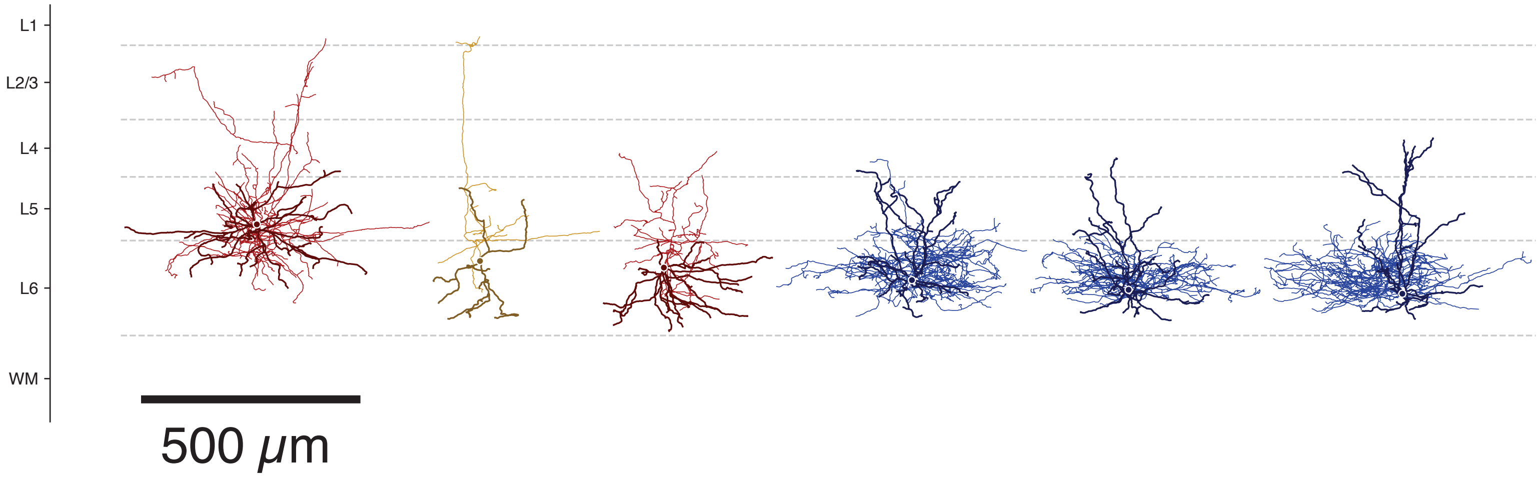

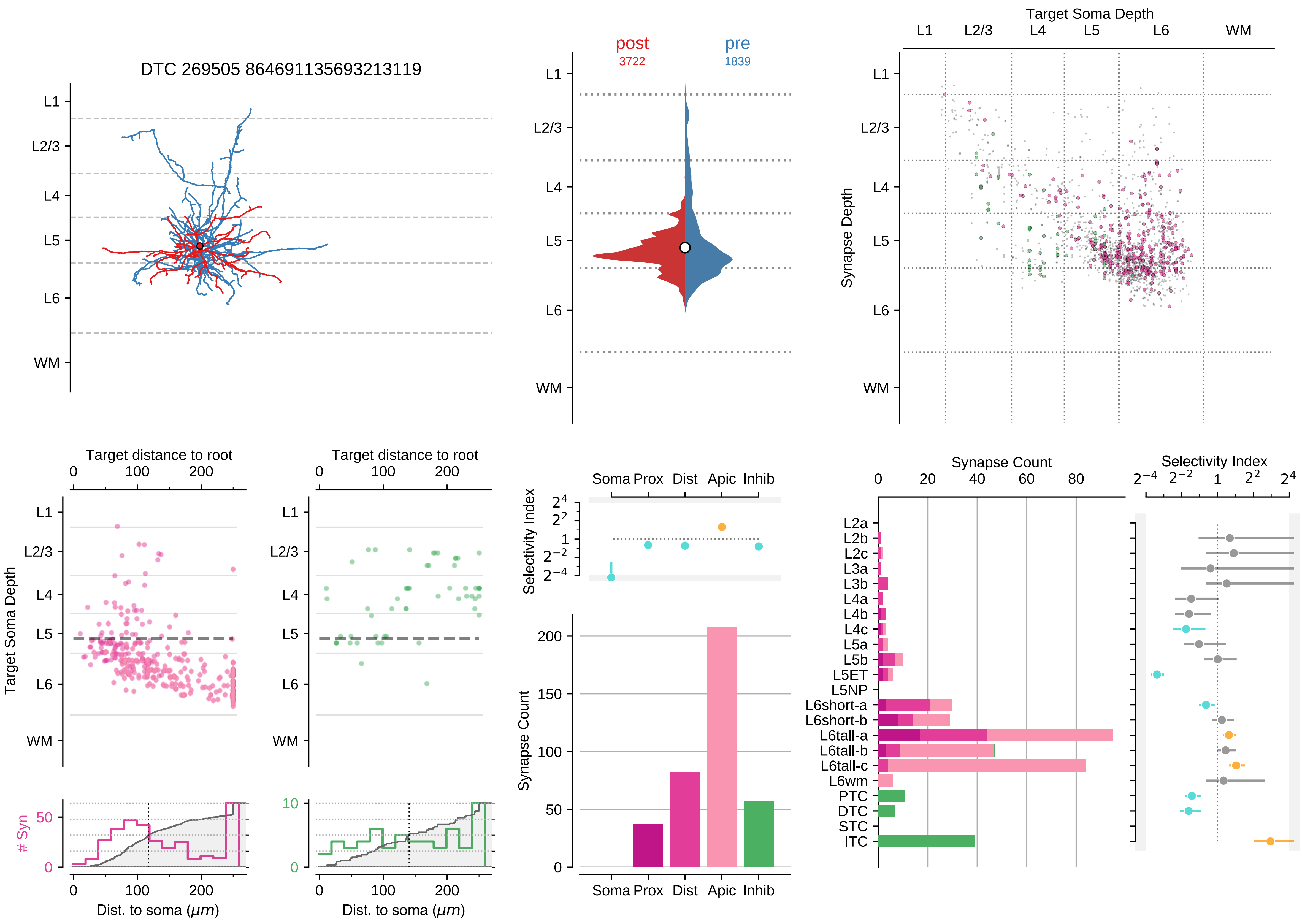

DTC 269505 864691135693213119
post
3722
pre
1839
Target Soma Depth
L1
L2/3
L4
L5
L6
WM
Synapse Depth
Target distance to root
Target Soma Depth
### Syn
Dist. to soma (μm)
Selectivity Index
Soma
Prox
Dist
Apic
Inhib
Synapse Count
Selectivity Index
L2a
L2b
L2c
L3a
L3b
L4a
L4b
L4c
L5a
L5b
L5ET
L5NP
L6short-a
L6short-b
L6tall-a
L6tall-b
L6tall-c
L6wm
PTC
DTC
STC
ITC

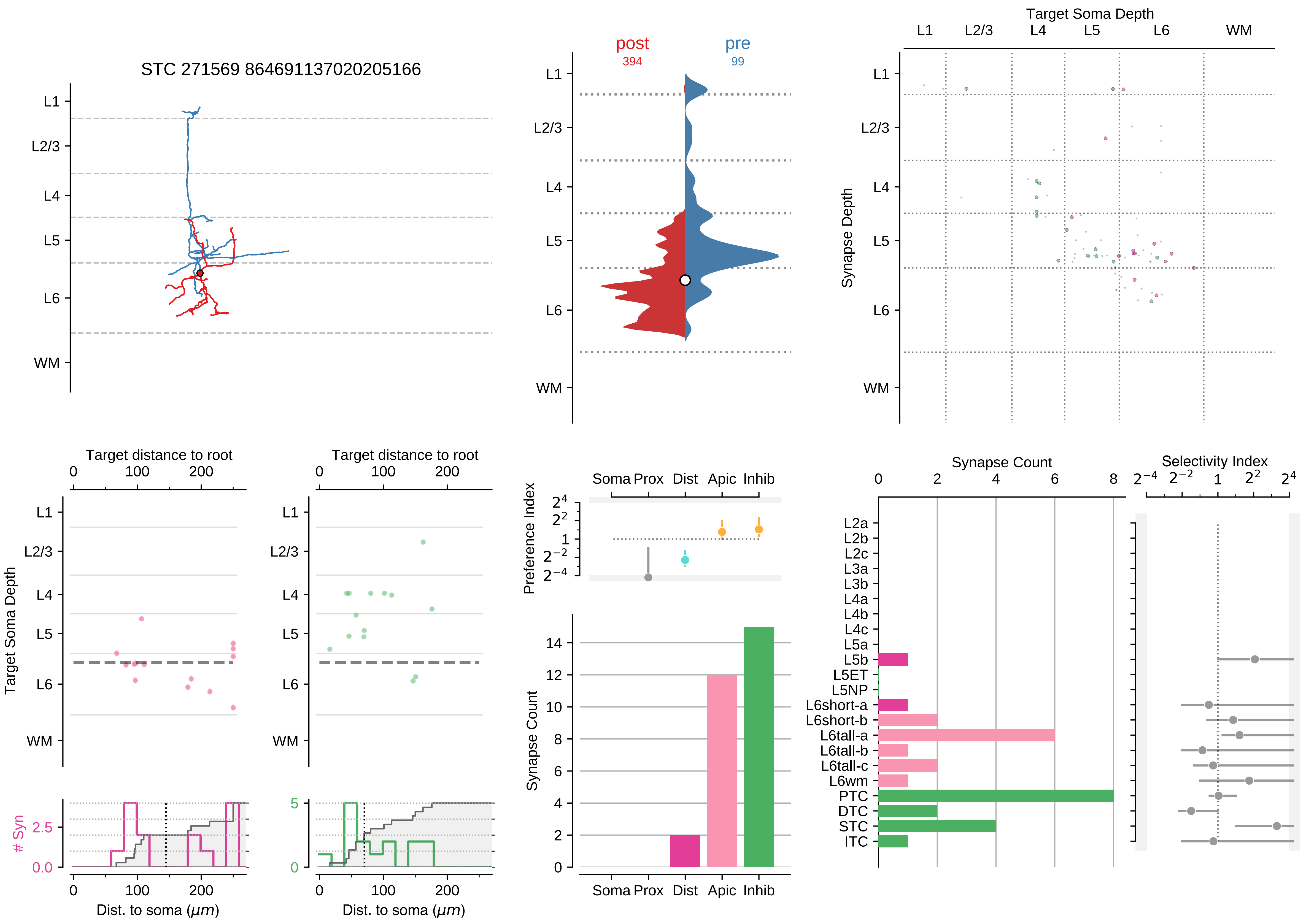

STC 271569 864691137020205166
L1
L2/3
L4
L5
L6
WM
post
394
pre
99
Target Soma Depth
Synapse Depth
Target distance to root
Target Soma Depth
### Syn
Dist. to soma ($\mu m$)
Soma Prox Dist Apic Inhib
Preference Index
Synapse Count
Selectivity Index
L2a
L2b
L2c
L3a
L3b
L4a
L4b
L4c
L5a
L5b
L5ET
L5NP
L6short-a
L6short-b
L6tall-a
L6tall-b
L6tall-c
L6wm
PTC
DTC
STC
ITC

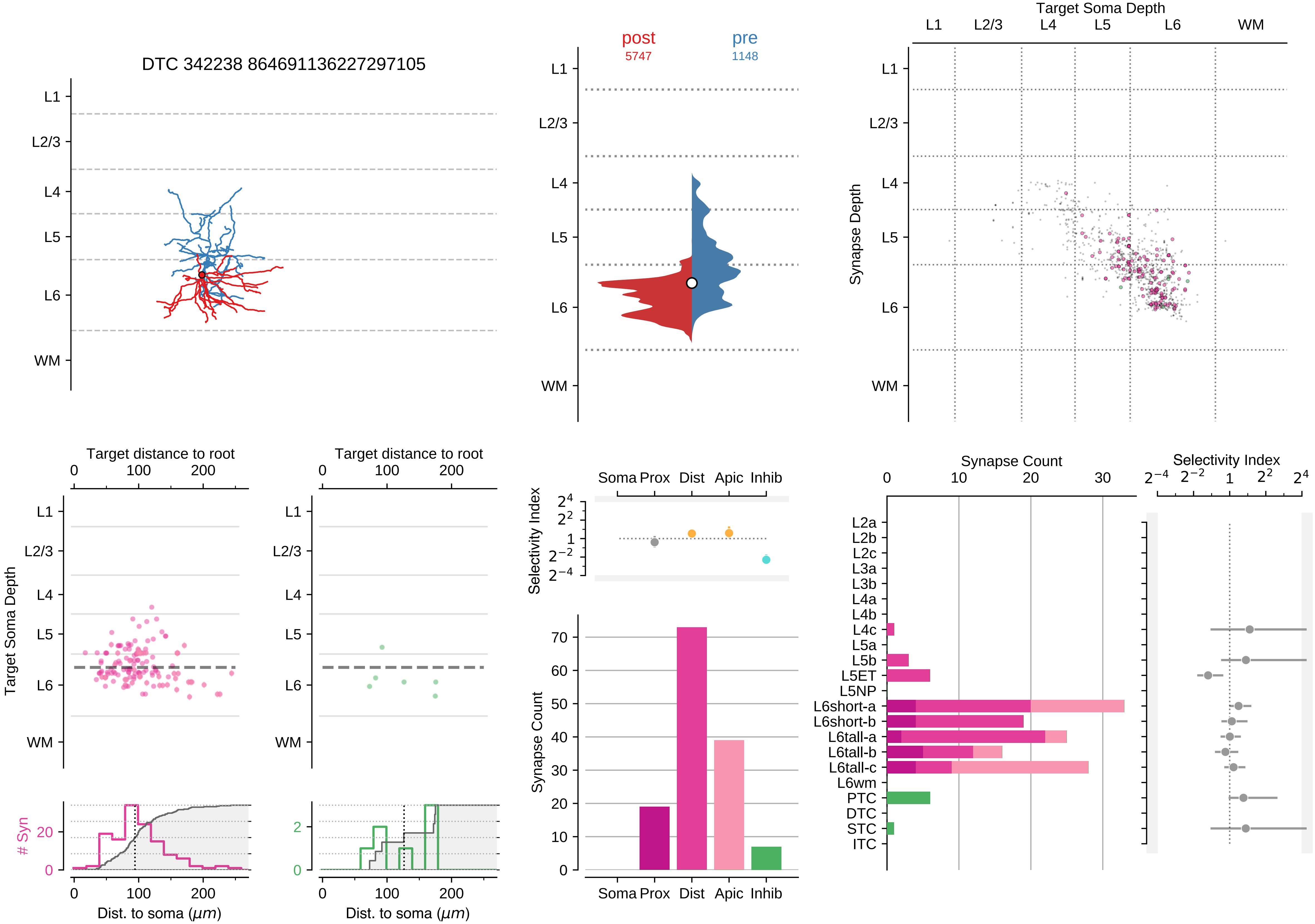

DTC 342238 864691136227297105
L1
L2/3
L4
L5
L6
WM
post
5747
pre
1148
Target Soma Depth
Synapse Depth
Target distance to root
0
100
200
Target Soma Depth
### Syn
Dist. to soma ($\mu m$)
Soma Prox Dist Apic Inhib
Selectivity Index
$2^4$
$2^2$
1
$2^{-2}$
$2^{-4}$
Synapse Count
0
10
20
30
40
50
60
70
L2a
L2b
L2c
L3a
L3b
L4a
L4b
L4c
L5a
L5b
L5ET
L5NP
L6short-a
L6short-b
L6tall-a
L6tall-b
L6tall-c
L6wm
PTC
DTC
STC
ITC

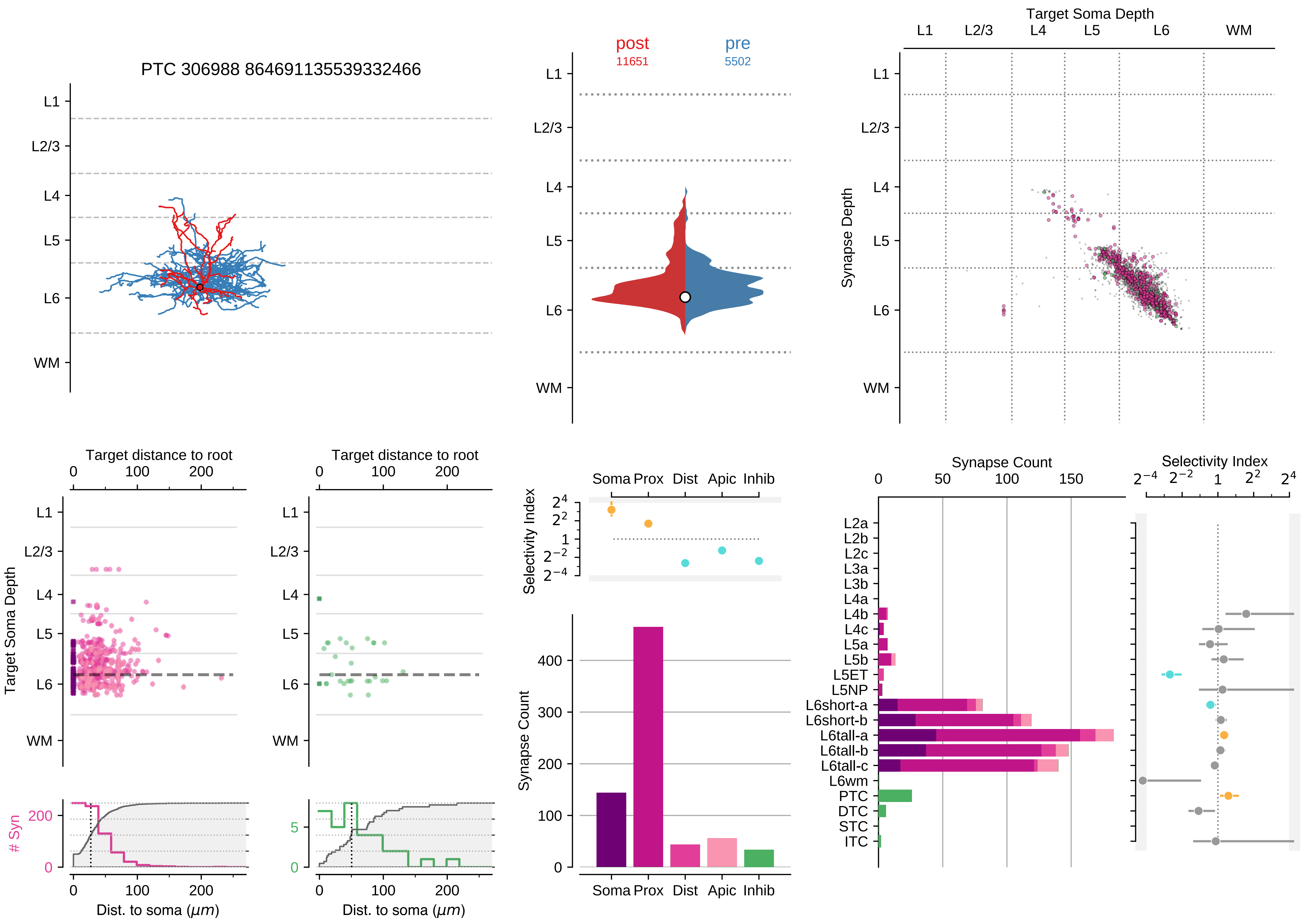

PTC 306988 864691135539332466
post
11651
pre
5502
Target Soma Depth
L1
L2/3
L4
L5
L6
WM
Synapse Depth
Target distance to root
Target Soma Depth
### Syn
Dist. to soma ($\mu m$)
Selectivity Index
Soma Prox Dist Apic Inhib
Synapse Count
L2a
L2b
L2c
L3a
L3b
L4a
L4b
L4c
L5a
L5b
L5ET
L5NP
L6short-a
L6short-b
L6tall-a
L6tall-b
L6tall-c
L6wm
PTC
DTC
STC
ITC

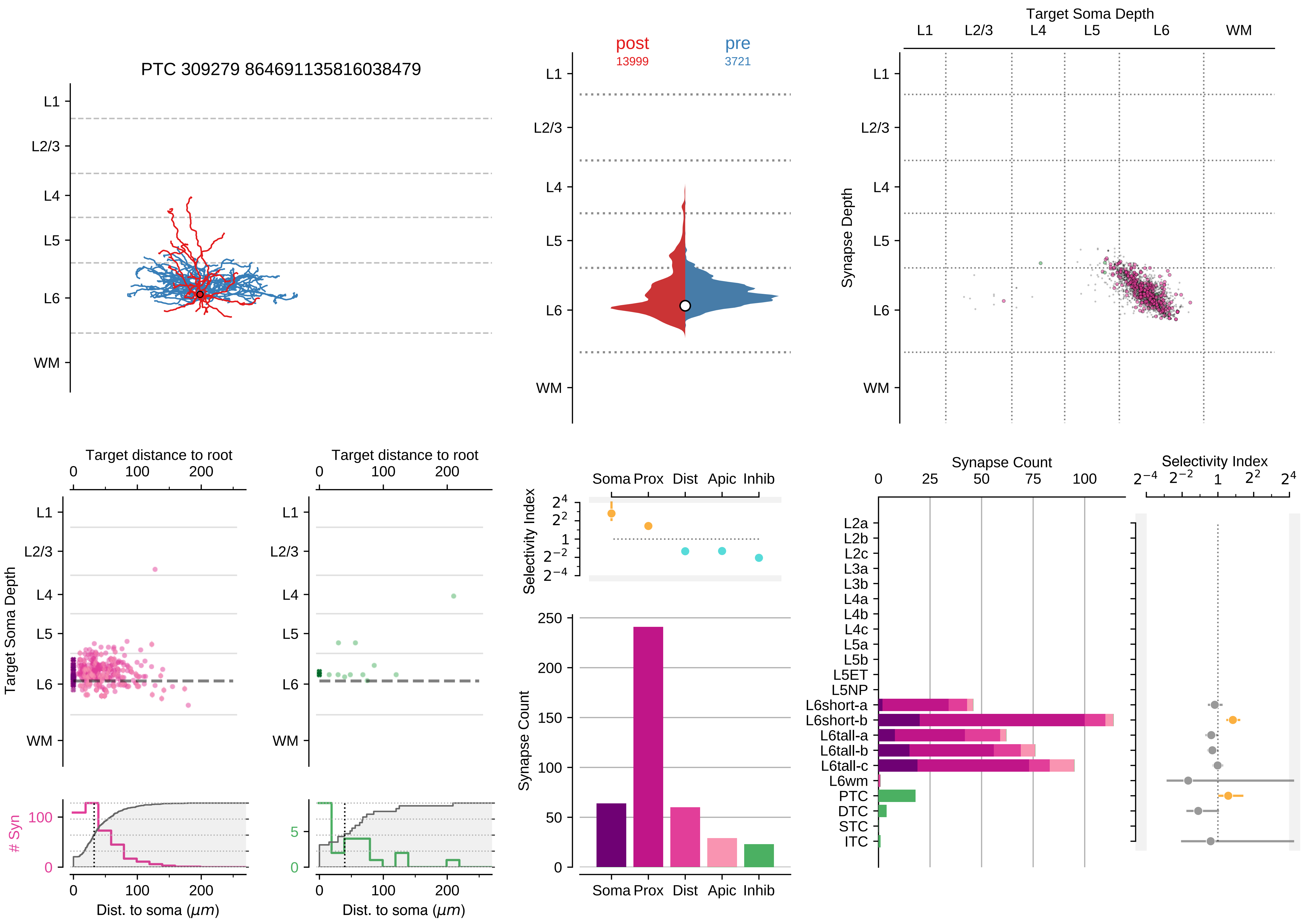

PTC 309279 864691135816038479
L1
L2/3
L4
L5
L6
WM
post
13999
pre
3721
Target Soma Depth
Synapse Depth
Target distance to root
Target Soma Depth
### Syn
Dist. to soma ($\mu m$)
Selectivity Index
Soma Prox Dist Apic Inhib
Synapse Count
L2a
L2b
L2c
L3a
L3b
L4a
L4b
L4c
L5a
L5b
L5ET
L5NP
L6short-a
L6short-b
L6tall-a
L6tall-b
L6tall-c
L6wm
PTC
DTC
STC
ITC

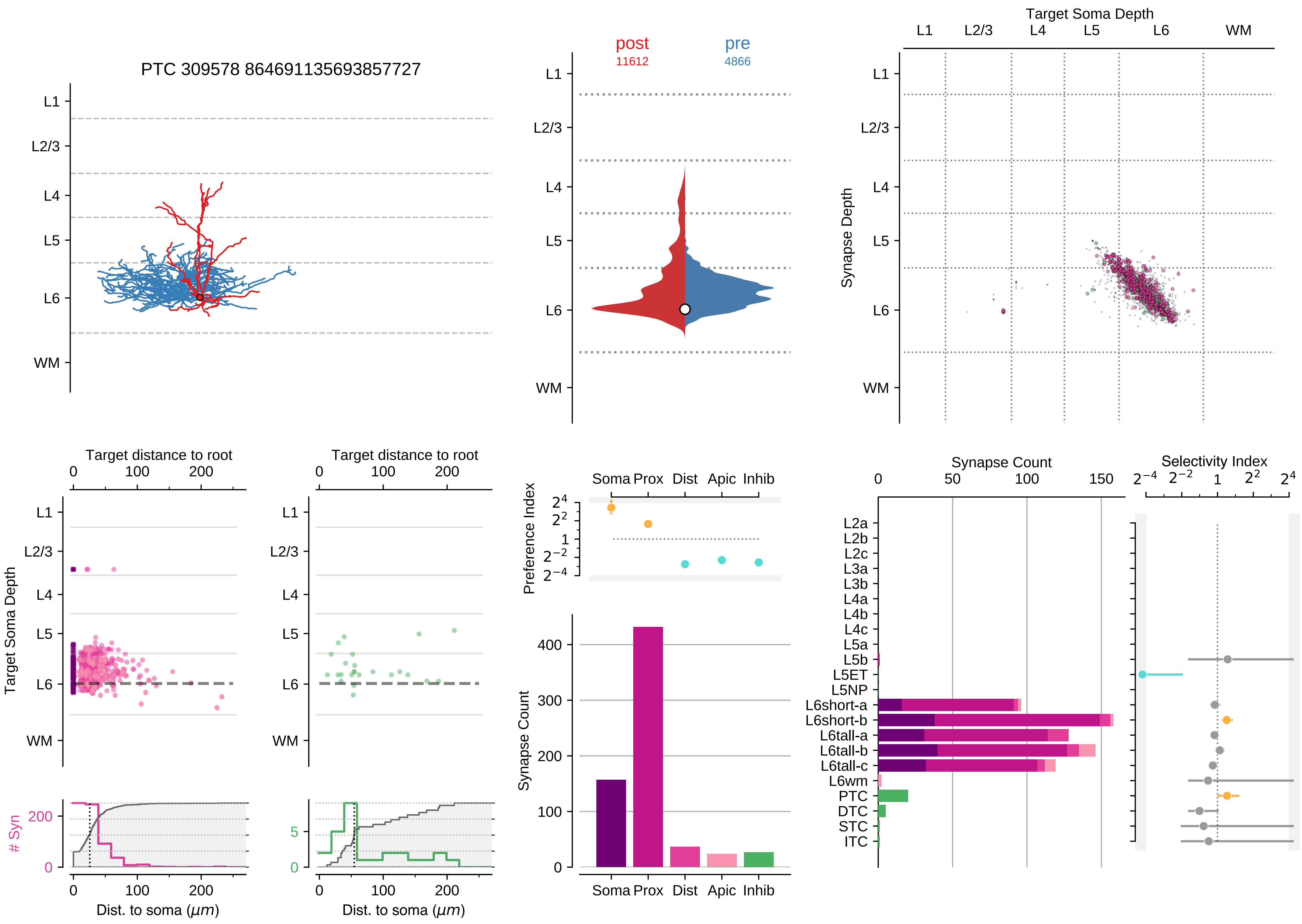

PTC 309578 864691135693857727
L1
L2/3
L4
L5
L6
WM
post
11612
pre
4866
Target Soma Depth
Synapse Depth
Target distance to root
Target Soma Depth
### Syn
Dist. to soma (μm)
Preference Index
Soma
Prox
Dist
Apic
Inhib
Synapse Count
Selectivity Index
L2a
L2b
L2c
L3a
L3b
L4a
L4b
L4c
L5a
L5b
L5ET
L5NP
L6short-a
L6short-b
L6tall-a
L6tall-b
L6tall-c
L6wm
PTC
DTC
STC
ITC

### Motif Group 17

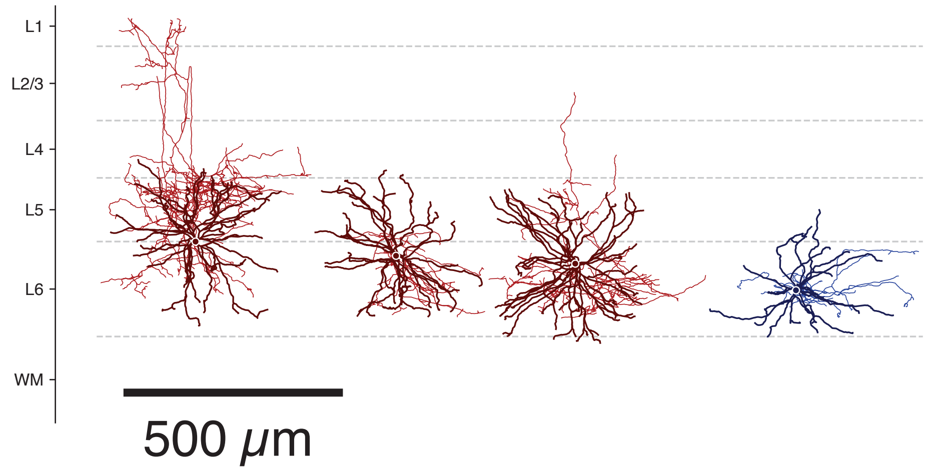

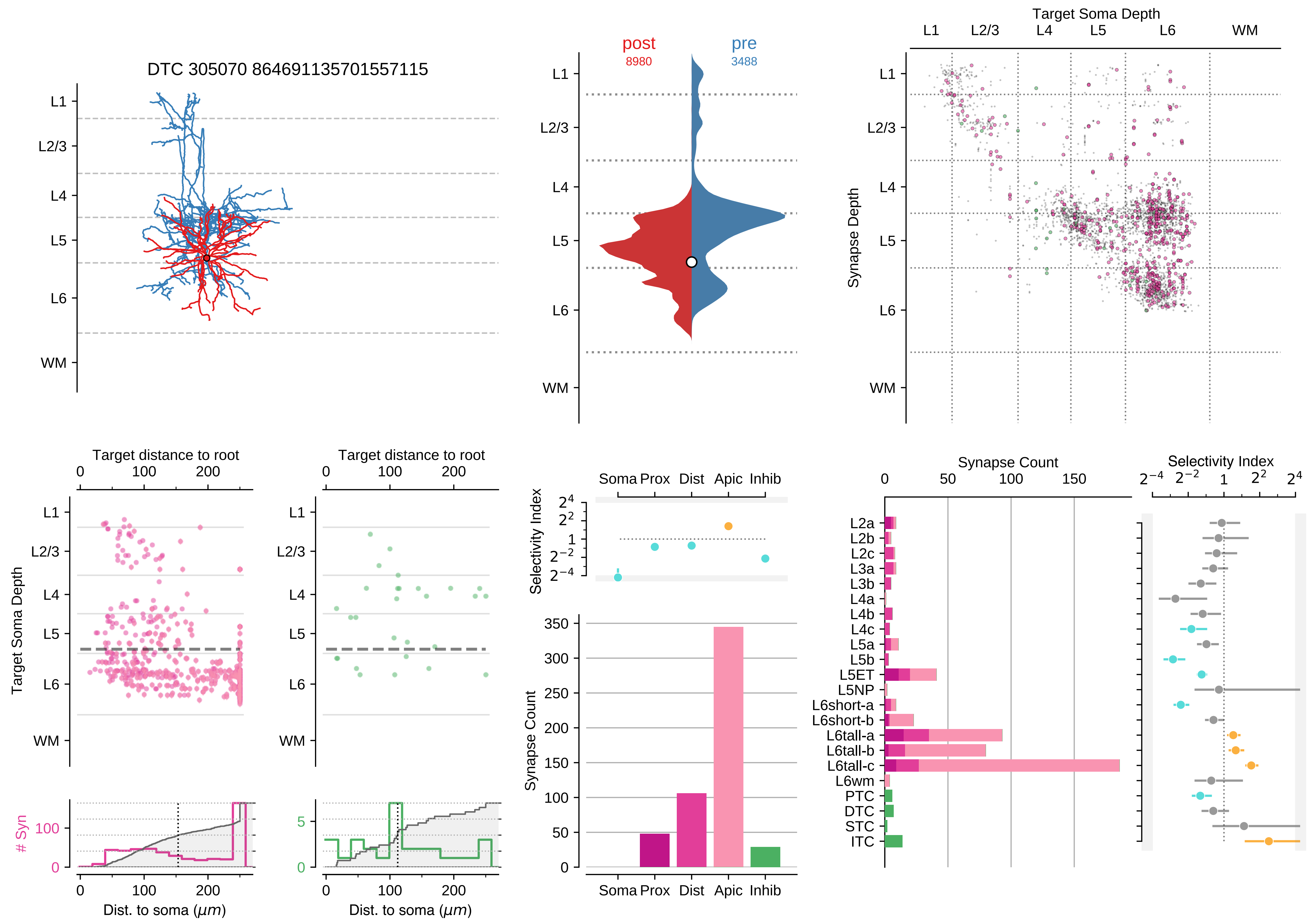

DTC 305070 864691135701557115
L1
L2/3
L4
L5
L6
WM
post
8980
pre
3488
Target Soma Depth
Synapse Depth
Target distance to root
Target Soma Depth
### Syn
Dist. to soma ($\mu m$)
Selectivity Index
Soma Prox Dist Apic Inhib
Synapse Count
L2a
L2b
L2c
L3a
L3b
L4a
L4b
L4c
L5a
L5b
L5ET
L5NP
L6short-a
L6short-b
L6tall-a
L6tall-b
L6tall-c
L6wm
PTC
DTC
STC
ITC

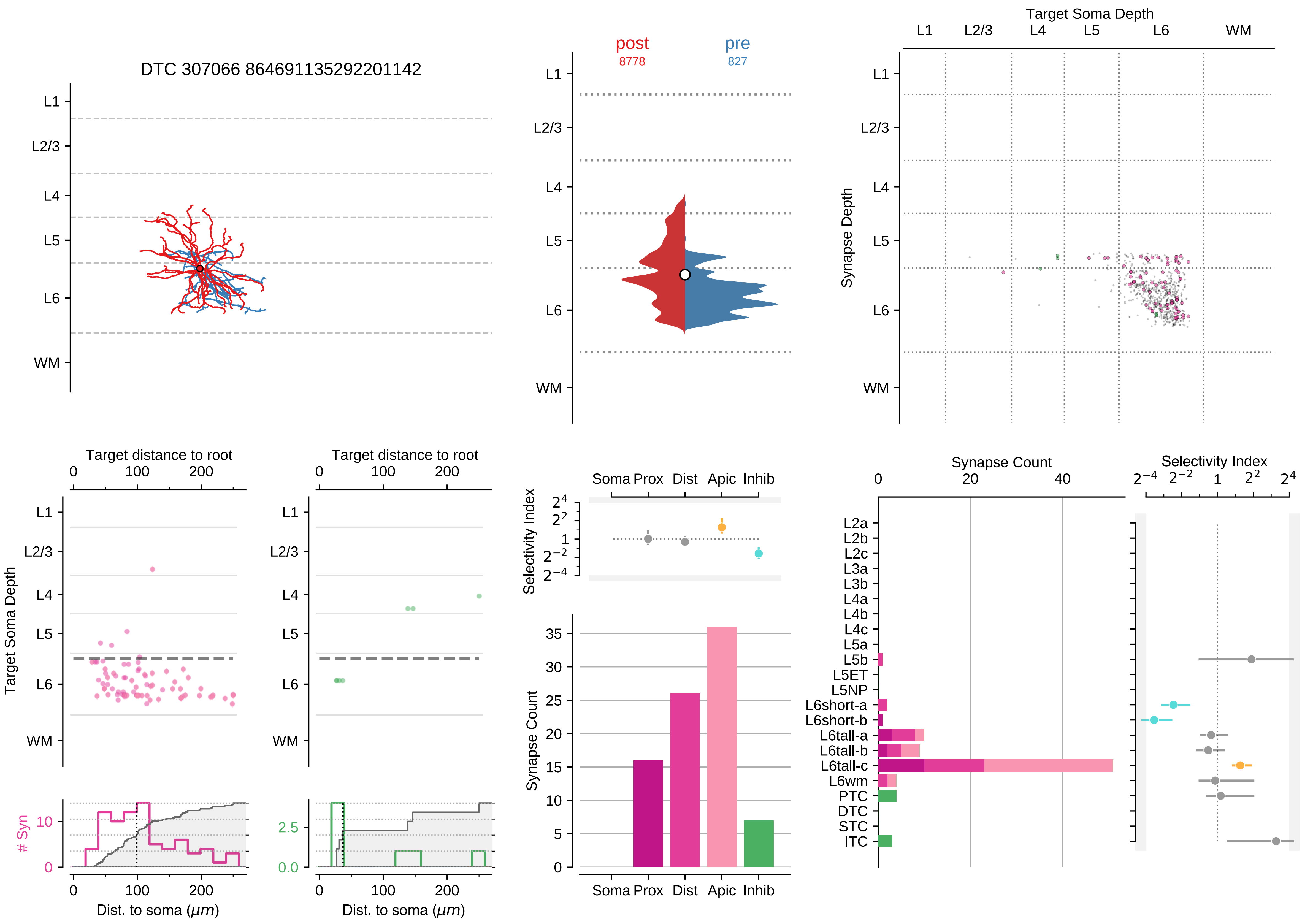

DTC 307066 864691135292201142
L1
L2/3
L4
L5
L6
WM
post
8778
pre
827
Target Soma Depth
Synapse Depth
Target distance to root
Target Soma Depth
### Syn
Dist. to soma ($\mu m$)
Selectivity Index
Soma Prox Dist Apic Inhib
Synapse Count
L2a
L2b
L2c
L3a
L3b
L4a
L4b
L4c
L5a
L5b
L5ET
L5NP
L6short-a
L6short-b
L6tall-a
L6tall-b
L6tall-c
L6wm
PTC
DTC
STC
ITC

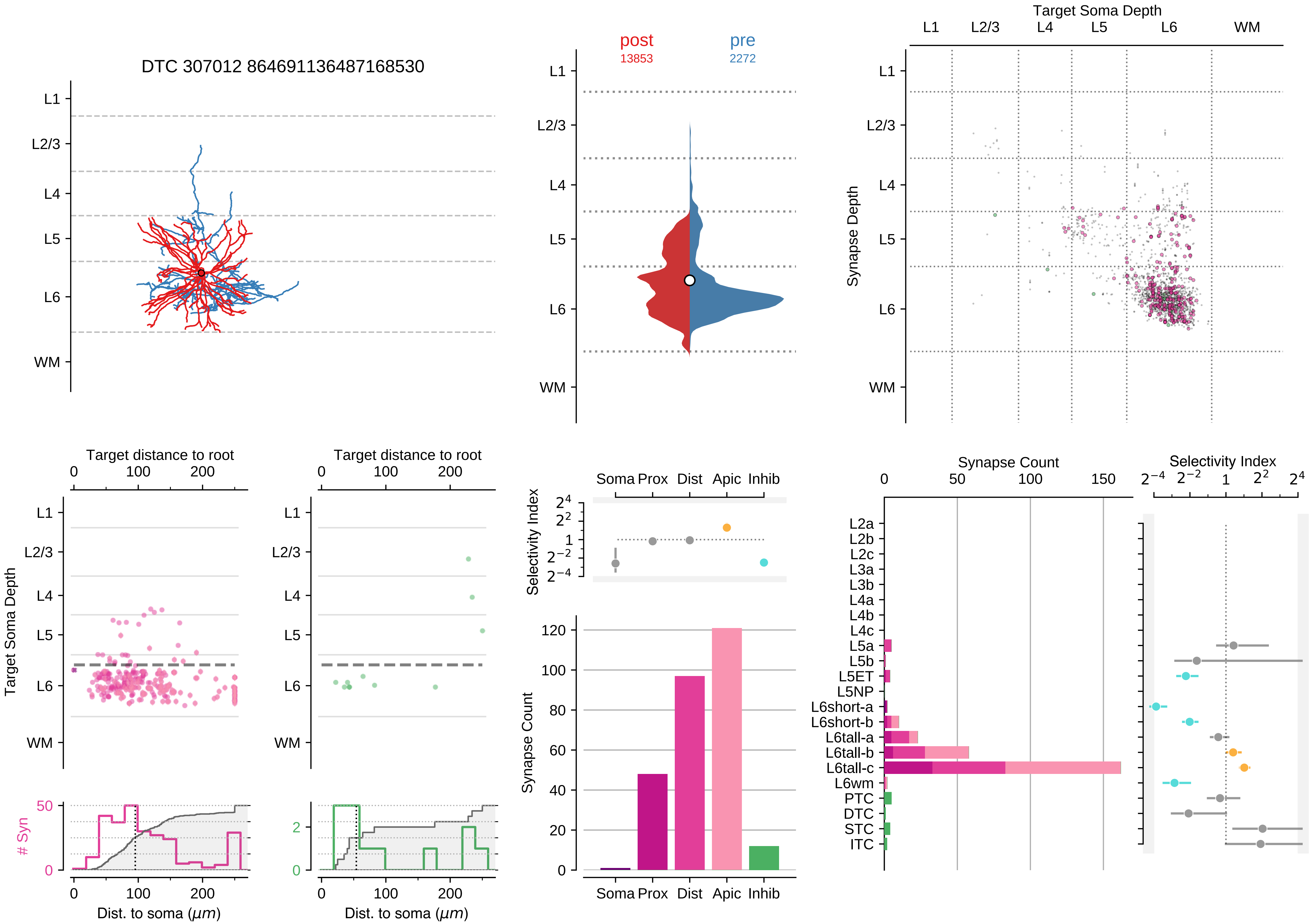

DTC 307012 864691136487168530
post
13853
pre
2272
Target Soma Depth
Synapse Depth
L1
L2/3
L4
L5
L6
WM
Target distance to root
Target Soma Depth
### Syn
Dist. to soma (μm)
Selectivity Index
Soma Prox Dist Apic Inhib
Synapse Count
Synapse Count
Selectivity Index
L2a
L2b
L2c
L3a
L3b
L4a
L4b
L4c
L5a
L5b
L5ET
L5NP
L6short-a
L6short-b
L6tall-a
L6tall-b
L6tall-c
L6wm
PTC
DTC
STC
ITC

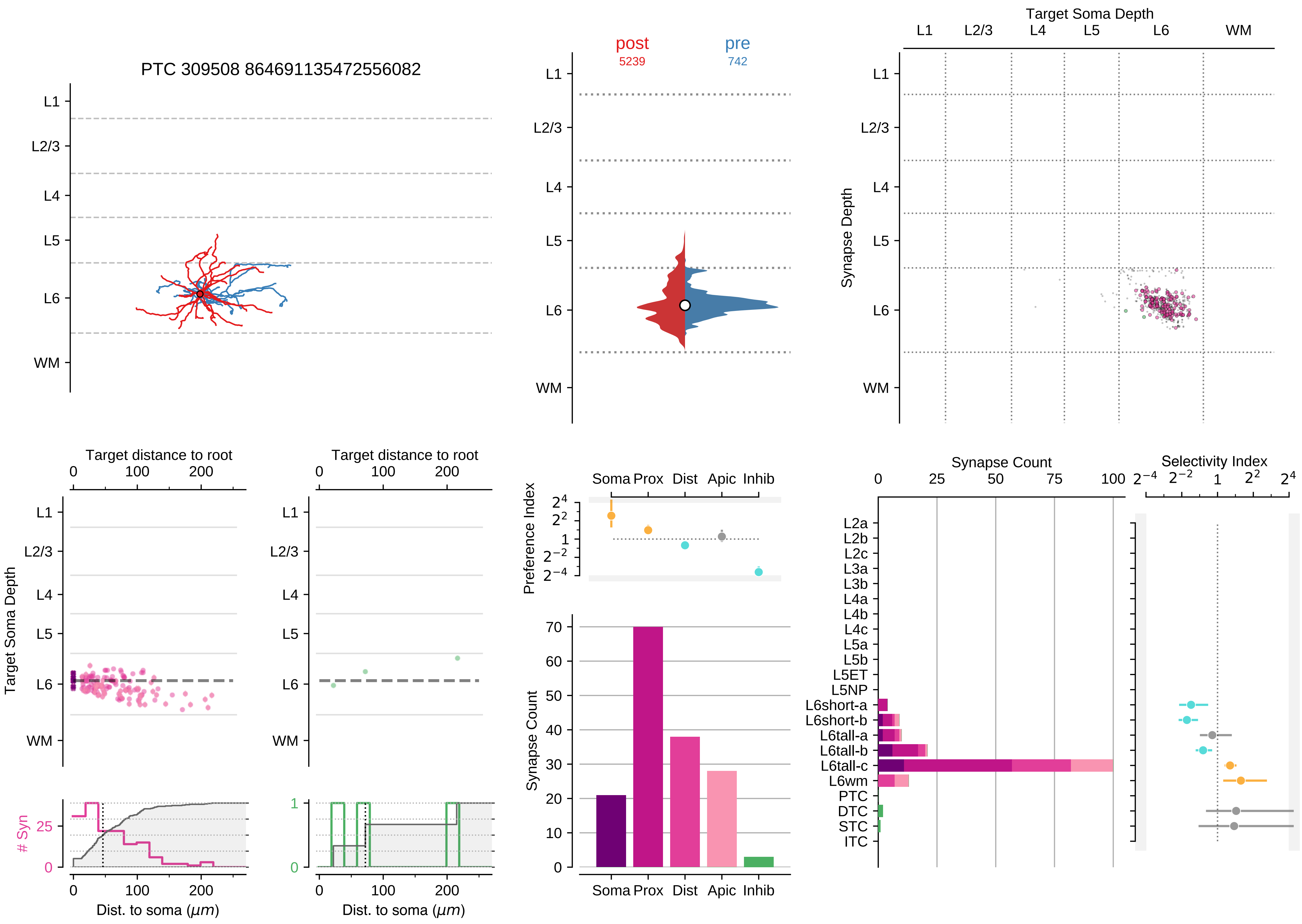
PTC 309508 864691135472556082
L1
L2/3
L4
L5
L6
WM
post
5239
pre
742
Target Soma Depth
Synapse Depth
Target distance to root
Target Soma Depth
Dist. to soma (μm)
### Syn
Preference Index
Soma
Prox
Dist
Apic
Inhib
Synapse Count
Selectivity Index
L2a
L2b
L2c
L3a
L3b
L4a
L4b
L4c
L5a
L5b
L5ET
L5NP
L6short-a
L6short-b
L6tall-a
L6tall-b
L6tall-c
L6wm
PTC
DTC
STC
ITC

### Motif Group 18

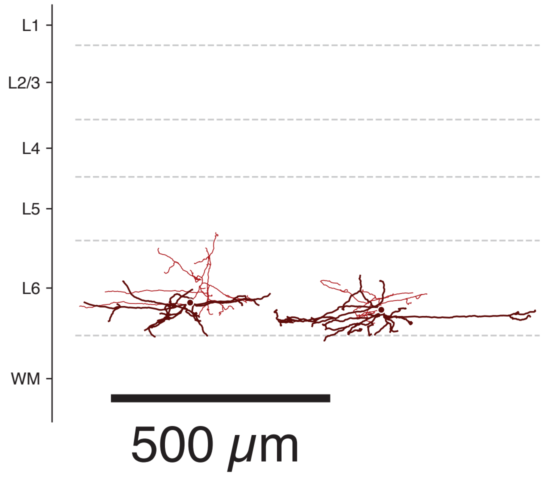

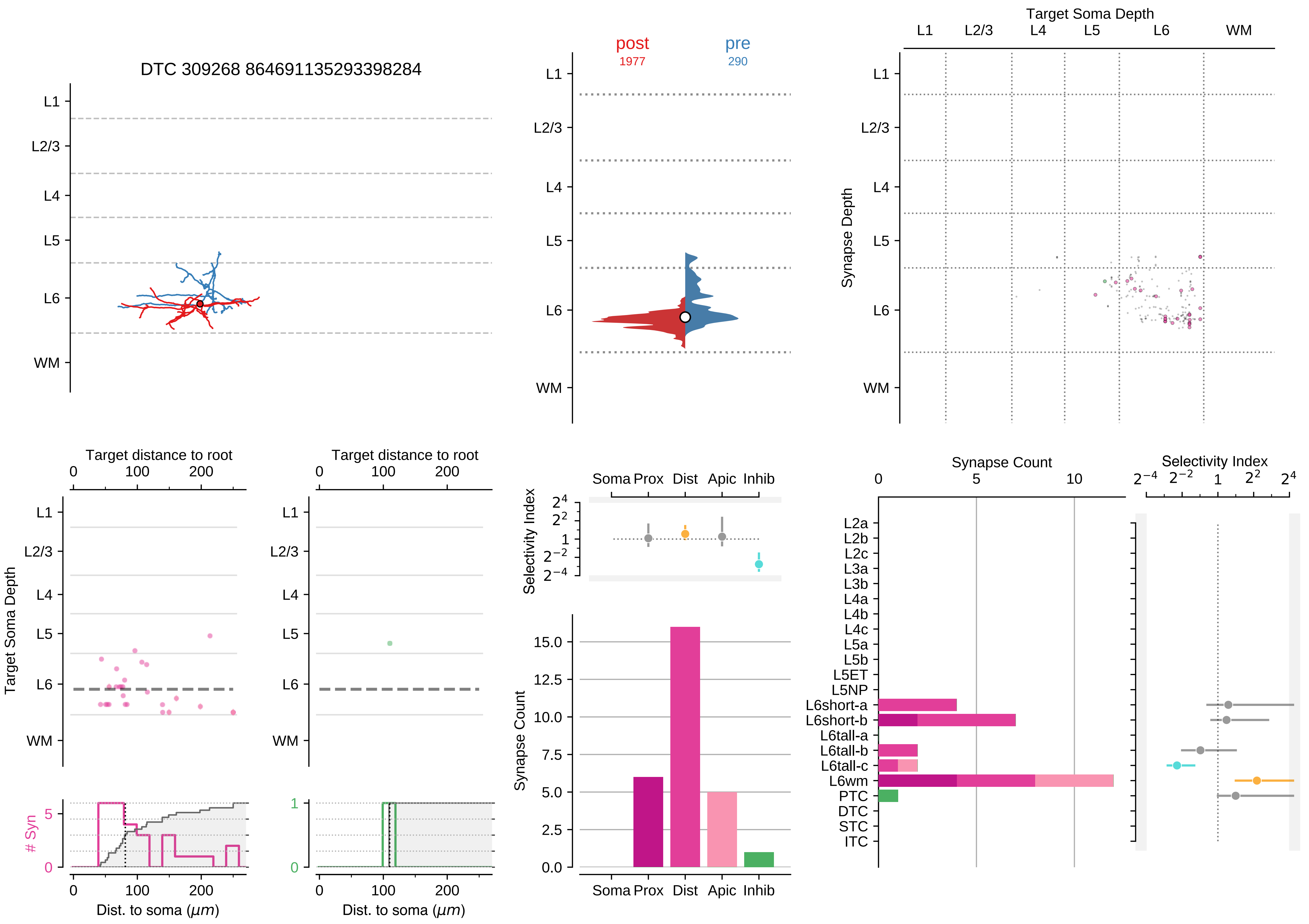

DTC 309268 864691135293398284
L1
L2/3
L4
L5
L6
WM
post
1977
pre
290
Target Soma Depth
Synapse Depth
Target distance to root
0
100
200
Target Soma Depth
### Syn
Dist. to soma (μm)
Soma Prox Dist Apic Inhib
Selectivity Index
Synapse Count
Synapse Count
Selectivity Index
L2a
L2b
L2c
L3a
L3b
L4a
L4b
L4c
L5a
L5b
L5ET
L5NP
L6short-a
L6short-b
L6tall-a
L6tall-b
L6tall-c
L6wm
PTC
DTC
STC
ITC

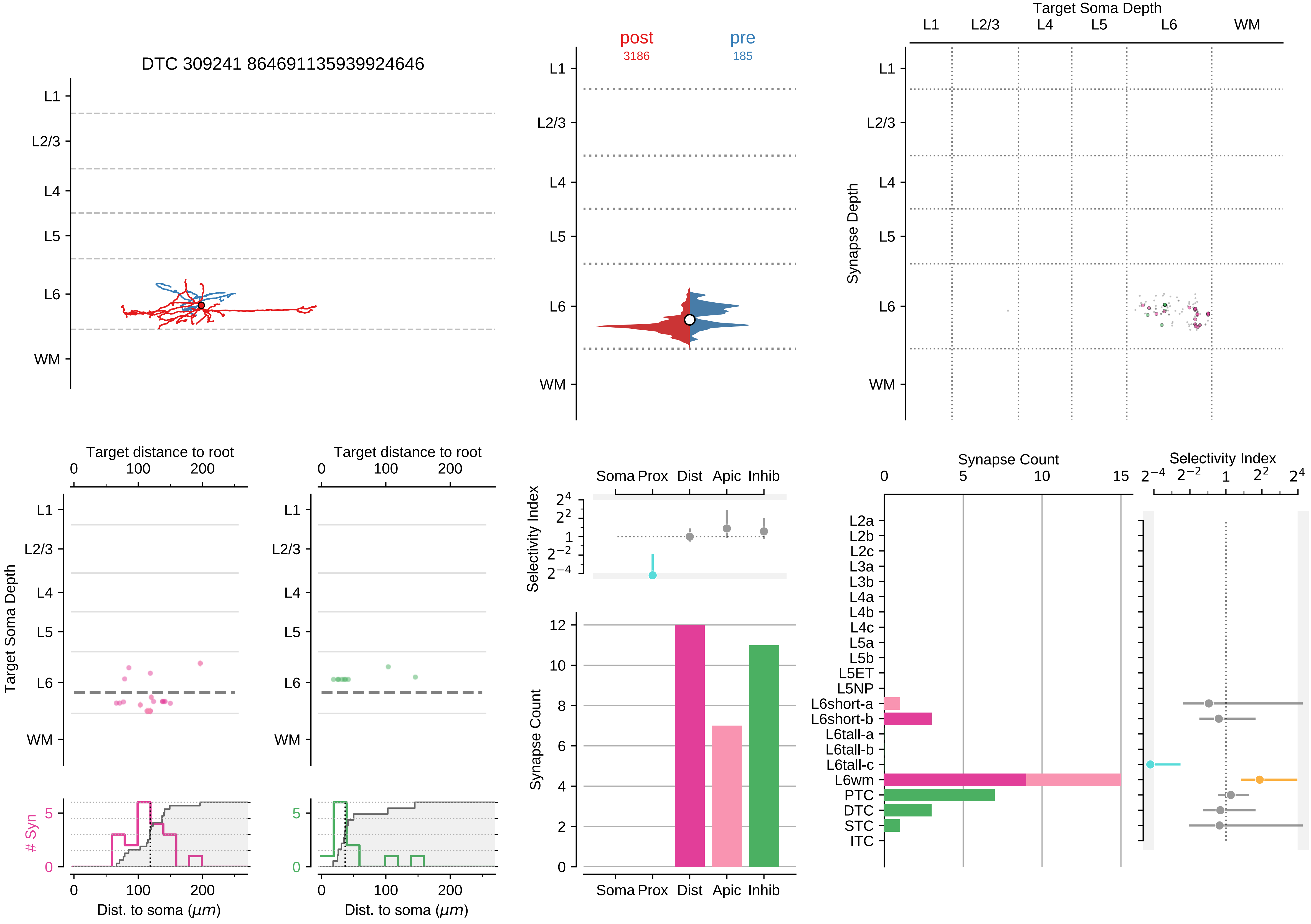

DTC 309241 864691135939924646
post
3186
pre
185
Target Soma Depth
L1
L2/3
L4
L5
L6
WM
Synapse Depth
Target distance to root
Target Soma Depth
### Syn
Dist. to soma ($\mu m$)
Selectivity Index
Soma Prox
Dist
Apic
Inhib
Synapse Count
L2a
L2b
L2c
L3a
L3b
L4a
L4b
L4c
L5a
L5b
L5ET
L5NP
L6short-a
L6short-b
L6tall-a
L6tall-b
L6tall-c
L6wm
PTC
DTC
STC
ITC

### Motif Group 19

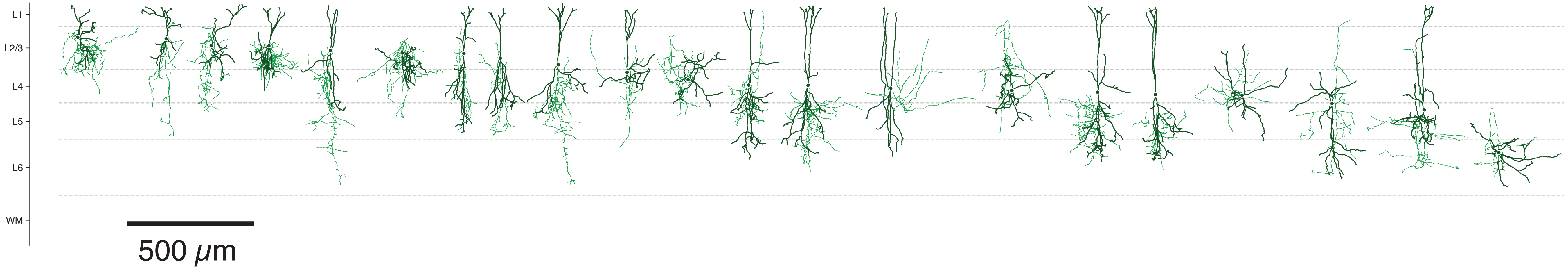

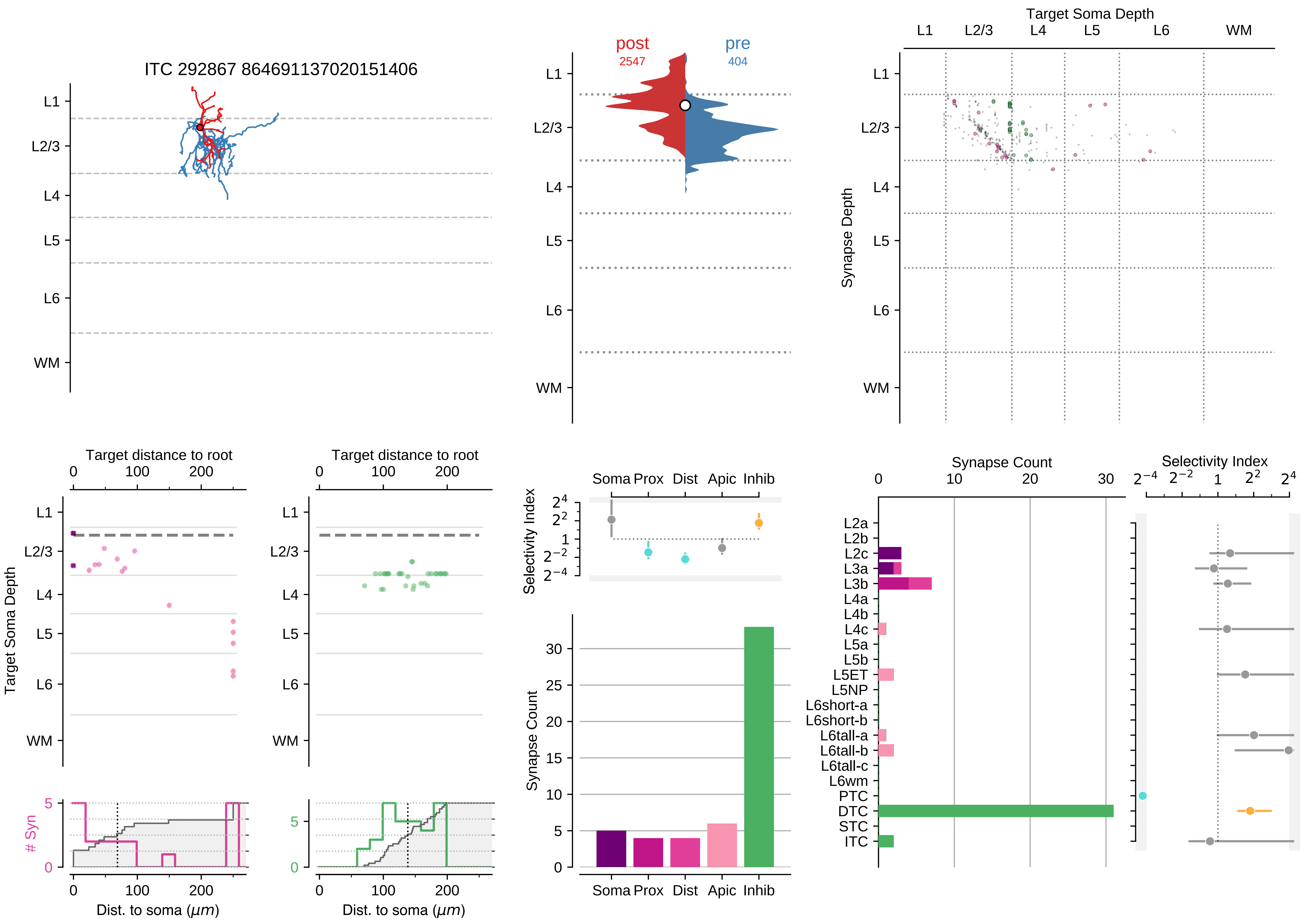

ITC 292867 864691137020151406
L1
L2/3
L4
L5
L6
WM
post
2547
pre
404
Target Soma Depth
Synapse Depth
Target distance to root
0
100
200
Target Soma Depth
### Syn
Dist. to soma ($\mu m$)
Selectivity Index
Soma Prox Dist Apic Inhib
Synapse Count
Synapse Count
Selectivity Index
L2a
L2b
L2c
L3a
L3b
L4a
L4b
L4c
L5a
L5b
L5ET
L5NP
L6short-a
L6short-b
L6tall-a
L6tall-b
L6tall-c
L6wm
PTC
DTC
STC
ITC

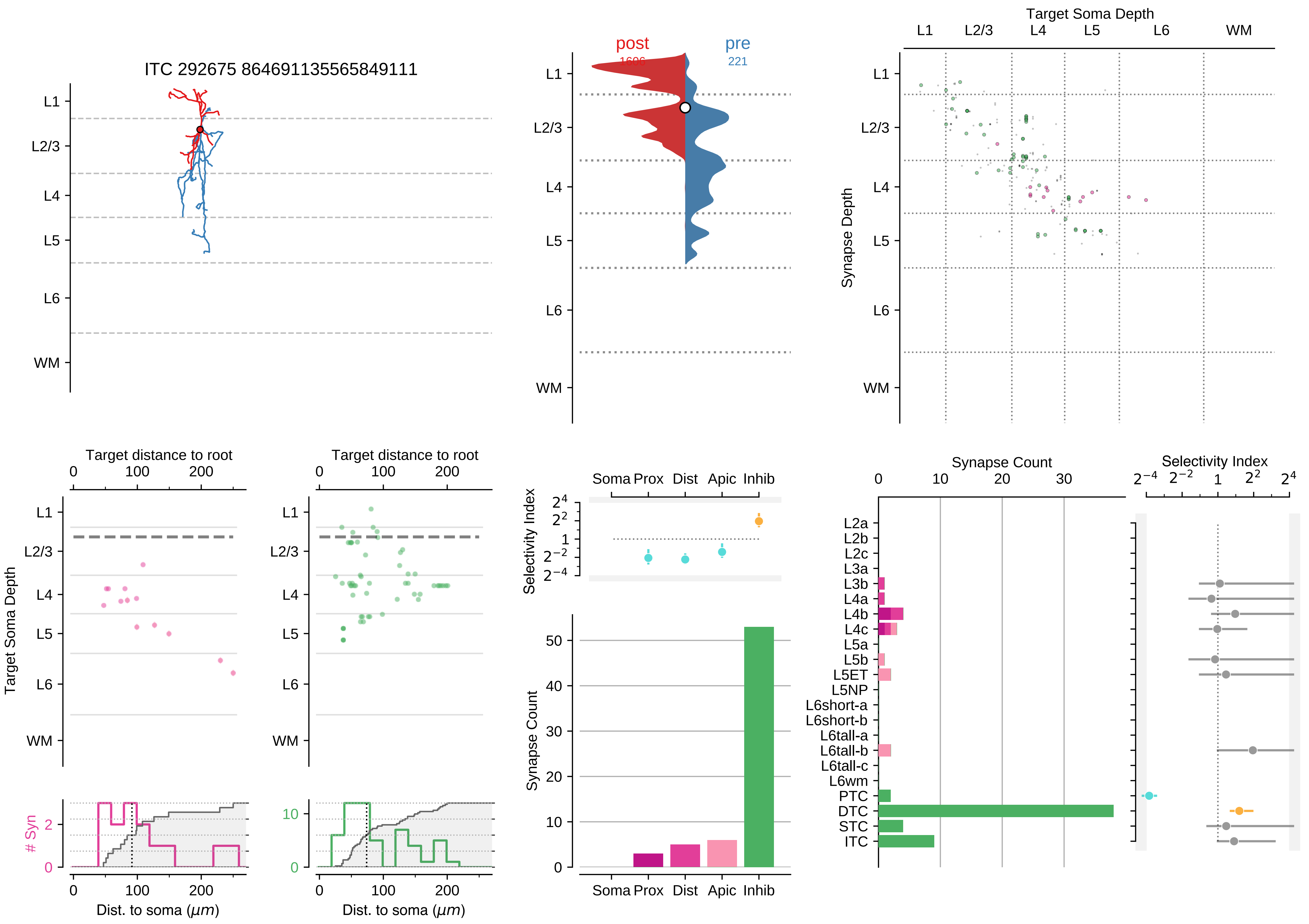

ITC 292675 864691135565849111
L1
L2/3
L4
L5
L6
WM
post
pre
221
Target Soma Depth
Synapse Depth
Target distance to root
Dist. to soma ($\mu m$)
### Syn
Selectivity Index
Soma Prox Dist Apic Inhib
Synapse Count
L2a
L2b
L2c
L3a
L3b
L4a
L4b
L4c
L5a
L5b
L5ET
L5NP
L6short-a
L6short-b
L6tall-a
L6tall-b
L6tall-c
L6wm
PTC
DTC
STC
ITC

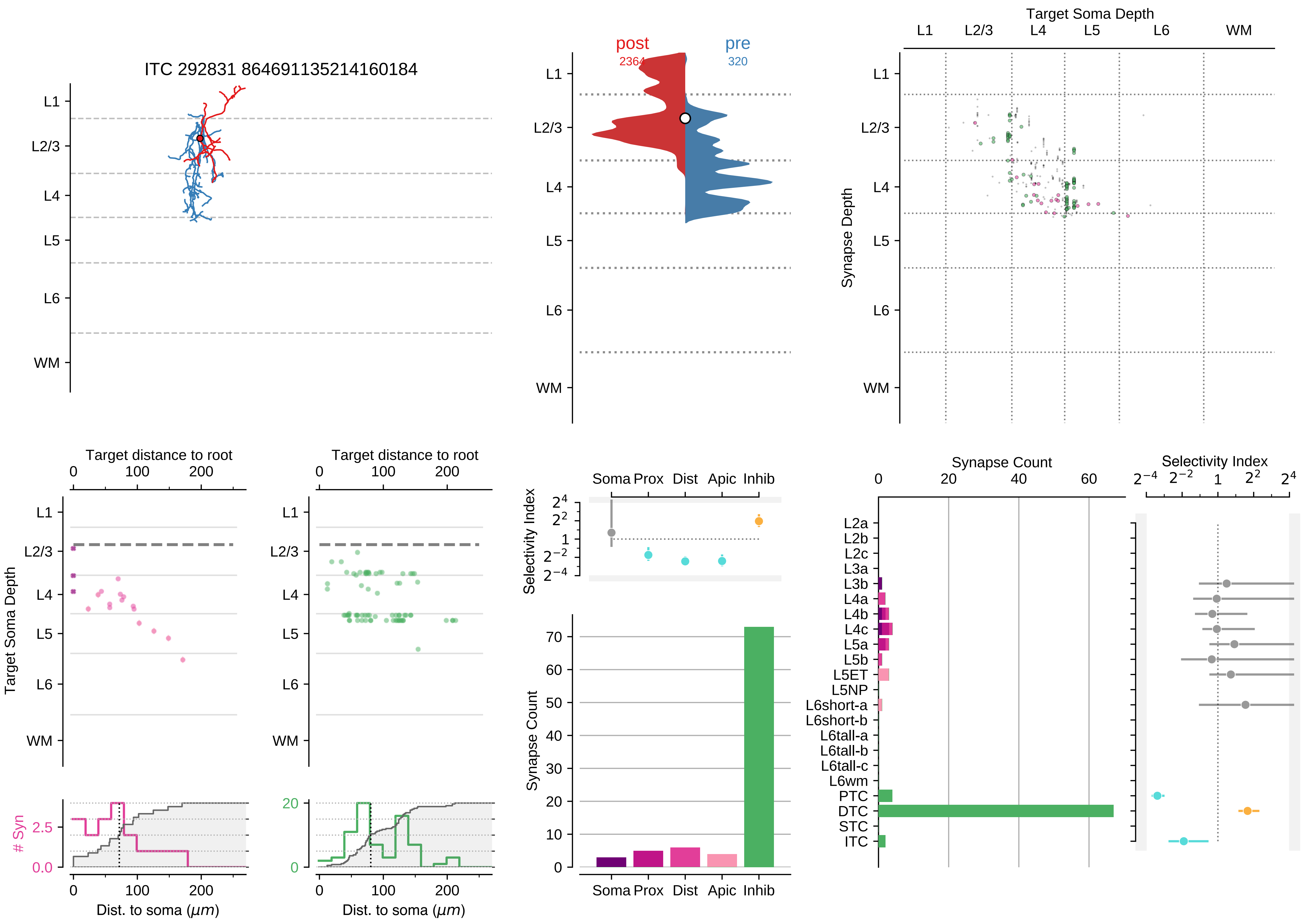

ITC 292831 864691135214160184
L1
L2/3
L4
L5
L6
WM
post
2364
pre
320
Target Soma Depth
Synapse Depth
Target distance to root
Target Soma Depth
### Syn
Dist. to soma (μm)
Selectivity Index
Soma
Prox
Dist
Apic
Inhib
Synapse Count
L2a
L2b
L2c
L3a
L3b
L4a
L4b
L4c
L5a
L5b
L5ET
L5NP
L6short-a
L6short-b
L6tall-a
L6tall-b
L6tall-c
L6wm
PTC
DTC
STC
ITC

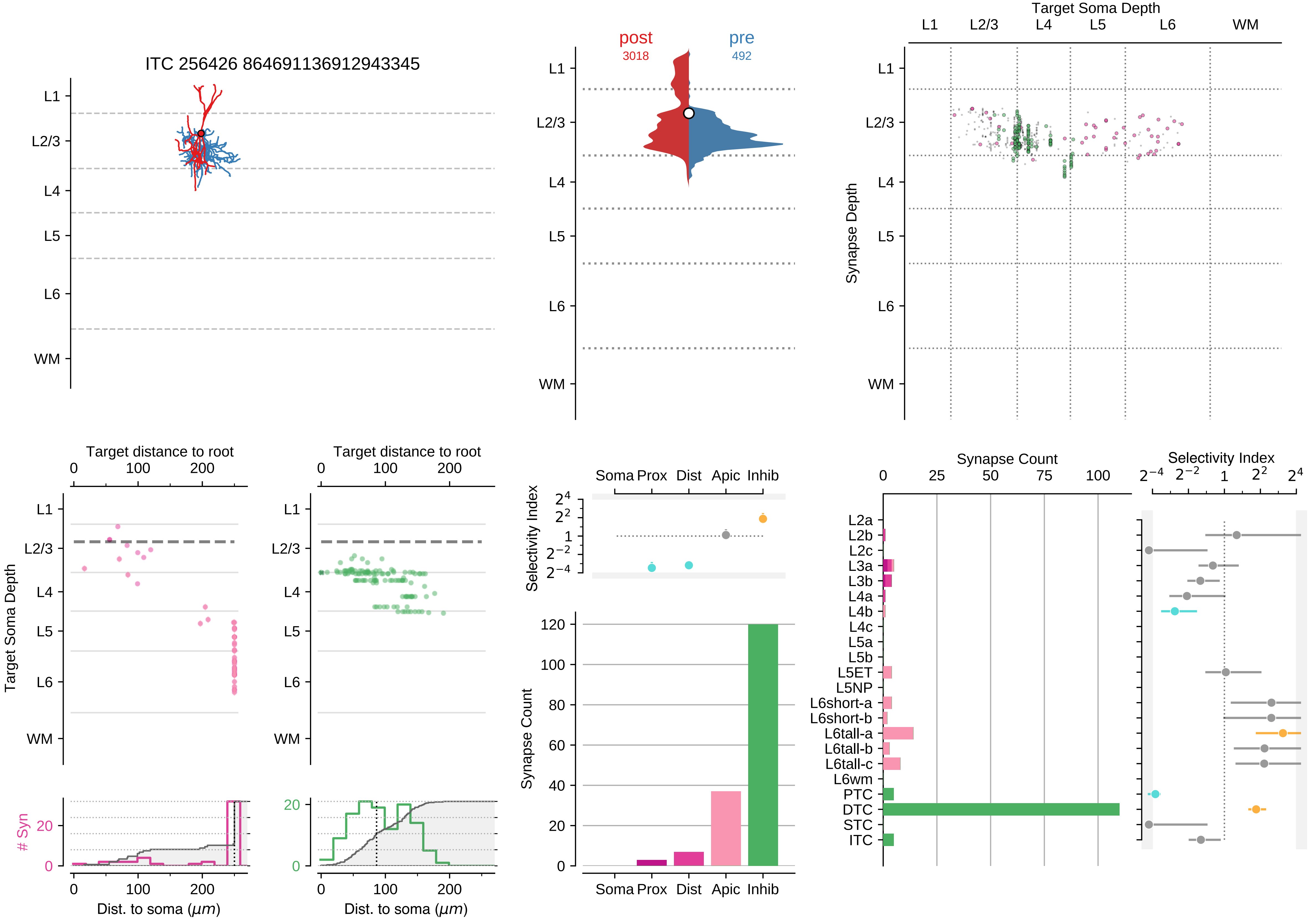

ITC 256426 864691136912943345
L1
L2/3
L4
L5
L6
WM
post
3018
pre
492
Target Soma Depth
Synapse Depth
Target distance to root
Target Soma Depth
### Syn
Dist. to soma (μm)
Selectivity Index
Soma Prox
Dist
Apic
Inhib
Synapse Count
L2a
L2b
L2c
L3a
L3b
L4a
L4b
L4c
L5a
L5b
L5ET
L5NP
L6short-a
L6short-b
L6tall-a
L6tall-b
L6tall-c
L6wm
PTC
DTC
STC
ITC

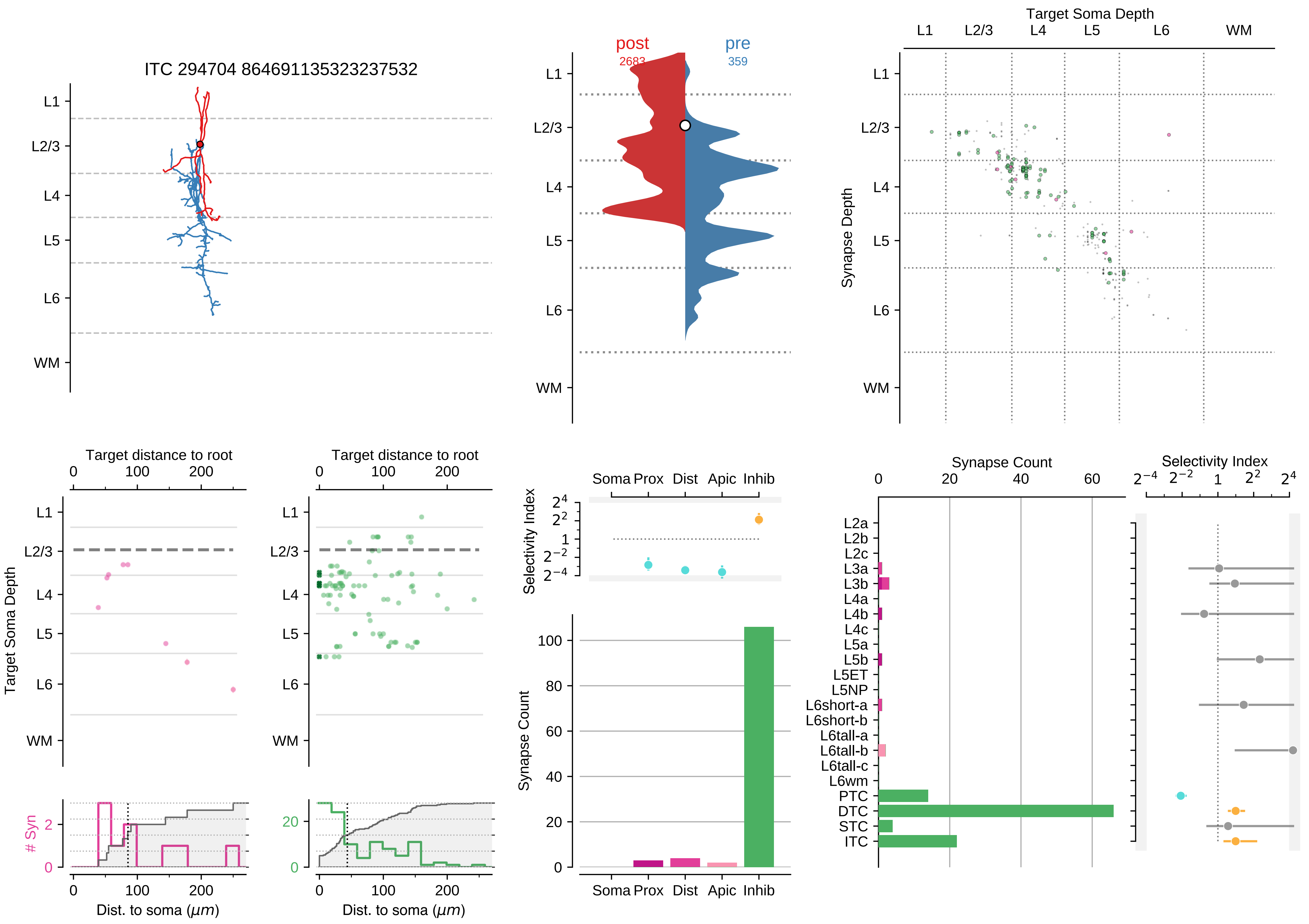

ITC 294704 864691135323237532
L1
L2/3
L4
L5
L6
WM
post
2683
pre
359
Target Soma Depth
Synapse Depth
Target distance to root
Target Soma Depth
### Syn
Dist. to soma ($\mu m$)
Selectivity Index
Soma Prox Dist Apic Inhib
Synapse Count
Synapse Count
Selectivity Index
L2a
L2b
L2c
L3a
L3b
L4a
L4b
L4c
L5a
L5b
L5ET
L5NP
L6short-a
L6short-b
L6tall-a
L6tall-b
L6tall-c
L6wm
PTC
DTC
STC
ITC

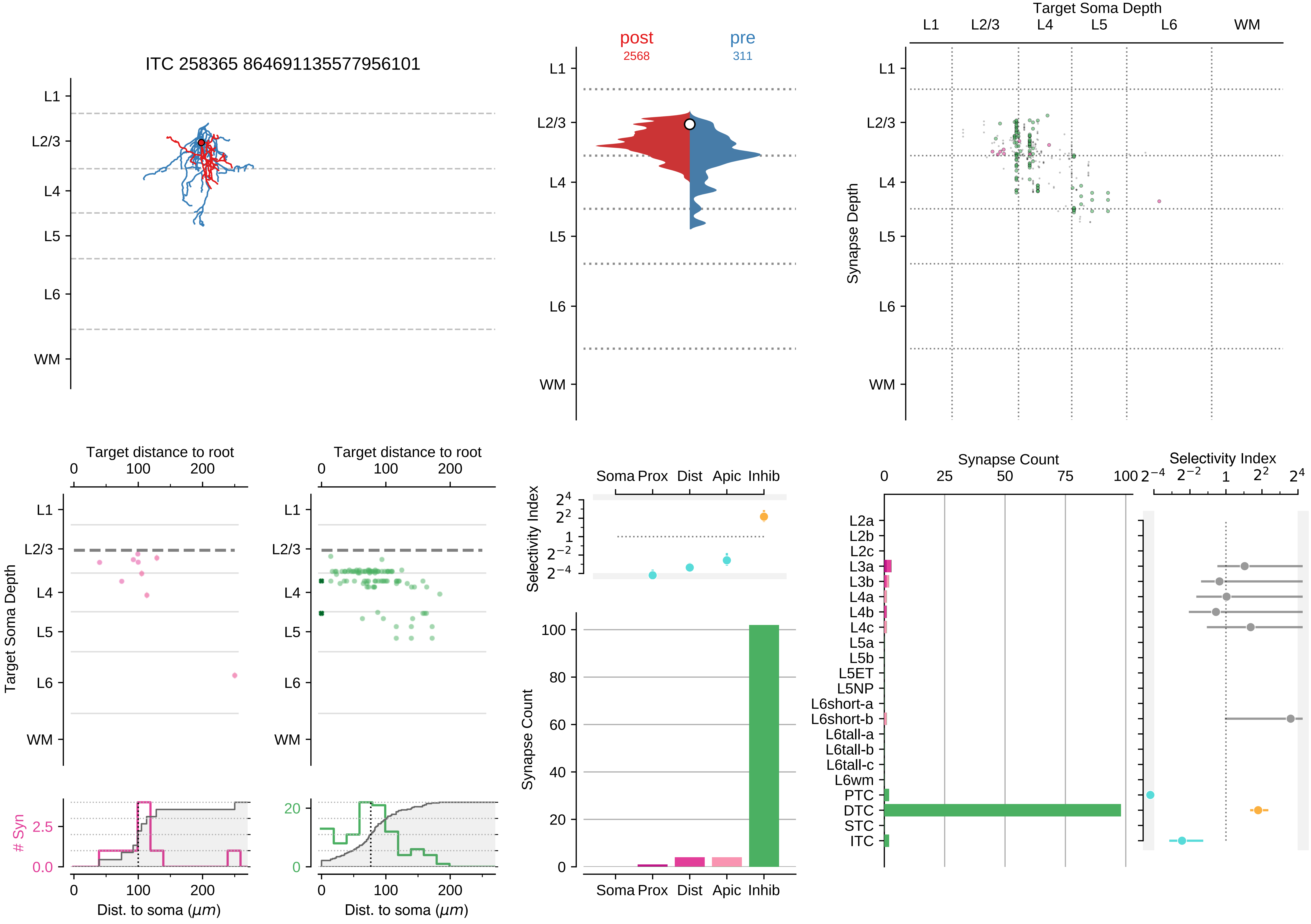

ITC 258365 864691135577956101
L1
L2/3
L4
L5
L6
WM
post
2568
pre
311
Target Soma Depth
Synapse Depth
Target distance to root
Target Soma Depth
### Syn
Dist. to soma (μm)
Soma Prox
Dist
Apic
Inhib
Selectivity Index
Synapse Count
L2a
L2b
L2c
L3a
L3b
L4a
L4b
L4c
L5a
L5b
L5ET
L5NP
L6short-a
L6short-b
L6tall-a
L6tall-b
L6tall-c
L6wm
PTC
DTC
STC
ITC

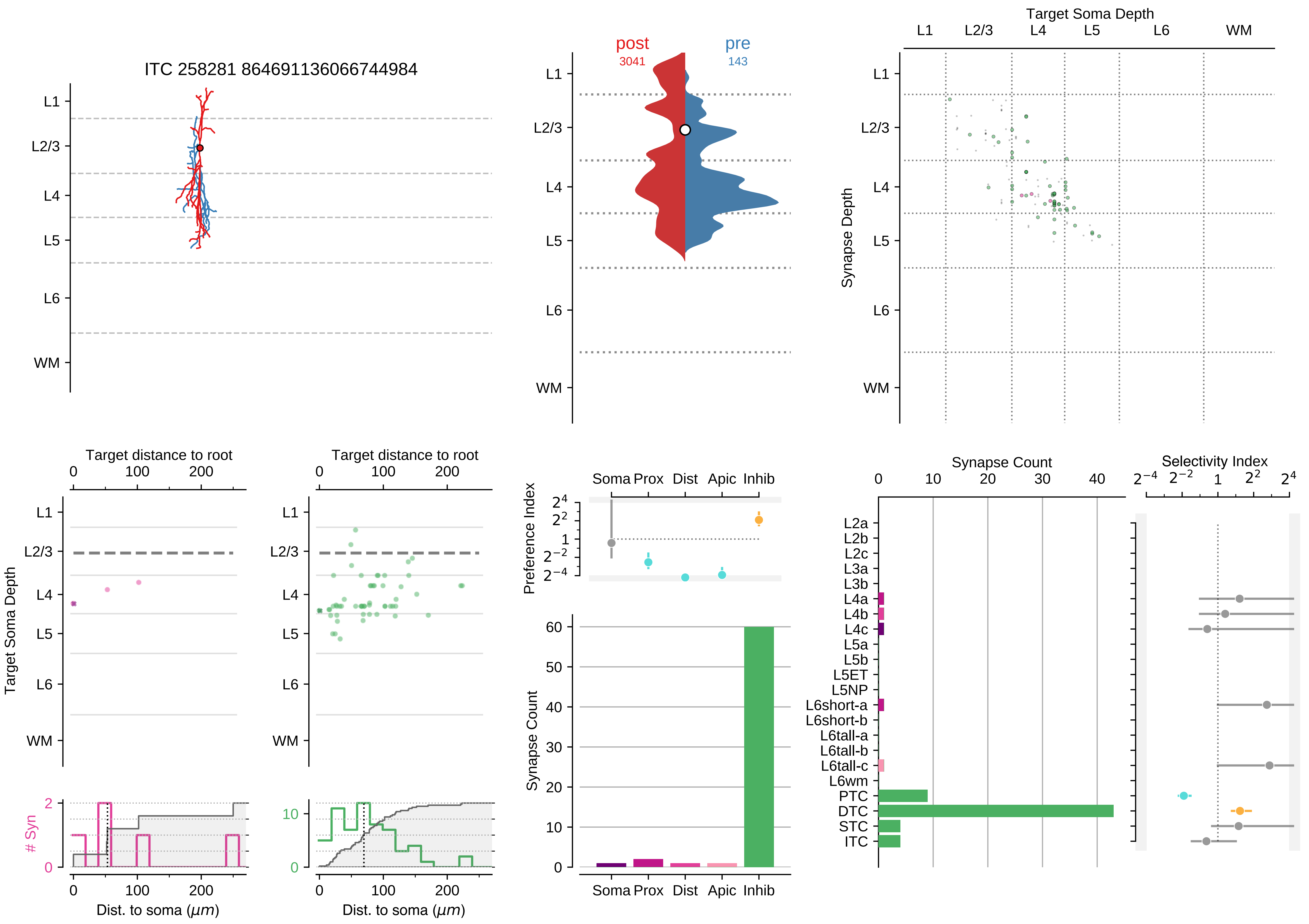

ITC 258281 864691136066744984
L1
L2/3
L4
L5
L6
WM
post
3041
pre
143
Target Soma Depth
Synapse Depth
Target distance to root
Target Soma Depth
### Syn
Dist. to soma ($\mu m$)
Preference Index
Soma Prox Dist Apic Inhib
Synapse Count
Selectivity Index
L2a
L2b
L2c
L3a
L3b
L4a
L4b
L4c
L5a
L5b
L5ET
L5NP
L6short-a
L6short-b
L6tall-a
L6tall-b
L6tall-c
L6wm
PTC
DTC
STC
ITC

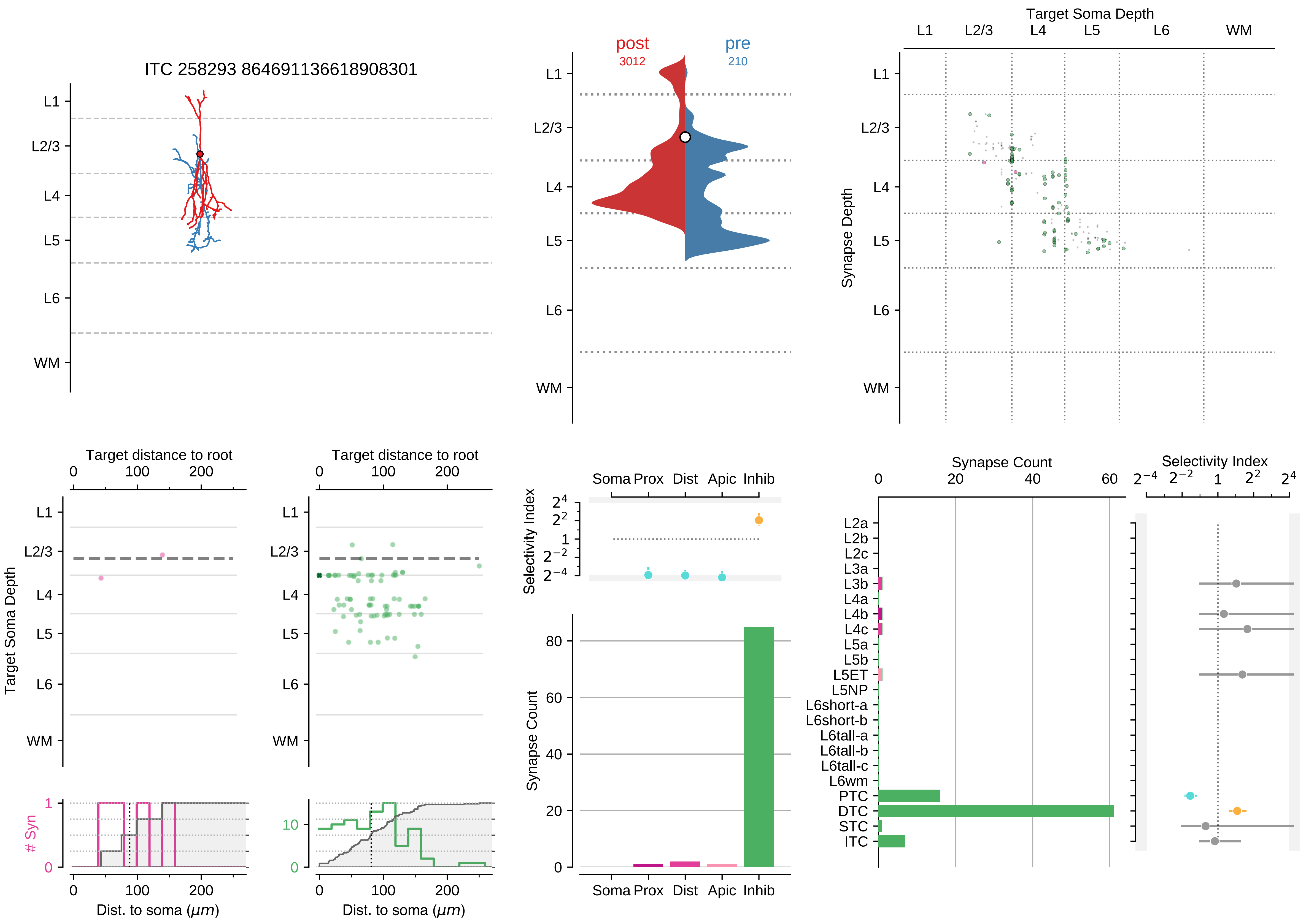

ITC 258293 864691136618908301
L1
L2/3
L4
L5
L6
WM
post
3012
pre
210
Target Soma Depth
Synapse Depth
Target distance to root
Target Soma Depth
### Syn
Dist. to soma ($\mu m$)
Soma Prox
Dist
Apic
Inhib
Selectivity Index
Synapse Count
L2a
L2b
L2c
L3a
L3b
L4a
L4b
L4c
L5a
L5b
L5ET
L5NP
L6short-a
L6short-b
L6tall-a
L6tall-b
L6tall-c
L6wm
PTC
DTC
STC
ITC

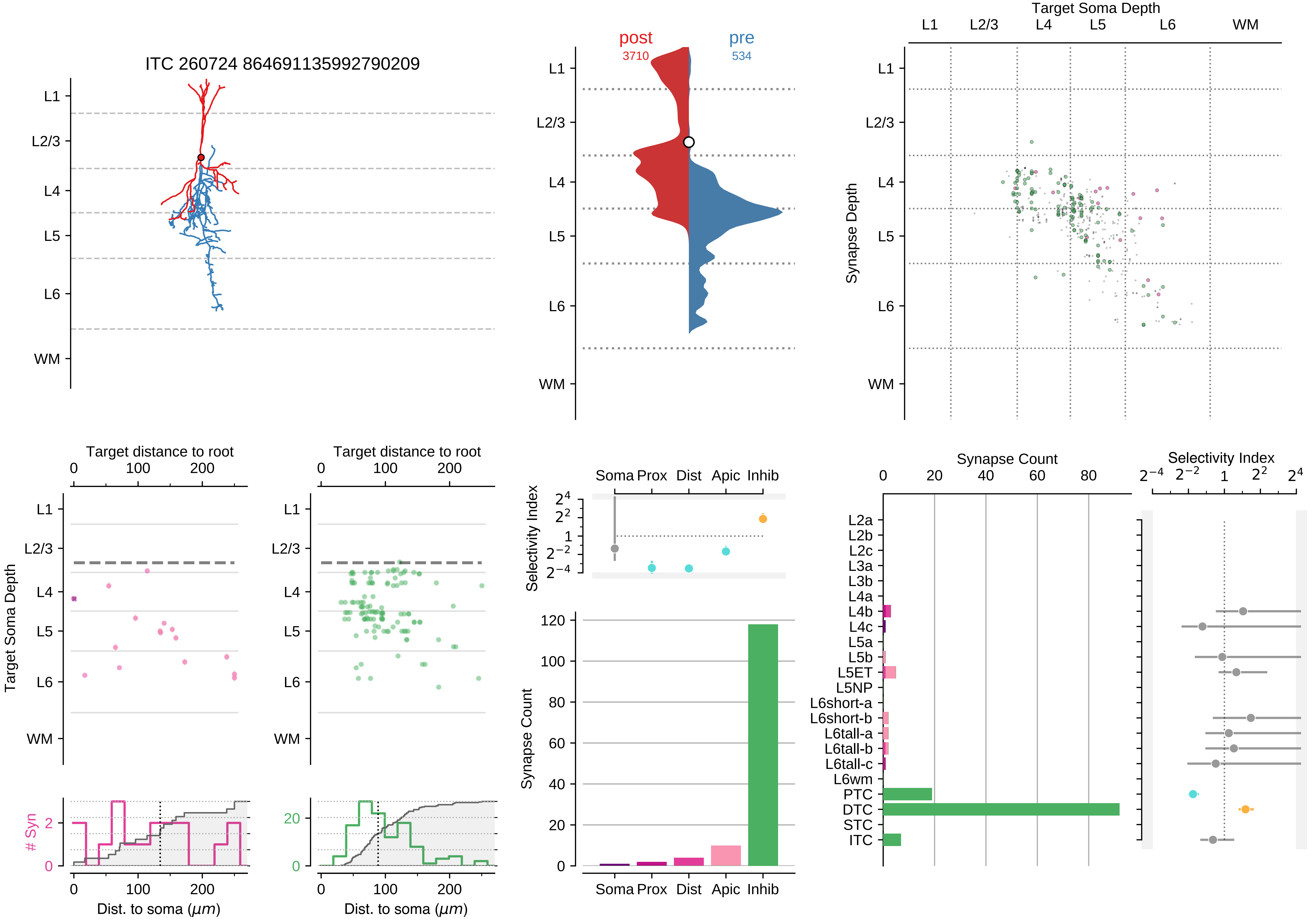
ITC 260724 864691135992790209
L1
L2/3
L4
L5
L6
WM
post
3710
pre
534
Target Soma Depth
Synapse Depth
Target distance to root
Target Soma Depth
### Syn
Dist. to soma ($\mu m$)
Selectivity Index
Soma Prox Dist Apic Inhib
Synapse Count
Synapse Count
Selectivity Index
L2a
L2b
L2c
L3a
L3b
L4a
L4b
L4c
L5a
L5b
L5ET
L5NP
L6short-a
L6short-b
L6tall-a
L6tall-b
L6tall-c
L6wm
PTC
DTC
STC
ITC

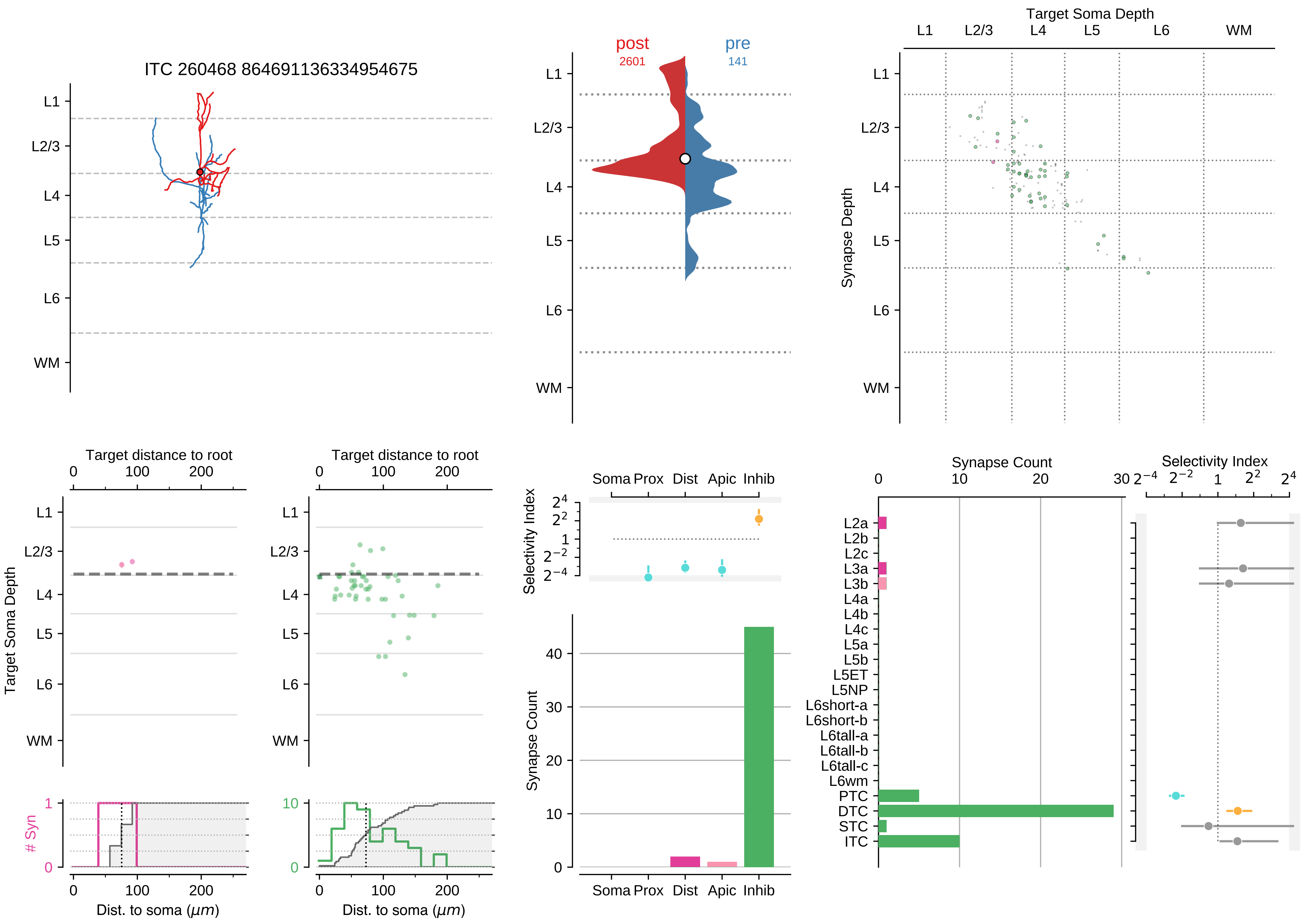

ITC 260468 864691136334954675
post
2601
pre
141
Target Soma Depth
Synapse Depth
L1
L2/3
L4
L5
L6
WM
Target distance to root
Target Soma Depth
### Syn
Dist. to soma ($\mu m$)
Selectivity Index
Soma Prox Dist Apic Inhib
Synapse Count
L2a
L2b
L2c
L3a
L3b
L4a
L4b
L4c
L5a
L5b
L5ET
L5NP
L6short-a
L6short-b
L6tall-a
L6tall-b
L6tall-c
L6wm
PTC
DTC
STC
ITC

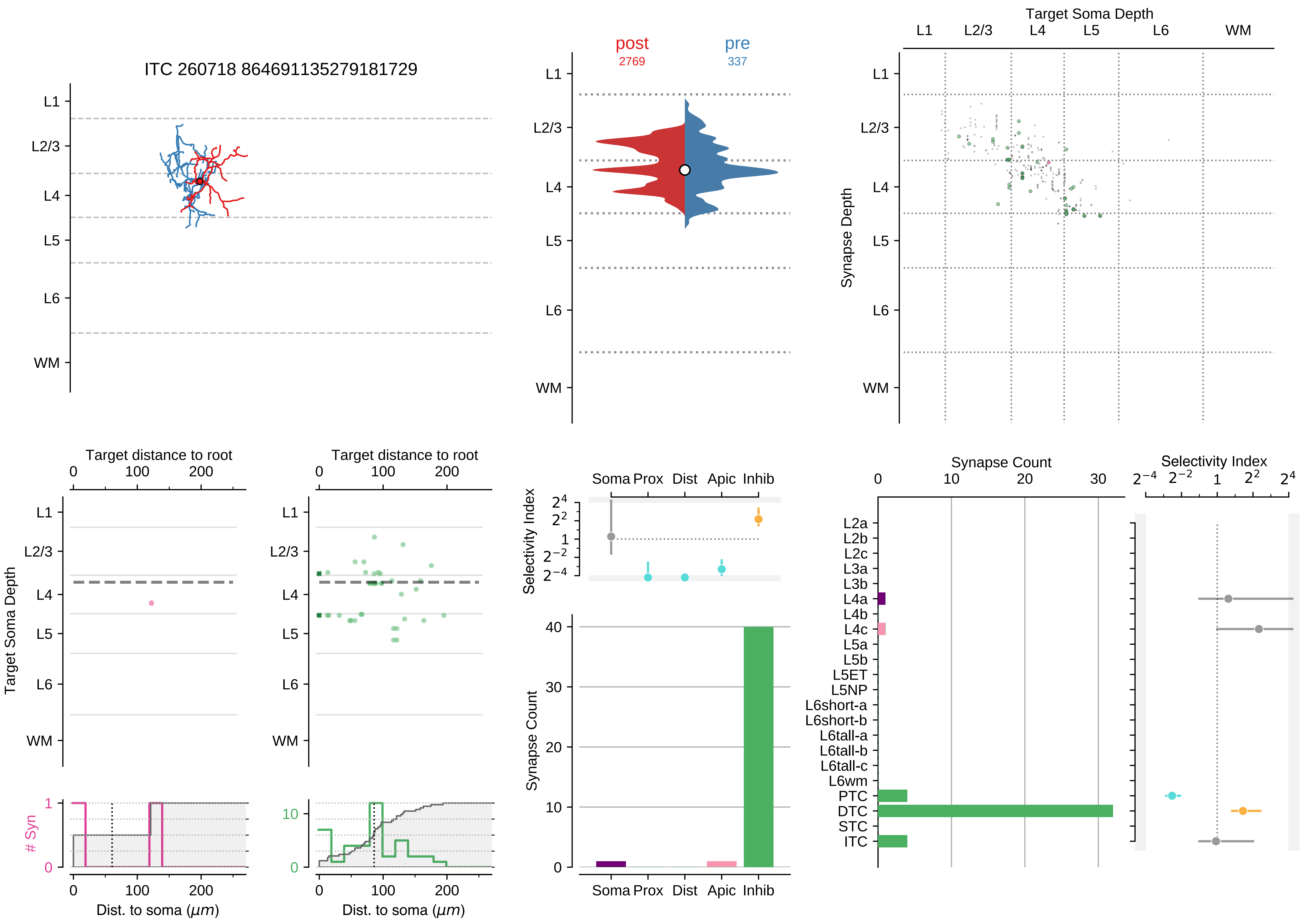

ITC 260718 864691135279181729
L1
L2/3
L4
L5
L6
WM
post
2769
pre
337
Target Soma Depth
Synapse Depth
Target distance to root
Target Soma Depth
### Syn
Dist. to soma (μm)
Soma
Prox
Dist
Apic
Inhib
Selectivity Index
Synapse Count
L2a
L2b
L2c
L3a
L3b
L4a
L4b
L4c
L5a
L5b
L5ET
L5NP
L6short-a
L6short-b
L6tall-a
L6tall-b
L6tall-c
L6wm
PTC
DTC
STC
ITC

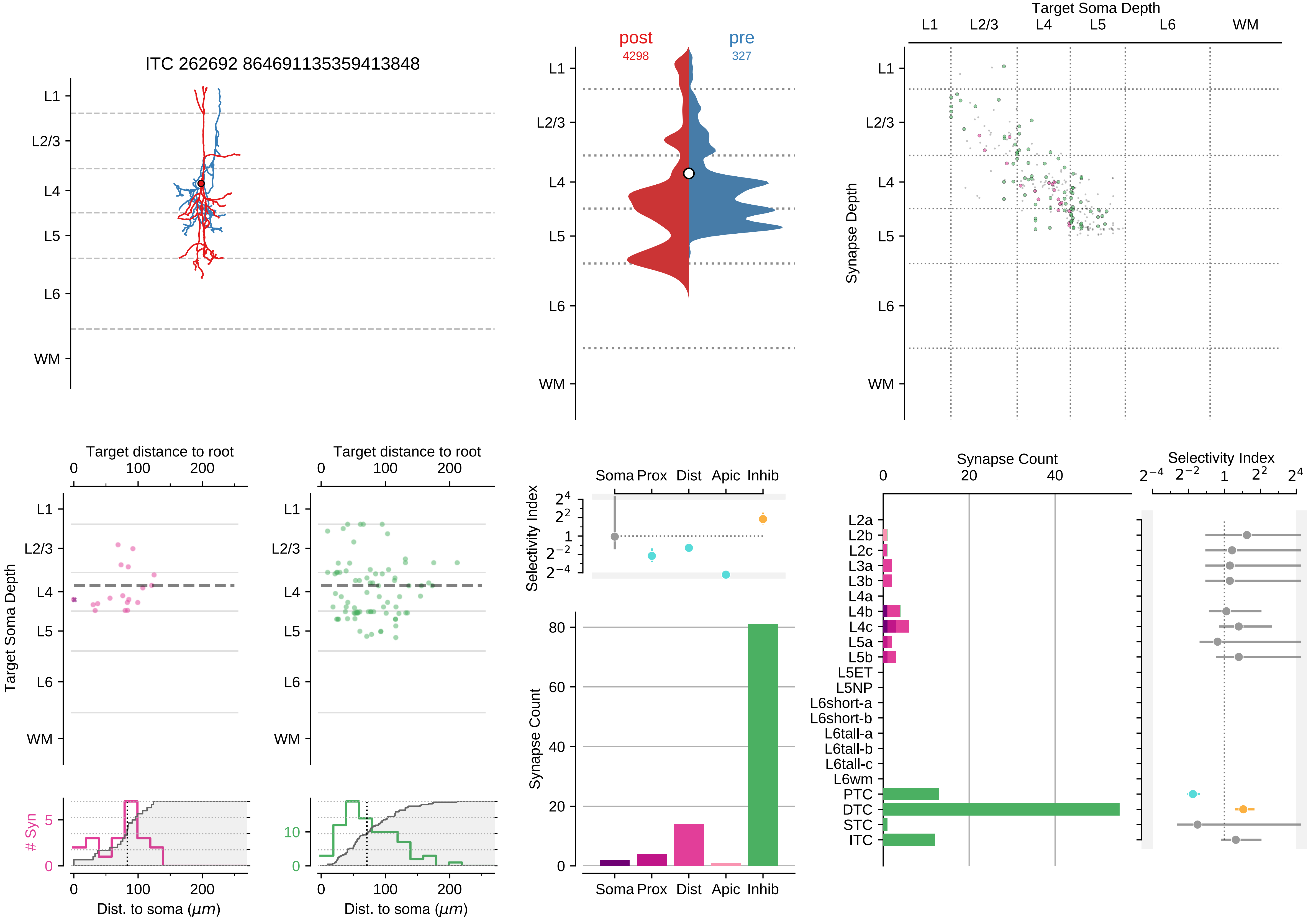

ITC 262692 864691135359413848
L1
L2/3
L4
L5
L6
WM
post
4298
pre
327
Target Soma Depth
Synapse Depth
Target distance to root
0
100
200
Target Soma Depth
### Syn
Dist. to soma ($\mu m$)
Selectivity Index
Soma Prox Dist Apic Inhib
Synapse Count
Synapse Count
Selectivity Index
L2a
L2b
L2c
L3a
L3b
L4a
L4b
L4c
L5a
L5b
L5ET
L5NP
L6short-a
L6short-b
L6tall-a
L6tall-b
L6tall-c
L6wm
PTC
DTC
STC
ITC

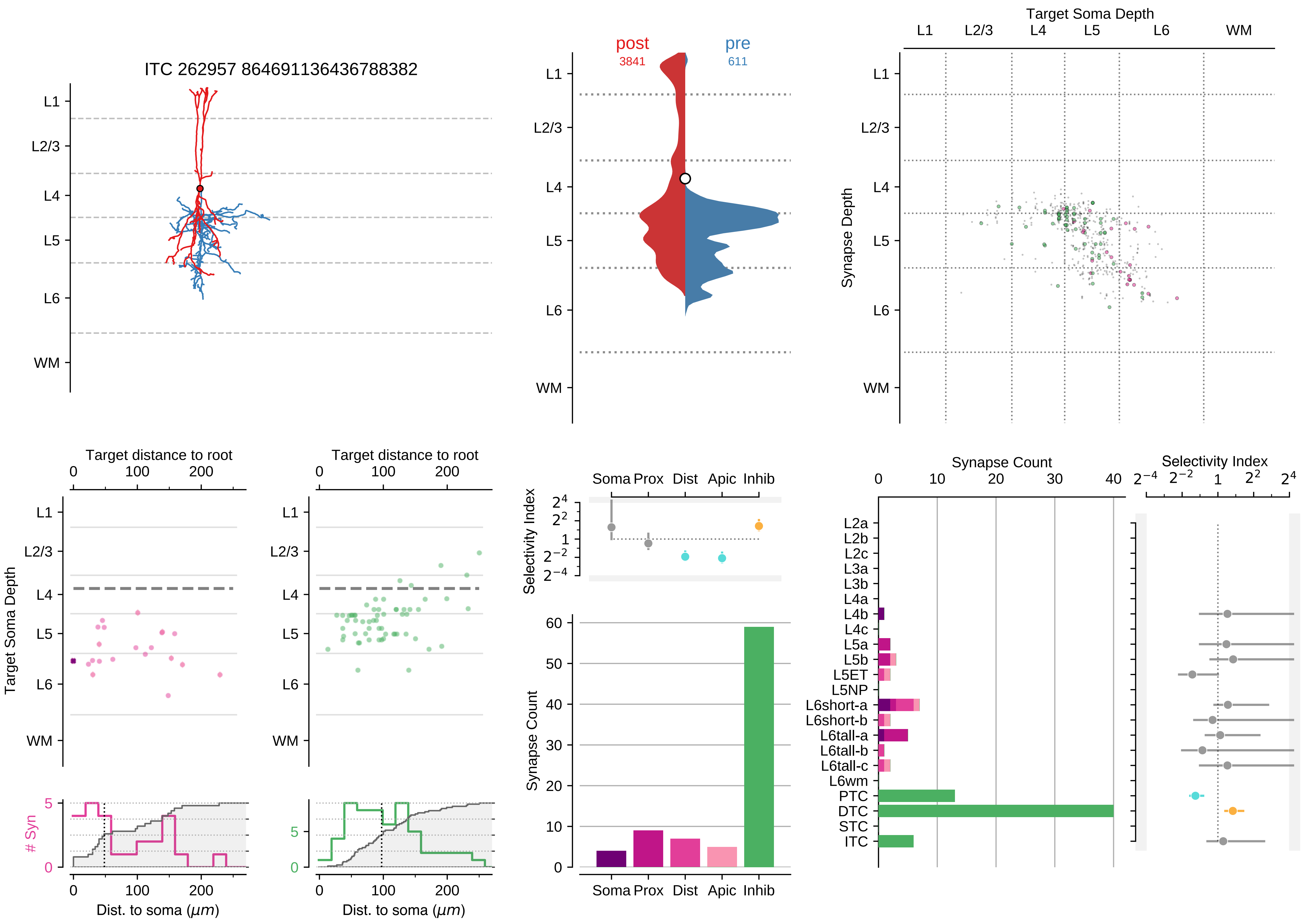

ITC 262957 864691136436788382
post
3841
pre
611
Target Soma Depth
Synapse Depth
L1
L2/3
L4
L5
L6
WM
Target distance to root
Target Soma Depth
### Syn
Dist. to soma ($\mu m$)
Selectivity Index
Soma Prox Dist Apic Inhib
Synapse Count
L2a
L2b
L2c
L3a
L3b
L4a
L4b
L4c
L5a
L5b
L5ET
L5NP
L6short-a
L6short-b
L6tall-a
L6tall-b
L6tall-c
L6wm
PTC
DTC
STC
ITC

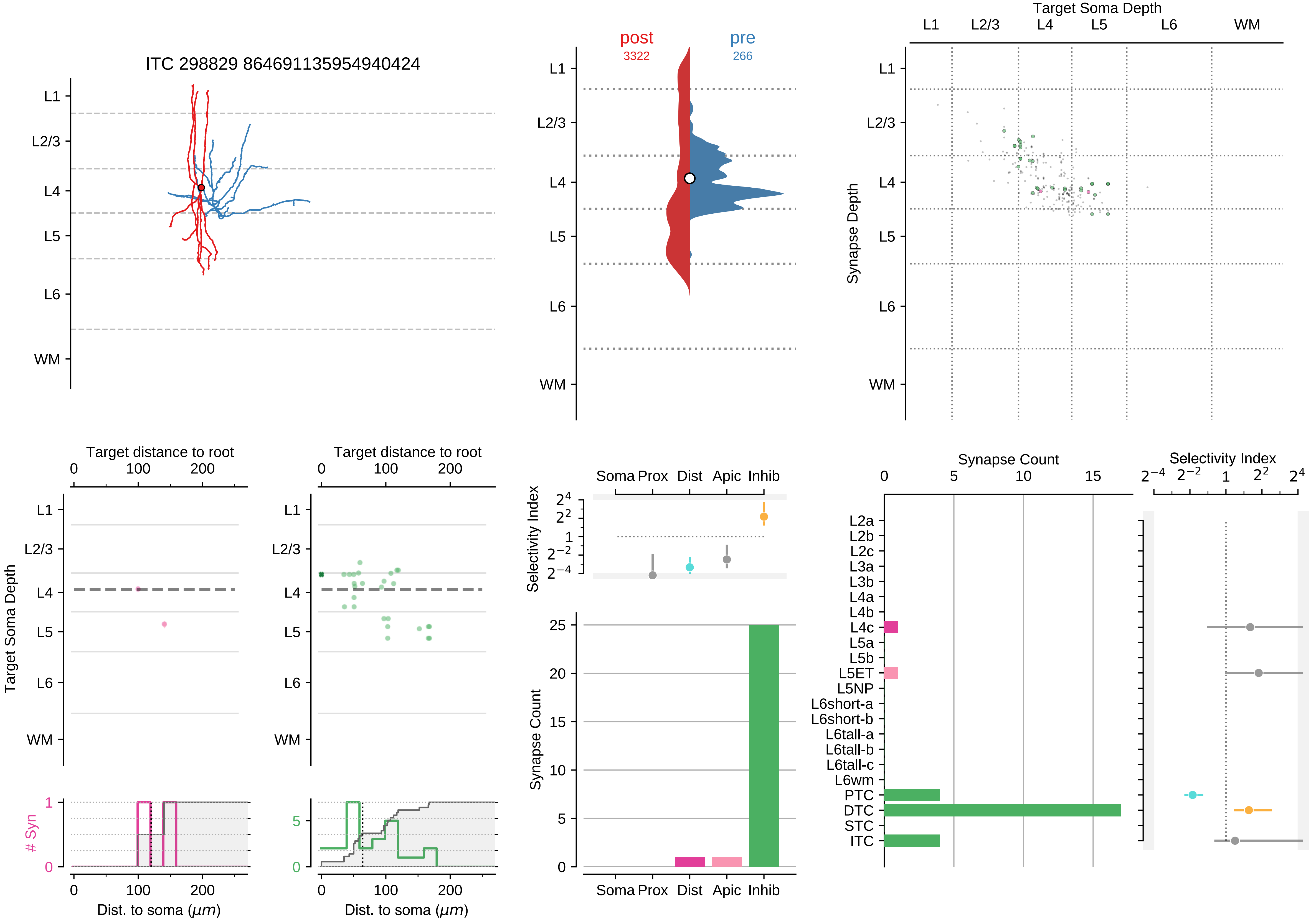

ITC 298829 864691135954940424
L1
L2/3
L4
L5
L6
WM
post
3322
pre
266
Target Soma Depth
Synapse Depth
Target distance to root
0
100
200
Target Soma Depth
### Syn
Dist. to soma (μm)
Soma Prox
Dist
Apic
Inhib
Selectivity Index
Synapse Count
L2a
L2b
L2c
L3a
L3b
L4a
L4b
L4c
L5a
L5b
L5ET
L5NP
L6short-a
L6short-b
L6tall-a
L6tall-b
L6tall-c
L6wm
PTC
DTC
STC
ITC

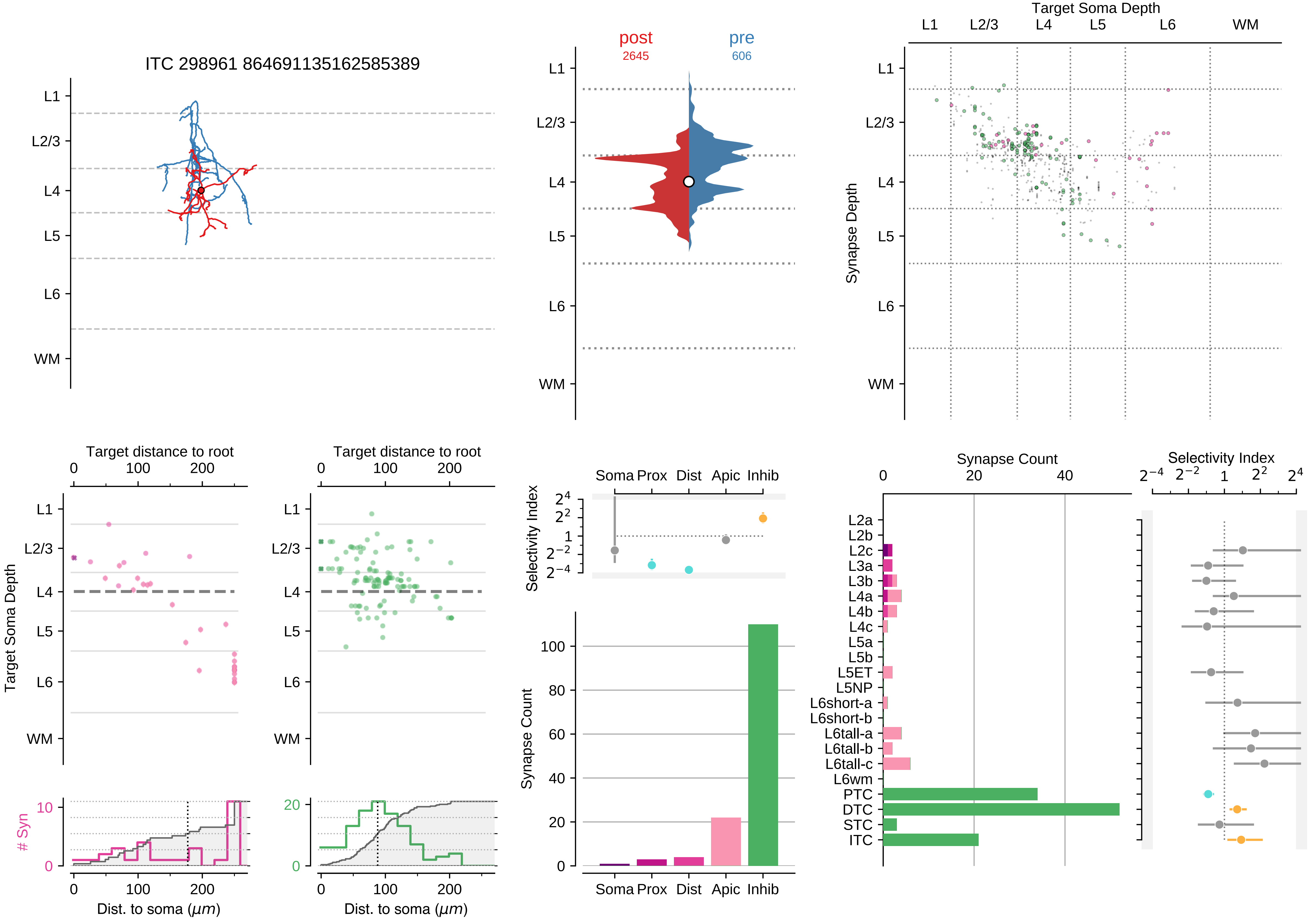

ITC 298961 864691135162585389
L1
L2/3
L4
L5
L6
WM
post
2645
pre
606
Target Soma Depth
Synapse Depth
Target distance to root
Target Soma Depth
Dist. to soma (μm)
### Syn
Selectivity Index
Soma Prox Dist Apic Inhib
Synapse Count
L2a
L2b
L2c
L3a
L3b
L4a
L4b
L4c
L5a
L5b
L5ET
L5NP
L6short-a
L6short-b
L6tall-a
L6tall-b
L6tall-c
L6wm
PTC
DTC
STC
ITC

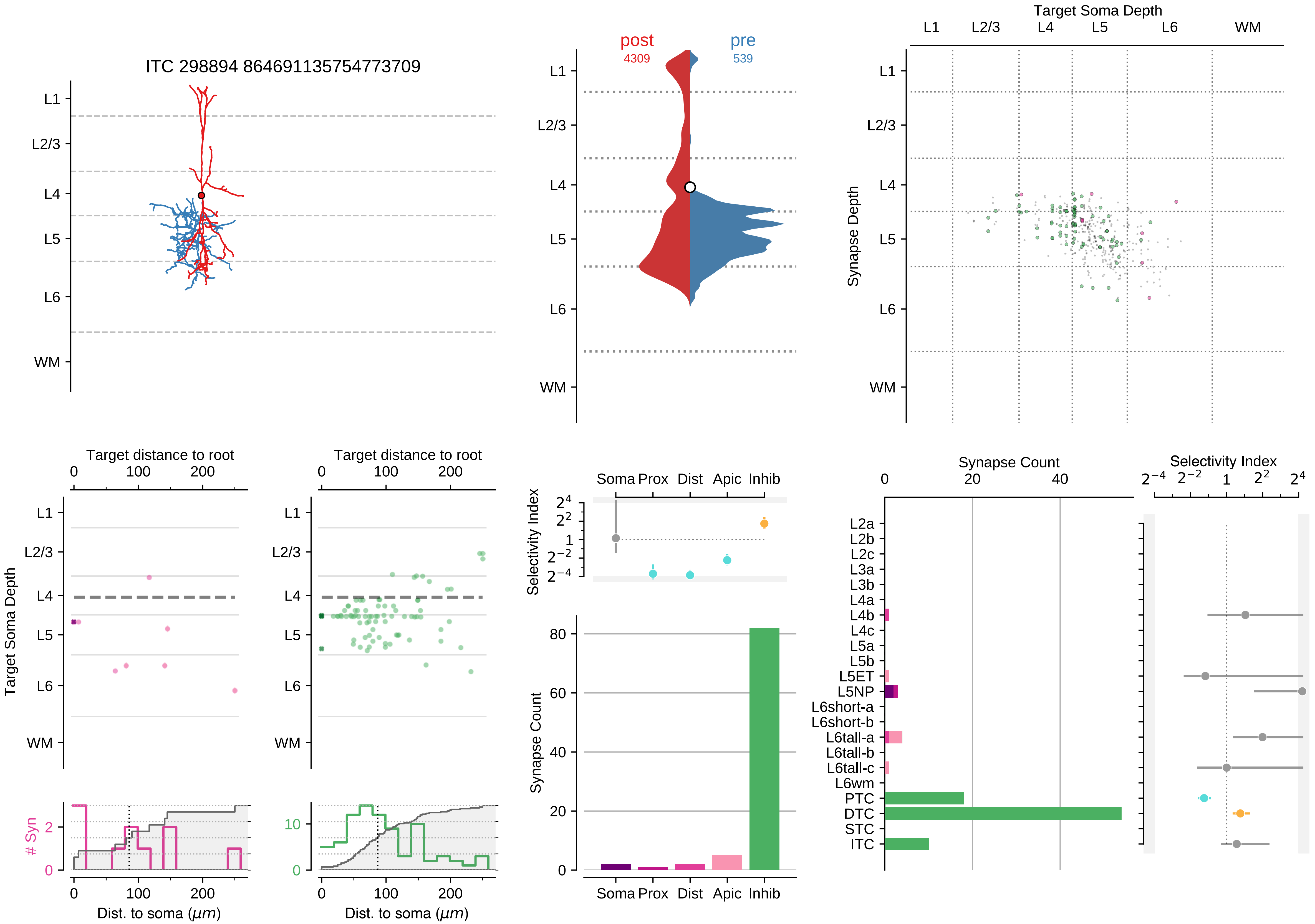
ITC 298894 864691135754773709
L1
L2/3
L4
L5
L6
WM
post
4309
pre
539
Target Soma Depth
Synapse Depth
Target distance to root
Target Soma Depth
### Syn
Dist. to soma (μm)
Selectivity Index
Soma
Prox
Dist
Apic
Inhib
Synapse Count
L2a
L2b
L2c
L3a
L3b
L4a
L4b
L4c
L5a
L5b
L5ET
L5NP
L6short-a
L6short-b
L6tall-a
L6tall-b
L6tall-c
L6wm
PTC
DTC
STC
ITC

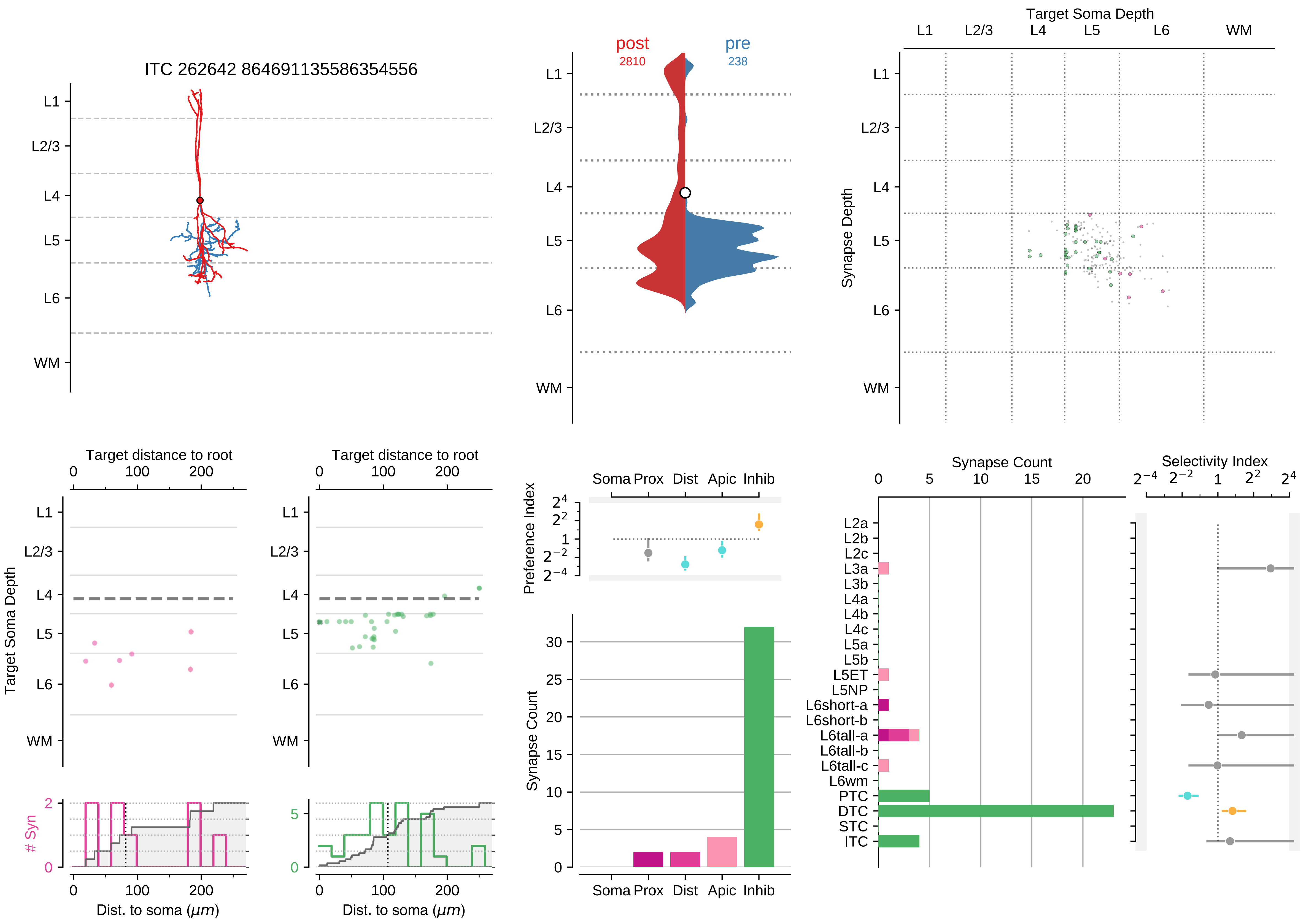

ITC 262642 864691135586354556
L1
L2/3
L4
L5
L6
WM
post
2810
pre
238
Target Soma Depth
Synapse Depth
Target distance to root
Target Soma Depth
### Syn
Dist. to soma (μm)
Preference Index
Soma Prox Dist Apic Inhib
Synapse Count
Selectivity Index
L2a
L2b
L2c
L3a
L3b
L4a
L4b
L4c
L5a
L5b
L5ET
L5NP
L6short-a
L6short-b
L6tall-a
L6tall-b
L6tall-c
L6wm
PTC
DTC
STC
ITC

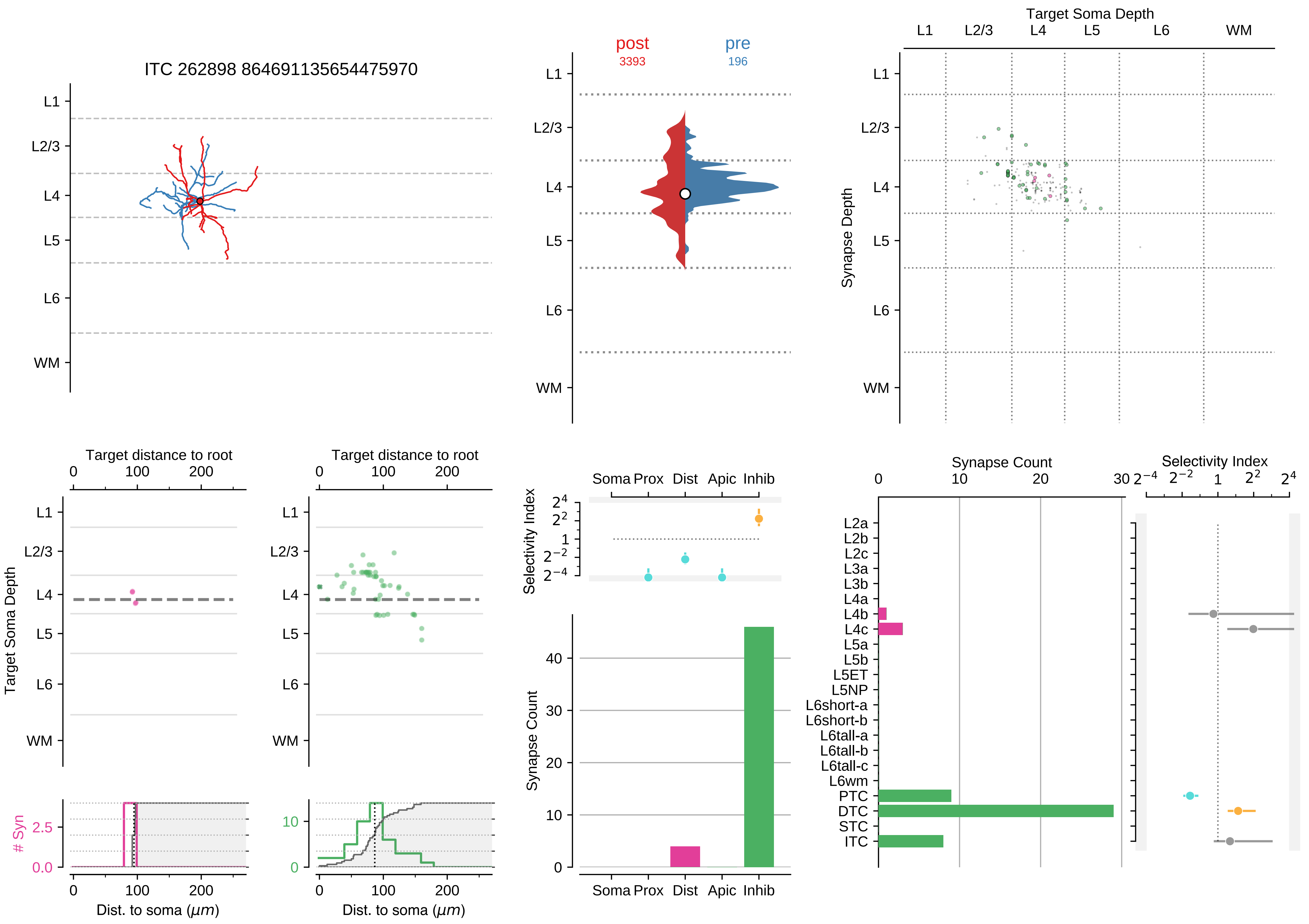

ITC 262898 864691135654475970
L1
L2/3
L4
L5
L6
WM
post
3393
pre
196
Target Soma Depth
Synapse Depth
Target distance to root
0
100
200
Target Soma Depth
### Syn
2.5
0.0
10
Dist. to soma ($\mu m$)
Selectivity Index
Soma Prox
Dist
Apic
Inhib
Synapse Count
Synapse Count
Selectivity Index
L2a
L2b
L2c
L3a
L3b
L4a
L4b
L4c
L5a
L5b
L5ET
L5NP
L6short-a
L6short-b
L6tall-a
L6tall-b
L6tall-c
L6wm
PTC
DTC
STC
ITC

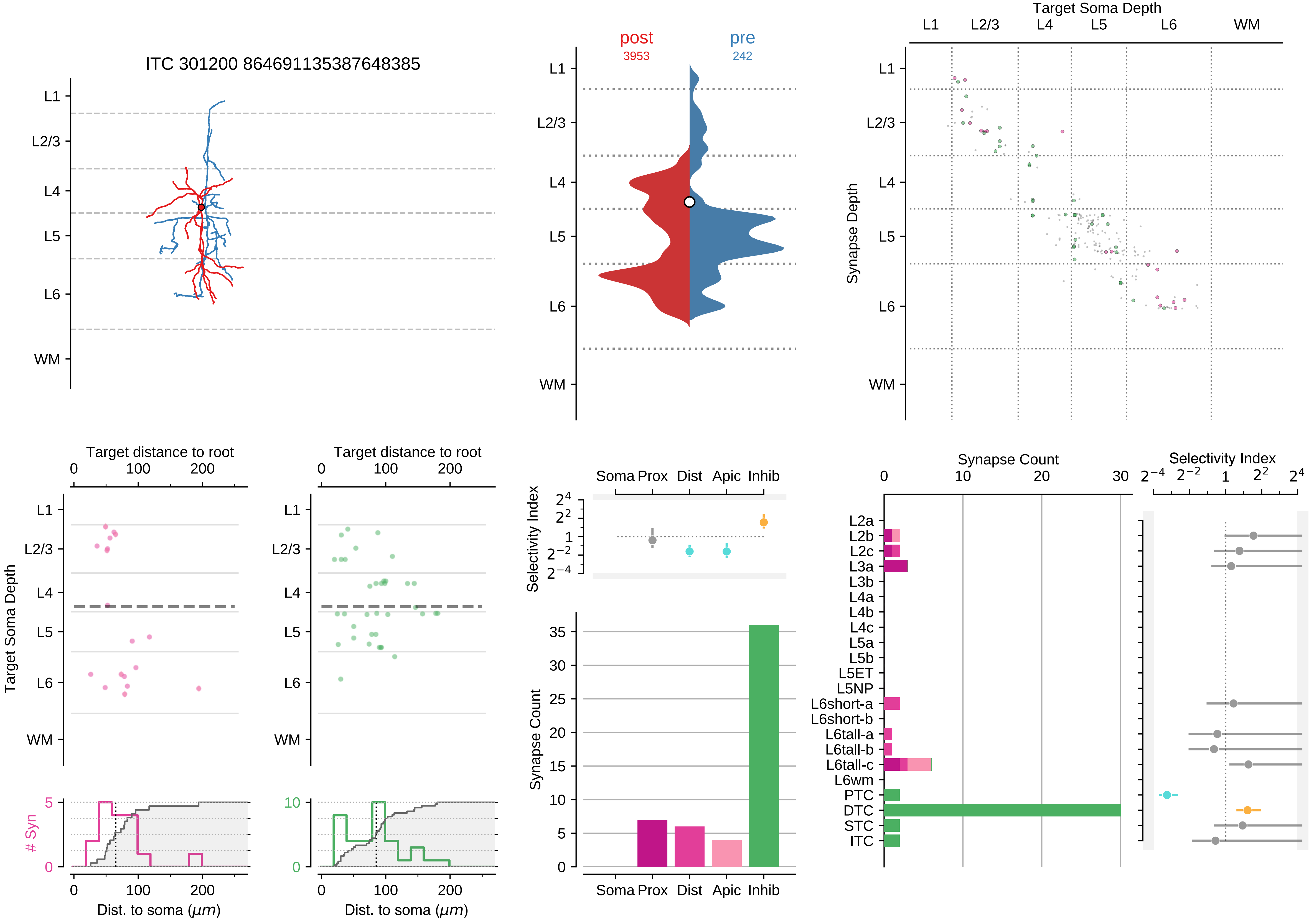

ITC 301200 864691135387648385
L1
L2/3
L4
L5
L6
WM
post
3953
pre
242
Target Soma Depth
Synapse Depth
Target distance to root
Target Soma Depth
### Syn
Dist. to soma ($\mu m$)
Soma Prox Dist Apic Inhib
Selectivity Index
Synapse Count
L2a
L2b
L2c
L3a
L3b
L4a
L4b
L4c
L5a
L5b
L5ET
L5NP
L6short-a
L6short-b
L6tall-a
L6tall-b
L6tall-c
L6wm
PTC
DTC
STC
ITC

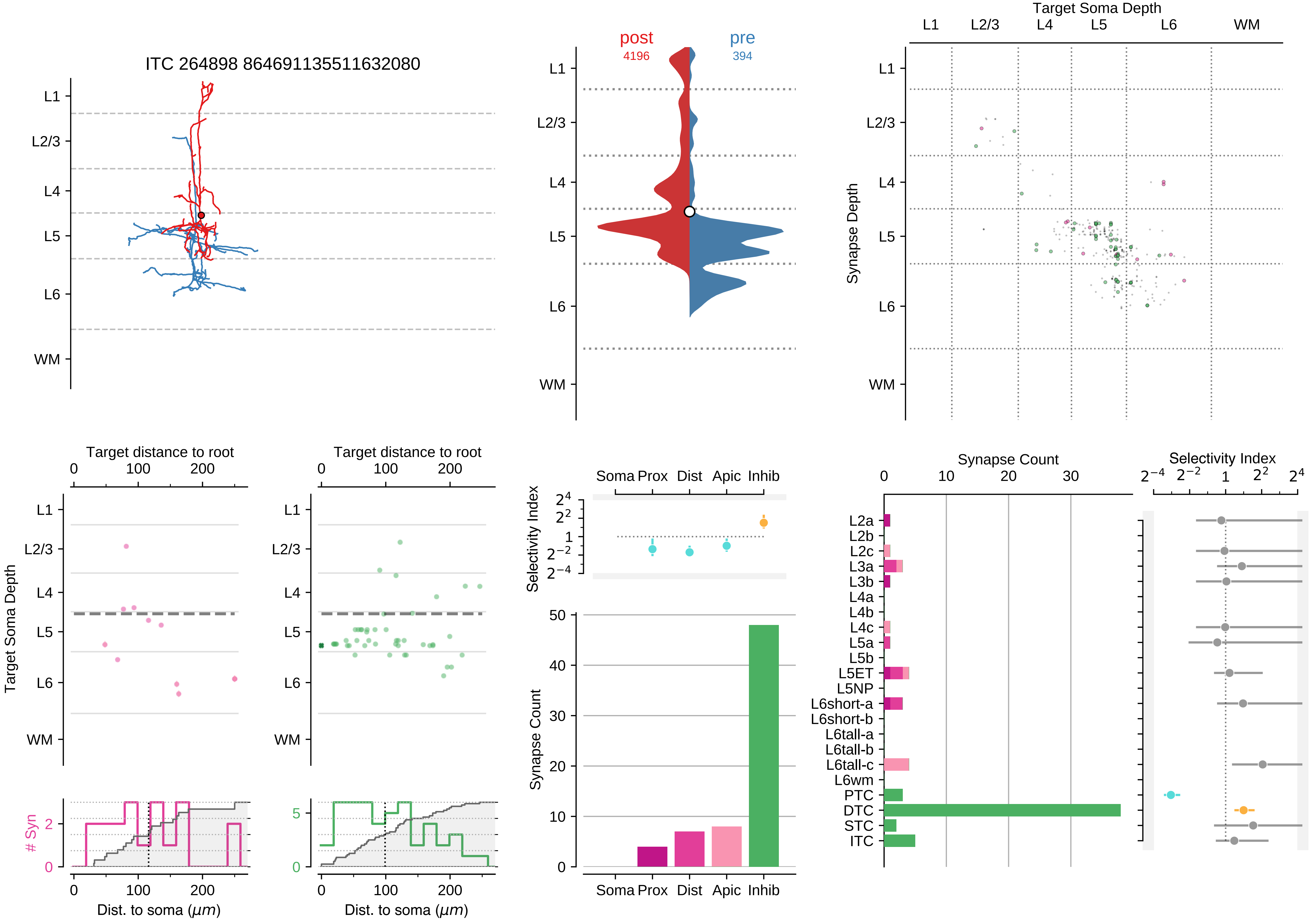

ITC 264898 864691135511632080
L1
L2/3
L4
L5
L6
WM
post
4196
pre
394
Target Soma Depth
Synapse Depth
Target distance to root
Target Soma Depth
### Syn
Dist. to soma ($\mu m$)
Soma
Prox
Dist
Apic
Inhib
Selectivity Index
Synapse Count
L2a
L2b
L2c
L3a
L3b
L4a
L4b
L4c
L5a
L5b
L5ET
L5NP
L6short-a
L6short-b
L6tall-a
L6tall-b
L6tall-c
L6wm
PTC
DTC
STC
ITC

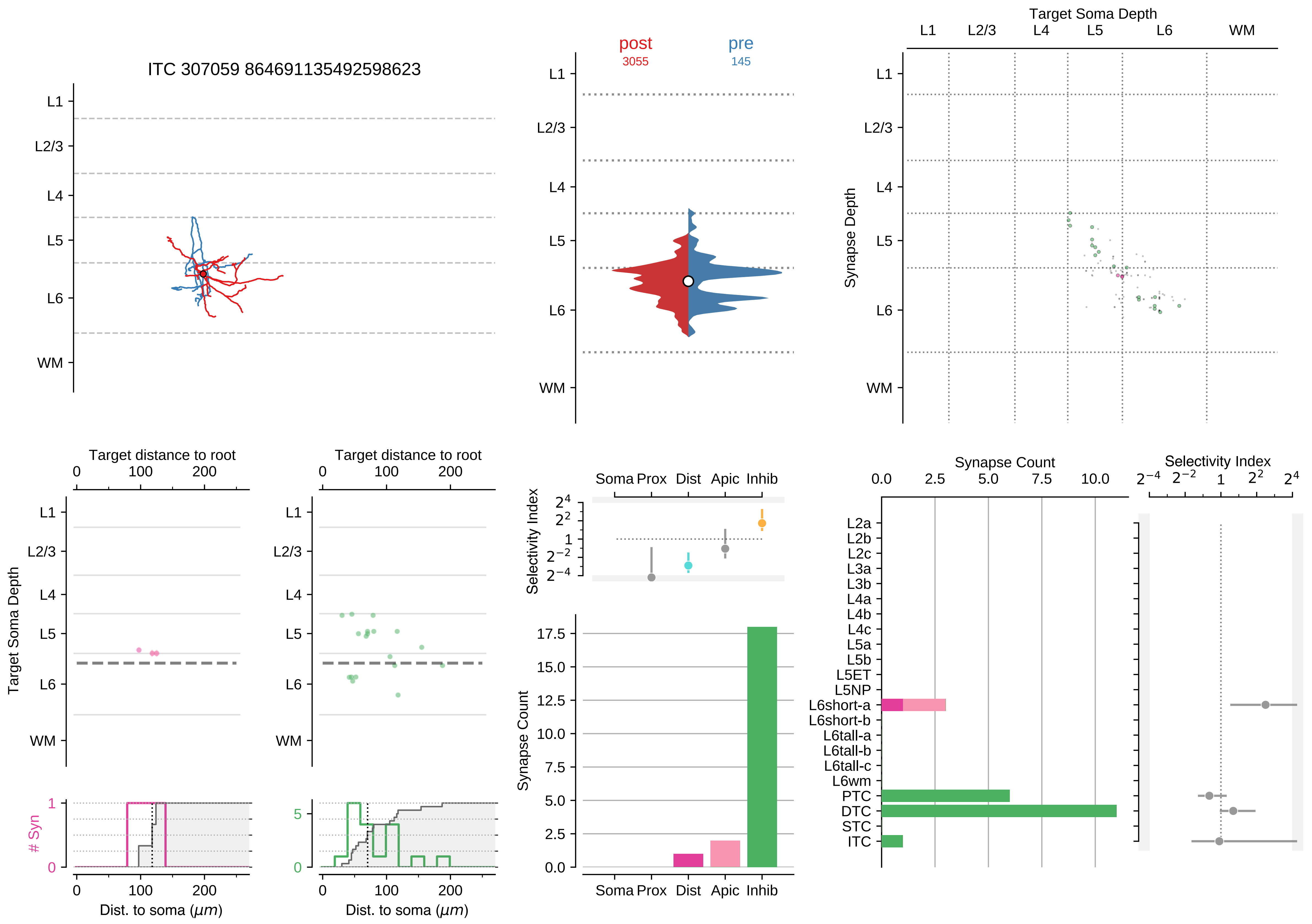

ITC 307059 864691135492598623
L1
L2/3
L4
L5
L6
WM
post
3055
pre
145
Target Soma Depth
Synapse Depth
Target distance to root
0
100
200
Target Soma Depth
### Syn
Dist. to soma ($\mu m$)
Soma Prox Dist Apic Inhib
Selectivity Index
Synapse Count
Synapse Count
Selectivity Index
L2a
L2b
L2c
L3a
L3b
L4a
L4b
L4c
L5a
L5b
L5ET
L5NP
L6short-a
L6short-b
L6tall-a
L6tall-b
L6tall-c
L6wm
PTC
DTC
STC
ITC

### Motif Group 20

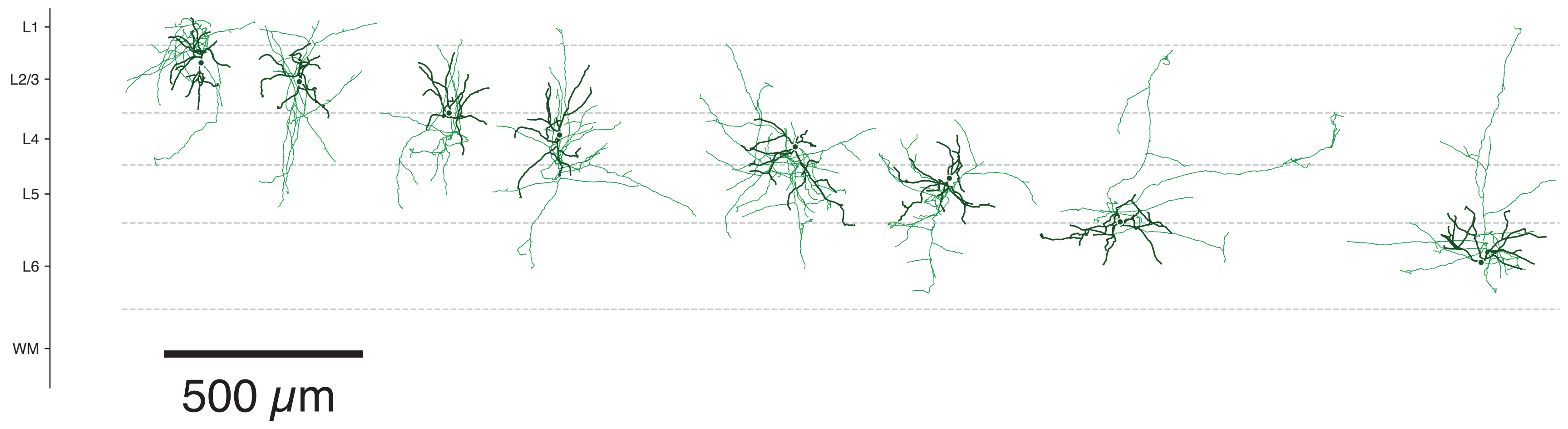

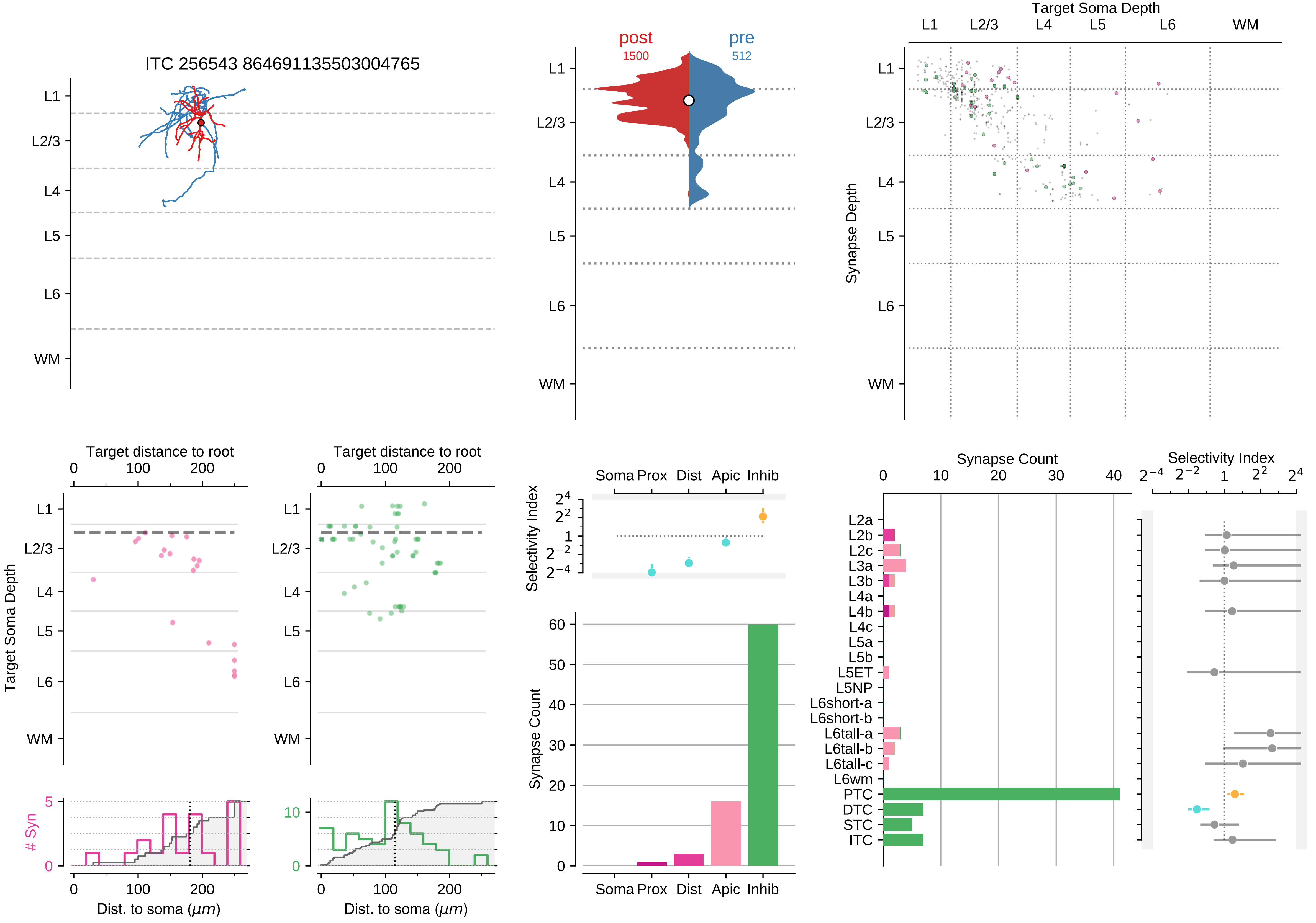
ITC 256543 864691135503004765
L1
L2/3
L4
L5
L6
WM
post
1500
pre
512
Target Soma Depth
Synapse Depth
Target distance to root
Target Soma Depth
### Syn
Dist. to soma (μm)
Soma Prox Dist Apic Inhib
Selectivity Index
Synapse Count
Synapse Count
Selectivity Index
L2a
L2b
L2c
L3a
L3b
L4a
L4b
L4c
L5a
L5b
L5ET
L5NP
L6short-a
L6short-b
L6tall-a
L6tall-b
L6tall-c
L6wm
PTC
DTC
STC
ITC

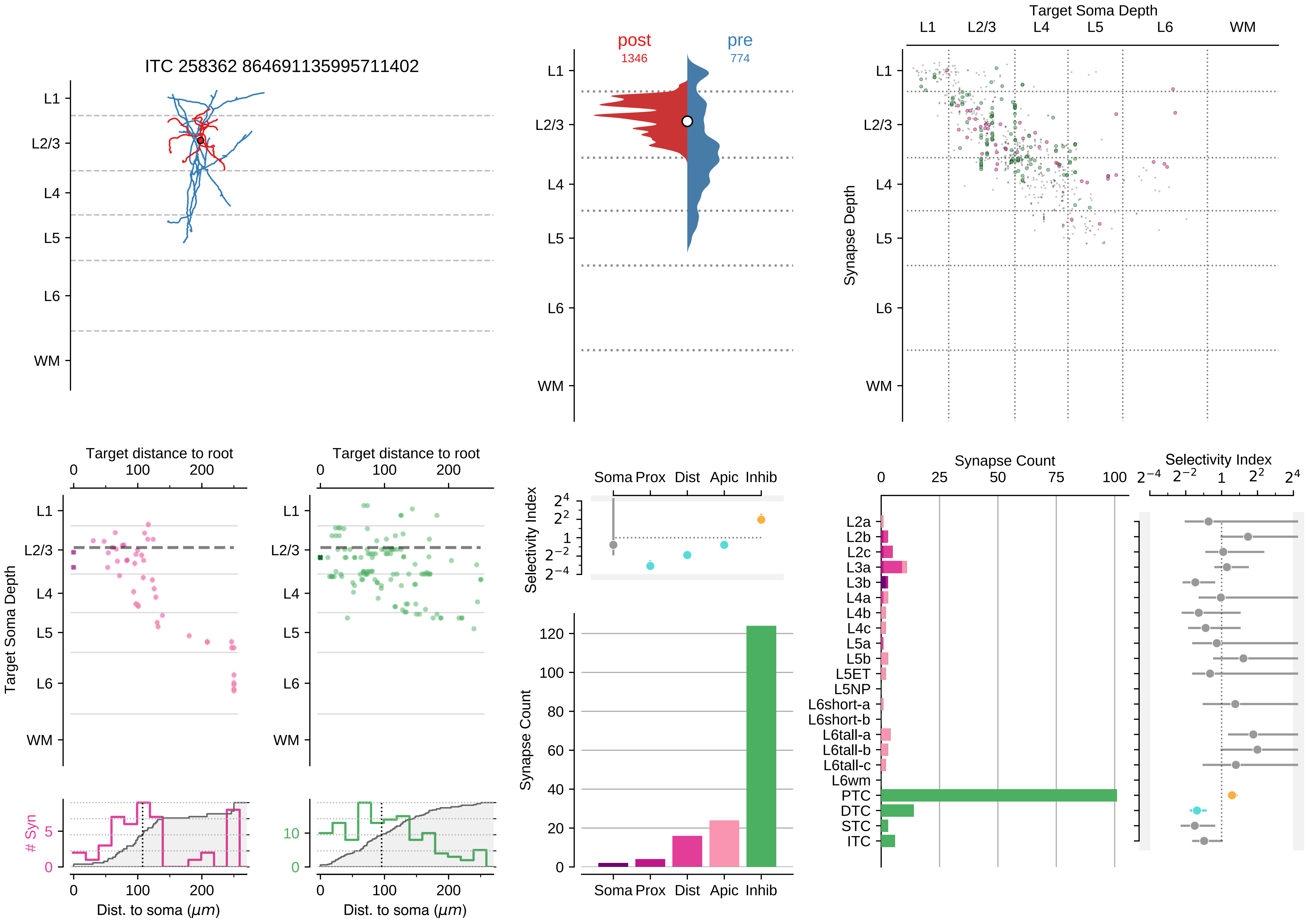

ITC 258362 864691135995711402
L1
L2/3
L4
L5
L6
WM
post
1346
pre
774
Target Soma Depth
Synapse Depth
Target distance to root
Target Soma Depth
### Syn
Dist. to soma ($\mu m$)
Selectivity Index
Soma Prox Dist Apic Inhib
Synapse Count
Synapse Count
Selectivity Index
L2a
L2b
L2c
L3a
L3b
L4a
L4b
L4c
L5a
L5b
L5ET
L5NP
L6short-a
L6short-b
L6tall-a
L6tall-b
L6tall-c
L6wm
PTC
DTC
STC
ITC

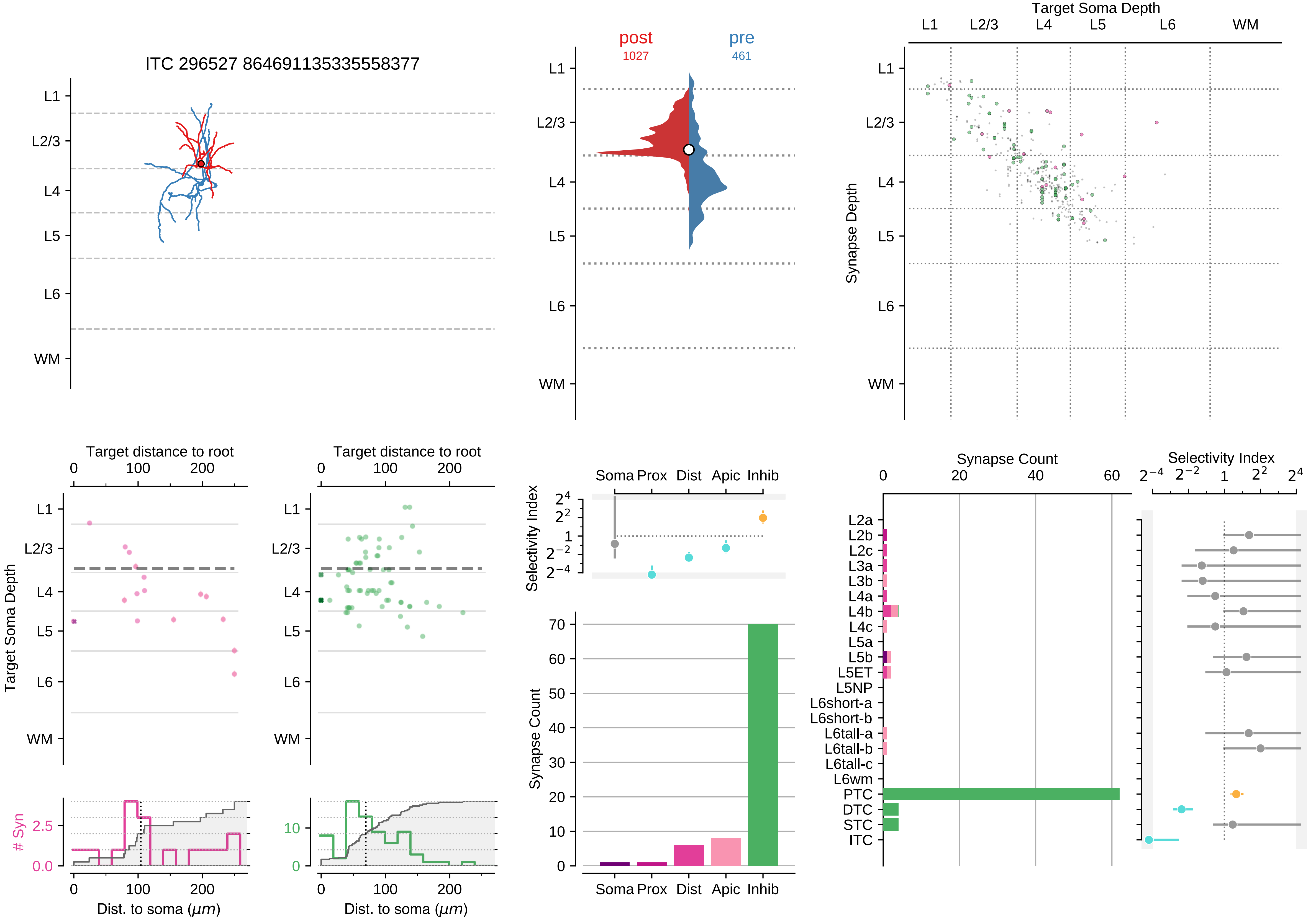

ITC 296527 864691135335558377
post
1027
pre
461
Target Soma Depth
Synapse Depth
L1
L2/3
L4
L5
L6
WM
Target distance to root
Target Soma Depth
### Syn
Dist. to soma ($\mu m$)
Selectivity Index
Soma Prox Dist Apic Inhib
Synapse Count
L2a
L2b
L2c
L3a
L3b
L4a
L4b
L4c
L5a
L5b
L5ET
L5NP
L6short-a
L6short-b
L6tall-a
L6tall-b
L6tall-c
L6wm
PTC
DTC
STC
ITC

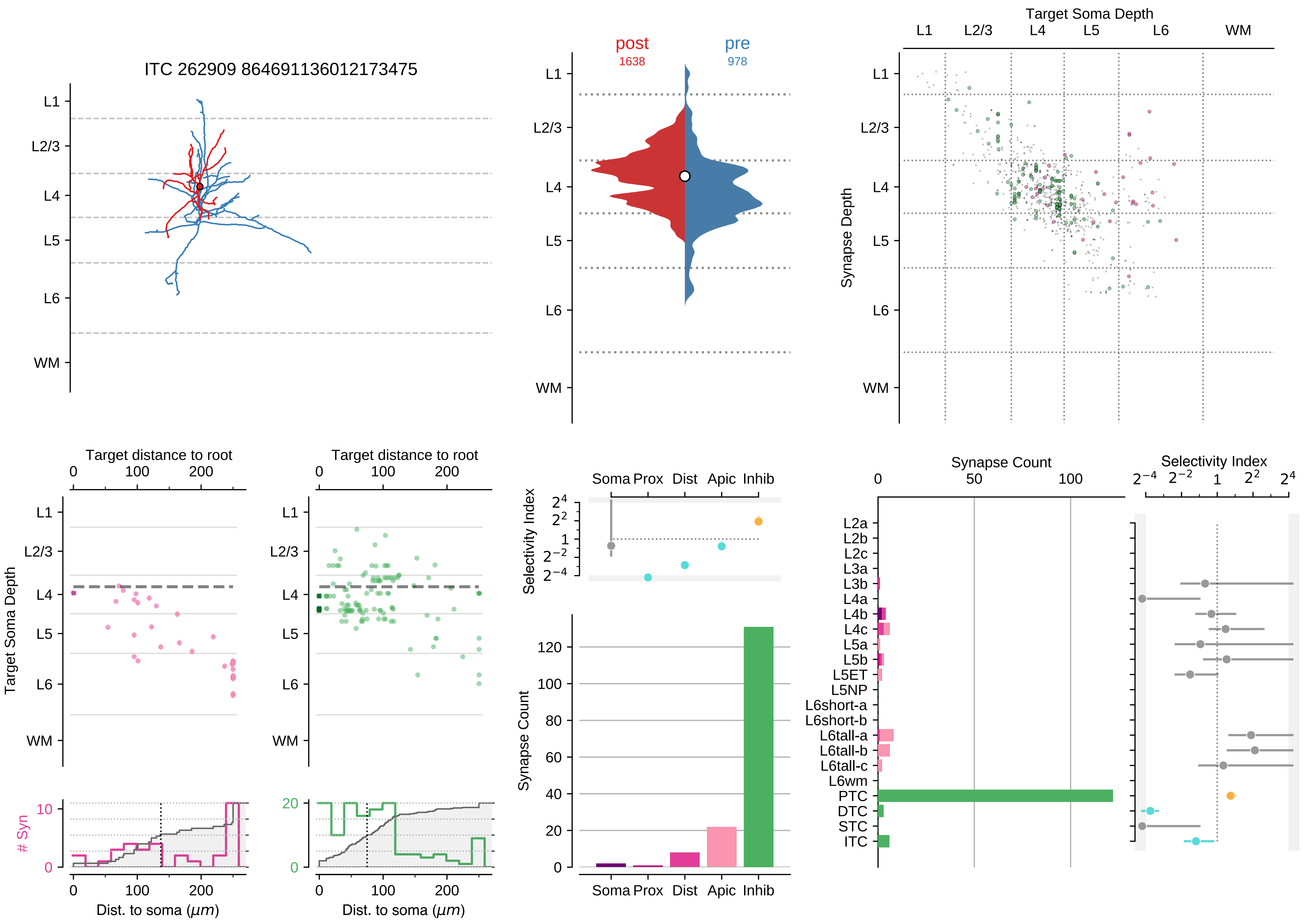

ITC 262909 864691136012173475
L1
L2/3
L4
L5
L6
WM
post
1638
pre
978
Target Soma Depth
Synapse Depth
Target distance to root
Target Soma Depth
### Syn
Dist. to soma (μm)
Selectivity Index
Soma Prox Dist Apic Inhib
Synapse Count
Selectivity Index
L2a
L2b
L2c
L3a
L3b
L4a
L4b
L4c
L5a
L5b
L5ET
L5NP
L6short-a
L6short-b
L6tall-a
L6tall-b
L6tall-c
L6wm
PTC
DTC
STC
ITC

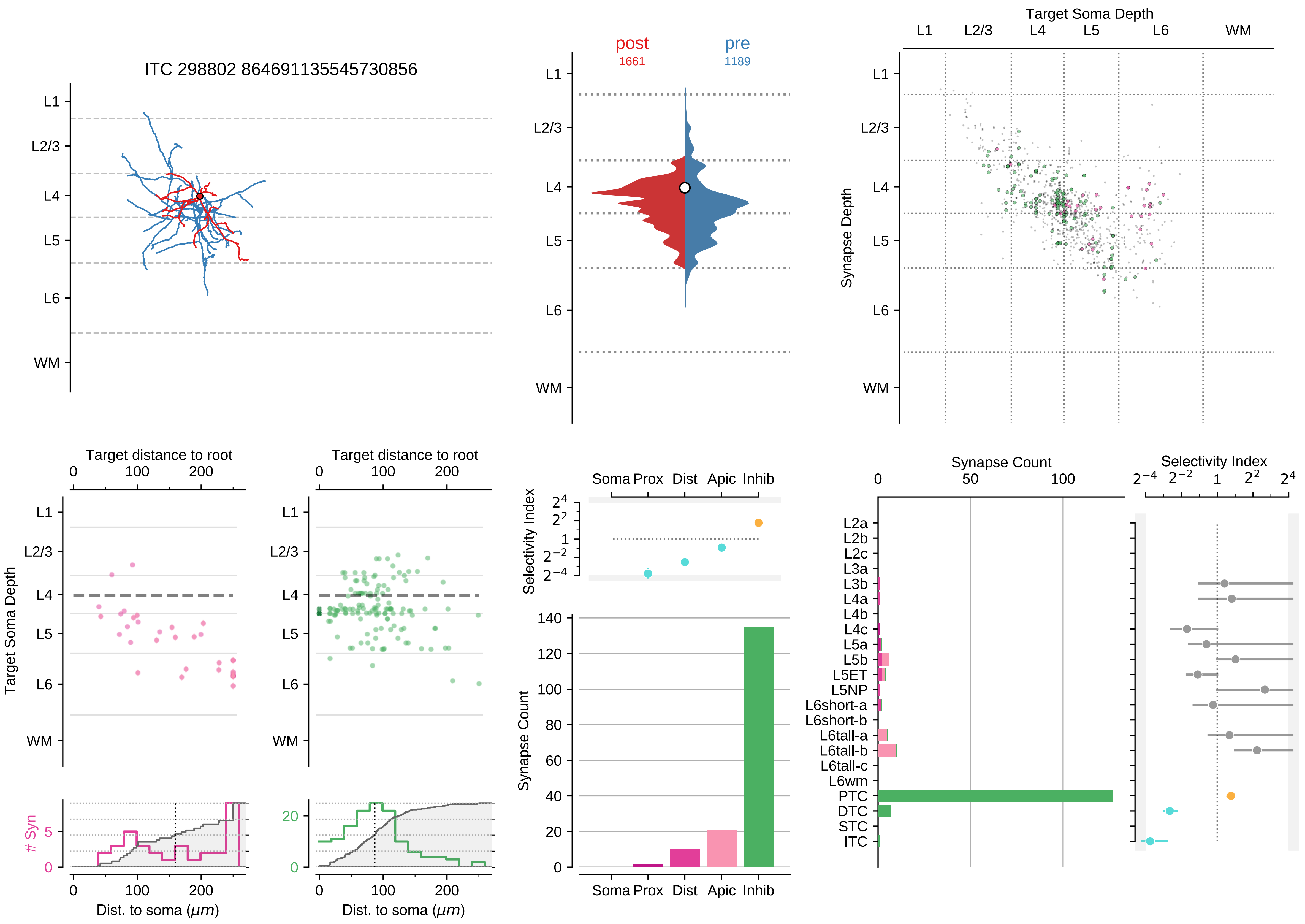

ITC 298802 864691135545730856
L1
L2/3
L4
L5
L6
WM
post
1661
pre
1189
Target Soma Depth
Synapse Depth
Target distance to root
Target Soma Depth
### Syn
Dist. to soma ($\mu m$)
Soma Prox Dist Apic Inhib
Selectivity Index
Synapse Count
L2a
L2b
L2c
L3a
L3b
L4a
L4b
L4c
L5a
L5b
L5ET
L5NP
L6short-a
L6short-b
L6tall-a
L6tall-b
L6tall-c
L6wm
PTC
DTC
STC
ITC

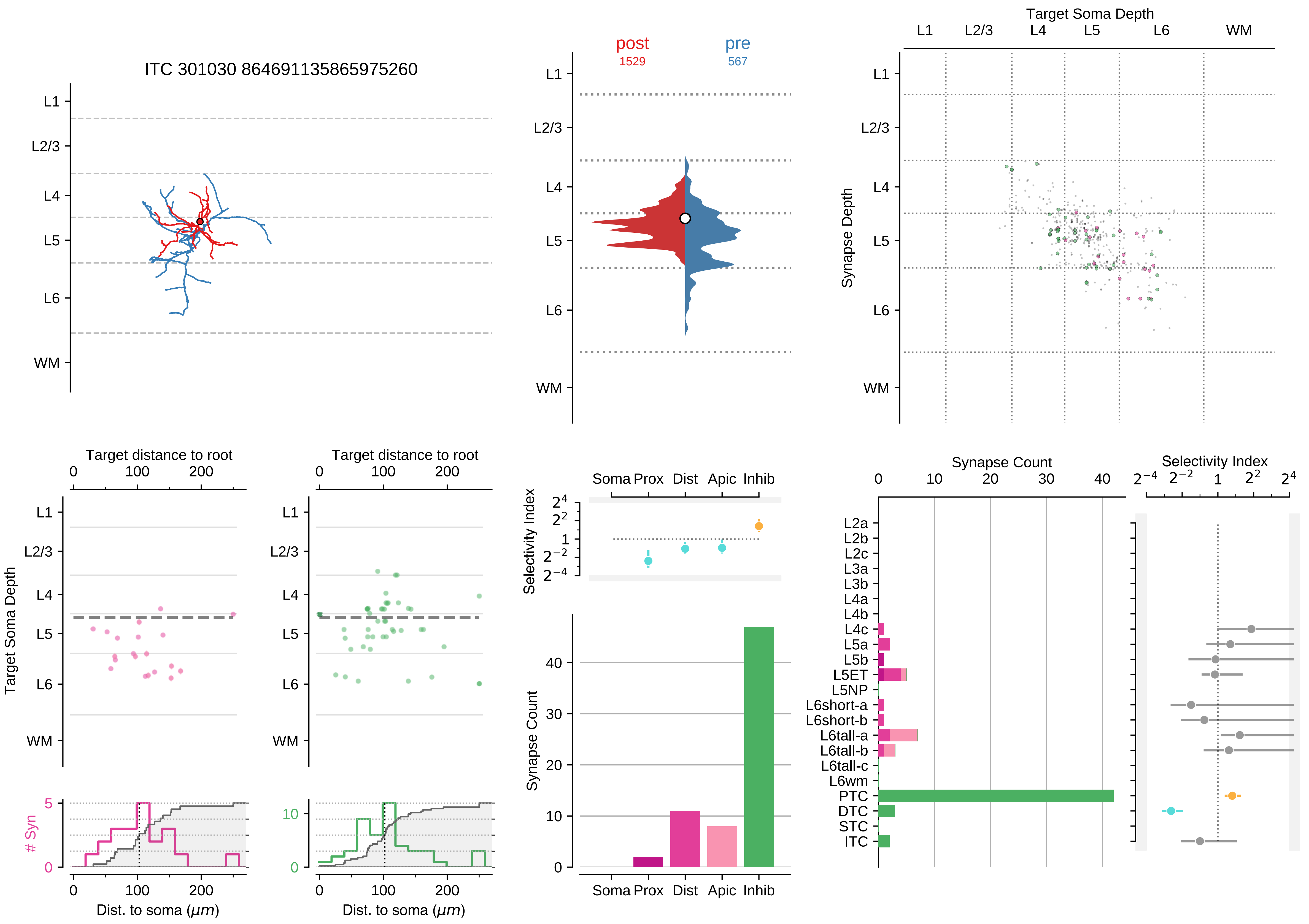

ITC 301030 864691135865975260
L1
L2/3
L4
L5
L6
WM
post
1529
pre
567
Target Soma Depth
Synapse Depth
Target distance to root
Dist. to soma ($\mu m$)
### Syn
Selectivity Index
Synapse Count
Soma Prox Dist Apic Inhib
L2a
L2b
L2c
L3a
L3b
L4a
L4b
L4c
L5a
L5b
L5ET
L5NP
L6short-a
L6short-b
L6tall-a
L6tall-b
L6tall-c
L6wm
PTC
DTC
STC
ITC

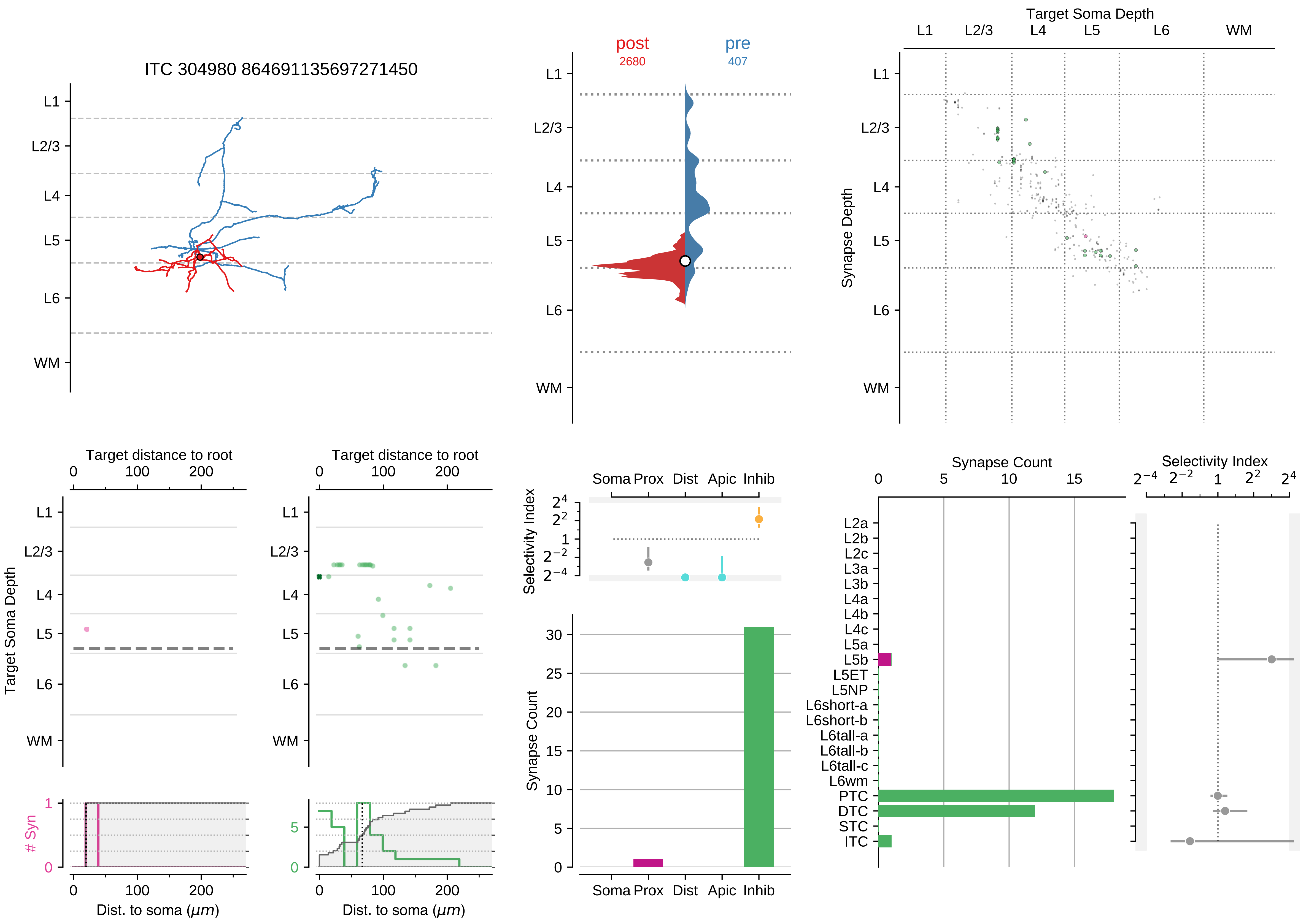

ITC 304980 864691135697271450
L1
L2/3
L4
L5
L6
WM
post
2680
pre
407
Target Soma Depth
Synapse Depth
Target distance to root
Target Soma Depth
### Syn
Dist. to soma ($\mu m$)
Soma Prox Dist Apic Inhib
Selectivity Index
Synapse Count
L2a
L2b
L2c
L3a
L3b
L4a
L4b
L4c
L5a
L5b
L5ET
L5NP
L6short-a
L6short-b
L6tall-a
L6tall-b
L6tall-c
L6wm
PTC
DTC
STC
ITC

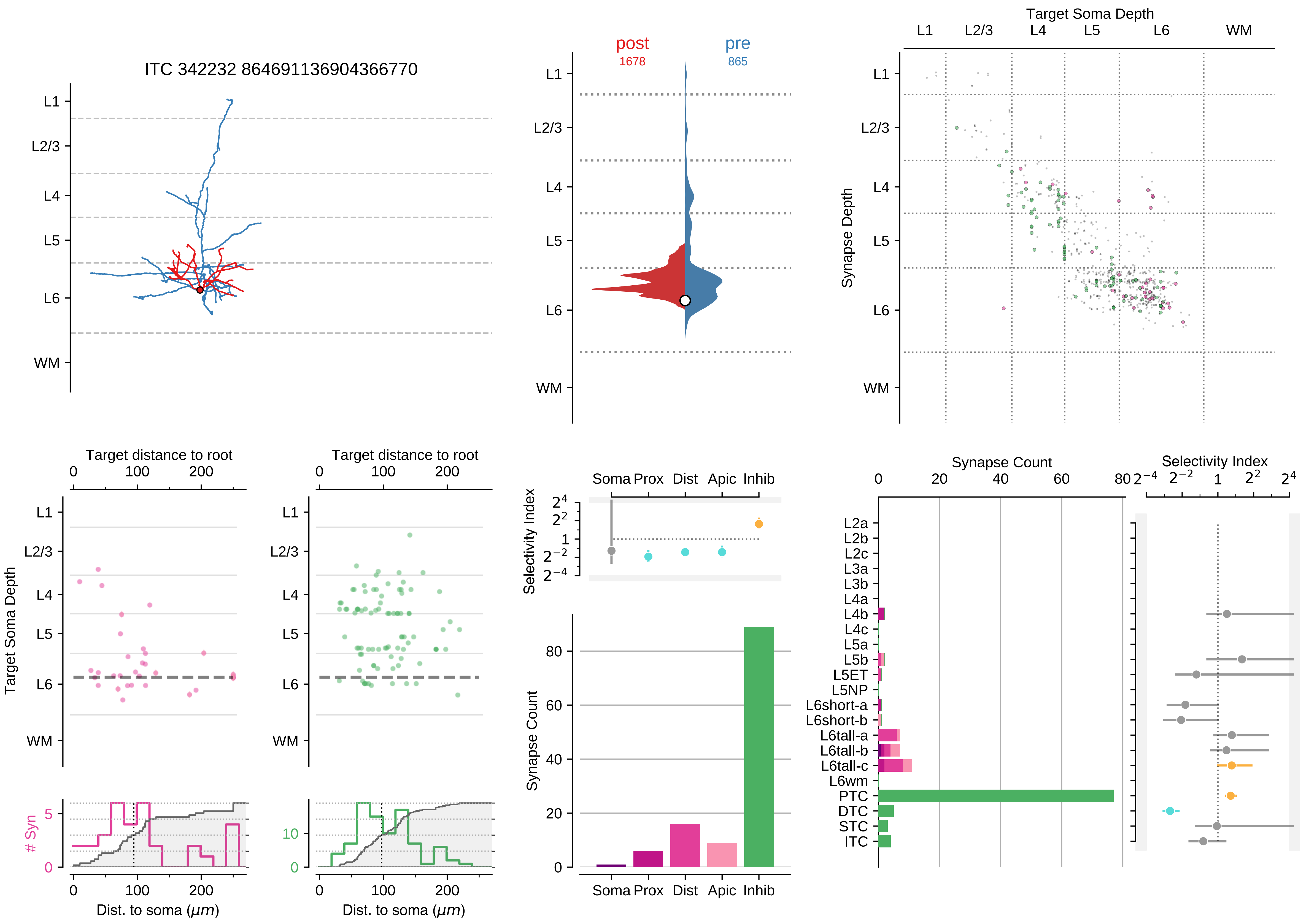

ITC 342232 864691136904366770
L1
L2/3
L4
L5
L6
WM
post
1678
pre
865
Target Soma Depth
Synapse Depth
Target distance to root
Dist. to soma (μm)
### Syn
Selectivity Index
Synapse Count
Soma
Prox
Dist
Apic
Inhib
L2a
L2b
L2c
L3a
L3b
L4a
L4b
L4c
L5a
L5b
L5ET
L5NP
L6short-a
L6short-b
L6tall-a
L6tall-b
L6tall-c
L6wm
PTC
DTC
STC
ITC
